# Deciphering the function of Com_YlbF domain-containing proteins in Staphylococcus aureus

**DOI:** 10.1101/2024.11.14.623514

**Authors:** Zayda Corredor-Rozo, Ricaurte Marquez-Ortiz, Myriam L. Velandia-Romero, Deisy Abril-Riaño, Johana Madroñero, Luisa F. Prada, Natasha Vanegas-Gomez, Begoña García, Maite Echeverz, María Angélica Calderón-Peláez, Jacqueline Chaparro-Olaya, Liliana Morales, Carlos Nieto-Clavijo, Javier Escobar-Perez

## Abstract

The com_ylbF domain-containing proteins that inhibit sporulation, competition, and biofilm formation by affecting the activity of Ribonuclease RNase-Y in *Bacillus subtilis*. Similar Com_YlbF proteins are found in *Staphylococcus aureus*, but their function is yet to be determined. This study investigates the role of com_ylbF domain-containing proteins (Qrp/YheA, YmcA, and YlbF) in *S. aureus* by evaluating the impact of *qrp/yheA*, *ymcA,* and *ylbF* gene deletion on biofilm formation, PIA/PNAG production, and hemolytic capacity. It was found that deletion of the three *qrp/yheA*, *ymcA,* and *ylbF* genes in *S. aureus* produced a decrease in biofilm formation, a slight decrease in PIA/PNAG production, and a decrease in its hemolytic capacity. Global transcriptional analysis in the mutant strain showed dysregulation of several genes associated with pathogenesis, notably a decreased expression of the *agrA* gene (quorum sensing system), the *delta*, *alpha,* and *gamma* hemolysins, as well as the *sdrC* genes, *eap/map* (adhesion proteins). It also showed-an attenuation of virulence, manifesting as enhanced survival among *Galleria mellonella* larvae, and BALB/c mice, possibly associated due to disrupted RNase-Y activity when the *qrp/yheA*, *ymcA,* and *ylbF* are deleted. In conclusion, this work supports the hypothesis that this new family of proteins containing a com_ylbF domain is involved in the regulation of genes related to biofilm formation, hemolysis, and virulence in *S. aureus* through a change in RNase-Y activity.

## INTRODUCTION

*Staphylococcus aureus* can cause a wide range of infections in humans, from superficial skin infections to severe and potentially life-threatening infections such as sepsis, bacteraemia, pneumonia, infective endocarditis, and osteomyelitis (1, 2). The ability of this bacterium to cause infection is due to the production of dozens of virulence factors such as hemolysins, leukocidins, modulins, proteases, surface-associated protein adhesins, exotoxins, superantigens, DNases, exopolysaccharides, host immune evasion molecules, and immune modulators (3, 4). In addition, *S. aureus* forms biofilms on indwelling medical devices or implanted materials, leading to chronic and recurrent infections that are challenging to treat (5). The expression of these virulence factors is tightly regulated by a complex and dynamic circuit that rapidly responds to changing environmental conditions.

YlbF, YmcA, and more recently YheA (also known as Qrp, glutamine-rich protein (6)), are a group of small proteins (10 to 15kDa) with a com_ylbF domain spanning through almost the entire protein (7), and their function in *S. aureus* is still unknown. *In-silico* analysis shows a wide distribution of these proteins in Gram-positive bacteria, suggesting their role in conserved biological processes. YmcA and YlbF have been characterized genetically and biochemically in *Bacillus subtilis,* another Gram-positive bacterium, where their deletion results in significant deficiencies in sporulation, competency, and biofilm formation (7, 8). In *B. subtilis,* YmcA, and YlbF form a stable ternary complex with the YaaT protein (named Y-complex), the latter lacking the com_ylbF domain required for spore and biofilm formation (9). The Y-complex modulates the function of two key regulatory transcriptional proteins: Spo0A and SinR. Increasing for Spo0A its phosphorylated-active form (9), and destabilizing SinR mRNA through cleavage by the endoribonuclease RNase-Y (9–12). Notably, the Y-complex physically interacts with RNase-Y and is necessary for the processing and regulation of many mRNA transcripts (11).

Information on the com_ylbF domain-harboring protein of *S. aureus* is very limited. A study performed by Deloughery *et al.* using end-enrichment RNA sequencing assay (Rend-seq) showed that deletion of the *ylbF* gene altered RNase-Y mRNA processing of the *cggR(gapR)-gapA* operon, and the riboswitches *SerS* and *ValS* (11). We also demonstrated that *S. aureus* expresses another com_ylbF domain-containing protein, Qrp/YheA, which shares a highly conserved 3D structure with YmcA and YlbF, and contains a putative conserved motif (QQKQMQ) located in the central region of the com_ylbF domain, which could be involved in its function (6). Here, we provide evidence that Qrp/YheA, YmcA, and YlbF proteins could modulate virulence and biofilm formation in *S. aureus* through a post-transcriptional regulation mediated by RNase-Y activity (13, 14).

## RESULTS

### The Qrp/YheA, YmcA, and YlbF proteins could participate in the transcription of genes associated with virulence and biofilms

In *B. subtilis,* Com_YlbF proteins have been associated with the regulation of numerous genes, by influencing the activation of master transcription factors or the activity of ribonucleases (11). To further understand the function of Com_YlbF proteins in *S. aureus*, we generated a triple mutant, deletion of the *qrp*/*yheA*, *ymcA*, and *ylbF* genes and compared the transcriptomic profile of this mutant with the wild-type strain. In this assay, we obtained data from 2,872 transcripts (with FPKMs > 0.5), covering 95% of the ORFs in the *S. aureus* genome. The results showed that 201 genes were differentially expressed (with absolute log2 fold change above the conservative threshold of 1.5) in the absence of *qrp*/*yheA*, *ymcA*, and *ylbF* (Figure 1A). Of these, 155 genes were downregulated and 46 were upregulated (Table S3). The top 17 biological processes associated with the differentially expressed genes in *S. aureus* are shown in Figure 1B. Notably, the GO annotations mainly include three major categories, ‘virulence’ (23 upregulated genes; 43 downregulated genes) [GO:0009405], ‘regulation of transcription DNA-dependent’ (18 up; 22 down) [GO:0006355], ‘membrane transport’ (15 up; 19 down) [GO:0055085]. Other noble subsets include ‘hemolysis in another organism’ (5 down) [GO:0044179] and ‘cell adhesion’ as factors involved in biofilm formation (5 up) [GO:0007155] (Figure 1A).

**Figure 1.**
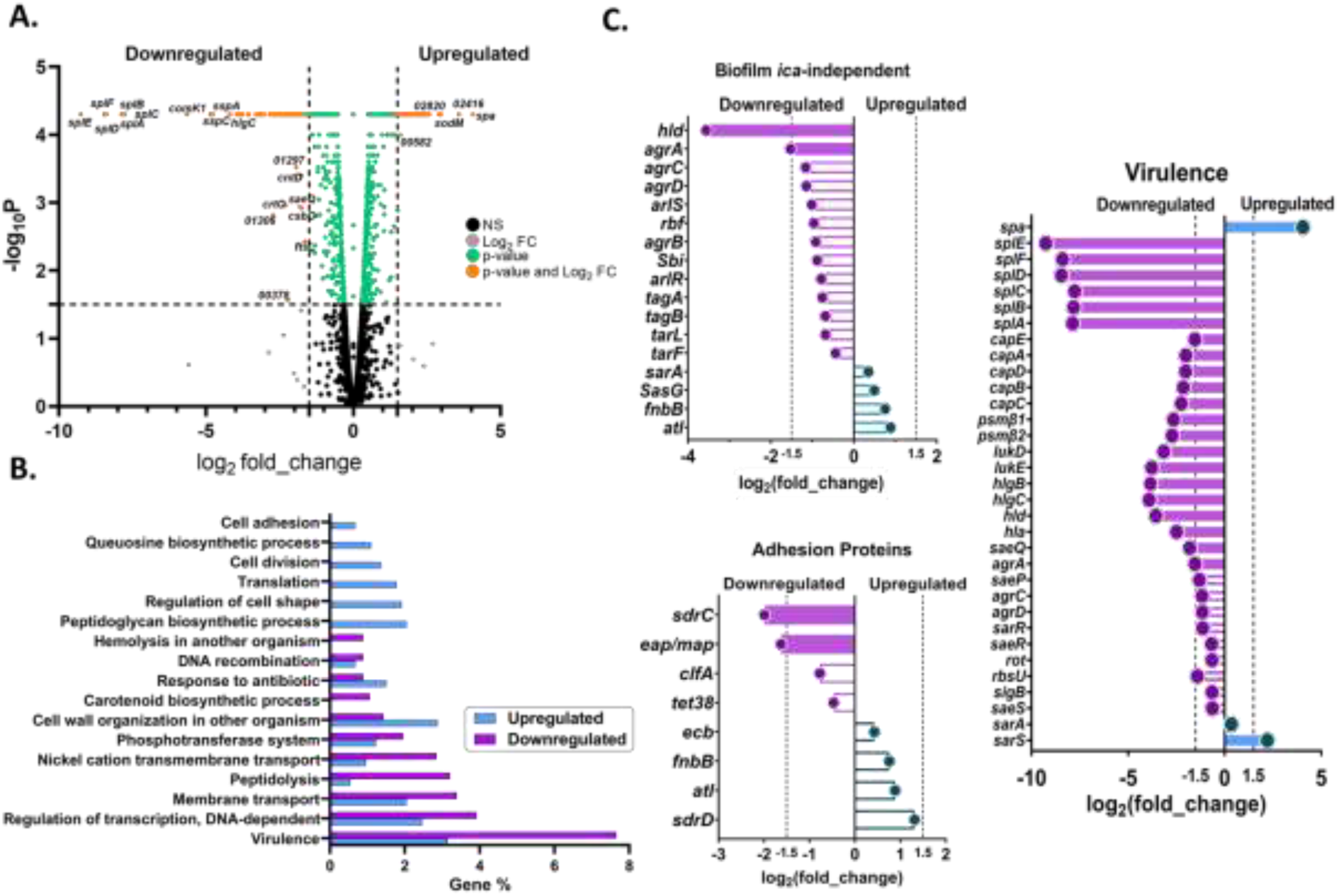
Differentially expressed genes (DEGs) in *S. aureus qrp*/*yheA*, *ymcA, ylbF* mutant. **A.** Volcano plot of the genes detected in the exponential phase. Each dot represents a gene, with orange dots highlighting differentially expressed genes (DEGs). Right and left sides indicate upregulated and downregulated genes, respectively. **B.** Distribution of differentially expressed transcripts. The graph shows the DEGs assigned to the 17 major GO Biological process categories. The percentage of upregulated genes is shown in blue and the percentage of downregulated genes is shown in purple. **C.** Differentially expressed genes involved in biofilm formation as *ica*-independent pathway, adhesion proteins, and virulence. The dotted lines indicate the detection limit for transcripts with genes exhibiting a log2 fold change |FC| ≥1.5 considered differentially expressed.

Notably, the transcripts for genes encoding virulence factors and biofilm formation were categorized under ‘virulence’. Specifically, the spl (serine-protease like) operon, that harbors six serine proteases (*splE*, *splD*, *splF*, *splA*, *splB*, and *splC*) was the most downregulated (7 to 9 fold decrease), followed by hemolysins (*lukE*, *lukD*, *hlgB*, *hlgC*, *hla,* and *hld*) with a 2.4 to 3.9-fold decrease, phenol-soluble modulins (*psmβ1*, *psmβ2*) with a 2.6 to 2.7 fold decrease, fibrinogen binding protein SdrC of the MSCRAMM group (*sdrC*) with a 1.9-fold decrease, extracellular adhesion protein Eap/Map of the SERAM group (*eap/map*) with a1.6-fold decrease, and capsular polysaccharide biosynthesis proteins (*capA*, *capB*, *capC*, *capD,* and *capE*) with a 1.5 to 2 fold decrease. Interestingly, the accessory gene regulatory Agr-system (*agrA*) also showed a decrease of 1.5-fold. On the other hand, among the up-regulated and pathogenesis-categorized genes, the SarA-homologous transcriptional regulator (*sarS*) and the immunoglobulin G binding protein A precursor (*spa*) showed the biggest change (4 fold increase) (Figure 1A, C).

### The Qrp/YheA, YmcA, and YlbF proteins promote biofilm formation in a PIA/PNAG-dependent manner

Based on the results of the transcriptomic analysis, biofilm formation is one of the phenotypes affected by the absence of Qrp/YheA, YmcA, and YlbF proteins. To determine whether the effect on biofilm was related to any individual protein, we constructed single and double mutants in the *qrp*/*yheA*, *ymcA*, and *ylbF* genes. Deletions of these genes had no impact on growth rate (Figure S1).

We assessed the biofilm-forming ability of the mutants under high glucose concentration (2.5% p/v) and high osmolarity (4% p/v NaCl) conditions. Most mutant strains showed reduced biofilm formation (P<0.05) under high glucose concentration, but not under high osmolarity (Figure 2A). Significant reductions in biofilm formation compared to the wild-type (WT) strain were observed in the single mutants Δ*ylbF* (30%, P < 0.01: p= 0.006) and Δ*ymcA* (28%, P < 0.05: p= 0.020). The double mutants Δ*qrp*/*yheA*Δ*ymcA* (31%, P < 0.05: p= 0.037), Δ*qrp*/*yheA*Δ*ylbF* (8%, P < 0.05: p= 0.004), and the triple mutant Δ*qrp*/*yheA*Δ*ymcA*Δ*ylbF* (26%, P < 0.05: p= 0.021) also showed significant reductions (Figure 2A).

**Figure 2.**
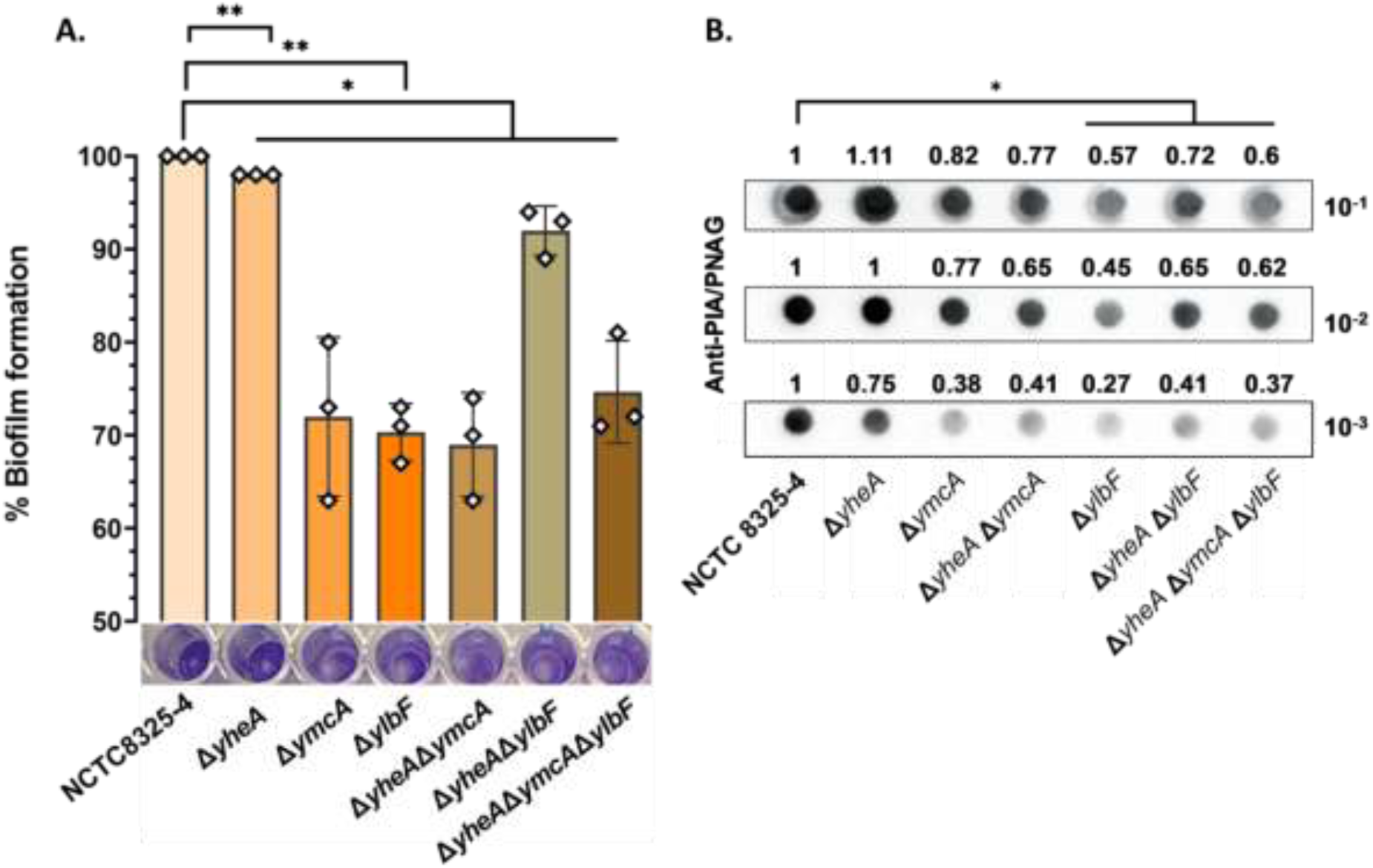
Mutants for *qrp/yheA*, *ymcA*, and *ylbF* showed a reduction of biofilm in *S. aureus*. **A.** Percentage of biofilm formation relative to WT, measured as OD (600 nm) after crystal violet staining after 24 h incubation in the presence of glucose. Significant reductions in biofilm formation compared to the wild-type (WT) strain were observed in Δ*ylbF* (30%, P < 0.01: p= 0.006), Δ*ymcA* (28%, P < 0.05: p= 0.020), Δ*qrp*/*yheA*Δ*ymcA* (31%, P < 0.05: p= 0.037), Δ*qrp*/*yheA*Δ*ylbF* (8%, P < 0.05: p= 0.004), and Δ*qrp*/*yheA*Δ*ymcA*Δ*ylbF* (26%, P < 0.05: p= 0.021). **B.** Dot-blot densitometric analysis of PIA-PNAG in *S. aureus* NCTC 8325-4 and mutants in the presence of glucose (0.25% p/v; TSBG). Significant reductions in PIA/PNAG production in Δ*ylbF* (57%, P < 0.05: p= 0.023), Δ*qrp*/*yheA*Δ*ylbF* (40%, P < 0.05: p= 0.048), and Δ*qrp*/*yheA*Δ*ymcA*Δ*ylbF* (47%, P < 0.05: p= 0.026). For dot blot analysis serial dilutions (1:10 to 1:1.000) of the total protein extraction samples were spotted onto nitrocellulose membranes, and PNAG production was detected with PIA/PNAG antibody. Relative quantification was obtained using the ImageJ and the values are expressed in densitometric proportions, normalized to wild-type (WT) strain. The assays were performed in two independent experiments, each conducted in triplicate. Δ*yheA=*Δ*qrp/yheA*. Statistical analysis was performed using a paired t-test. *P < 0.05, **P < 0.01.

Production of PIA/PNAG by the *icaADBC* operon is tightly regulated at the transcriptional level (15). We hypothesized that the role of Com_YlbF proteins (Qrp/YheA, YmcA, YlbF) in biofilm formation could be related to the regulation of PIA/PNAG exopolysaccharide expression. To assess changes in PIA/PNAG production, we measured the levels of this exopolysaccharide in strains cultivated in TSB supplemented with glucose (0,25% p/v; TSB_G_) using dot-blot experiments with an anti-PIA/PNAG polyclonal antibody (16, 17). Our results showed significant reductions in PIA/PNAG production by 57% (P < 0.05: p= 0.023), 40% (P < 0.05: p= 0.048), and 47% (P < 0.05: p= 0.026) in Δ*ylbF*, Δ*qrp/yheA*Δ*ylbF* and Δ*qrp/yheA*Δ*ymcA*Δ*ylbF* mutants, respectively (Figure 2B). These findings suggest that YmcA and YlbF proteins are involved in the regulation of the synthesis of PIA/PNAG exopolysaccharide synthesis.

### The Qrp/YheA, YmcA, and YlbF proteins contribute to the hemolytic activity of S. aureus

The transcriptomic analysis revealed a decrease in hemolysins expression in the mutants. Red blood cells (RBCs) lysis is an important ability that *S. aureus* possesses and that has been associated with the production of different hemolysins and leukocidins, such as ED (18). We investigated the role of *qrp/yheA*, *ymcA,* and *ylbF* mutants in the hemolytic activity of the bacteria using qualitative and quantitative methods. Qualitative analysis was performed by observing opacity around individual colonies in blood-agar assays (19), while quantitative analysis involved measuring the absorbance of the released hemoglobin from hemolyzed RBCs. Interestingly, in the qualitative assay, we found that the hemolytic activity of the triple-mutant strain (*qrp/yheA, ymcA,* and *ylbF* deletion) was significantly reduced in 74% (P < 0.01: p= 0,0024) versus the wild-type strain NCTC8325-4. The triple-mutant strain did not produce a halo around colonies, unlike the wild-type strain, which exhibited a transparent halo approximately 1 cm in radius (Figure 3A).

**Figure 3.**
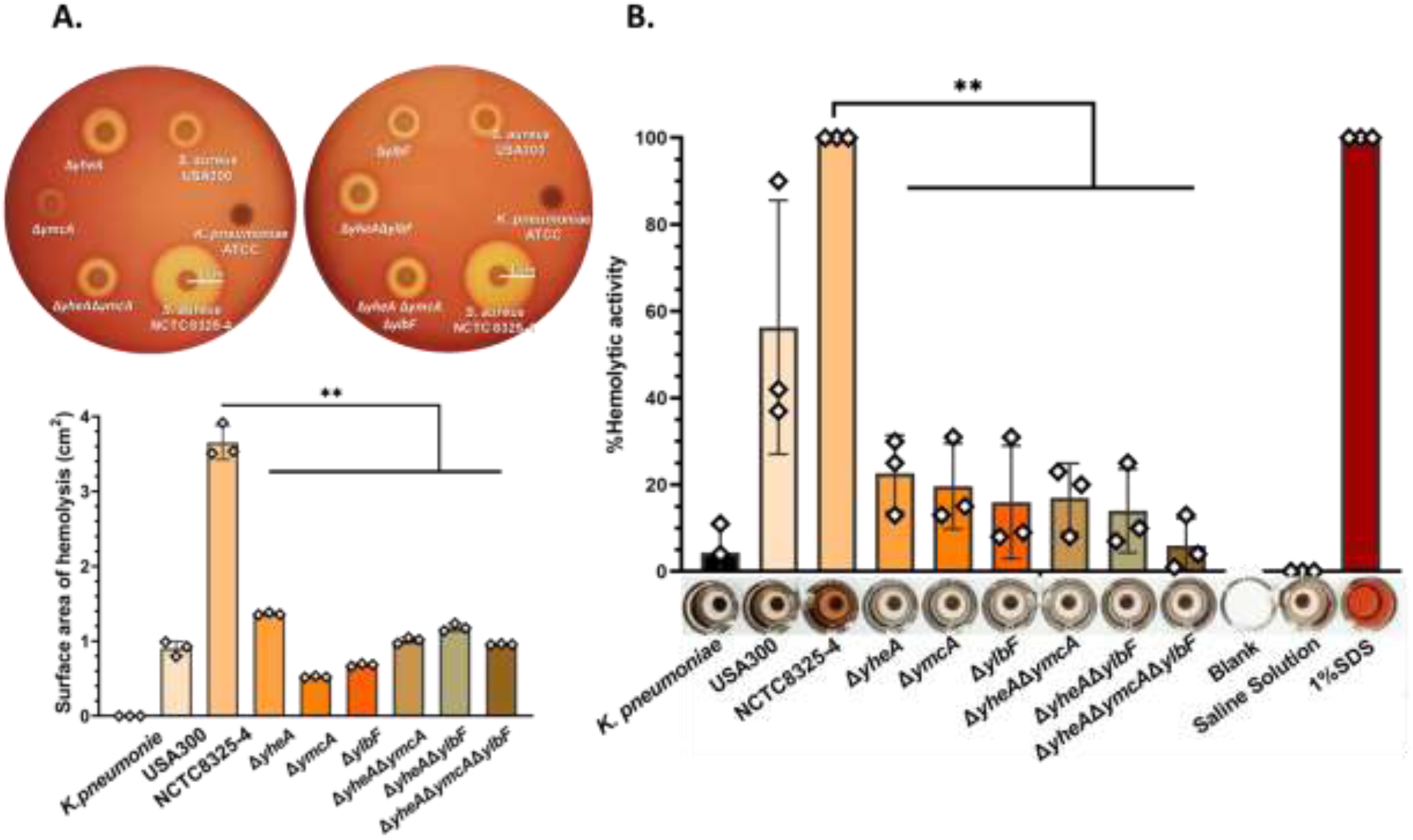
Mutants for *qrp/yheA*, *ymcA*, and *ylbF* showed a reduction of hemolysis for *S. aureus*. Assessment of the hemolytic capability by qualitative analysis was significantly reduced in Δ*yheA* (63%, P < 0.01: p= 0.0035), Δ*ymcA* (86%, P < 0.01: p= 0.0019), Δ*ylbF* (81%, P < 0.01: p= 0.0019), Δ*qrp*/*yheA*Δ*ymcA* (72%, P < 0.01: p= 0.0033), Δ*qrp*/*yheA*Δ*ylbF* (68%, P < 0.01: p= 0.0017), and Δ*qrp*/*yheA*Δ*ymcA*Δ*ylbF* (74%, P < 0.01: p= 0.0024) (**A**) and by quantitative analysis was significantly reduced in Δ*yheA* (77%, P < 0.01: p= 0.0044), Δ*ymcA* (80%, P < 0.01: p= 0.0048), Δ*ylbF* (84%, P < 0.01: p= 0.0076), Δ*qrp*/*yheA*Δ*ymcA* (83%, P < 0.01: p= 0.0028), Δ*qrp*/*yheA*Δ*ylbF* (86%, P < 0.01: p= 0.0039), and Δ*qrp*/*yheA*Δ*ymcA*Δ*ylbF* (94%, P < 0.01: p= 0.0014) (**B**). The hemolytic capability of the *qrp*/*yheA, ymcA,* and *ylbF* mutants of *S. aureus* was drastically reduced after 24 hours of incubation. *K. pneumoniae*, non-hemolytic control. *S. aureus* USA300, positive control Incomplete Hemolytic Phenotype (SIHP). *S. aureus* NCTC 8325-4, positive control Complete Hemolytic Phenotype (SCHP). Δ*yheA=*Δ*qrp/yheA*. Blank, empty well. 0.98% Saline Solution, negative control. 1% SDS, hemolytic positive control. Hemolytic capability quantification in blood-agar plates was obtained using the ImageJ. The assay was conducted once, in triplicate. The scale bar represents 1 cm. Quantification of the hemolytic capability was performed by measuring the hemoglobin released from red blood cells at 540 nm. The assay was performed in three independent experiments, each conducted in triplicate. Statistical analysis was performed using a paired t-test **P < 0.01.

Quantitatively, hemolytic activity was reduced by 94% (P < 0.01; p = 0.0085), with the triple-mutant strain showing minimal hemolysis of RBCs (Figure 3B). Both results suggest that Qrp/YheA, YmcA, and YlbF proteins are involved in the regulation of hemolysins or leukocidin ED expression, probably via up-regulation (18). Single mutant strains showed an average reduction of hemolytic halo by 77% (P < 0.01: p= 0.0076) and of released hemoglobin by 80% (P < 0.01: p= 0.0044), while double mutant strains an average reduced hemolytic halo by 70% (P < 0.01: p= 0.0074) and released hemoglobin by 85% (P < 0.01: p= 0.0066) (Figure 3).

### The Qrp/YheA, YmcA, and YlbF proteins contribute to the transcriptional and post-transcriptional regulation of genes through RNase-Y activity

According to the transcriptome results, Qrp/YheA, YmcA, and YlbF may be associated with transcriptional regulation of some virulence genes. To assess this possibility, a reporter plasmid pCN52 with a transcriptional fusion of *gfp* to the Gamma-hemolysin component C promoter (P*hlgC*), was constructed (20). The transcriptional-fusion constructs were transformed into all deletion strains to monitor promoter activity by Western Blotting using an anti-GFP antibody. Analysis of the expression of *hlgC* by promoter fusion assay using *qrp/yheA*, *ymcA, ylbF* mutant strain revealed an approximately 8,3-fold reduction compared to the wild-type strain. These results suggested that the promoter-activating expression of Gamma-hemolysin (HlgC) could be modulated by Com_YlbF proteins (Figure 4A).

**Figure 4.**
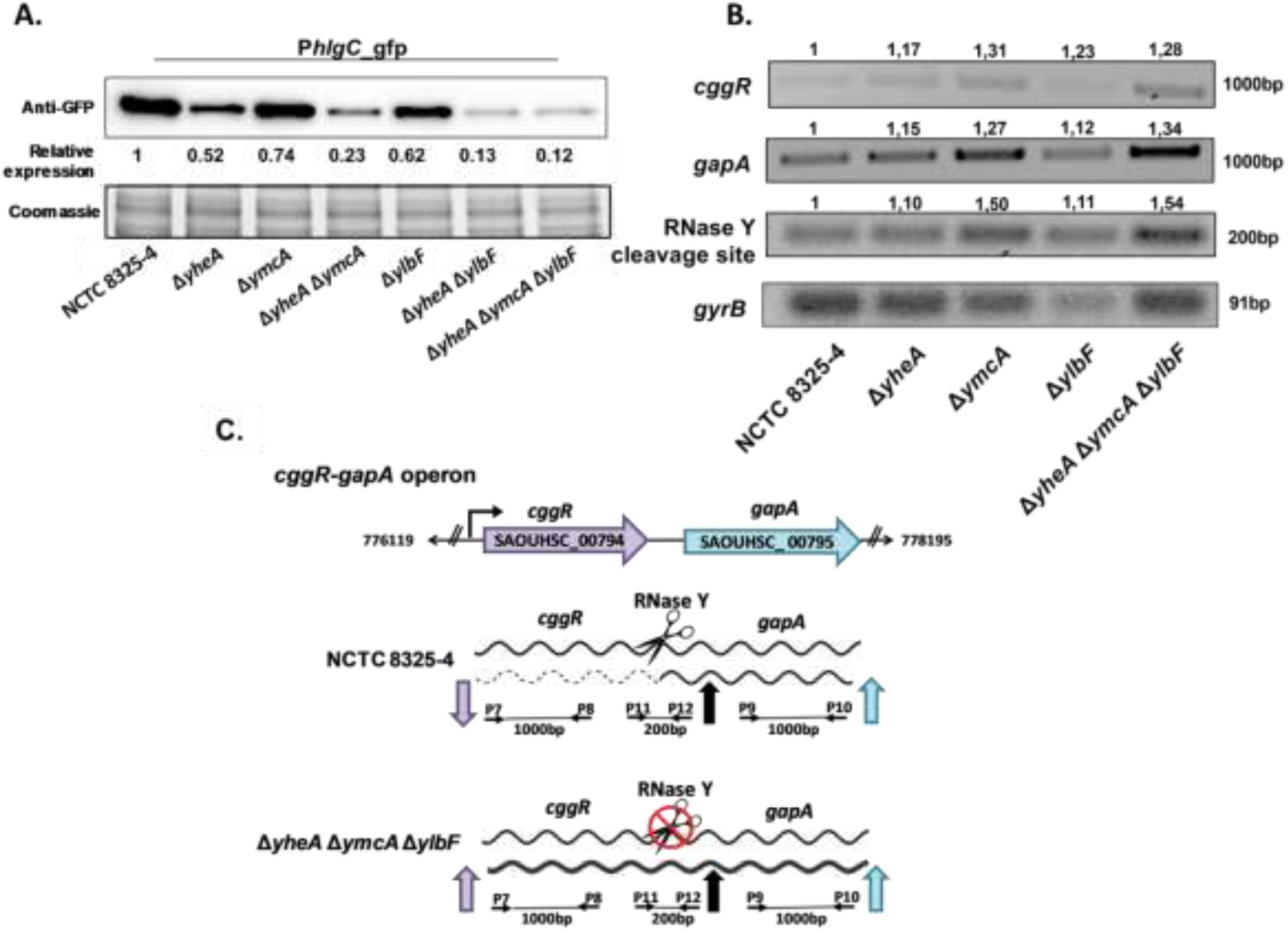
Assessment of the RNase-Y activity and *hlgC* promoter activity in *S. aureus* NCTC 8325-4 and Δ*qrp*/*yheA*, Δ*ymcA*, Δ*ylbF* strains. **A.** Expression of GFP under the control of the *hlgC* promoter at stationary phase (pCN52-P*hlgC*_*gfp*) determined by Western blotting using monoclonal antibodies. Δ*yheA=*Δ*qrp/yheA*. Relative quantification of the GFP protein was obtained using the ImageJ software. The Coomassie-stained gel portion was used as a loading control. **B.** Densitometric analysis of the RT-PCR of the wild-type strain NCTC8325-4 and the simple, double, and triple deletions of *qrp/yheA*, *ymcA,* and *ylbF* genes, indicating variations of the amplified regions in the *cggR-gapA* operon. The housekeeping gene *gyrB* was used for densitometric normalization. Numbers indicate the relative quantification of the PCR products obtained using the ImageJ program. The assay was performed in independent duplicates. **C.** Natural scheme of bicistronic transcription regulation of the *cggR-gapA* operon by RNase-Y mediated processing in *S. aureus* NCTC 8325-4 compared to mutant strains.

In *B. subtilis*, YmcA and YlbF regulate the expression of many mRNA transcripts through the interaction with RNase-Y (10). We speculated that Com_YlbF proteins might also play a role in the post-transcriptional regulation of the virulence genes by collaborating with the activity of RNase-Y in *S. aureus*. We confirmed that the RNase-Y activity was altered in the triple mutant, as this was previously observed in the *ylbF* mutant by DeLoughery A *et al*. (11). This was evidenced by the diminished processing of the glycolytic *gapA* operon, one of the most widely studied targets of RNase-Y(21), and an increased presence of the *cggR* transcript (Figure 4B, C).

### Com_YlbF proteins are intervening in the S. aureus pathogenesis in vivo models

In order to assess whether the phenotypic alterations found in vitro could affect the S. aureus pathogenesis, we performed a *Galleria mellonella* infection model and a murine peritonitis model. For the larval model, they were exposed to different inoculum sizes of both the wild-type *S. aureus* strain and corresponding isogenic mutants to qrp/yheA, ymcA, and ylbF. Mortality rates were monitored over 4 days. The larvae were injected with 5 µl containing 1×10^5 CFU allowed us to calculate a 50% mortality rate by the fourth day and enabled observation of larval health indicators during the study. Survival rates in *G. mellonella* with 5×10^5 CFU, revealing a 40% mortality rate among larvae exposed to the mutant strains compared to 80% mortality with the wild-type strain (Figure 5A). The changes in survival rates were visibly different between the wild-type and triple mutant strains, and significantly reduced in 40% (P < 0.05: p= 0,0397).

**Figure 5.**
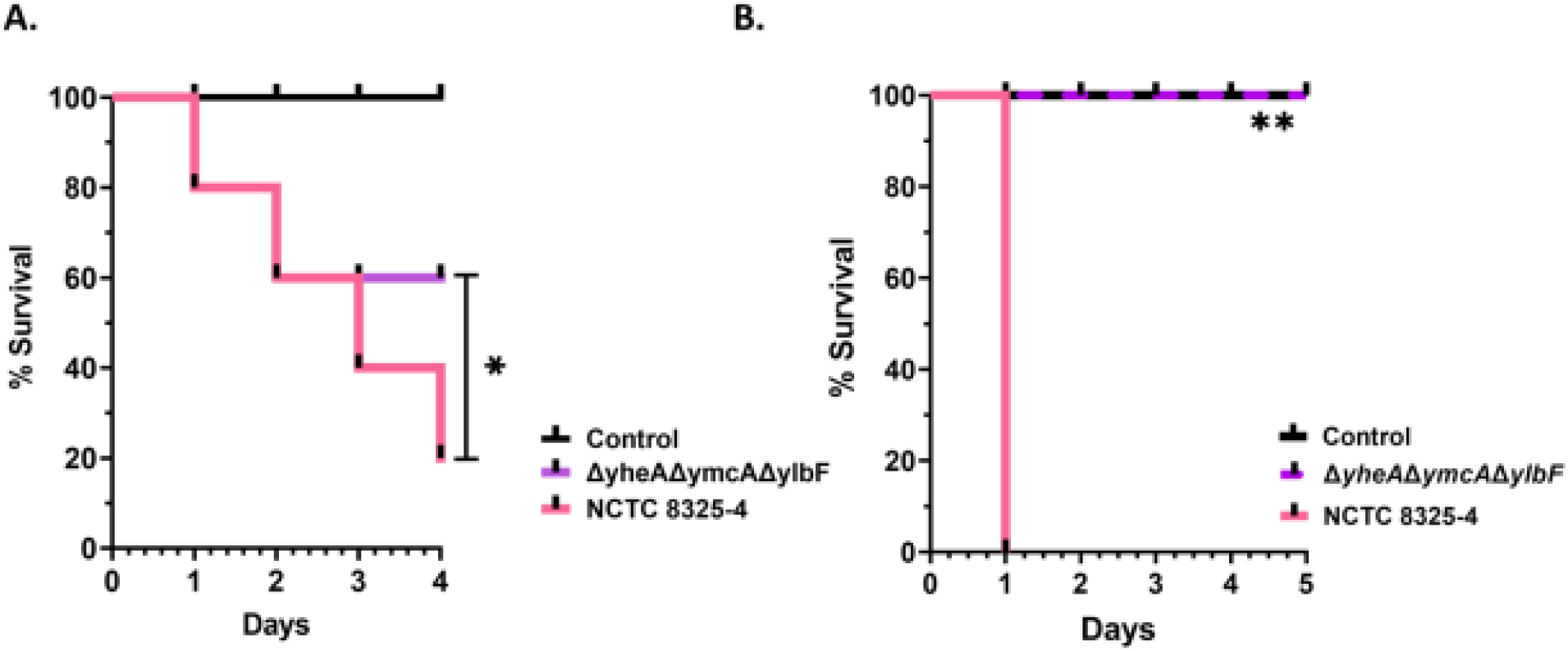
Survival analysis of in vivo models infected with *S. aureus*. **A.** Probability of larvae survival (n = 10 for each group) was evaluated over 4 days following inoculation. The larvae were inoculated with 5 µl of 1×10^5 CFU of the *S. aureus* NCTC8325-4 and *qrp/yheA*, *ymcA* and *ylbF* deletion strain (OD 0.1). Statistical analysis was performed using log-rank test *P < 0.05. This assay was performed in three independent experimental replicates. **B.** Probability of mice survival (n = 5 for each group) was evaluated over 5 days following intraperitoneal inoculation with 150 µl of 5×10^8 CFU of the *S. aureus* NCTC8325-4 and *qrp/yheA*, *ymcA* and *ylbF* deletion strain (OD 0.7). Statistical analysis was performed by log-rank (Mantel-Cox) test **P < 0.01. This assay was performed once. Δ*yheA=*Δ*qrp/yheA*. The inoculation with saline solution (0.98%) was used as a control.

The mice were injected intraperitoneally (i.p.) with 150 µl containing 5×10^8 CFU of the mutant strain Δ*qrp*/*yheA*Δ*ymcA*Δ*ylbF and* the wild-type NCTC8325-4 strain (22–24). The mice inoculated with the mutant strain Δ*qrp*/*yheA*Δ*ymcA*Δ*ylbF* showed similar results to the negative control, with 100% survival over the 5-day observation period. In contrast, injection of the wild-type strain NCTC8325-4 resulted in all infected mice developing a lethal infection within 24 h (Figure 5B). The changes in survival rates were visibly different between the wild-type and triple deletion strains, and significantly reduced in 100% (P < 0.01: p= 0,0009). Both in vivo models showed that the deletion of Com_YlbF (Qrp/YheA, YmcA, and YlbF) proteins in *S. aureus* increases significantly the survival rate of larvae and mice.

## DISCUSSION

In the present study we reveal some roles that Qrp/YheA, YmcA, and YlbF proteins may have in *S. aureus*, they seem be participating in a RNase-Y involved way, necessary to stimulate the adequate production of virulence factors such as hemolysins, proteases, adhesion proteins, and PIA/PNAG. Most of the studies have been focused on deciphering the role of the proteins YmcA and YlbF in *B. subtilis*, showing that these are necessary for the K-state, sporulation, and normal biofilm formation (7, 8, 10, 11, 25). However, in *S. aureus*, the knowledge on these Com_YlbF proteins is scarce. Only it has been reported that the full *ylbF* gene deletion affects the enzymatic activity of the endoribonuclease RNase-Y (11).

Additionally, we examined in both *G. mellonella* and murine peritonitis models the effect of Qrp/YheA, YmcA and YlbF proteins in *S. aureus* and the possible relationship of the expression of *S. aureus* virulence factors during infection. Notably, the triple mutant strain had also attenuated virulence in both *G. mellonella* and mice infection assays perhaps due to lack of bacterium ability to generate some virulence factors. To highlight, similar results were reported by Kaito *et a*l. in 2005, when the *rny*/*cvfA* gene (encoding RNase-Y protein) was disrupted in the CK501 strain of *S. aureus*, a decreased secretion of exotoxins, reduced expression of RNAII, RNAIII, and other virulence genes, and virulence attenuation in silkworm infection model was observed (26). Several studies have showed the important role that the diverse virulence factors have during the course of Staphylococcal infection and their ability to cause disease in human patients and animal models-infection (27). The survival results suggest that the mRNA levels regulated by RNase-Y in the mutant strain could be related to low in larvae or not in mice mortality. Although *G. mellonella* lacks an adaptive immune system, it has been an informative model for pathogen-host interactions, since in its innate system (cellular and humoral response), hemocyte immune cells resemble mammalian neutrophils (28, 29). We hypothesize that Com_YlbF proteins could be involved in systemic involvement in an infectious model of *G. mellonella*. This is supported by the increased survival time observed in larvae infected with the mutant strain of *S. aureus*, compared with the WT, possibly the decreased expression of pathogen virulence factors such as serine proteases and capsular polysaccharide proteins, could stimulating the cellular and humoral response.

In addition, the transcriptomic results showed the most relevant number of altered transcripts were in the category ‘virulence’ (5% decreased; 2% increased), with significantly deregulated genes, mainly associated with regulators such as RNAIII and the *agr* accessory gene regulator system (agrA), or virulence factors and the biofilm formation process, including serine proteases, capsular polysaccharide biosynthesis proteins, hemolysins, phenol-soluble modulins, the MSCRAMM and SERAM group. Our transcriptional analysis also identified a large decrease in the *spl* operon in the triple mutant (between 7-9fold). Interestingly, the expression of serine proteases encoding in this operon (that has only been found in *S. aureus*) has been associated with affectations of the host immune responses. For instance, Paharik *et al.*, examining the behavior of a *spl* mutant in a rabbit pneumonia model, revealed a complex virulence phenotype, where the Spl absence did not produce attenuated lethality but was able to induce lung-specific severe damage. In addition, they also found that the SplA was able to cleave human mucin 16 (first host substrate identified for an Spl protein), an important protect protein of the epithelia (30). This evidence suggests the role the *spl* operon as a virulence factor in the *S. aureus* disease and might correlate to the no mortality mice found by us.

Other groups of genes playing an important role in the virulence, resistance, and survival of *S. aureus* are genes encoding capsular polysaccharide biosynthesis proteins (CP), major factors in bacterial evasion of the host immune defenses (31, 32). The *S. aureus* CPs have been shown to possess antiphagocytic properties, allowing the bacterium to persist in the blood and tissues of infected hosts (33). Iqbal Z *et al*. described the transcriptome of a clinical strain derived from animals, compared it with a human *S. aureus* virulent strain, and reported a significant upregulation in *cap* genes, highlighting their pathogenicity in a murine model (34). Similarly, our transcriptional analyses of the Com_YlbF mutant showed that the cap operon expression levels were significantly decreased (between 1.5 and 2-fold), suggesting favorable conditions for phagocytosis. This may enhance the cellular defense response by facilitating the internalization and destruction of the pathogen by hemocyte immune cells.

*Staphylococcus aureus* strains encode between 16 and 17 different two-component systems (35–37). Some of them were affected in our transcriptional analyses such as *LytRS* (murein hydrolase activity), *VraSR* (cell wall biosynthesis), *saeRS* (secreted factors mainly involved in immune evasion), and *arlRS*; but with deregulation in *KdpED* (potassium transport) and *agrAC* (cell wall protein synthesis, detection of quorum sensing). The best-studied of these regulatory systems is the accessory gene regulator *agr* which acts as a master virulence regulator (38), as its importance in biofilm formation in both *S. aureus* and *S. epidermidis* (39, 40). Although numerous biofilm-related genes were repressed, many genes did not pass the threshold; but we identified the greatest transcriptional impact associated with this master regulator as the SarA protein family of transcriptional regulators (SarARXZ, ArlRS, CcpA, CodY, Rot, SrrAB), alternative sigma factor (SigB) (Table S3)(41–44).

The Com_YlbF proteins (YmcA and YlbF) gained attention when its essential role in the biofilm production in *B. subtilis* was discovered (9). Then, we assessed whether this same impact could be seen in *S. aureus*. Two alternative biofilm synthesis mechanisms that allow to *S. aureus* to adhere to and colonize different surfaces have been described. The *ica*-dependent pathway based on a PIA/PNAG production pathway under osmotic stress conditions (NaCl) (5, 45–47) and the *ica*-independent pathway lacking to the ability to express a variety of adhesion proteins under conditions of acidity (glucose). Here, we have found that single, double, and triple mutant strains do not completely null the biofilm formation, but whether to reduce it, with a decrease differently the PIA production and PIA-dependent biofilm. However, a marked reduction of hemolytic ability was observed.

One of the main attributes of *S. aureus* is the ability to evade both innate and adaptive immune responses involving several virulence factors, including exotoxins (48). *In vitro* and *in vivo* animal studies have shown that pore-forming toxins (PFTs) are the main virulence factors involved in the pathophysiology of Staphylococcal infections, however, their regulatory mechanisms still need to be further explored. To confirm the hypothesis of com_ylbF proteins domain and their participation in virulence, we evaluated the triple mutant strain in a qualitative and quantitative hemolytic assay using sheep red blood cells, demonstrating their high participation in virulence hemolytic capacity of *S. aureus*. Our data suggest that the Qrp/YheA, YmcA, and YlbF proteins act by modulating the virulence and pathogenicity in *S. aureus*. Since a reduction of biofilm coincides with reduced activity in hemolysis; it is an aspect associated with the contribution to biofilm integrity, a model proposed by Graf *et al.*, who discovered how various secreted virulence factors such as hemolysins, leukotoxins, lipases, and ribosomal proteins are key players with a moonlighting function in the model of biofilm integrity (49).

In the gene expression processes, ribonucleases (RNases) are the proteins responsible for post-transcriptional regulation of RNA maturation (50, 51), with RNase-Y being the main ribonuclease implied the degradation or stability (in several cases) of RNA transcripts (14, 51, 52). Regarding to *S. aureus,* an ortholog of RNase-Y (*rny*/*cvfA*) was discovered by Kaito *et al*. in 2005 (26, 53) but, to date, its characterization is very limited. The few studies carried out have shown that at least 99 genes could be processed for this endoribonuclease, including mRNA, tRNA, and non-coding RNAs (11). The enzyme participates more frequently in the degradation of many RNAs but also in the stabilization of others (52, 54), and their deletion causes a decrease in the virulence of the bacteria *in vivo* models (decreased hemolysis, adhesion and activation of some metabolic pathways) (20, 55, 56). To highlighting, some studies carried out by Wolz lab have found that RNase Y is important for the stabilization of the immature transcript of the operon SaePQRS and other virulence-related small RNAs (20, 55, 57). The operon SaePQRS is a two-component system that acts as a global regulator of the expression of major virulence genes in *S. aureus*, including hemotoxins (58–60).

In *B. subtilis*, an example of a multigene transcript that undergoes endoribonuclease-dependent maturation is the *cggR-gapA* glycolysis operon (11). With Rend-seq assay in *S. aureus*, DeLoughery *et al*., demonstrated that generation of one of the three mRNA isoforms of the *cggR*(*gapR*)-*gapA* operon requires the YlbF ortholog (61–63). Recently, Le Scornet *et al*. identified critical factors for precise and efficient RNA cleavage by RNase Y (for instance a secondary structure a few nucleotides downstream of the RNase y-cleavage site)(64). Our results in *S. aureus* indicated that in the triple deletional strain the transcript of the *cggR-gapA* operon is not adequately processed; suggesting a lower activity of RNase-Y ribonuclease. As it was previously mentioned, this change in the regulation of the *cggR-gapA* operon in *S. aureus* is likely a consequence of an increase in the half-life of the co-transcript as has also been reported in *B subtilis* (57, 65).

Taking together our results, it is plausible to suggest that the reduced virulence found in the triple mutant strain could be explained for an alteration in the RNase-Y activity, which in turn affects the stabilization of operon SaePQRS and other small RNAs, producing a reduced secretion of virulence factors. The interest in exploring the role of Com_YlbF proteins has been increasing, since phylogenetic analyses have suggested these proteins have co-evolved among firmicutes in the Bacilli order, especially in *Bacillus subtilis* (12). In this sense, our study shows the importance of this new family of small proteins, which, although are not essential for the survival of the bacterium, whether seem be involved in the regulation of its biofilm formation, hemolysis, and virulence.

## MATERIALS AND METHODS

### Bacterial strains, plasmids, culture conditions and oligonucleotides

Bacterial strains and plasmids used in this study are listed in Table S1. *Escherichia coli* strains were grown in Luria Bertani broth (LB) (BD Difco™) and *S. aureus* strains were grown in Trypticase Soy Broth (TSB) (BD Difco™). The Media were supplemented when appropriate with 10 μl/ml erythromycin, 100 μg/ml ampicillin, 80 μg/mL 5-bromo-4-chloro-3-indolyl-β-D-galactopiranósido (X-Gal), 0.25% wt/vol glucose (TSB_G_) or NaCl 3% (TSB_N_). Oligonucleotides are listed in Table S2 and were synthesized by the Oligonucleotide Synthesis Service (Macrogen Inc., Seoul, Korea).

### DNA manipulations and bacterial transformation

Plasmids were purified using the NucleoSpin Plasmid miniprep kit (Macherey-Nagel) according to the manufacturer’s protocol. FastDigest restriction enzymes and a Rapid DNA ligation kit (Thermo Scientific) were used according to the manufacturer’s instructions. Plasmids were transformed into *E. coli* IM01B strain and *S. aureus* by electroporation. Staphylococcal electrocompetent cells were generated as has previously been described (66, 67). Deletion mutants were generated via allelic replacement using the vector pMAD and conditions described by Arnaud, M *et al.*, and Valle, J., *et al.* (68, 69).

### Allelic exchange of chromosomal genes

To construct the deletions, two fragments of approximately 500 bp that flanked the left and the right of the sequence targeted for deletion were amplified by PCR using the A, B C, and D corresponding oligonucleotides (Table S2). Oligonucleotides to overlap have a 20-base complementary region to allow the products of the first PCR to anneal at their overlapping region and a second PCR was performed to obtain a single fragment. The fusion products were purified and cloned into the plasmid pJET® (Thermo Scientific). The fragment was then cloned into the polylinker sites of the shuttle plasmid pMAD (68) using the corresponding restriction enzymes (Table S2). The resulting pMAD (pMADr), was transformed into *E. coli* IM01B by electroporation. Plasmid pMADr contains a temperature-sensitive origin of replication and an erythromycin resistance gene. The plasmid was integrated into the chromosome through homologous recombination growing bacteria at the nonpermissive temperature (42°C) in the presence of erythromycin. From the 42°C plate, one to five colonies were picked into 10 ml of TSB and incubated for 2 -3 days at 28°C. Tenfold serial dilutions of this culture in sterile TSB were plated on TSA containing X-Gal. White colonies, which no longer contained the pMAD plasmid, were tested to confirm the replacement by PCR using E and F corresponding oligonucleotides (Table S2) and by sequencing.

### *RNA isolation, construction* of cDNA libraries and sequencing by Illumina

A bacterial pre-inoculum was prepared for the strains *S. aureus* NCTC8325-4 and *qrp/yheA*, *ymcA,* and *ylbF* mutant. They were cultivated overnight at 37 °C without shaking in TSB supplemented with glucose (0,25% p/v; TSB_G_). Then for RNA isolation, cultures were incubated with shaking until the mid-exponential phase at an OD600 of ∼0.7. RNA was purified using either RNeasy Mini Kit (Qiagen) for RNA-seq and RT-PCR. The quality and quantity of the isolated RNA were determined by agarose gel electrophoresis and confirmed by measuring the absorbance at 260 nm using a Nanodrop spectrophotometer NP80 (Implen GmbH). The RNA isolation was performed on three replicates, in two independent experiments. All samples were submitted to the TruSeq total RNA library construction for microbe with Ribo-zero kit previously to be sequenced using the NovaSeq 6000 (Illumina, CA) platform at Macrogen.

### Differentially Expressed Genes Identification and Annotation

Raw reads were generated from image data and stored in FASTQ format. Raw data were filtered to remove adaptor-contaminated and low-quality sequences and obtain clean reads. Clean reads were quality examined by Trimmomatic program V. 0.39 (Phred score ≤ 20) and aligned to the reference genome of *S. aureus* NCTC 8325 (NCBI Reference Sequence: NC_007795.1) using HiSat2 v. 2.1.0. Gene coverage was calculated by the percentage of genes covered by reads FPKMs (Fragments Per Kilobase of transcript, per Million mapped reads) and gene functional annotation was performed through Cuffnorm program included in Cufflinks v2.2. The differentially expressed genes (DEGs) were identified using R DeSeq2. The genes with an P value < 0.05 and adjusted |log2(fold change) | > 1,5 and < -1,5 were identified as DEGs, compared with the repository of the *Staphylococcus aureus* research & annotation community (AureoWiki) (70).

### Biofilm formation assay

The biofilm formation assays were performed in microtiter wells as previously described (17, 71). *S. aureus* strains were cultivated overnight in TSB supplemented with glucose (0,25% p/v; TSB_G_) to evaluate under acid conditions and with NaCl (3% w/v; TSB_N_) to evaluate under osmotic stress conditions. Optical density (OD600) was adjusted at 0.1 absorbance units with 0.98% Saline Solution, and 5 µl was inoculated with 195 µl sterile TSB_G_ or TSB_N_ in 96-well polystyrene microtiter plates (NEST). After culturing for 24 hours at 37°C, the wells were gently washed twice with 200 ml of sterile phosphate-buffered saline (PBS). Plates were air-dried and remaining cells adsorbed to the surface of individual wells were stained with Crystal violet-stained. The cells were quantified by solubilizing the dye with 200 μl of ethanol-acetone (80:20, vol/vol) and determining the optical density at 595 nm (OD595). Each assay was performed in triplicate and repeated at least three times.

### PIA/PNAG detection

*S. aureus* strains were cultivated overnight in TSB supplemented with glucose (0,25% p/v; TSBG) and incubated at 37 °C with shaking, as described by Gerke *et al* (72). PIA/PNAG production was detected with an anti-*S. aureus* PIA/PNAG (D. McKenney, Boston, USA) diluted 1:10,000 (16, 17, 73) on a nitrocellulose membrane using the Bio-Dot Microfiltration kit (Bio-Rad). Bound antibodies were detected with a peroxidase-conjugated Goat anti-rabbit immunoglobulin G (IgG) antibody (Jackson Immuno Research Laboratories, Inc., PA, USA) diluted 1:5,000, and Luminol Western Blotting Reagent, from the Chemiluminescent Nucleic Acid Detection Kit (Thermo Scientific).

### Hemolysis assay

The hemolytic capability was evaluated using qualitative and quantitative analysis. For the qualitative assay, overnight cultures of *S. aureus* strains were prepared by inoculating 3 mL of TSB with a single colony of each strain independently and incubated at 37 °C without shaking. The culture absorbance was measured at 620 nm and adjusted in saline solution at OD 0.4 (McFarland 2 standard). A drop (2 µL) of this adjusted resuspension of each *S. aureus* strain was put in blood agar plates (5% sheep blood in tryptic soy agar (TSA), Hardy Diagnostics, CA, USA) at separation distances of 3 cm. Plates were incubated at 37 °C and the hemolytic activity was recorded at 24 h. The presence of a distinct translucent halo around the inoculum site was considered indicative of positive hemolytic activity. The diameters of the zones of lysis and the colony were measured using a computerized image analysis system to estimate the hemolytic area.

For hemolytic capability quantitative assays, overnight cultures of *S. aureus* strains were prepared by inoculating 3 mL of TSB with a single colony of each strain independently and incubated at 37 °C without shaking. The culture absorbance was measured at 620 nm and adjusted with 0.98% saline solution at OD 0.4 (McFarland 2 standard) and were performed in sheep blood. Sheep blood was defibrinated by rinsing with saline solution to obtain a solution of blood 3%. 270μl of prepared blood was mixed with 30μl of each previously adjusted culture of *S. aureus* strains in 96-well plates. Plates were incubated at 37 °C without shaking. Hemolysis was visually manifested as the profit of the red/orange color detected by absorbance measured at 540 nm, representing the release of hemoglobin from hemolyzed RBCs. The positive control for lysis was 1% SDS and the negative control was 0.98% saline solution. The assay was performed three times in three independent experiments.

### Analysis of post-transcriptional processing

To determine the possible involvement of Com_YlbF proteins (Qrp/YheA, YmcA, YlbF) with processes at the post-transcriptional processes, the processing of the *cggR*-*gapA* operon transcript was evaluated (74, 75). RNA was purified using RNeasy Mini Kit (Qiagen), and the quality and quantity of the isolated RNA was determined as previously described. Contaminating DNA in the RNA preparations was removed using “RNase-free DNase I” (Thermo Scientific). Purified RNA samples were converted to cDNA using Moloney murine leukemia virus (M-MLV) reverse transcriptase enzyme (Promega), following the manufacturer’s instructions. cDNA (50 ng) was then used directly used as template for RT-PCR with respective primers (Table S2). The RT-PCR products were examined by agarose gel electrophoresis and stained with ethidium bromide. The gene expression was compared between *S. aureus* NCTC8325-4 and its respective single, double, and triple *qrp/yheA*, *ymcA,* and *ylbF* mutant strains. The relative densitometry analysis was performed using ImageJ software.

### Generation of transcriptional fusions with Gfp

To obtain transcriptional fusions the promoter of *hlgC* was amplified using the respective primers (Table S2) and cloned into pCN52 plasmid (Charpentier et al., 2004), generating plasmids pCN52-*phlgC*_*gfp* plasmids. Plasmids were transformed into *S. aureus* NCTC8325-4 and *qrp/yheA*, *ymcA,* and *ylbF* mutant strains to test for *hlgC* promoter activity. The strains were cultivated overnight at 37 °C with shaking in TSB. Cells were pelleted by centrifugation at 4500 rpm for 30 min at 4 °C, washed with 1 mL of PBS, suspended in 400 μL PBS, and lysed using a FastPrep-24TM 5G homogenizer (MP Biomedicals, LLC, Irvine, CA, USA). Total protein extracts were recovered, and analyzed by SDS-PAGE and Western blot as described by Morales-Laverde *et al*. (76). GFP was detected using an anti-green Fluorescent Protein (GFP) Monoclonal antibody (SIGMA®) at a 1:2,500 dilution in 0.1% PBS-Tween 5% skimmed milk. As secondary antibody, ECL Mouse IgG conjugated with peroxidase (HRP) (Amersham®) was used, diluted 1:5000 in 0.1% PBS-Tween 5% skimmed milk. The Chemiluminescent Nucleic Acid Detection Kit (Thermo Scientific) was used as a substrate.

### Galleria mellonella survival assay

A total of 90 *Galleria mellonella* larvae, 30 per experiment (n = 10 for each group) were used and randomly assigned to a specific experimental group. The groups evaluated were: negative control (0.98% Saline Solution), positive control (wild-type strain NCTC8325-4), and mutant strain (Δ*qrp*/*yheA*Δ*ymcA*Δ*ylbF*). The larvae, corresponding to the sixth instar, healthy and weighing between 200-300 milligrams, with no specific exclusions, were obtained from the Biological Control Laboratory at the Department of Biology, Pontificia Universidad Javeriana de Bogotá (77, 78). The strains *S. aureus* NCTC8325-4 and Δ*qrp/yheA*Δ*ymcA*Δ*ylbF* mutant were prepared independently by inoculating 3 mL of TSB with a single colony of each strain independently and incubated at 37 °C without shaking. The optical density of the cultures in TSB was measured at 600 nm and adjusted with 0.98% saline solution to OD 0,1 until complete the same inoculum of 1×10^^5^ CFU (77, 79). Before handling the larvae, disinfection washes were carried out with hypochlorite (0.1%) and sterile distilled water. Aliquots of 5 µl of each bacterial dilution were injected into the second middle left proleg of the *G. mellonella* larvae. Control larvae were injected with the same volume (5 µl) of saline solution to monitor any problem associated with the injection process. Following injection, larvae were placed in glass petri dishes and stored in the dark at 28°C for 4 days. For each group of larvae their survival and appearance were evaluated in 24 hours intervals. Three independent experimental replicates were performed in triplicate. Larvae were considered dead if they did not respond to touch by moving if they did not respond to touch by moving (80). Along with the survival test, the melanization changes were recorded according to the scoring system published by Champion *et al.* (80).

The larvae survival experiments were advised by Dr. Adriana Sáenz Aponte, Master of Agricultural Sciences - Entomology. the experiments were conducted in accordance with the ethical considerations of the management of invertebrate animals (81) and in accordance with the protocols described in the literature (77).

### Mice survival assay

The experiments were performed using female BALB/c mice, aged 5–6 weeks (17–22 g), with no specific exclusions, purchased from the Central Bioterium of the Universidad Nacional de Colombia. The peritonitis model in BALB/c mouse strains was adapted from models previously described in the literature (22–24). A total of 15 mice, (n = 5 for each group evaluated), were used and randomly assigned to a specific experimental group. The groups evaluated were: negative control (0.98% Saline Solution), positive control (wild-type strain NCTC8325-4) and mutant strain (Δ*qrp*/*yheA*Δ*ymcA*Δ*ylbF)*. For the bacterial inoculum, *S. aureus* strains were prepared by cultivating a single colony of each strain in 5 mL of TSB and incubating at 37 °C without shaking. The optical density of the cultures in TSB was measured at 600 nm and adjusted to mid-exponential phase OD 0.7 with 0.98% saline solution. The mice were intraperitoneally (i.p.) injected with 150 µl 5×10^8 CFU of the *S. aureus* NCTC8325-4 and Δ*qrp*/*yheA*Δ*ymcA*Δ*ylbF* mutant strain according to the experimental group.

The animals were housed in the animal facility at Universidad Nacional de Colombia, in an isolated room meeting all the necessary requirements for proper animal care. Groups of five female mice were kept in appropriately sized ventilated cages (500 cm² of floor space, model NexGen500, Allentown), covered with sterile wood chip (Aspen Chip and Lab Grande Aspen, NEPCO). The top of the cage held an external plastic 250 mL water bottle and a Whatman filter that allowed clean air exchange and protected the food (placed in a half pocket wire bar lid) with all the nutritional requirements for mice’s survival. The cage had an enrichment (60 mm x 78 mm) for mice entertainment. All home cages were properly labeled. During the experimental period, the animals moved to clean cages with new food and water once a day and were kept in the experimental 22°C, on a 12-h light-dark cycle.

The health status of the postinfection mice was observed for a maximum period of 5 days to minimize suffering, according to the scoring system published by Carstens, *et al.* (82). If mice showed severe signs of illness, they were euthanized following Guidelines-Humane Endpoints for Research, Teaching and Testing Animals (83). The mouse survival experiments were directed by Dr. Myriam L. Velandia-Romero, author, and in charge of the animals throughout the experimental process. who is certified by the Colombian Association for the Science and Welfare of Laboratory Animals (ACCBAL), the Institutional Committees for the Care and Use of Animals (CICUA) of the Pontificia Universidad Javeriana and the Universidad de los Andes, and the Research Ethics Committee of the National University of Colombia.

All experiments were conducted in accordance with the ethical principles of animal experimentation set by ICLAS, the Comité Institucional de Ética en Investigación of Universidad El Bosque No. 017 of 2022, the International Council for Laboratory Animal Science, and Resolution No. 008430 of 1993 from the Ministry of Health of the Republic of Colombia.

### Image quantification and statistics

GraphPad Prism software, version 9.0, was used to perform the non-parametric statistical analyses showing the behavior of the variables through histograms and curves. Densitometry analysis were performed using ImageJ software (http://imagej.nih.gov/ij/). Rectangles were drawn around different bands and the average pixel intensity was measured. The same size of the rectangle was used to quantify the bands being compared, and the density of the background of the same size was subtracted from the measurement. Two-tailed P values were determined based on unpaired t-tests or log-rank test. In images, statistical significance is indicated as *P < 0.05, **P < 0.01.

## Supporting information

Table S3_Transcriptomic analysis of the deletion of the qrp_yheA, ymcA and ylbF genes in NCTC8325_4 of Staphylococcus aureus

## ACKNOWLEDGMENTS

We thank Dr. Miguel Otero for his invaluable support. We also extend our gratitude to Dr. Adriana Sáenz Aponte from the Biological Control Laboratory, Department of Biology, Pontificia Universidad Javeriana de Bogotá, for providing the *G. mellonella* larvae and sharing her expertise on the model. We acknowledge Dr. Jesus Alfredo Cortes, Director of Bioterio Central at Universidad Nacional de Colombia, for supplying the BALB/c mouse strains and sharing his expertise in the model. Additionally, we acknowledge Dr Nancy Mugridge Senir, for your review and editing of the manuscript.

## Author Contributions

Zayda Corredor-Rozo, Formal analysis, Investigation, Murine model, project administration, writing—original draft preparation, Writing – review and editing. Dr. Ricaurte Marquez-Ortiz Data curation, Software, Formal analysis, Writing – review and editing. Deisy Abril-Riaño, Murine model, review. Dr. Johana Madroñero, Data curation, Software, Formal analysis, Writing – review and editing. Dr. Luisa F. Prada Murine model, review. Dr. Natasha Vanegas-Gomez, Writing – review and editing. Dr. Begoña García, Investigation, Formal analysis, Writing – review and editing. Dr. Maite Echeverz, Investigation, Formal analysis, review, and editing. Dr. Myriam L. Velandia-Romero, director and validation of the Murine model, Formal analysis, Writing – review and editing. María Angélica Calderón-Peláez, Murine model, Formal analysis, review. Dr. Jacqueline Chaparro-Olaya, Murine model, Writing – review and editing. Liliana Morales, Murine model, review. Carlos Nieto-Clavijo, Murine model, review. Dr. Javier Escobar-Perez, Director, Conceptualization, Supervision, Validation, Investigation, project administration, funding acquisition, writing—original draft preparation, Writing – review, and editing. All authors have read and agreed to the published version of the manuscript.

## Tables and supplementary figures

**Table S1.** Bacterial strains and plasmids were used in this study.

| Strains | Relevant properties | Reference |
| --- | --- | --- |
| ***Staphylococcus aureus*** |  |  |
| 8325-4 | *Staphylococcus aureus* NCTC 8325-4 prophage curing and harbors an 11 bp deletion in the *rsbU* gene | (84) |
| $\Delta yheA$ | *S. aureus* 8325-4 with *qrp/yheA* gene deleted | This study. |
| $\Delta yheA \Delta ylbF$ | *S. aureus* 8325-4 with *qrp/yheA*, and *ylbF* genes deleted | This study |
| $\Delta yheA \Delta ymcA$ | *S. aureus* 8325-4 with *qrp/yheA*. and *ymcA* genes deleted | This study |
| $\Delta yheA \Delta ymcA \Delta ylbF$ | *S. aureus* 8325-4 with *qrp/yheA*, *ymcA*, and *ylbF* genes deleted | This study |
| 8325-4 pCN52-Pgfp | *S. aureus* 8325-4, with pCN52 and expression of GFP | This study |
| 8325-4 pCN52-PhlgC | *S. aureus* 8325-4, with pCN52 plasmid with the gamma hemolysin C subunit promoter with the GFP reporter gene. | This study |
| $\Delta yheA \Delta ymcA \Delta ylbF$ pCN52-Pgfp | *S. aureus* 8325-4 *qrp/yheA*, *ymcA* y *ylbF* mutant, with pCN52 and expression of GFP under the control of the respective promoter. | This study |
| $\Delta yheA \Delta ymcA \Delta ylbF$ pCN52-PhlgC | *S. aureus* 8325-4 *qrp/yheA*, *ymcA* y *ylbF* mutant, with pCN52 plasmid with the gamma hemolysin C subunit promoter with the GFP reporter gene. | This study |
| ***Escherichia coli*** |  |  |
| IM01B | *E. coli* K12 DH10B | (85) |
| Plasmids | Relevant properties | Reference |
| pCN52 | Plasmid for promoter activity coupled to the green fluorescent protein (GFP) reporter gene. | (86) |
| pMAD | Thermosensitive plasmid for homologous recombination in *Staphylococcus aureus*. | (68) |

**Table S2.**
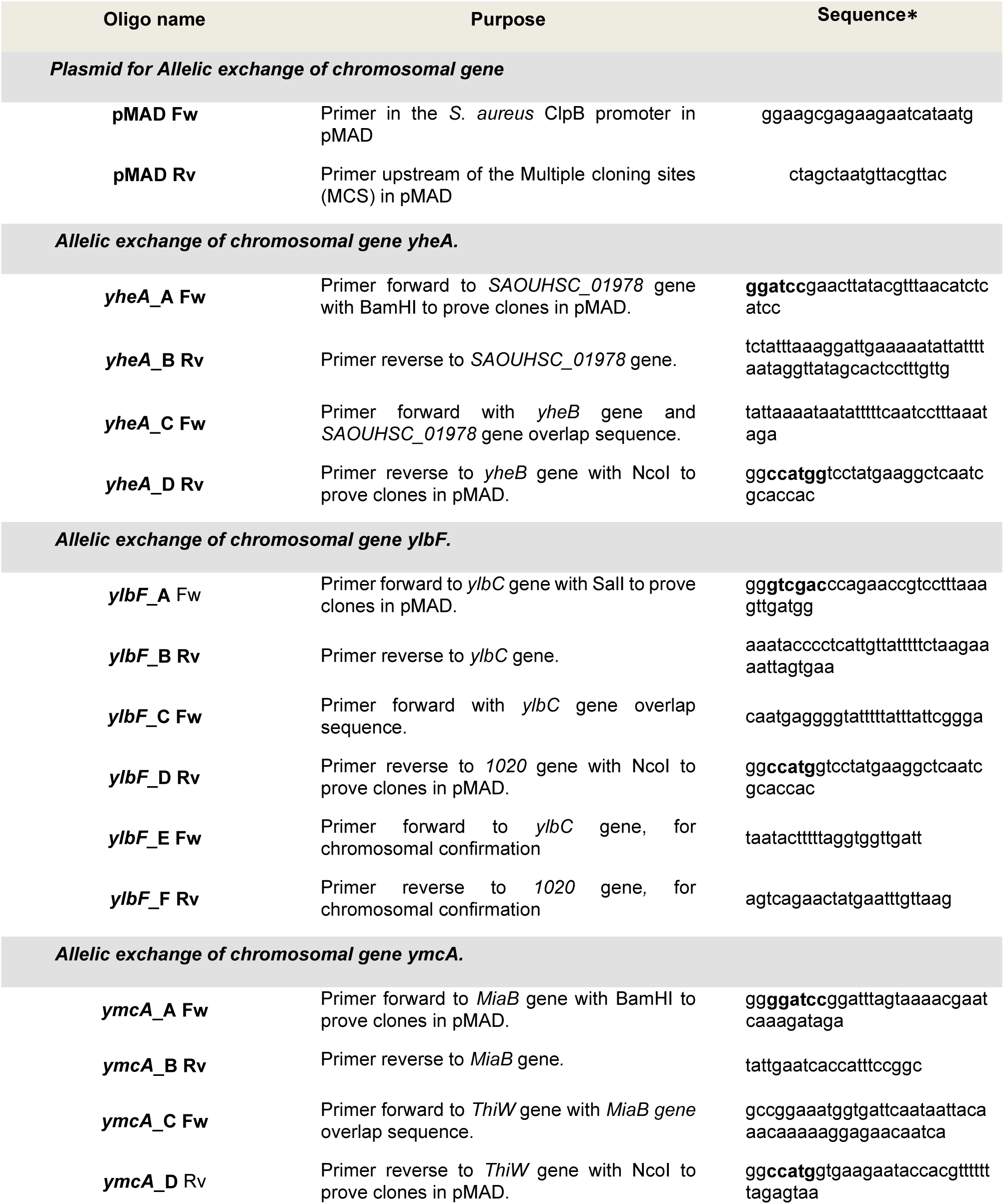
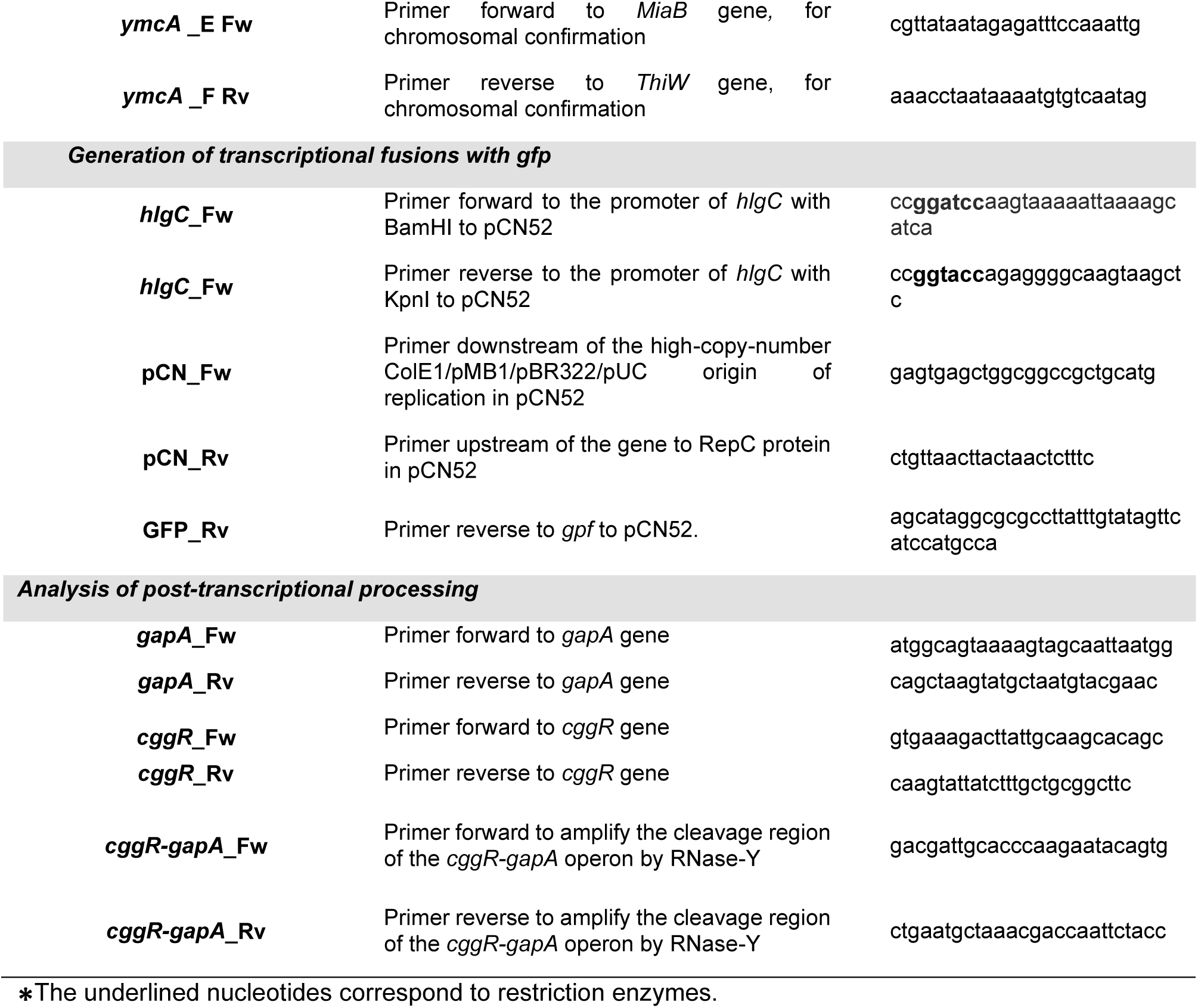
Oligonucleotides used in this study.

**Figure S1.**
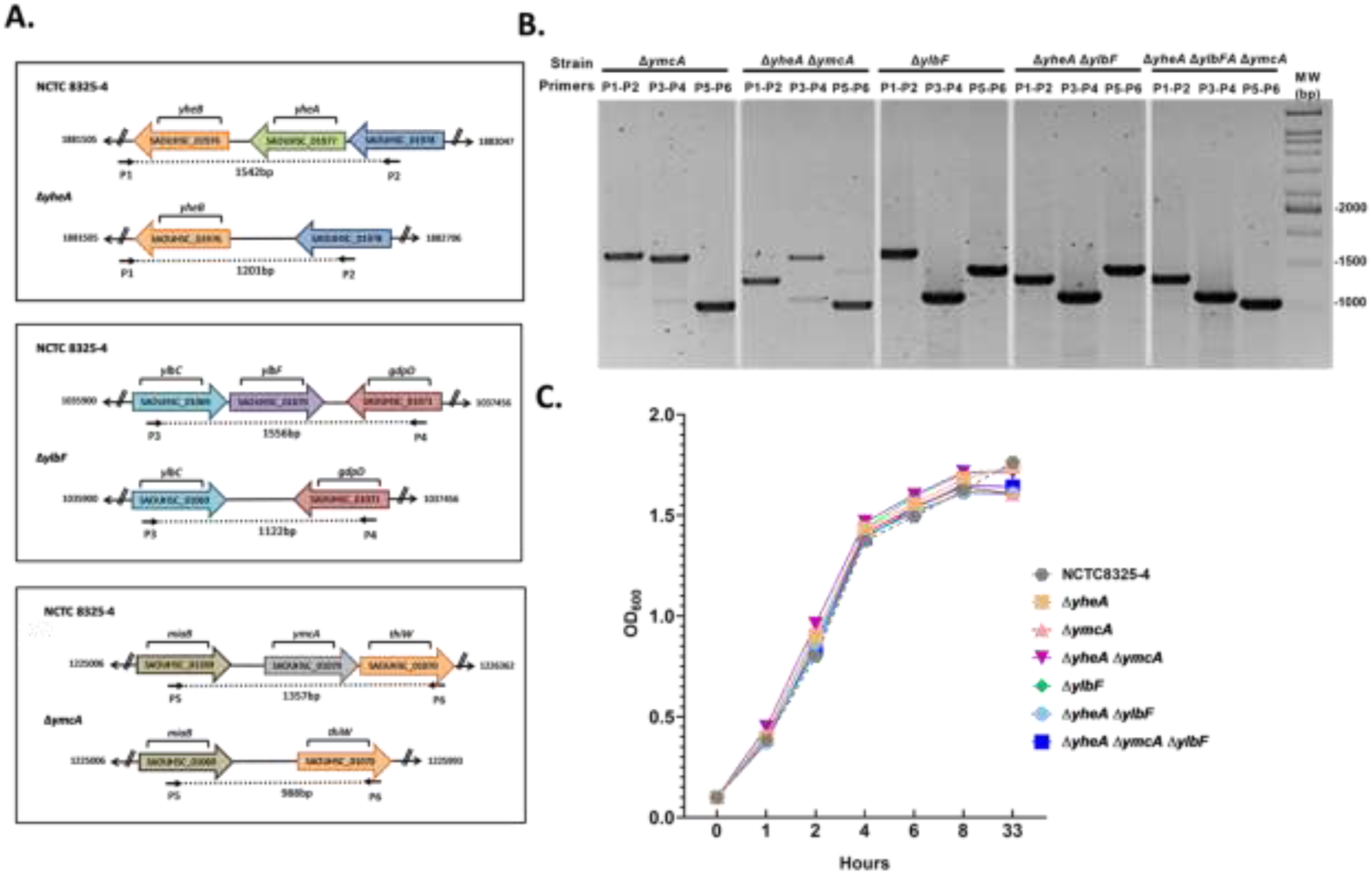
**A.** Diagram showing the positions of primers used for the identification of mutants in *S. aureus* NCTC8325-4. **B.** Confirmation of the deletion of the genes by agarose gel electrophoresis of PCR products Lane MW, 1000 bp marker. **C.** The *in vitro* growth curves of *S. aureus* NCTC8325-4 and mutation strains. The strains were cultivated in TSB overnight at 37°C. The bacterial solution was diluted to an optical density at 600 nm (OD600) of 0.1 and cultivated again. The OD600 was then measured at 1, 2, 4, 6, 8, and 33 h to draw the curves.

## REFERENCES

1. Tong SY, Davis JS, Eichenberger E, Holland TL, Fowler VG, Jr. Staphylococcus aureus infections: epidemiology, pathophysiology, clinical manifestations, and management. Clin Microbiol Rev. 2015;28(3):603–61.

2. Balasubramanian D, Harper L, Shopsin B, Torres VJ. Staphylococcus aureus pathogenesis in diverse host environments. Pathog Dis. 2017;75(1).

3. Koymans KJ, Vrieling M, Gorham RD, Jr., van Strijp JAG. Staphylococcal Immune Evasion Proteins: Structure, Function, and Host Adaptation. Curr Top Microbiol Immunol. 2017;409:441–89.

4. Foster TJ, Geoghegan JA, Ganesh VK, Hook M. Adhesion, invasion and evasion: the many functions of the surface proteins of Staphylococcus aureus. Nat Rev Microbiol. 2014;12(1):49–62.

5. Arciola CR, Campoccia D, Speziale P, Montanaro L, Costerton JW. Biofilm formation in Staphylococcus implant infections. A review of molecular mechanisms and implications for biofilm-resistant materials. Biomaterials. 2012;33(26):5967–82.

6. Escobar-Perez J, Ospina-Garcia K, Rozo ZLC, Marquez-Ortiz RA, Castellanos JE, Gomez NV. Identification and in silico Structural Analysis of the Glutamine-rich Protein Qrp (YheA) in Staphylococcus Aureus. The Open Bioinformatics Journal. 2019;12:18–29.

7. Tortosa P, Albano M, Dubnau D. Characterization of ylbF, a new gene involved in competence development and sporulation in Bacillus subtilis. 2000;35(5):1110–9.

8. Branda SS, Gonzalez-Pastor JE, Dervyn E, Ehrlich SD, Losick R, Kolter R. Genes involved in formation of structured multicellular communities by Bacillus subtilis. J Bacteriol. 2004;186(12):3970–9.

9. Carabetta VJ, Tanner AW, Greco TM, Defrancesco M, Cristea IM, Dubnau D. A complex of YlbF, YmcA and YaaT regulates sporulation, competence and biofilm formation by accelerating the phosphorylation of Spo0A. Mol Microbiol. 2013;88(2):283–300.

10. DeLoughery A, Dengler V, Chai Y, Losick R. Biofilm formation by Bacillus subtilis requires an endoribonuclease-containing multisubunit complex that controls mRNA levels for the matrix gene repressor SinR. Mol Microbiol. 2016;99(2):425–37.

11. DeLoughery A, Lalanne JB, Losick R, Li GW. Maturation of polycistronic mRNAs by the endoribonuclease RNase Y and its associated Y-complex in Bacillus subtilis. Proc Natl Acad Sci U S A. 2018;115(24):E5585–E94.

12. Tanner AW, Carabetta VJ, Martinie RJ, Mashruwala AA, Boyd JM, Krebs C, et al. The RicAFT (YmcA-YlbF-YaaT) complex carries two [4Fe-4S](2+) clusters and may respond to redox changes. Mol Microbiol. 2017;104(5):837–50.

13. Khemici V, Prados J, Linder P, Redder P. Decay-Initiating Endoribonucleolytic Cleavage by RNase Y Is Kept under Tight Control via Sequence Preference and Sub-cellular Localisation. PLOS Genetics. 2015;11(10):e1005577.

14. Shahbabian K, Jamalli A, Zig L, Putzer H. RNase Y, a novel endoribonuclease, initiates riboswitch turnover in Bacillus subtilis. The EMBO journal. 2009;28(22):3523–33.

15. Cue D, Lei MG, Lee CY. Genetic regulation of the intercellular adhesion locus in staphylococci. Frontiers in cellular and infection microbiology. 2012;2:38.

16. McKenney D, Pouliot KL, Wang Y, Murthy V, Ulrich M, Doring G, et al. Broadly protective vaccine for Staphylococcus aureus based on an in vivo-expressed antigen. Science. 1999;284(5419):1523-7.

17. Cucarella C, Solano C, Valle J, Amorena B, Lasa Ini, Penadés JRJJob. Bap, a Staphylococcus aureus surface protein involved in biofilm formation. 2001;183(9):2888–96.

18. Das S, Lindemann C, Young BC, Muller J, Osterreich B, Ternette N, et al. Natural mutations in a Staphylococcus aureus virulence regulator attenuate cytotoxicity but permit bacteremia and abscess formation. Proc Natl Acad Sci U S A. 2016;113(22):E3101–10.

19. Moriano A, Serra DO, Hoard A, Montana S, Degrossi J, Bonomo RA, et al. Staphylococcus aureus Potentiates the Hemolytic Activity of Burkholderia cepacia Complex (Bcc) Bacteria. Curr Microbiol. 2021;78(5):1864–70.

20. Marincola G, Schafer T, Behler J, Bernhardt J, Ohlsen K, Goerke C, et al. RNase Y of Staphylococcus aureus and its role in the activation of virulence genes. Mol Microbiol. 2012;85(5):817–32.

21. Lehnik-Habrink M, Lewis RJ, Mader U, Stulke J. RNA degradation in Bacillus subtilis: an interplay of essential endo- and exoribonucleases. Mol Microbiol. 2012;84(6):1005–17.

22. Rauch S, DeDent AC, Kim HK, Bubeck Wardenburg J, Missiakas DM, Schneewind O. Abscess formation and alpha-hemolysin induced toxicity in a mouse model of Staphylococcus aureus peritoneal infection. Infect Immun. 2012;80(10):3721–32.

23. Domenech A, Ribes S, Cabellos C, Dominguez MA, Montero A, Linares J, et al. A mouse peritonitis model for the study of glycopeptide efficacy in GISA infections. Microb Drug Resist. 2004;10(4):346–53.

24. Frimodt-Moller N. The mouse peritonitis model: present and future use. J Antimicrob Chemother. 1993;31 Suppl D:55–60.

25. Dubnau EJ, Carabetta VJ, Tanner AW, Miras M, Diethmaier C, Dubnau D. A protein complex supports the production of Spo0A-P and plays additional roles for biofilms and the K-state in Bacillus subtilis. Mol Microbiol. 2016;101(4):606–24.

26. Kaito C, Kurokawa K, Matsumoto Y, Terao Y, Kawabata S, Hamada S, et al. Silkworm pathogenic bacteria infection model for identification of novel virulence genes. 2005;56(4):934–44.

27. Kolar SL, Ibarra JA, Rivera FE, Mootz JM, Davenport JE, Stevens SM, et al. Extracellular proteases are key mediators of Staphylococcus aureus virulence via the global modulation of virulence-determinant stability. MicrobiologyOpen. 2013;2(1):18–34.

28. Sheehan G, Dixon A, Kavanagh K. Utilization of Galleria mellonella larvae to characterize the development of Staphylococcus aureus infection. 2019;165(8):863–75.

29. Ménard G, Rouillon A, Cattoir V, Donnio PY. Galleria mellonella as a Suitable Model of Bacterial Infection: Past, Present and Future. Frontiers in cellular and infection microbiology. 2021;11:782733.

30. Paharik AE, Salgado-Pabon W, Meyerholz DK, White MJ, Schlievert PM, Horswill AR. The Spl Serine Proteases Modulate Staphylococcus aureus Protein Production and Virulence in a Rabbit Model of Pneumonia. mSphere. 2016;1(5).

31. O’Riordan K, Lee JC. Staphylococcus aureus capsular polysaccharides. Clin Microbiol Rev. 2004;17(1):218–34.

32. Keinhörster D, Salzer A, Duque-Jaramillo A, George SE, Marincola G, Lee JC, et al. Revisiting the regulation of the capsular polysaccharide biosynthesis gene cluster in Staphylococcus aureus. Mol Microbiol. 2019;112(4):1083–99.

33. Weidenmaier C, Lee JC. Structure and Function of Surface Polysaccharides of Staphylococcus aureus. Curr Top Microbiol Immunol. 2017;409:57–93.

34. Iqbal Z, Seleem MN, Hussain HI, Huang L, Hao H, Yuan Z. Comparative virulence studies and transcriptome analysis of Staphylococcus aureus strains isolated from animals. Scientific reports. 2016;6:35442.

35. Haag AF, Bagnoli F. The Role of Two-Component Signal Transduction Systems in Staphylococcus aureus Virulence Regulation. Curr Top Microbiol Immunol. 2017;409:145–98.

36. Somerville GA, Proctor RA. At the crossroads of bacterial metabolism and virulence factor synthesis in Staphylococci. Microbiology and molecular biology reviews : MMBR. 2009;73(2):233–48.

37. Villanueva M, García B, Valle J, Rapún B, Ruiz de los Mozos I, Solano C, et al. Sensory deprivation in Staphylococcus aureus. Nature Communications. 2018;9(1):523.

38. Recsei P, Kreiswirth B, O’Reilly M, Schlievert P, Gruss A, Novick RP. Regulation of exoprotein gene expression in Staphylococcus aureus by agar. Molecular & general genetics : MGG. 1986;202(1):58–61.

39. Vuong C, Saenz HL, Götz F, Otto M. Impact of the agr Quorum-Sensing System on Adherence to Polystyrene in Staphylococcus aureus. The Journal of Infectious Diseases. 2000;182(6):1688–93.

40. Otto M. Staphylococcus aureus and Staphylococcus epidermidis peptide pheromones produced by the accessory gene regulator agr system. Peptides. 2001;22(10):1603–8.

41. Jenul C, Horswill AR. Regulation of Staphylococcus aureus Virulence. Microbiology spectrum. 2019;7(2).

42. Resch A, Leicht S, Saric M, Pásztor L, Jakob A, Götz F, et al. Comparative proteome analysis of Staphylococcus aureus biofilm and planktonic cells and correlation with transcriptome profiling. Proteomics. 2006;6(6):1867–77.

43. Resch A, Rosenstein R, Nerz C, Götz F. Differential gene expression profiling of Staphylococcus aureus cultivated under biofilm and planktonic conditions. Applied and environmental microbiology. 2005;71(5):2663–76.

44. Foulston L, Elsholz AK, DeFrancesco AS, Losick R. The extracellular matrix of Staphylococcus aureus biofilms comprises cytoplasmic proteins that associate with the cell surface in response to decreasing pH. mBio. 2014;5(5):e01667–14.

45. Valle J, Echeverz M, Lasa I. σ(B) Inhibits Poly-N-Acetylglucosamine Exopolysaccharide Synthesis and Biofilm Formation in Staphylococcus aureus. J Bacteriol. 2019;201(11).

46. Vergara-Irigaray M, Valle J, Merino N, Latasa C, García B, Ruiz de Los Mozos I, et al. Relevant role of fibronectin-binding proteins in Staphylococcus aureus biofilm-associated foreign-body infections. Infect Immun. 2009;77(9):3978–91.

47. Houston P, Rowe SE, Pozzi C, Waters EM, O’Gara JP. Essential role for the major autolysin in the fibronectin-binding protein-mediated Staphylococcus aureus biofilm phenotype. Infect Immun. 2011;79(3):1153–65.

48. Dinges MM, Orwin PM, Schlievert PM. Exotoxins of Staphylococcus aureus. Clin Microbiol Rev. 2000;13(1):16–34, table of contents.

49. Graf AC, Leonard A, Schäuble M, Rieckmann LM, Hoyer J, Maass S, et al. Virulence Factors Produced by Staphylococcus aureus Biofilms Have a Moonlighting Function Contributing to Biofilm Integrity. Molecular & cellular proteomics : MCP. 2019;18(6):1036–53.

50. Deutscher MP. The metabolic role of RNases. Trends in Biochemical Sciences. 1988;13(4):136–9.

51. Durand S, Gilet L, Bessières P, Nicolas P, Condon C. Three essential ribonucleases-RNase Y, J1, and III-control the abundance of a majority of Bacillus subtilis mRNAs. PLoS Genet. 2012;8(3):e1002520.

52. Cho KH. The Structure and Function of the Gram-Positive Bacterial RNA Degradosome. Frontiers in microbiology. 2017;8:154.

53. Kang SO, Caparon MG, Cho KH. Virulence gene regulation by CvfA, a putative RNase: the CvfA-enolase complex in Streptococcus pyogenes links nutritional stress, growth-phase control, and virulence gene expression. Infect Immun. 2010;78(6):2754–67.

54. Richards J, Liu Q, Pellegrini O, Celesnik H, Yao S, Bechhofer DH, et al. An RNA pyrophosphohydrolase triggers 5’-exonucleolytic degradation of mRNA in Bacillus subtilis. Molecular cell. 2011;43(6):940–9.

55. Marincola G, Wolz C. Downstream element determines RNase Y cleavage of the saePQRS operon in Staphylococcus aureus. Nucleic Acids Res. 2017;45(10):5980–94.

56. Mathy N, Hébert A, Mervelet P, Bénard L, Dorléans A, Li de la Sierra-Gallay I, et al. Bacillus subtilis ribonucleases J1 and J2 form a complex with altered enzyme behaviour. Mol Microbiol. 2010;75(2):489–98.

57. Bonnin RA, Bouloc P. RNA Degradation in Staphylococcus aureus: Diversity of Ribonucleases and Their Impact. International journal of genomics. 2015;2015:395753.

58. Geiger T, Goerke C, Mainiero M, Kraus D, Wolz C. The virulence regulator Sae of Staphylococcus aureus: promoter activities and response to phagocytosis-related signals. J Bacteriol. 2008;190(10):3419–28.

59. Adhikari RP, Novick RP. Regulatory organization of the staphylococcal sae locus. Microbiology (Reading, England). 2008;154(Pt 3):949–59.

60. Steinhuber A, Goerke C, Bayer MG, Döring G, Wolz C. Molecular architecture of the regulatory Locus sae of Staphylococcus aureus and its impact on expression of virulence factors. J Bacteriol. 2003;185(21):6278–86.

61. Commichau FM, Rothe FM, Herzberg C, Wagner E, Hellwig D, Lehnik-Habrink M, et al. Novel activities of glycolytic enzymes in Bacillus subtilis: interactions with essential proteins involved in mRNA processing. Molecular & cellular proteomics : MCP. 2009;8(6):1350–60.

62. Lehnik-Habrink M, Pförtner H, Rempeters L, Pietack N, Herzberg C, Stülke J. The RNA degradosome in Bacillus subtilis: identification of CshA as the major RNA helicase in the multiprotein complex. Mol Microbiol. 2010;77(4):958–71.

63. Liu B, Deikus G, Bree A, Durand S, Kearns DB, Bechhofer DH. Global analysis of mRNA decay intermediates in Bacillus subtilis wild-type and polynucleotide phosphorylase-deletion strains. Mol Microbiol. 2014;94(1):41–55.

64. Le Scornet A, Jousselin A, Baumas K, Kostova G, Durand S, Poljak L, et al. Critical factors for precise and efficient RNA cleavage by RNase Y in Staphylococcus aureus. PLOS Genetics. 2024;20(8):e1011349.

65. Meinken C, Blencke HM, Ludwig H, Stülke J. Expression of the glycolytic gapA operon in Bacillus subtilis: differential syntheses of proteins encoded by the operon. Microbiology (Reading, England). 2003;149(Pt 3):751–61.

66. Lee JC. Electrotransformation of Staphylococci. Methods in molecular biology (Clifton, NJ). 1995;47:209–16.

67. Schenk S, Laddaga RAJFML. Improved method for electroporation of Staphylococcus aureus. 1992;94(1-2):133–8.

68. Arnaud M, Chastanet A, Debarbouille M. New vector for efficient allelic replacement in naturally nontransformable, low-GC-content, gram-positive bacteria. Applied and environmental microbiology. 2004;70(11):6887–91.

69. Valle J, Toledo-Arana A, Berasain C, Ghigo JM, Amorena B, Penadés JR, et al. SarA and not σB is essential for biofilm development by Staphylococcus aureus. 2003;48(4):1075–87.

70. Fuchs S, Mehlan H, Bernhardt J, Hennig A, Michalik S, Surmann K, et al. AureoWiki The repository of the Staphylococcus aureus research and annotation community. Int J Med Microbiol. 2018;308(6):558–68.

71. Pfaller M, Davenport D, Bale M, Barrett M, Koontz F, Massanari RM. Development of the quantitative micro-test for slime production by coagulase-negative staphylococci. Eur J Clin Microbiol Infect Dis. 1988;7(1):30–3.

72. Gerke C, Kraft A, Süßmuth R, Schweitzer O, Götz FJJoBC. Characterization of theN-Acetylglucosaminyltransferase Activity Involved in the Biosynthesis of the Staphylococcus epidermidisPolysaccharide Intercellular Adhesin. 1998;273(29):18586–93.

73. Tormo MaAn, Martí M, Valle J, Manna AC, Cheung AL, Lasa I, et al. SarA is an essential positive regulator of Staphylococcus epidermidis biofilm development. 2005;187(7):2348–56.

74. Bronner S, Stoessel P, Gravet A, Monteil H, Prevost G. Variable expressions of Staphylococcus aureus bicomponent leucotoxins semiquantified by competitive reverse transcription-PCR. Applied and environmental microbiology. 2000;66(9):3931–8.

75. Atshan SS, Nor Shamsudin M, Sekawi Z, Lung LT, Hamat RA, Karunanidhi A, et al. Prevalence of adhesion and regulation of biofilm-related genes in different clones of Staphylococcus aureus. J Biomed Biotechnol. 2012;2012:976972.

76. Morales-Laverde L, Trobos M, Echeverz M, Solano C, Lasa I. Functional analysis of intergenic regulatory regions of genes encoding surface adhesins in Staphylococcus aureus isolates from periprosthetic joint infections. Biofilm. 2022;4:100093.

77. Ramarao N, Nielsen-Leroux C, Lereclus D. The insect Galleria mellonella as a powerful infection model to investigate bacterial pathogenesis. Journal of visualized experiments : JoVE. 2012(70):e4392.

78. Torres M, Pinzón EN, Rey FM, Martinez H, Parra Giraldo CM, Celis Ramírez AM. Galleria mellonella as a Novelty in vivo Model of Host-Pathogen Interaction for Malassezia furfur CBS 1878 and Malassezia pachydermatis CBS 1879. Frontiers in cellular and infection microbiology. 2020;10:199-.

79. Viegas SC, Mil-Homens D, Fialho AM, Arraiano CM. The virulence of Salmonella enterica Serovar Typhimurium in the insect model galleria mellonella is impaired by mutations in RNase E and RNase III. Applied and environmental microbiology. 2013;79(19):6124–33.

80. Champion OL, Titball RW, Bates S. Standardization of G. mellonella Larvae to Provide Reliable and Reproducible Results in the Study of Fungal Pathogens. J Fungi (Basel). 2018;4(3).

81. Crespi-Abril A-C, Rubilar T. Moving forward in the ethical consideration of invertebrates in experimentation: Beyond the Three R’s Principle. Revista de Biología Tropical. 2021;69(S1):S346–S57.

82. Carstens E, Moberg GP. Recognizing pain and distress in laboratory animals. ILAR J. 2000;41(2):62–71.

83. Stokes WS. Humane Endpoints for Laboratory Animals Used in Regulatory Testing. ILAR Journal. 2002;43(Suppl_1):S31–S8.

84. Bæk KT, Frees D, Renzoni A, Barras C, Rodriguez N, Manzano C, et al. Genetic Variation in the Staphylococcus aureus 8325 Strain Lineage Revealed by Whole-Genome Sequencing. PLOS ONE. 2013;8(9):e77122.

85. Monk IR, Tree JJ, Howden BP, Stinear TP, Foster TJ. Complete Bypass of Restriction Systems for Major Staphylococcus aureus Lineages. mBio. 2015;6(3):e00308–15.

86. Charpentier E, Anton AI, Barry P, Alfonso B, Fang Y, Novick RP. Novel cassette-based shuttle vector system for gram-positive bacteria. Applied and environmental microbiology. 2004;70(10):6076–85.

