## Supplementary material for "Deciphering the function of Com_YlbF domain-containing proteins in Staphylococcus aureus": Table S3_Transcriptomic analysis of the deletion of the qrp_yheA, ymcA and ylbF genes in NCTC8325_4 of Staphylococcus aureus

| test_id | gene_id | | gene | locus |
|---|---|---|---|---|
| gene1567 | gene1567 | | SAOUHSC_01687 | NC_007795.1:1597625-1597754 |
| gene2393 | gene2393 | | SAOUHSC_02533 | NC_007795.1:2335480-2335651 |
| gene2566 | gene2566 | | SAOUHSC_02721 | NC_007795.1:2502609-2502717 |
| gene1005 | gene1005 | ***ylbF*** | SAOUHSC_01070 | NC_007795.1:1036529-1036964 |
| gene1187 | gene1187 | ***ymcA*** | SAOUHSC_01270 | NC_007795.1:1225494-1225860 |
| gene1842 | gene1842 | ***yheA*** | SAOUHSC_01977 | NC_007795.1:1882163-1882508 |
| gene63 | gene63 | | SAOUHSC_00069 | NC_007795.1:73428-74979 |
| gene2287 | gene2287 | | SAOUHSC_02416 | NC_007795.1:2244538-2244724 |
| gene86 | gene86 | | SAOUHSC_00093 | NC_007795.1:99487-100087 |
| gene2655 | gene2655 | | SAOUHSC_02820 | NC_007795.1:2598081-2598777 |
| gene2366 | gene2366 | | SAOUHSC_02504 | NC_007795.1:2313309-2313943 |
| gene2361 | gene2361 | | rpsN | NC_007795.1:2311495-2311681 |
| gene2367 | gene2367 | | rplP | NC_007795.1:2313309-2313943 |
| gene2363 | gene2363 | | rplX | NC_007795.1:2312269-2312587 |
| gene2370 | gene2370 | | rpsS | NC_007795.1:2315004-2315283 |
| gene2364 | gene2364 | | rplN | NC_007795.1:2312622-2312991 |
| gene2358 | gene2358 | | rplR | NC_007795.1:2310114-2310474 |
| gene2368 | gene2368 | | rpsC | NC_007795.1:2313945-2314599 |
| gene2844 | gene2844 | | SAOUHSC_03024 | NC_007795.1:2797472-2798429 |
| gene2362 | gene2362 | | rplE | NC_007795.1:2311703-2312243 |
| gene2347 | gene2347 | | rplQ | NC_007795.1:2304223-2304592 |
| gene2426 | gene2426 | | SAOUHSC_02571 | NC_007795.1:2361997-2362801 |
| gene2365 | gene2365 | | rpsQ | NC_007795.1:2313022-2313286 |
| gene2369 | gene2369 | | rplV | NC_007795.1:2314622-2314976 |
| gene2348 | gene2348 | | SAOUHSC_02485 | NC_007795.1:2304608-2305553 |
| gene2360 | gene2360 | | rpsH | NC_007795.1:2311065-2311464 |
| gene470 | gene470 | | SAOUHSC_00499 | NC_007795.1:500137-501025 |
| gene2349 | gene2349 | | SAOUHSC_02486 | NC_007795.1:2305627-2306017 |
| gene2359 | gene2359 | | rplF | NC_007795.1:2310504-2311041 |
| gene2350 | gene2350 | | rpsM | NC_007795.1:2306040-2306406 |
| gene2656 | gene2656 | | SAOUHSC_02821 | NC_007795.1:2598778-2599549 |
| gene2357 | gene2357 | | rpsE | NC_007795.1:2309593-2310094 |
| gene2374 | gene2374 | | rplC | NC_007795.1:2317140-2317803 |
| gene2716 | gene2716 | | SAOUHSC_02887 | NC_007795.1:2660683-2661385 |
| gene64 | gene64 | | SAOUHSC_00070 | NC_007795.1:75399-76152 |
| gene2372 | gene2372 | | rplW | NC_007795.1:2316215-2317114 |
| gene2353 | gene2353 | | adk | NC_007795.1:2306984-2307632 |
| gene471 | gene471 | | SAOUHSC_00500 | NC_007795.1:501028-501589 |
| gene2665 | gene2665 | | SAOUHSC_02830 | NC_007795.1:2608102-2609050 |
| gene228 | gene228 | | SAOUHSC_00246 | NC_007795.1:264824-266201 |
| gene2352 | gene2352 | | infA | NC_007795.1:2306573-2306792 |
| gene2355 | gene2355 | | rplO | NC_007795.1:2307648-2309381 |
| gene2356 | gene2356 | | rpmD | NC_007795.1:2309397-2309577 |
| gene2354 | gene2354 | | secY | NC_007795.1:2307648-2309381 |
| gene1131 | gene1131 | | rpsP | NC_007795.1:1159734-1160010 |
| gene2375 | gene2375 | | SAOUHSC_02512a | NC_007795.1:2317830-2318139 |
| gene2371 | gene2371 | | rplB | NC_007795.1:2315349-2316183 |
| gene2373 | gene2373 | | rplD | NC_007795.1:2316215-2317114 |
| gene2434 | gene2434 | | SAOUHSC_02580 | NC_007795.1:2370059-2370698 |

| | | | |
|---|---|---|---|
| gene490 | gene490 | rplJ | NC_007795.1:520250-520751 |
| gene322 | gene322 | rpsF | NC_007795.1:360629-360926 |
| gene6 | gene6 | SAOUHSC_00007 | NC_007795.1:9754-10456 |
| gene2351 | gene2351 | rpmJ | NC_007795.1:2306428-2306542 |
| gene323 | gene323 | SAOUHSC_00349 | NC_007795.1:360946-361450 |
| gene1753 | gene1753 | SAOUHSC_01887 | NC_007795.1:1799087-1800269 |
| gene1752 | gene1752 | ribH | NC_007795.1:1798610-1799075 |
| gene1754 | gene1754 | SAOUHSC_01888 | NC_007795.1:1800279-1800912 |
| gene1396 | gene1396 | SAOUHSC_01504 | NC_007795.1:1457754-1457979 |
| gene491 | gene491 | rplL | NC_007795.1:520793-521162 |
| gene324 | gene324 | rpsR | NC_007795.1:361501-361744 |
| gene234 | gene234 | SAOUHSC_00253 | NC_007795.1:271578-273102 |
| gene1658 | gene1658 | rpmI | NC_007795.1:1682851-1683052 |
| gene1558 | gene1558 | rpsU | NC_007795.1:1587819-1587996 |
| gene936 | gene936 | SAOUHSC_00995 | NC_007795.1:968090-968525 |
| gene2570 | gene2570 | SAOUHSC_02725 | NC_007795.1:2505547-2506759 |
| gene2684 | gene2684 | SAOUHSC_02851 | NC_007795.1:2625066-2626144 |
| gene2682 | gene2682 | SAOUHSC_02849 | NC_007795.1:2623280-2625020 |
| gene2650 | gene2650 | SAOUHSC_02815 | NC_007795.1:2593501-2594779 |
| gene1659 | gene1659 | infC | NC_007795.1:1683080-1683608 |
| gene994 | gene994 | SAOUHSC_01058 | NC_007795.1:1023661-1025509 |
| gene969 | gene969 | SAOUHSC_01031 | NC_007795.1:1000776-1003154 |
| gene545 | gene545 | SAOUHSC_00581 | NC_007795.1:583051-584777 |
| gene1657 | gene1657 | rplT | NC_007795.1:1682383-1682805 |
| gene2612 | gene2612 | SAOUHSC_02773 | NC_007795.1:2550087-2551626 |
| gene2341 | gene2341 | rpsI | NC_007795.1:2299326-2299725 |
| gene2229 | gene2229 | rpmE2 | NC_007795.1:2182601-2182856 |
| gene1745 | gene1745 | SAOUHSC_01877 | NC_007795.1:1792813-1794327 |
| gene2683 | gene2683 | SAOUHSC_02850 | NC_007795.1:2625066-2626144 |
| gene546 | gene546 | SAOUHSC_00582 | NC_007795.1:583051-584777 |
| gene1755 | gene1755 | SAOUHSC_01889 | NC_007795.1:1800918-1801923 |
| gene861 | gene861 | SAOUHSC_00920 | NC_007795.1:891144-892086 |
| gene822 | gene822 | SAOUHSC_00880 | NC_007795.1:845024-846341 |
| gene494 | gene494 | SAOUHSC_00525 | NC_007795.1:525874-529471 |
| gene2698 | gene2698 | SAOUHSC_02866 | NC_007795.1:2638639-2641108 |
| gene2750 | gene2750 | SAOUHSC_02923 | NC_007795.1:2687991-2689440 |
| gene236 | gene236 | SAOUHSC_00255 | NC_007795.1:274059-274647 |
| gene2342 | gene2342 | rplM | NC_007795.1:2299738-2300176 |
| gene403 | gene403 | gltD | NC_007795.1:437848-439312 |
| gene402 | gene402 | SAOUHSC_00435 | NC_007795.1:433331-437831 |
| gene2860 | gene2860 | SAOUHSC_03042 | NC_007795.1:2812083-2812434 |
| gene1858 | gene1858 | SAOUHSC_01993 | NC_007795.1:1897328-1898648 |
| gene2256 | gene2256 | SAOUHSC_02388 | NC_007795.1:2209006-2209327 |
| gene1348 | gene1348 | SAOUHSC_01451 | NC_007795.1:1407996-1409037 |
| gene1133 | gene1133 | trmD | NC_007795.1:1160197-1161438 |
| gene2242 | gene2242 | SAOUHSC_02374 | NC_007795.1:2195933-2198311 |
| gene1758 | gene1758 | SAOUHSC_01893 | NC_007795.1:1804467-1806074 |
| gene2658 | gene2658 | SAOUHSC_02823 | NC_007795.1:2602403-2603309 |
| gene2412 | gene2412 | SAOUHSC_02554 | NC_007795.1:2348592-2349501 |
| gene2764 | gene2764 | SAOUHSC_02937 | NC_007795.1:2703338-2704961 |
| gene2786 | gene2786 | SAOUHSC_02963 | NC_007795.1:2722967-2725601 |

| | | | |
|---|---|---|---|
| gene347 | gene347 | SAOUHSC_00373 | NC_007795.1:378328-380175 |
| gene1349 | gene1349 | SAOUHSC_01452 | NC_007795.1:1409131-1410250 |
| gene1696 | gene1696 | SAOUHSC_01825 | NC_007795.1:1729303-1731675 |
| gene1300 | gene1300 | SAOUHSC_01392 | NC_007795.1:1334366-1335968 |
| gene1132 | gene1132 | rimM | NC_007795.1:1160197-1161438 |
| gene2237 | gene2237 | SAOUHSC_02369 | NC_007795.1:2192004-2192535 |
| gene1294 | gene1294 | SAOUHSC_01385 | NC_007795.1:1328428-1329280 |
| gene2823 | gene2823 | SAOUHSC_03002 | NC_007795.1:2775173-2778589 |
| gene346 | gene346 | SAOUHSC_00372 | NC_007795.1:378328-380175 |
| gene2274 | gene2274 | SAOUHSC_02410 | NC_007795.1:2237283-2238603 |
| gene2257 | gene2257 | SAOUHSC_02389 | NC_007795.1:2209328-2210309 |
| gene395 | gene395 | SAOUHSC_00427 | NC_007795.1:427608-428613 |
| gene2480 | gene2480 | SAOUHSC_02630 | NC_007795.1:2418794-2419442 |
| gene1746 | gene1746 | SAOUHSC_01878 | NC_007795.1:1792813-1794327 |
| gene497 | gene497 | SAOUHSC_00528 | NC_007795.1:530438-530909 |
| gene997 | gene997 | SAOUHSC_01062 | NC_007795.1:1026440-1026716 |
| gene1856 | gene1856 | SAOUHSC_01991 | NC_007795.1:1893344-1895517 |
| gene446 | gene446 | SAOUHSC_00483 | NC_007795.1:480714-481116 |
| gene489 | gene489 | rplA | NC_007795.1:519286-519979 |
| gene1243 | gene1243 | SAOUHSC_01330 | NC_007795.1:1272662-1273741 |
| gene2435 | gene2435 | SAOUHSC_A02450 | NC_007795.1:2370857-2371091 |
| gene1773 | gene1773 | SAOUHSC_01909 | NC_007795.1:1817569-1818763 |
| gene544 | gene544 | SAOUHSC_00580 | NC_007795.1:582658-583000 |
| gene2241 | gene2241 | SAOUHSC_02373 | NC_007795.1:2195933-2198311 |
| gene2590 | gene2590 | SAOUHSC_02750 | NC_007795.1:2525681-2527319 |
| gene700 | gene700 | SAOUHSC_00748 | NC_007795.1:731043-733716 |
| gene515 | gene515 | SAOUHSC_00545 | NC_007795.1:551104-555154 |
| gene970 | gene970 | SAOUHSC_01032 | NC_007795.1:1000776-1003154 |
| gene1981 | gene1981 | SAOUHSC_02093 | NC_007795.1:1970257-1970458 |
| gene2444 | gene2444 | SAOUHSC_02590 | NC_007795.1:2381767-2383147 |
| gene227 | gene227 | SAOUHSC_00244 | NC_007795.1:263759-264152 |
| gene1295 | gene1295 | SAOUHSC_01386 | NC_007795.1:1329326-1330244 |
| gene450 | gene450 | hslO | NC_007795.1:485713-486595 |
| gene1695 | gene1695 | SAOUHSC_01824 | NC_007795.1:1729303-1731675 |
| gene1342 | gene1342 | SAOUHSC_01440 | NC_007795.1:1374385-1375090 |
| gene1264 | gene1264 | SAOUHSC_01354 | NC_007795.1:1296848-1298309 |
| gene1296 | gene1296 | SAOUHSC_01387 | NC_007795.1:1330245-1331172 |
| gene2829 | gene2829 | SAOUHSC_03009 | NC_007795.1:2782170-2786388 |
| gene987 | gene987 | SAOUHSC_01049 | NC_007795.1:1016321-1018204 |
| gene38 | gene38 | SAOUHSC_00039 | NC_007795.1:42680-43613 |
| gene947 | gene947 | SAOUHSC_01008 | NC_007795.1:979384-980915 |
| gene2660 | gene2660 | SAOUHSC_02825 | NC_007795.1:2604028-2604901 |
| gene493 | gene493 | rpoB | NC_007795.1:522159-525711 |
| gene796 | gene796 | SAOUHSC_00853 | NC_007795.1:823485-823800 |
| gene2540 | gene2540 | SAOUHSC_02695 | NC_007795.1:2478588-2478939 |
| gene1757 | gene1757 | SAOUHSC_01891 | NC_007795.1:1804467-1806074 |
| gene1143 | gene1143 | SAOUHSC_01222 | NC_007795.1:1172817-1174893 |
| gene1759 | gene1759 | SAOUHSC_01894 | NC_007795.1:1806091-1806487 |
| gene362 | gene362 | SAOUHSC_00391 | NC_007795.1:394120-394816 |
| gene983 | gene983 | SAOUHSC_01045 | NC_007795.1:1013879-1014419 |
| gene2288 | gene2288 | SAOUHSC_02417 | NC_007795.1:2244812-2245879 |

| | | | |
|---|---|---|---|
| gene492 | gene492 | SAOUHSC_00523 | NC_007795.1:521336-521945 |
| gene1534 | gene1534 | rpmG | NC_007795.1:1564680-1564830 |
| gene805 | gene805 | SAOUHSC_00863 | NC_007795.1:831125-831392 |
| gene654 | gene654 | SAOUHSC_00701 | NC_007795.1:685709-686608 |
| gene852 | gene852 | SAOUHSC_00910 | NC_007795.1:879805-880072 |
| gene2830 | gene2830 | hisH | NC_007795.1:2782170-2786388 |
| gene1604 | gene1604 | SAOUHSC_01727 | NC_007795.1:1632958-1634101 |
| gene2479 | gene2479 | SAOUHSC_02629 | NC_007795.1:2416850-2418782 |
| gene525 | gene525 | SAOUHSC_00556 | NC_007795.1:564127-565528 |
| gene895 | gene895 | SAOUHSC_00954 | NC_007795.1:928482-931770 |
| gene496 | gene496 | rpsL | NC_007795.1:529959-530373 |
| gene1700 | gene1700 | rpsD | NC_007795.1:1734576-1735179 |
| gene488 | gene488 | rplK | NC_007795.1:518656-519079 |
| gene2187 | gene2187 | SAOUHSC_02316 | NC_007795.1:2148166-2149687 |
| gene2142 | gene2142 | SAOUHSC_02272 | NC_007795.1:2102462-2103449 |
| gene1293 | gene1293 | SAOUHSC_01384 | NC_007795.1:1327780-1328422 |
| gene1134 | gene1134 | rplS | NC_007795.1:1161540-1161891 |
| gene1628 | gene1628 | SAOUHSC_01756 | NC_007795.1:1658866-1659187 |
| gene1167 | gene1167 | rpsO | NC_007795.1:1201451-1201721 |
| gene1706 | gene1706 | SAOUHSC_01835 | NC_007795.1:1739622-1740809 |
| gene2582 | gene2582 | SAOUHSC_02740 | NC_007795.1:2519027-2520428 |
| gene968 | gene968 | SAOUHSC_01030 | NC_007795.1:1000345-1000579 |
| gene2664 | gene2664 | SAOUHSC_02829 | NC_007795.1:2607116-2607788 |
| gene2870 | gene2870 | rnpA | NC_007795.1:2820535-2820889 |
| gene2539 | gene2539 | SAOUHSC_02694 | NC_007795.1:2477970-2478570 |
| gene1201 | gene1201 | SAOUHSC_01285 | NC_007795.1:1242420-1242789 |
| gene2312 | gene2312 | SAOUHSC_02448 | NC_007795.1:2272019-2272904 |
| gene439 | gene439 | SAOUHSC_00474 | NC_007795.1:472096-472842 |
| gene2377 | gene2377 | SAOUHSC_02516 | NC_007795.1:2319076-2320255 |
| gene653 | gene653 | SAOUHSC_00700 | NC_007795.1:685709-686608 |
| gene2575 | gene2575 | SAOUHSC_02731 | NC_007795.1:2510826-2512905 |
| gene2006 | gene2006 | SAOUHSC_02119 | NC_007795.1:1991872-1993411 |
| gene1229 | gene1229 | SAOUHSC_01316 | NC_007795.1:1259355-1259889 |
| gene1694 | gene1694 | SAOUHSC_01823 | NC_007795.1:1728489-1729260 |
| gene701 | gene701 | SAOUHSC_00749 | NC_007795.1:733834-734863 |
| gene2715 | gene2715 | SAOUHSC_02886 | NC_007795.1:2659919-2660408 |
| gene1144 | gene1144 | gid | NC_007795.1:1175048-1176356 |
| gene2141 | gene2141 | SAOUHSC_02271 | NC_007795.1:2102178-2102403 |
| gene2462 | gene2462 | SAOUHSC_02611 | NC_007795.1:2401430-2402690 |
| gene1582 | gene1582 | SAOUHSC_01702 | NC_007795.1:1609898-1610585 |
| gene2472 | gene2472 | SAOUHSC_02622 | NC_007795.1:2410443-2411652 |
| gene317 | gene317 | SAOUHSC_00342 | NC_007795.1:356883-357729 |
| gene2659 | gene2659 | SAOUHSC_02824 | NC_007795.1:2603413-2604007 |
| gene787 | gene787 | SAOUHSC_00844 | NC_007795.1:814850-815672 |
| gene785 | gene785 | SAOUHSC_00842 | NC_007795.1:813119-814833 |
| gene823 | gene823 | SAOUHSC_00881 | NC_007795.1:846359-846734 |
| gene2871 | gene2871 | rpmH | NC_007795.1:2821009-2821147 |
| gene897 | gene897 | prfC | NC_007795.1:928482-931770 |
| gene984 | gene984 | SAOUHSC_01046 | NC_007795.1:1014431-1016316 |
| gene673 | gene673 | SAOUHSC_00721 | NC_007795.1:704265-704934 |
| gene971 | gene971 | SAOUHSC_01034 | NC_007795.1:1003286-1003949 |

| | | | |
|---|---|---|---|
| gene2504 | gene2504 | SAOUHSC_02658 | NC_007795.1:2441790-2443068 |
| gene1613 | gene1613 | hisS | NC_007795.1:1641089-1642352 |
| gene1849 | gene1849 | SAOUHSC_01984 | NC_007795.1:1889484-1889880 |
| gene2192 | gene2192 | SAOUHSC_02321 | NC_007795.1:2154695-2154905 |
| gene2627 | gene2627 | SAOUHSC_02789 | NC_007795.1:2560848-2561550 |
| gene1183 | gene1183 | SAOUHSC_01266 | NC_007795.1:1220801-1222562 |
| gene1629 | gene1629 | rplU | NC_007795.1:1659192-1659501 |
| gene2835 | gene2835 | hisZ | NC_007795.1:2788278-2789097 |
| gene959 | gene959 | SAOUHSC_01021 | NC_007795.1:990820-993020 |
| gene2390 | gene2390 | SAOUHSC_02530 | NC_007795.1:2333256-2334901 |
| gene1580 | gene1580 | SAOUHSC_01700 | NC_007795.1:1608250-1609351 |
| gene958 | gene958 | SAOUHSC_01019 | NC_007795.1:990820-993020 |
| gene2869 | gene2869 | trmE | NC_007795.1:2819017-2820397 |
| gene2832 | gene2832 | SAOUHSC_03012 | NC_007795.1:2782170-2786388 |
| gene1603 | gene1603 | SAOUHSC_01726 | NC_007795.1:1631840-1632958 |
| gene986 | gene986 | SAOUHSC_01048 | NC_007795.1:1016321-1018204 |
| gene672 | gene672 | SAOUHSC_00720 | NC_007795.1:703844-704264 |
| gene2868 | gene2868 | gidA | NC_007795.1:2816354-2818951 |
| gene2431 | gene2431 | SAOUHSC_02576 | NC_007795.1:2366717-2367218 |
| gene1182 | gene1182 | SAOUHSC_01265 | NC_007795.1:1219864-1220662 |
| gene613 | gene613 | SAOUHSC_00660 | NC_007795.1:647302-648370 |
| gene1744 | gene1744 | SAOUHSC_01876 | NC_007795.1:1791522-1792704 |
| gene720 | gene720 | SAOUHSC_00771 | NC_007795.1:753020-753896 |
| gene2123 | gene2123 | SAOUHSC_02250 | NC_007795.1:2085258-2086058 |
| gene1543 | gene1543 | SAOUHSC_01661 | NC_007795.1:1573689-1574367 |
| gene908 | gene908 | SAOUHSC_00965 | NC_007795.1:941383-941986 |
| gene26 | gene26 | SAOUHSC_00027 | NC_007795.1:33554-34170 |
| gene2212 | gene2212 | SAOUHSC_02341 | NC_007795.1:2167862-2169275 |
| gene1581 | gene1581 | SAOUHSC_01701 | NC_007795.1:1609351-1609879 |
| gene1899 | gene1899 | SAOUHSC_02005 | NC_007795.1:1914996-1916034 |
| gene2024 | gene2024 | SAOUHSC_02139 | NC_007795.1:2012963-2013524 |
| gene1594 | gene1594 | SAOUHSC_01716 | NC_007795.1:1620428-1622320 |
| gene869 | gene869 | SAOUHSC_00928 | NC_007795.1:899922-901638 |
| gene1013 | gene1013 | SAOUHSC_01078 | NC_007795.1:1041898-1042073 |
| gene671 | gene671 | SAOUHSC_00719 | NC_007795.1:703127-703841 |
| gene2666 | gene2666 | SAOUHSC_02831 | NC_007795.1:2609298-2610099 |
| gene2240 | gene2240 | SAOUHSC_02372 | NC_007795.1:2194863-2195535 |
| gene2618 | gene2618 | SAOUHSC_02779 | NC_007795.1:2555391-2555794 |
| gene230 | gene230 | SAOUHSC_00248 | NC_007795.1:267752-268703 |
| gene1595 | gene1595 | SAOUHSC_01717 | NC_007795.1:1622331-1623255 |
| gene2828 | gene2828 | SAOUHSC_03008 | NC_007795.1:2782170-2786388 |
| gene2800 | gene2800 | SAOUHSC_02978 | NC_007795.1:2741076-2744058 |
| gene1329 | gene1329 | SAOUHSC_01425 | NC_007795.1:1364791-1365301 |
| gene2867 | gene2867 | gidB | NC_007795.1:2816354-2818951 |
| gene1101 | gene1101 | gmk | NC_007795.1:1126850-1127692 |
| gene907 | gene907 | SAOUHSC_00964 | NC_007795.1:941192-941369 |
| gene2725 | gene2725 | SAOUHSC_02896 | NC_007795.1:2666559-2666859 |
| gene601 | gene601 | SAOUHSC_00646 | NC_007795.1:634529-635825 |
| gene2193 | gene2193 | SAOUHSC_02322 | NC_007795.1:2154916-2155210 |
| gene2211 | gene2211 | atpC | NC_007795.1:2167438-2167843 |
| gene519 | gene519 | SAOUHSC_00550 | NC_007795.1:559531-560197 |

| | | | |
|---|---|---|---|
| gene537 | gene537 | SAOUHSC_00572 | NC_007795.1:574839-575490 |
| gene1897 | gene1897 | SAOUHSC_02003 | NC_007795.1:1912125-1913862 |
| gene2230 | gene2230 | rho | NC_007795.1:2182973-2184290 |
| gene2143 | gene2143 | SAOUHSC_02273 | NC_007795.1:2103845-2104481 |
| gene1843 | gene1843 | SAOUHSC_01978 | NC_007795.1:1882576-1883701 |
| gene815 | gene815 | SAOUHSC_00873 | NC_007795.1:838631-838874 |
| gene1612 | gene1612 | aspS | NC_007795.1:1639307-1641074 |
| gene742 | gene742 | SAOUHSC_00794 | NC_007795.1:776118-777132 |
| gene1855 | gene1855 | SAOUHSC_01990 | NC_007795.1:1893344-1895517 |
| gene2526 | gene2526 | SAOUHSC_02681 | NC_007795.1:2463890-2467580 |
| gene2760 | gene2760 | SAOUHSC_02933 | NC_007795.1:2699894-2701385 |
| gene1536 | gene1536 | SAOUHSC_01653 | NC_007795.1:1567138-1567738 |
| gene1560 | gene1560 | SAOUHSC_01680 | NC_007795.1:1589641-1590394 |
| gene985 | gene985 | SAOUHSC_01047 | NC_007795.1:1014431-1016316 |
| gene1857 | gene1857 | SAOUHSC_01992 | NC_007795.1:1895774-1896836 |
| gene1593 | gene1593 | SAOUHSC_01715 | NC_007795.1:1620428-1622320 |
| gene1763 | gene1763 | SAOUHSC_01898 | NC_007795.1:1808917-1809791 |
| gene704 | gene704 | SAOUHSC_00753 | NC_007795.1:736561-737080 |
| gene786 | gene786 | SAOUHSC_00843 | NC_007795.1:813119-814833 |
| gene365 | gene365 | SAOUHSC_00394 | NC_007795.1:397353-398052 |
| gene2456 | gene2456 | SAOUHSC_02605 | NC_007795.1:2393767-2394889 |
| gene316 | gene316 | SAOUHSC_00341 | NC_007795.1:349924-356220 |
| gene348 | gene348 | SAOUHSC_00374 | NC_007795.1:380212-381679 |
| gene1678 | gene1678 | SAOUHSC_01807 | NC_007795.1:1709744-1710668 |
| gene2326 | gene2326 | SAOUHSC_02461 | NC_007795.1:2284052-2284469 |
| gene2651 | gene2651 | SAOUHSC_02816 | NC_007795.1:2595325-2595937 |
| gene1210 | gene1210 | SAOUHSC_01295 | NC_007795.1:1248419-1248671 |
| gene349 | gene349 | guaA | NC_007795.1:381703-383245 |
| gene2746 | gene2746 | panB | NC_007795.1:2682682-2684318 |
| gene2198 | gene2198 | SAOUHSC_02327 | NC_007795.1:2158347-2159220 |
| gene2332 | gene2332 | SAOUHSC_02468 | NC_007795.1:2291063-2292728 |
| gene1627 | gene1627 | rpmA | NC_007795.1:1658570-1658855 |
| gene699 | gene699 | SAOUHSC_00747 | NC_007795.1:731043-733716 |
| gene57 | gene57 | SAOUHSC_00060 | NC_007795.1:64264-65926 |
| gene520 | gene520 | SAOUHSC_00551 | NC_007795.1:560210-560573 |
| gene2437 | gene2437 | SAOUHSC_02582 | NC_007795.1:2371318-2374746 |
| gene1136 | gene1136 | rbgA | NC_007795.1:1165141-1166777 |
| gene2433 | gene2433 | SAOUHSC_02579 | NC_007795.1:2368624-2369749 |
| gene1592 | gene1592 | greA | NC_007795.1:1619924-1620401 |
| gene935 | gene935 | SAOUHSC_00994 | NC_007795.1:964092-967863 |
| gene383 | gene383 | SAOUHSC_00413 | NC_007795.1:415008-417714 |
| gene518 | gene518 | SAOUHSC_00549 | NC_007795.1:558640-559519 |
| gene1544 | gene1544 | SAOUHSC_01662 | NC_007795.1:1574497-1575604 |
| gene500 | gene500 | SAOUHSC_00531 | NC_007795.1:534795-535971 |
| gene1340 | gene1340 | SAOUHSC_01438 | NC_007795.1:1373683-1373935 |
| gene1559 | gene1559 | SAOUHSC_01679 | NC_007795.1:1588288-1589635 |
| gene1262 | gene1262 | SAOUHSC_01352 | NC_007795.1:1292205-1296599 |
| gene2217 | gene2217 | SAOUHSC_02349 | NC_007795.1:2172981-2173194 |
| gene978 | gene978 | SAOUHSC_01040 | NC_007795.1:1008406-1009519 |
| gene854 | gene854 | SAOUHSC_00912 | NC_007795.1:882635-885245 |
| gene1853 | gene1853 | SAOUHSC_01988 | NC_007795.1:1891591-1892062 |

| | | | |
|---|---|---|---|
| gene1907 | gene1907 | SAOUHSC_02014 | NC_007795.1:1922967-1923129 |
| gene305 | gene305 | SAOUHSC_00330 | NC_007795.1:342787-343177 |
| gene1389 | gene1389 | cmk | NC_007795.1:1450480-1451140 |
| gene9 | gene9 | SAOUHSC_00010 | NC_007795.1:14721-15743 |
| gene396 | gene396 | SAOUHSC_00428 | NC_007795.1:428797-429067 |
| gene1255 | gene1255 | SAOUHSC_1342a | NC_007795.1:1284056-1284419 |
| gene1980 | gene1980 | SAOUHSC_02092 | NC_007795.1:1968998-1970255 |
| gene1012 | gene1012 | SAOUHSC_01077 | NC_007795.1:1041261-1041819 |
| gene247 | gene247 | SAOUHSC_00267 | NC_007795.1:286254-287560 |
| gene1121 | gene1121 | SAOUHSC_01197 | NC_007795.1:1148669-1151302 |
| gene960 | gene960 | SAOUHSC_01022 | NC_007795.1:993034-993610 |
| gene1123 | gene1123 | SAOUHSC_01199 | NC_007795.1:1148669-1151302 |
| gene1854 | gene1854 | SAOUHSC_01989 | NC_007795.1:1892066-1893209 |
| gene623 | gene623 | SAOUHSC_00670 | NC_007795.1:657936-658944 |
| gene2213 | gene2213 | SAOUHSC_02343 | NC_007795.1:2169296-2170163 |
| gene1577 | gene1577 | SAOUHSC_01697 | NC_007795.1:1605990-1607134 |
| gene948 | gene948 | SAOUHSC_01009 | NC_007795.1:979384-980915 |
| gene2126 | gene2126 | groES | NC_007795.1:2088451-2088736 |
| gene615 | gene615 | SAOUHSC_00662 | NC_007795.1:649892-650321 |
| gene2535 | gene2535 | SAOUHSC_02690 | NC_007795.1:2475335-2476886 |
| gene318 | gene318 | SAOUHSC_00344 | NC_007795.1:357885-358767 |
| gene536 | gene536 | SAOUHSC_00571 | NC_007795.1:574374-574827 |
| gene299 | gene299 | SAOUHSC_00324 | NC_007795.1:337066-337612 |
| gene456 | gene456 | lysS | NC_007795.1:490079-491567 |
| gene1341 | gene1341 | SAOUHSC_01439 | NC_007795.1:1373946-1374168 |
| gene1137 | gene1137 | rnhB | NC_007795.1:1165141-1166777 |
| gene2726 | gene2726 | SAOUHSC_02897 | NC_007795.1:2666989-2668244 |
| gene1384 | gene1384 | engA | NC_007795.1:1447061-1448372 |
| gene1089 | gene1089 | SAOUHSC_01163 | NC_007795.1:1111793-1113202 |
| gene245 | gene245 | SAOUHSC_00265 | NC_007795.1:286254-287560 |
| gene2525 | gene2525 | SAOUHSC_02680 | NC_007795.1:2461103-2463787 |
| gene622 | gene622 | SAOUHSC_00669 | NC_007795.1:657303-657921 |
| gene1548 | gene1548 | SAOUHSC_01666 | NC_007795.1:1579474-1580866 |
| gene2478 | gene2478 | SAOUHSC_02628 | NC_007795.1:2415966-2416605 |
| gene1743 | gene1743 | leuS | NC_007795.1:1788817-1791235 |
| gene4 | gene4 | SAOUHSC_00005 | NC_007795.1:5033-6968 |
| gene2206 | gene2206 | SAOUHSC_02335 | NC_007795.1:2164231-2164735 |
| gene931 | gene931 | SAOUHSC_00989 | NC_007795.1:960953-962108 |
| gene1261 | gene1261 | SAOUHSC_01351 | NC_007795.1:1292205-1296599 |
| gene1564 | gene1564 | SAOUHSC_01684 | NC_007795.1:1594513-1595140 |
| gene862 | gene862 | SAOUHSC_00921 | NC_007795.1:892097-893342 |
| gene1256 | gene1256 | SAOUHSC_01346 | NC_007795.1:1284622-1286269 |
| gene1383 | gene1383 | gpsA | NC_007795.1:1446046-1447045 |
| gene2748 | gene2748 | SAOUHSC_02921 | NC_007795.1:2685363-2686068 |
| gene1650 | gene1650 | engB | NC_007795.1:1677129-1677720 |
| gene722 | gene722 | SAOUHSC_00773 | NC_007795.1:754304-755144 |
| gene685 | gene685 | SAOUHSC_00733 | NC_007795.1:718985-720044 |
| gene1184 | gene1184 | SAOUHSC_01267 | NC_007795.1:1222562-1223429 |
| gene1898 | gene1898 | SAOUHSC_02004 | NC_007795.1:1914152-1914695 |
| gene1589 | gene1589 | SAOUHSC_01710 | NC_007795.1:1617413-1617863 |
| gene1520 | gene1520 | SAOUHSC_01635 | NC_007795.1:1556066-1556727 |

| | | | |
|---|---|---|---|
| gene1024 | gene1024 | SAOUHSC_01091 | NC_007795.1:1050761-1051502 |
| gene1122 | gene1122 | SAOUHSC_01198 | NC_007795.1:1148669-1151302 |
| gene2639 | gene2639 | SAOUHSC_02802 | NC_007795.1:2577878-2580632 |
| gene2125 | gene2125 | groEL | NC_007795.1:2086759-2088376 |
| gene2531 | gene2531 | SAOUHSC_02686 | NC_007795.1:2472606-2473143 |
| gene531 | gene531 | SAOUHSC_00562 | NC_007795.1:570075-570906 |
| gene1725 | gene1725 | murC | NC_007795.1:1761821-1763135 |
| gene2227 | gene2227 | prfA | NC_007795.1:2180578-2181655 |
| gene1569 | gene1569 | rpsT | NC_007795.1:1600105-1600357 |
| gene972 | gene972 | SAOUHSC_A00992 | NC_007795.1:1003966-1004122 |
| gene874 | gene874 | SAOUHSC_00933 | NC_007795.1:905547-906537 |
| gene2381 | gene2381 | SAOUHSC_02520 | NC_007795.1:2323913-2324777 |
| gene750 | gene750 | SAOUHSC_00802 | NC_007795.1:784574-785315 |
| gene1610 | gene1610 | SAOUHSC_01735 | NC_007795.1:1638073-1638975 |
| gene979 | gene979 | SAOUHSC_01041 | NC_007795.1:1009522-1010500 |
| gene1339 | gene1339 | SAOUHSC_01437 | NC_007795.1:1372521-1373646 |
| gene2487 | gene2487 | SAOUHSC_02640 | NC_007795.1:2426130-2427851 |
| gene2759 | gene2759 | SAOUHSC_02932 | NC_007795.1:2697924-2699634 |
| gene1563 | gene1563 | dnaK | NC_007795.1:1592612-1594445 |
| gene932 | gene932 | SAOUHSC_00991 | NC_007795.1:962303-963314 |
| gene266 | gene266 | SAOUHSC_00290 | NC_007795.1:302335-303416 |
| gene1608 | gene1608 | SAOUHSC_01732 | NC_007795.1:1636130-1636554 |
| gene1158 | gene1158 | polC | NC_007795.1:1189524-1193835 |
| gene1202 | gene1202 | SAOUHSC_01287 | NC_007795.1:1242807-1244148 |
| gene409 | gene409 | SAOUHSC_00441 | NC_007795.1:444666-445191 |
| gene10 | gene10 | SAOUHSC_00012 | NC_007795.1:14721-15743 |
| gene2866 | gene2866 | SAOUHSC_03049 | NC_007795.1:2815472-2816312 |
| gene2681 | gene2681 | SAOUHSC_02848 | NC_007795.1:2620913-2622980 |
| gene2541 | gene2541 | SAOUHSC_02696 | NC_007795.1:2479205-2480456 |
| gene1772 | gene1772 | SAOUHSC_01908 | NC_007795.1:1816536-1817445 |
| gene443 | gene443 | SAOUHSC_00480 | NC_007795.1:473152-480200 |
| gene2408 | gene2408 | SAOUHSC_02550 | NC_007795.1:2344766-2345564 |
| gene2331 | gene2331 | SAOUHSC_02467 | NC_007795.1:2290322-2291027 |
| gene2762 | gene2762 | SAOUHSC_02935 | NC_007795.1:2701637-2702201 |
| gene1579 | gene1579 | SAOUHSC_01699 | NC_007795.1:1607430-1608237 |
| gene419 | gene419 | SAOUHSC_00452 | NC_007795.1:456713-457043 |
| gene2577 | gene2577 | SAOUHSC_02733 | NC_007795.1:2512983-2514874 |
| gene275 | gene275 | SAOUHSC_00299 | NC_007795.1:312594-313908 |
| gene1541 | gene1541 | SAOUHSC_01659 | NC_007795.1:1571126-1572473 |
| gene319 | gene319 | SAOUHSC_00345 | NC_007795.1:358796-359000 |
| gene1151 | gene1151 | tsf | NC_007795.1:1181769-1182651 |
| gene698 | gene698 | SAOUHSC_00746 | NC_007795.1:731043-733716 |
| gene3 | gene3 | recF | NC_007795.1:3669-5024 |
| gene1562 | gene1562 | SAOUHSC_01682 | NC_007795.1:1591337-1592477 |
| gene2745 | gene2745 | panC | NC_007795.1:2682682-2684318 |
| gene8 | gene8 | SAOUHSC_00009 | NC_007795.1:12785-14072 |
| gene363 | gene363 | SAOUHSC_00392 | NC_007795.1:395236-395932 |
| gene1328 | gene1328 | murG | NC_007795.1:1363709-1364780 |
| gene1576 | gene1576 | SAOUHSC_01696 | NC_007795.1:1605990-1607134 |
| gene1596 | gene1596 | SAOUHSC_01718 | NC_007795.1:1623257-1623896 |
| gene2236 | gene2236 | pyrG | NC_007795.1:2190058-2191669 |

| | | | |
|---|---|---|---|
| gene2216 | gene2216 | SAOUHSC_02347 | NC_007795.1:2171723-2172784 |
| gene2530 | gene2530 | SAOUHSC_02685 | NC_007795.1:2471645-2472377 |
| gene410 | gene410 | SAOUHSC_00442 | NC_007795.1:445283-446957 |
| gene1130 | gene1130 | SAOUHSC_01207 | NC_007795.1:1157932-1159300 |
| gene2833 | gene2833 | SAOUHSC_03013 | NC_007795.1:2786403-2788261 |
| gene445 | gene445 | SAOUHSC_00482 | NC_007795.1:480217-480610 |
| gene248 | gene248 | SAOUHSC_00268 | NC_007795.1:287569-289414 |
| gene714 | gene714 | SAOUHSC_00764 | NC_007795.1:746396-746780 |
| gene235 | gene235 | SAOUHSC_00254 | NC_007795.1:273372-274026 |
| gene2208 | gene2208 | SAOUHSC_02337 | NC_007795.1:2165267-2166533 |
| gene2043 | gene2043 | SAOUHSC_02158 | NC_007795.1:2027686-2028973 |
| gene655 | gene655 | SAOUHSC_00702 | NC_007795.1:686793-687216 |
| gene1245 | gene1245 | SAOUHSC_01332 | NC_007795.1:1273814-1274678 |
| gene1651 | gene1651 | clpX | NC_007795.1:1677873-1679136 |
| gene2687 | gene2687 | SAOUHSC_02855 | NC_007795.1:2628009-2628441 |
| gene803 | gene803 | SAOUHSC_00861 | NC_007795.1:829616-830534 |
| gene2692 | gene2692 | SAOUHSC_02860 | NC_007795.1:2631567-2632734 |
| gene2845 | gene2845 | SAOUHSC_03025 | NC_007795.1:2798580-2799219 |
| gene868 | gene868 | SAOUHSC_00927 | NC_007795.1:898055-899711 |
| gene356 | gene356 | SAOUHSC_00383 | NC_007795.1:388293-388974 |
| gene161 | gene161 | SAOUHSC_00174 | NC_007795.1:190529-191108 |
| gene2796 | gene2796 | SAOUHSC_02973 | NC_007795.1:2735411-2735870 |
| gene1116 | gene1116 | rpmB | NC_007795.1:1142973-1143162 |
| gene2228 | gene2228 | SAOUHSC_02360 | NC_007795.1:2181655-2182255 |
| gene2209 | gene2209 | SAOUHSC_02338 | NC_007795.1:2166643-2166877 |
| gene253 | gene253 | SAOUHSC_00274 | NC_007795.1:292050-292551 |
| gene2214 | gene2214 | SAOUHSC_02345 | NC_007795.1:2170193-2171702 |
| gene1979 | gene1979 | SAOUHSC_02091 | NC_007795.1:1968378-1968909 |
| gene1001 | gene1001 | SAOUHSC_01066 | NC_007795.1:1033752-1034664 |
| gene1277 | gene1277 | trpD | NC_007795.1:1312411-1313410 |
| gene2008 | gene2008 | SAOUHSC_02122 | NC_007795.1:1994711-1996715 |
| gene2009 | gene2009 | SAOUHSC_02123 | NC_007795.1:1996718-1999600 |
| gene1318 | gene1318 | SAOUHSC_01413 | NC_007795.1:1353323-1354115 |
| gene1200 | gene1200 | SAOUHSC_01284 | NC_007795.1:1240939-1242178 |
| gene2219 | gene2219 | SAOUHSC_02351 | NC_007795.1:2173985-2174438 |
| gene2215 | gene2215 | SAOUHSC_02346 | NC_007795.1:2171723-2172784 |
| gene2140 | gene2140 | SAOUHSC_02270 | NC_007795.1:2100720-2101971 |
| gene474 | gene474 | SAOUHSC_00503 | NC_007795.1:503726-505212 |
| gene660 | gene660 | SAOUHSC_00707 | NC_007795.1:690241-691923 |
| gene2473 | gene2473 | SAOUHSC_02623 | NC_007795.1:2411699-2412749 |
| gene495 | gene495 | SAOUHSC_00526 | NC_007795.1:529607-529862 |
| gene2529 | gene2529 | SAOUHSC_02684 | NC_007795.1:2469174-2471580 |
| gene1666 | gene1666 | SAOUHSC_01794 | NC_007795.1:1691106-1692132 |
| gene2547 | gene2547 | SAOUHSC_02702 | NC_007795.1:2484493-2484598 |
| gene2713 | gene2713 | SAOUHSC_02883 | NC_007795.1:2656468-2657236 |
| gene1168 | gene1168 | SAOUHSC_01251 | NC_007795.1:1202090-1204187 |
| gene2297 | gene2297 | SAOUHSC_02427 | NC_007795.1:2254505-2256502 |
| gene1472 | gene1472 | SAOUHSC_01585 | NC_007795.1:1511733-1514191 |
| gene1561 | gene1561 | prmA | NC_007795.1:1590395-1591334 |
| gene2841 | gene2841 | SAOUHSC_03021 | NC_007795.1:2794365-2795214 |
| gene681 | gene681 | SAOUHSC_00729 | NC_007795.1:712363-714247 |

| | | | |
|---|---|---|---|
| gene1540 | gene1540 | SAOUHSC_01658 | NC_007795.1:1570226-1571117 |
| gene2466 | gene2466 | SAOUHSC_02615 | NC_007795.1:2405553-2405880 |
| gene2488 | gene2488 | SAOUHSC_02641 | NC_007795.1:2426130-2427851 |
| gene2581 | gene2581 | SAOUHSC_02739 | NC_007795.1:2517866-2518802 |
| gene2545 | gene2545 | SAOUHSC_02700 | NC_007795.1:2482880-2484239 |
| gene1565 | gene1565 | SAOUHSC_01685 | NC_007795.1:1595171-1596149 |
| gene218 | gene218 | SAOUHSC_00235 | NC_007795.1:256219-257011 |
| gene933 | gene933 | SAOUHSC_00992 | NC_007795.1:963466-963886 |
| gene48 | gene48 | SAOUHSC_00050 | NC_007795.1:53567-54050 |
| gene980 | gene980 | SAOUHSC_01042 | NC_007795.1:1010590-1011883 |
| gene2346 | gene2346 | cbiO | NC_007795.1:2301223-2303686 |
| gene1762 | gene1762 | SAOUHSC_01897 | NC_007795.1:1808334-1808805 |
| gene261 | gene261 | SAOUHSC_00282 | NC_007795.1:297407-298685 |
| gene2157 | gene2157 | SAOUHSC_02286 | NC_007795.1:2118247-2119294 |
| gene2389 | gene2389 | SAOUHSC_02529 | NC_007795.1:2332087-2333095 |
| gene1846 | gene1846 | SAOUHSC_01981 | NC_007795.1:1885344-1886457 |
| gene2218 | gene2218 | SAOUHSC_02350 | NC_007795.1:2173236-2173965 |
| gene2859 | gene2859 | SAOUHSC_03041 | NC_007795.1:2811507-2812005 |
| gene1793 | gene1793 | SAOUHSC_01930 | NC_007795.1:1833475-1833835 |
| gene1338 | gene1338 | SAOUHSC_01436 | NC_007795.1:1372068-1372506 |
| gene682 | gene682 | SAOUHSC_00730 | NC_007795.1:714258-716040 |
| gene428 | gene428 | SAOUHSC_00463 | NC_007795.1:464660-465197 |
| gene2751 | gene2751 | SAOUHSC_02924 | NC_007795.1:2689520-2690858 |
| gene1371 | gene1371 | SAOUHSC_01479 | NC_007795.1:1435594-1436747 |
| gene2391 | gene2391 | SAOUHSC_02531 | NC_007795.1:2333256-2334901 |
| gene1742 | gene1742 | SAOUHSC_01874 | NC_007795.1:1788484-1788796 |
| gene2155 | gene2155 | SAOUHSC_02284 | NC_007795.1:2115681-2116686 |
| gene1298 | gene1298 | SAOUHSC_01389 | NC_007795.1:1331362-1332346 |
| gene1359 | gene1359 | SAOUHSC_01467 | NC_007795.1:1420877-1423735 |
| gene1617 | gene1617 | SAOUHSC_01743 | NC_007795.1:1646765-1647284 |
| gene1350 | gene1350 | SAOUHSC_01454 | NC_007795.1:1410724-1411603 |
| gene614 | gene614 | SAOUHSC_00661 | NC_007795.1:648520-649564 |
| gene1905 | gene1905 | SAOUHSC_02012 | NC_007795.1:1921171-1921981 |
| gene418 | gene418 | tmk | NC_007795.1:456068-456686 |
| gene119 | gene119 | SAOUHSC_00129 | NC_007795.1:134914-136060 |
| gene2542 | gene2542 | SAOUHSC_02697 | NC_007795.1:2480552-2482760 |
| gene12 | gene12 | SAOUHSC_00014 | NC_007795.1:17364-18303 |
| gene1363 | gene1363 | asnC | NC_007795.1:1426309-1427602 |
| gene2723 | gene2723 | SAOUHSC_02894 | NC_007795.1:2665324-2665702 |
| gene2156 | gene2156 | SAOUHSC_02285 | NC_007795.1:2116715-2118245 |
| gene1239 | gene1239 | SAOUHSC_01326 | NC_007795.1:1268367-1269822 |
| gene2727 | gene2727 | SAOUHSC_02898 | NC_007795.1:2666989-2668244 |
| gene751 | gene751 | SAOUHSC_00803 | NC_007795.1:785348-787721 |
| gene1250 | gene1250 | SAOUHSC_01338 | NC_007795.1:1278773-1279146 |
| gene981 | gene981 | SAOUHSC_01043 | NC_007795.1:1011886-1013293 |
| gene2592 | gene2592 | SAOUHSC_02752 | NC_007795.1:2529340-2530468 |
| gene1346 | gene1346 | SAOUHSC_01448 | NC_007795.1:1405095-1406487 |
| gene1302 | gene1302 | SAOUHSC_01395 | NC_007795.1:1338096-1339086 |
| gene1624 | gene1624 | ruvA | NC_007795.1:1655839-1656442 |
| gene1626 | gene1626 | obgE | NC_007795.1:1656923-1658216 |
| gene1739 | gene1739 | SAOUHSC_01871 | NC_007795.1:1777436-1779790 |

| | | | |
|---|---|---|---|
| gene1710 | gene1710 | SAOUHSC_01839 | NC_007795.1:1744666-1745929 |
| gene2670 | gene2670 | SAOUHSC_02836 | NC_007795.1:2612765-2613257 |
| gene927 | gene927 | SAOUHSC_00985 | NC_007795.1:953071-957547 |
| gene2699 | gene2699 | SAOUHSC_02867 | NC_007795.1:2641273-2641822 |
| gene1640 | gene1640 | valS | NC_007795.1:1665675-1668306 |
| gene2724 | gene2724 | SAOUHSC_02895 | NC_007795.1:2665718-2666540 |
| gene1178 | gene1178 | SAOUHSC_01261 | NC_007795.1:1214906-1216058 |
| gene1128 | gene1128 | SAOUHSC_01205 | NC_007795.1:1152771-1157907 |
| gene2804 | gene2804 | SAOUHSC_02983 | NC_007795.1:2749271-2752131 |
| gene2498 | gene2498 | SAOUHSC_02652 | NC_007795.1:2438501-2439404 |
| gene995 | gene995 | SAOUHSC_01060 | NC_007795.1:1025648-1025816 |
| gene1616 | gene1616 | SAOUHSC_01742 | NC_007795.1:1644148-1646338 |
| gene1199 | gene1199 | SAOUHSC_01283 | NC_007795.1:1239682-1240921 |
| gene246 | gene246 | SAOUHSC_00266 | NC_007795.1:286254-287560 |
| gene398 | gene398 | SAOUHSC_00430 | NC_007795.1:429215-430086 |
| gene624 | gene624 | SAOUHSC_00671 | NC_007795.1:659531-660329 |
| gene2298 | gene2298 | SAOUHSC_02428 | NC_007795.1:2254505-2256502 |
| gene2007 | gene2007 | SAOUHSC_02121 | NC_007795.1:1993499-1994699 |
| gene1011 | gene1011 | SAOUHSC_01076 | NC_007795.1:1039995-1041135 |
| gene656 | gene656 | SAOUHSC_00703 | NC_007795.1:687424-688591 |
| gene2159 | gene2159 | leuD | NC_007795.1:2120678-2121251 |
| gene2385 | gene2385 | SAOUHSC_02524 | NC_007795.1:2326645-2326963 |
| gene1764 | gene1764 | SAOUHSC_01899 | NC_007795.1:1808917-1809791 |
| gene1283 | gene1283 | SAOUHSC_01374 | NC_007795.1:1318356-1319616 |
| gene1317 | gene1317 | SAOUHSC_01412 | NC_007795.1:1351423-1353310 |
| gene1120 | gene1120 | SAOUHSC_01196 | NC_007795.1:1148092-1148665 |
| gene1084 | gene1084 | SAOUHSC_01156 | NC_007795.1:1105461-1106268 |
| gene2158 | gene2158 | SAOUHSC_02287 | NC_007795.1:2119307-2120678 |
| gene1575 | gene1575 | SAOUHSC_01695 | NC_007795.1:1605636-1605990 |
| gene1516 | gene1516 | SAOUHSC_01630 | NC_007795.1:1551234-1551621 |
| gene2527 | gene2527 | SAOUHSC_02682 | NC_007795.1:2467888-2469171 |
| gene320 | gene320 | ychF | NC_007795.1:359011-360109 |
| gene758 | gene758 | SAOUHSC_00811 | NC_007795.1:792192-792723 |
| gene988 | gene988 | SAOUHSC_01050 | NC_007795.1:1018277-1019297 |
| gene97 | gene97 | SAOUHSC_00106 | NC_007795.1:110292-111837 |
| gene1118 | gene1118 | SAOUHSC_01193 | NC_007795.1:1143993-1145640 |
| gene293 | gene293 | SAOUHSC_00317 | NC_007795.1:331345-332704 |
| gene2744 | gene2744 | SAOUHSC_02916 | NC_007795.1:2682297-2682669 |
| gene181 | gene181 | SAOUHSC_00195 | NC_007795.1:214245-215430 |
| gene1002 | gene1002 | SAOUHSC_01067 | NC_007795.1:1034688-1035150 |
| gene2543 | gene2543 | SAOUHSC_02698 | NC_007795.1:2480552-2482760 |
| gene993 | gene993 | SAOUHSC_01055a | NC_007795.1:1023368-1023560 |
| gene1327 | gene1327 | SAOUHSC_01423 | NC_007795.1:1363078-1363693 |
| gene1113 | gene1113 | SAOUHSC_01188 | NC_007795.1:1140424-1141300 |
| gene1010 | gene1010 | coaD | NC_007795.1:1039451-1039934 |
| gene454 | gene454 | SAOUHSC_00491 | NC_007795.1:487921-489541 |
| gene1760 | gene1760 | SAOUHSC_01895 | NC_007795.1:1806781-1807636 |
| gene1578 | gene1578 | SAOUHSC_01698 | NC_007795.1:1607136-1607427 |
| gene692 | gene692 | SAOUHSC_00740 | NC_007795.1:725113-725848 |
| gene2468 | gene2468 | SAOUHSC_02618 | NC_007795.1:2405928-2408222 |
| gene1397 | gene1397 | SAOUHSC_01505 | NC_007795.1:1458084-1458630 |

| | | | |
|---|---|---|---|
| gene2763 | gene2763 | SAOUHSC_02936 | NC_007795.1:2702388-2703075 |
| gene19 | gene19 | SAOUHSC_00020 | NC_007795.1:24930-25632 |
| gene677 | gene677 | SAOUHSC_00725 | NC_007795.1:707686-707893 |
| gene2189 | gene2189 | ddl | NC_007795.1:2151576-2152647 |
| gene374 | gene374 | SAOUHSC_00404 | NC_007795.1:406301-407087 |
| gene382 | gene382 | SAOUHSC_00412 | NC_007795.1:413511-414996 |
| gene2465 | gene2465 | SAOUHSC_02614 | NC_007795.1:2404504-2405524 |
| gene616 | gene616 | SAOUHSC_00663 | NC_007795.1:650529-651036 |
| gene1641 | gene1641 | SAOUHSC_01768 | NC_007795.1:1668708-1669269 |
| gene1085 | gene1085 | SAOUHSC_01158 | NC_007795.1:1106291-1106909 |
| gene2226 | gene2226 | SAOUHSC_02358 | NC_007795.1:2179755-2180508 |
| gene364 | gene364 | SAOUHSC_00393 | NC_007795.1:396275-396974 |
| gene1665 | gene1665 | nrdR | NC_007795.1:1690426-1690897 |
| gene154 | gene154 | SAOUHSC_00167 | NC_007795.1:180401-183429 |
| gene866 | gene866 | SAOUHSC_00925 | NC_007795.1:896023-898037 |
| gene2207 | gene2207 | fabZ | NC_007795.1:2164793-2165234 |
| gene2635 | SasG | SAOUHSC_02798 | NC_007795.1:2570008-2574892 |
| gene2514 | gene2514 | SAOUHSC_02668 | NC_007795.1:2453507-2453873 |
| gene2296 | gene2296 | SAOUHSC_02426 | NC_007795.1:2252899-2254093 |
| gene749 | gene749 | secG | NC_007795.1:784225-784459 |
| gene1035 | gene1035 | SAOUHSC_01103 | NC_007795.1:1064669-1065284 |
| gene254 | gene254 | SAOUHSC_00275 | NC_007795.1:292561-293062 |
| gene2743 | gene2743 | SAOUHSC_02915 | NC_007795.1:2680486-2682169 |
| gene2451 | gene2451 | SAOUHSC_02597 | NC_007795.1:2386752-2388357 |
| gene16 | gene16 | SAOUHSC_00019 | NC_007795.1:22443-23727 |
| gene966 | gene966 | SAOUHSC_01028 | NC_007795.1:998121-998388 |
| gene1639 | gene1639 | SAOUHSC_01766 | NC_007795.1:1664391-1665663 |
| gene2025 | gene2025 | SAOUHSC_02140 | NC_007795.1:2013576-2014506 |
| gene2378 | gene2378 | SAOUHSC_02517 | NC_007795.1:2320523-2322659 |
| gene818 | gene818 | SAOUHSC_00876 | NC_007795.1:840737-840974 |
| gene2740 | gene2740 | SAOUHSC_02912 | NC_007795.1:2679236-2679686 |
| gene1078 | gene1078 | SAOUHSC_01149 | NC_007795.1:1100156-1101569 |
| gene366 | gene366 | SAOUHSC_00395 | NC_007795.1:398417-399101 |
| gene690 | gene690 | SAOUHSC_00738 | NC_007795.1:722747-724253 |
| gene427 | gene427 | SAOUHSC_00462 | NC_007795.1:463720-464494 |
| gene1519 | gene1519 | gcvT | NC_007795.1:1554816-1555908 |
| gene2803 | gene2803 | SAOUHSC_02982 | NC_007795.1:2747122-2749030 |
| gene2475 | gene2475 | SAOUHSC_02625 | NC_007795.1:2413823-2414731 |
| gene2805 | gene2805 | SAOUHSC_02984 | NC_007795.1:2749271-2752131 |
| gene310 | gene310 | SAOUHSC_00335 | NC_007795.1:346193-347327 |
| gene2000 | gene2000 | SAOUHSC_02113 | NC_007795.1:1985146-1986508 |
| gene2387 | gene2387 | SAOUHSC_02527 | NC_007795.1:2330433-2331699 |
| gene867 | gene867 | SAOUHSC_00926 | NC_007795.1:896023-898037 |
| gene2739 | gene2739 | SAOUHSC_02911 | NC_007795.1:2678160-2678883 |
| gene1906 | gene1906 | SAOUHSC_02013 | NC_007795.1:1922321-1922837 |
| gene277 | gene277 | SAOUHSC_00301 | NC_007795.1:316639-317467 |
| gene1845 | gene1845 | SAOUHSC_01980 | NC_007795.1:1884699-1885323 |
| gene1301 | gene1301 | SAOUHSC_01394 | NC_007795.1:1336827-1338033 |
| gene2396 | gene2396 | SAOUHSC_02536 | NC_007795.1:2336045-2337068 |
| gene2515 | gene2515 | SAOUHSC_02669 | NC_007795.1:2454270-2454717 |
| gene436 | gene436 | glmU | NC_007795.1:469574-470927 |

| | | | | |
|---|---|---|---|---|
| gene996 | gene996 | | SAOUHSC_01061 | NC_007795.1:1025817-1026300 |
| gene1997 | gene1997 | | SAOUHSC_02110 | NC_007795.1:1982507-1983062 |
| gene1542 | gene1542 | | SAOUHSC_01660 | NC_007795.1:1572586-1573687 |
| gene619 | gene619 | | SAOUHSC_00666 | NC_007795.1:652087-653795 |
| gene475 | gene475 | | SAOUHSC_00504 | NC_007795.1:503726-505212 |
| gene373 | gene373 | | SAOUHSC_00402 | NC_007795.1:405464-406283 |
| gene967 | gene967 | | SAOUHSC_01029 | NC_007795.1:998390-1000109 |
| gene1372 | gene1372 | | SAOUHSC_01480 | NC_007795.1:1436760-1437864 |
| gene2840 | gene2840 | | SAOUHSC_03020 | NC_007795.1:2793782-2794337 |
| gene982 | gene982 | | SAOUHSC_01044 | NC_007795.1:1013460-1013736 |
| gene1894 | gene1894 | | SAOUHSC_02000 | NC_007795.1:1909258-1910548 |
| gene2313 | gene2313 | | SAOUHSC_02449 | NC_007795.1:2273154-2274567 |
| gene5 | gene5 | | SAOUHSC_00006 | NC_007795.1:7004-9668 |
| gene1260 | gene1260 | | SAOUHSC_01350 | NC_007795.1:1291391-1292000 |
| gene502 | gene502 | | SAOUHSC_00533 | NC_007795.1:537594-538473 |
| gene23 | gene23 | | SAOUHSC_00024 | NC_007795.1:29976-30777 |
| gene2152 | gene2152 | | SAOUHSC_02281 | NC_007795.1:2111805-2113494 |
| gene2001 | gene2001 | | SAOUHSC_02114 | NC_007795.1:1986588-1987536 |
| gene11 | gene11 | | SAOUHSC_00013 | NC_007795.1:16105-17074 |
| gene1186 | gene1186 | | SAOUHSC_01269 | NC_007795.1:1223949-1225494 |
| gene429 | gene429 | | ksgA | NC_007795.1:465207-466101 |
| gene1588 | gene1588 | | SAOUHSC_01709 | NC_007795.1:1615286-1617400 |
| gene661 | gene661 | | SAOUHSC_00708 | NC_007795.1:691928-693887 |
| gene309 | gene309 | | SAOUHSC_00334 | NC_007795.1:344355-345827 |
| gene949 | gene949 | | SAOUHSC_01010 | NC_007795.1:980918-981886 |
| gene706 | gene706 | | SAOUHSC_00755 | NC_007795.1:738231-738552 |
| gene384 | gene384 | | SAOUHSC_00414 | NC_007795.1:417875-418238 |
| gene1623 | gene1623 | | ruvB | NC_007795.1:1654803-1655808 |
| gene212 | gene212 | | SAOUHSC_00229 | NC_007795.1:250590-251265 |
| gene752 | gene752 | | smpB | NC_007795.1:787742-788207 |
| gene1079 | gene1079 | | SAOUHSC_01150 | NC_007795.1:1101601-1102774 |
| gene1165 | gene1165 | | SAOUHSC_01248 | NC_007795.1:1199433-1200351 |
| gene1272 | gene1272 | | SAOUHSC_01363 | NC_007795.1:1306258-1307521 |
| gene1242 | gene1242 | | rpsN | NC_007795.1:1272236-1272506 |
| gene423 | gene423 | | SAOUHSC_00457 | NC_007795.1:459624-460591 |
| gene1620 | gene1620 | *yajC preprote* | SAOUHSC_01747 | NC_007795.1:1652335-1652596 |
| gene2386 | gene2386 | | SAOUHSC_02525 | NC_007795.1:2327149-2330317 |
| gene827 | gene827 | | SAOUHSC_00885 | NC_007795.1:848192-849299 |
| gene2580 | gene2580 | | SAOUHSC_02737 | NC_007795.1:2516693-2517545 |
| gene1007 | gene1007 | | SAOUHSC_01072 | NC_007795.1:1038191-1038446 |
| gene2693 | gene2693 | | SAOUHSC_02861 | NC_007795.1:2632903-2633425 |
| gene1042 | gene1042 | | SAOUHSC_01110 | NC_007795.1:1070536-1070866 |
| gene1125 | gene1125 | | acpP | NC_007795.1:1151544-1151778 |
| gene2506 | gene2506 | | SAOUHSC_02660 | NC_007795.1:2443897-2444842 |
| gene2822 | *icaR* | | SAOUHSC_03001 | NC_007795.1:2774449-2775010 |
| gene691 | gene691 | | SAOUHSC_00739 | NC_007795.1:724593-725094 |
| gene1117 | gene1117 | | SAOUHSC_01192 | NC_007795.1:1143604-1143979 |
| gene1390 | gene1390 | | SAOUHSC_01497 | NC_007795.1:1451216-1452185 |
| gene845 | gene845 | | SAOUHSC_00903 | NC_007795.1:867743-868211 |
| gene1270 | gene1270 | | SAOUHSC_01361 | NC_007795.1:1304854-1305838 |
| gene505 | gene505 | | SAOUHSC_00536 | NC_007795.1:541782-543005 |

| | | | |
|---|---|---|---|
| gene973 | gene973 | SAOUHSC_01035 | NC_007795.1:1004297-1006213 |
| gene2250 | gene2250 | SAOUHSC_02382 | NC_007795.1:2203638-2204052 |
| gene2754 | gene2754 | SAOUHSC_02927 | NC_007795.1:2692737-2694234 |
| gene976 | gene976 | def | NC_007795.1:1006693-1007245 |
| gene1282 | gene1282 | SAOUHSC_01373 | NC_007795.1:1317075-1318338 |
| gene2325 | gene2325 | SAOUHSC_02460 | NC_007795.1:2283134-2283983 |
| gene255 | gene255 | SAOUHSC_00276 | NC_007795.1:293072-293573 |
| gene820 | gene820 | SAOUHSC_00878 | NC_007795.1:841799-843008 |
| gene1622 | gene1622 | queA | NC_007795.1:1653776-1654802 |
| gene1166 | gene1166 | SAOUHSC_01249 | NC_007795.1:1200365-1201337 |
| gene1347 | gene1347 | SAOUHSC_01450 | NC_007795.1:1406643-1407966 |
| gene2730 | gene2730 | SAOUHSC_02901 | NC_007795.1:2670348-2671239 |
| gene1233 | gene1233 | SAOUHSC_01320 | NC_007795.1:1263418-1264699 |
| gene636 | gene636 | SAOUHSC_00683 | NC_007795.1:670048-670738 |
| gene2474 | gene2474 | SAOUHSC_02624 | NC_007795.1:2412779-2413727 |
| gene139 | gene139 | SAOUHSC_00150 | NC_007795.1:161056-162241 |
| gene797 | gene797 | SAOUHSC_00854 | NC_007795.1:824164-825205 |
| gene2747 | gene2747 | SAOUHSC_02920 | NC_007795.1:2684417-2685278 |
| gene1677 | gene1677 | SAOUHSC_01806 | NC_007795.1:1707965-1709723 |
| gene473 | gene473 | SAOUHSC_00502 | NC_007795.1:503168-503630 |
| gene20 | gene20 | SAOUHSC_00021 | NC_007795.1:25644-27471 |
| gene1661 | gene1661 | thrS | NC_007795.1:1685755-1687693 |
| gene833 | gene833 | SAOUHSC_00891 | NC_007795.1:854534-855128 |
| gene1625 | gene1625 | SAOUHSC_01752 | NC_007795.1:1656455-1656911 |
| gene816 | gene816 | SAOUHSC_00874 | NC_007795.1:838974-839298 |
| gene772 | gene772 | SAOUHSC_00828 | NC_007795.1:805314-805932 |
| gene2144 | gene2144 | SAOUHSC_02274 | NC_007795.1:2104733-2106662 |
| gene412 | gene412 | recR | NC_007795.1:447370-447967 |
| gene1036 | gene1036 | sdhA | NC_007795.1:1065335-1067917 |
| gene2705 | gene2705 | SAOUHSC_02874 | NC_007795.1:2647868-2648075 |
| gene1726 | gene1726 | SAOUHSC_01857 | NC_007795.1:1763158-1766983 |
| gene144 | gene144 | SAOUHSC_00155 | NC_007795.1:166582-168660 |
| gene922 | gene922 | SAOUHSC_00980 | NC_007795.1:951801-952740 |
| gene21 | gene21 | SAOUHSC_00022 | NC_007795.1:27514-28798 |
| gene278 | gene278 | SAOUHSC_00302 | NC_007795.1:317539-318739 |
| gene1652 | gene1652 | tig | NC_007795.1:1679286-1680588 |
| gene1684 | gene1684 | SAOUHSC_01813 | NC_007795.1:1719076-1720375 |
| gene1635 | gene1635 | SAOUHSC_01762 | NC_007795.1:1662379-1662664 |
| gene905 | gene905 | SAOUHSC_00962 | NC_007795.1:939708-939927 |
| gene392 | gene392 | SAOUHSC_00423 | NC_007795.1:424709-425735 |
| gene2343 | gene2343 | truA | NC_007795.1:2300415-2301219 |
| gene1831 | gene1831 | SAOUHSC_01964 | NC_007795.1:1870107-1870611 |
| gene1198 | gene1198 | SAOUHSC_01282 | NC_007795.1:1239094-1239571 |
| gene2852 | gene2852 | SAOUHSC_03033 | NC_007795.1:2805005-2806022 |
| gene244 | gene244 | SAOUHSC_00264 | NC_007795.1:285846-286239 |
| gene644 | gene644 | SAOUHSC_00691 | NC_007795.1:674808-675684 |
| gene2016 | gene2016 | SAOUHSC_02131 | NC_007795.1:2004153-2004910 |
| gene828 | gene828 | SAOUHSC_00886 | NC_007795.1:849300-851119 |
| gene1683 | gene1683 | SAOUHSC_01812 | NC_007795.1:1717874-1718816 |
| gene1713 | gene1713 | SAOUHSC_01843 | NC_007795.1:1747418-1750194 |
| gene2238 | gene2238 | SAOUHSC_02370 | NC_007795.1:2192646-2193507 |

| | | | |
|---|---|---|---|
| gene890 | gene890 | SAOUHSC_00949 | NC_007795.1:922422-923991 |
| gene1375 | gene1375 | SAOUHSC_01483 | NC_007795.1:1440409-1441576 |
| gene2836 | gene2836 | SAOUHSC_03016 | NC_007795.1:2789401-2790526 |
| gene15 | gene15 | SAOUHSC_00018 | NC_007795.1:20765-22166 |
| gene76 | gene76 | SAOUHSC_00083 | NC_007795.1:90202-90967 |
| gene2445 | gene2445 | SAOUHSC_02591 | NC_007795.1:2383426-2383965 |
| gene1535 | gene1535 | SAOUHSC_01652 | NC_007795.1:1564942-1567018 |
| gene957 | gene957 | SAOUHSC_01018 | NC_007795.1:989308-990556 |
| gene686 | gene686 | SAOUHSC_00734 | NC_007795.1:720394-720937 |
| gene2251 | gene2251 | SAOUHSC_02383 | NC_007795.1:2204509-2205883 |
| gene2398 | gene2398 | SAOUHSC_02538 | NC_007795.1:2337693-2337927 |
| gene1305 | gene1305 | SAOUHSC_01398 | NC_007795.1:1340720-1341440 |
| gene1030 | gene1030 | SAOUHSC_01098 | NC_007795.1:1057683-1059396 |
| gene2477 | gene2477 | SAOUHSC_02627 | NC_007795.1:2414838-2415741 |
| gene1621 | gene1621 | tgt | NC_007795.1:1652614-1653754 |
| gene449 | gene449 | SAOUHSC_00486 | NC_007795.1:483391-485485 |
| gene579 | gene579 | SAOUHSC_00620 | NC_007795.1:611300-611675 |
| gene875 | gene875 | spxA | NC_007795.1:906831-907227 |
| gene814 | gene814 | SAOUHSC_00872 | NC_007795.1:836960-838369 |
| gene2464 | gene2464 | SAOUHSC_02613 | NC_007795.1:2403615-2404269 |
| gene670 | gene670 | SAOUHSC_00718 | NC_007795.1:702444-703029 |
| gene260 | gene260 | SAOUHSC_00281 | NC_007795.1:296346-297171 |
| gene1539 | gene1539 | SAOUHSC_01657 | NC_007795.1:1569315-1570101 |
| gene1303 | gene1303 | SAOUHSC_01396 | NC_007795.1:1339087-1339975 |
| gene2661 | gene2661 | SAOUHSC_02826 | NC_007795.1:2605074-2605509 |
| gene1026 | gene1026 | pheT | NC_007795.1:1051882-1055343 |
| gene889 | gene889 | SAOUHSC_00948 | NC_007795.1:920995-922081 |
| gene1761 | gene1761 | SAOUHSC_01896 | NC_007795.1:1807911-1808136 |
| gene1038 | gene1038 | SAOUHSC_01106 | NC_007795.1:1068152-1068953 |
| gene1110 | gene1110 | SAOUHSC_01185 | NC_007795.1:1136361-1137456 |
| gene2397 | gene2397 | mobA | NC_007795.1:2337087-2337687 |
| gene1008 | gene1008 | SAOUHSC_01073 | NC_007795.1:1038448-1038838 |
| gene851 | gene851 | SAOUHSC_00909 | NC_007795.1:878825-879650 |
| gene940 | gene940 | SAOUHSC_00999 | NC_007795.1:972595-972874 |
| gene426 | gene426 | SAOUHSC_00461 | NC_007795.1:461716-463690 |
| gene900 | gene900 | SAOUHSC_00959 | NC_007795.1:935434-936793 |
| gene47 | gene47 | SAOUHSC_00049 | NC_007795.1:50358-53511 |
| gene535 | gene535 | SAOUHSC_00569 | NC_007795.1:572748-574233 |
| gene2428 | gene2428 | SAOUHSC_02573 | NC_007795.1:2363233-2364634 |
| gene790 | gene790 | SAOUHSC_00847 | NC_007795.1:817631-818393 |
| gene1188 | gene1188 | SAOUHSC_01271 | NC_007795.1:1225886-1226378 |
| gene2756 | gene2756 | SAOUHSC_02929 | NC_007795.1:2694918-2696517 |
| gene2788 | gene2788 | SAOUHSC_02965 | NC_007795.1:2726753-2727695 |
| gene1171 | gene1171 | SAOUHSC_01254 | NC_007795.1:1208725-1209439 |
| gene151 | gene151 | SAOUHSC_00163 | NC_007795.1:177270-178113 |
| gene285 | gene285 | SAOUHSC_00309 | NC_007795.1:323457-324483 |
| gene577 | gene577 | SAOUHSC_00618 | NC_007795.1:609376-610093 |
| gene53 | gene53 | SAOUHSC_00055 | NC_007795.1:57954-58722 |
| gene1719 | gene1719 | SAOUHSC_01850 | NC_007795.1:1756345-1757335 |
| gene2294 | gene2294 | SAOUHSC_02424 | NC_007795.1:2251815-2252331 |
| gene54 | gene54 | SAOUHSC_00056 | NC_007795.1:58854-61053 |

| | | | | |
|---|---|---|---|---|
| gene990 | gene990 | | SAOUHSC_01053 | NC_007795.1:1020080-1021433 |
| gene937 | gene937 | | SAOUHSC_00996 | NC_007795.1:968679-969150 |
| gene192 | gene192 | | SAOUHSC_00206 | NC_007795.1:228481-229291 |
| gene1892 | gene1892 | | SAOUHSC_01998 | NC_007795.1:1907766-1908717 |
| gene899 | gene899 | | SAOUHSC_00958 | NC_007795.1:933093-935418 |
| gene1193 | gene1193 | | glpK | NC_007795.1:1233280-1234777 |
| gene588 | gene588 | | SAOUHSC_00632 | NC_007795.1:620305-620764 |
| gene2701 | gene2701 | | SAOUHSC_02869 | NC_007795.1:2642177-2643722 |
| gene2706 | gene2706 | | SAOUHSC_02875 | NC_007795.1:2648164-2649163 |
| gene627 | gene627 | | SAOUHSC_00674 | NC_007795.1:664260-664686 |
| gene695 | gene695 | | nrdF | NC_007795.1:729364-730336 |
| gene1126 | gene1126 | | rnc | NC_007795.1:1151893-1152625 |
| gene688 | gene688 | | SAOUHSC_00736 | NC_007795.1:721560-722478 |
| gene2406 | gene2406 | | SAOUHSC_02547 | NC_007795.1:2343009-2343681 |
| gene1332 | gene1332 | | SAOUHSC_01430 | NC_007795.1:1367382-1368104 |
| gene2035 | gene2035 | | SAOUHSC_02150 | NC_007795.1:2022518-2023082 |
| gene2484 | gene2484 | | SAOUHSC_02635 | NC_007795.1:2421819-2423202 |
| gene747 | gene747 | | eno | NC_007795.1:782056-783361 |
| gene612 | gene612 | | SAOUHSC_00659 | NC_007795.1:646349-646847 |
| gene1669 | gene1669 | | SAOUHSC_01797 | NC_007795.1:1693828-1696459 |
| gene1402 | gene1402 | | SAOUHSC_01512 | NC_007795.1:1461362-1462280 |
| gene1904 | gene1904 | | recX | NC_007795.1:1920092-1920911 |
| gene189 | gene189 | | SAOUHSC_00203 | NC_007795.1:226420-226594 |
| gene1314 | gene1314 | | SAOUHSC_01408 | NC_007795.1:1347707-1348844 |
| gene2266 | gene2266 | | SAOUHSC_02401 | NC_007795.1:2221119-2223252 |
| gene2779 | gene2779 | | nsaS | NC_007795.1:2717696-2718584 |
| gene504 | gene504 | | SAOUHSC_00535 | NC_007795.1:540482-541448 |
| gene2023 | gene2023 | | SAOUHSC_02138 | NC_007795.1:2011494-2012592 |
| gene884 | gene884 | | ppnK | NC_007795.1:914832-916493 |
| gene1223 | gene1223 | | SAOUHSC_01310 | NC_007795.1:1253957-1255439 |
| gene376 | gene376 | | SAOUHSC_00406 | NC_007795.1:407344-409720 |
| gene1147 | gene1147 | | hslU | NC_007795.1:1178276-1179680 |
| gene563 | gene563 | | SAOUHSC_00604 | NC_007795.1:596703-597240 |
| gene992 | gene992 | | SAOUHSC_01055 | NC_007795.1:1022387-1023215 |
| gene2027 | gene2027 | | SAOUHSC_02142 | NC_007795.1:2014925-2016305 |
| gene2425 | gene2425 | | SAOUHSC_02570 | NC_007795.1:2359432-2361388 |
| gene2302 | gene2302 | | SAOUHSC_02434 | NC_007795.1:2258965-2261903 |
| gene1380 | gene1380 | | SAOUHSC_01488 | NC_007795.1:1444600-1445414 |
| gene303 | gene303 | | SAOUHSC_00328 | NC_007795.1:341745-342402 |
| gene1230 | gene1230 | | SAOUHSC_01317 | NC_007795.1:1260031-1260886 |
| gene1648 | gene1648 | | SAOUHSC_01775 | NC_007795.1:1674729-1675545 |
| gene1715 | gene1715 | | SAOUHSC_01845 | NC_007795.1:1750439-1752107 |
| gene434 | gene434 | SpoVG | SAOUHSC_00469 | NC_007795.1:468928-469231 |
| gene808 | gene808 | | SAOUHSC_00866 | NC_007795.1:832810-833695 |
| gene1890 | gene1890 | | SAOUHSC_R0003 | NC_007795.1:1904930-1906485 |
| gene2849 | gene2849 | | SAOUHSC_03030 | NC_007795.1:2801301-2802720 |
| gene708 | gene708 | | SAOUHSC_00757 | NC_007795.1:740289-741516 |
| gene1826 | gene1826 | | SAOUHSC_01958 | NC_007795.1:1865525-1866080 |
| gene725 | gene725 | | SAOUHSC_00776 | NC_007795.1:756469-757501 |
| gene2235 | gene2235 | | SAOUHSC_02367 | NC_007795.1:2189428-2189950 |
| gene794 | gene794 | | SAOUHSC_00851 | NC_007795.1:821758-823156 |

| | | | |
|---|---|---|---|
| gene1602 | gene1602 | SAOUHSC_01724 | NC_007795.1:1630710-1631379 |
| gene120 | gene120 | SAOUHSC_00130 | NC_007795.1:136124-136451 |
| gene2737 | gene2737 | SAOUHSC_02909 | NC_007795.1:2676480-2677545 |
| gene2273 | gene2273 | SAOUHSC_02409 | NC_007795.1:2236149-2237058 |
| gene728 | gene728 | uvrA | NC_007795.1:759661-762508 |
| gene186 | gene186 | SAOUHSC_00200 | NC_007795.1:222778-223921 |
| gene433 | gene433 | SAOUHSC_00468 | NC_007795.1:468476-468857 |
| gene123 | gene123 | SAOUHSC_00133 | NC_007795.1:139400-140360 |
| gene2423 | gene2423 | SAOUHSC_02568 | NC_007795.1:2358273-2358600 |
| gene2195 | gene2195 | SAOUHSC_02324 | NC_007795.1:2156887-2157535 |
| gene1668 | gene1668 | SAOUHSC_01796 | NC_007795.1:1692940-1693813 |
| gene1654 | gene1654 | SAOUHSC_01781 | NC_007795.1:1680750-1681680 |
| gene2579 | gene2579 | SAOUHSC_02736 | NC_007795.1:2515105-2516602 |
| gene2432 | gene2432 | SAOUHSC_02577 | NC_007795.1:2367579-2368533 |
| gene207 | gene207 | SAOUHSC_00223 | NC_007795.1:243804-244974 |
| gene1322 | gene1322 | sucA | NC_007795.1:1357198-1359997 |
| gene860 | gene860 | SAOUHSC_00919 | NC_007795.1:890664-890850 |
| gene1716 | gene1716 | SAOUHSC_01846 | NC_007795.1:1752546-1754253 |
| gene91 | gene91 | SAOUHSC_00100 | NC_007795.1:104507-105170 |
| gene1062 | gene1062 | SAOUHSC_01133 | NC_007795.1:1086146-1087094 |
| gene1735 | gene1735 | SAOUHSC_01867 | NC_007795.1:1773872-1774721 |
| gene1765 | gene1765 | SAOUHSC_01900 | NC_007795.1:1810086-1810722 |
| gene1690 | gene1690 | SAOUHSC_01819 | NC_007795.1:1724791-1725292 |
| gene2802 | gene2802 | SAOUHSC_02980 | NC_007795.1:2746394-2746955 |
| gene507 | gene507 | SAOUHSC_00538 | NC_007795.1:543108-543792 |
| gene571 | gene571 | SAOUHSC_00612 | NC_007795.1:604015-604651 |
| gene50 | gene50 | SAOUHSC_00052 | NC_007795.1:55461-56232 |
| gene2676 | gene2676 | SAOUHSC_02842 | NC_007795.1:2617233-2617437 |
| gene1774 | gene1774 | SAOUHSC_01910 | NC_007795.1:1819133-1820726 |
| gene2032 | gene2032 | SAOUHSC_02147 | NC_007795.1:2020417-2021739 |
| gene1730 | gene1730 | SAOUHSC_01861 | NC_007795.1:1768961-1770038 |
| gene1196 | gene1196 | miaA | NC_007795.1:1237689-1238625 |
| gene2336 | gene2336 | SAOUHSC_02472 | NC_007795.1:2295452-2296589 |
| gene2413 | gene2413 | SAOUHSC_02555 | NC_007795.1:2349714-2350880 |
| gene90 | gene90 | SAOUHSC_00099 | NC_007795.1:103074-104427 |
| gene2376 | gene2376 | SAOUHSC_02515 | NC_007795.1:2318495-2318885 |
| gene1192 | gene1192 | SAOUHSC_01275 | NC_007795.1:1232333-1233152 |
| gene239 | gene239 | SAOUHSC_00258 | NC_007795.1:276307-280009 |
| gene1232 | gene1232 | SAOUHSC_01319 | NC_007795.1:1261845-1263228 |
| gene839 | gene839 | SAOUHSC_00897 | NC_007795.1:860763-861693 |
| gene2790 | gene2790 | SAOUHSC_02968 | NC_007795.1:2729232-2730243 |
| gene372 | gene372 | SAOUHSC_00401 | NC_007795.1:404818-405127 |
| gene832 | gene832 | SAOUHSC_00890 | NC_007795.1:854087-854471 |
| gene1499 | gene1499 | SAOUHSC_01612 | NC_007795.1:1535923-1537899 |
| gene1311 | gene1311 | SAOUHSC_01405 | NC_007795.1:1346287-1346596 |
| gene1004 | gene1004 | SAOUHSC_01069 | NC_007795.1:1035361-1036514 |
| gene2850 | gene2850 | SAOUHSC_03031 | NC_007795.1:2803132-2804041 |
| gene1335 | gene1335 | SAOUHSC_01433 | NC_007795.1:1369155-1369995 |
| gene1999 | gene1999 | SAOUHSC_02112 | NC_007795.1:1984446-1984977 |
| gene87 | gene87 | SAOUHSC_00094 | NC_007795.1:100454-101054 |
| gene1498 | gene1498 | SAOUHSC_01611 | NC_007795.1:1534636-1535911 |

| | | | | |
|---|---|---|---|---|
| gene328 | gene328 | | SAOUHSC_00354 | NC_007795.1:363893-364505 |
| gene730 | gene730 | | SAOUHSC_00782 | NC_007795.1:764107-764947 |
| gene2794 | gene2794 | aur | SAOUHSC_02971 | NC_007795.1:2732673-2734170 |
| gene2200 | gene2200 | | SAOUHSC_02329 | NC_007795.1:2159948-2162236 |
| gene29 | gene29 | | SAOUHSC_00030 | NC_007795.1:34711-36457 |
| gene530 | gene530 | | SAOUHSC_00561 | NC_007795.1:569401-569569 |
| gene1148 | gene1148 | | SAOUHSC_01228 | NC_007795.1:1179704-1180478 |
| gene2686 | gene2686 | | SAOUHSC_02853 | NC_007795.1:2627401-2627620 |
| gene159 | gene159 | | SAOUHSC_00172 | NC_007795.1:188731-189505 |
| gene1321 | gene1321 | | SAOUHSC_01416 | NC_007795.1:1355916-1357185 |
| gene2606 | gene2606 | | SAOUHSC_02767 | NC_007795.1:2542856-2544455 |
| gene130 | gene130 | | SAOUHSC_00141 | NC_007795.1:145404-145764 |
| gene60 | gene60 | | SAOUHSC_00064 | NC_007795.1:69185-70583 |
| gene36 | gene36 | | SAOUHSC_00037 | NC_007795.1:40820-42014 |
| gene1900 | gene1900 | | SAOUHSC_02006 | NC_007795.1:1916185-1917163 |
| gene216 | gene216 | | SAOUHSC_00233 | NC_007795.1:254121-255259 |
| gene723 | gene723 | | SAOUHSC_00774 | NC_007795.1:755317-756201 |
| gene526 | gene526 | | SAOUHSC_00557 | NC_007795.1:566071-567448 |
| gene2139 | gene2139 | | SAOUHSC_02269 | NC_007795.1:2099587-2100538 |
| gene1319 | gene1319 | | SAOUHSC_01414 | NC_007795.1:1354295-1354499 |
| gene1633 | gene1633 | | SAOUHSC_01761 | NC_007795.1:1661432-1661906 |
| gene1601 | gene1601 | | SAOUHSC_01723 | NC_007795.1:1628231-1630709 |
| gene773 | gene773 | | SAOUHSC_00830 | NC_007795.1:806086-806581 |
| gene2791 | gene2791 | | SAOUHSC_02969 | NC_007795.1:2730275-2731511 |
| gene1095 | gene1095 | | carB | NC_007795.1:1117856-1122123 |
| gene242 | gene242 | | SAOUHSC_00261 | NC_007795.1:280021-281356 |
| gene297 | gene297 | | SAOUHSC_00322 | NC_007795.1:335652-337054 |
| gene2440 | gene2440 | | SAOUHSC_02585 | NC_007795.1:2377609-2378302 |
| gene2729 | gene2729 | | SAOUHSC_02900 | NC_007795.1:2669371-2670202 |
| gene1817 | gene1817 | | SAOUHSC_01957 | NC_007795.1:1862280-1863069 |
| gene1702 | gene1702 | | SAOUHSC_01831 | NC_007795.1:1736217-1736664 |
| gene1237 | gene1237 | | SAOUHSC_01324 | NC_007795.1:1267835-1268229 |
| gene1693 | gene1693 | | tpx | NC_007795.1:1727897-1728392 |
| gene406 | gene406 | | SAOUHSC_00438 | NC_007795.1:441634-443275 |
| gene1075 | gene1075 | | mraY | NC_007795.1:1096399-1097365 |
| gene1779 | gene1779 | | SAOUHSC_01915 | NC_007795.1:1823011-1824267 |
| gene533 | gene533 | | SAOUHSC_00565 | NC_007795.1:571746-572127 |
| gene1364 | gene1364 | | SAOUHSC_01472 | NC_007795.1:1427923-1430617 |
| gene2810 | gene2810 | | secY | NC_007795.1:2758587-2759799 |
| gene945 | gene945 | | SAOUHSC_01005 | NC_007795.1:977137-977455 |
| gene1647 | gene1647 | | hemC | NC_007795.1:1673761-1674688 |
| gene360 | gene360 | | SAOUHSC_00389 | NC_007795.1:391680-392607 |
| gene2777 | gene2777 | | SAOUHSC_02953 | NC_007795.1:2714836-2717589 |
| gene2856 | gene2856 | | SAOUHSC_03037 | NC_007795.1:2807987-2810616 |
| gene2028 | gene2028 | | SAOUHSC_02143 | NC_007795.1:2016424-2017453 |
| gene2262 | gene2262 | | SAOUHSC_02396 | NC_007795.1:2215121-2215979 |
| gene1259 | gene1259 | | SAOUHSC_01349 | NC_007795.1:1290719-1291016 |
| gene1195 | gene1195 | | SAOUHSC_01279 | NC_007795.1:1236757-1237672 |
| gene2407 | gene2407 | | SAOUHSC_02549 | NC_007795.1:2343694-2344477 |
| gene2438 | gene2438 | | SAOUHSC_02583 | NC_007795.1:2375347-2376271 |
| gene259 | gene259 | | SAOUHSC_00280 | NC_007795.1:295732-296131 |

| | | | |
|---|---|---|---|
| gene559 | gene559 | SAOUHSC_00599 | NC_007795.1:593788-594635 |
| gene2721 | gene2721 | SAOUHSC_02892 | NC_007795.1:2664224-2664725 |
| gene1998 | gene1998 | SAOUHSC_02111 | NC_007795.1:1983129-1984170 |
| gene1501 | gene1501 | SAOUHSC_01614 | NC_007795.1:1537914-1539336 |
| gene1027 | gene1027 | SAOUHSC_01095 | NC_007795.1:1055513-1056452 |
| gene2816 | gene2816 | SAOUHSC_02995 | NC_007795.1:2769561-2770173 |
| gene472 | gene472 | SAOUHSC_00501 | NC_007795.1:501796-503011 |
| gene1833 | gene1833 | SAOUHSC_01967 | NC_007795.1:1870734-1872691 |
| gene2736 | gene2736 | SAOUHSC_02908 | NC_007795.1:2675396-2676263 |
| gene1986 | gene1986 | SAOUHSC_02099 | NC_007795.1:1972961-1975322 |
| gene478 | gene478 | SAOUHSC_00508 | NC_007795.1:509573-510629 |
| gene24 | gene24 | SAOUHSC_00025 | NC_007795.1:31004-33323 |
| gene663 | gene663 | SAOUHSC_00711 | NC_007795.1:695596-696946 |
| gene77 | gene77 | SAOUHSC_00084 | NC_007795.1:91162-91570 |
| gene2572 | gene2572 | SAOUHSC_02727 | NC_007795.1:2507554-2508124 |
| gene1218 | gene1218 | SAOUHSC_01304 | NC_007795.1:1251075-1251270 |
| gene1672 | gene1672 | SAOUHSC_01800 | NC_007795.1:1698738-1701104 |
| gene341 | gene341 | SAOUHSC_00367 | NC_007795.1:373164-374553 |
| gene1728 | gene1728 | SAOUHSC_01859 | NC_007795.1:1767628-1768486 |
| gene589 | gene589 | SAOUHSC_00633 | NC_007795.1:621142-623146 |
| gene2496 | gene2496 | SAOUHSC_02650 | NC_007795.1:2436750-2437380 |
| gene96 | gene96 | SAOUHSC_00105 | NC_007795.1:109107-110064 |
| gene2789 | gene2789 | SAOUHSC_02967 | NC_007795.1:2727711-2729142 |
| gene2548 | gene2548 | gpmA | NC_007795.1:2484751-2485438 |
| gene1173 | gene1173 | SAOUHSC_01256 | NC_007795.1:1209469-1212725 |
| gene863 | gene863 | SAOUHSC_00922 | NC_007795.1:893396-893768 |
| gene2741 | gene2741 | SAOUHSC_02913 | NC_007795.1:2679766-2680430 |
| gene2679 | gene2679 | SAOUHSC_02845 | NC_007795.1:2619737-2620082 |
| gene2817 | gene2817 | SAOUHSC_02996 | NC_007795.1:2770251-2770728 |
| gene918 | gene918 | SAOUHSC_00976 | NC_007795.1:949715-950579 |
| gene407 | gene407 | SAOUHSC_00439 | NC_007795.1:443299-444028 |
| gene250 | gene250 | SAOUHSC_00270 | NC_007795.1:290123-290807 |
| gene1837 | gene1837 | SAOUHSC_01972 | NC_007795.1:1875248-1876211 |
| gene666 | gene666 | SAOUHSC_00714 | NC_007795.1:699193-701383 |
| gene115 | gene115 | SAOUHSC_00125 | NC_007795.1:130861-132067 |
| gene2247 | gene2247 | SAOUHSC_02379 | NC_007795.1:2201187-2201850 |
| gene760 | gene760 | SAOUHSC_00816 | NC_007795.1:797865-798888 |
| gene1751 | gene1751 | SAOUHSC_01884 | NC_007795.1:1797487-1798489 |
| gene1836 | gene1836 | SAOUHSC_01971 | NC_007795.1:1874486-1875044 |
| gene586 | gene586 | SAOUHSC_00628 | NC_007795.1:614928-619570 |
| gene1356 | gene1356 | SAOUHSC_01463 | NC_007795.1:1419482-1420389 |
| gene2169 | gene2169 | SAOUHSC_02298 | NC_007795.1:2131928-2133481 |
| gene2260 | gene2260 | SAOUHSC_02393 | NC_007795.1:2212899-2214036 |
| gene580 | gene580 | SAOUHSC_00621 | NC_007795.1:612622-613516 |
| gene1747 | gene1747 | SAOUHSC_01879 | NC_007795.1:1794446-1794848 |
| gene2309 | gene2309 | SAOUHSC_02444 | NC_007795.1:2267631-2269194 |
| gene872 | gene872 | SAOUHSC_00931 | NC_007795.1:902677-904613 |
| gene561 | gene561 | SAOUHSC_00602 | NC_007795.1:594658-595285 |
| gene880 | gene880 | SAOUHSC_00939 | NC_007795.1:912568-912886 |
| gene2301 | gene2301 | SAOUHSC_02433 | NC_007795.1:2257888-2258959 |
| gene2662 | gene2662 | SAOUHSC_02827 | NC_007795.1:2605735-2606020 |

| | | | |
|---|---|---|---|
| gene667 | gene667 | SAOUHSC_00715 | NC_007795.1:699193-701383 |
| gene1532 | gene1532 | SAOUHSC_01649 | NC_007795.1:1560341-1563921 |
| gene1399 | gene1399 | SAOUHSC_01508 | NC_007795.1:1459226-1460141 |
| gene1673 | gene1673 | SAOUHSC_01801 | NC_007795.1:1701493-1702762 |
| gene175 | gene175 | SAOUHSC_00188 | NC_007795.1:208179-209233 |
| gene452 | gene452 | SAOUHSC_00489 | NC_007795.1:487921-489541 |
| gene625 | gene625 | SAOUHSC_00672 | NC_007795.1:660689-661334 |
| gene1357 | gene1357 | SAOUHSC_01464 | NC_007795.1:1419482-1420389 |
| gene2245 | gene2245 | SAOUHSC_02377 | NC_007795.1:2199607-2201021 |
| gene1211 | gene1211 | SAOUHSC_01296 | NC_007795.1:1248716-1249301 |
| gene727 | gene727 | SAOUHSC_00779 | NC_007795.1:758679-759654 |
| gene206 | gene206 | SAOUHSC_00222 | NC_007795.1:241835-243377 |
| gene2392 | gene2392 | SAOUHSC_02532 | NC_007795.1:2335004-2335355 |
| gene1671 | gene1671 | SAOUHSC_01799 | NC_007795.1:1698738-1701104 |
| gene598 | gene598 | SAOUHSC_00643 | NC_007795.1:631790-633952 |
| gene558 | gene558 | SAOUHSC_00598 | NC_007795.1:593182-593752 |
| gene65 | gene65 | SAOUHSC_00071 | NC_007795.1:76520-77435 |
| gene2864 | gene2864 | SAOUHSC_03047 | NC_007795.1:2814202-2814598 |
| gene2614 | gene2614 | SAOUHSC_02775 | NC_007795.1:2552611-2553031 |
| gene2607 | gene2607 | SAOUHSC_02768 | NC_007795.1:2544597-2545812 |
| gene2634 | gene2634 | SAOUHSC_02797 | NC_007795.1:2568217-2569625 |
| gene217 | gene217 | SAOUHSC_00234 | NC_007795.1:255366-256071 |
| gene2555 | gene2555 | SAOUHSC_02711 | NC_007795.1:2492703-2493171 |
| gene2767 | gene2767 | SAOUHSC_02941 | NC_007795.1:2705479-2707863 |
| gene1063 | gene1063 | SAOUHSC_01134 | NC_007795.1:1087342-1087531 |
| gene339 | gene339 | SAOUHSC_00365 | NC_007795.1:371267-371837 |
| gene1670 | gene1670 | SAOUHSC_01798 | NC_007795.1:1696752-1698240 |
| gene129 | gene129 | SAOUHSC_00139 | NC_007795.1:144156-145188 |
| gene58 | gene58 | SAOUHSC_00061 | NC_007795.1:66251-68027 |
| gene906 | gene906 | SAOUHSC_00963 | NC_007795.1:940007-940994 |
| gene194 | gene194 | SAOUHSC_00209 | NC_007795.1:229501-231140 |
| gene1253 | gene1253 | SAOUHSC_01341 | NC_007795.1:1279830-1280952 |
| gene2224 | gene2224 | SAOUHSC_02356 | NC_007795.1:2178209-2179723 |
| gene2146 | gene2146 | SAOUHSC_02276 | NC_007795.1:2107023-2108634 |
| gene41 | gene41 | SAOUHSC_00042 | NC_007795.1:44595-46360 |
| gene2620 | gene2620 | SAOUHSC_02781 | NC_007795.1:2555932-2556124 |
| gene1365 | gene1365 | SAOUHSC_01473 | NC_007795.1:1430640-1432801 |
| gene2645 | gene2645 | SAOUHSC_02809 | NC_007795.1:2587958-2588639 |
| gene2291 | gene2291 | SAOUHSC_02420 | NC_007795.1:2248362-2249706 |
| gene1401 | gene1401 | SAOUHSC_01511 | NC_007795.1:1460155-1461245 |
| gene1503 | gene1503 | SAOUHSC_01617 | NC_007795.1:1541181-1541634 |
| gene2163 | gene2163 | SAOUHSC_R0004 | NC_007795.1:2126399-2127954 |
| gene2335 | gene2335 | SAOUHSC_02471 | NC_007795.1:2293917-2295324 |
| gene1901 | gene1901 | SAOUHSC_02008 | NC_007795.1:1917423-1918260 |
| gene79 | gene79 | SAOUHSC_00086 | NC_007795.1:92457-93234 |
| gene1729 | gene1729 | SAOUHSC_01860 | NC_007795.1:1768585-1768897 |
| gene1701 | gene1701 | SAOUHSC_01830 | NC_007795.1:1735394-1736138 |
| gene649 | gene649 | SAOUHSC_00696 | NC_007795.1:681029-681938 |
| gene595 | gene595 | SAOUHSC_00640 | NC_007795.1:628913-629678 |
| gene855 | gene855 | SAOUHSC_00913 | NC_007795.1:885303-886173 |
| gene2801 | gene2801 | SAOUHSC_02979 | NC_007795.1:2744268-2746128 |

| | | | | |
|---|---|---|---|---|
| gene2500 | gene2500 | | SAOUHSC_02654 | NC_007795.1:2439886-2440921 |
| gene1717 | gene1717 | | SAOUHSC_01847 | NC_007795.1:1754421-1755054 |
| gene1832 | gene1832 | | SAOUHSC_01966 | NC_007795.1:1870734-1872691 |
| gene757 | gene757 | | SAOUHSC_00810 | NC_007795.1:791532-792054 |
| gene759 | gene759 | | SAOUHSC_00812 | NC_007795.1:792985-795769 |
| gene1285 | gene1285 | | SAOUHSC_01376 | NC_007795.1:1321113-1321587 |
| gene1155 | gene1155 | | SAOUHSC_01238 | NC_007795.1:1185232-1186015 |
| gene157 | gene157 | | SAOUHSC_00170 | NC_007795.1:184614-186390 |
| gene128 | gene128 | | SAOUHSC_00138 | NC_007795.1:142411-144144 |
| gene1989 | gene1989 | | SAOUHSC_02102 | NC_007795.1:1975939-1976698 |
| gene1045 | gene1045 | | SAOUHSC_01113 | NC_007795.1:1072307-1072814 |
| gene1324 | gene1324 | | SAOUHSC_01420 | NC_007795.1:1360280-1362292 |
| gene650 | gene650 | | SAOUHSC_00697 | NC_007795.1:681972-682260 |
| gene1093 | gene1093 | | pyrC | NC_007795.1:1116580-1117855 |
| gene338 | gene338 | | SAOUHSC_00364 | NC_007795.1:369728-371252 |
| gene291 | gene291 | | SAOUHSC_00315 | NC_007795.1:329363-330719 |
| gene2205 | gene2205 | | SAOUHSC_02334 | NC_007795.1:2163705-2164101 |
| gene2022 | gene2022 | | SAOUHSC_02137 | NC_007795.1:2009727-2011290 |
| gene665 | gene665 | | SAOUHSC_00713 | NC_007795.1:698138-699122 |
| gene1895 | gene1895 | | SAOUHSC_02001 | NC_007795.1:1910891-1911935 |
| gene2837 | gene2837 | | SAOUHSC_03017 | NC_007795.1:2790643-2791141 |
| gene420 | gene420 | | SAOUHSC_00454 | NC_007795.1:457343-458183 |
| gene2714 | gene2714 | | SAOUHSC_02885 | NC_007795.1:2657596-2659408 |
| gene807 | gene807 | | SAOUHSC_00865 | NC_007795.1:831489-832703 |
| gene715 | gene715 | | SAOUHSC_00765 | NC_007795.1:747287-748893 |
| gene399 | gene399 | | SAOUHSC_00431 | NC_007795.1:430254-432152 |
| gene2329 | gene2329 | | SAOUHSC_02464 | NC_007795.1:2287359-2288214 |
| gene2005 | gene2005 | | gatC | NC_007795.1:1991203-1991506 |
| gene1222 | gene1222 | | SAOUHSC_1307a | NC_007795.1:1253702-1253900 |
| gene1736 | gene1736 | | SAOUHSC_01868 | NC_007795.1:1774724-1776134 |
| gene225 | gene225 | rbsR | SAOUHSC_00242 | NC_007795.1:262381-263380 |
| gene503 | gene503 | | SAOUHSC_00534 | NC_007795.1:538631-540269 |
| gene2244 | gene2244 | | SAOUHSC_02376 | NC_007795.1:2199243-2199594 |
| gene2694 | gene2694 | | SAOUHSC_02862 | NC_007795.1:2633833-2635939 |
| gene281 | gene281 | | SAOUHSC_00305 | NC_007795.1:320075-320408 |
| gene2202 | gene2202 | | SAOUHSC_02331 | NC_007795.1:2159948-2162236 |
| gene252 | gene252 | | SAOUHSC_00272 | NC_007795.1:291700-291919 |
| gene183 | gene183 | | SAOUHSC_00197 | NC_007795.1:217906-219118 |
| gene2571 | gene2571 | | SAOUHSC_02726 | NC_007795.1:2506881-2507355 |
| gene2328 | gene2328 | | SAOUHSC_02463 | NC_007795.1:2284712-2287300 |
| gene2862 | gene2862 | | SAOUHSC_03045 | NC_007795.1:2813019-2813247 |
| gene2493 | gene2493 | | SAOUHSC_02647 | NC_007795.1:2431150-2432629 |
| gene2183 | gene2183 | | SAOUHSC_02312 | NC_007795.1:2142497-2144174 |
| gene1711 | gene1711 | | SAOUHSC_01840 | NC_007795.1:1746310-1747288 |
| gene1017 | gene1017 | | SAOUHSC_01084 | NC_007795.1:1045628-1047388 |
| gene401 | gene401 | | SAOUHSC_00434 | NC_007795.1:432266-433151 |
| gene1016 | gene1016 | | SAOUHSC_01082 | NC_007795.1:1045628-1047388 |
| gene1790 | gene1790 | | SAOUHSC_01926 | NC_007795.1:1831475-1831859 |
| gene2029 | gene2029 | | SAOUHSC_02144 | NC_007795.1:2017722-2018124 |
| gene2450 | gene2450 | | SAOUHSC_02596 | NC_007795.1:2386156-2386687 |
| gene2749 | gene2749 | | SAOUHSC_02922 | NC_007795.1:2686345-2687305 |

| | | | |
|---|---|---|---|
| gene1512 | gene1512 | SAOUHSC_01626 | NC_007795.1:1548203-1549265 |
| gene378 | gene378 | SAOUHSC_00408 | NC_007795.1:409738-410053 |
| gene769 | gene769 | SAOUHSC_00825 | NC_007795.1:803999-804436 |
| gene283 | gene283 | SAOUHSC_00307 | NC_007795.1:320408-323143 |
| gene1377 | gene1377 | ndk | NC_007795.1:1442370-1442802 |
| gene92 | gene92 | SAOUHSC_00101 | NC_007795.1:105197-106376 |
| gene878 | gene878 | SAOUHSC_00937 | NC_007795.1:909471-911280 |
| gene2793 | gene2793 | SAOUHSC_02970 | NC_007795.1:2731857-2732307 |
| gene2605 | gene2605 | SAOUHSC_02766 | NC_007795.1:2539488-2542844 |
| gene1141 | gene1141 | SAOUHSC_01220 | NC_007795.1:1170354-1171599 |
| gene171 | gene171 | SAOUHSC_00184 | NC_007795.1:202047-205320 |
| gene1493 | gene1493 | SAOUHSC_01606 | NC_007795.1:1531152-1532286 |
| gene135 | gene135 | SAOUHSC_00146 | NC_007795.1:157204-157684 |
| gene2551 | Sbi | SAOUHSC_02706 | NC_007795.1:2487382-2488693 |
| gene251 | gene251 | SAOUHSC_00271 | NC_007795.1:290953-291565 |
| gene1207 | gene1207 | SAOUHSC_01292 | NC_007795.1:1247489-1247675 |
| gene1902 | gene1902 | SAOUHSC_02009 | NC_007795.1:1918268-1919786 |
| gene7 | gene7 | SAOUHSC_00008 | NC_007795.1:10892-12407 |
| gene1385 | gene1385 | rpsA | NC_007795.1:1448593-1449769 |
| gene1780 | gene1780 | SAOUHSC_01916 | NC_007795.1:1824271-1825750 |
| gene66 | gene66 | SAOUHSC_00072 | NC_007795.1:77515-78511 |
| gene385 | gene385 | SAOUHSC_00415 | NC_007795.1:418466-418883 |
| gene767 | gene767 | SAOUHSC_00823 | NC_007795.1:803181-803370 |
| gene2819 | gene2819 | SAOUHSC_02998 | NC_007795.1:2771465-2772922 |
| gene551 | gene551 | SAOUHSC_00587 | NC_007795.1:588391-588937 |
| gene1988 | gene1988 | SAOUHSC_02101 | NC_007795.1:1975336-1975723 |
| gene1768 | gene1768 | SAOUHSC_01903 | NC_007795.1:1812692-1813408 |
| gene185 | gene185 | SAOUHSC_00199 | NC_007795.1:220760-222338 |
| gene2132 | gene2132 | SAOUHSC_02261 | NC_007795.1:2093873-2094580 |
| gene2601 | gene2601 | SAOUHSC_02762 | NC_007795.1:2538283-2539477 |
| gene370 | gene370 | SAOUHSC_00399 | NC_007795.1:402507-403185 |
| gene2795 | gene2795 | SAOUHSC_02972 | NC_007795.1:2734626-2735154 |
| gene184 | gene184 | SAOUHSC_00198 | NC_007795.1:219229-220735 |
| gene2499 | gene2499 | SAOUHSC_02653 | NC_007795.1:2439449-2439785 |
| gene859 | gene859 | SAOUHSC_00917 | NC_007795.1:886282-889663 |
| gene1219 | gene1219 | SAOUHSC_01305 | NC_007795.1:1251319-1251661 |
| gene1485 | gene1485 | SAOUHSC_01598 | NC_007795.1:1522314-1523235 |
| gene2611 | gene2611 | SAOUHSC_02772 | NC_007795.1:2549036-2549756 |
| gene2034 | gene2034 | SAOUHSC_02149 | NC_007795.1:2021873-2022398 |
| gene1050 | gene1050 | SAOUHSC_A01081 | NC_007795.1:1075290-1075509 |
| gene1991 | gene1991 | SAOUHSC_02104 | NC_007795.1:1978642-1979056 |
| gene214 | gene214 | SAOUHSC_00231 | NC_007795.1:253268-254009 |
| gene709 | gene709 | SAOUHSC_00758 | NC_007795.1:741529-742024 |
| gene1891 | gene1891 | SAOUHSC_01997 | NC_007795.1:1907223-1907670 |
| gene1533 | gene1533 | SAOUHSC_01650 | NC_007795.1:1563932-1564472 |
| gene2600 | gene2600 | SAOUHSC_02761 | NC_007795.1:2537310-2538060 |
| gene294 | gene294 | SAOUHSC_00318 | NC_007795.1:333011-333938 |
| gene1055 | gene1055 | SAOUHSC_01124 | NC_007795.1:1078815-1079532 |
| gene1071 | gene1071 | SAOUHSC_01142 | NC_007795.1:1092095-1092527 |
| gene169 | gene169 | SAOUHSC_00182 | NC_007795.1:199582-200155 |
| gene1067 | gene1067 | SAOUHSC_01137 | NC_007795.1:1088891-1089587 |

| | | | |
|---|---|---|---|
| gene1676 | gene1676 | SAOUHSC_01805 | NC_007795.1:1705885-1707325 |
| gene2768 | gene2768 | SAOUHSC_02942 | NC_007795.1:2705479-2707863 |
| gene662 | gene662 | SAOUHSC_00710 | NC_007795.1:694194-695376 |
| gene626 | gene626 | SAOUHSC_00673 | NC_007795.1:662030-664181 |
| gene1018 | gene1018 | SAOUHSC_01085 | NC_007795.1:1047414-1048263 |
| gene2330 | gene2330 | SAOUHSC_02465 | NC_007795.1:2288424-2289240 |
| gene2678 | gene2678 | SAOUHSC_02844 | NC_007795.1:2618636-2619605 |
| gene1394 | gene1394 | SAOUHSC_01502 | NC_007795.1:1455300-1457623 |
| gene188 | gene188 | SAOUHSC_00202 | NC_007795.1:225907-226264 |
| gene2117 | gene2117 | SAOUHSC_02244 | NC_007795.1:2078220-2079444 |
| gene881 | gene881 | SAOUHSC_00940 | NC_007795.1:913037-913631 |
| gene379 | gene379 | SAOUHSC_00409 | NC_007795.1:410342-410603 |
| gene2842 | gene2842 | SAOUHSC_03022 | NC_007795.1:2795593-2796109 |
| gene2292 | gene2292 | SAOUHSC_02422 | NC_007795.1:2249882-2250566 |
| gene1634 | gene1634 | SAOUHSC_01761a | NC_007795.1:1662065-1662263 |
| gene2599 | gene2599 | SAOUHSC_02760 | NC_007795.1:2535538-2537116 |
| gene2290 | gene2290 | SAOUHSC_02419 | NC_007795.1:2247790-2248264 |
| gene2268 | gene2268 | SAOUHSC_02403 | NC_007795.1:2223263-2224804 |
| gene2667 | gene2667 | SAOUHSC_02832 | NC_007795.1:2610419-2611055 |
| gene1286 | gene1286 | SAOUHSC_01377 | NC_007795.1:1321710-1324974 |
| gene916 | gene916 | SAOUHSC_00973 | NC_007795.1:947204-948687 |
| gene939 | gene939 | SAOUHSC_00998 | NC_007795.1:970865-972059 |
| gene1323 | gene1323 | SAOUHSC_01419 | NC_007795.1:1360280-1362292 |
| gene2509 | gene2509 | SAOUHSC_02663 | NC_007795.1:2446554-2446938 |
| gene302 | gene302 | SAOUHSC_00327 | NC_007795.1:339970-341653 |
| gene2685 | gene2685 | SAOUHSC_02852 | NC_007795.1:2626356-2627184 |
| gene1006 | gene1006 | SAOUHSC_01071 | NC_007795.1:1037026-1037953 |
| gene554 | gene554 | SAOUHSC_00591 | NC_007795.1:589836-590430 |
| gene2563 | gene2563 | SAOUHSC_02718 | NC_007795.1:2498460-2500194 |
| gene567 | gene567 | adhA | NC_007795.1:600267-601278 |
| gene400 | gene400 | SAOUHSC_00433 | NC_007795.1:430254-432152 |
| gene2249 | gene2249 | SAOUHSC_02381 | NC_007795.1:2202995-2203439 |
| gene1019 | gene1019 | SAOUHSC_01086 | NC_007795.1:1048275-1049240 |
| gene335 | gene335 | SAOUHSC_00360 | NC_007795.1:368019-368403 |
| gene576 | gene576 | SAOUHSC_00617 | NC_007795.1:608708-609215 |
| gene1984 | gene1984 | SAOUHSC_02097 | NC_007795.1:1971628-1972846 |
| gene770 | gene770 | SAOUHSC_00826 | NC_007795.1:803999-804436 |
| gene1135 | gene1135 | SAOUHSC_01213 | NC_007795.1:1162134-1164741 |
| gene1020 | gene1020 | SAOUHSC_01087 | NC_007795.1:1048275-1049240 |
| gene2615 | gene2615 | SAOUHSC_02776 | NC_007795.1:2553048-2553468 |
| gene2190 | gene2190 | SAOUHSC_02319 | NC_007795.1:2152964-2154167 |
| gene1057 | gene1057 | SAOUHSC_01127 | NC_007795.1:1080459-1081185 |
| gene237 | gene237 | SAOUHSC_00256 | NC_007795.1:274789-275683 |
| gene282 | gene282 | SAOUHSC_00306 | NC_007795.1:320408-323143 |
| gene1788 | gene1788 | SAOUHSC_01924 | NC_007795.1:1831101-1831287 |
| gene1995 | gene1995 | SAOUHSC_02108 | NC_007795.1:1981618-1982119 |
| gene1056 | gene1056 | SAOUHSC_01125 | NC_007795.1:1079639-1080365 |
| gene917 | gene917 | SAOUHSC_00975 | NC_007795.1:948775-949132 |
| gene639 | gene639 | SAOUHSC_00686 | NC_007795.1:671907-672207 |
| gene296 | gene296 | SAOUHSC_00320 | NC_007795.1:335026-335593 |
| gene295 | gene295 | SAOUHSC_00319 | NC_007795.1:333951-335013 |

| | | | |
|---|---|---|---|
| gene774 | gene774 | SAOUHSC_00831 | NC_007795.1:806979-807402 |
| gene288 | gene288 | SAOUHSC_00312 | NC_007795.1:326222-326666 |
| gene609 | gene609 | SAOUHSC_00655 | NC_007795.1:644284-645253 |
| gene430 | gene430 | SAOUHSC_00465 | NC_007795.1:466200-466464 |
| gene664 | gene664 | SAOUHSC_00712 | NC_007795.1:697167-698007 |
| gene2815 | gene2815 | SAOUHSC_02994 | NC_007795.1:2769117-2769477 |
| gene1216 | gene1216 | SAOUHSC_01301 | NC_007795.1:1249946-1250347 |
| gene2598 | gene2598 | SAOUHSC_02759 | NC_007795.1:2534915-2535371 |
| gene271 | gene271 | SAOUHSC_00295 | NC_007795.1:308682-309564 |
| gene915 | gene915 | SAOUHSC_00972 | NC_007795.1:946911-947199 |
| gene879 | gene879 | SAOUHSC_00938 | NC_007795.1:911739-912546 |
| gene2492 | gene2492 | SAOUHSC_02646 | NC_007795.1:2430189-2431085 |
| gene2422 | gene2422 | SAOUHSC_02566 | NC_007795.1:2357498-2357846 |
| gene2452 | gene2452 | SAOUHSC_02599 | NC_007795.1:2388686-2389451 |
| gene2593 | gene2593 | SAOUHSC_02753 | NC_007795.1:2530923-2532355 |
| gene287 | gene287 | SAOUHSC_00311 | NC_007795.1:325936-326221 |
| gene1194 | gene1194 | SAOUHSC_01278 | NC_007795.1:1234934-1236608 |
| gene2460 | gene2460 | SAOUHSC_02609 | NC_007795.1:2399530-2399950 |
| gene2853 | gene2853 | SAOUHSC_03034 | NC_007795.1:2806264-2807059 |
| gene111 | gene111 | SAOUHSC_00121 | NC_007795.1:126742-127369 |
| gene1786 | gene1786 | SAOUHSC_01922 | NC_007795.1:1829577-1830129 |
| gene112 | gene112 | SAOUHSC_00122 | NC_007795.1:127373-128483 |
| gene2703 | gene2703 | SAOUHSC_02872 | NC_007795.1:2644749-2644941 |
| gene2414 | gene2414 | SAOUHSC_02557 | NC_007795.1:2351096-2351987 |
| gene753 | gene753 | SAOUHSC_00806 | NC_007795.1:789528-789852 |
| gene2647 | gene2647 | SAOUHSC_02811 | NC_007795.1:2589721-2590414 |
| gene1483 | gene1483 | SAOUHSC_01596 | NC_007795.1:1520497-1521253 |
| gene2119 | gene2119 | SAOUHSC_02246 | NC_007795.1:2080722-2081634 |
| gene556 | gene556 | SAOUHSC_00593 | NC_007795.1:591276-591905 |
| gene2134 | gene2134 | SAOUHSC_02264 | NC_007795.1:2094652-2095897 |
| gene270 | gene270 | SAOUHSC_00294 | NC_007795.1:307110-308643 |
| gene351 | gene351 | SAOUHSC_00377 | NC_007795.1:384286-384991 |
| gene2286 | gene2286 | SAOUHSC_R0005 | NC_007795.1:2242799-2244356 |
| gene2455 | gene2455 | SAOUHSC_02604 | NC_007795.1:2392775-2393657 |
| gene2118 | gene2118 | SAOUHSC_02245 | NC_007795.1:2079815-2080661 |
| gene1990 | gene1990 | SAOUHSC_02103 | NC_007795.1:1977225-1978212 |
| gene2646 | gene2646 | SAOUHSC_02810 | NC_007795.1:2588792-2589557 |
| gene114 | gene114 | SAOUHSC_00124 | NC_007795.1:128496-130861 |
| gene220 | gene220 | SAOUHSC_00237 | NC_007795.1:258895-259657 |
| gene173 | gene173 | SAOUHSC_00186 | NC_007795.1:202047-205320 |
| gene1344 | gene1344 | SAOUHSC_01443 | NC_007795.1:1375630-1376032 |
| gene806 | gene806 | SAOUHSC_00864 | NC_007795.1:831489-832703 |
| gene110 | gene110 | SAOUHSC_00120 | NC_007795.1:125615-126740 |
| gene1776 | gene1776 | SAOUHSC_01912 | NC_007795.1:1821721-1822468 |
| gene1142 | gene1142 | SAOUHSC_01221 | NC_007795.1:1171903-1172644 |
| gene1571 | gene1571 | SAOUHSC_01691 | NC_007795.1:1601432-1603580 |
| gene301 | gene301 | SAOUHSC_00326 | NC_007795.1:337879-339960 |
| gene710 | gene710 | SAOUHSC_00759 | NC_007795.1:742041-742803 |
| gene1217 | gene1217 | SAOUHSC_01303 | NC_007795.1:1250603-1251065 |
| gene800 | gene800 | SAOUHSC_00858 | NC_007795.1:826498-827347 |
| gene755 | gene755 | SAOUHSC_00808 | NC_007795.1:790205-790934 |

| | | | |
|---|---|---|---|
| gene1502 | gene1502 | SAOUHSC_01615 | NC_007795.1:1539486-1541166 |
| gene1709 | gene1709 | SAOUHSC_01838 | NC_007795.1:1743298-1744573 |
| gene754 | gene754 | SAOUHSC_00807 | NC_007795.1:789867-790014 |
| gene1756 | gene1756 | SAOUHSC_01890 | NC_007795.1:1802442-1803945 |
| gene289 | gene289 | SAOUHSC_00313 | NC_007795.1:326670-328626 |
| gene1106 | gene1106 | SAOUHSC_01181 | NC_007795.1:1133136-1133415 |
| gene156 | gene156 | SAOUHSC_00169 | NC_007795.1:183434-184598 |
| gene61 | gene61 | SAOUHSC_00065 | NC_007795.1:70772-71246 |
| gene1092 | gene1092 | pyrB | NC_007795.1:1115681-1116563 |
| gene610 | gene610 | SAOUHSC_00656 | NC_007795.1:645294-646234 |
| gene802 | gene802 | SAOUHSC_00860 | NC_007795.1:828213-829533 |
| gene555 | gene555 | SAOUHSC_00592 | NC_007795.1:590588-591230 |
| gene590 | gene590 | SAOUHSC_00634 | NC_007795.1:624571-627080 |
| gene42 | gene42 | SAOUHSC_00043 | NC_007795.1:44595-46360 |
| gene557 | gene557 | SAOUHSC_00596 | NC_007795.1:592357-592987 |
| gene669 | gene669 | SAOUHSC_00717 | NC_007795.1:701724-702165 |
| gene528 | gene528 | SAOUHSC_00559 | NC_007795.1:567449-568929 |
| gene1482 | gene1482 | SAOUHSC_01595 | NC_007795.1:1520233-1520482 |
| gene1240 | gene1240 | SAOUHSC_01327 | NC_007795.1:1270121-1271543 |
| gene1288 | gene1288 | SAOUHSC_01379 | NC_007795.1:1321710-1324974 |
| gene280 | gene280 | SAOUHSC_00304 | NC_007795.1:319040-320042 |
| gene2040 | gene2040 | SAOUHSC_02155 | NC_007795.1:2025437-2026711 |
| gene2863 | gene2863 | SAOUHSC_03046 | NC_007795.1:2813373-2813943 |
| gene1769 | gene1769 | SAOUHSC_01904 | NC_007795.1:1812692-1813408 |
| gene1526 | gene1526 | SAOUHSC_01641 | NC_007795.1:1558273-1559194 |
| gene1718 | gene1718 | SAOUHSC_01849 | NC_007795.1:1755078-1756248 |
| gene172 | gene172 | SAOUHSC_00185 | NC_007795.1:202047-205320 |
| gene821 | gene821 | SAOUHSC_00879 | NC_007795.1:843138-844614 |
| gene2630 | gene2630 | SAOUHSC_02793 | NC_007795.1:2565299-2567108 |
| gene2458 | gene2458 | SAOUHSC_02607 | NC_007795.1:2395165-2398065 |
| gene2459 | gene2459 | SAOUHSC_02608 | NC_007795.1:2398224-2399109 |
| gene2457 | gene2457 | SAOUHSC_02606 | NC_007795.1:2395165-2398065 |
| gene1090 | gene1090 | SAOUHSC_01164 | NC_007795.1:1113601-1114129 |
| gene337 | gene337 | SAOUHSC_00363 | NC_007795.1:369356-369650 |
| gene67 | gene67 | SAOUHSC_00074 | NC_007795.1:78526-79519 |
| gene2797 | gene2797 | SAOUHSC_02974 | NC_007795.1:2736109-2737984 |
| gene1287 | gene1287 | SAOUHSC_01378 | NC_007795.1:1321710-1324974 |
| gene1140 | gene1140 | SAOUHSC_01219 | NC_007795.1:1169208-1170327 |
| gene1172 | gene1172 | SAOUHSC_01255 | NC_007795.1:1209469-1212725 |
| gene583 | gene583 | SAOUHSC_00625 | NC_007795.1:614928-619570 |
| gene1488 | gene1488 | SAOUHSC_01601 | NC_007795.1:1526210-1527860 |
| gene801 | gene801 | SAOUHSC_00859 | NC_007795.1:827359-828187 |
| gene224 | gene224 | SAOUHSC_00241 | NC_007795.1:261268-262150 |
| gene2491 | gene2491 | SAOUHSC_02645 | NC_007795.1:2430189-2431085 |
| gene345 | gene345 | SAOUHSC_00371 | NC_007795.1:377408-377816 |
| gene2127 | gene2127 | SAOUHSC_02256 | NC_007795.1:2088910-2089654 |
| gene113 | gene113 | SAOUHSC_00123 | NC_007795.1:128496-130861 |
| gene1022 | gene1022 | SAOUHSC_01089 | NC_007795.1:1050054-1050378 |
| gene300 | gene300 | SAOUHSC_00325 | NC_007795.1:337879-339960 |
| gene109 | gene109 | SAOUHSC_00119 | NC_007795.1:124502-125612 |
| gene2586 | gene2586 | SAOUHSC_02744 | NC_007795.1:2522374-2524233 |

| | | | |
|---|---|---|---|
| gene1692 | gene1692 | SAOUHSC_01821 | NC_007795.1:1726827-1727775 |
| gene1806 | gene1806 | SAOUHSC_01945 | NC_007795.1:1849364-1851510 |
| gene2441 | gene2441 | SAOUHSC_02587 | NC_007795.1:2378948-2379686 |
| gene1289 | gene1289 | SAOUHSC_01380 | NC_007795.1:1321710-1324974 |
| gene2044 | gene2044 | SAOUHSC_02160 | NC_007795.1:2029296-2029587 |
| gene2610 | gene2610 | SAOUHSC_02771 | NC_007795.1:2548408-2548774 |
| gene187 | gene187 | SAOUHSC_00201 | NC_007795.1:224234-225647 |
| gene2848 | gene2848 | SAOUHSC_03028 | NC_007795.1:2800599-2801064 |
| gene131 | gene131 | SAOUHSC_00142 | NC_007795.1:145936-146962 |
| gene550 | gene550 | SAOUHSC_00586 | NC_007795.1:587590-588232 |
| gene724 | gene724 | SAOUHSC_00775 | NC_007795.1:755317-756201 |
| gene2631 | gene2631 | SAOUHSC_02794 | NC_007795.1:2567391-2567658 |
| gene2708 | gene2708 | SAOUHSC_02877 | NC_007795.1:2650766-2652275 |
| gene611 | gene611 | SAOUHSC_00658 | NC_007795.1:645294-646234 |
| gene2583 | gene2583 | SAOUHSC_02741 | NC_007795.1:2520721-2522358 |
| gene2854 | gene2854 | SAOUHSC_03035 | NC_007795.1:2807192-2807705 |
| gene1091 | gene1091 | SAOUHSC_01165 | NC_007795.1:1114346-1115654 |
| gene552 | gene552 | SAOUHSC_00588 | NC_007795.1:589011-589641 |
| gene1785 | gene1785 | SAOUHSC_01921 | NC_007795.1:1829135-1829480 |
| gene2602 | gene2602 | SAOUHSC_02763 | NC_007795.1:2539488-2542844 |
| gene2135 | gene2135 | SAOUHSC_02265 | NC_007795.1:2096005-2096632 |
| gene991 | gene991 | SAOUHSC_01054 | NC_007795.1:1021616-1022231 |
| gene78 | gene78 | SAOUHSC_00085 | NC_007795.1:91619-92246 |
| gene108 | gene108 | SAOUHSC_00118 | NC_007795.1:121648-124490 |
| gene602 | gene602 | SAOUHSC_00647 | NC_007795.1:636245-637973 |
| gene1787 | gene1787 | SAOUHSC_01923 | NC_007795.1:1830346-1830988 |
| gene2037 | gene2037 | SAOUHSC_02152 | NC_007795.1:2023147-2025437 |
| gene223 | gene223 | SAOUHSC_00240 | NC_007795.1:260849-261254 |
| gene2564 | gene2564 | SAOUHSC_02719 | NC_007795.1:2500218-2501982 |
| gene1044 | gene1044 | SAOUHSC_01112 | NC_007795.1:1071182-1071636 |
| gene640 | gene640 | SAOUHSC_00687 | NC_007795.1:672286-672829 |
| gene286 | gene286 | ulaA | NC_007795.1:324575-325922 |
| gene2585 | gene2585 | SAOUHSC_02743 | NC_007795.1:2522374-2524233 |
| gene2019 | gene2019 | SAOUHSC_02134 | NC_007795.1:2007647-2008724 |
| gene1784 | gene1784 | SAOUHSC_01920 | NC_007795.1:1828468-1829095 |
| gene1021 | gene1021 | SAOUHSC_01088 | NC_007795.1:1049301-1050036 |
| gene2039 | gene2039 | SAOUHSC_02154 | NC_007795.1:2025437-2026711 |
| gene2591 | gene2591 | SAOUHSC_02751 | NC_007795.1:2527926-2529279 |
| gene163 | gene163 | SAOUHSC_00176 | NC_007795.1:192600-193872 |
| gene2827 | gene2827 | SAOUHSC_03006 | NC_007795.1:2779035-2781078 |
| gene2045 | gene2045 | SAOUHSC_02161 | NC_007795.1:2029597-2031352 |
| gene2584 | gene2584 | SAOUHSC_02742 | NC_007795.1:2520721-2522358 |
| gene2604 | gene2604 | SAOUHSC_02765 | NC_007795.1:2539488-2542844 |
| gene1073 | gene1073 | SAOUHSC_01144 | NC_007795.1:1093491-1096108 |
| gene2617 | gene2617 | SAOUHSC_02778 | NC_007795.1:2554407-2555103 |
| gene2552 | gene2552 | SAOUHSC_02708 | NC_007795.1:2489228-2490158 |
| gene1807 | gene1807 | SAOUHSC_01947 | NC_007795.1:1849364-1851510 |
| gene1060 | gene1060 | SAOUHSC_01130 | NC_007795.1:1083816-1085307 |
| gene2657 | gene2657 | SAOUHSC_02822 | NC_007795.1:2599921-2601886 |
| gene1059 | gene1059 | SAOUHSC_01129 | NC_007795.1:1082646-1083579 |
| gene2421 | gene2421 | SAOUHSC_02565 | NC_007795.1:2355859-2357310 |

| | | | |
|---|---|---|---|
| gene1607 | gene1607 | SAOUHSC_01730 | NC_007795.1:1635848-1636031 |
| gene2603 | gene2603 | SAOUHSC_02764 | NC_007795.1:2539488-2542844 |
| gene2014 | gene2014 | SAOUHSC_02129 | NC_007795.1:2003535-2003862 |
| gene167 | gene167 | SAOUHSC_00180 | NC_007795.1:197158-198199 |
| gene1813 | gene1813 | SAOUHSC_01953 | NC_007795.1:1857744-1857888 |
| gene1636 | gene1636 | SAOUHSC_01763 | NC_007795.1:1662731-1663388 |
| gene134 | gene134 | SAOUHSC_00145 | NC_007795.1:156232-156877 |
| gene168 | gene168 | SAOUHSC_00181 | NC_007795.1:198253-199222 |
| gene2648 | gene2648 | SAOUHSC_02812 | NC_007795.1:2590644-2591061 |
| gene668 | gene668 | SAOUHSC_00716 | NC_007795.1:699193-701383 |
| gene2420 | gene2420 | SAOUHSC_02564 | NC_007795.1:2355859-2357310 |
| gene2304 | gene2304 | SAOUHSC_02436 | NC_007795.1:2262004-2263981 |
| gene2594 | gene2594 | SAOUHSC_02754 | NC_007795.1:2530923-2532355 |
| gene164 | gene164 | SAOUHSC_00177 | NC_007795.1:193874-195143 |
| gene1231 | gene1231 | SAOUHSC_01318 | NC_007795.1:1261216-1261792 |
| gene1606 | gene1606 | SAOUHSC_01729 | NC_007795.1:1635662-1635809 |
| gene2036 | gene2036 | SAOUHSC_02151 | NC_007795.1:2023147-2025437 |
| gene2843 | gene2843 | SAOUHSC_03023 | NC_007795.1:2796257-2797229 |
| gene1527 | gene1527 | SAOUHSC_01643 | NC_007795.1:1559315-1560290 |
| gene222 | gene222 | SAOUHSC_00239 | NC_007795.1:259907-260822 |
| gene2020 | gene2020 | SAOUHSC_02135 | NC_007795.1:2008743-2009538 |
| gene2038 | gene2038 | SAOUHSC_02153 | NC_007795.1:2023147-2025437 |
| gene2616 | gene2616 | SAOUHSC_02777 | NC_007795.1:2553734-2554157 |
| gene2643 | gene2643 | SAOUHSC_02806 | NC_007795.1:2584906-2586265 |
| gene591 | gene591 | SAOUHSC_00636 | NC_007795.1:624571-627080 |
| gene549 | gene549 | SAOUHSC_00585 | NC_007795.1:586907-587465 |
| gene2649 | gene2649 | SAOUHSC_02813 | NC_007795.1:2591199-2593093 |
| gene2613 | gene2613 | SAOUHSC_02774 | NC_007795.1:2551737-2552160 |
| gene1808 | gene1808 | SAOUHSC_01948 | NC_007795.1:1849364-1851510 |
| gene1809 | gene1809 | SAOUHSC_01949 | NC_007795.1:1851532-1852906 |
| gene1212 | gene1212 | SAOUHSC_01297 | NC_007795.1:1249421-1249670 |
| gene166 | gene166 | SAOUHSC_00179 | NC_007795.1:196057-197134 |
| gene2533 | gene2533 | SAOUHSC_02688 | NC_007795.1:2474416-2474599 |
| gene2697 | gene2697 | SAOUHSC_02865 | NC_007795.1:2638196-2638424 |
| gene514 | gene514 | SAOUHSC_00544 | NC_007795.1:547607-550738 |
| gene592 | gene592 | SAOUHSC_00637 | NC_007795.1:624571-627080 |
| gene1015 | gene1015 | SAOUHSC_01081 | NC_007795.1:1044367-1045420 |
| gene104 | capA | SAOUHSC_00114 | NC_007795.1:119491-120160 |
| gene1637 | gene1637 | SAOUHSC_01764 | NC_007795.1:1663414-1664122 |
| gene107 | gene107 | SAOUHSC_00117 | NC_007795.1:121648-124490 |
| gene1292 | gene1292 | SAOUHSC_01383 | NC_007795.1:1325826-1327641 |
| gene380 | gene380 | SAOUHSC_00410 | NC_007795.1:410721-411924 |
| gene771 | gene771 | SAOUHSC_00827 | NC_007795.1:804681-805242 |
| gene1221 | gene1221 | SAOUHSC_01307 | NC_007795.1:1252407-1253433 |
| gene2255 | gene2255 | SAOUHSC_02387 | NC_007795.1:2208189-2208855 |
| gene2696 | gene2696 | SAOUHSC_02864 | NC_007795.1:2636190-2638185 |
| gene162 | gene162 | SAOUHSC_00175 | NC_007795.1:191490-192588 |
| gene2380 | gene2380 | SAOUHSC_02519 | NC_007795.1:2322845-2323733 |
| gene2185 | gene2185 | SAOUHSC_02314 | NC_007795.1:2144445-2147798 |
| gene330 | gene330 | SAOUHSC_00356 | NC_007795.1:365422-365995 |
| gene1777 | gene1777 | SAOUHSC_01913 | NC_007795.1:1822472-1822946 |

| | | | |
|---|---|---|---|
| gene105 | gene105 | SAOUHSC_00115 | NC_007795.1:120175-120862 |
| gene133 | gene133 | SAOUHSC_00144 | NC_007795.1:149044-156220 |
| gene2709 | gene2709 | SAOUHSC_02879 | NC_007795.1:2652286-2653150 |
| gene2512 | gene2512 | SAOUHSC_02666 | NC_007795.1:2451288-2451645 |
| gene165 | gene165 | SAOUHSC_00178 | NC_007795.1:195144-195984 |
| gene2419 | gene2419 | SAOUHSC_02563 | NC_007795.1:2354712-2355847 |
| gene2695 | gene2695 | SAOUHSC_02863 | NC_007795.1:2636007-2636178 |
| gene2644 | gene2644 | SAOUHSC_02808 | NC_007795.1:2586381-2587935 |
| gene352 | gene352 | SAOUHSC_00378 | NC_007795.1:385389-385662 |
| gene1489 | gene1489 | SAOUHSC_01602 | NC_007795.1:1527875-1528895 |
| gene106 | gene106 | SAOUHSC_00116 | NC_007795.1:120864-121629 |
| gene2799 | gene2799 | SAOUHSC_02976 | NC_007795.1:2740026-2740965 |
| gene2673 | gene2673 | SAOUHSC_02839 | NC_007795.1:2614043-2614943 |
| gene332 | gene332 | SAOUHSC_00358 | NC_007795.1:366695-366947 |
| gene262 | gene262 | SAOUHSC_00284 | NC_007795.1:299298-300189 |
| gene2798 | gene2798 | SAOUHSC_02975 | NC_007795.1:2738060-2740013 |
| gene1058 | gene1058 | SAOUHSC_01128 | NC_007795.1:1081622-1082624 |
| gene276 | gene276 | SAOUHSC_00300 | NC_007795.1:314325-316398 |
| gene2712 | gene2712 | SAOUHSC_02882 | NC_007795.1:2654319-2656294 |
| gene195 | gene195 | SAOUHSC_00211 | NC_007795.1:231501-232437 |
| gene2710 | gene2710 | SAOUHSC_02880 | NC_007795.1:2653186-2654314 |
| gene582 | gene582 | SAOUHSC_00624 | NC_007795.1:614349-614910 |
| gene2418 | gene2418 | ureE | NC_007795.1:2354712-2355847 |
| gene762 | gene762 | SAOUHSC_00818 | NC_007795.1:800033-800720 |
| gene2415 | gene2415 | ureA | NC_007795.1:2352261-2352564 |
| gene132 | gene132 | SAOUHSC_00143 | NC_007795.1:147347-148598 |
| gene146 | gene146 | murQ | NC_007795.1:169308-171245 |
| gene1810 | gene1810 | SAOUHSC_01950 | NC_007795.1:1852915-1853434 |
| gene1352 | gene1352 | SAOUHSC_01456 | NC_007795.1:1415366-1416707 |
| gene2186 | gene2186 | SAOUHSC_02315 | NC_007795.1:2144445-2147798 |
| gene1052 | gene1052 | SAOUHSC_01121 | NC_007795.1:1076412-1077372 |
| gene1811 | gene1811 | SAOUHSC_01951 | NC_007795.1:1853449-1857680 |
| gene2416 | gene2416 | ureB | NC_007795.1:2352577-2354700 |
| gene2632 | gene2632 | SAOUHSC_02795 | NC_007795.1:2567801-2568110 |
| gene529 | gene529 | SAOUHSC_00560 | NC_007795.1:568931-569201 |
| gene2675 | gene2675 | SAOUHSC_02841 | NC_007795.1:2615639-2616635 |
| gene2013 | gene2013 | SAOUHSC_02127 | NC_007795.1:2002338-2003505 |
| gene2417 | gene2417 | ureC | NC_007795.1:2352577-2354700 |
| gene2674 | gene2674 | SAOUHSC_02840 | NC_007795.1:2614956-2615637 |
| gene1065 | gene1065 | SAOUHSC_01135 | NC_007795.1:1088442-1088577 |
| gene2711 | gene2711 | SAOUHSC_02881 | NC_007795.1:2654319-2656294 |
| gene2115 | gene2115 | SAOUHSC_02241 | NC_007795.1:2075692-2076709 |
| gene147 | gene147 | SAOUHSC_00158 | NC_007795.1:171256-172711 |
| gene1066 | gene1066 | SAOUHSC_01136 | NC_007795.1:1088633-1088768 |
| gene1220 | gene1220 | SAOUHSC_01306 | NC_007795.1:1251667-1251859 |
| gene2306 | gene2306 | SAOUHSC_02441 | NC_007795.1:2266096-2266606 |
| gene1812 | gene1812 | SAOUHSC_01952 | NC_007795.1:1853449-1857680 |
| gene145 | gene145 | SAOUHSC_00156 | NC_007795.1:169308-171245 |
| gene148 | gene148 | SAOUHSC_00160 | NC_007795.1:172713-173589 |
| gene2116 | gene2116 | SAOUHSC_02243 | NC_007795.1:2076730-2077786 |
| gene1814 | gene1814 | SAOUHSC_01954 | NC_007795.1:1859009-1859993 |

| | | | |
|---|---|---|---|
| gene2308 | gene2308 | SAOUHSC_02443 | NC_007795.1:2266920-2267457 |
| gene2307 | gene2307 | SAOUHSC_02442 | NC_007795.1:2266668-2266908 |
| gene788 | gene788 | SAOUHSC_00845 | NC_007795.1:816025-816220 |
| gene2131 | gene2131 | SAOUHSC_02260 | NC_007795.1:2093503-2093638 |
| gene103 | gene103 | SAOUHSC_00113 | NC_007795.1:116537-119147 |
| gene1815 | gene1815 | SAOUHSC_01955 | NC_007795.1:1859994-1860900 |
| gene2554 | gene2554 | SAOUHSC_02710 | NC_007795.1:2491674-2492652 |
| gene2553 | gene2553 | SAOUHSC_02709 | NC_007795.1:2490725-2491673 |
| gene1805 | gene1805 | SAOUHSC_01944 | NC_007795.1:1848324-1848891 |
| gene581 | gene581 | SAOUHSC_00622 | NC_007795.1:613746-613971 |
| gene930 | gene930 | SAOUHSC_00988 | NC_007795.1:959415-960426 |
| gene928 | gene928 | SAOUHSC_00986 | NC_007795.1:957785-958115 |
| gene49 | gene49 | SAOUHSC_00051 | NC_007795.1:54254-55241 |
| gene929 | gene929 | SAOUHSC_00987 | NC_007795.1:958152-959334 |
| gene904 | gene904 | SAOUHSC_00961 | NC_007795.1:938929-939460 |
| gene1802 | gene1802 | SAOUHSC_01939 | NC_007795.1:1845029-1845749 |
| gene1803 | gene1803 | SAOUHSC_01941 | NC_007795.1:1845806-1846529 |
| gene1804 | gene1804 | SAOUHSC_01942 | NC_007795.1:1846653-1847361 |
| gene1798 | gene1798 | SAOUHSC_01935 | NC_007795.1:1842445-1843165 |
| gene1801 | gene1801 | SAOUHSC_01938 | NC_007795.1:1844133-1844909 |
| gene1799 | gene1799 | SAOUHSC_01936 | NC_007795.1:1843315-1844032 |

| | | | NCTC82325-4 | MUTANT | | |
|---|---|---|---|---|---|---|
| sample_1 | sample_2 | status | value_1 | value_2 | log2(fold_cha | test_stat |
| mut_A | wt_R | OK | 0 | 27.1583 | inf | -nan |
| mut_A | wt_R | OK | 0 | 3.30855 | inf | -nan |
| mut_A | wt_R | OK | 0 | 134.407 | inf | -nan |
| mut_A | wt_R | OK | 179.999 | 0 | -inf | -nan |
| mut_A | wt_R | OK | 312.801 | 0 | -inf | -nan |
| mut_A | wt_R | OK | 2647.05 | 0 | -inf | -nan |
| mut_A | wt_R | OK | 195.696 | 3255.41 | 4.05615 | 16.7546 |
| mut_A | wt_R | OK | 22.9439 | 276.812 | 3.59272 | 6.52882 |
| mut_A | wt_R | OK | 306.813 | 2403.5 | 2.96971 | 12.1867 |
| mut_A | wt_R | OK | 9.08806 | 68.5361 | 2.91482 | 9.4386 |
| mut_A | wt_R | OK | 2524.7 | 15010.1 | 2.57175 | 3.83527 |
| mut_A | wt_R | OK | 3011.51 | 17334.9 | 2.52512 | 10.1783 |
| mut_A | wt_R | OK | 2669.08 | 15160.8 | 2.50593 | 9.50721 |
| mut_A | wt_R | OK | 2247.57 | 12742.8 | 2.50325 | 10.1494 |
| mut_A | wt_R | OK | 3184.59 | 17830.4 | 2.48516 | 10.0955 |
| mut_A | wt_R | OK | 2489.06 | 13900.6 | 2.48147 | 10.0891 |
| mut_A | wt_R | OK | 2285.03 | 12610.9 | 2.46438 | 9.93336 |
| mut_A | wt_R | OK | 2664.57 | 14331.3 | 2.4272 | 9.92546 |
| mut_A | wt_R | OK | 72.3003 | 388.027 | 2.42408 | 9.88015 |
| mut_A | wt_R | OK | 2268.16 | 12013.3 | 2.40504 | 9.86175 |
| mut_A | wt_R | OK | 1738.25 | 9143.82 | 2.39516 | 9.83897 |
| mut_A | wt_R | OK | 49.9851 | 261.399 | 2.38668 | 9.51847 |
| mut_A | wt_R | OK | 3024.07 | 15764.5 | 2.38211 | 9.73466 |
| mut_A | wt_R | OK | 2578.81 | 13376.4 | 2.37491 | 9.69514 |
| mut_A | wt_R | OK | 1721.24 | 8808.25 | 2.35541 | 9.65368 |
| mut_A | wt_R | OK | 2606.91 | 13078.9 | 2.32682 | 9.54029 |
| mut_A | wt_R | OK | 211.35 | 1043.11 | 2.30318 | 9.48112 |
| mut_A | wt_R | OK | 1964.09 | 9670.1 | 2.29967 | 9.41993 |
| mut_A | wt_R | OK | 2408.19 | 11832.7 | 2.29675 | 9.41515 |
| mut_A | wt_R | OK | 3194.53 | 15609.6 | 2.28876 | 9.39936 |
| mut_A | wt_R | OK | 8.61057 | 41.9795 | 2.28551 | 7.39334 |
| mut_A | wt_R | OK | 3127.97 | 15211.5 | 2.28187 | 9.33187 |
| mut_A | wt_R | OK | 1459.44 | 6951.13 | 2.25183 | 9.23539 |
| mut_A | wt_R | OK | 117.92 | 560.681 | 2.24937 | 9.18681 |
| mut_A | wt_R | OK | 18.3886 | 85.5803 | 2.21847 | 8.0579 |
| mut_A | wt_R | OK | 1884.12 | 8755.96 | 2.21637 | 3.72273 |
| mut_A | wt_R | OK | 2427.25 | 11269.1 | 2.21497 | 9.07485 |
| mut_A | wt_R | OK | 271.89 | 1259.81 | 2.21211 | 9.06692 |
| mut_A | wt_R | OK | 171.833 | 789.217 | 2.19942 | 9.07467 |
| mut_A | wt_R | OK | 19.9601 | 91.3027 | 2.19354 | 8.63719 |
| mut_A | wt_R | OK | 3716.16 | 16971.2 | 2.19121 | 8.83906 |
| mut_A | wt_R | OK | 2381.97 | 10863.3 | 2.18923 | 3.76172 |
| mut_A | wt_R | OK | 4806.78 | 21653.3 | 2.17145 | 8.79613 |
| mut_A | wt_R | OK | 2587.6 | 11554.7 | 2.15879 | 7.96564 |
| mut_A | wt_R | OK | 581.525 | 2573.21 | 2.14566 | 8.69875 |
| mut_A | wt_R | OK | 2328.51 | 10303.2 | 2.14562 | 8.81164 |
| mut_A | wt_R | OK | 2743.04 | 12077.4 | 2.13846 | 8.72597 |
| mut_A | wt_R | OK | 2093.41 | 9123.91 | 2.1238 | 7.87732 |
| mut_A | wt_R | OK | 72.9631 | 314.806 | 2.10923 | 8.42071 |

| | | | | | | |
|---|---|---|---|---|---|---|
| mut_A | wt_R | OK | 2807.17 | 11485.9 | 2.03267 | 8.33463 |
| mut_A | wt_R | OK | 2116.2 | 8631.55 | 2.02815 | 8.27803 |
| mut_A | wt_R | OK | 331.456 | 1345.21 | 2.02095 | 8.28988 |
| mut_A | wt_R | OK | 11271.9 | 44950.6 | 1.9956 | 6.9178 |
| mut_A | wt_R | OK | 2209.01 | 8672.14 | 1.97299 | 8.07524 |
| mut_A | wt_R | OK | 20.7329 | 80.0338 | 1.94869 | 7.60235 |
| mut_A | wt_R | OK | 29.1286 | 111.587 | 1.93766 | 6.8574 |
| mut_A | wt_R | OK | 25.802 | 97.7484 | 1.92159 | 7.11941 |
| mut_A | wt_R | OK | 135.149 | 509.517 | 1.91458 | 6.60275 |
| mut_A | wt_R | OK | 2783.95 | 10238 | 1.87873 | 7.74717 |
| mut_A | wt_R | OK | 916.178 | 3292.91 | 1.84566 | 7.45517 |
| mut_A | wt_R | OK | 47.37 | 166.317 | 1.81189 | 7.39008 |
| mut_A | wt_R | OK | 1605.36 | 5546.97 | 1.78881 | 7.17147 |
| mut_A | wt_R | OK | 6509.42 | 22474.9 | 1.78771 | 7.23181 |
| mut_A | wt_R | OK | 27.0405 | 91.773 | 1.76295 | 6.00004 |
| mut_A | wt_R | OK | 200.843 | 674.284 | 1.74729 | 7.21221 |
| mut_A | wt_R | OK | 139.742 | 461.794 | 1.72449 | 3.49157 |
| mut_A | wt_R | OK | 247.537 | 816.494 | 1.7218 | 7.06309 |
| mut_A | wt_R | OK | 42.5985 | 140.345 | 1.7201 | 6.95518 |
| mut_A | wt_R | OK | 877.808 | 2886.81 | 1.7175 | 7.06322 |
| mut_A | wt_R | OK | 420.362 | 1368.01 | 1.70237 | 7.01478 |
| mut_A | wt_R | OK | 113.763 | 366.494 | 1.68776 | 5.3864 |
| mut_A | wt_R | OK | 29.0582 | 93.2719 | 1.6825 | 6.26348 |
| mut_A | wt_R | OK | 721.608 | 2306.66 | 1.67652 | 6.84073 |
| mut_A | wt_R | OK | 116.335 | 367.75 | 1.66044 | 6.83752 |
| mut_A | wt_R | OK | 2034.08 | 6394.74 | 1.65251 | 6.79421 |
| mut_A | wt_R | OK | 3503.79 | 10854.1 | 1.63125 | 6.72758 |
| mut_A | wt_R | OK | 64.3811 | 199.233 | 1.62975 | 5.52191 |
| mut_A | wt_R | OK | 177.466 | 541.001 | 1.60808 | 5.67082 |
| mut_A | wt_R | OK | 19.919 | 60.3984 | 1.60037 | 2.19515 |
| mut_A | wt_R | OK | 40.9755 | 123.797 | 1.59514 | 6.39026 |
| mut_A | wt_R | OK | 176.008 | 531.612 | 1.59474 | 6.53893 |
| mut_A | wt_R | OK | 33.6507 | 101.186 | 1.5883 | 6.38343 |
| mut_A | wt_R | OK | 1648.22 | 4927.18 | 1.57985 | 6.48266 |
| mut_A | wt_R | OK | 10.594 | 30.9026 | 1.54449 | 6.07791 |
| mut_A | wt_R | OK | 14.0127 | 40.6753 | 1.53742 | 5.86864 |
| mut_A | wt_R | OK | 18.6761 | 54.0742 | 1.53375 | 5.29532 |
| mut_A | wt_R | OK | 1912.23 | 5518.05 | 1.5289 | 6.30875 |
| mut_A | wt_R | OK | 50.4583 | 144.706 | 1.51996 | 6.18364 |
| mut_A | wt_R | OK | 21.9898 | 63.0382 | 1.51939 | 6.23198 |
| mut_A | wt_R | OK | 5.21324 | 14.8756 | 1.51269 | 2.89536 |
| mut_A | wt_R | OK | 0.427284 | 1.2124 | 1.5046 | 2.26334 |
| mut_A | wt_R | OK | 40.3193 | 113.89 | 1.4981 | 4.85983 |
| mut_A | wt_R | OK | 290.148 | 818.001 | 1.49531 | 6.16067 |
| mut_A | wt_R | OK | 129.04 | 355.249 | 1.46101 | 5.01725 |
| mut_A | wt_R | OK | 55.2502 | 151.429 | 1.45459 | 3.94188 |
| mut_A | wt_R | OK | 20.7918 | 56.9135 | 1.45276 | 5.14843 |
| mut_A | wt_R | OK | 33.3822 | 91.0264 | 1.44721 | 5.67875 |
| mut_A | wt_R | OK | 140.186 | 382.165 | 1.44685 | 5.94418 |
| mut_A | wt_R | OK | 15.1112 | 41.0593 | 1.44209 | 5.62358 |
| mut_A | wt_R | OK | 140.392 | 380.615 | 1.43887 | 5.89184 |

| | | | | | | |
|---|---|---|---|---|---|---|
| mut_A | wt_R | OK | 193.527 | 523.011 | 1.4343 | 5.18371 |
| mut_A | wt_R | OK | 190.67 | 512.653 | 1.4269 | 5.83874 |
| mut_A | wt_R | OK | 135.332 | 361.713 | 1.41834 | 3.66083 |
| mut_A | wt_R | OK | 400.841 | 1067.42 | 1.41303 | 5.84099 |
| mut_A | wt_R | OK | 95.8352 | 253.995 | 1.40617 | 3.03866 |
| mut_A | wt_R | OK | 297.155 | 785.794 | 1.40294 | 5.73269 |
| mut_A | wt_R | OK | 12.7608 | 33.6802 | 1.40019 | 4.91863 |
| mut_A | wt_R | OK | 2.34643 | 6.1756 | 1.39612 | 2.48466 |
| mut_A | wt_R | OK | 162.549 | 427.487 | 1.39501 | 2.73029 |
| mut_A | wt_R | OK | 0.415289 | 1.08876 | 1.3905 | 2.02644 |
| mut_A | wt_R | OK | 31.118 | 81.4942 | 1.38895 | 5.46694 |
| mut_A | wt_R | OK | 184.714 | 480.459 | 1.37912 | 5.63973 |
| mut_A | wt_R | OK | 413.512 | 1062.85 | 1.36194 | 5.60737 |
| mut_A | wt_R | OK | 63.0665 | 161.614 | 1.3576 | 3.02213 |
| mut_A | wt_R | OK | 6241.96 | 15978.5 | 1.35606 | 5.54732 |
| mut_A | wt_R | OK | 992.115 | 2531.95 | 1.35167 | 5.55738 |
| mut_A | wt_R | OK | 44.1098 | 112.543 | 1.35131 | 4.36212 |
| mut_A | wt_R | OK | 1422.37 | 3618.92 | 1.34726 | 5.53577 |
| mut_A | wt_R | OK | 1392.27 | 3529.46 | 1.34201 | 5.52181 |
| mut_A | wt_R | OK | 82.629 | 209.451 | 1.3419 | 5.44503 |
| mut_A | wt_R | OK | 36.5843 | 92.3736 | 1.33625 | 3.39921 |
| mut_A | wt_R | OK | 230.815 | 582.097 | 1.33452 | 5.50516 |
| mut_A | wt_R | OK | 95.7709 | 240.69 | 1.32952 | 5.01976 |
| mut_A | wt_R | OK | 71.26 | 177.957 | 1.32036 | 4.00563 |
| mut_A | wt_R | OK | 79.8607 | 199.055 | 1.31761 | 5.40751 |
| mut_A | wt_R | OK | 12.4981 | 31.0813 | 1.31434 | 2.51246 |
| mut_A | wt_R | OK | 242.968 | 603.911 | 1.31357 | 5.41649 |
| mut_A | wt_R | OK | 120.534 | 298.055 | 1.30614 | 3.29071 |
| mut_A | wt_R | OK | 71.2455 | 176.151 | 1.30595 | 3.48398 |
| mut_A | wt_R | OK | 128.891 | 317.859 | 1.30224 | 5.34644 |
| mut_A | wt_R | OK | 79.7163 | 195.958 | 1.2976 | 4.91415 |
| mut_A | wt_R | OK | 12.071 | 29.6696 | 1.29744 | 4.56728 |
| mut_A | wt_R | OK | 140.546 | 344.98 | 1.29547 | 5.31506 |
| mut_A | wt_R | OK | 188.915 | 463.593 | 1.29512 | 4.06119 |
| mut_A | wt_R | OK | 83.8684 | 205.618 | 1.29377 | 5.2142 |
| mut_A | wt_R | OK | 331.867 | 812.279 | 1.29137 | 5.32807 |
| mut_A | wt_R | OK | 10.0586 | 24.6066 | 1.29062 | 4.40669 |
| mut_A | wt_R | OK | 4.7706 | 11.6516 | 1.28829 | 1.79576 |
| mut_A | wt_R | OK | 191.984 | 468.678 | 1.28761 | 4.3275 |
| mut_A | wt_R | OK | 245.149 | 598.227 | 1.28703 | 5.26737 |
| mut_A | wt_R | OK | 28.9332 | 70.4725 | 1.28433 | 1.48192 |
| mut_A | wt_R | OK | 2.39355 | 5.81107 | 1.27965 | 2.95666 |
| mut_A | wt_R | OK | 1378.76 | 3344.38 | 1.27837 | 5.24631 |
| mut_A | wt_R | OK | 11.3355 | 27.3865 | 1.27262 | 2.83687 |
| mut_A | wt_R | OK | 159.299 | 383.103 | 1.26599 | 5.00924 |
| mut_A | wt_R | OK | 47.253 | 113.327 | 1.26201 | 2.26389 |
| mut_A | wt_R | OK | 303.054 | 725.047 | 1.2585 | 5.20265 |
| mut_A | wt_R | OK | 39.0062 | 92.8432 | 1.25109 | 4.35928 |
| mut_A | wt_R | OK | 3.88072 | 9.20243 | 1.24569 | 3.05504 |
| mut_A | wt_R | OK | 69.0998 | 163.694 | 1.24425 | 4.88304 |
| mut_A | wt_R | OK | 318.78 | 752.553 | 1.23923 | 5.12396 |

| | | | | | | |
|---|---|---|---|---|---|---|
| mut_A | wt_R | OK | 285.045 | 672.192 | 1.23768 | 5.08224 |
| mut_A | wt_R | OK | 5662.9 | 13327 | 1.23474 | 4.99756 |
| mut_A | wt_R | OK | 26.4847 | 62.3263 | 1.23468 | 3.16437 |
| mut_A | wt_R | OK | 49.4668 | 115.661 | 1.22537 | 4.00708 |
| mut_A | wt_R | OK | 67.7719 | 157.69 | 1.21833 | 4.00362 |
| mut_A | wt_R | OK | 5.13398 | 11.9226 | 1.21555 | 1.46967 |
| mut_A | wt_R | OK | 179.884 | 416.506 | 1.21127 | 4.95389 |
| mut_A | wt_R | OK | 694.918 | 1608.27 | 1.21059 | 4.99941 |
| mut_A | wt_R | OK | 49.3198 | 114.136 | 1.21052 | 4.93462 |
| mut_A | wt_R | OK | 77.6226 | 179.55 | 1.20983 | 2.56394 |
| mut_A | wt_R | OK | 6893.2 | 15938 | 1.20923 | 4.95004 |
| mut_A | wt_R | OK | 2199.03 | 5084 | 1.2091 | 4.99665 |
| mut_A | wt_R | OK | 4542 | 10443.1 | 1.20115 | 4.9439 |
| mut_A | wt_R | OK | 830.41 | 1906.92 | 1.19935 | 4.93328 |
| mut_A | wt_R | OK | 43.0341 | 98.7177 | 1.19783 | 4.79421 |
| mut_A | wt_R | OK | 26.0865 | 59.8097 | 1.19708 | 4.38229 |
| mut_A | wt_R | OK | 3803.7 | 8696.28 | 1.19299 | 4.91142 |
| mut_A | wt_R | OK | 2439.14 | 5552.85 | 1.18686 | 4.81427 |
| mut_A | wt_R | OK | 3340.38 | 7582.34 | 1.18263 | 4.83438 |
| mut_A | wt_R | OK | 50.2585 | 113.496 | 1.1752 | 4.74813 |
| mut_A | wt_R | OK | 85.7202 | 193.28 | 1.17298 | 4.81642 |
| mut_A | wt_R | OK | 188.173 | 423.678 | 1.17091 | 4.26097 |
| mut_A | wt_R | OK | 163.584 | 367.362 | 1.16717 | 4.74383 |
| mut_A | wt_R | OK | 47.225 | 105.674 | 1.162 | 4.0161 |
| mut_A | wt_R | OK | 142.877 | 319.124 | 1.15935 | 4.72574 |
| mut_A | wt_R | OK | 404.393 | 902.39 | 1.15799 | 4.72632 |
| mut_A | wt_R | OK | 46.2121 | 103.086 | 1.15751 | 4.61487 |
| mut_A | wt_R | OK | 2047.36 | 4560.05 | 1.15529 | 4.75366 |
| mut_A | wt_R | OK | 270.378 | 601.18 | 1.15282 | 4.76996 |
| mut_A | wt_R | OK | 55.6998 | 123.683 | 1.15091 | 2.32886 |
| mut_A | wt_R | OK | 44.4841 | 98.4773 | 1.1465 | 4.68488 |
| mut_A | wt_R | OK | 113.98 | 251.429 | 1.14137 | 4.68639 |
| mut_A | wt_R | OK | 136.996 | 301.138 | 1.1363 | 4.59347 |
| mut_A | wt_R | OK | 137.482 | 301.961 | 1.13512 | 4.63646 |
| mut_A | wt_R | OK | 29.5817 | 64.6888 | 1.12881 | 4.43412 |
| mut_A | wt_R | OK | 1.00592 | 2.19785 | 1.12758 | 1.33692 |
| mut_A | wt_R | OK | 352.794 | 769.223 | 1.12458 | 4.64414 |
| mut_A | wt_R | OK | 123.551 | 268.793 | 1.12139 | 3.69585 |
| mut_A | wt_R | OK | 102.25 | 222.076 | 1.11895 | 4.59475 |
| mut_A | wt_R | OK | 145.567 | 314.705 | 1.11231 | 4.54436 |
| mut_A | wt_R | OK | 82.8031 | 178.781 | 1.11043 | 4.52356 |
| mut_A | wt_R | OK | 55.8969 | 120.475 | 1.10789 | 4.43283 |
| mut_A | wt_R | OK | 6.24652 | 13.419 | 1.10316 | 2.89498 |
| mut_A | wt_R | OK | 57.9129 | 124.203 | 1.10074 | 4.34642 |
| mut_A | wt_R | OK | 94.3441 | 202.331 | 1.10071 | 3.85432 |
| mut_A | wt_R | OK | 213.917 | 456.829 | 1.0946 | 4.41287 |
| mut_A | wt_R | OK | 4237.84 | 9048.77 | 1.09439 | 4.2293 |
| mut_A | wt_R | OK | 209.272 | 445.847 | 1.09117 | 3.80071 |
| mut_A | wt_R | OK | 56.9923 | 121.242 | 1.08905 | 3.01221 |
| mut_A | wt_R | OK | 64.9092 | 137.75 | 1.08555 | 4.31089 |
| mut_A | wt_R | OK | 189.932 | 402.136 | 1.0822 | 4.42836 |

| | | | | | | |
|---|---|---|---|---|---|---|
| mut_A | wt_R | OK | 22.7 | 47.9568 | 1.07904 | 4.23281 |
| mut_A | wt_R | OK | 131.913 | 277.515 | 1.07298 | 4.40661 |
| mut_A | wt_R | OK | 57.1979 | 119.271 | 1.06021 | 3.87506 |
| mut_A | wt_R | OK | 264.807 | 551.962 | 1.05963 | 3.85061 |
| mut_A | wt_R | OK | 23.7469 | 49.4696 | 1.0588 | 3.87219 |
| mut_A | wt_R | OK | 168.065 | 348.833 | 1.05352 | 4.35876 |
| mut_A | wt_R | OK | 4016.12 | 8333.43 | 1.05311 | 4.31436 |
| mut_A | wt_R | OK | 1.77851 | 3.68972 | 1.05284 | 2.11415 |
| mut_A | wt_R | OK | 1715.81 | 3557.57 | 1.052 | 3.81277 |
| mut_A | wt_R | OK | 19.8511 | 41.0909 | 1.0496 | 1.90525 |
| mut_A | wt_R | OK | 190.412 | 393.842 | 1.0485 | 4.30246 |
| mut_A | wt_R | OK | 981.971 | 2028.21 | 1.04645 | 2.01343 |
| mut_A | wt_R | OK | 112.671 | 232.5 | 1.04511 | 4.3043 |
| mut_A | wt_R | OK | 4.44946 | 9.17124 | 1.04349 | 1.71943 |
| mut_A | wt_R | OK | 297.599 | 612.544 | 1.04144 | 4.29528 |
| mut_A | wt_R | OK | 151.952 | 312.408 | 1.03981 | 2.45198 |
| mut_A | wt_R | OK | 139.194 | 284.746 | 1.03258 | 4.09223 |
| mut_A | wt_R | OK | 151.38 | 308.873 | 1.02883 | 3.5548 |
| mut_A | wt_R | OK | 28.3524 | 57.6284 | 1.02331 | 3.56648 |
| mut_A | wt_R | OK | 117.838 | 239.499 | 1.02321 | 4.16348 |
| mut_A | wt_R | OK | 198.326 | 402.625 | 1.02157 | 4.18818 |
| mut_A | wt_R | OK | 72.7331 | 147.58 | 1.02081 | 4.1696 |
| mut_A | wt_R | OK | 200.723 | 407.203 | 1.02054 | 4.20388 |
| mut_A | wt_R | OK | 6.20687 | 12.5805 | 1.01925 | 2.53539 |
| mut_A | wt_R | OK | 73.9267 | 149.407 | 1.01508 | 4.07009 |
| mut_A | wt_R | OK | 224.682 | 454.033 | 1.01491 | 4.13644 |
| mut_A | wt_R | OK | 115.128 | 232.272 | 1.01258 | 4.01572 |
| mut_A | wt_R | OK | 2564.88 | 5171.67 | 1.01174 | 4.15692 |
| mut_A | wt_R | OK | 214.988 | 432.497 | 1.00844 | 4.12436 |
| mut_A | wt_R | OK | 25.075 | 50.3813 | 1.00664 | 3.89819 |
| mut_A | wt_R | OK | 647.126 | 1297.52 | 1.00364 | 4.14328 |
| mut_A | wt_R | OK | 257.921 | 516.681 | 1.00234 | 3.20639 |
| mut_A | wt_R | OK | 1.69668 | 3.39649 | 1.00133 | 2.59518 |
| mut_A | wt_R | OK | 2107.09 | 4216.9 | 1.00093 | 3.99754 |
| mut_A | wt_R | OK | 126.211 | 252.241 | 0.998972 | 4.07464 |
| mut_A | wt_R | OK | 68.3411 | 136.4 | 0.997017 | 4.01666 |
| mut_A | wt_R | OK | 170.56 | 339.962 | 0.995095 | 4.05521 |
| mut_A | wt_R | OK | 572.169 | 1139.59 | 0.994 | 4.02082 |
| mut_A | wt_R | OK | 48.5688 | 96.6277 | 0.992408 | 3.97816 |
| mut_A | wt_R | OK | 149.233 | 296.729 | 0.991578 | 4.06653 |
| mut_A | wt_R | OK | 8.24197 | 16.3593 | 0.989053 | 2.57733 |
| mut_A | wt_R | OK | 223.552 | 442.03 | 0.983535 | 4.06894 |
| mut_A | wt_R | OK | 101.376 | 200.177 | 0.981555 | 3.91432 |
| mut_A | wt_R | OK | 207.252 | 408.805 | 0.980028 | 2.20627 |
| mut_A | wt_R | OK | 593.177 | 1169.28 | 0.979084 | 3.61243 |
| mut_A | wt_R | OK | 190.171 | 374.034 | 0.975872 | 2.86 |
| mut_A | wt_R | OK | 35.934 | 70.612 | 0.974562 | 2.9258 |
| mut_A | wt_R | OK | 45.8099 | 89.8873 | 0.972458 | 3.92572 |
| mut_A | wt_R | OK | 159.774 | 313.186 | 0.970988 | 3.7142 |
| mut_A | wt_R | OK | 2585.62 | 5052.21 | 0.966405 | 3.9831 |
| mut_A | wt_R | OK | 503.318 | 982.697 | 0.965277 | 3.97795 |

| | | | | | | |
|---|---|---|---|---|---|---|
| mut_A | wt_R | OK | 66.2268 | 129.23 | 0.964458 | 3.82853 |
| mut_A | wt_R | OK | 129.889 | 253.216 | 0.96309 | 3.95815 |
| mut_A | wt_R | OK | 304.004 | 591.384 | 0.960005 | 3.95232 |
| mut_A | wt_R | OK | 522.829 | 1015.6 | 0.957923 | 3.95767 |
| mut_A | wt_R | OK | 32.103 | 62.3557 | 0.957815 | 3.78902 |
| mut_A | wt_R | OK | 404.844 | 782.682 | 0.951061 | 3.74688 |
| mut_A | wt_R | OK | 246.876 | 476.248 | 0.947929 | 3.91008 |
| mut_A | wt_R | OK | 3632.07 | 6980.39 | 0.942514 | 3.87057 |
| mut_A | wt_R | OK | 67.6997 | 130.078 | 0.94215 | 2.27607 |
| mut_A | wt_R | OK | 583.182 | 1118.96 | 0.940146 | 3.79831 |
| mut_A | wt_R | OK | 9.68142 | 18.563 | 0.939139 | 3.4349 |
| mut_A | wt_R | OK | 2018.48 | 3869.42 | 0.938849 | 3.88517 |
| mut_A | wt_R | OK | 177.018 | 338.152 | 0.933778 | 3.81763 |
| mut_A | wt_R | OK | 100.093 | 191.042 | 0.932547 | 2.77279 |
| mut_A | wt_R | OK | 82.6518 | 157.434 | 0.92963 | 3.80201 |
| mut_A | wt_R | OK | 428.214 | 815.211 | 0.928843 | 2.37912 |
| mut_A | wt_R | OK | 49.353 | 93.6425 | 0.924028 | 2.44999 |
| mut_A | wt_R | OK | 45.1968 | 85.7555 | 0.924008 | 3.44645 |
| mut_A | wt_R | OK | 55.8504 | 105.818 | 0.921944 | 1.8751 |
| mut_A | wt_R | OK | 5.51258 | 10.4436 | 0.921814 | 2.45189 |
| mut_A | wt_R | OK | 14.8155 | 28.0172 | 0.919204 | 3.40245 |
| mut_A | wt_R | OK | 8.28645 | 15.6601 | 0.918268 | 1.38334 |
| mut_A | wt_R | OK | 659.137 | 1245.03 | 0.91753 | 3.79484 |
| mut_A | wt_R | OK | 959.726 | 1811.2 | 0.91625 | 3.76559 |
| mut_A | wt_R | OK | 33.3982 | 63.0001 | 0.915583 | 3.08524 |
| mut_A | wt_R | OK | 51.5002 | 97.0121 | 0.913588 | 3.54037 |
| mut_A | wt_R | OK | 23.2574 | 43.804 | 0.913373 | 2.10236 |
| mut_A | wt_R | OK | 688.673 | 1296.93 | 0.913212 | 3.77358 |
| mut_A | wt_R | OK | 372.093 | 700.154 | 0.91201 | 2.59412 |
| mut_A | wt_R | OK | 638.141 | 1198.16 | 0.908869 | 3.75004 |
| mut_A | wt_R | OK | 1370.13 | 2565.33 | 0.904838 | 3.73191 |
| mut_A | wt_R | OK | 7005.75 | 13115.1 | 0.904617 | 3.7132 |
| mut_A | wt_R | OK | 16.4633 | 30.7908 | 0.903242 | 2.17908 |
| mut_A | wt_R | OK | 106.178 | 198.33 | 0.901413 | 3.68551 |
| mut_A | wt_R | OK | 499.295 | 928.961 | 0.895726 | 3.64607 |
| mut_A | wt_R | OK | 401.951 | 747.246 | 0.894564 | 3.51598 |
| mut_A | wt_R | OK | 167.901 | 310.708 | 0.887949 | 2.40068 |
| mut_A | wt_R | OK | 162.216 | 299.874 | 0.886445 | 3.64901 |
| mut_A | wt_R | OK | 524.437 | 968.196 | 0.884529 | 3.64537 |
| mut_A | wt_R | OK | 168.842 | 311.535 | 0.88372 | 3.65452 |
| mut_A | wt_R | OK | 184.313 | 339.742 | 0.882279 | 3.64734 |
| mut_A | wt_R | OK | 625.657 | 1152.93 | 0.881863 | 3.64282 |
| mut_A | wt_R | OK | 581.596 | 1070.45 | 0.880127 | 3.6371 |
| mut_A | wt_R | OK | 93.0156 | 170.559 | 0.874727 | 3.57264 |
| mut_A | wt_R | OK | 78.703 | 143.886 | 0.870438 | 2.79617 |
| mut_A | wt_R | OK | 347.154 | 633.699 | 0.868224 | 3.58898 |
| mut_A | wt_R | OK | 346.861 | 632.1 | 0.865798 | 2.96585 |
| mut_A | wt_R | OK | 1133.74 | 2054.51 | 0.857702 | 3.45085 |
| mut_A | wt_R | OK | 2928.94 | 5304.93 | 0.856956 | 3.5246 |
| mut_A | wt_R | OK | 290.641 | 525.705 | 0.855018 | 3.52406 |
| mut_A | wt_R | OK | 87.6462 | 158.532 | 0.855012 | 3.34774 |

| | | | | | | |
|---|---|---|---|---|---|---|
| mut_A | wt_R | OK | 312.641 | 563.926 | 0.850999 | 2.52001 |
| mut_A | wt_R | OK | 107.518 | 193.781 | 0.849853 | 3.27687 |
| mut_A | wt_R | OK | 153.515 | 276.557 | 0.849202 | 3.46399 |
| mut_A | wt_R | OK | 16.6233 | 29.845 | 0.844281 | 2.49389 |
| mut_A | wt_R | OK | 218.317 | 391.876 | 0.843974 | 3.25303 |
| mut_A | wt_R | OK | 355.206 | 636.837 | 0.842268 | 3.41935 |
| mut_A | wt_R | OK | 266.581 | 477.355 | 0.840488 | 3.46931 |
| mut_A | wt_R | OK | 2312.71 | 4140.86 | 0.84035 | 3.45064 |
| mut_A | wt_R | OK | 112.185 | 200.777 | 0.839709 | 1.50239 |
| mut_A | wt_R | OK | 570.727 | 1016.98 | 0.833419 | 1.97873 |
| mut_A | wt_R | OK | 1205.52 | 2142.87 | 0.82989 | 3.42769 |
| mut_A | wt_R | OK | 821.375 | 1455.91 | 0.825809 | 1.93817 |
| mut_A | wt_R | OK | 116.646 | 206.74 | 0.825687 | 3.37324 |
| mut_A | wt_R | OK | 522.989 | 926.324 | 0.824737 | 3.40888 |
| mut_A | wt_R | OK | 2460.47 | 4350.97 | 0.822403 | 3.38325 |
| mut_A | wt_R | OK | 260.016 | 459.348 | 0.820986 | 2.28519 |
| mut_A | wt_R | OK | 83.9636 | 148.157 | 0.819288 | 3.17721 |
| mut_A | wt_R | OK | 200.418 | 353.462 | 0.818542 | 3.16465 |
| mut_A | wt_R | OK | 87.3649 | 153.744 | 0.815404 | 3.14783 |
| mut_A | wt_R | OK | 70.2115 | 123.43 | 0.813916 | 3.32379 |
| mut_A | wt_R | OK | 129.313 | 227.176 | 0.812946 | 3.32108 |
| mut_A | wt_R | OK | 96.8815 | 169.959 | 0.810893 | 3.18283 |
| mut_A | wt_R | OK | 74.6881 | 130.933 | 0.809879 | 3.17 |
| mut_A | wt_R | OK | 607.495 | 1064.17 | 0.808784 | 3.34951 |
| mut_A | wt_R | OK | 84.7528 | 148.373 | 0.807895 | 2.37482 |
| mut_A | wt_R | OK | 271.329 | 474.782 | 0.807223 | 2.51353 |
| mut_A | wt_R | OK | 17.221 | 30.0911 | 0.805172 | 1.68481 |
| mut_A | wt_R | OK | 343.007 | 598.112 | 0.802178 | 3.31292 |
| mut_A | wt_R | OK | 407.401 | 709.067 | 0.799474 | 3.22991 |
| mut_A | wt_R | OK | 105.975 | 184.296 | 0.798302 | 1.36652 |
| mut_A | wt_R | OK | 669.726 | 1163.74 | 0.797121 | 2.58841 |
| mut_A | wt_R | OK | 376.802 | 654.716 | 0.797063 | 3.27494 |
| mut_A | wt_R | OK | 1085.39 | 1885.73 | 0.796918 | 3.29109 |
| mut_A | wt_R | OK | 87.7174 | 152.165 | 0.794702 | 3.18664 |
| mut_A | wt_R | OK | 287.755 | 498.993 | 0.79418 | 3.27391 |
| mut_A | wt_R | OK | 529.858 | 918.053 | 0.792971 | 3.28015 |
| mut_A | wt_R | OK | 44.889 | 77.4858 | 0.787569 | 2.91687 |
| mut_A | wt_R | OK | 57.1517 | 98.5299 | 0.785765 | 3.17017 |
| mut_A | wt_R | OK | 190.341 | 327.929 | 0.784797 | 1.78273 |
| mut_A | wt_R | OK | 827.907 | 1424.55 | 0.782969 | 3.23299 |
| mut_A | wt_R | OK | 635.529 | 1092.7 | 0.781869 | 3.2072 |
| mut_A | wt_R | OK | 228.568 | 392.579 | 0.780358 | 3.21501 |
| mut_A | wt_R | OK | 556.921 | 955.717 | 0.779111 | 3.22564 |
| mut_A | wt_R | OK | 138.781 | 237.476 | 0.774973 | 3.16348 |
| mut_A | wt_R | OK | 206.831 | 352.936 | 0.770956 | 3.15571 |
| mut_A | wt_R | OK | 17.4077 | 29.6705 | 0.769307 | 2.76306 |
| mut_A | wt_R | OK | 14.8221 | 25.235 | 0.767679 | 2.80576 |
| mut_A | wt_R | OK | 233.308 | 397.075 | 0.767177 | 3.16143 |
| mut_A | wt_R | OK | 208.601 | 354.948 | 0.766864 | 3.13477 |
| mut_A | wt_R | OK | 37.2478 | 63.1667 | 0.762009 | 2.63842 |
| mut_A | wt_R | OK | 99.4027 | 168.282 | 0.759525 | 3.02945 |

| | | | | | | |
|---|---|---|---|---|---|---|
| mut_A | wt_R | OK | 188.894 | 319.294 | 0.757313 | 3.09945 |
| mut_A | wt_R | OK | 624.253 | 1054.24 | 0.756003 | 1.78991 |
| mut_A | wt_R | OK | 14.66 | 24.751 | 0.755598 | 3.01077 |
| mut_A | wt_R | OK | 308.745 | 521.23 | 0.755506 | 3.11919 |
| mut_A | wt_R | OK | 65.029 | 109.682 | 0.754173 | 2.92663 |
| mut_A | wt_R | OK | 212.299 | 358.014 | 0.753919 | 3.08107 |
| mut_A | wt_R | OK | 305.24 | 514.435 | 0.753043 | 3.10655 |
| mut_A | wt_R | OK | 183.523 | 309.171 | 0.752446 | 3.09509 |
| mut_A | wt_R | OK | 6237.36 | 10485.7 | 0.749421 | 3.09422 |
| mut_A | wt_R | OK | 405.284 | 681.086 | 0.748904 | 2.18836 |
| mut_A | wt_R | OK | 430.734 | 723.846 | 0.748887 | 3.08442 |
| mut_A | wt_R | OK | 236.024 | 396.346 | 0.747823 | 3.08295 |
| mut_A | wt_R | OK | 188.777 | 316.71 | 0.746479 | 3.05242 |
| mut_A | wt_R | OK | 226.849 | 380.464 | 0.746027 | 3.07294 |
| mut_A | wt_R | OK | 3654.93 | 6128.65 | 0.745726 | 3.06131 |
| mut_A | wt_R | OK | 178.166 | 298.737 | 0.745655 | 3.0626 |
| mut_A | wt_R | OK | 8.61158 | 14.4379 | 0.745514 | 1.7953 |
| mut_A | wt_R | OK | 7.06896 | 11.8508 | 0.745417 | 2.64919 |
| mut_A | wt_R | OK | 1198.69 | 2006.33 | 0.743098 | 3.06508 |
| mut_A | wt_R | OK | 25.7221 | 42.9384 | 0.739259 | 2.80397 |
| mut_A | wt_R | OK | 185.775 | 310.106 | 0.739207 | 3.04539 |
| mut_A | wt_R | OK | 236.284 | 393.713 | 0.73662 | 2.95408 |
| mut_A | wt_R | OK | 88.9779 | 147.932 | 0.733414 | 3.01972 |
| mut_A | wt_R | OK | 988.274 | 1642.95 | 0.733303 | 3.02683 |
| mut_A | wt_R | OK | 95.1566 | 158.185 | 0.733236 | 2.91419 |
| mut_A | wt_R | OK | 24.8408 | 41.2529 | 0.731783 | 1.34508 |
| mut_A | wt_R | OK | 467.758 | 775.66 | 0.72966 | 3.00214 |
| mut_A | wt_R | OK | 1239.42 | 2054.38 | 0.729038 | 2.95896 |
| mut_A | wt_R | OK | 13.3134 | 22.0343 | 0.726868 | 2.68562 |
| mut_A | wt_R | OK | 415.661 | 687.885 | 0.72676 | 2.98362 |
| mut_A | wt_R | OK | 252.975 | 418.593 | 0.726556 | 1.42715 |
| mut_A | wt_R | OK | 53.531 | 88.5028 | 0.725349 | 2.87985 |
| mut_A | wt_R | OK | 1658.93 | 2742.37 | 0.725172 | 3.00236 |
| mut_A | wt_R | OK | 159.081 | 262.335 | 0.721649 | 2.92659 |
| mut_A | wt_R | OK | 140.772 | 231.155 | 0.715501 | 2.92565 |
| mut_A | wt_R | OK | 780.108 | 1279.82 | 0.714192 | 2.91303 |
| mut_A | wt_R | OK | 1108.71 | 1818.69 | 0.714015 | 2.94107 |
| mut_A | wt_R | OK | 1.31101 | 2.15046 | 0.713971 | 1.51837 |
| mut_A | wt_R | OK | 271.602 | 445.298 | 0.713277 | 2.94068 |
| mut_A | wt_R | OK | 170.946 | 280.122 | 0.71252 | 2.31984 |
| mut_A | wt_R | OK | 2004.72 | 3281.65 | 0.711022 | 2.93523 |
| mut_A | wt_R | OK | 15.6939 | 25.6409 | 0.708247 | 1.61439 |
| mut_A | wt_R | OK | 472.055 | 770.681 | 0.707181 | 2.73776 |
| mut_A | wt_R | OK | 336.388 | 548.119 | 0.704365 | 2.905 |
| mut_A | wt_R | OK | 384.751 | 626.2 | 0.702698 | 2.05657 |
| mut_A | wt_R | OK | 580.75 | 944.416 | 0.701506 | 2.90195 |
| mut_A | wt_R | OK | 11.0552 | 17.9771 | 0.701429 | 2.20727 |
| mut_A | wt_R | OK | 284.214 | 462.072 | 0.701139 | 2.89344 |
| mut_A | wt_R | OK | 292.215 | 474.963 | 0.700787 | 2.0706 |
| mut_A | wt_R | OK | 102.36 | 166.02 | 0.697703 | 2.80801 |
| mut_A | wt_R | OK | 508.114 | 824.042 | 0.697565 | 2.88703 |

| | | | | | | |
|---|---|---|---|---|---|---|
| mut_A | wt_R | OK | 1755.89 | 2843.29 | 0.695358 | 1.90183 |
| mut_A | wt_R | OK | 394.489 | 638.461 | 0.694615 | 2.81267 |
| mut_A | wt_R | OK | 224.997 | 364.118 | 0.694498 | 2.86433 |
| mut_A | wt_R | OK | 400.072 | 646.789 | 0.693034 | 2.8655 |
| mut_A | wt_R | OK | 3.96872 | 6.41346 | 0.692428 | 1.78508 |
| mut_A | wt_R | OK | 464.968 | 749.637 | 0.68906 | 2.81513 |
| mut_A | wt_R | OK | 115.404 | 185.991 | 0.68853 | 2.82849 |
| mut_A | wt_R | OK | 460.14 | 741.551 | 0.688472 | 2.81387 |
| mut_A | wt_R | OK | 5.49939 | 8.85272 | 0.686849 | 1.74439 |
| mut_A | wt_R | OK | 446.517 | 717.425 | 0.684114 | 2.82812 |
| mut_A | wt_R | OK | 389.163 | 623.784 | 0.680673 | 2.81422 |
| mut_A | wt_R | OK | 55.4272 | 88.7287 | 0.678807 | 2.48078 |
| mut_A | wt_R | OK | 258.55 | 413.415 | 0.677144 | 2.78205 |
| mut_A | wt_R | OK | 949.374 | 1516.03 | 0.675248 | 2.78865 |
| mut_A | wt_R | OK | 35.2509 | 56.2418 | 0.673984 | 2.31669 |
| mut_A | wt_R | OK | 686.7 | 1095.22 | 0.673465 | 2.78856 |
| mut_A | wt_R | OK | 557.74 | 886.436 | 0.668424 | 2.76352 |
| mut_A | wt_R | OK | 32.7309 | 52.0014 | 0.667899 | 2.47177 |
| mut_A | wt_R | OK | 60.6698 | 96.3214 | 0.666878 | 2.69182 |
| mut_A | wt_R | OK | 10.8081 | 17.1589 | 0.666846 | 2.06623 |
| mut_A | wt_R | OK | 34.966 | 55.4829 | 0.66609 | 2.42998 |
| mut_A | wt_R | OK | 100.945 | 160.147 | 0.66583 | 2.61395 |
| mut_A | wt_R | OK | 21172.5 | 33583.2 | 0.665547 | 2.72118 |
| mut_A | wt_R | OK | 124.259 | 197.04 | 0.665133 | 2.68637 |
| mut_A | wt_R | OK | 245.064 | 388.336 | 0.664146 | 2.46479 |
| mut_A | wt_R | OK | 55.8264 | 88.3769 | 0.662722 | 2.5055 |
| mut_A | wt_R | OK | 1652.82 | 2616.32 | 0.662612 | 2.72115 |
| mut_A | wt_R | OK | 369.414 | 584.56 | 0.662111 | 2.71857 |
| mut_A | wt_R | OK | 780.183 | 1233.42 | 0.660784 | 2.73306 |
| mut_A | wt_R | OK | 7.85081 | 12.3994 | 0.659359 | 2.11902 |
| mut_A | wt_R | OK | 138.857 | 218.823 | 0.656159 | 2.7003 |
| mut_A | wt_R | OK | 101.342 | 159.613 | 0.655351 | 2.48897 |
| mut_A | wt_R | OK | 132.661 | 207.876 | 0.647983 | 2.64989 |
| mut_A | wt_R | OK | 149.002 | 233.354 | 0.647189 | 2.66662 |
| mut_A | wt_R | OK | 1585.8 | 2482.35 | 0.646502 | 2.67229 |
| mut_A | wt_R | OK | 2139.75 | 3345.7 | 0.644867 | 1.98239 |
| mut_A | wt_R | OK | 4.95394 | 7.74456 | 0.644607 | 1.98263 |
| mut_A | wt_R | OK | 397.396 | 620.897 | 0.643778 | 1.43646 |
| mut_A | wt_R | OK | 353.819 | 552.673 | 0.643415 | 2.01194 |
| mut_A | wt_R | OK | 171.052 | 267.023 | 0.642532 | 2.64679 |
| mut_A | wt_R | OK | 1585.48 | 2473.06 | 0.641373 | 2.60557 |
| mut_A | wt_R | OK | 898.622 | 1400.5 | 0.640152 | 2.59117 |
| mut_A | wt_R | OK | 1422.38 | 2215.56 | 0.639364 | 2.61595 |
| mut_A | wt_R | OK | 7610.1 | 11832.1 | 0.636719 | 1.34344 |
| mut_A | wt_R | OK | 6.95517 | 10.8025 | 0.635204 | 1.82873 |
| mut_A | wt_R | OK | 798.248 | 1239.25 | 0.634561 | 2.6172 |
| mut_A | wt_R | OK | 53.9464 | 83.611 | 0.632166 | 1.97051 |
| mut_A | wt_R | OK | 610.651 | 946.409 | 0.632115 | 1.66245 |
| mut_A | wt_R | OK | 412.185 | 638.715 | 0.631881 | 2.60652 |
| mut_A | wt_R | OK | 3.49547 | 5.41549 | 0.631605 | 1.53068 |
| mut_A | wt_R | OK | 85.2317 | 132.048 | 0.631597 | 2.59885 |

| | | | | | | |
|---|---|---|---|---|---|---|
| mut_A | wt_R | OK | 348.209 | 539.208 | 0.63089 | 2.60212 |
| mut_A | wt_R | OK | 291.263 | 450.989 | 0.630767 | 2.53271 |
| mut_A | wt_R | OK | 3.95222 | 6.11878 | 0.630579 | 1.34496 |
| mut_A | wt_R | OK | 196.136 | 303.634 | 0.630477 | 2.59309 |
| mut_A | wt_R | OK | 115.295 | 178.452 | 0.630208 | 2.57932 |
| mut_A | wt_R | OK | 440.186 | 681.024 | 0.629594 | 2.56823 |
| mut_A | wt_R | OK | 41.7352 | 64.5038 | 0.628118 | 2.44688 |
| mut_A | wt_R | OK | 11.6798 | 18.0379 | 0.627009 | 1.62512 |
| mut_A | wt_R | OK | 70.7283 | 109.015 | 0.624166 | 2.39978 |
| mut_A | wt_R | OK | 3611.01 | 5560.26 | 0.622749 | 2.55436 |
| mut_A | wt_R | OK | 148.271 | 228.161 | 0.621821 | 1.32353 |
| mut_A | wt_R | OK | 47.2134 | 72.6405 | 0.621576 | 2.27409 |
| mut_A | wt_R | OK | 38.4807 | 59.1468 | 0.620166 | 2.48872 |
| mut_A | wt_R | OK | 14.6579 | 22.5274 | 0.620002 | 2.23785 |
| mut_A | wt_R | OK | 44.3897 | 68.1761 | 0.619042 | 2.16885 |
| mut_A | wt_R | OK | 37.0546 | 56.9101 | 0.619033 | 2.45713 |
| mut_A | wt_R | OK | 2317.87 | 3558.17 | 0.61834 | 2.55227 |
| mut_A | wt_R | OK | 6.40719 | 9.82233 | 0.616374 | 1.42287 |
| mut_A | wt_R | OK | 63.7969 | 97.7 | 0.614871 | 2.17648 |
| mut_A | wt_R | OK | 183.96 | 281.647 | 0.614491 | 2.47313 |
| mut_A | wt_R | OK | 140.096 | 214.347 | 0.613531 | 2.52355 |
| mut_A | wt_R | OK | 145.786 | 222.732 | 0.611457 | 2.4742 |
| mut_A | wt_R | OK | 10.0952 | 15.4068 | 0.609895 | 2.16141 |
| mut_A | wt_R | OK | 122.175 | 186.3 | 0.608677 | 1.91011 |
| mut_A | wt_R | OK | 18.401 | 28.015 | 0.606416 | 2.02998 |
| mut_A | wt_R | OK | 403.709 | 613.468 | 0.603671 | 2.44013 |
| mut_A | wt_R | OK | 11.1596 | 16.9521 | 0.603177 | 2.06495 |
| mut_A | wt_R | OK | 23.1832 | 35.1524 | 0.600539 | 2.26978 |
| mut_A | wt_R | OK | 1132.53 | 1715.11 | 0.598758 | 2.3143 |
| mut_A | wt_R | OK | 360.034 | 545.071 | 0.59831 | 2.44864 |
| mut_A | wt_R | OK | 175.734 | 266.009 | 0.598085 | 2.45159 |
| mut_A | wt_R | OK | 40.2726 | 60.9207 | 0.597132 | 2.37224 |
| mut_A | wt_R | OK | 205.835 | 311.288 | 0.596759 | 2.4484 |
| mut_A | wt_R | OK | 45.5309 | 68.8129 | 0.595833 | 2.27452 |
| mut_A | wt_R | OK | 58.9118 | 88.9062 | 0.593727 | 2.40605 |
| mut_A | wt_R | OK | 1952.5 | 2938.2 | 0.589605 | 1.52512 |
| mut_A | wt_R | OK | 118.834 | 178.791 | 0.589331 | 2.41258 |
| mut_A | wt_R | OK | 1154.35 | 1734.05 | 0.587071 | 2.42131 |
| mut_A | wt_R | OK | 61.9465 | 93.0057 | 0.586297 | 2.1052 |
| mut_A | wt_R | OK | 11.8964 | 17.8378 | 0.58442 | 2.15394 |
| mut_A | wt_R | OK | 93.1811 | 139.635 | 0.583549 | 2.3891 |
| mut_A | wt_R | OK | 28.9455 | 43.372 | 0.583423 | 1.83537 |
| mut_A | wt_R | OK | 381.283 | 571.291 | 0.583364 | 2.41258 |
| mut_A | wt_R | OK | 1282.55 | 1919.54 | 0.581748 | 2.34517 |
| mut_A | wt_R | OK | 5119.66 | 7647.6 | 0.578959 | 2.37465 |
| mut_A | wt_R | OK | 70.9169 | 105.859 | 0.577949 | 2.35268 |
| mut_A | wt_R | OK | 89.504 | 133.383 | 0.57555 | 2.336 |
| mut_A | wt_R | OK | 45.3643 | 67.5683 | 0.574789 | 2.15247 |
| mut_A | wt_R | OK | 360.569 | 536.907 | 0.574399 | 2.36371 |
| mut_A | wt_R | OK | 436.329 | 649.492 | 0.573893 | 2.37538 |
| mut_A | wt_R | OK | 218.469 | 324.887 | 0.572507 | 1.94528 |

| | | | | | | |
|---|---|---|---|---|---|---|
| mut_A | wt_R | OK | 144.742 | 215.12 | 0.571661 | 2.35352 |
| mut_A | wt_R | OK | 78.948 | 117.271 | 0.570868 | 2.21787 |
| mut_A | wt_R | OK | 1063.38 | 1579.21 | 0.570551 | 1.93352 |
| mut_A | wt_R | OK | 63.3891 | 93.989 | 0.568257 | 2.19694 |
| mut_A | wt_R | OK | 415.449 | 615.807 | 0.567809 | 2.34515 |
| mut_A | wt_R | OK | 30.66 | 45.44 | 0.567603 | 2.16006 |
| mut_A | wt_R | OK | 80.9039 | 119.831 | 0.566714 | 2.31499 |
| mut_A | wt_R | OK | 200.823 | 297.402 | 0.566491 | 1.45741 |
| mut_A | wt_R | OK | 131.084 | 193.936 | 0.565088 | 1.74365 |
| mut_A | wt_R | OK | 235.965 | 349.094 | 0.56504 | 2.32368 |
| mut_A | wt_R | OK | 249.789 | 369.278 | 0.563999 | 1.60901 |
| mut_A | wt_R | OK | 243.818 | 359.423 | 0.559878 | 2.31022 |
| mut_A | wt_R | OK | 76.9871 | 113.412 | 0.558878 | 2.28246 |
| mut_A | wt_R | OK | 104.798 | 154.361 | 0.558699 | 1.72779 |
| mut_A | wt_R | OK | 154.611 | 227.683 | 0.558388 | 1.90126 |
| mut_A | wt_R | OK | 377.117 | 555.263 | 0.558158 | 2.29517 |
| mut_A | wt_R | OK | 36.172 | 53.2392 | 0.557615 | 1.45257 |
| mut_A | wt_R | OK | 201.013 | 295.546 | 0.556093 | 2.28877 |
| mut_A | wt_R | OK | 55.4916 | 81.552 | 0.555453 | 2.24987 |
| mut_A | wt_R | OK | 46.9662 | 68.9919 | 0.554803 | 2.22993 |
| mut_A | wt_R | OK | 11.0556 | 16.2006 | 0.551266 | 1.59363 |
| mut_A | wt_R | OK | 128.732 | 188.63 | 0.551191 | 2.07607 |
| mut_A | wt_R | OK | 52.259 | 76.5211 | 0.550179 | 1.38621 |
| mut_A | wt_R | OK | 470.373 | 688.278 | 0.549188 | 2.27166 |
| mut_A | wt_R | OK | 115.35 | 168.501 | 0.546735 | 2.25018 |
| mut_A | wt_R | OK | 439.134 | 641.196 | 0.546105 | 2.24961 |
| mut_A | wt_R | OK | 114.577 | 167.26 | 0.545772 | 2.22369 |
| mut_A | wt_R | OK | 13.3308 | 19.4377 | 0.544096 | 2.01737 |
| mut_A | wt_R | OK | 155.436 | 226.578 | 0.543694 | 2.12204 |
| mut_A | wt_R | OK | 1428.94 | 2081.06 | 0.542373 | 2.23172 |
| mut_A | wt_R | OK | 1092.07 | 1588.38 | 0.540483 | 2.00738 |
| mut_A | wt_R | OK | 359.494 | 522.703 | 0.540024 | 2.22492 |
| mut_A | wt_R | OK | 44.9333 | 65.2782 | 0.538815 | 1.99635 |
| mut_A | wt_R | OK | 695.405 | 1009.14 | 0.537204 | 2.22419 |
| mut_A | wt_R | OK | 10.8561 | 15.7507 | 0.536907 | 1.98033 |
| mut_A | wt_R | OK | 563.808 | 817.676 | 0.536326 | 2.21919 |
| mut_A | wt_R | OK | 128.698 | 186.409 | 0.534485 | 2.14845 |
| mut_A | wt_R | OK | 460.719 | 666.161 | 0.531985 | 2.16954 |
| mut_A | wt_R | OK | 11.6027 | 16.7678 | 0.531236 | 1.89226 |
| mut_A | wt_R | OK | 623.864 | 901.078 | 0.530421 | 2.18417 |
| mut_A | wt_R | OK | 1731.91 | 2499.39 | 0.529208 | 1.26723 |
| mut_A | wt_R | OK | 207.859 | 299.773 | 0.528265 | 1.63555 |
| mut_A | wt_R | OK | 267.746 | 383.889 | 0.519824 | 2.13023 |
| mut_A | wt_R | OK | 195.556 | 280.363 | 0.519713 | 2.13508 |
| mut_A | wt_R | OK | 279.863 | 401.065 | 0.519116 | 2.12415 |
| mut_A | wt_R | OK | 239.503 | 342.816 | 0.517388 | 1.48088 |
| mut_A | wt_R | OK | 610.247 | 873.411 | 0.517269 | 2.13486 |
| mut_A | wt_R | OK | 148.519 | 212.538 | 0.517069 | 1.93415 |
| mut_A | wt_R | OK | 219.634 | 313.915 | 0.51527 | 2.1093 |
| mut_A | wt_R | OK | 66.333 | 94.7516 | 0.514424 | 1.80004 |
| mut_A | wt_R | OK | 365.124 | 521.459 | 0.514169 | 2.10914 |

| | | | | | | |
|---|---|---|---|---|---|---|
| mut_A | wt_R | OK | 7.7327 | 11.0394 | 0.513622 | 1.44548 |
| mut_A | wt_R | OK | 238.324 | 340.226 | 0.513569 | 2.10224 |
| mut_A | wt_R | OK | 268.487 | 382.818 | 0.511806 | 1.81565 |
| mut_A | wt_R | OK | 428.981 | 610.831 | 0.509859 | 2.10143 |
| mut_A | wt_R | OK | 73.7193 | 104.942 | 0.509482 | 2.04389 |
| mut_A | wt_R | OK | 117.681 | 167.495 | 0.509229 | 2.08705 |
| mut_A | wt_R | OK | 639.528 | 910.043 | 0.508926 | 2.1072 |
| mut_A | wt_R | OK | 185.175 | 263.208 | 0.50731 | 2.05103 |
| mut_A | wt_R | OK | 263.087 | 373.515 | 0.505629 | 2.06797 |
| mut_A | wt_R | OK | 1246.09 | 1767.89 | 0.504624 | 2.08815 |
| mut_A | wt_R | OK | 300.989 | 426.591 | 0.503143 | 2.06855 |
| mut_A | wt_R | OK | 7.51455 | 10.6462 | 0.502585 | 1.40941 |
| mut_A | wt_R | OK | 745.396 | 1055.39 | 0.501699 | 2.06349 |
| mut_A | wt_R | OK | 57.5928 | 81.4775 | 0.500514 | 1.91259 |
| mut_A | wt_R | OK | 57.3863 | 81.0637 | 0.49835 | 1.34992 |
| mut_A | wt_R | OK | 707.472 | 999.113 | 0.497976 | 2.05162 |
| mut_A | wt_R | OK | 30.57 | 43.0796 | 0.494887 | 2.03085 |
| mut_A | wt_R | OK | 235.891 | 332.391 | 0.494762 | 1.98446 |
| mut_A | wt_R | OK | 66.631 | 93.7901 | 0.493241 | 1.99176 |
| mut_A | wt_R | OK | 671.56 | 944.683 | 0.492314 | 1.97031 |
| mut_A | wt_R | OK | 763.34 | 1073.33 | 0.4917 | 2.01841 |
| mut_A | wt_R | OK | 22.4265 | 31.5134 | 0.490763 | 1.59155 |
| mut_A | wt_R | OK | 31.5466 | 44.3183 | 0.490415 | 1.97573 |
| mut_A | wt_R | OK | 90.0265 | 126.373 | 0.489264 | 1.98974 |
| mut_A | wt_R | OK | 200.317 | 281.189 | 0.489252 | 2.00941 |
| mut_A | wt_R | OK | 6255.19 | 8765.62 | 0.486802 | 2.01768 |
| mut_A | wt_R | OK | 506.017 | 709.09 | 0.486783 | 2.01715 |
| mut_A | wt_R | OK | 1612.37 | 2258.68 | 0.486303 | 2.00834 |
| mut_A | wt_R | OK | 93.8195 | 131.359 | 0.485555 | 2.00038 |
| mut_A | wt_R | OK | 763.212 | 1068.49 | 0.48542 | 1.95627 |
| mut_A | wt_R | OK | 277.337 | 388.177 | 0.485076 | 1.96749 |
| mut_A | wt_R | OK | 1565.63 | 2189.85 | 0.48409 | 1.99945 |
| mut_A | wt_R | OK | 14.3614 | 20.0774 | 0.48338 | 1.57443 |
| mut_A | wt_R | OK | 897.594 | 1253.42 | 0.481737 | 1.9902 |
| mut_A | wt_R | OK | 813.78 | 1136.24 | 0.481553 | 1.99078 |
| mut_A | wt_R | OK | 154.561 | 215.735 | 0.481085 | 1.97134 |
| mut_A | wt_R | OK | 537.734 | 750.412 | 0.480791 | 1.98836 |
| mut_A | wt_R | OK | 148.988 | 207.794 | 0.479957 | 1.29181 |
| mut_A | wt_R | OK | 92.2427 | 128.622 | 0.479628 | 1.28024 |
| mut_A | wt_R | OK | 12.9943 | 18.1079 | 0.478744 | 1.71798 |
| mut_A | wt_R | OK | 79.2689 | 110.414 | 0.478102 | 1.95166 |
| mut_A | wt_R | OK | 228.244 | 317.676 | 0.47698 | 1.96593 |
| mut_A | wt_R | OK | 74.0487 | 103.055 | 0.476866 | 1.35807 |
| mut_A | wt_R | OK | 51.0019 | 70.836 | 0.473932 | 1.85453 |
| mut_A | wt_R | OK | 488.548 | 677.679 | 0.472103 | 1.93735 |
| mut_A | wt_R | OK | 62.3701 | 86.4718 | 0.471375 | 1.88263 |
| mut_A | wt_R | OK | 68.9859 | 95.584 | 0.470468 | 1.85328 |
| mut_A | wt_R | OK | 53.1221 | 73.5696 | 0.469798 | 1.78266 |
| mut_A | wt_R | OK | 287.093 | 397.549 | 0.469612 | 1.93615 |
| mut_A | wt_R | OK | 1037.71 | 1436.85 | 0.469507 | 1.93323 |
| mut_A | wt_R | OK | 506.091 | 700.733 | 0.469468 | 1.94486 |

| | | | | | | |
|---|---|---|---|---|---|---|
| mut_A | wt_R | OK | 180.535 | 249.707 | 0.467959 | 1.89443 |
| mut_A | wt_R | OK | 207.891 | 287.462 | 0.467543 | 1.90778 |
| mut_A | wt_R | OK | 117.259 | 161.926 | 0.46563 | 1.907 |
| mut_A | wt_R | OK | 63.1048 | 87.0949 | 0.464839 | 1.54931 |
| mut_A | wt_R | OK | 365.758 | 504.313 | 0.463431 | 1.59433 |
| mut_A | wt_R | OK | 20.8367 | 28.7175 | 0.462804 | 1.67305 |
| mut_A | wt_R | OK | 3756.49 | 5169.1 | 0.460527 | 1.89938 |
| mut_A | wt_R | OK | 290.407 | 399.575 | 0.460388 | 1.90037 |
| mut_A | wt_R | OK | 13.3297 | 18.34 | 0.460343 | 1.36509 |
| mut_A | wt_R | OK | 144.911 | 199.147 | 0.458669 | 1.67981 |
| mut_A | wt_R | OK | 200.249 | 275.162 | 0.458491 | 1.8888 |
| mut_A | wt_R | OK | 11.0466 | 15.1651 | 0.457142 | 1.662 |
| mut_A | wt_R | OK | 592.926 | 813.88 | 0.456962 | 1.88831 |
| mut_A | wt_R | OK | 484.452 | 664.954 | 0.4569 | 1.88183 |
| mut_A | wt_R | OK | 1634.7 | 2243.03 | 0.456421 | 1.88377 |
| mut_A | wt_R | OK | 53.862 | 73.8732 | 0.455784 | 1.80273 |
| mut_A | wt_R | OK | 11.6113 | 15.8965 | 0.45318 | 1.65804 |
| mut_A | wt_R | OK | 324.916 | 444.319 | 0.451532 | 1.85858 |
| mut_A | wt_R | OK | 31.3026 | 42.7988 | 0.451287 | 1.73862 |
| mut_A | wt_R | OK | 257.046 | 351.237 | 0.450418 | 1.85414 |
| mut_A | wt_R | OK | 90.5125 | 123.637 | 0.449919 | 1.8299 |
| mut_A | wt_R | OK | 44.8171 | 61.0936 | 0.44697 | 1.4134 |
| mut_A | wt_R | OK | 691.041 | 941.785 | 0.446627 | 1.83755 |
| mut_A | wt_R | OK | 184.268 | 251.021 | 0.446004 | 1.41555 |
| mut_A | wt_R | OK | 56.6393 | 77.1546 | 0.445948 | 1.62135 |
| mut_A | wt_R | OK | 326.168 | 443.786 | 0.444248 | 1.78659 |
| mut_A | wt_R | OK | 492.599 | 669.5 | 0.442672 | 1.80988 |
| mut_A | wt_R | OK | 483.475 | 655.857 | 0.439939 | 1.81537 |
| mut_A | wt_R | OK | 470.154 | 637.213 | 0.438642 | 1.80714 |
| mut_A | wt_R | OK | 427.383 | 579.055 | 0.438169 | 1.79153 |
| mut_A | wt_R | OK | 2572.3 | 3482.97 | 0.437262 | 1.80228 |
| mut_A | wt_R | OK | 65.5783 | 88.7761 | 0.436954 | 1.75937 |
| mut_A | wt_R | OK | 41.7913 | 56.5617 | 0.436623 | 1.75515 |
| mut_A | wt_R | OK | 158.058 | 213.825 | 0.43598 | 1.5923 |
| mut_A | wt_R | OK | 46.972 | 63.5155 | 0.435308 | 1.50464 |
| mut_A | wt_R | OK | 903.434 | 1221.35 | 0.434981 | 1.77611 |
| mut_A | wt_R | OK | 119.695 | 161.678 | 0.43376 | 1.78681 |
| mut_A | wt_R | OK | 235.626 | 317.759 | 0.431435 | 1.30336 |
| mut_A | wt_R | OK | 284.9 | 383.834 | 0.430027 | 1.76806 |
| mut_A | wt_R | OK | 99.7483 | 134.352 | 0.42965 | 1.4259 |
| mut_A | wt_R | OK | 301.966 | 406.706 | 0.429598 | 1.75695 |
| mut_A | wt_R | OK | 271.022 | 364.838 | 0.428845 | 1.71786 |
| mut_A | wt_R | OK | 5727.73 | 7680.41 | 0.42322 | 1.74409 |
| mut_A | wt_R | OK | 145.491 | 194.901 | 0.421809 | 1.72761 |
| mut_A | wt_R | OK | 211.512 | 283.313 | 0.421655 | 1.71381 |
| mut_A | wt_R | OK | 313.997 | 420.293 | 0.420644 | 1.72011 |
| mut_A | wt_R | OK | 345.789 | 462.762 | 0.420378 | 1.71154 |
| mut_A | wt_R | OK | 192.411 | 257.49 | 0.420326 | 1.72741 |
| mut_A | wt_R | OK | 340.588 | 455.786 | 0.420326 | 1.72172 |
| mut_A | wt_R | OK | 296.502 | 396.721 | 0.420085 | 1.72301 |
| mut_A | wt_R | OK | 317.142 | 424.245 | 0.419767 | 1.72644 |

| | | | | | | |
|---|---|---|---|---|---|---|
| mut_A | wt_R | OK | 738.705 | 988.019 | 0.419541 | 1.71939 |
| mut_A | wt_R | OK | 403.029 | 538.467 | 0.417974 | 1.70087 |
| mut_A | wt_R | OK | 6286.87 | 8390.23 | 0.416368 | 1.7091 |
| mut_A | wt_R | OK | 449.141 | 599.303 | 0.416116 | 1.70975 |
| mut_A | wt_R | OK | 185.37 | 247.31 | 0.415916 | 1.70261 |
| mut_A | wt_R | OK | 117.901 | 157.209 | 0.415102 | 1.69162 |
| mut_A | wt_R | OK | 19.1363 | 25.5018 | 0.414286 | 1.30171 |
| mut_A | wt_R | OK | 2009.98 | 2674.15 | 0.411897 | 1.69086 |
| mut_A | wt_R | OK | 355.911 | 473.461 | 0.411729 | 1.69643 |
| mut_A | wt_R | OK | 232.476 | 308.743 | 0.409324 | 1.68506 |
| mut_A | wt_R | OK | 215.352 | 285.546 | 0.407024 | 1.65674 |
| mut_A | wt_R | OK | 11.8271 | 15.6712 | 0.406014 | 1.36123 |
| mut_A | wt_R | OK | 29.0142 | 38.4324 | 0.405566 | 1.59019 |
| mut_A | wt_R | OK | 23.6255 | 31.2903 | 0.405373 | 1.44123 |
| mut_A | wt_R | OK | 70.2471 | 92.8894 | 0.403076 | 1.62562 |
| mut_A | wt_R | OK | 12.957 | 17.1302 | 0.402808 | 1.45157 |
| mut_A | wt_R | OK | 44.4296 | 58.6888 | 0.401565 | 1.59237 |
| mut_A | wt_R | OK | 144.111 | 190.238 | 0.400627 | 1.64139 |
| mut_A | wt_R | OK | 2825.24 | 3721.62 | 0.397555 | 1.63527 |
| mut_A | wt_R | OK | 510.624 | 672.441 | 0.397145 | 1.62501 |
| mut_A | wt_R | OK | 373.587 | 491.388 | 0.395421 | 1.63737 |
| mut_A | wt_R | OK | 1011.64 | 1329.86 | 0.394575 | 1.62953 |
| mut_A | wt_R | OK | 333.947 | 438.915 | 0.394325 | 1.61753 |
| mut_A | wt_R | OK | 487.192 | 639.591 | 0.39266 | 1.61214 |
| mut_A | wt_R | OK | 207.325 | 272.092 | 0.392205 | 1.52277 |
| mut_A | wt_R | OK | 21.4352 | 28.1047 | 0.390825 | 1.33607 |
| mut_A | wt_R | OK | 64.8073 | 84.9221 | 0.389984 | 1.59764 |
| mut_A | wt_R | OK | 852.871 | 1116.37 | 0.388418 | 1.5991 |
| mut_A | wt_R | OK | 1065.88 | 1394.98 | 0.388197 | 1.28583 |
| mut_A | wt_R | OK | 207.793 | 271.388 | 0.385208 | 1.28401 |
| mut_A | wt_R | OK | 227.316 | 296.827 | 0.384922 | 1.59273 |
| mut_A | wt_R | OK | 616.48 | 803.868 | 0.382904 | 1.56095 |
| mut_A | wt_R | OK | 267.243 | 348.168 | 0.381632 | 1.57266 |
| mut_A | wt_R | OK | 456.028 | 593.935 | 0.381181 | 1.57533 |
| mut_A | wt_R | OK | 18.0779 | 23.5424 | 0.381035 | 1.43142 |
| mut_A | wt_R | OK | 2683.31 | 3493.18 | 0.380531 | 1.56978 |
| mut_A | wt_R | OK | 92.6012 | 120.399 | 0.378714 | 1.55184 |
| mut_A | wt_R | OK | 650.155 | 844.661 | 0.377589 | 1.53447 |
| mut_A | wt_R | OK | 1590.55 | 2065.35 | 0.376854 | 1.4559 |
| mut_A | wt_R | OK | 34.2614 | 44.4777 | 0.376499 | 1.44339 |
| mut_A | wt_R | OK | 191.27 | 247.996 | 0.374709 | 1.5359 |
| mut_A | wt_R | OK | 2419.88 | 3136.95 | 0.374427 | 1.55002 |
| mut_A | wt_R | OK | 467.559 | 606.012 | 0.374199 | 1.53429 |
| mut_A | wt_R | OK | 15.5905 | 20.2062 | 0.37413 | 1.34545 |
| mut_A | wt_R | OK | 53.5157 | 69.3202 | 0.373313 | 1.31802 |
| mut_A | wt_R | OK | 151.025 | 195.574 | 0.372926 | 1.52651 |
| mut_A | wt_R | OK | 513.572 | 664.925 | 0.372625 | 1.50356 |
| mut_A | wt_R | OK | 174.677 | 225.861 | 0.37075 | 1.36378 |
| mut_A | wt_R | OK | 231.072 | 298.458 | 0.369186 | 1.51631 |
| mut_A | wt_R | OK | 7.90036 | 10.198 | 0.368294 | 1.39328 |
| mut_A | wt_R | OK | 58.827 | 75.9282 | 0.368158 | 1.46811 |

| | | | | | | |
|---|---|---|---|---|---|---|
| mut_A | wt_R | OK | 10.1033 | 13.0363 | 0.367698 | 1.33328 |
| mut_A | wt_R | OK | 108.567 | 140.059 | 0.367445 | 1.50649 |
| mut_A | wt_R | OK | 22.3819 | 28.8126 | 0.364368 | 1.38628 |
| mut_A | wt_R | OK | 443.236 | 569.261 | 0.361016 | 1.49153 |
| mut_A | wt_R | OK | 21.6585 | 27.8003 | 0.360168 | 1.29093 |
| mut_A | wt_R | OK | 167.349 | 214.784 | 0.360023 | 1.29863 |
| mut_A | wt_R | OK | 265.69 | 340.51 | 0.357955 | 1.48137 |
| mut_A | wt_R | OK | 439.69 | 563.455 | 0.357815 | 1.47745 |
| mut_A | wt_R | OK | 47.4168 | 60.6735 | 0.355668 | 1.32414 |
| mut_A | wt_R | OK | 284.855 | 364.235 | 0.354641 | 1.45919 |
| mut_A | wt_R | OK | 373.385 | 477.088 | 0.353592 | 1.35711 |
| mut_A | wt_R | OK | 58.7058 | 74.9821 | 0.353043 | 1.35435 |
| mut_A | wt_R | OK | 172.278 | 219.965 | 0.35253 | 1.455 |
| mut_A | wt_R | OK | 439.257 | 559.947 | 0.350225 | 1.44098 |
| mut_A | wt_R | OK | 374.79 | 477.72 | 0.350081 | 1.43855 |
| mut_A | wt_R | OK | 929.831 | 1184.94 | 0.349777 | 1.44483 |
| mut_A | wt_R | OK | 1538.51 | 1960.58 | 0.349746 | 1.4324 |
| mut_A | wt_R | OK | 11914.5 | 15181 | 0.349547 | 1.43982 |
| mut_A | wt_R | OK | 1086.31 | 1383.65 | 0.34904 | 1.40295 |
| mut_A | wt_R | OK | 93.8043 | 118.827 | 0.34114 | 1.36598 |
| mut_A | wt_R | OK | 130.626 | 165.268 | 0.339364 | 1.34704 |
| mut_A | wt_R | OK | 614.9 | 776.442 | 0.336528 | 1.37354 |
| mut_A | wt_R | OK | 223.621 | 282.104 | 0.335169 | 1.37256 |
| mut_A | wt_R | OK | 60.2317 | 75.9643 | 0.334799 | 1.2637 |
| mut_A | wt_R | OK | 97.2011 | 122.483 | 0.333532 | 1.28861 |
| mut_A | wt_R | OK | 361.857 | 455.865 | 0.333186 | 1.22999 |
| mut_A | wt_R | OK | 343.565 | 432.652 | 0.332624 | 1.36827 |
| mut_A | wt_R | OK | 553.676 | 696.4 | 0.330874 | 1.30073 |
| mut_A | wt_R | OK | 206.682 | 259.224 | 0.326785 | 1.33856 |
| mut_A | wt_R | OK | 755.526 | 947.125 | 0.326074 | 1.35211 |
| mut_A | wt_R | OK | 243.858 | 305.086 | 0.323172 | 1.32279 |
| mut_A | wt_R | OK | 145.558 | 182.044 | 0.322693 | 1.26781 |
| mut_A | wt_R | OK | 148.366 | 185.55 | 0.322648 | 1.31764 |
| mut_A | wt_R | OK | 7622.38 | 9531.64 | 0.322483 | 1.3212 |
| mut_A | wt_R | OK | 830.306 | 1034.83 | 0.317678 | 1.31295 |
| mut_A | wt_R | OK | 220.818 | 275.15 | 0.31736 | 1.3091 |
| mut_A | wt_R | OK | 36.1584 | 44.9779 | 0.314883 | 1.28982 |
| mut_A | wt_R | OK | 292.691 | 363.723 | 0.313464 | 1.29192 |
| mut_A | wt_R | OK | 94.1177 | 116.816 | 0.311695 | 1.27879 |
| mut_A | wt_R | OK | 356.16 | 441.141 | 0.308718 | 1.27137 |
| mut_A | wt_R | OK | 574.937 | 710.558 | 0.305549 | 1.25429 |
| mut_A | wt_R | OK | 220.294 | 178.544 | -0.303152 | -1.25214 |
| mut_A | wt_R | OK | 138.815 | 112.242 | -0.30655 | -1.2513 |
| mut_A | wt_R | OK | 133.905 | 108.132 | -0.308426 | -1.24017 |
| mut_A | wt_R | OK | 448.143 | 361.616 | -0.309504 | -1.25654 |
| mut_A | wt_R | OK | 155.48 | 125.093 | -0.313731 | -1.28296 |
| mut_A | wt_R | OK | 349.357 | 280.081 | -0.318861 | -1.29116 |
| mut_A | wt_R | OK | 75.0427 | 59.9552 | -0.323827 | -1.28022 |
| mut_A | wt_R | OK | 1204 | 961.623 | -0.324286 | -1.34173 |
| mut_A | wt_R | OK | 363.343 | 290.196 | -0.324304 | -1.32718 |
| mut_A | wt_R | OK | 97.8497 | 78.1274 | -0.32474 | -1.33255 |

| | | | | | | |
|---|---|---|---|---|---|---|
| mut_A | wt_R | OK | 50.6654 | 40.4442 | -0.325067 | -1.3068 |
| mut_A | wt_R | OK | 1113.05 | 888.31 | -0.325378 | -1.3434 |
| mut_A | wt_R | OK | 14435 | 11508.8 | -0.326831 | -1.34439 |
| mut_A | wt_R | OK | 471.72 | 375.982 | -0.327268 | -1.34737 |
| mut_A | wt_R | OK | 52.5419 | 41.7408 | -0.332011 | -1.35382 |
| mut_A | wt_R | OK | 544.864 | 431.976 | -0.334945 | -1.36387 |
| mut_A | wt_R | OK | 80.283 | 63.6391 | -0.33518 | -1.24923 |
| mut_A | wt_R | OK | 323.3 | 255.572 | -0.339143 | -1.37378 |
| mut_A | wt_R | OK | 40.7164 | 32.1817 | -0.339368 | -1.31395 |
| mut_A | wt_R | OK | 1660.42 | 1309.85 | -0.342147 | -1.40478 |
| mut_A | wt_R | OK | 1267.84 | 998.673 | -0.344284 | -1.42601 |
| mut_A | wt_R | OK | 82.1663 | 64.6885 | -0.345038 | -1.36802 |
| mut_A | wt_R | OK | 48.6494 | 38.2956 | -0.345244 | -1.34244 |
| mut_A | wt_R | OK | 494.811 | 388.156 | -0.350242 | -1.43946 |
| mut_A | wt_R | OK | 2475.72 | 1940.94 | -0.351095 | -1.34425 |
| mut_A | wt_R | OK | 1513.96 | 1186.92 | -0.351102 | -1.45177 |
| mut_A | wt_R | OK | 132.183 | 103.577 | -0.351836 | -1.44409 |
| mut_A | wt_R | OK | 11812.8 | 9232.18 | -0.355607 | -1.45597 |
| mut_A | wt_R | OK | 1295.78 | 1010.61 | -0.358591 | -1.4668 |
| mut_A | wt_R | OK | 520.719 | 405.735 | -0.359966 | -1.48753 |
| mut_A | wt_R | OK | 61.9178 | 48.2251 | -0.360569 | -1.42937 |
| mut_A | wt_R | OK | 138.02 | 107.494 | -0.360618 | -1.46638 |
| mut_A | wt_R | OK | 2953.23 | 2292.59 | -0.365314 | -1.47917 |
| mut_A | wt_R | OK | 657.745 | 509.528 | -0.368368 | -1.52035 |
| mut_A | wt_R | OK | 142.225 | 109.998 | -0.370697 | -1.49835 |
| mut_A | wt_R | OK | 464.29 | 358.981 | -0.371121 | -1.52724 |
| mut_A | wt_R | OK | 608.15 | 469.049 | -0.37469 | -1.5288 |
| mut_A | wt_R | OK | 33.6011 | 25.87 | -0.377225 | -1.45423 |
| mut_A | wt_R | OK | 977.874 | 752.799 | -0.377383 | -1.38479 |
| mut_A | wt_R | OK | 466.195 | 358.291 | -0.379802 | -1.56742 |
| mut_A | wt_R | OK | 413.1 | 317.161 | -0.381274 | -1.38931 |
| mut_A | wt_R | OK | 1142.33 | 875.315 | -0.384099 | -1.57765 |
| mut_A | wt_R | OK | 567.773 | 434.831 | -0.38486 | -1.57748 |
| mut_A | wt_R | OK | 1740.66 | 1329.25 | -0.389023 | -1.60094 |
| mut_A | wt_R | OK | 272.832 | 208.298 | -0.389361 | -1.60248 |
| mut_A | wt_R | OK | 7.83939 | 5.98344 | -0.389765 | -1.35989 |
| mut_A | wt_R | OK | 81.5192 | 62.1912 | -0.390429 | -1.47978 |
| mut_A | wt_R | OK | 79.2138 | 60.2976 | -0.393651 | -1.51213 |
| mut_A | wt_R | OK | 65.1649 | 49.3832 | -0.400075 | -1.53773 |
| mut_A | wt_R | OK | 1044.99 | 791.723 | -0.400421 | -1.65212 |
| mut_A | wt_R | OK | 371.115 | 280.974 | -0.401433 | -1.6503 |
| mut_A | wt_R | OK | 739.295 | 559.431 | -0.402191 | -1.66411 |
| mut_A | wt_R | OK | 2887.03 | 2184.02 | -0.402602 | -1.66168 |
| mut_A | wt_R | OK | 488.402 | 369.31 | -0.403236 | -1.66123 |
| mut_A | wt_R | OK | 30.0759 | 22.6932 | -0.406344 | -1.5132 |
| mut_A | wt_R | OK | 909.003 | 684.474 | -0.409289 | -1.69046 |
| mut_A | wt_R | OK | 563.178 | 423.958 | -0.40967 | -1.69315 |
| mut_A | wt_R | OK | 89.6576 | 67.4115 | -0.411431 | -1.59395 |
| mut_A | wt_R | OK | 430.956 | 321.858 | -0.421117 | -1.73658 |
| mut_A | wt_R | OK | 78.0026 | 58.1478 | -0.423797 | -1.60591 |
| mut_A | wt_R | OK | 1688.96 | 1258.63 | -0.424279 | -1.75268 |

| | | | | | | |
|---|---|---|---|---|---|---|
| mut_A | wt_R | OK | 93.0829 | 69.3203 | -0.425238 | -1.68659 |
| mut_A | wt_R | OK | 148.065 | 110.185 | -0.426303 | -1.58455 |
| mut_A | wt_R | OK | 158.459 | 117.476 | -0.431749 | -1.74509 |
| mut_A | wt_R | OK | 203.63 | 150.923 | -0.432138 | -1.76957 |
| mut_A | wt_R | OK | 746.23 | 552.471 | -0.433721 | -1.79256 |
| mut_A | wt_R | OK | 56.8579 | 42.015 | -0.436458 | -1.74516 |
| mut_A | wt_R | OK | 1273.58 | 940.874 | -0.436813 | -1.80021 |
| mut_A | wt_R | OK | 38.186 | 28.2099 | -0.436842 | -1.67488 |
| mut_A | wt_R | OK | 380.447 | 280.865 | -0.43782 | -1.7403 |
| mut_A | wt_R | OK | 155.431 | 114.723 | -0.438126 | -1.77251 |
| mut_A | wt_R | OK | 399.733 | 294.944 | -0.438596 | -1.80446 |
| mut_A | wt_R | OK | 349.147 | 256.89 | -0.442687 | -1.82494 |
| mut_A | wt_R | OK | 93.2127 | 68.5714 | -0.442919 | -1.80854 |
| mut_A | wt_R | OK | 1189.36 | 874.597 | -0.443492 | -1.83515 |
| mut_A | wt_R | OK | 161.956 | 119.087 | -0.443579 | -1.81328 |
| mut_A | wt_R | OK | 976.761 | 717.738 | -0.444548 | -1.83604 |
| mut_A | wt_R | OK | 3399.19 | 2497.27 | -0.444839 | -1.81548 |
| mut_A | wt_R | OK | 45.7836 | 33.6293 | -0.445113 | -1.79457 |
| mut_A | wt_R | OK | 5604.5 | 4111.65 | -0.446869 | -1.8371 |
| mut_A | wt_R | OK | 149.206 | 109.435 | -0.447236 | -1.82609 |
| mut_A | wt_R | OK | 680.257 | 498.07 | -0.44973 | -1.85072 |
| mut_A | wt_R | OK | 22.9862 | 16.8269 | -0.449996 | -1.47728 |
| mut_A | wt_R | OK | 7762.33 | 5675.99 | -0.451617 | -1.85037 |
| mut_A | wt_R | OK | 1020.88 | 746.188 | -0.452207 | -1.86408 |
| mut_A | wt_R | OK | 128.09 | 93.371 | -0.456107 | -1.83563 |
| mut_A | wt_R | OK | 13.8187 | 10.0671 | -0.456969 | -1.33185 |
| mut_A | wt_R | OK | 31.4759 | 22.9036 | -0.458675 | -1.66494 |
| mut_A | wt_R | OK | 425.492 | 309.139 | -0.460877 | -1.63246 |
| mut_A | wt_R | OK | 798.405 | 579.444 | -0.46245 | -1.88777 |
| mut_A | wt_R | OK | 164.505 | 119.317 | -0.463328 | -1.75886 |
| mut_A | wt_R | OK | 227.232 | 164.626 | -0.464979 | -1.90687 |
| mut_A | wt_R | OK | 152.706 | 110.557 | -0.465969 | -1.90333 |
| mut_A | wt_R | OK | 256.737 | 185.494 | -0.468917 | -1.91064 |
| mut_A | wt_R | OK | 198.788 | 143.452 | -0.47067 | -1.93297 |
| mut_A | wt_R | OK | 4346.75 | 3135.39 | -0.471291 | -1.92873 |
| mut_A | wt_R | OK | 304.402 | 219.499 | -0.471762 | -1.87856 |
| mut_A | wt_R | OK | 2143.56 | 1543.33 | -0.473964 | -1.93903 |
| mut_A | wt_R | OK | 168.787 | 121.449 | -0.474856 | -1.85031 |
| mut_A | wt_R | OK | 6.06517 | 4.36343 | -0.475085 | -1.43838 |
| mut_A | wt_R | OK | 113.481 | 81.5389 | -0.476894 | -1.88515 |
| mut_A | wt_R | OK | 96.1721 | 69.0968 | -0.476999 | -1.93054 |
| mut_A | wt_R | OK | 182.576 | 131.036 | -0.47854 | -1.79172 |
| mut_A | wt_R | OK | 50.297 | 36.0806 | -0.479248 | -1.56719 |
| mut_A | wt_R | OK | 759.963 | 544.096 | -0.482068 | -1.42219 |
| mut_A | wt_R | OK | 318.349 | 227.875 | -0.482367 | -1.89138 |
| mut_A | wt_R | OK | 175.896 | 125.811 | -0.483465 | -1.97036 |
| mut_A | wt_R | OK | 70.2103 | 50.1904 | -0.48427 | -1.92531 |
| mut_A | wt_R | OK | 412.93 | 294.952 | -0.485418 | -1.99701 |
| mut_A | wt_R | OK | 211.762 | 151.217 | -0.485821 | -1.9491 |
| mut_A | wt_R | OK | 15767 | 11241.9 | -0.488016 | -2.0039 |
| mut_A | wt_R | OK | 854.337 | 608.998 | -0.488368 | -2.02349 |

| | | | | | | |
|---|---|---|---|---|---|---|
| mut_A | wt_R | OK | 51.0256 | 36.3213 | -0.490408 | -1.8198 |
| mut_A | wt_R | OK | 242.123 | 172.198 | -0.491671 | -2.0126 |
| mut_A | wt_R | OK | 76.7144 | 54.5085 | -0.493016 | -2.00452 |
| mut_A | wt_R | OK | 118.552 | 84.1393 | -0.494667 | -1.27212 |
| mut_A | wt_R | OK | 38.0609 | 27.0113 | -0.494745 | -1.95887 |
| mut_A | wt_R | OK | 10296.3 | 7267.74 | -0.502552 | -1.99636 |
| mut_A | wt_R | OK | 1408.68 | 993.279 | -0.504077 | -2.08178 |
| mut_A | wt_R | OK | 35703.4 | 25166.2 | -0.504574 | -2.0462 |
| mut_A | wt_R | OK | 89.8112 | 63.2665 | -0.505455 | -2.01151 |
| mut_A | wt_R | OK | 755.39 | 531.944 | -0.505948 | -2.0901 |
| mut_A | wt_R | OK | 71.5204 | 50.3555 | -0.506206 | -2.0573 |
| mut_A | wt_R | OK | 1418.42 | 998.65 | -0.50623 | -2.05369 |
| mut_A | wt_R | OK | 7.19525 | 5.05499 | -0.509339 | -1.59938 |
| mut_A | wt_R | OK | 54.2094 | 38.0686 | -0.50994 | -2.02595 |
| mut_A | wt_R | OK | 418.046 | 293.557 | -0.510019 | -2.09041 |
| mut_A | wt_R | OK | 520.403 | 364.855 | -0.512305 | -1.81401 |
| mut_A | wt_R | OK | 193.768 | 135.747 | -0.513408 | -1.98116 |
| mut_A | wt_R | OK | 6.16447 | 4.31839 | -0.513482 | -1.55735 |
| mut_A | wt_R | OK | 27.8011 | 19.4723 | -0.513723 | -1.8831 |
| mut_A | wt_R | OK | 1776.23 | 1241.82 | -0.516364 | -2.08231 |
| mut_A | wt_R | OK | 325.41 | 227.12 | -0.518807 | -2.1113 |
| mut_A | wt_R | OK | 115.283 | 80.4485 | -0.519036 | -2.13735 |
| mut_A | wt_R | OK | 1439.66 | 1004.51 | -0.519239 | -2.1374 |
| mut_A | wt_R | OK | 96.6171 | 67.325 | -0.521135 | -2.11366 |
| mut_A | wt_R | OK | 347.775 | 242.088 | -0.522621 | -1.53012 |
| mut_A | wt_R | OK | 60.9166 | 42.3208 | -0.525469 | -2.11775 |
| mut_A | wt_R | OK | 67.6231 | 46.8711 | -0.528818 | -2.09497 |
| mut_A | wt_R | OK | 574.526 | 398.065 | -0.529369 | -2.17852 |
| mut_A | wt_R | OK | 113.992 | 78.9717 | -0.529526 | -2.12703 |
| mut_A | wt_R | OK | 305.425 | 211.081 | -0.53302 | -2.13816 |
| mut_A | wt_R | OK | 287.833 | 198.816 | -0.533801 | -2.16089 |
| mut_A | wt_R | OK | 1742.12 | 1201.87 | -0.535561 | -2.18718 |
| mut_A | wt_R | OK | 2109.87 | 1449.77 | -0.541325 | -2.24008 |
| mut_A | wt_R | OK | 410.259 | 280.959 | -0.546176 | -2.22058 |
| mut_A | wt_R | OK | 132.711 | 90.6666 | -0.549645 | -2.23628 |
| mut_A | wt_R | OK | 146.664 | 100.164 | -0.550145 | -1.84655 |
| mut_A | wt_R | OK | 116.381 | 79.3737 | -0.552126 | -2.05254 |
| mut_A | wt_R | OK | 172.977 | 117.401 | -0.559134 | -2.30646 |
| mut_A | wt_R | OK | 22.5765 | 15.3118 | -0.560174 | -2.0845 |
| mut_A | wt_R | OK | 1678.95 | 1137.17 | -0.562107 | -2.31264 |
| mut_A | wt_R | OK | 363.133 | 245.72 | -0.563485 | -2.31404 |
| mut_A | wt_R | OK | 5.02466 | 3.39935 | -0.563766 | -1.41013 |
| mut_A | wt_R | OK | 128.841 | 87.1478 | -0.564057 | -1.95464 |
| mut_A | wt_R | OK | 10.1603 | 6.86217 | -0.566203 | -1.73836 |
| mut_A | wt_R | OK | 299.705 | 202.408 | -0.566274 | -2.33211 |
| mut_A | wt_R | OK | 470.148 | 317.38 | -0.566904 | -2.31374 |
| mut_A | wt_R | OK | 228.351 | 154.107 | -0.567316 | -2.11888 |
| mut_A | wt_R | OK | 129.2 | 87.1398 | -0.568201 | -2.30758 |
| mut_A | wt_R | OK | 457.222 | 307.467 | -0.572464 | -2.35697 |
| mut_A | wt_R | OK | 994.89 | 668.863 | -0.572826 | -2.37204 |
| mut_A | wt_R | OK | 741.206 | 497.607 | -0.574867 | -2.35362 |

| | | | | | | |
|---|---|---|---|---|---|---|
| mut_A | wt_R | OK | 41.2007 | 27.599 | -0.578054 | -1.55397 |
| mut_A | wt_R | OK | 83.6455 | 55.9612 | -0.579859 | -2.18341 |
| mut_A | wt_R | OK | 123.211 | 82.4042 | -0.580343 | -2.3543 |
| mut_A | wt_R | OK | 577.793 | 385.579 | -0.583528 | -2.41452 |
| mut_A | wt_R | OK | 71.3502 | 47.6046 | -0.583817 | -2.32763 |
| mut_A | wt_R | OK | 53.4669 | 35.6494 | -0.584767 | -2.16935 |
| mut_A | wt_R | OK | 290.244 | 193.349 | -0.586059 | -2.40918 |
| mut_A | wt_R | OK | 216.855 | 144.437 | -0.586291 | -1.38586 |
| mut_A | wt_R | OK | 50.8234 | 33.8427 | -0.586648 | -2.26663 |
| mut_A | wt_R | OK | 309.883 | 206.216 | -0.587569 | -1.78139 |
| mut_A | wt_R | OK | 554.863 | 368.998 | -0.58852 | -2.42094 |
| mut_A | wt_R | OK | 46.8628 | 31.1176 | -0.590714 | -2.40411 |
| mut_A | wt_R | OK | 338.182 | 224.468 | -0.59129 | -2.43852 |
| mut_A | wt_R | OK | 58.4711 | 38.7793 | -0.592436 | -2.01986 |
| mut_A | wt_R | OK | 543.177 | 359.757 | -0.594401 | -2.44204 |
| mut_A | wt_R | OK | 385.732 | 255.392 | -0.594887 | -1.9848 |
| mut_A | wt_R | OK | 313.19 | 206.641 | -0.599915 | -1.3156 |
| mut_A | wt_R | OK | 1921.59 | 1266.53 | -0.60142 | -2.48356 |
| mut_A | wt_R | OK | 219.912 | 144.812 | -0.602744 | -2.46498 |
| mut_A | wt_R | OK | 298.183 | 196.127 | -0.604405 | -2.49226 |
| mut_A | wt_R | OK | 949.067 | 624.179 | -0.60455 | -2.48732 |
| mut_A | wt_R | OK | 11.9908 | 7.88585 | -0.604583 | -1.92069 |
| mut_A | wt_R | OK | 90.9532 | 59.806 | -0.604834 | -2.46668 |
| mut_A | wt_R | OK | 2201.17 | 1439.85 | -0.612353 | -2.5218 |
| mut_A | wt_R | OK | 135.45 | 88.3543 | -0.616393 | -1.65669 |
| mut_A | wt_R | OK | 35.2933 | 23.0057 | -0.617399 | -1.81343 |
| mut_A | wt_R | OK | 67.6561 | 43.973 | -0.621603 | -2.34818 |
| mut_A | wt_R | OK | 821.478 | 533.415 | -0.622965 | -2.5342 |
| mut_A | wt_R | OK | 17.254 | 11.2015 | -0.623243 | -1.68749 |
| mut_A | wt_R | OK | 15.3432 | 9.95458 | -0.624172 | -2.02714 |
| mut_A | wt_R | OK | 480.996 | 311.309 | -0.627679 | -2.54085 |
| mut_A | wt_R | OK | 91.3413 | 59.1161 | -0.627717 | -2.44886 |
| mut_A | wt_R | OK | 1055.15 | 680.246 | -0.633317 | -2.61609 |
| mut_A | wt_R | OK | 1142.87 | 736.38 | -0.634144 | -1.82973 |
| mut_A | wt_R | OK | 31.9509 | 20.5471 | -0.636922 | -2.44638 |
| mut_A | wt_R | OK | 944.92 | 607.077 | -0.638312 | -2.63096 |
| mut_A | wt_R | OK | 17.5372 | 11.2589 | -0.639343 | -2.21842 |
| mut_A | wt_R | OK | 320.689 | 205.822 | -0.639779 | -2.59245 |
| mut_A | wt_R | OK | 11.8283 | 7.5907 | -0.639939 | -1.66146 |
| mut_A | wt_R | OK | 34.845 | 22.3549 | -0.64036 | -1.4639 |
| mut_A | wt_R | OK | 841.974 | 539.985 | -0.640857 | -2.03802 |
| mut_A | wt_R | OK | 532.967 | 341.494 | -0.642188 | -2.22854 |
| mut_A | wt_R | OK | 90.1269 | 57.6563 | -0.64448 | -2.59127 |
| mut_A | wt_R | OK | 40.8536 | 26.1046 | -0.646161 | -2.44521 |
| mut_A | wt_R | OK | 1186.04 | 756.195 | -0.649318 | -2.67711 |
| mut_A | wt_R | OK | 9.52316 | 6.06528 | -0.650865 | -2.19946 |
| mut_A | wt_R | OK | 11.4891 | 7.30772 | -0.652775 | -1.79861 |
| mut_A | wt_R | OK | 35.5185 | 22.5375 | -0.656243 | -2.3027 |
| mut_A | wt_R | OK | 513.047 | 324.793 | -0.659571 | -2.64879 |
| mut_A | wt_R | OK | 57.7768 | 36.4572 | -0.664285 | -2.63092 |
| mut_A | wt_R | OK | 732.822 | 461.59 | -0.666849 | -2.67571 |

| | | | | | | |
|---|---|---|---|---|---|---|
| mut_A | wt_R | OK | 1471.9 | 926.126 | -0.668403 | -1.6609 |
| mut_A | wt_R | OK | 386.101 | 242.824 | -0.669064 | -1.52404 |
| mut_A | wt_R | OK | 73.8166 | 46.3934 | -0.670026 | -2.66201 |
| mut_A | wt_R | OK | 462.002 | 289.82 | -0.672744 | -2.7692 |
| mut_A | wt_R | OK | 10483.3 | 6573.27 | -0.673411 | -2.75973 |
| mut_A | wt_R | OK | 117.437 | 73.5576 | -0.674936 | -1.58768 |
| mut_A | wt_R | OK | 233.151 | 146.016 | -0.675138 | -2.74907 |
| mut_A | wt_R | OK | 1207.65 | 756.045 | -0.675654 | -1.69634 |
| mut_A | wt_R | OK | 709.67 | 443.02 | -0.679776 | -2.80944 |
| mut_A | wt_R | OK | 6.58423 | 4.10713 | -0.680885 | -1.49826 |
| mut_A | wt_R | OK | 527.088 | 327.831 | -0.685092 | -2.83035 |
| mut_A | wt_R | OK | 111.98 | 69.6414 | -0.685226 | -2.7936 |
| mut_A | wt_R | OK | 283.472 | 176.195 | -0.686038 | -2.69238 |
| mut_A | wt_R | OK | 282.199 | 175.2 | -0.687714 | -2.38934 |
| mut_A | wt_R | OK | 88.4507 | 54.817 | -0.69025 | -1.57413 |
| mut_A | wt_R | OK | 14.9193 | 9.16816 | -0.702476 | -1.96038 |
| mut_A | wt_R | OK | 40.4676 | 24.8174 | -0.705416 | -2.67613 |
| mut_A | wt_R | OK | 253.635 | 154.604 | -0.714181 | -2.84126 |
| mut_A | wt_R | OK | 112.907 | 68.8169 | -0.714299 | -2.68445 |
| mut_A | wt_R | OK | 7.55563 | 4.60333 | -0.714874 | -2.14011 |
| mut_A | wt_R | OK | 40.0775 | 24.4146 | -0.71505 | -2.77206 |
| mut_A | wt_R | OK | 103.564 | 63.0015 | -0.717065 | -2.84291 |
| mut_A | wt_R | OK | 9.04618 | 5.48533 | -0.72173 | -1.55652 |
| mut_A | wt_R | OK | 337.637 | 204.462 | -0.72364 | -1.44816 |
| mut_A | wt_R | OK | 62.7518 | 37.9773 | -0.724521 | -1.36718 |
| mut_A | wt_R | OK | 7823.45 | 4733.4 | -0.724927 | -2.99491 |
| mut_A | wt_R | OK | 22.0974 | 13.3365 | -0.728491 | -2.72954 |
| mut_A | wt_R | OK | 419.912 | 253.307 | -0.729197 | -2.97494 |
| mut_A | wt_R | OK | 33.1365 | 19.9673 | -0.730784 | -2.89067 |
| mut_A | wt_R | OK | 325.377 | 196.009 | -0.731191 | -3.01309 |
| mut_A | wt_R | OK | 90.9995 | 54.7864 | -0.73204 | -2.96776 |
| mut_A | wt_R | OK | 70.651 | 42.4644 | -0.734457 | -2.94451 |
| mut_A | wt_R | OK | 101.253 | 60.8546 | -0.734528 | -1.31816 |
| mut_A | wt_R | OK | 31.7785 | 19.0979 | -0.73464 | -2.8821 |
| mut_A | wt_R | OK | 26.7754 | 16.0695 | -0.736581 | -2.57035 |
| mut_A | wt_R | OK | 21887 | 13112.7 | -0.739112 | -2.99904 |
| mut_A | wt_R | OK | 169.116 | 101.276 | -0.739721 | -1.73912 |
| mut_A | wt_R | OK | 561.174 | 335.886 | -0.740475 | -2.9748 |
| mut_A | wt_R | OK | 189.628 | 113.227 | -0.743955 | -3.05948 |
| mut_A | wt_R | OK | 23.0061 | 13.7305 | -0.744637 | -2.52059 |
| mut_A | wt_R | OK | 406.966 | 241.692 | -0.751739 | -3.04792 |
| mut_A | wt_R | OK | 3.56432 | 2.11223 | -0.754856 | -1.94183 |
| mut_A | wt_R | OK | 123.467 | 73.1449 | -0.755301 | -3.01253 |
| mut_A | wt_R | OK | 129.138 | 76.4356 | -0.7566 | -3.02797 |
| mut_A | wt_R | OK | 1375.03 | 813.666 | -0.756958 | -3.07038 |
| mut_A | wt_R | OK | 726.291 | 429.359 | -0.758365 | -3.08066 |
| mut_A | wt_R | OK | 77.8376 | 46.0048 | -0.758684 | -2.95428 |
| mut_A | wt_R | OK | 183.807 | 108.62 | -0.75891 | -3.09842 |
| mut_A | wt_R | OK | 81.9318 | 48.4058 | -0.759243 | -2.98807 |
| mut_A | wt_R | OK | 5.0355 | 2.97029 | -0.76153 | -1.8153 |
| mut_A | wt_R | OK | 931.268 | 549.116 | -0.762085 | -3.15194 |

| | | | | | | |
|---|---|---|---|---|---|---|
| mut_A | wt_R | OK | 53.7352 | 31.6447 | -0.763904 | -2.99838 |
| mut_A | wt_R | OK | 170.394 | 100.321 | -0.76425 | -3.06902 |
| mut_A | wt_R | OK | 242.34 | 142.387 | -0.767222 | -2.56809 |
| mut_A | wt_R | OK | 9.57105 | 5.61409 | -0.769624 | -1.77585 |
| mut_A | wt_R | OK | 179.72 | 105.414 | -0.769688 | -3.17436 |
| mut_A | wt_R | OK | 71.4447 | 41.775 | -0.774189 | -2.80536 |
| mut_A | wt_R | OK | 153.663 | 89.7839 | -0.775237 | -3.14149 |
| mut_A | wt_R | OK | 26.174 | 15.2896 | -0.77559 | -3.01463 |
| mut_A | wt_R | OK | 298.948 | 174.456 | -0.777033 | -2.24987 |
| mut_A | wt_R | OK | 417.465 | 243.286 | -0.779005 | -3.20823 |
| mut_A | wt_R | OK | 145.52 | 84.6779 | -0.781155 | -2.98868 |
| mut_A | wt_R | OK | 690.191 | 400.855 | -0.783915 | -1.927 |
| mut_A | wt_R | OK | 75.1285 | 43.6145 | -0.784553 | -2.33182 |
| mut_A | wt_R | OK | 279.838 | 162.217 | -0.786665 | -2.79066 |
| mut_A | wt_R | OK | 6207.64 | 3595.38 | -0.787901 | -3.24646 |
| mut_A | wt_R | OK | 35.0472 | 20.2829 | -0.789034 | -3.06757 |
| mut_A | wt_R | OK | 75.3873 | 43.5546 | -0.791496 | -2.75684 |
| mut_A | wt_R | OK | 36.9902 | 21.3668 | -0.791772 | -3.12138 |
| mut_A | wt_R | OK | 215.898 | 124.539 | -0.793748 | -3.24007 |
| mut_A | wt_R | OK | 637.454 | 367.518 | -0.794504 | -3.26572 |
| mut_A | wt_R | OK | 18.0792 | 10.4086 | -0.796555 | -2.17491 |
| mut_A | wt_R | OK | 82.9905 | 47.7384 | -0.797797 | -3.16643 |
| mut_A | wt_R | OK | 390.704 | 224.331 | -0.800448 | -3.30213 |
| mut_A | wt_R | OK | 323.343 | 185.532 | -0.801402 | -2.87801 |
| mut_A | wt_R | OK | 5.40992 | 3.1039 | -0.801525 | -1.70583 |
| mut_A | wt_R | OK | 159.838 | 91.2587 | -0.808575 | -2.62736 |
| mut_A | wt_R | OK | 222.746 | 127.123 | -0.809176 | -3.285 |
| mut_A | wt_R | OK | 178.994 | 102.147 | -0.809271 | -2.95125 |
| mut_A | wt_R | OK | 58.8179 | 33.5546 | -0.809746 | -1.56115 |
| mut_A | wt_R | OK | 341.699 | 194.79 | -0.810807 | -3.3392 |
| mut_A | wt_R | OK | 30.7607 | 17.4827 | -0.815163 | -3.01618 |
| mut_A | wt_R | OK | 47.3229 | 26.7724 | -0.821792 | -3.25445 |
| mut_A | wt_R | OK | 1774.11 | 1002.74 | -0.823142 | -3.38064 |
| mut_A | wt_R | OK | 149.435 | 84.2789 | -0.826272 | -3.3994 |
| mut_A | wt_R | OK | 76.1623 | 42.9191 | -0.827457 | -2.70737 |
| mut_A | wt_R | OK | 69.136 | 38.9079 | -0.829372 | -1.37537 |
| mut_A | wt_R | OK | 233.107 | 131.153 | -0.82974 | -2.70206 |
| mut_A | wt_R | OK | 9.86031 | 5.54682 | -0.829972 | -2.62124 |
| mut_A | wt_R | OK | 15.8895 | 8.9344 | -0.830632 | -2.12782 |
| mut_A | wt_R | OK | 56.0332 | 31.479 | -0.831891 | -3.36979 |
| mut_A | wt_R | OK | 4808.41 | 2700.14 | -0.832523 | -3.37167 |
| mut_A | wt_R | OK | 401.833 | 225.533 | -0.83326 | -3.44172 |
| mut_A | wt_R | OK | 6.38584 | 3.58305 | -0.833686 | -2.60713 |
| mut_A | wt_R | OK | 46.2909 | 25.9718 | -0.83378 | -3.18208 |
| mut_A | wt_R | OK | 16.0519 | 9.003 | -0.834262 | -2.23367 |
| mut_A | wt_R | OK | 16.4003 | 9.19698 | -0.83449 | -2.69815 |
| mut_A | wt_R | OK | 20.4195 | 11.4356 | -0.836414 | -1.8575 |
| mut_A | wt_R | OK | 158.458 | 88.4075 | -0.841856 | -3.18613 |
| mut_A | wt_R | OK | 124.801 | 69.6079 | -0.842308 | -3.15553 |
| mut_A | wt_R | OK | 35.4462 | 19.6785 | -0.849015 | -2.79294 |
| mut_A | wt_R | OK | 1009.82 | 560.18 | -0.850139 | -3.49872 |

| | | | | | | |
|---|---|---|---|---|---|---|
| mut_A | wt_R | OK | 541.766 | 299.668 | -0.854307 | -3.52904 |
| mut_A | wt_R | OK | 253.31 | 139.951 | -0.85598 | -3.27495 |
| mut_A | wt_R | OK | 208.871 | 115.391 | -0.856085 | -1.90117 |
| mut_A | wt_R | OK | 39.375 | 21.7229 | -0.858064 | -1.63297 |
| mut_A | wt_R | OK | 562.019 | 309.57 | -0.860354 | -3.49552 |
| mut_A | wt_R | OK | 8832.23 | 4853.8 | -0.863663 | -3.53974 |
| mut_A | wt_R | OK | 133.521 | 73.3333 | -0.864528 | -3.54459 |
| mut_A | wt_R | OK | 262.885 | 143.657 | -0.871808 | -3.47933 |
| mut_A | wt_R | OK | 32.6084 | 17.8033 | -0.873101 | -2.08804 |
| mut_A | wt_R | OK | 28.3659 | 15.4588 | -0.875733 | -3.29367 |
| mut_A | wt_R | OK | 133.192 | 72.4591 | -0.878266 | -2.21038 |
| mut_A | wt_R | OK | 324.738 | 176.514 | -0.879494 | -3.60647 |
| mut_A | wt_R | OK | 5045.5 | 2738.73 | -0.881491 | -3.59297 |
| mut_A | wt_R | OK | 439.12 | 238.135 | -0.882835 | -3.6403 |
| mut_A | wt_R | OK | 65.5888 | 35.5161 | -0.884977 | -3.30903 |
| mut_A | wt_R | OK | 53.7639 | 29.1077 | -0.885238 | -1.49973 |
| mut_A | wt_R | OK | 86.0501 | 46.5725 | -0.885698 | -3.6044 |
| mut_A | wt_R | OK | 67.0104 | 36.1763 | -0.889339 | -3.58258 |
| mut_A | wt_R | OK | 3883.19 | 2094.87 | -0.890382 | -3.67399 |
| mut_A | wt_R | OK | 82.7153 | 44.2704 | -0.901812 | -3.6566 |
| mut_A | wt_R | OK | 26.1406 | 13.9831 | -0.902614 | -3.24993 |
| mut_A | wt_R | OK | 17.9073 | 9.57577 | -0.903088 | -2.23632 |
| mut_A | wt_R | OK | 442.314 | 236.156 | -0.90533 | -2.94673 |
| mut_A | wt_R | OK | 17.9892 | 9.57611 | -0.909617 | -2.33006 |
| mut_A | wt_R | OK | 47.714 | 25.3777 | -0.910851 | -3.18323 |
| mut_A | wt_R | OK | 96.7924 | 51.4693 | -0.911181 | -3.2364 |
| mut_A | wt_R | OK | 22.5062 | 11.9559 | -0.9126 | -1.66666 |
| mut_A | wt_R | OK | 36.9758 | 19.631 | -0.913447 | -3.58581 |
| mut_A | wt_R | OK | 9880.19 | 5228.28 | -0.918203 | -3.75397 |
| mut_A | wt_R | OK | 57.5488 | 30.4256 | -0.9195 | -3.63277 |
| mut_A | wt_R | OK | 13.2213 | 6.98001 | -0.921561 | -2.59532 |
| mut_A | wt_R | OK | 985.006 | 519.684 | -0.922498 | -3.74225 |
| mut_A | wt_R | OK | 25.1644 | 13.2632 | -0.923951 | -3.50175 |
| mut_A | wt_R | OK | 22.4125 | 11.7893 | -0.926824 | -2.14803 |
| mut_A | wt_R | OK | 27.0291 | 14.2043 | -0.928191 | -1.77845 |
| mut_A | wt_R | OK | 86.9647 | 45.6665 | -0.929296 | -3.01933 |
| mut_A | wt_R | OK | 124.841 | 65.5092 | -0.930317 | -3.75972 |
| mut_A | wt_R | OK | 19.9469 | 10.4515 | -0.932452 | -2.9591 |
| mut_A | wt_R | OK | 49.9183 | 26.0976 | -0.935652 | -3.25702 |
| mut_A | wt_R | OK | 77.2697 | 40.3349 | -0.937875 | -2.14912 |
| mut_A | wt_R | OK | 17.5075 | 9.13818 | -0.937997 | -2.27247 |
| mut_A | wt_R | OK | 53.9695 | 28.1132 | -0.940895 | -3.51724 |
| mut_A | wt_R | OK | 262.528 | 136.317 | -0.945501 | -3.81255 |
| mut_A | wt_R | OK | 3998.03 | 2073.04 | -0.947541 | -3.90884 |
| mut_A | wt_R | OK | 421.214 | 218.077 | -0.949716 | -3.82434 |
| mut_A | wt_R | OK | 49.0531 | 25.3778 | -0.950777 | -3.52363 |
| mut_A | wt_R | OK | 53.5646 | 27.6739 | -0.952753 | -3.61782 |
| mut_A | wt_R | OK | 14.5978 | 7.53547 | -0.953978 | -2.7891 |
| mut_A | wt_R | OK | 6753.64 | 3484.32 | -0.954787 | -3.95174 |
| mut_A | wt_R | OK | 124.849 | 64.4074 | -0.954886 | -3.69446 |
| mut_A | wt_R | OK | 280.745 | 144.798 | -0.955212 | -3.90266 |

| | | | | | | |
|---|---|---|---|---|---|---|
| mut_A | wt_R | OK | 1.66117 | 0.856748 | -0.955259 | -1.84609 |
| mut_A | wt_R | OK | 250.813 | 128.868 | -0.960726 | -3.46405 |
| mut_A | wt_R | OK | 282.085 | 144.803 | -0.962039 | -3.95353 |
| mut_A | wt_R | OK | 18.5707 | 9.5314 | -0.962268 | -3.67348 |
| mut_A | wt_R | OK | 27.0902 | 13.8891 | -0.963816 | -3.37351 |
| mut_A | wt_R | OK | 242.88 | 124.439 | -0.964805 | -3.92425 |
| mut_A | wt_R | OK | 870.125 | 445.688 | -0.965188 | -3.98506 |
| mut_A | wt_R | OK | 122.024 | 62.4893 | -0.965486 | -3.39786 |
| mut_A | wt_R | OK | 145.541 | 74.5033 | -0.966048 | -3.53725 |
| mut_A | wt_R | OK | 72.7905 | 37.1005 | -0.97231 | -3.88655 |
| mut_A | wt_R | OK | 354.503 | 180.304 | -0.975366 | -3.95863 |
| mut_A | wt_R | OK | 325.836 | 164.959 | -0.982044 | -3.57543 |
| mut_A | wt_R | OK | 473.979 | 239.87 | -0.982568 | -3.86004 |
| mut_A | wt_R | OK | 253.65 | 127.629 | -0.990882 | -4.03688 |
| mut_A | wt_R | OK | 50.2203 | 25.2604 | -0.991391 | -1.76153 |
| mut_A | wt_R | OK | 159.282 | 79.9994 | -0.993524 | -4.07947 |
| mut_A | wt_R | OK | 249.618 | 125.243 | -0.994997 | -3.95621 |
| mut_A | wt_R | OK | 176.73 | 88.5725 | -0.996615 | -3.5134 |
| mut_A | wt_R | OK | 6.82155 | 3.4137 | -0.998762 | -2.20099 |
| mut_A | wt_R | OK | 156.83 | 78.2684 | -1.0027 | -2.14334 |
| mut_A | wt_R | OK | 320.405 | 159.722 | -1.00434 | -4.12587 |
| mut_A | wt_R | OK | 350.529 | 174.25 | -1.00838 | -4.13425 |
| mut_A | wt_R | OK | 541.949 | 268.38 | -1.01388 | -3.3407 |
| mut_A | wt_R | OK | 100.886 | 49.935 | -1.0146 | -3.60681 |
| mut_A | wt_R | OK | 33.0528 | 16.3341 | -1.01688 | -3.96566 |
| mut_A | wt_R | OK | 81.4465 | 40.2109 | -1.01827 | -3.99694 |
| mut_A | wt_R | OK | 158.472 | 78.1939 | -1.0191 | -4.11584 |
| mut_A | wt_R | OK | 32.2366 | 15.895 | -1.02013 | -3.34151 |
| mut_A | wt_R | OK | 13.6716 | 6.73355 | -1.02174 | -3.64866 |
| mut_A | wt_R | OK | 7984.49 | 3932.26 | -1.02184 | -4.20523 |
| mut_A | wt_R | OK | 62.5871 | 30.8078 | -1.02257 | -2.45899 |
| mut_A | wt_R | OK | 2615.34 | 1287.28 | -1.02267 | -4.21744 |
| mut_A | wt_R | OK | 16.4056 | 8.07248 | -1.02311 | -2.92591 |
| mut_A | wt_R | OK | 108.615 | 53.3785 | -1.02489 | -3.68063 |
| mut_A | wt_R | OK | 10620.2 | 5213.46 | -1.0265 | -4.21376 |
| mut_A | wt_R | OK | 95.0237 | 46.6193 | -1.02736 | -4.1559 |
| mut_A | wt_R | OK | 141.544 | 69.4262 | -1.0277 | -2.36918 |
| mut_A | wt_R | OK | 62.7769 | 30.7652 | -1.02893 | -4.1793 |
| mut_A | wt_R | OK | 23.047 | 11.2517 | -1.03444 | -1.4793 |
| mut_A | wt_R | OK | 46.786 | 22.8141 | -1.03615 | -3.27509 |
| mut_A | wt_R | OK | 138.441 | 67.3258 | -1.04004 | -4.23401 |
| mut_A | wt_R | OK | 9.06051 | 4.39188 | -1.04475 | -2.64338 |
| mut_A | wt_R | OK | 133.125 | 64.1592 | -1.05305 | -4.24719 |
| mut_A | wt_R | OK | 91.3297 | 44.0145 | -1.05311 | -2.60838 |
| mut_A | wt_R | OK | 43.1161 | 20.5686 | -1.06778 | -1.64096 |
| mut_A | wt_R | OK | 6153.27 | 2931.73 | -1.0696 | -4.41921 |
| mut_A | wt_R | OK | 7.94434 | 3.75008 | -1.083 | -2.64605 |
| mut_A | wt_R | OK | 163.391 | 77.0966 | -1.08359 | -4.02406 |
| mut_A | wt_R | OK | 152.431 | 71.9068 | -1.08396 | -3.779 |
| mut_A | wt_R | OK | 95.2683 | 44.9011 | -1.08525 | -4.10297 |
| mut_A | wt_R | OK | 100.032 | 47.1426 | -1.08535 | -4.32084 |

| | | | | | | |
|---|---|---|---|---|---|---|
| mut_A | wt_R | OK | 68.4067 | 32.2347 | -1.08552 | -3.68905 |
| mut_A | wt_R | OK | 159.658 | 75.1381 | -1.08736 | -4.14919 |
| mut_A | wt_R | OK | 513.119 | 240.723 | -1.09192 | -4.45896 |
| mut_A | wt_R | OK | 1603.01 | 750.274 | -1.09529 | -4.41501 |
| mut_A | wt_R | OK | 310.753 | 145.142 | -1.0983 | -4.48975 |
| mut_A | wt_R | OK | 166.495 | 77.6148 | -1.10107 | -4.06317 |
| mut_A | wt_R | OK | 534.028 | 248.128 | -1.10583 | -2.2421 |
| mut_A | wt_R | OK | 4021.01 | 1868.07 | -1.10601 | -4.54276 |
| mut_A | wt_R | OK | 25.8372 | 12.0032 | -1.10602 | -3.78391 |
| mut_A | wt_R | OK | 157.263 | 72.9097 | -1.10899 | -3.80823 |
| mut_A | wt_R | OK | 674.711 | 311.253 | -1.11618 | -4.56811 |
| mut_A | wt_R | OK | 104.561 | 48.0677 | -1.1212 | -3.15083 |
| mut_A | wt_R | OK | 2903.01 | 1330.16 | -1.12595 | -4.63474 |
| mut_A | wt_R | OK | 22.9331 | 10.5028 | -1.12666 | -3.68068 |
| mut_A | wt_R | OK | 34.1413 | 15.6175 | -1.12836 | -3.31996 |
| mut_A | wt_R | OK | 165.647 | 75.6399 | -1.13089 | -3.86209 |
| mut_A | wt_R | OK | 3314.07 | 1505.36 | -1.1385 | -4.60465 |
| mut_A | wt_R | OK | 7.86152 | 3.56784 | -1.13976 | -2.04969 |
| mut_A | wt_R | OK | 256.736 | 116.292 | -1.14253 | -4.62653 |
| mut_A | wt_R | OK | 52.3567 | 23.6951 | -1.14378 | -4.10619 |
| mut_A | wt_R | OK | 96.649 | 43.6828 | -1.14569 | -4.33771 |
| mut_A | wt_R | OK | 41.3879 | 18.651 | -1.14995 | -4.38211 |
| mut_A | wt_R | OK | 914.525 | 410.939 | -1.1541 | -4.21229 |
| mut_A | wt_R | OK | 48.4266 | 21.75 | -1.15478 | -4.35547 |
| mut_A | wt_R | OK | 547.825 | 245.925 | -1.1555 | -4.60665 |
| mut_A | wt_R | OK | 108.376 | 48.6334 | -1.15603 | -4.53055 |
| mut_A | wt_R | OK | 61.0562 | 27.3655 | -1.15778 | -4.3732 |
| mut_A | wt_R | OK | 5.68685 | 2.54544 | -1.15971 | -2.7557 |
| mut_A | wt_R | OK | 26.6036 | 11.9009 | -1.16055 | -3.67088 |
| mut_A | wt_R | OK | 5157.81 | 2300.9 | -1.16456 | -4.80613 |
| mut_A | wt_R | OK | 23.093 | 10.2972 | -1.1652 | -4.29858 |
| mut_A | wt_R | OK | 6.18431 | 2.75491 | -1.16661 | -2.5398 |
| mut_A | wt_R | OK | 1.69671 | 0.755567 | -1.16711 | -2.20846 |
| mut_A | wt_R | OK | 110.922 | 49.3462 | -1.16854 | -4.66048 |
| mut_A | wt_R | OK | 21.8643 | 9.68438 | -1.17485 | -3.85852 |
| mut_A | wt_R | OK | 22.5844 | 9.99092 | -1.17664 | -4.04152 |
| mut_A | wt_R | OK | 326.686 | 143.889 | -1.18294 | -4.83167 |
| mut_A | wt_R | OK | 17.7542 | 7.81313 | -1.18419 | -3.07364 |
| mut_A | wt_R | OK | 48.7174 | 21.3152 | -1.19256 | -4.37227 |
| mut_A | wt_R | OK | 52.9736 | 23.1368 | -1.19509 | -2.11517 |
| mut_A | wt_R | OK | 9.03825 | 3.93639 | -1.19917 | -2.20161 |
| mut_A | wt_R | OK | 272.669 | 117.804 | -1.21076 | -2.40856 |
| mut_A | wt_R | OK | 53.5832 | 23.1202 | -1.21263 | -4.71969 |
| mut_A | wt_R | OK | 50.008 | 21.5755 | -1.21277 | -4.45507 |
| mut_A | wt_R | OK | 10.2443 | 4.41763 | -1.21348 | -3.15755 |
| mut_A | wt_R | OK | 15.7088 | 6.77086 | -1.21416 | -4.50016 |
| mut_A | wt_R | OK | 46.8995 | 20.2035 | -1.21497 | -3.43772 |
| mut_A | wt_R | OK | 95.5555 | 41.1394 | -1.21582 | -4.76531 |
| mut_A | wt_R | OK | 38.4013 | 16.4553 | -1.2226 | -3.71114 |
| mut_A | wt_R | OK | 96.7659 | 41.4165 | -1.22429 | -4.84255 |
| mut_A | wt_R | OK | 425.309 | 181.92 | -1.22521 | -5.03003 |

| | | | | | | |
|---|---|---|---|---|---|---|
| mut_A | wt_R | OK | 431.617 | 183.928 | -1.23061 | -5.07805 |
| mut_A | wt_R | OK | 461.618 | 194.336 | -1.24815 | -5.14627 |
| mut_A | wt_R | OK | 189.222 | 79.6152 | -1.24896 | -2.03679 |
| mut_A | wt_R | OK | 12.3276 | 5.185 | -1.24948 | -4.1993 |
| mut_A | wt_R | OK | 97.5483 | 40.994 | -1.2507 | -5.0472 |
| mut_A | wt_R | OK | 1233.89 | 517.422 | -1.2538 | -5.06745 |
| mut_A | wt_R | OK | 18.3963 | 7.63483 | -1.26875 | -4.29743 |
| mut_A | wt_R | OK | 77.9845 | 32.1927 | -1.27645 | -4.51802 |
| mut_A | wt_R | OK | 308.815 | 127.429 | -1.27705 | -4.61925 |
| mut_A | wt_R | OK | 359.913 | 148.364 | -1.27851 | -3.70343 |
| mut_A | wt_R | OK | 234.771 | 96.7587 | -1.27879 | -5.23918 |
| mut_A | wt_R | OK | 37.7384 | 15.53 | -1.28097 | -4.3267 |
| mut_A | wt_R | OK | 2057.57 | 840.169 | -1.29219 | -3.79946 |
| mut_A | wt_R | OK | 16.1431 | 6.57021 | -1.2969 | -1.73908 |
| mut_A | wt_R | OK | 5.64938 | 2.28841 | -1.30375 | -2.53126 |
| mut_A | wt_R | OK | 913.813 | 368.926 | -1.30857 | -5.34753 |
| mut_A | wt_R | OK | 43.1828 | 17.4275 | -1.30909 | -1.71259 |
| mut_A | wt_R | OK | 108.515 | 43.7895 | -1.30924 | -3.67765 |
| mut_A | wt_R | OK | 1353.6 | 544.973 | -1.31254 | -5.43155 |
| mut_A | wt_R | OK | 127.704 | 51.3783 | -1.31357 | -2.70265 |
| mut_A | wt_R | OK | 48.079 | 19.3398 | -1.31383 | -4.98275 |
| mut_A | wt_R | OK | 204.826 | 81.6408 | -1.32704 | -2.58165 |
| mut_A | wt_R | OK | 109.374 | 43.5858 | -1.32734 | -5.04865 |
| mut_A | wt_R | OK | 45.9792 | 18.1916 | -1.33771 | -3.13906 |
| mut_A | wt_R | OK | 4.01017 | 1.58556 | -1.33867 | -2.76454 |
| mut_A | wt_R | OK | 135.193 | 53.3767 | -1.34074 | -5.42854 |
| mut_A | wt_R | OK | 62.6209 | 24.72 | -1.34097 | -3.33355 |
| mut_A | wt_R | OK | 295.615 | 116.41 | -1.34451 | -5.52511 |
| mut_A | wt_R | OK | 83.8589 | 32.9859 | -1.34612 | -5.46668 |
| mut_A | wt_R | OK | 90.0776 | 35.4286 | -1.34625 | -3.86921 |
| mut_A | wt_R | OK | 48.8829 | 19.1926 | -1.34878 | -4.98707 |
| mut_A | wt_R | OK | 124.171 | 48.5198 | -1.35568 | -3.83986 |
| mut_A | wt_R | OK | 314.776 | 122.874 | -1.35714 | -5.08757 |
| mut_A | wt_R | OK | 76.8815 | 29.7569 | -1.36941 | -3.96186 |
| mut_A | wt_R | OK | 52.513 | 20.3179 | -1.36993 | -5.21258 |
| mut_A | wt_R | OK | 61.8764 | 23.8758 | -1.37384 | -5.48564 |
| mut_A | wt_R | OK | 125.465 | 48.344 | -1.37587 | -2.64573 |
| mut_A | wt_R | OK | 43.2946 | 16.6114 | -1.38202 | -5.20963 |
| mut_A | wt_R | OK | 83.3386 | 31.9751 | -1.38204 | -2.46372 |
| mut_A | wt_R | OK | 39.7652 | 15.1666 | -1.39061 | -3.95645 |
| mut_A | wt_R | OK | 509.012 | 193.654 | -1.39422 | -5.68658 |
| mut_A | wt_R | OK | 165.324 | 62.7054 | -1.39863 | -5.60806 |
| mut_A | wt_R | OK | 310.107 | 116.926 | -1.40717 | -5.75407 |
| mut_A | wt_R | OK | 80.6859 | 30.2699 | -1.41443 | -3.10259 |
| mut_A | wt_R | OK | 1259.19 | 471.055 | -1.41853 | -5.82551 |
| mut_A | wt_R | OK | 447.732 | 167.154 | -1.42145 | -5.71041 |
| mut_A | wt_R | OK | 18.8518 | 7.00894 | -1.42743 | -3.5651 |
| mut_A | wt_R | OK | 20.7374 | 7.68765 | -1.43162 | -2.9424 |
| mut_A | wt_R | OK | 47.9558 | 17.7688 | -1.43236 | -2.95626 |
| mut_A | wt_R | OK | 65.7044 | 24.3179 | -1.43397 | -5.60503 |
| mut_A | wt_R | OK | 257.426 | 95.1491 | -1.4359 | -4.88644 |

| | | | | | | |
|---|---|---|---|---|---|---|
| mut_A | wt_R | OK | 181.825 | 67.164 | -1.43679 | -5.83508 |
| mut_A | wt_R | OK | 43.0533 | 15.8967 | -1.43739 | -2.50352 |
| mut_A | wt_R | OK | 89.2256 | 32.9127 | -1.43881 | -5.51625 |
| mut_A | wt_R | OK | 101.863 | 37.4545 | -1.44342 | -2.86842 |
| mut_A | wt_R | OK | 112.184 | 41.1503 | -1.4469 | -4.51727 |
| mut_A | wt_R | OK | 19.2869 | 7.06981 | -1.44788 | -3.12843 |
| mut_A | wt_R | OK | 189.276 | 69.2027 | -1.45159 | -5.94321 |
| mut_A | wt_R | OK | 36.6991 | 13.3974 | -1.45379 | -4.27912 |
| mut_A | wt_R | OK | 81.781 | 29.6092 | -1.46572 | -5.68934 |
| mut_A | wt_R | OK | 44.0575 | 15.9168 | -1.46884 | -4.92745 |
| mut_A | wt_R | OK | 202.598 | 73.0233 | -1.47219 | -1.74648 |
| mut_A | wt_R | OK | 95.0469 | 34.2038 | -1.47448 | -4.16545 |
| mut_A | wt_R | OK | 54.0366 | 19.3139 | -1.48429 | -5.84502 |
| mut_A | wt_R | OK | 835.765 | 298.628 | -1.48475 | -4.26845 |
| mut_A | wt_R | OK | 380.787 | 135.985 | -1.48554 | -4.02423 |
| mut_A | wt_R | OK | 94.5716 | 33.6803 | -1.4895 | -5.40228 |
| mut_A | wt_R | OK | 184.164 | 65.5239 | -1.4909 | -5.44478 |
| mut_A | wt_R | OK | 28.5917 | 10.142 | -1.49526 | -4.63544 |
| mut_A | wt_R | OK | 64.8022 | 22.9793 | -1.49571 | -4.46163 |
| mut_A | wt_R | OK | 41.3378 | 14.5978 | -1.50172 | -2.95593 |
| mut_A | wt_R | OK | 8380.57 | 2901.37 | -1.53032 | -6.30266 |
| mut_A | wt_R | OK | 455.747 | 157.703 | -1.53102 | -6.23934 |
| mut_A | wt_R | OK | 59.4873 | 20.5458 | -1.53374 | -5.45054 |
| mut_A | wt_R | OK | 44.5584 | 15.3751 | -1.5351 | -3.54585 |
| mut_A | wt_R | OK | 2396.66 | 822.244 | -1.54339 | -6.36301 |
| mut_A | wt_R | OK | 33.6677 | 11.5182 | -1.54744 | -4.92142 |
| mut_A | wt_R | OK | 284.404 | 97.0149 | -1.55166 | -4.45664 |
| mut_A | wt_R | OK | 471.561 | 160.622 | -1.55378 | -6.24375 |
| mut_A | wt_R | OK | 9.90979 | 3.34984 | -1.56476 | -5.03224 |
| mut_A | wt_R | OK | 195.72 | 66.1195 | -1.56564 | -5.90077 |
| mut_A | wt_R | OK | 31.5383 | 10.5457 | -1.58045 | -4.73484 |
| mut_A | wt_R | OK | 231.248 | 76.9547 | -1.58736 | -6.40231 |
| mut_A | wt_R | OK | 275.237 | 91.2561 | -1.59268 | -3.57639 |
| mut_A | wt_R | OK | 572.75 | 189.893 | -1.59272 | -6.51463 |
| mut_A | wt_R | OK | 36.0435 | 11.9403 | -1.5939 | -5.13921 |
| mut_A | wt_R | OK | 9.59275 | 3.1561 | -1.6038 | -3.84679 |
| mut_A | wt_R | OK | 187.272 | 61.6124 | -1.60384 | -5.55114 |
| mut_A | wt_R | OK | 656.262 | 215.303 | -1.6079 | -6.6617 |
| mut_A | wt_R | OK | 126.547 | 41.3088 | -1.61515 | -6.3535 |
| mut_A | wt_R | OK | 388.644 | 126.48 | -1.61954 | -6.54677 |
| mut_A | wt_R | OK | 96.9731 | 31.4053 | -1.62658 | -6.57783 |
| mut_A | wt_R | OK | 321.686 | 103.54 | -1.63547 | -4.8598 |
| mut_A | wt_R | OK | 19.4322 | 6.212 | -1.64532 | -2.59701 |
| mut_A | wt_R | OK | 1015.65 | 323.841 | -1.64904 | -1.93778 |
| mut_A | wt_R | OK | 270.004 | 84.9334 | -1.66858 | -6.72227 |
| mut_A | wt_R | OK | 38.405 | 11.6909 | -1.71591 | -6.00114 |
| mut_A | wt_R | OK | 57.4189 | 17.2622 | -1.73391 | -3.63359 |
| mut_A | wt_R | OK | 162.038 | 48.6688 | -1.73526 | -6.75093 |
| mut_A | wt_R | OK | 188.679 | 56.658 | -1.73558 | -6.98342 |
| mut_A | wt_R | OK | 441.569 | 132.509 | -1.73654 | -6.73361 |
| mut_A | wt_R | OK | 310.997 | 93.1889 | -1.73867 | -5.60794 |

| | | | | | | |
|---|---|---|---|---|---|---|
| mut_A | wt_R | OK | 48.4644 | 14.2989 | -1.76102 | -2.30625 |
| mut_A | wt_R | OK | 16.7358 | 4.90859 | -1.76956 | -2.32972 |
| mut_A | wt_R | OK | 117.197 | 34.3422 | -1.77088 | -5.6467 |
| mut_A | wt_R | OK | 97.2152 | 28.4286 | -1.77384 | -6.78175 |
| mut_A | wt_R | OK | 896.145 | 262.058 | -1.77385 | -4.3008 |
| mut_A | wt_R | OK | 18.0296 | 5.26878 | -1.77483 | -4.78362 |
| mut_A | wt_R | OK | 445.927 | 130.261 | -1.7754 | -7.22818 |
| mut_A | wt_R | OK | 117.877 | 34.2728 | -1.78214 | -6.88406 |
| mut_A | wt_R | OK | 138.487 | 39.8362 | -1.7976 | -6.46646 |
| mut_A | wt_R | OK | 842.16 | 241.589 | -1.80154 | -2.06979 |
| mut_A | wt_R | OK | 278.427 | 79.1578 | -1.8145 | -4.36074 |
| mut_A | wt_R | OK | 203.122 | 57.448 | -1.82202 | -7.46095 |
| mut_A | wt_R | OK | 23.6721 | 6.69302 | -1.82246 | -3.28093 |
| mut_A | wt_R | OK | 186.114 | 52.5924 | -1.82326 | -7.22201 |
| mut_A | wt_R | OK | 160.068 | 45.0865 | -1.82792 | -6.98083 |
| mut_A | wt_R | OK | 837.736 | 235.939 | -1.82808 | -4.52677 |
| mut_A | wt_R | OK | 157.273 | 44.1139 | -1.83397 | -3.36114 |
| mut_A | wt_R | OK | 188.55 | 52.5708 | -1.84261 | -7.39319 |
| mut_A | wt_R | OK | 21.7432 | 6.00014 | -1.85749 | -5.90366 |
| mut_A | wt_R | OK | 407.912 | 112.305 | -1.86084 | -7.62987 |
| mut_A | wt_R | OK | 374.296 | 100.02 | -1.9039 | -7.71247 |
| mut_A | wt_R | OK | 88.7519 | 23.7104 | -1.90426 | -2.57889 |
| mut_A | wt_R | OK | 217.411 | 57.9126 | -1.90848 | -7.19225 |
| mut_A | wt_R | OK | 837.862 | 223.044 | -1.90938 | -7.68918 |
| mut_A | wt_R | OK | 1373.52 | 365.066 | -1.91165 | -3.78137 |
| mut_A | wt_R | OK | 8.76834 | 2.32928 | -1.91242 | -3.62785 |
| mut_A | wt_R | OK | 102.246 | 27.083 | -1.91659 | -7.73661 |
| mut_A | wt_R | OK | 105.902 | 27.8012 | -1.92952 | -6.49653 |
| mut_A | wt_R | OK | 106.411 | 27.8535 | -1.93371 | -5.11772 |
| mut_A | wt_R | OK | 100.229 | 26.1285 | -1.9396 | -7.71499 |
| mut_A | wt_R | OK | 20.3213 | 5.28775 | -1.94226 | -2.82906 |
| mut_A | wt_R | OK | 122.306 | 31.4392 | -1.95986 | -7.58388 |
| mut_A | wt_R | OK | 205.351 | 52.1151 | -1.97832 | -4.43851 |
| mut_A | wt_R | OK | 251.291 | 63.6213 | -1.98178 | -5.9667 |
| mut_A | wt_R | OK | 2896.54 | 728.422 | -1.99149 | -8.16065 |
| mut_A | wt_R | OK | 1506.62 | 378.337 | -1.99357 | -3.80816 |
| mut_A | wt_R | OK | 117.118 | 29.2876 | -1.9996 | -7.90126 |
| mut_A | wt_R | OK | 53.9104 | 13.4363 | -2.00443 | -6.80575 |
| mut_A | wt_R | OK | 9.51841 | 2.36934 | -2.00624 | -4.43282 |
| mut_A | wt_R | OK | 52.2189 | 12.9246 | -2.01445 | -6.53562 |
| mut_A | wt_R | OK | 239.922 | 58.8581 | -2.02725 | -8.28772 |
| mut_A | wt_R | OK | 140.074 | 34.0462 | -2.04062 | -8.19651 |
| mut_A | wt_R | OK | 468.842 | 113.265 | -2.0494 | -8.2713 |
| mut_A | wt_R | OK | 92.4814 | 22.2754 | -2.05371 | -7.94591 |
| mut_A | wt_R | OK | 70.401 | 16.8014 | -2.06701 | -7.2345 |
| mut_A | wt_R | OK | 126.567 | 30.0282 | -2.07551 | -8.43841 |
| mut_A | wt_R | OK | 30.4026 | 7.07907 | -2.10256 | -6.98117 |
| mut_A | wt_R | OK | 377.88 | 87.7048 | -2.1072 | -8.58673 |
| mut_A | wt_R | OK | 77.6756 | 17.9153 | -2.11627 | -7.80104 |
| mut_A | wt_R | OK | 374.343 | 85.9568 | -2.12268 | -8.36942 |
| mut_A | wt_R | OK | 36.8241 | 8.44903 | -2.12379 | -5.75652 |

| | | | | | | |
|---|---|---|---|---|---|---|
| mut_A | wt_R | OK | 89.0232 | 20.2152 | -2.13874 | -7.82673 |
| mut_A | wt_R | OK | 530.855 | 120.336 | -2.14125 | -8.83247 |
| mut_A | wt_R | OK | 40.7822 | 9.18246 | -2.15099 | -7.28821 |
| mut_A | wt_R | OK | 621.258 | 138.969 | -2.16044 | -8.42995 |
| mut_A | wt_R | OK | 168.278 | 37.502 | -2.1658 | -8.32054 |
| mut_A | wt_R | OK | 319.133 | 70.5952 | -2.17651 | -6.83418 |
| mut_A | wt_R | OK | 473.95 | 102.726 | -2.20594 | -5.58714 |
| mut_A | wt_R | OK | 320.433 | 69.1463 | -2.2123 | -8.86939 |
| mut_A | wt_R | OK | 7.02085 | 1.50354 | -2.22329 | -1.8125 |
| mut_A | wt_R | OK | 285.02 | 60.5031 | -2.23598 | -9.02477 |
| mut_A | wt_R | OK | 85.2354 | 18.0643 | -2.23831 | -8.22517 |
| mut_A | wt_R | OK | 2008.49 | 423.23 | -2.2466 | -9.14825 |
| mut_A | wt_R | OK | 823.667 | 172.759 | -2.2533 | -9.20132 |
| mut_A | wt_R | OK | 704.656 | 146.669 | -2.26436 | -8.39801 |
| mut_A | wt_R | OK | 592.66 | 122.475 | -2.27472 | -9.2624 |
| mut_A | wt_R | OK | 1776.6 | 365.745 | -2.28021 | -9.2897 |
| mut_A | wt_R | OK | 338.884 | 69.4889 | -2.28594 | -8.78394 |
| mut_A | wt_R | OK | 845.431 | 169.643 | -2.31719 | -9.55443 |
| mut_A | wt_R | OK | 14.2038 | 2.84258 | -2.321 | -2.25264 |
| mut_A | wt_R | OK | 96.0708 | 19.062 | -2.3334 | -8.8591 |
| mut_A | wt_R | OK | 35.3579 | 6.93459 | -2.35015 | -7.99891 |
| mut_A | wt_R | OK | 25.6712 | 5.00664 | -2.35823 | -5.98769 |
| mut_A | wt_R | OK | 398.416 | 76.1546 | -2.38727 | -5.78595 |
| mut_A | wt_R | OK | 52.6275 | 10.0555 | -2.38783 | -7.78887 |
| mut_A | wt_R | OK | 176.711 | 33.7378 | -2.38895 | -7.4767 |
| mut_A | wt_R | OK | 898.957 | 171.459 | -2.39039 | -9.81991 |
| mut_A | wt_R | OK | 185.994 | 34.8683 | -2.41527 | -5.64883 |
| mut_A | wt_R | OK | 51.8544 | 9.56723 | -2.43829 | -7.16874 |
| mut_A | wt_R | OK | 705.476 | 128.614 | -2.45555 | -10.0601 |
| mut_A | wt_R | OK | 89.0358 | 16.1219 | -2.46537 | -4.09467 |
| mut_A | wt_R | OK | 10389.3 | 1855.62 | -2.48513 | -10.2548 |
| mut_A | wt_R | OK | 64.8333 | 11.4722 | -2.4986 | -4.7162 |
| mut_A | wt_R | OK | 375.159 | 65.1554 | -2.52554 | -3.87137 |
| mut_A | wt_R | OK | 15.6119 | 2.68344 | -2.54049 | -3.46342 |
| mut_A | wt_R | OK | 113.52 | 19.278 | -2.55792 | -6.45476 |
| mut_A | wt_R | OK | 850.652 | 142.767 | -2.5749 | -10.4889 |
| mut_A | wt_R | OK | 116.919 | 18.9626 | -2.62428 | -10.1917 |
| mut_A | wt_R | OK | 347.626 | 55.811 | -2.63892 | -9.96797 |
| mut_A | wt_R | OK | 648.149 | 104.005 | -2.63968 | -10.614 |
| mut_A | wt_R | OK | 48108.7 | 7713.6 | -2.64082 | -10.4281 |
| mut_A | wt_R | OK | 30.2576 | 4.78792 | -2.65983 | -8.43356 |
| mut_A | wt_R | OK | 1961.34 | 302.378 | -2.69741 | -11.0759 |
| mut_A | wt_R | OK | 174.534 | 26.8247 | -2.70187 | -10.677 |
| mut_A | wt_R | OK | 33635.4 | 5124.59 | -2.71447 | -10.7019 |
| mut_A | wt_R | OK | 42.372 | 6.39563 | -2.72795 | -2.93604 |
| mut_A | wt_R | OK | 973.21 | 143.508 | -2.76162 | -11.1353 |
| mut_A | wt_R | OK | 94.6178 | 13.9116 | -2.76582 | -9.86253 |
| mut_A | wt_R | OK | 331.261 | 46.2714 | -2.83978 | -9.14485 |
| mut_A | wt_R | OK | 226.63 | 27.7473 | -3.02992 | -11.6345 |
| mut_A | wt_R | OK | 1516.88 | 175.37 | -3.11263 | -12.7209 |
| mut_A | wt_R | OK | 56.1343 | 6.30958 | -3.15327 | -10.3896 |

| | | | | | | |
|---|---|---|---|---|---|---|
| mut_A | wt_R | OK | 703.533 | 78.2347 | -3.16874 | -12.4214 |
| mut_A | wt_R | OK | 700.348 | 75.642 | -3.21081 | -10.5843 |
| mut_A | wt_R | OK | 3399.74 | 345.716 | -3.29776 | -12.3159 |
| mut_A | wt_R | OK | 202532 | 17173.3 | -3.55991 | -14.5762 |
| mut_A | wt_R | OK | 2039.5 | 171.308 | -3.57355 | -14.1415 |
| mut_A | wt_R | OK | 40.0359 | 2.92741 | -3.7736 | -10.2546 |
| mut_A | wt_R | OK | 710.44 | 49.6686 | -3.83831 | -15.4736 |
| mut_A | wt_R | OK | 640.818 | 42.8504 | -3.90254 | -15.5741 |
| mut_A | wt_R | OK | 105.347 | 6.80098 | -3.95327 | -11.2253 |
| mut_A | wt_R | OK | 133.018 | 7.26972 | -4.19358 | -6.37317 |
| mut_A | wt_R | OK | 2231.8 | 81.5208 | -4.7749 | -19.5074 |
| mut_A | wt_R | OK | 2466.3 | 86.1855 | -4.83876 | -18.3226 |
| mut_A | wt_R | OK | 697.944 | 24.2409 | -4.8476 | -18.9662 |
| mut_A | wt_R | OK | 2210.92 | 76.6098 | -4.85097 | -19.8652 |
| mut_A | wt_R | OK | 563.03 | 11.1716 | -5.6553 | -17.9499 |
| mut_A | wt_R | OK | 3823.21 | 17.5841 | -7.76437 | -28.3053 |
| mut_A | wt_R | OK | 2918.41 | 12.9372 | -7.81752 | -27.4723 |
| mut_A | wt_R | OK | 3160.45 | 13.525 | -7.86836 | -27.5681 |
| mut_A | wt_R | OK | 3836.11 | 11.4284 | -8.39088 | -29.0079 |
| mut_A | wt_R | OK | 4177.95 | 11.9149 | -8.45389 | -29.4092 |
| mut_A | wt_R | OK | 7015.9 | 11.4073 | -9.26452 | -31.8054 |

| p_value | q_value | significant | | |
|---|---|---|---|---|
| 0.02395 | 0.0414616 | yes | | |
| 0.00215 | 0.00455164 | yes | | |
| 0.00465 | 0.00923245 | yes | | |
| 5.00E-05 | 0.00013743 | yes | | |
| 5.00E-05 | 0.00013743 | yes | | |
| 5.00E-05 | 0.00013743 | yes | | |
| 5.00E-05 | 0.00013743 | yes | | |
| 5.00E-05 | 0.00013743 | yes | | |
| 5.00E-05 | 0.00013743 | yes | | |
| 5.00E-05 | 0.00013743 | yes | | |
| 5.00E-05 | 0.00013743 | yes | | |
| 5.00E-05 | 0.00013743 | yes | | |
| 5.00E-05 | 0.00013743 | yes | | |
| 5.00E-05 | 0.00013743 | yes | | |
| 5.00E-05 | 0.00013743 | yes | | |
| 5.00E-05 | 0.00013743 | yes | | |
| 5.00E-05 | 0.00013743 | yes | | |
| 5.00E-05 | 0.00013743 | yes | | |
| 5.00E-05 | 0.00013743 | yes | 5.37846632 | 2.42719484 |
| 5.00E-05 | 0.00013743 | yes | | |
| 5.00E-05 | 0.00013743 | yes | | |
| 5.00E-05 | 0.00013743 | yes | | |
| 5.00E-05 | 0.00013743 | yes | | |
| 5.00E-05 | 0.00013743 | yes | | |
| 5.00E-05 | 0.00013743 | yes | | |
| 5.00E-05 | 0.00013743 | yes | | |
| 5.00E-05 | 0.00013743 | yes | | |
| 5.00E-05 | 0.00013743 | yes | | |
| 5.00E-05 | 0.00013743 | yes | | |
| 5.00E-05 | 0.00013743 | yes | | |
| 5.00E-05 | 0.00013743 | yes | | |
| 5.00E-05 | 0.00013743 | yes | | |
| 5.00E-05 | 0.00013743 | yes | | |
| 5.00E-05 | 0.00013743 | yes | | |
| 5.00E-05 | 0.00013743 | yes | | |
| 5.00E-05 | 0.00013743 | yes | | |
| 5.00E-05 | 0.00013743 | yes | | |
| 5.00E-05 | 0.00013743 | yes | | |
| 5.00E-05 | 0.00013743 | yes | | |
| 5.00E-05 | 0.00013743 | yes | | |
| 5.00E-05 | 0.00013743 | yes | | |
| 5.00E-05 | 0.00013743 | yes | | |
| 5.00E-05 | 0.00013743 | yes | | |
| 5.00E-05 | 0.00013743 | yes | | |
| 5.00E-05 | 0.00013743 | yes | | |
| 5.00E-05 | 0.00013743 | yes | | |
| 5.00E-05 | 0.00013743 | yes | | |
| 5.00E-05 | 0.00013743 | yes | | |
| 5.00E-05 | 0.00013743 | yes | | |

| | | |
|---|---|---|
| 5.00E-05 | 0.00013743 | yes |
| 5.00E-05 | 0.00013743 | yes |
| 5.00E-05 | 0.00013743 | yes |
| 5.00E-05 | 0.00013743 | yes |
| 5.00E-05 | 0.00013743 | yes |
| 5.00E-05 | 0.00013743 | yes |
| 5.00E-05 | 0.00013743 | yes |
| 5.00E-05 | 0.00013743 | yes |
| 5.00E-05 | 0.00013743 | yes |
| 5.00E-05 | 0.00013743 | yes |
| 5.00E-05 | 0.00013743 | yes |
| 5.00E-05 | 0.00013743 | yes |
| 5.00E-05 | 0.00013743 | yes |
| 5.00E-05 | 0.00013743 | yes |
| 5.00E-05 | 0.00013743 | yes |
| 5.00E-05 | 0.00013743 | yes |
| 5.00E-05 | 0.00013743 | yes |
| 5.00E-05 | 0.00013743 | yes |
| 5.00E-05 | 0.00013743 | yes |
| 5.00E-05 | 0.00013743 | yes |
| 5.00E-05 | 0.00013743 | yes |
| 5.00E-05 | 0.00013743 | yes |
| 5.00E-05 | 0.00013743 | yes |
| 5.00E-05 | 0.00013743 | yes |
| 5.00E-05 | 0.00013743 | yes |
| 5.00E-05 | 0.00013743 | yes |
| 5.00E-05 | 0.00013743 | yes |
| 5.00E-05 | 0.00013743 | yes |
| 5.00E-05 | 0.00013743 | yes |
| 0.0001 | 0.00026595 | yes |
| 5.00E-05 | 0.00013743 | yes |
| 5.00E-05 | 0.00013743 | yes |
| 5.00E-05 | 0.00013743 | yes |
| 5.00E-05 | 0.00013743 | yes |
| 5.00E-05 | 0.00013743 | yes |
| 5.00E-05 | 0.00013743 | yes |
| 5.00E-05 | 0.00013743 | yes |
| 5.00E-05 | 0.00013743 | yes |
| 5.00E-05 | 0.00013743 | yes |
| 5.00E-05 | 0.00013743 | yes |
| 5.00E-05 | 0.00013743 | yes |
| 0.0013 | 0.00286816 | yes |
| 5.00E-05 | 0.00013743 | yes |
| 5.00E-05 | 0.00013743 | yes |
| 5.00E-05 | 0.00013743 | yes |
| 5.00E-05 | 0.00013743 | yes |
| 5.00E-05 | 0.00013743 | yes |
| 5.00E-05 | 0.00013743 | yes |
| 5.00E-05 | 0.00013743 | yes |
| 5.00E-05 | 0.00013743 | yes |
| 5.00E-05 | 0.00013743 | yes |

| | | |
|---|---|---|
| 5.00E-05 | 0.00013743 | yes |
| 5.00E-05 | 0.00013743 | yes |
| 5.00E-05 | 0.00013743 | yes |
| 5.00E-05 | 0.00013743 | yes |
| 5.00E-05 | 0.00013743 | yes |
| 5.00E-05 | 0.00013743 | yes |
| 5.00E-05 | 0.00013743 | yes |
| 0.0001 | 0.00026595 | yes |
| 5.00E-05 | 0.00013743 | yes |
| 0.00225 | 0.00474293 | yes |
| 5.00E-05 | 0.00013743 | yes |
| 5.00E-05 | 0.00013743 | yes |
| 5.00E-05 | 0.00013743 | yes |
| 5.00E-05 | 0.00013743 | yes |
| 5.00E-05 | 0.00013743 | yes |
| 5.00E-05 | 0.00013743 | yes |
| 5.00E-05 | 0.00013743 | yes |
| 5.00E-05 | 0.00013743 | yes |
| 5.00E-05 | 0.00013743 | yes |
| 5.00E-05 | 0.00013743 | yes |
| 5.00E-05 | 0.00013743 | yes |
| 5.00E-05 | 0.00013743 | yes |
| 5.00E-05 | 0.00013743 | yes |
| 5.00E-05 | 0.00013743 | yes |
| 5.00E-05 | 0.00013743 | yes |
| 5.00E-05 | 0.00013743 | yes |
| 5.00E-05 | 0.00013743 | yes |
| 5.00E-05 | 0.00013743 | yes |
| 5.00E-05 | 0.00013743 | yes |
| 5.00E-05 | 0.00013743 | yes |
| 5.00E-05 | 0.00013743 | yes |
| 5.00E-05 | 0.00013743 | yes |
| 5.00E-05 | 0.00013743 | yes |
| 5.00E-05 | 0.00013743 | yes |
| 5.00E-05 | 0.00013743 | yes |
| 5.00E-05 | 0.00013743 | yes |
| 5.00E-05 | 0.00013743 | yes |
| 0.0027 | 0.00560506 | yes |
| 5.00E-05 | 0.00013743 | yes |
| 5.00E-05 | 0.00013743 | yes |
| 0.0125 | 0.0229993 | yes |
| 5.00E-05 | 0.00013743 | yes |
| 5.00E-05 | 0.00013743 | yes |
| 0.0001 | 0.00026595 | yes |
| 5.00E-05 | 0.00013743 | yes |
| 5.00E-05 | 0.00013743 | yes |
| 5.00E-05 | 0.00013743 | yes |
| 5.00E-05 | 0.00013743 | yes |
| 5.00E-05 | 0.00013743 | yes |
| 5.00E-05 | 0.00013743 | yes |
| 5.00E-05 | 0.00013743 | yes |

| | | |
|---|---|---|
| 5.00E-05 | 0.00013743 | yes |
| 5.00E-05 | 0.00013743 | yes |
| 5.00E-05 | 0.00013743 | yes |
| 5.00E-05 | 0.00013743 | yes |
| 5.00E-05 | 0.00013743 | yes |
| 0.0139 | 0.0253101 | yes |
| 5.00E-05 | 0.00013743 | yes |
| 5.00E-05 | 0.00013743 | yes |
| 5.00E-05 | 0.00013743 | yes |
| 5.00E-05 | 0.00013743 | yes |
| 5.00E-05 | 0.00013743 | yes |
| 5.00E-05 | 0.00013743 | yes |
| 5.00E-05 | 0.00013743 | yes |
| 5.00E-05 | 0.00013743 | yes |
| 5.00E-05 | 0.00013743 | yes |
| 5.00E-05 | 0.00013743 | yes |
| 5.00E-05 | 0.00013743 | yes |
| 5.00E-05 | 0.00013743 | yes |
| 5.00E-05 | 0.00013743 | yes |
| 5.00E-05 | 0.00013743 | yes |
| 5.00E-05 | 0.00013743 | yes |
| 5.00E-05 | 0.00013743 | yes |
| 5.00E-05 | 0.00013743 | yes |
| 5.00E-05 | 0.00013743 | yes |
| 5.00E-05 | 0.00013743 | yes |
| 5.00E-05 | 0.00013743 | yes |
| 5.00E-05 | 0.00013743 | yes |
| 5.00E-05 | 0.00013743 | yes |
| 5.00E-05 | 0.00013743 | yes |
| 0.0001 | 0.00026595 | yes |
| 5.00E-05 | 0.00013743 | yes |
| 5.00E-05 | 0.00013743 | yes |
| 5.00E-05 | 0.00013743 | yes |
| 5.00E-05 | 0.00013743 | yes |
| 5.00E-05 | 0.00013743 | yes |
| 0.02935 | 0.049897 | yes |
| 5.00E-05 | 0.00013743 | yes |
| 5.00E-05 | 0.00013743 | yes |
| 5.00E-05 | 0.00013743 | yes |
| 5.00E-05 | 0.00013743 | yes |
| 5.00E-05 | 0.00013743 | yes |
| 5.00E-05 | 0.00013743 | yes |
| 5.00E-05 | 0.00013743 | yes |
| 5.00E-05 | 0.00013743 | yes |
| 5.00E-05 | 0.00013743 | yes |
| 5.00E-05 | 0.00013743 | yes |
| 5.00E-05 | 0.00013743 | yes |
| 5.00E-05 | 0.00013743 | yes |
| 5.00E-05 | 0.00013743 | yes |
| 5.00E-05 | 0.00013743 | yes |
| 5.00E-05 | 0.00013743 | yes |

| | | |
|---|---|---|
| 5.00E-05 | 0.00013743 | yes |
| 5.00E-05 | 0.00013743 | yes |
| 5.00E-05 | 0.00013743 | yes |
| 5.00E-05 | 0.00013743 | yes |
| 5.00E-05 | 0.00013743 | yes |
| 5.00E-05 | 0.00013743 | yes |
| 5.00E-05 | 0.00013743 | yes |
| 0.00115 | 0.00256018 | yes |
| 5.00E-05 | 0.00013743 | yes |
| 0.0015 | 0.00327128 | yes |
| 5.00E-05 | 0.00013743 | yes |
| 0.00065 | 0.00151996 | yes |
| 5.00E-05 | 0.00013743 | yes |
| 0.0026 | 0.00542494 | yes |
| 5.00E-05 | 0.00013743 | yes |
| 5.00E-05 | 0.00013743 | yes |
| 5.00E-05 | 0.00013743 | yes |
| 5.00E-05 | 0.00013743 | yes |
| 5.00E-05 | 0.00013743 | yes |
| 5.00E-05 | 0.00013743 | yes |
| 5.00E-05 | 0.00013743 | yes |
| 5.00E-05 | 0.00013743 | yes |
| 5.00E-05 | 0.00013743 | yes |
| 0.00015 | 0.00038924 | yes |
| 5.00E-05 | 0.00013743 | yes |
| 5.00E-05 | 0.00013743 | yes |
| 5.00E-05 | 0.00013743 | yes |
| 5.00E-05 | 0.00013743 | yes |
| 5.00E-05 | 0.00013743 | yes |
| 5.00E-05 | 0.00013743 | yes |
| 5.00E-05 | 0.00013743 | yes |
| 5.00E-05 | 0.00013743 | yes |
| 5.00E-05 | 0.00013743 | yes |
| 5.00E-05 | 0.00013743 | yes |
| 5.00E-05 | 0.00013743 | yes |
| 5.00E-05 | 0.00013743 | yes |
| 5.00E-05 | 0.00013743 | yes |
| 5.00E-05 | 0.00013743 | yes |
| 5.00E-05 | 0.00013743 | yes |
| 5.00E-05 | 0.00013743 | yes |
| 5.00E-05 | 0.00013743 | yes |
| 5.00E-05 | 0.00013743 | yes |
| 5.00E-05 | 0.00013743 | yes |
| 0.0001 | 0.00026595 | yes |
| 5.00E-05 | 0.00013743 | yes |
| 5.00E-05 | 0.00013743 | yes |
| 5.00E-05 | 0.00013743 | yes |
| 5.00E-05 | 0.00013743 | yes |
| 5.00E-05 | 0.00013743 | yes |
| 5.00E-05 | 0.00013743 | yes |
| 5.00E-05 | 0.00013743 | yes |

| | | |
|---|---|---|
| 5.00E-05 | 0.00013743 | yes |
| 5.00E-05 | 0.00013743 | yes |
| 5.00E-05 | 0.00013743 | yes |
| 5.00E-05 | 0.00013743 | yes |
| 5.00E-05 | 0.00013743 | yes |
| 5.00E-05 | 0.00013743 | yes |
| 5.00E-05 | 0.00013743 | yes |
| 5.00E-05 | 0.00013743 | yes |
| 5.00E-05 | 0.00013743 | yes |
| 5.00E-05 | 0.00013743 | yes |
| 5.00E-05 | 0.00013743 | yes |
| 5.00E-05 | 0.00013743 | yes |
| 5.00E-05 | 0.00013743 | yes |
| 5.00E-05 | 0.00013743 | yes |
| 5.00E-05 | 0.00013743 | yes |
| 5.00E-05 | 0.00013743 | yes |
| 5.00E-05 | 0.00013743 | yes |
| 5.00E-05 | 0.00013743 | yes |
| 0.00105 | 0.0023589 | yes |
| 5.00E-05 | 0.00013743 | yes |
| 5.00E-05 | 0.00013743 | yes |
| 0.0206 | 0.036223 | yes |
| 5.00E-05 | 0.00013743 | yes |
| 5.00E-05 | 0.00013743 | yes |
| 5.00E-05 | 0.00013743 | yes |
| 5.00E-05 | 0.00013743 | yes |
| 0.00055 | 0.00130598 | yes |
| 5.00E-05 | 0.00013743 | yes |
| 5.00E-05 | 0.00013743 | yes |
| 5.00E-05 | 0.00013743 | yes |
| 5.00E-05 | 0.00013743 | yes |
| 5.00E-05 | 0.00013743 | yes |
| 5.00E-05 | 0.00013743 | yes |
| 5.00E-05 | 0.00013743 | yes |
| 5.00E-05 | 0.00013743 | yes |
| 5.00E-05 | 0.00013743 | yes |
| 5.00E-05 | 0.00013743 | yes |
| 5.00E-05 | 0.00013743 | yes |
| 5.00E-05 | 0.00013743 | yes |
| 5.00E-05 | 0.00013743 | yes |
| 5.00E-05 | 0.00013743 | yes |
| 5.00E-05 | 0.00013743 | yes |
| 5.00E-05 | 0.00013743 | yes |
| 5.00E-05 | 0.00013743 | yes |
| 5.00E-05 | 0.00013743 | yes |
| 5.00E-05 | 0.00013743 | yes |
| 5.00E-05 | 0.00013743 | yes |
| 5.00E-05 | 0.00013743 | yes |
| 5.00E-05 | 0.00013743 | yes |
| 5.00E-05 | 0.00013743 | yes |
| 5.00E-05 | 0.00013743 | yes |

| | | |
|---|---|---|
| 5.00E-05 | 0.00013743 | yes |
| 5.00E-05 | 0.00013743 | yes |
| 5.00E-05 | 0.00013743 | yes |
| 5.00E-05 | 0.00013743 | yes |
| 5.00E-05 | 0.00013743 | yes |
| 5.00E-05 | 0.00013743 | yes |
| 5.00E-05 | 0.00013743 | yes |
| 5.00E-05 | 0.00013743 | yes |
| 0.00995 | 0.0186705 | yes |
| 0.00055 | 0.00130598 | yes |
| 5.00E-05 | 0.00013743 | yes |
| 0.0008 | 0.00183411 | yes |
| 5.00E-05 | 0.00013743 | yes |
| 5.00E-05 | 0.00013743 | yes |
| 5.00E-05 | 0.00013743 | yes |
| 0.0002 | 0.00051037 | yes |
| 5.00E-05 | 0.00013743 | yes |
| 5.00E-05 | 0.00013743 | yes |
| 5.00E-05 | 0.00013743 | yes |
| 5.00E-05 | 0.00013743 | yes |
| 5.00E-05 | 0.00013743 | yes |
| 5.00E-05 | 0.00013743 | yes |
| 5.00E-05 | 0.00013743 | yes |
| 5.00E-05 | 0.00013743 | yes |
| 5.00E-05 | 0.00013743 | yes |
| 5.00E-05 | 0.00013743 | yes |
| 0.00305 | 0.0062525 | yes |
| 5.00E-05 | 0.00013743 | yes |
| 5.00E-05 | 0.00013743 | yes |
| 0.01875 | 0.0333514 | yes |
| 5.00E-05 | 0.00013743 | yes |
| 5.00E-05 | 0.00013743 | yes |
| 5.00E-05 | 0.00013743 | yes |
| 5.00E-05 | 0.00013743 | yes |
| 5.00E-05 | 0.00013743 | yes |
| 5.00E-05 | 0.00013743 | yes |
| 5.00E-05 | 0.00013743 | yes |
| 5.00E-05 | 0.00013743 | yes |
| 0.0021 | 0.00446114 | yes |
| 5.00E-05 | 0.00013743 | yes |
| 5.00E-05 | 0.00013743 | yes |
| 5.00E-05 | 0.00013743 | yes |
| 5.00E-05 | 0.00013743 | yes |
| 5.00E-05 | 0.00013743 | yes |
| 5.00E-05 | 0.00013743 | yes |
| 5.00E-05 | 0.00013743 | yes |
| 5.00E-05 | 0.00013743 | yes |
| 5.00E-05 | 0.00013743 | yes |
| 5.00E-05 | 0.00013743 | yes |
| 5.00E-05 | 0.00013743 | yes |
| 5.00E-05 | 0.00013743 | yes |

| | | |
|---|---|---|
| 5.00E-05 | 0.00013743 | yes |
| 0.0021 | 0.00446114 | yes |
| 5.00E-05 | 0.00013743 | yes |
| 5.00E-05 | 0.00013743 | yes |
| 5.00E-05 | 0.00013743 | yes |
| 5.00E-05 | 0.00013743 | yes |
| 5.00E-05 | 0.00013743 | yes |
| 5.00E-05 | 0.00013743 | yes |
| 5.00E-05 | 0.00013743 | yes |
| 0.00025 | 0.00063012 | yes |
| 5.00E-05 | 0.00013743 | yes |
| 5.00E-05 | 0.00013743 | yes |
| 5.00E-05 | 0.00013743 | yes |
| 5.00E-05 | 0.00013743 | yes |
| 5.00E-05 | 0.00013743 | yes |
| 5.00E-05 | 0.00013743 | yes |
| 0.00225 | 0.00474293 | yes |
| 5.00E-05 | 0.00013743 | yes |
| 5.00E-05 | 0.00013743 | yes |
| 5.00E-05 | 0.00013743 | yes |
| 5.00E-05 | 0.00013743 | yes |
| 5.00E-05 | 0.00013743 | yes |
| 5.00E-05 | 0.00013743 | yes |
| 5.00E-05 | 0.00013743 | yes |
| 5.00E-05 | 0.00013743 | yes |
| 0.02205 | 0.0385522 | yes |
| 5.00E-05 | 0.00013743 | yes |
| 5.00E-05 | 0.00013743 | yes |
| 5.00E-05 | 0.00013743 | yes |
| 5.00E-05 | 0.00013743 | yes |
| 0.01405 | 0.0255643 | yes |
| 5.00E-05 | 0.00013743 | yes |
| 5.00E-05 | 0.00013743 | yes |
| 5.00E-05 | 0.00013743 | yes |
| 5.00E-05 | 0.00013743 | yes |
| 5.00E-05 | 0.00013743 | yes |
| 5.00E-05 | 0.00013743 | yes |
| 0.0097 | 0.0182431 | yes |
| 5.00E-05 | 0.00013743 | yes |
| 0.00015 | 0.00038924 | yes |
| 5.00E-05 | 0.00013743 | yes |
| 0.0056 | 0.0109594 | yes |
| 5.00E-05 | 0.00013743 | yes |
| 5.00E-05 | 0.00013743 | yes |
| 0.00035 | 0.00086186 | yes |
| 5.00E-05 | 0.00013743 | yes |
| 0.0002 | 0.00051037 | yes |
| 5.00E-05 | 0.00013743 | yes |
| 0.00015 | 0.00038924 | yes |
| 5.00E-05 | 0.00013743 | yes |
| 5.00E-05 | 0.00013743 | yes |

| | | |
|---|---|---|
| 0.0009 | 0.00204621 | yes |
| 5.00E-05 | 0.00013743 | yes |
| 5.00E-05 | 0.00013743 | yes |
| 5.00E-05 | 0.00013743 | yes |
| 0.00325 | 0.00664037 | yes |
| 5.00E-05 | 0.00013743 | yes |
| 5.00E-05 | 0.00013743 | yes |
| 5.00E-05 | 0.00013743 | yes |
| 0.0035 | 0.0071157 | yes |
| 5.00E-05 | 0.00013743 | yes |
| 5.00E-05 | 0.00013743 | yes |
| 5.00E-05 | 0.00013743 | yes |
| 5.00E-05 | 0.00013743 | yes |
| 5.00E-05 | 0.00013743 | yes |
| 5.00E-05 | 0.00013743 | yes |
| 5.00E-05 | 0.00013743 | yes |
| 5.00E-05 | 0.00013743 | yes |
| 5.00E-05 | 0.00013743 | yes |
| 5.00E-05 | 0.00013743 | yes |
| 0.0006 | 0.00141244 | yes |
| 5.00E-05 | 0.00013743 | yes |
| 5.00E-05 | 0.00013743 | yes |
| 5.00E-05 | 0.00013743 | yes |
| 5.00E-05 | 0.00013743 | yes |
| 5.00E-05 | 0.00013743 | yes |
| 5.00E-05 | 0.00013743 | yes |
| 5.00E-05 | 0.00013743 | yes |
| 5.00E-05 | 0.00013743 | yes |
| 5.00E-05 | 0.00013743 | yes |
| 0.0005 | 0.00119766 | yes |
| 5.00E-05 | 0.00013743 | yes |
| 5.00E-05 | 0.00013743 | yes |
| 5.00E-05 | 0.00013743 | yes |
| 5.00E-05 | 0.00013743 | yes |
| 5.00E-05 | 0.00013743 | yes |
| 0.00055 | 0.00130598 | yes |
| 0.0008 | 0.00183411 | yes |
| 0.0141 | 0.0256364 | yes |
| 0.0003 | 0.00074621 | yes |
| 5.00E-05 | 0.00013743 | yes |
| 5.00E-05 | 0.00013743 | yes |
| 5.00E-05 | 0.00013743 | yes |
| 5.00E-05 | 0.00013743 | yes |
| 0.02635 | 0.0452345 | yes |
| 0.0012 | 0.00266186 | yes |
| 5.00E-05 | 0.00013743 | yes |
| 0.0005 | 0.00119766 | yes |
| 0.00445 | 0.00886397 | yes |
| 5.00E-05 | 0.00013743 | yes |
| 0.00925 | 0.0174368 | yes |
| 5.00E-05 | 0.00013743 | yes |

| | | |
|---|---|---|
| 5.00E-05 | 0.00013743 | yes |
| 5.00E-05 | 0.00013743 | yes |
| 0.021 | 0.0368474 | yes |
| 5.00E-05 | 0.00013743 | yes |
| 5.00E-05 | 0.00013743 | yes |
| 5.00E-05 | 0.00013743 | yes |
| 0.0001 | 0.00026595 | yes |
| 0.0065 | 0.0125806 | yes |
| 5.00E-05 | 0.00013743 | yes |
| 5.00E-05 | 0.00013743 | yes |
| 0.0224 | 0.0390809 | yes |
| 5.00E-05 | 0.00013743 | yes |
| 5.00E-05 | 0.00013743 | yes |
| 5.00E-05 | 0.00013743 | yes |
| 5.00E-05 | 0.00013743 | yes |
| 5.00E-05 | 0.00013743 | yes |
| 5.00E-05 | 0.00013743 | yes |
| 0.0153 | 0.0276547 | yes |
| 0.0002 | 0.00051037 | yes |
| 5.00E-05 | 0.00013743 | yes |
| 5.00E-05 | 0.00013743 | yes |
| 5.00E-05 | 0.00013743 | yes |
| 0.0001 | 0.00026595 | yes |
| 0.001 | 0.00225275 | yes |
| 0.0005 | 0.00119766 | yes |
| 5.00E-05 | 0.00013743 | yes |
| 0.00055 | 0.00130598 | yes |
| 0.0001 | 0.00026595 | yes |
| 0.0001 | 0.00026595 | yes |
| 5.00E-05 | 0.00013743 | yes |
| 5.00E-05 | 0.00013743 | yes |
| 0.0001 | 0.00026595 | yes |
| 5.00E-05 | 0.00013743 | yes |
| 0.00015 | 0.00038924 | yes |
| 5.00E-05 | 0.00013743 | yes |
| 0.00805 | 0.0153155 | yes |
| 5.00E-05 | 0.00013743 | yes |
| 5.00E-05 | 0.00013743 | yes |
| 0.0001 | 0.00026595 | yes |
| 0.0002 | 0.00051037 | yes |
| 0.00015 | 0.00038924 | yes |
| 0.0015 | 0.00327128 | yes |
| 5.00E-05 | 0.00013743 | yes |
| 5.00E-05 | 0.00013743 | yes |
| 5.00E-05 | 0.00013743 | yes |
| 5.00E-05 | 0.00013743 | yes |
| 5.00E-05 | 0.00013743 | yes |
| 0.00035 | 0.00086186 | yes |
| 5.00E-05 | 0.00013743 | yes |
| 5.00E-05 | 0.00013743 | yes |
| 0.00065 | 0.00151996 | yes |

| | | |
|---|---|---|
| 5.00E-05 | 0.00013743 | yes |
| 0.00015 | 0.00038924 | yes |
| 0.001 | 0.00225275 | yes |
| 0.0002 | 0.00051037 | yes |
| 5.00E-05 | 0.00013743 | yes |
| 0.00015 | 0.00038924 | yes |
| 0.00015 | 0.00038924 | yes |
| 0.01115 | 0.0207795 | yes |
| 0.0023 | 0.00483177 | yes |
| 5.00E-05 | 0.00013743 | yes |
| 0.00575 | 0.0112351 | yes |
| 0.0001 | 0.00026595 | yes |
| 0.0001 | 0.00026595 | yes |
| 0.0029 | 0.00598992 | yes |
| 0.0007 | 0.00162914 | yes |
| 5.00E-05 | 0.00013743 | yes |
| 0.0116 | 0.0215366 | yes |
| 5.00E-05 | 0.00013743 | yes |
| 0.00015 | 0.00038924 | yes |
| 0.00015 | 0.00038924 | yes |
| 0.00685 | 0.0131958 | yes |
| 0.0003 | 0.00074621 | yes |
| 0.01755 | 0.0314672 | yes |
| 5.00E-05 | 0.00013743 | yes |
| 5.00E-05 | 0.00013743 | yes |
| 0.00015 | 0.00038924 | yes |
| 5.00E-05 | 0.00013743 | yes |
| 0.0006 | 0.00141244 | yes |
| 0.00015 | 0.00038924 | yes |
| 0.0001 | 0.00026595 | yes |
| 0.0004 | 0.00097813 | yes |
| 5.00E-05 | 0.00013743 | yes |
| 0.00075 | 0.00173402 | yes |
| 0.00025 | 0.00063012 | yes |
| 0.0004 | 0.00097813 | yes |
| 0.00015 | 0.00038924 | yes |
| 0.0001 | 0.00026595 | yes |
| 0.00015 | 0.00038924 | yes |
| 0.001 | 0.00225275 | yes |
| 5.00E-05 | 0.00013743 | yes |
| 0.02795 | 0.0477811 | yes |
| 0.0055 | 0.0107809 | yes |
| 5.00E-05 | 0.00013743 | yes |
| 0.0003 | 0.00074621 | yes |
| 0.0002 | 0.00051037 | yes |
| 0.01105 | 0.0206088 | yes |
| 0.00015 | 0.00038924 | yes |
| 0.0013 | 0.00286816 | yes |
| 0.00025 | 0.00063012 | yes |
| 0.00195 | 0.00417493 | yes |
| 0.00035 | 0.00086186 | yes |

| | | |
|---|---|---|
| 0.01325 | 0.0241982 | yes |
| 0.0002 | 0.00051037 | yes |
| 0.00175 | 0.00376311 | yes |
| 0.0002 | 0.00051037 | yes |
| 0.00035 | 0.00086186 | yes |
| 0.0004 | 0.00097813 | yes |
| 0.00025 | 0.00063012 | yes |
| 0.0003 | 0.00074621 | yes |
| 0.00035 | 0.00086186 | yes |
| 5.00E-05 | 0.00013743 | yes |
| 0.0003 | 0.00074621 | yes |
| 0.01565 | 0.0282665 | yes |
| 0.0004 | 0.00097813 | yes |
| 0.00075 | 0.00173402 | yes |
| 0.0199 | 0.0351176 | yes |
| 0.00015 | 0.00038924 | yes |
| 0.0003 | 0.00074621 | yes |
| 0.0005 | 0.00119766 | yes |
| 0.00075 | 0.00173402 | yes |
| 0.00065 | 0.00151996 | yes |
| 0.0003 | 0.00074621 | yes |
| 0.0067 | 0.0129474 | yes |
| 0.00045 | 0.0010885 | yes |
| 0.0006 | 0.00141244 | yes |
| 0.00045 | 0.0010885 | yes |
| 0.00035 | 0.00086186 | yes |
| 0.00045 | 0.0010885 | yes |
| 0.0006 | 0.00141244 | yes |
| 0.0005 | 0.00119766 | yes |
| 0.0006 | 0.00141244 | yes |
| 0.0007 | 0.00162914 | yes |
| 0.0005 | 0.00119766 | yes |
| 0.00795 | 0.0151605 | yes |
| 0.00065 | 0.00151996 | yes |
| 0.00055 | 0.00130598 | yes |
| 0.0007 | 0.00162914 | yes |
| 0.00065 | 0.00151996 | yes |
| 0.0232 | 0.0402482 | yes |
| 0.0273 | 0.0467674 | yes |
| 0.0024 | 0.00502896 | yes |
| 0.00075 | 0.00173402 | yes |
| 0.00085 | 0.0019415 | yes |
| 0.01805 | 0.0322228 | yes |
| 0.0013 | 0.00286816 | yes |
| 0.001 | 0.00225275 | yes |
| 0.0014 | 0.003075 | yes |
| 0.0011 | 0.00246224 | yes |
| 0.00265 | 0.0055199 | yes |
| 0.0009 | 0.00204621 | yes |
| 0.0008 | 0.00183411 | yes |
| 0.00095 | 0.0021559 | yes |

| | | |
|---|---|---|
| 0.00135 | 0.00297048 | yes |
| 0.001 | 0.00225275 | yes |
| 0.0006 | 0.00141244 | yes |
| 0.0059 | 0.0115099 | yes |
| 0.0055 | 0.0107809 | yes |
| 0.0034 | 0.00692958 | yes |
| 0.001 | 0.00225275 | yes |
| 0.001 | 0.00225275 | yes |
| 0.0189 | 0.0335697 | yes |
| 0.00405 | 0.00810659 | yes |
| 0.00085 | 0.0019415 | yes |
| 0.0044 | 0.00877859 | yes |
| 0.0011 | 0.00246224 | yes |
| 0.0008 | 0.00183411 | yes |
| 0.00155 | 0.00336243 | yes |
| 0.00215 | 0.00455164 | yes |
| 0.0037 | 0.0074852 | yes |
| 0.00125 | 0.00277027 | yes |
| 0.0026 | 0.00542494 | yes |
| 0.0014 | 0.003075 | yes |
| 0.00175 | 0.00376311 | yes |
| 0.0139 | 0.0253101 | yes |
| 0.0008 | 0.00183411 | yes |
| 0.01285 | 0.0235727 | yes |
| 0.00455 | 0.00905583 | yes |
| 0.0027 | 0.00560506 | yes |
| 0.00175 | 0.00376311 | yes |
| 0.00145 | 0.00317914 | yes |
| 0.0016 | 0.00346174 | yes |
| 0.0013 | 0.00286816 | yes |
| 0.002 | 0.00427826 | yes |
| 0.0021 | 0.00446114 | yes |
| 0.00215 | 0.00455164 | yes |
| 0.0056 | 0.0109594 | yes |
| 0.0083 | 0.0157546 | yes |
| 0.00205 | 0.00437002 | yes |
| 0.00155 | 0.00336243 | yes |
| 0.02265 | 0.039461 | yes |
| 0.0019 | 0.00407498 | yes |
| 0.01415 | 0.0256893 | yes |
| 0.0034 | 0.00692958 | yes |
| 0.00275 | 0.00569444 | yes |
| 0.00245 | 0.00512936 | yes |
| 0.00235 | 0.00493259 | yes |
| 0.0028 | 0.0057931 | yes |
| 0.0036 | 0.00730091 | yes |
| 0.0023 | 0.00483177 | yes |
| 0.00275 | 0.00569444 | yes |
| 0.0027 | 0.00560506 | yes |
| 0.0031 | 0.00634971 | yes |
| 0.00305 | 0.0062525 | yes |

| | | |
|---|---|---|
| 0.0025 | 0.00522959 | yes |
| 0.00275 | 0.00569444 | yes |
| 0.0037 | 0.0074852 | yes |
| 0.00305 | 0.0062525 | yes |
| 0.003 | 0.00617057 | yes |
| 0.00345 | 0.00701985 | yes |
| 0.02505 | 0.0432746 | yes |
| 0.0027 | 0.00560506 | yes |
| 0.00355 | 0.0072114 | yes |
| 0.00345 | 0.00701985 | yes |
| 0.00385 | 0.00775676 | yes |
| 0.018 | 0.0321569 | yes |
| 0.006 | 0.0116865 | yes |
| 0.01305 | 0.0238684 | yes |
| 0.004 | 0.0080261 | yes |
| 0.01185 | 0.021918 | yes |
| 0.0055 | 0.0107809 | yes |
| 0.00445 | 0.00886397 | yes |
| 0.004 | 0.0080261 | yes |
| 0.00405 | 0.00810659 | yes |
| 0.0059 | 0.0115099 | yes |
| 0.00535 | 0.0105204 | yes |
| 0.00505 | 0.0100024 | yes |
| 0.00485 | 0.00962177 | yes |
| 0.00905 | 0.0170729 | yes |
| 0.0198 | 0.0350165 | yes |
| 0.00495 | 0.00981225 | yes |
| 0.0053 | 0.0104304 | yes |
| 0.0274 | 0.0468734 | yes |
| 0.02295 | 0.0399272 | yes |
| 0.00605 | 0.0117652 | yes |
| 0.00705 | 0.0135598 | yes |
| 0.00645 | 0.0124937 | yes |
| 0.0063 | 0.0122128 | yes |
| 0.01275 | 0.0234242 | yes |
| 0.0068 | 0.01312 | yes |
| 0.0069 | 0.0132817 | yes |
| 0.00685 | 0.0131958 | yes |
| 0.0125 | 0.0229993 | yes |
| 0.01305 | 0.0238684 | yes |
| 0.00755 | 0.0144312 | yes |
| 0.0084 | 0.0159322 | yes |
| 0.008 | 0.0152322 | yes |
| 0.01915 | 0.0339892 | yes |
| 0.02145 | 0.0375833 | yes |
| 0.0079 | 0.0150885 | yes |
| 0.0071 | 0.0136453 | yes |
| 0.0166 | 0.0298509 | yes |
| 0.0073 | 0.0139642 | yes |
| 0.01635 | 0.0294228 | yes |
| 0.011 | 0.0205311 | yes |

| | | |
|---|---|---|
| 0.0215 | 0.0376441 | yes |
| 0.0083 | 0.0157546 | yes |
| 0.0158 | 0.0284747 | yes |
| 0.0089 | 0.0168157 | yes |
| 0.02405 | 0.0416055 | yes |
| 0.0221 | 0.0386122 | yes |
| 0.0096 | 0.0180689 | yes |
| 0.0098 | 0.0184031 | yes |
| 0.02 | 0.0352688 | yes |
| 0.01155 | 0.02146 | yes |
| 0.01675 | 0.0300986 | yes |
| 0.0196 | 0.0347378 | yes |
| 0.0115 | 0.0213832 | yes |
| 0.0112 | 0.0208569 | yes |
| 0.01235 | 0.0227573 | yes |
| 0.0109 | 0.0203599 | yes |
| 0.01285 | 0.0235727 | yes |
| 0.0123 | 0.0226822 | yes |
| 0.01415 | 0.0256893 | yes |
| 0.0179 | 0.0320247 | yes |
| 0.01755 | 0.0314672 | yes |
| 0.01765 | 0.0316235 | yes |
| 0.01825 | 0.0325091 | yes |
| 0.02935 | 0.049897 | yes |
| 0.02635 | 0.0452345 | yes |
| 0.0291 | 0.0495748 | yes |
| 0.0182 | 0.0324435 | yes |
| 0.0256 | 0.04407 | yes |
| 0.021 | 0.0368474 | yes |
| 0.01815 | 0.0323778 | yes |
| 0.02135 | 0.0374348 | yes |
| 0.02625 | 0.0451258 | yes |
| 0.0232 | 0.0402482 | yes |
| 0.02155 | 0.0377048 | yes |
| 0.0197 | 0.0348898 | yes |
| 0.023 | 0.0399859 | yes |
| 0.02305 | 0.0400445 | yes |
| 0.0255 | 0.0439594 | yes |
| 0.02515 | 0.0434168 | yes |
| 0.02595 | 0.0446413 | yes |
| 0.02885 | 0.049183 | yes |
| 0.0285 | 0.0486537 | yes |
| 0.0283 | 0.0483458 | yes |
| 0.02915 | 0.0496256 | yes |
| 0.02735 | 0.0468205 | yes |
| 0.0233 | 0.0403932 | yes |
| 0.02535 | 0.0437314 | yes |
| 0.0256 | 0.04407 | yes |
| 0.01975 | 0.0349532 | yes |
| 0.0203 | 0.035721 | yes |
| 0.0203 | 0.035721 | yes |

| | | |
|---|---|---|
| 0.02335 | 0.0404514 | yes |
| 0.01855 | 0.0330195 | yes |
| 0.02095 | 0.0368121 | yes |
| 0.01795 | 0.0320908 | yes |
| 0.0172 | 0.0308847 | yes |
| 0.0177 | 0.03169 | yes |
| 0.0288 | 0.0491318 | yes |
| 0.01465 | 0.0265383 | yes |
| 0.02245 | 0.0391403 | yes |
| 0.0146 | 0.0264672 | yes |
| 0.0127 | 0.0233498 | yes |
| 0.0161 | 0.0289941 | yes |
| 0.0199 | 0.0351176 | yes |
| 0.0105 | 0.0196426 | yes |
| 0.01935 | 0.0343194 | yes |
| 0.0138 | 0.0251653 | yes |
| 0.01335 | 0.0243628 | yes |
| 0.01065 | 0.0199081 | yes |
| 0.01025 | 0.0191895 | yes |
| 0.0086 | 0.0162738 | yes |
| 0.0132 | 0.0241248 | yes |
| 0.00975 | 0.0183231 | yes |
| 0.0101 | 0.0189231 | yes |
| 0.00825 | 0.0156839 | yes |
| 0.00895 | 0.0168972 | yes |
| 0.0073 | 0.0139642 | yes |
| 0.0067 | 0.0129474 | yes |
| 0.01175 | 0.0217822 | yes |
| 0.0157 | 0.028336 | yes |
| 0.0063 | 0.0122128 | yes |
| 0.0145 | 0.0263053 | yes |
| 0.0062 | 0.0120379 | yes |
| 0.00525 | 0.0103403 | yes |
| 0.00525 | 0.0103403 | yes |
| 0.00525 | 0.0103403 | yes |
| 0.0189 | 0.0335697 | yes |
| 0.0088 | 0.0166395 | yes |
| 0.0072 | 0.0138159 | yes |
| 0.00795 | 0.0151605 | yes |
| 0.00465 | 0.00923245 | yes |
| 0.00365 | 0.00739621 | yes |
| 0.00395 | 0.00793873 | yes |
| 0.00465 | 0.00923245 | yes |
| 0.00375 | 0.00757389 | yes |
| 0.008 | 0.0152322 | yes |
| 0.0029 | 0.00598992 | yes |
| 0.0036 | 0.00730091 | yes |
| 0.00565 | 0.0110485 | yes |
| 0.00305 | 0.0062525 | yes |
| 0.0052 | 0.0102747 | yes |
| 0.0022 | 0.00464948 | yes |

| | | |
|---|---|---|
| 0.003 | 0.00617057 | yes |
| 0.00615 | 0.0119502 | yes |
| 0.003 | 0.00617057 | yes |
| 0.00205 | 0.00437002 | yes |
| 0.00155 | 0.00336243 | yes |
| 0.00255 | 0.00532965 | yes |
| 0.0012 | 0.00266186 | yes |
| 0.0041 | 0.0082 | yes |
| 0.0024 | 0.00502896 | yes |
| 0.0019 | 0.00407498 | yes |
| 0.0022 | 0.00464948 | yes |
| 0.00115 | 0.00256018 | yes |
| 0.0011 | 0.00246224 | yes |
| 0.00115 | 0.00256018 | yes |
| 0.0015 | 0.00327128 | yes |
| 0.00135 | 0.00297048 | yes |
| 0.0021 | 0.00446114 | yes |
| 0.00205 | 0.00437002 | yes |
| 0.00145 | 0.00317914 | yes |
| 0.0017 | 0.00366842 | yes |
| 0.00075 | 0.00173402 | yes |
| 0.01125 | 0.0209342 | yes |
| 0.00135 | 0.00297048 | yes |
| 0.0008 | 0.00183411 | yes |
| 0.0017 | 0.00366842 | yes |
| 0.0245 | 0.0423542 | yes |
| 0.00385 | 0.00775676 | yes |
| 0.00395 | 0.00793873 | yes |
| 0.0009 | 0.00204621 | yes |
| 0.0023 | 0.00483177 | yes |
| 0.00085 | 0.0019415 | yes |
| 0.0012 | 0.00266186 | yes |
| 0.00115 | 0.00256018 | yes |
| 0.0006 | 0.00141244 | yes |
| 0.0008 | 0.00183411 | yes |
| 0.00115 | 0.00256018 | yes |
| 0.00045 | 0.0010885 | yes |
| 0.00105 | 0.0023589 | yes |
| 0.0151 | 0.0273132 | yes |
| 0.0005 | 0.00119766 | yes |
| 0.0007 | 0.00162914 | yes |
| 0.0016 | 0.00346174 | yes |
| 0.0073 | 0.0139642 | yes |
| 0.01185 | 0.021918 | yes |
| 0.00095 | 0.0021559 | yes |
| 0.00055 | 0.00130598 | yes |
| 0.0008 | 0.00183411 | yes |
| 0.00045 | 0.0010885 | yes |
| 0.00085 | 0.0019415 | yes |
| 0.00055 | 0.00130598 | yes |
| 0.00045 | 0.0010885 | yes |

| | | |
|---|---|---|
| 0.00155 | 0.00336243 | yes |
| 0.0008 | 0.00183411 | yes |
| 0.0005 | 0.00119766 | yes |
| 0.027 | 0.046318 | yes |
| 0.00065 | 0.00151996 | yes |
| 0.0009 | 0.00204621 | yes |
| 0.0003 | 0.00074621 | yes |
| 0.00025 | 0.00063012 | yes |
| 0.00075 | 0.00173402 | yes |
| 0.0003 | 0.00074621 | yes |
| 0.0004 | 0.00097813 | yes |
| 0.00025 | 0.00063012 | yes |
| 0.00605 | 0.0117652 | yes |
| 0.0007 | 0.00162914 | yes |
| 0.0002 | 0.00051037 | yes |
| 0.0015 | 0.00327128 | yes |
| 0.0003 | 0.00074621 | yes |
| 0.00715 | 0.0137307 | yes |
| 0.0015 | 0.00327128 | yes |
| 0.0002 | 0.00051037 | yes |
| 0.00025 | 0.00063012 | yes |
| 5.00E-05 | 0.00013743 | yes |
| 0.00015 | 0.00038924 | yes |
| 0.00015 | 0.00038924 | yes |
| 0.0118 | 0.0218584 | yes |
| 0.00015 | 0.00038924 | yes |
| 0.00035 | 0.00086186 | yes |
| 5.00E-05 | 0.00013743 | yes |
| 0.00015 | 0.00038924 | yes |
| 0.00025 | 0.00063012 | yes |
| 0.0002 | 0.00051037 | yes |
| 0.0001 | 0.00026595 | yes |
| 0.0002 | 0.00051037 | yes |
| 0.0001 | 0.00026595 | yes |
| 0.00025 | 0.00063012 | yes |
| 0.00225 | 0.00474293 | yes |
| 0.00055 | 0.00130598 | yes |
| 0.0001 | 0.00026595 | yes |
| 0.00045 | 0.0010885 | yes |
| 5.00E-05 | 0.00013743 | yes |
| 0.0001 | 0.00026595 | yes |
| 0.0148 | 0.0267903 | yes |
| 0.00075 | 0.00173402 | yes |
| 0.00315 | 0.00644676 | yes |
| 5.00E-05 | 0.00013743 | yes |
| 5.00E-05 | 0.00013743 | yes |
| 0.00045 | 0.0010885 | yes |
| 5.00E-05 | 0.00013743 | yes |
| 5.00E-05 | 0.00013743 | yes |
| 5.00E-05 | 0.00013743 | yes |
| 0.0001 | 0.00026595 | yes |

| | | |
|---|---|---|
| 0.0086 | 0.0162738 | yes |
| 0.0002 | 0.00051037 | yes |
| 5.00E-05 | 0.00013743 | yes |
| 5.00E-05 | 0.00013743 | yes |
| 0.0001 | 0.00026595 | yes |
| 5.00E-05 | 0.00013743 | yes |
| 5.00E-05 | 0.00013743 | yes |
| 0.01575 | 0.0284054 | yes |
| 5.00E-05 | 0.00013743 | yes |
| 0.0023 | 0.00483177 | yes |
| 5.00E-05 | 0.00013743 | yes |
| 5.00E-05 | 0.00013743 | yes |
| 5.00E-05 | 0.00013743 | yes |
| 0.00065 | 0.00151996 | yes |
| 5.00E-05 | 0.00013743 | yes |
| 0.00045 | 0.0010885 | yes |
| 0.0224 | 0.0390809 | yes |
| 5.00E-05 | 0.00013743 | yes |
| 5.00E-05 | 0.00013743 | yes |
| 5.00E-05 | 0.00013743 | yes |
| 5.00E-05 | 0.00013743 | yes |
| 0.00105 | 0.0023589 | yes |
| 5.00E-05 | 0.00013743 | yes |
| 0.0001 | 0.00026595 | yes |
| 0.003 | 0.00617057 | yes |
| 0.00205 | 0.00437002 | yes |
| 5.00E-05 | 0.00013743 | yes |
| 5.00E-05 | 0.00013743 | yes |
| 0.0042 | 0.00839318 | yes |
| 0.00025 | 0.00063012 | yes |
| 5.00E-05 | 0.00013743 | yes |
| 5.00E-05 | 0.00013743 | yes |
| 5.00E-05 | 0.00013743 | yes |
| 0.0017 | 0.00366842 | yes |
| 0.0001 | 0.00026595 | yes |
| 5.00E-05 | 0.00013743 | yes |
| 0.0002 | 0.00051037 | yes |
| 5.00E-05 | 0.00013743 | yes |
| 0.0038 | 0.00766858 | yes |
| 0.01225 | 0.0226069 | yes |
| 0.0006 | 0.00141244 | yes |
| 0.0001 | 0.00026595 | yes |
| 5.00E-05 | 0.00013743 | yes |
| 5.00E-05 | 0.00013743 | yes |
| 5.00E-05 | 0.00013743 | yes |
| 0.0001 | 0.00026595 | yes |
| 0.00175 | 0.00376311 | yes |
| 5.00E-05 | 0.00013743 | yes |
| 5.00E-05 | 0.00013743 | yes |
| 5.00E-05 | 0.00013743 | yes |
| 5.00E-05 | 0.00013743 | yes |

| | | |
|---|---|---|
| 0.00405 | 0.00810659 | yes |
| 0.01005 | 0.0188437 | yes |
| 5.00E-05 | 0.00013743 | yes |
| 5.00E-05 | 0.00013743 | yes |
| 5.00E-05 | 0.00013743 | yes |
| 0.00675 | 0.0130338 | yes |
| 5.00E-05 | 0.00013743 | yes |
| 0.00335 | 0.006839 | yes |
| 5.00E-05 | 0.00013743 | yes |
| 0.0117 | 0.0217059 | yes |
| 5.00E-05 | 0.00013743 | yes |
| 5.00E-05 | 0.00013743 | yes |
| 5.00E-05 | 0.00013743 | yes |
| 5.00E-05 | 0.00013743 | yes |
| 0.00725 | 0.013901 | yes |
| 0.0016 | 0.00346174 | yes |
| 5.00E-05 | 0.00013743 | yes |
| 5.00E-05 | 0.00013743 | yes |
| 5.00E-05 | 0.00013743 | yes |
| 0.00025 | 0.00063012 | yes |
| 5.00E-05 | 0.00013743 | yes |
| 5.00E-05 | 0.00013743 | yes |
| 0.0095 | 0.0178943 | yes |
| 0.0129 | 0.0236468 | yes |
| 0.02015 | 0.0355079 | yes |
| 5.00E-05 | 0.00013743 | yes |
| 5.00E-05 | 0.00013743 | yes |
| 5.00E-05 | 0.00013743 | yes |
| 5.00E-05 | 0.00013743 | yes |
| 5.00E-05 | 0.00013743 | yes |
| 5.00E-05 | 0.00013743 | yes |
| 5.00E-05 | 0.00013743 | yes |
| 0.0228 | 0.0396943 | yes |
| 5.00E-05 | 0.00013743 | yes |
| 0.0001 | 0.00026595 | yes |
| 5.00E-05 | 0.00013743 | yes |
| 0.00265 | 0.0055199 | yes |
| 5.00E-05 | 0.00013743 | yes |
| 5.00E-05 | 0.00013743 | yes |
| 5.00E-05 | 0.00013743 | yes |
| 5.00E-05 | 0.00013743 | yes |
| 0.001 | 0.00225275 | yes |
| 5.00E-05 | 0.00013743 | yes |
| 5.00E-05 | 0.00013743 | yes |
| 5.00E-05 | 0.00013743 | yes |
| 5.00E-05 | 0.00013743 | yes |
| 5.00E-05 | 0.00013743 | yes |
| 5.00E-05 | 0.00013743 | yes |
| 5.00E-05 | 0.00013743 | yes |
| 0.00195 | 0.00417493 | yes |
| 5.00E-05 | 0.00013743 | yes |

| | | |
|---|---|---|
| 5.00E-05 | 0.00013743 | yes |
| 5.00E-05 | 0.00013743 | yes |
| 5.00E-05 | 0.00013743 | yes |
| 0.00295 | 0.00608809 | yes |
| 5.00E-05 | 0.00013743 | yes |
| 5.00E-05 | 0.00013743 | yes |
| 5.00E-05 | 0.00013743 | yes |
| 5.00E-05 | 0.00013743 | yes |
| 0.0001 | 0.00026595 | yes |
| 5.00E-05 | 0.00013743 | yes |
| 5.00E-05 | 0.00013743 | yes |
| 0.0005 | 0.00119766 | yes |
| 5.00E-05 | 0.00013743 | yes |
| 5.00E-05 | 0.00013743 | yes |
| 5.00E-05 | 0.00013743 | yes |
| 5.00E-05 | 0.00013743 | yes |
| 5.00E-05 | 0.00013743 | yes |
| 5.00E-05 | 0.00013743 | yes |
| 5.00E-05 | 0.00013743 | yes |
| 5.00E-05 | 0.00013743 | yes |
| 0.0005 | 0.00119766 | yes |
| 5.00E-05 | 0.00013743 | yes |
| 5.00E-05 | 0.00013743 | yes |
| 5.00E-05 | 0.00013743 | yes |
| 0.00395 | 0.00793873 | yes |
| 5.00E-05 | 0.00013743 | yes |
| 5.00E-05 | 0.00013743 | yes |
| 5.00E-05 | 0.00013743 | yes |
| 0.01215 | 0.0224392 | yes |
| 5.00E-05 | 0.00013743 | yes |
| 5.00E-05 | 0.00013743 | yes |
| 5.00E-05 | 0.00013743 | yes |
| 5.00E-05 | 0.00013743 | yes |
| 5.00E-05 | 0.00013743 | yes |
| 5.00E-05 | 0.00013743 | yes |
| 0.0199 | 0.0351176 | yes |
| 5.00E-05 | 0.00013743 | yes |
| 5.00E-05 | 0.00013743 | yes |
| 0.00035 | 0.00086186 | yes |
| 5.00E-05 | 0.00013743 | yes |
| 5.00E-05 | 0.00013743 | yes |
| 5.00E-05 | 0.00013743 | yes |
| 5.00E-05 | 0.00013743 | yes |
| 5.00E-05 | 0.00013743 | yes |
| 0.0001 | 0.00026595 | yes |
| 5.00E-05 | 0.00013743 | yes |
| 0.0013 | 0.00286816 | yes |
| 5.00E-05 | 0.00013743 | yes |
| 5.00E-05 | 0.00013743 | yes |
| 5.00E-05 | 0.00013743 | yes |
| 5.00E-05 | 0.00013743 | yes |

| | | |
|---|---|---|
| 5.00E-05 | 0.00013743 | yes |
| 5.00E-05 | 0.00013743 | yes |
| 0.0019 | 0.00407498 | yes |
| 0.0052 | 0.0102747 | yes |
| 5.00E-05 | 0.00013743 | yes |
| 5.00E-05 | 0.00013743 | yes |
| 5.00E-05 | 0.00013743 | yes |
| 5.00E-05 | 0.00013743 | yes |
| 0.00045 | 0.0010885 | yes |
| 5.00E-05 | 0.00013743 | yes |
| 0.00025 | 0.00063012 | yes |
| 5.00E-05 | 0.00013743 | yes |
| 5.00E-05 | 0.00013743 | yes |
| 5.00E-05 | 0.00013743 | yes |
| 5.00E-05 | 0.00013743 | yes |
| 0.0121 | 0.0223636 | yes |
| 5.00E-05 | 0.00013743 | yes |
| 5.00E-05 | 0.00013743 | yes |
| 5.00E-05 | 0.00013743 | yes |
| 5.00E-05 | 0.00013743 | yes |
| 5.00E-05 | 0.00013743 | yes |
| 0.0002 | 0.00051037 | yes |
| 5.00E-05 | 0.00013743 | yes |
| 0.00015 | 0.00038924 | yes |
| 5.00E-05 | 0.00013743 | yes |
| 5.00E-05 | 0.00013743 | yes |
| 0.00595 | 0.0115983 | yes |
| 5.00E-05 | 0.00013743 | yes |
| 5.00E-05 | 0.00013743 | yes |
| 5.00E-05 | 0.00013743 | yes |
| 5.00E-05 | 0.00013743 | yes |
| 5.00E-05 | 0.00013743 | yes |
| 5.00E-05 | 0.00013743 | yes |
| 0.0004 | 0.00097813 | yes |
| 0.00215 | 0.00455164 | yes |
| 5.00E-05 | 0.00013743 | yes |
| 5.00E-05 | 0.00013743 | yes |
| 5.00E-05 | 0.00013743 | yes |
| 5.00E-05 | 0.00013743 | yes |
| 0.0006 | 0.00141244 | yes |
| 0.00035 | 0.00086186 | yes |
| 5.00E-05 | 0.00013743 | yes |
| 5.00E-05 | 0.00013743 | yes |
| 5.00E-05 | 0.00013743 | yes |
| 5.00E-05 | 0.00013743 | yes |
| 5.00E-05 | 0.00013743 | yes |
| 5.00E-05 | 0.00013743 | yes |
| 5.00E-05 | 0.00013743 | yes |
| 5.00E-05 | 0.00013743 | yes |
| 5.00E-05 | 0.00013743 | yes |
| 5.00E-05 | 0.00013743 | yes |

| | | |
|---|---|---|
| 0.00325 | 0.00664037 | yes |
| 5.00E-05 | 0.00013743 | yes |
| 5.00E-05 | 0.00013743 | yes |
| 5.00E-05 | 0.00013743 | yes |
| 5.00E-05 | 0.00013743 | yes |
| 5.00E-05 | 0.00013743 | yes |
| 5.00E-05 | 0.00013743 | yes |
| 5.00E-05 | 0.00013743 | yes |
| 5.00E-05 | 0.00013743 | yes |
| 5.00E-05 | 0.00013743 | yes |
| 5.00E-05 | 0.00013743 | yes |
| 5.00E-05 | 0.00013743 | yes |
| 5.00E-05 | 0.00013743 | yes |
| 5.00E-05 | 0.00013743 | yes |
| 0.00525 | 0.0103403 | yes |
| 5.00E-05 | 0.00013743 | yes |
| 5.00E-05 | 0.00013743 | yes |
| 5.00E-05 | 0.00013743 | yes |
| 0.00045 | 0.0010885 | yes |
| 0.0003 | 0.00074621 | yes |
| 5.00E-05 | 0.00013743 | yes |
| 5.00E-05 | 0.00013743 | yes |
| 5.00E-05 | 0.00013743 | yes |
| 5.00E-05 | 0.00013743 | yes |
| 5.00E-05 | 0.00013743 | yes |
| 5.00E-05 | 0.00013743 | yes |
| 5.00E-05 | 0.00013743 | yes |
| 5.00E-05 | 0.00013743 | yes |
| 5.00E-05 | 0.00013743 | yes |
| 5.00E-05 | 0.00013743 | yes |
| 5.00E-05 | 0.00013743 | yes |
| 5.00E-05 | 0.00013743 | yes |
| 5.00E-05 | 0.00013743 | yes |
| 5.00E-05 | 0.00013743 | yes |
| 5.00E-05 | 0.00013743 | yes |
| 5.00E-05 | 0.00013743 | yes |
| 0.0002 | 0.00051037 | yes |
| 5.00E-05 | 0.00013743 | yes |
| 0.01305 | 0.0238684 | yes |
| 5.00E-05 | 0.00013743 | yes |
| 5.00E-05 | 0.00013743 | yes |
| 5.00E-05 | 0.00013743 | yes |
| 5.00E-05 | 0.00013743 | yes |
| 5.00E-05 | 0.00013743 | yes |
| 0.0085 | 0.0161094 | yes |
| 5.00E-05 | 0.00013743 | yes |
| 5.00E-05 | 0.00013743 | yes |
| 5.00E-05 | 0.00013743 | yes |
| 5.00E-05 | 0.00013743 | yes |
| 5.00E-05 | 0.00013743 | yes |
| 5.00E-05 | 0.00013743 | yes |

| | | |
|---|---|---|
| 5.00E-05 | 0.00013743 | yes |
| 5.00E-05 | 0.00013743 | yes |
| 5.00E-05 | 0.00013743 | yes |
| 5.00E-05 | 0.00013743 | yes |
| 5.00E-05 | 0.00013743 | yes |
| 5.00E-05 | 0.00013743 | yes |
| 0.00055 | 0.00130598 | yes |
| 5.00E-05 | 0.00013743 | yes |
| 5.00E-05 | 0.00013743 | yes |
| 5.00E-05 | 0.00013743 | yes |
| 5.00E-05 | 0.00013743 | yes |
| 5.00E-05 | 0.00013743 | yes |
| 5.00E-05 | 0.00013743 | yes |
| 5.00E-05 | 0.00013743 | yes |
| 5.00E-05 | 0.00013743 | yes |
| 5.00E-05 | 0.00013743 | yes |
| 5.00E-05 | 0.00013743 | yes |
| 0.0015 | 0.00327128 | yes |
| 5.00E-05 | 0.00013743 | yes |
| 5.00E-05 | 0.00013743 | yes |
| 5.00E-05 | 0.00013743 | yes |
| 5.00E-05 | 0.00013743 | yes |
| 5.00E-05 | 0.00013743 | yes |
| 5.00E-05 | 0.00013743 | yes |
| 5.00E-05 | 0.00013743 | yes |
| 5.00E-05 | 0.00013743 | yes |
| 5.00E-05 | 0.00013743 | yes |
| 0.0001 | 0.00026595 | yes |
| 5.00E-05 | 0.00013743 | yes |
| 5.00E-05 | 0.00013743 | yes |
| 5.00E-05 | 0.00013743 | yes |
| 0.00015 | 0.00038924 | yes |
| 0.0009 | 0.00204621 | yes |
| 5.00E-05 | 0.00013743 | yes |
| 5.00E-05 | 0.00013743 | yes |
| 5.00E-05 | 0.00013743 | yes |
| 5.00E-05 | 0.00013743 | yes |
| 5.00E-05 | 0.00013743 | yes |
| 5.00E-05 | 0.00013743 | yes |
| 0.0003 | 0.00074621 | yes |
| 0.00035 | 0.00086186 | yes |
| 5.00E-05 | 0.00013743 | yes |
| 5.00E-05 | 0.00013743 | yes |
| 5.00E-05 | 0.00013743 | yes |
| 5.00E-05 | 0.00013743 | yes |
| 5.00E-05 | 0.00013743 | yes |
| 5.00E-05 | 0.00013743 | yes |
| 5.00E-05 | 0.00013743 | yes |
| 5.00E-05 | 0.00013743 | yes |
| 5.00E-05 | 0.00013743 | yes |
| 5.00E-05 | 0.00013743 | yes |

| | | |
|---|---|---|
| 5.00E-05 | 0.00013743 | yes |
| 5.00E-05 | 0.00013743 | yes |
| 0.00155 | 0.00336243 | yes |
| 5.00E-05 | 0.00013743 | yes |
| 5.00E-05 | 0.00013743 | yes |
| 5.00E-05 | 0.00013743 | yes |
| 5.00E-05 | 0.00013743 | yes |
| 5.00E-05 | 0.00013743 | yes |
| 5.00E-05 | 0.00013743 | yes |
| 5.00E-05 | 0.00013743 | yes |
| 5.00E-05 | 0.00013743 | yes |
| 5.00E-05 | 0.00013743 | yes |
| 5.00E-05 | 0.00013743 | yes |
| 0.00425 | 0.0084862 | yes |
| 0.00015 | 0.00038924 | yes |
| 5.00E-05 | 0.00013743 | yes |
| 0.0055 | 0.0107809 | yes |
| 5.00E-05 | 0.00013743 | yes |
| 5.00E-05 | 0.00013743 | yes |
| 5.00E-05 | 0.00013743 | yes |
| 5.00E-05 | 0.00013743 | yes |
| 5.00E-05 | 0.00013743 | yes |
| 5.00E-05 | 0.00013743 | yes |
| 5.00E-05 | 0.00013743 | yes |
| 5.00E-05 | 0.00013743 | yes |
| 5.00E-05 | 0.00013743 | yes |
| 5.00E-05 | 0.00013743 | yes |
| 5.00E-05 | 0.00013743 | yes |
| 5.00E-05 | 0.00013743 | yes |
| 5.00E-05 | 0.00013743 | yes |
| 5.00E-05 | 0.00013743 | yes |
| 5.00E-05 | 0.00013743 | yes |
| 5.00E-05 | 0.00013743 | yes |
| 5.00E-05 | 0.00013743 | yes |
| 5.00E-05 | 0.00013743 | yes |
| 5.00E-05 | 0.00013743 | yes |
| 5.00E-05 | 0.00013743 | yes |
| 5.00E-05 | 0.00013743 | yes |
| 0.0001 | 0.00026595 | yes |
| 5.00E-05 | 0.00013743 | yes |
| 5.00E-05 | 0.00013743 | yes |
| 5.00E-05 | 0.00013743 | yes |
| 5.00E-05 | 0.00013743 | yes |
| 5.00E-05 | 0.00013743 | yes |
| 5.00E-05 | 0.00013743 | yes |
| 5.00E-05 | 0.00013743 | yes |
| 5.00E-05 | 0.00013743 | yes |
| 5.00E-05 | 0.00013743 | yes |
| 5.00E-05 | 0.00013743 | yes |
| 5.00E-05 | 0.00013743 | yes |
| 5.00E-05 | 0.00013743 | yes |

| | | |
|---|---|---|
| 5.00E-05 | 0.00013743 | yes |
| 5.00E-05 | 0.00013743 | yes |
| 5.00E-05 | 0.00013743 | yes |
| 5.00E-05 | 0.00013743 | yes |
| 5.00E-05 | 0.00013743 | yes |
| 5.00E-05 | 0.00013743 | yes |
| 5.00E-05 | 0.00013743 | yes |
| 5.00E-05 | 0.00013743 | yes |
| 5.00E-05 | 0.00013743 | yes |
| 5.00E-05 | 0.00013743 | yes |
| 0.0051 | 0.0100933 | yes |
| 5.00E-05 | 0.00013743 | yes |
| 5.00E-05 | 0.00013743 | yes |
| 5.00E-05 | 0.00013743 | yes |
| 5.00E-05 | 0.00013743 | yes |
| 5.00E-05 | 0.00013743 | yes |
| 5.00E-05 | 0.00013743 | yes |
| 5.00E-05 | 0.00013743 | yes |
| 5.00E-05 | 0.00013743 | yes |
| 5.00E-05 | 0.00013743 | yes |
| 5.00E-05 | 0.00013743 | yes |
| 5.00E-05 | 0.00013743 | yes |
| 5.00E-05 | 0.00013743 | yes |
| 5.00E-05 | 0.00013743 | yes |
| 5.00E-05 | 0.00013743 | yes |
| 5.00E-05 | 0.00013743 | yes |
| 5.00E-05 | 0.00013743 | yes |
| 5.00E-05 | 0.00013743 | yes |
| 5.00E-05 | 0.00013743 | yes |
| 5.00E-05 | 0.00013743 | yes |
| 5.00E-05 | 0.00013743 | yes |
| 5.00E-05 | 0.00013743 | yes |
| 5.00E-05 | 0.00013743 | yes |
| 5.00E-05 | 0.00013743 | yes |
| 5.00E-05 | 0.00013743 | yes |
| 5.00E-05 | 0.00013743 | yes |
| 5.00E-05 | 0.00013743 | yes |
| 5.00E-05 | 0.00013743 | yes |
| 5.00E-05 | 0.00013743 | yes |
| 5.00E-05 | 0.00013743 | yes |
| 5.00E-05 | 0.00013743 | yes |
| 5.00E-05 | 0.00013743 | yes |
| 5.00E-05 | 0.00013743 | yes |
| 0.00375 | 0.00757389 | yes |
| 5.00E-05 | 0.00013743 | yes |
| 5.00E-05 | 0.00013743 | yes |
| 5.00E-05 | 0.00013743 | yes |
| 5.00E-05 | 0.00013743 | yes |
| 5.00E-05 | 0.00013743 | yes |
| 5.00E-05 | 0.00013743 | yes |
| 5.00E-05 | 0.00013743 | yes |

| | | |
|---|---|---|
| 0.0012 | 0.00266186 | yes |
| 0.0004 | 0.00097813 | yes |
| 5.00E-05 | 0.00013743 | yes |
| 5.00E-05 | 0.00013743 | yes |
| 5.00E-05 | 0.00013743 | yes |
| 5.00E-05 | 0.00013743 | yes |
| 5.00E-05 | 0.00013743 | yes |
| 5.00E-05 | 0.00013743 | yes |
| 5.00E-05 | 0.00013743 | yes |
| 0.00115 | 0.00256018 | yes |
| 5.00E-05 | 0.00013743 | yes |
| 5.00E-05 | 0.00013743 | yes |
| 5.00E-05 | 0.00013743 | yes |
| 5.00E-05 | 0.00013743 | yes |
| 5.00E-05 | 0.00013743 | yes |
| 5.00E-05 | 0.00013743 | yes |
| 5.00E-05 | 0.00013743 | yes |
| 5.00E-05 | 0.00013743 | yes |
| 5.00E-05 | 0.00013743 | yes |
| 5.00E-05 | 0.00013743 | yes |
| 5.00E-05 | 0.00013743 | yes |
| 5.00E-05 | 0.00013743 | yes |
| 5.00E-05 | 0.00013743 | yes |
| 5.00E-05 | 0.00013743 | yes |
| 5.00E-05 | 0.00013743 | yes |
| 5.00E-05 | 0.00013743 | yes |
| 5.00E-05 | 0.00013743 | yes |
| 5.00E-05 | 0.00013743 | yes |
| 5.00E-05 | 0.00013743 | yes |
| 5.00E-05 | 0.00013743 | yes |
| 0.0003 | 0.00074621 | yes |
| 5.00E-05 | 0.00013743 | yes |
| 5.00E-05 | 0.00013743 | yes |
| 5.00E-05 | 0.00013743 | yes |
| 5.00E-05 | 0.00013743 | yes |
| 5.00E-05 | 0.00013743 | yes |
| 5.00E-05 | 0.00013743 | yes |
| 5.00E-05 | 0.00013743 | yes |
| 5.00E-05 | 0.00013743 | yes |
| 5.00E-05 | 0.00013743 | yes |
| 5.00E-05 | 0.00013743 | yes |
| 5.00E-05 | 0.00013743 | yes |
| 5.00E-05 | 0.00013743 | yes |
| 5.00E-05 | 0.00013743 | yes |
| 5.00E-05 | 0.00013743 | yes |
| 5.00E-05 | 0.00013743 | yes |
| 5.00E-05 | 0.00013743 | yes |
| 5.00E-05 | 0.00013743 | yes |
| 5.00E-05 | 0.00013743 | yes |
| 5.00E-05 | 0.00013743 | yes |
| 5.00E-05 | 0.00013743 | yes |

| | | |
|---|---|---|
| 5.00E-05 | 0.00013743 | yes |
| 5.00E-05 | 0.00013743 | yes |
| 5.00E-05 | 0.00013743 | yes |
| 5.00E-05 | 0.00013743 | yes |
| 5.00E-05 | 0.00013743 | yes |
| 5.00E-05 | 0.00013743 | yes |
| 5.00E-05 | 0.00013743 | yes |
| 5.00E-05 | 0.00013743 | yes |
| 0.0271 | 0.0464571 | yes |
| 5.00E-05 | 0.00013743 | yes |
| 5.00E-05 | 0.00013743 | yes |
| 5.00E-05 | 0.00013743 | yes |
| 5.00E-05 | 0.00013743 | yes |
| 5.00E-05 | 0.00013743 | yes |
| 5.00E-05 | 0.00013743 | yes |
| 5.00E-05 | 0.00013743 | yes |
| 5.00E-05 | 0.00013743 | yes |
| 5.00E-05 | 0.00013743 | yes |
| 0.0011 | 0.00246224 | yes |
| 5.00E-05 | 0.00013743 | yes |
| 5.00E-05 | 0.00013743 | yes |
| 5.00E-05 | 0.00013743 | yes |
| 5.00E-05 | 0.00013743 | yes |
| 5.00E-05 | 0.00013743 | yes |
| 5.00E-05 | 0.00013743 | yes |
| 5.00E-05 | 0.00013743 | yes |
| 5.00E-05 | 0.00013743 | yes |
| 5.00E-05 | 0.00013743 | yes |
| 5.00E-05 | 0.00013743 | yes |
| 5.00E-05 | 0.00013743 | yes |
| 5.00E-05 | 0.00013743 | yes |
| 5.00E-05 | 0.00013743 | yes |
| 5.00E-05 | 0.00013743 | yes |
| 5.00E-05 | 0.00013743 | yes |
| 5.00E-05 | 0.00013743 | yes |
| 5.00E-05 | 0.00013743 | yes |
| 5.00E-05 | 0.00013743 | yes |
| 5.00E-05 | 0.00013743 | yes |
| 5.00E-05 | 0.00013743 | yes |
| 5.00E-05 | 0.00013743 | yes |
| 5.00E-05 | 0.00013743 | yes |
| 5.00E-05 | 0.00013743 | yes |
| 5.00E-05 | 0.00013743 | yes |
| 5.00E-05 | 0.00013743 | yes |
| 0.00155 | 0.00336243 | yes |
| 5.00E-05 | 0.00013743 | yes |
| 5.00E-05 | 0.00013743 | yes |
| 5.00E-05 | 0.00013743 | yes |
| 5.00E-05 | 0.00013743 | yes |
| 5.00E-05 | 0.00013743 | yes |
| 5.00E-05 | 0.00013743 | yes |

| | | |
|---|---|---|
| 5.00E-05 | 0.00013743 | yes |
| 5.00E-05 | 0.00013743 | yes |
| 5.00E-05 | 0.00013743 | yes |
| 5.00E-05 | 0.00013743 | yes |
| 5.00E-05 | 0.00013743 | yes |
| 5.00E-05 | 0.00013743 | yes |
| 5.00E-05 | 0.00013743 | yes |
| 5.00E-05 | 0.00013743 | yes |
| 5.00E-05 | 0.00013743 | yes |
| 5.00E-05 | 0.00013743 | yes |
| 5.00E-05 | 0.00013743 | yes |
| 5.00E-05 | 0.00013743 | yes |
| 5.00E-05 | 0.00013743 | yes |
| 5.00E-05 | 0.00013743 | yes |
| 5.00E-05 | 0.00013743 | yes |
| 5.00E-05 | 0.00013743 | yes |
| 5.00E-05 | 0.00013743 | yes |
| 5.00E-05 | 0.00013743 | yes |
| 5.00E-05 | 0.00013743 | yes |
| 5.00E-05 | 0.00013743 | yes |
| 5.00E-05 | 0.00013743 | yes |

| | |
|---|---|
| protein A | spa |
| | hypothetical protein |
| superoxide dismutase | sodM |
| | hypothetical protein |
| 50S ribosomal protein L29 | rpmC |
| | hypothetical protein |
| secretory antigen | ssaA |
| DNA-directed RNA polymerase subunit alpha | rpoA |
| pyridoxal biosynthesis lyase PdxS | pdxS |
| 30S ribosomal protein S11 | rpsK |
| | membrane spanning protein |
| immunodominant antigen A | isaA |
| accessory regulator-like protein | sarS |
| glutamine amidotransferase subunit PdxT | pdxT |
| D-lactate dehydrogenase | ddh |
| | drug transporter |
| 30S ribosomal protein S10 | rpsJ |
| | N-acetylmuramoyl-L-alanine amidase |
| hypothetical protein | nnrD |
| bacteriophage L54a single-stranded DNA binding proteir | ssb |
| riboflavin biosynthesis protein | ribA |
| riboflavin synthase subunit alpha | ribB |
| ferredoxin | fer |
| hypothetical protein | hypothetical protein |
| hypothetical protein | hypothetical protein |
| hypothetical protein | hypothetical protein |
| hypothetical protein | cidA |
| pyruvate oxidase | cidC |
| hypothetical protein | hypothetical protein |
| GTP-binding protein TypA | typA |
| cytochrome d ubiquinol oxidase subunit I | cydA |
| hypothetical protein | hypothetical protein |
| | transporter |
| hypothetical protein | hypothetical protein |
| hypothetical protein | cidB |
| hypothetical protein | hypothetical protein |
| riboflavin biosynthesis protein RibD | ribD |
| 3-oxoacyl-ACP synthase III | fabH |
| hypothetical protein | hypothetical protein |
| DNA-directed RNA polymerase subunit beta' | rpoC |
| hypothetical protein | farE |
| hypothetical protein | hypothetical protein |
| hypothetical protein | hypothetical protein |
| glutamate synthase large subunit | gltB |
| integrase/recombinase core subunit | integrase/recombinase core subunit |
| hypothetical protein | tnp3 |
| hypothetical protein | czrA |
| threonine dehydratase | ilvA1 |
| aminobenzoyl-glutamate utilization protein B | hmrA |

| | |
|---|---|
| Ars family arsenical pump | arsB |
| hypothetical protein | hypothetical protein |
| hypothetical protein | fhuD2 |
| choline transporter | cudT |
| clumping factor B | clfB |
| xanthine permease | pbuX |
| alanine dehydrogenase | ald1 |
| alanine dehydrogenase | ald1 |
| | ABC transporter ATP-binding protein |
| DNA-directed RNA polymerase subunit delta | rpoE |
| phosphate transporter ATP-binding protein | pstB |
| N-glycosyltransferase | icaA |
| xanthine phosphoribosyltransferase | xpt |
| | tnp4 |
| cation efflux family protein | czrB |
| autolysin | sle1 |
| hypothetical protein | hypothetical protein |
| hypothetical protein | hypothetical protein |
| 30S ribosomal protein S7 | rpsG |
| hypothetical protein | hypothetical protein |
| ABC transporter permease | glnP |
| hypothetical protein | hypothetical protein |
| guanosine 5'-monophosphate oxidoreductase | guaC |
| hypothetical protein | hypothetical protein |
| S-adenosylmethionine synthetase | metK |
| hypothetical protein | hypothetical protein |
| hypothetical protein | hypothetical protein |
| hypothetical protein | hypothetical protein |
| | sstC |
| fibrinogen-binding protein SdrD | sdrD |
| cytochrome d ubiquinol oxidase subunit II | cydB |
| hypothetical protein | hypothetical protein |
| hypothetical protein | hypothetical protein |
| hypothetical protein | hypothetical protein |
| phosphate ABC transporter permease | pstA |
| thiamine biosynthesis protein ThiI | thiI |
| hypothetical protein | hypothetical protein |
| sodium:alanine symporter family protein | alsT |
| hypothetical protein | pstC |
| 1-(5-phosphoribosyl)-5-[(5- phosphoribosylamino)methy | hisA |
| spermidine/putrescine ABC transporter substrate-bindin | potD |
| hypothetical protein | hypothetical protein |
| 5-(carboxyamino)imidazole ribonucleotide mutase | purE |
| hypothetical protein | mhqD |
| DNA-directed RNA polymerase subunit beta | rpoB |
| hypothetical protein | hypothetical protein |
| hypothetical protein | hypothetical protein |
| arsenate operon regulator | arsR |

| | |
|---|---|
| DNA topoisomerase I | topA |
| arsenate reductase | arsC |
| superantigen-like protein | ssl6 |
| hypothetical protein | hypothetical protein |
| hypothetical protein | salA |
| hypothetical protein | rsmC |
| 50S ribosomal protein L33 | rpmG |
| hypothetical protein | hypothetical protein |
| hypothetical protein | hypothetical protein |
| hypothetical protein | sufT |
| imidazole glycerol phosphate synthase subunit HisH | hisH |
| hypothetical protein | iscS |
| EmrB/QacA family drug resistance transporter | emrB |
| proline/betaine transporter | proP |
| UDP-N-acetylmuramoylalanyl-D-glutamate--L-lysine liga | murE |
| DEAD-box ATP dependent DNA helicase | cshA |
| hypothetical protein | hypothetical protein |
| PhoU family transcriptional regulator | phoU |
| hypothetical protein | hypothetical protein |
| hypothetical protein | hypothetical protein |
| | drug transporter |
| hypothetical protein | hypothetical protein |
| NAD(P)H-flavin oxidoreductase | frp |
| hypothetical protein | hypothetical protein |
| glutamine synthetase repressor | glnR |
| hypothetical protein | hypothetical protein |
| 50S ribosomal protein L25/general stress protein Ctc | rplY |
| hypothetical protein | hypothetical protein |
| hypothetical protein | hypothetical protein |
| high affinity proline permease | putP |
| thermonuclease | nuc2 |
| hypothetical protein | hypothetical protein |
| hypothetical protein | sstD |
| hypothetical protein | hypothetical protein |
| hypothetical protein | hypothetical protein |
| hypothetical protein | lyrA |
| 5'-methylthioadenosine/S-adenosylhomocysteine nucleo | mtnN |
| sodium/glutamate symporter | gltS |
| ParB family chromosome partioning protein | parB |
| hypothetical protein | mhqE |
| hypothetical protein | metQ |
| ABC transporter ATP-binding protein | metN |
| hypothetical protein | hypothetical protein |

| | |
|---|---|
| ABC transporter | potA |
| 7-cyano-7-deazaguanine synthase QueC | queC |
| hypothetical protein | ktrA |
| hypothetical protein | hypothetical protein |
| hypothetical protein | hypothetical protein |
| hypothetical protein | hypothetical protein |
| hypothetical protein | hypothetical protein |
| 50S ribosomal protein L21 | rplU |
| ATP phosphoribosyltransferase regulatory subunit | hisZ |
| hypothetical protein | thiW |
| | hypothetical protein |
| GTP-binding protein YqeH | GTP-binding protein YqeH |
| hypothetical protein | thiX |
| tRNA modification GTPase TrmE | trmE |
| histidinol-phosphate aminotransferase | histidinol-phosphate aminotransferase |
| hypothetical protein | mnmA |
| spermidine/putrescine ABC transporter permease | potC |
| 6-pyruvoyl tetrahydropterin synthase | queD |
| tRNA uridine 5-carboxymethylaminomethyl modificatior | gidA |
| secretory antigen SsaA | secretory antigen SsaA |
| hypothetical protein | hypothetical protein |
| hypothetical protein | hypothetical protein |
| hypothetical protein | hypothetical protein |
| peptide chain release factor 2 | prfB |
| phage terminase small subunit | phage terminase small subunit |
| hypothetical protein | trmK |
| hypothetical protein | hypothetical protein |
| 23S rRNA (pseudouridine(1915)-N(3))-methyltransferase | orfX |
| F0F1 ATP synthase subunit beta | atpD |
| hypothetical protein | hypothetical protein |
| A/G-specific adenine glycosylase | mutY |
| pyrazinamidase/nicotinamidase | pncA |
| hypothetical protein | hypothetical protein |
| oligopeptide ABC transporter substrate-binding protein | opp-4A |
| hypothetical protein | rpmF |
| 7-cyano-7-deazaguanosine (preQ0) biosynthesis protein | queE |
| hypothetical protein | hypothetical protein |
| hypothetical protein | hypothetical protein |
| hypothetical protein | hypothetical protein |
| peptidoglycan hydrolase | lytM |
| hypothetical protein | hypothetical protein |
| multifunctional imidazole glycerol phosphate synthase su | hisF |
| phage infection protein | phage infection protein |
| hypothetical protein | hypothetical protein |
| 16S rRNA methyltransferase GidB | gidB |
| guanylate kinase | gmk |
| hypothetical protein | hypothetical protein |

| | |
|---|---|
| hypothetical protein | hypothetical protein |
| penicillin-binding protein 4 | pbp4 |
| hypothetical protein | hypothetical protein |
| F0F1 ATP synthase subunit epsilon | atpC |
| hypothetical protein | bshB2 |
| hypothetical protein | hypothetical protein |
| ABC transporter ATP-binding/permease | ABC transporter ATP-binding/permease |
| transcription termination factor Rho | rho |
| redox-sensing transcriptional repressor Rex | rex |
| hypothetical protein | hypothetical protein |
| hypothetical protein | nfu |
| aspartyl-tRNA synthetase | aspS |
| glycolytic operon regulator | cggR |
| amino acid ABC transporter ATP-binding protein | glnQ |
| cytidylate kinase | cmk |
| betaine aldehyde dehydrogenase | betB |
| superoxide dismutase | sodA |
| 16S rRNA (uracil(1498)-N(3))-methyltransferase | rsmE |
| spermidine/putrescine ABC transporter permease | potB |
| PTS system transporter subunit IIC domain-containing p | PTS system transporter subunit IIC dom |
| uridine kinase | udk |
| hypothetical protein | comK2 |
| hypothetical protein | hypothetical protein |
| hypothetical protein | metP |
| superantigen-like protein | ssl9 |
| hypothetical protein | hypothetical protein |
| hypothetical protein | metI |
| inosine-5'-monophosphate dehydrogenase | guaB |
| 6-phosphofructokinase | pfkA |
| MerR family transcriptional regulator | MerR family transcriptional regulator |
| hypothetical protein | hypothetical protein |
| hypothetical protein | hypothetical protein |
| GMP synthase | guaA |
| 3-methyl-2-oxobutanoate hydroxymethyltransferase | panB |
| hypothetical protein | yidC |
| acetolactate synthase | budB |
| 50S ribosomal protein L27 | rpmA |
| hypothetical protein | sstB |
| hypothetical protein | nptA |
| hypothetical protein | hypothetical protein |
| formate dehydrogenase subunit alpha | fdhA |
| ribosomal biogenesis GTPase | rbgA |
| hypothetical protein | hypothetical protein |
| transcription elongation factor GreA | greA |
| bifunctional autolysin | atl |
| hypothetical protein | mpsB |
| GTP cyclohydrolase | folE2 |
| RNA polymerase sigma factor RpoD | sigA |
| hypothetical protein | hypothetical protein |
| hypothetical protein | hypothetical protein |
| hypothetical protein | mtaB |

| | |
|---|---|
| DNA topoisomerase IV subunit A | parC |
| F0F1 ATP synthase subunit C | atpE |
| pyruvate dehydrogenase complex, E1 component subu | pdhA |
| ATP-dependent Clp protease ATP-binding subunit ClpB | clpB |
| hypothetical protein | cspR |
| hypothetical protein | hypothetical protein |
| hypothetical protein | hypothetical protein |
| cytidylate kinase | cmk |
| hypothetical protein | hypothetical protein |
| hypothetical protein | hypothetical protein |
| large-conductance mechanosensitive channel | large-conductance mechanosensitive ch |
| aminopeptidase PepS | ampS |
| hypothetical protein | hypothetical protein |
| hypothetical protein | esxD |
| glycerol-3-phosphate acyltransferase PlsX | plsX |
| hypothetical protein | thiV |
| 3-oxoacyl-(acyl-carrier-protein) reductase | fabG |
| hypothetical protein | queG |
| hypothetical protein | pitA |
| F0F1 ATP synthase subunit gamma | atpG |
| nicotinate (nicotinamide) nucleotide adenylyltransferase | nadD |
| 5-(carboxyamino)imidazole ribonucleotide synthase | purK |
| : co-chaperonin GroES | groES |
| hypothetical protein | hypothetical protein |
| hypothetical protein | hypothetical protein |
| hypothetical protein | hypothetical protein |
| hypothetical protein | hypothetical protein |
| 50S ribosomal protein L7 serine acetyltransferase | 50S ribosomal protein L7 serine acetyltra |
| lysyl-tRNA synthetase | lysS |
| hypothetical protein | hypothetical protein |
| ribonuclease HII | rnhB |
| glucose induced biofilm accessory gene | gbaA |
| GTP-binding protein EngA | engA |
| hypothetical protein | hypothetical protein |
| hypothetical protein | esxB |
| nitrate reductase subunit beta | narH |
| hypothetical protein | pitR |
| glycyl-tRNA synthetase | glyS |
| hypothetical protein | hypothetical protein |
| leucyl-tRNA synthetase | leuS |
| DNA gyrase subunit B | gyrB |
| hypothetical protein | ywpF |
| hypothetical protein | patA |
| DNA topoisomerase IV subunit B | parE |
| heat shock protein GrpE | grpE |
| 3-oxoacyl- synthase | fabF |
| glycine betaine transporter | opuD1 |
| NAD(P)H-dependent glycerol-3-phosphate dehydrogena | gpsA |
| alpha-acetolactate decarboxylase | budA2 |
| ribosome biogenesis GTP-binding protein YsxC | engB |
| LysM domain-containing protein | LysM domain-containing protein |

| | |
|---|---|
| histidinol-phosphate aminotransferase | hisC |
| 2-oxoglutarate ferredoxin oxidoreductase subunit beta | 2-oxoglutarate ferredoxin oxidoreductase |
| hypothetical protein | hypothetical protein |
| acetyl-CoA carboxylase biotin carboxyl carrier protein su | acetyl-CoA carboxylase biotin carboxyl c |
| shikimate kinase | aroK |
| hypothetical protein | hypothetical protein |
| malonyl CoA-acyl carrier protein transacylase | fabD |
| fibronectin binding protein B | fnbB |
| chaperonin GroEL | groEL |
| hypothetical protein | hypothetical protein |
| phosphomethylpyrimidine kinase | pdxK |
| UDP-N-acetylmuramate--L-alanine ligase | murC |
| peptide chain release factor 1 | prfA |
| 30S ribosomal protein S20 | rpsT |
| hypothetical protein | hypothetical protein |
| tryptophanyl-tRNA synthetase | trpS |
| sugar transporter | glcU |
| carboxylesterase | est |
| hypothetical protein | tcdA |
| pyruvate dehydrogenase complex, E1 component subur | pdhB |
| hypothetical protein | cvfC |
| hypothetical protein | hrtA |
| choline dehydrogenase | betA |
| molecular chaperone DnaK | dnaK |
| hypothetical protein | hypothetical protein |
| hypothetical protein | hypothetical protein |
| hypothetical protein | cymR |
| DNA polymerase III PolC | polC |
| glutamine synthetase | glnA |
| hypothetical protein | hypothetical protein |
| hypothetical protein | hypothetical protein |
| hypothetical protein | noc |
| PTS system glucose-specific transporter subunit IIABC | glcB |
| methicillin resistance determinant protein FmhA | fmhA |
| hypothetical protein | hypothetical protein |
| hypothetical protein | hypothetical protein |
| formate dehydrogenase accessory protein | fdhD |
| alpha-acetolactate decarboxylase | budA1 |
| hypothetical protein | hypothetical protein |
| shikimate 5-dehydrogenase | aroE |
| hypothetical protein | darA/pstA |
| hypothetical protein | kimA |
| hypothetical protein | hypothetical protein |
| hypothetical protein | cshB |
| hypothetical protein | hypothetical protein |
| elongation factor Ts | tsf |
| hypothetical protein | sstA |
| recombination protein F | recF |
| chaperone protein DnaJ | dnaJ |
| pantoate--beta-alanine ligase | panC |
| seryl-tRNA synthetase | serS |

Up

| | |
|---|---|
| superantigen-like protein 7 | ssl7 |
| undecaprenyldiphospho-muramoylpentapeptide beta-N- | murG |
| hypothetical protein | hypothetical protein |
| hypothetical protein | hypothetical protein |
| CTP synthetase | pyrG |
| F0F1 ATP synthase subunit B | atpF |
| hypothetical protein | nirR |
| DNA polymerase III subunits gamma and tau | dnaX |
| signal recognition particle protein | ffh |
| histidinol dehydrogenase | hisD |
| hypothetical protein | divIC |
| hypothetical protein | esaD |
| hypothetical protein | fakB1 |
| hypothetical protein | hypothetical protein |
| UDP-N-acetylglucosamine 1-carboxyvinyltransferase | murA1 |
| hypothetical protein | aspB |
| hypothetical protein | hypothetical protein |
| hypothetical protein | hypothetical protein |
| ATP-dependent protease ATP-binding subunit ClpX | clpX |
| LysM domain-containing protein | LysM domain-containing protein |
| lipoyl synthase | lipA |
| HMG-CoA synthase | mvaS |
| pyrrolidone-carboxylate peptidase | pcp |
| oligopeptide ABC transporter substrate-binding protein | opp-3A |
| superantigen-like protein | ssl1 |
| M23/M37 peptidase domain-containing protein | M23/M37 peptidase domain-containing p |
| hypothetical protein | hypothetical protein |
| 50S ribosomal protein L28 | rpmB |
| thymidine kinase | tdk |
| hypothetical protein | hypothetical protein |
| hypothetical protein | hypothetical protein |
| F0F1 ATP synthase subunit alpha | atpA |
| acyl-CoA thioesterase | acyl-CoA thioesterase |
| protoheme IX farnesyltransferase | ctaB |
| anthranilate phosphoribosyltransferase | trpD |
| NAD-dependent DNA ligase | ligA |
| ATP-dependent DNA helicase PcrA | pcrA |
| hypothetical protein | hypothetical protein |
| hypothetical protein | hypothetical protein |
| hypothetical protein | atpI |
| F0F1 ATP synthase subunit delta | atpH |
| ammonium transporter | nrgA |
| UvrB/UvrC motif-containing protein | mcsA |
| 1-phosphofructokinase | fruB |
| isopentenyl pyrophosphate isomerase | fni |
| 50S ribosomal protein L7Ae-like protein | rplGB |
| assimilatory nitrite reductase [NAD(P)H] large subunit | nasD |
| glyceraldehyde 3-phosphate dehydrogenase 2 | gapB |
| hypothetical protein | hypothetical protein |
| LysM domain-containing protein | LysM domain-containing protein |
| polynucleotide phosphorylase/polyadenylase | pnpA |

| | |
|---|---|
| hypothetical protein | htsC |
| respiratory response protein SrrB | srrB |
| 50S ribosomal protein L11 methyltransferase | prmA |
| hypothetical protein | hypothetical protein |
| ABC transporter ATP-binding protein | ABC transporter ATP-binding protein |
| endonuclease IV | nfo |
| hypothetical protein | hypothetical protein |
| permease domain-containing protein | hrtB |
| 2-dehydropantoate 2-reductase | 2-dehydropantoate 2-reductase |
| hypothetical protein | mdeA |
| heat-inducible transcription repressor HrcA | hrcA |
| hypothetical protein | hypothetical protein |
| MarR family transcriptional regulator | MarR family transcriptional regulator |
| hypothetical protein | hypothetical protein |
| branched-chain alpha-keto acid dehydrogenase subunit | pdhC |
| cobalt transporter ATP-binding subunit | cbiO |
| hypothetical protein | sigS |
| branched-chain amino acid transport system II carrier pr | brnQ2 |
| 3-isopropylmalate dehydrogenase | leuB |
| hypothetical protein | hypothetical protein |
| sensor histidine kinase | airS |
| F0F1 ATP synthase subunit A | atpB |
| hypothetical protein | hypothetical protein |
| hypothetical protein | hypothetical protein |
| hypothetical protein | hypothetical protein |
| ATP-dependent DNA helicase RecQ | recQ1 |
| hypothetical protein | rnmV |
| 4-aminobutyrate aminotransferase | 4-aminobutyrate aminotransferase |
| hypothetical protein | hypothetical protein |
| hypothetical protein | hypothetical protein |
| hypothetical protein | hypothetical protein |
| ketol-acid reductoisomerase | ilvC |
| phosphate ABC transporter substrate-binding protein | pstS |
| penicillin-binding protein 2 | pbp2 |
| adenine phosphoribosyltransferase | apt |
| hypothetical protein | hypothetical protein |
| hypothetical protein | hypothetical protein |
| glycosyltransferase | sgtB |
| thymidylate kinase | tmk |
| UDP-N-acetylglucosamine 2-epimerase | capP |
| amino acid ABC transporter ATP-binding protein | tcyC |
| hypothetical protein | hypothetical protein |
| asparaginyl-tRNA synthetase | asnC |
| hypothetical protein | hypothetical protein |
| 2-isopropylmalate synthase | leuA |
| hypothetical protein | hypothetical protein |
| glucose induced biofilm accessory gene | gbaB |
| ribonuclease R | rnr |
| hypothetical protein | hypothetical protein |
| dihydrolipoamide dehydrogenase | pdhD |
| hypothetical protein | hypothetical protein |

| | |
|---|---|
| hypothetical protein | norB |
| aspartate semialdehyde dehydrogenase | asd |
| Holliday junction DNA helicase RuvA | ruvA |
| GTPase ObgE | obgE |
| polysaccharide biosynthesis protein | polysaccharide biosynthesis protein |
| tyrosyl-tRNA synthetase | tyrS |
| hypothetical protein | hypothetical protein |
| 1,4-dihydroxy-2-naphthoyl-CoA synthase | menB |
| hypothetical protein | farR |
| valyl-tRNA synthetase | valS |
| hypothetical protein | hypothetical protein |
| competence-damage inducible protein cinA | cinA |
| signal recognition particle-docking protein FtsY | ftsY |
| accessory Sec system glycosylation chaperone GtfB | gtfB |
| hypothetical protein | hypothetical protein |
| hypothetical protein | hypothetical protein |
| GTP pyrophosphokinase | relA |
| hypothetical protein | hflX |
| hypothetical protein | esaE |
| hypothetical protein | hypothetical protein |
| secretory antigen SsaA-like protein | secretory antigen SsaA-like protein |
| hypothetical protein | htsB |
| hypothetical protein | hypothetical protein |
| hypothetical protein | hypothetical protein |
| quinolone resistance protein NorA | norA |
| 3-isopropylmalate dehydratase small subunit | leuD |
| hypothetical protein | mspA |
| hypothetical protein | hypothetical protein |
| methicillin resistance factor | femB |
| hypothetical protein | hypothetical protein |
| fatty acid biosynthesis transcriptional regulator | fapR |
| hypothetical protein | hypothetical protein |
| isopropylmalate isomerase large subunit | leuC |
| hypothetical protein | rsfS |
| hypothetical protein | hypothetical protein |
| uroporphyrin-III C-methyltransferase | nasF |
| GTP-dependent nucleic acid-binding protein EngD | ychF |
| hypothetical protein | hypothetical protein |
| hypothetical protein | hypothetical protein |
| hypothetical protein | hypothetical protein |
| hypothetical protein | fakA |
| glycerol-3-phosphate transporter | glpT |
| aspartate alpha-decarboxylase | panD |
| acetyl-CoA acetyltransferase | fadA |
| hypothetical protein | ctaM |
| amino acid ABC transporter permease | tcyB |
| membrane protein | membrane protein |
| hypothetical protein | pgpB |
| hypothetical protein | rsgA |
| phosphopantetheine adenylyltransferase | coaD |
| 2-amino-4-hydroxy-6- hydroxymethyldihydropteridine py | folK |

| | |
|---|---|
| hypothetical protein | hypothetical protein |
| hypothetical protein | hypothetical protein |
| hypothetical protein | hypothetical protein |
| hypothetical protein | hypothetical protein |
| Riboflavin transporter | ribU |
| hypothetical protein | hypothetical protein |
| two-component response regulator | walR |
| hypothetical protein | hypothetical protein |
| D-alanyl-alanine synthetase A | ddl |
| hypothetical protein | lpl8 |
| NADH dehydrogenase subunit 5 | mpsA |
| aldose 1-epimerase | galM |
| hypothetical protein | hypothetical protein |
| hypothetical protein | tag |
| hypothetical protein | hypothetical protein |
| HemK family modification methylase | prmC |
| superantigen-like protein | ssl8 |
| transcriptional regulator NrdR | nrdR |
| peptide ABC transporter ATP-binding protein | peptide ABC transporter ATP-binding prc |
| hypothetical protein | opp-3D |
| (3R)-hydroxymyristoyl-ACP dehydratase | fabZ |
| hypothetical protein | sasG |
| hypothetical protein | hypothetical protein |
| hypothetical protein | hypothetical protein |
| preprotein translocase subunit SecG | secG |
| succinate dehydrogenase cytochrome b-558 subunit | sdhC |
| hypothetical protein | hypothetical protein |
| hypothetical protein | hypothetical protein |
| PTS system transporter | glvC |
| adenylosuccinate synthetase | purA |
| phosphocarrier protein HPr | ptsH |
| folylpolyglutamate synthase/dihydrofolate synthase | folC |
| manganese-dependent inorganic pyrophosphatase | ppaC |
| DNA topoisomerase III | topB |
| hypothetical protein | hypothetical protein |
| hypothetical protein | hypothetical protein |
| cell division protein | ftsA |
| superantigen-like protein | ssl10 |
| hypothetical protein | dtpT |
| hypothetical protein | hypothetical protein |
| glycine cleavage system aminomethyltransferase T | gcvT |
| hypothetical protein | sasF |
| hypothetical protein | hypothetical protein |
| accessory Sec system glycosyltransferase GtfA | gtfA |
| hypothetical protein | hypothetical protein |
| RNA methyltransferase | rumA |
| peptidoglycan pentaglycine interpeptide biosynthesis prc | femX |
| oligopeptide ABC transporter ATP-binding protein | opp-3F |
| hypothetical protein | hypothetical protein |
| hypothetical protein | hypothetical protein |
| hypothetical protein | hypothetical protein |

| | |
|---|---|
| DNA-binding response regulator | airR |
| aspartate kinase | lysC |
| molybdenum cofactor biosynthesis protein A | moaA |
| | sarZ |
| bifunctional N-acetylglucosamine-1-phosphate uridyltran | glmU |
| hypothetical protein | hypothetical protein |
| hypothetical protein | hypothetical protein |
| hypothetical protein | hypothetical protein |
| hypothetical protein | graS |
| ATP:guanido phosphotransferase | mcsB |
| hypothetical protein | lpl3 |
| hypothetical protein | hypothetical protein |
| hypothetical protein | hypothetical protein |
| hypothetical protein | hypothetical protein |
| hypothetical protein | hypothetical protein |
| glutamate-1-semialdehyde aminotransferase | gsaB |
| 6-phospho-beta-galactosidase | lacG |
| DNA gyrase subunit A | gyrA |
| hypothetical protein | plsY |
| chaperone protein HchA | hchA |
| hypothetical protein | walJ |
| dihydroxy-acid dehydratase | ilvD |
| lipid kinase | dgkB |
| hypothetical protein | metX |
| (dimethylallyl)adenosine tRNA methylthiotransferase | miaB |
| dimethyladenosine transferase | ksgA |
| acetyl-CoA carboxylase biotin carboxylase | acetyl-CoA carboxylase biotin carboxylas |
| fructose specific permease | fruA |
| hypothetical protein | hypothetical protein |
| phosphoribosylaminoimidazole-succinocarboxamide sy | purC |
| hypothetical protein | hypothetical protein |
| hypothetical protein | mpsC |
| Holliday junction DNA helicase RuvB | ruvB |
| cell wall biosynthesis protein ScdA | scdA |
| SsrA-binding protein | smpB |
| cell division protein FtsZ | ftsZ |
| tRNA pseudouridine synthase B | truB |
| hypothetical protein | umuC |
| 30S ribosomal protein S14 | rpsN |
| hypothetical protein | hypothetical protein |
| preprotein translocase subunit YajC | yajC |
| hypothetical protein | hypothetical protein |
| monovalent cation/H+ antiporter subunit E | mnhE |
| epimerase/dehydratase | epimerase/dehydratase |
| hypothetical protein | hypothetical protein |
| methylated-DNA--protein-cysteine methyltransferase | methylated-DNA--protein-cysteine methy |
| fibrinogen-binding protein-like protein | ecb |
| acyl carrier protein | acpP |
| hypothetical protein | corA2 |
| ica operon transcriptional regulator IcaR | icaR |
| 7-cyano-7-deazaguanine reductase | queF |

| | |
|---|---|
| hypothetical protein | hypothetical protein |
| L-asparaginase | ansA |
| signal peptidase IB | spsB |
| transcriptional regulator | msrR |
| branched-chain amino acid aminotransferase | ilvE |
| RNase J1 | rnjA |
| hypothetical protein | hypothetical protein |
| malate:quinone oxidoreductase | lqo |
| peptide deformylase | def |
| methicillin resistance factor FemA | femA |
| hypothetical protein | hypothetical protein |
| hypothetical protein | hypothetical protein |
| hypothetical protein | ndh2 |
| S-adenosylmethionine:tRNA ribosyltransferase-isomeras | queA |
| riboflavin biosynthesis protein RibF | ribC |
| hypothetical protein | hypothetical protein |
| hypothetical protein | hypothetical protein |
| homoserine dehydrogenase | hom |
| hypothetical protein | hypothetical protein |
| hypothetical protein | corA |
| ornithine aminotransferase | argD |
| hypothetical protein | mpfB |
| 2-dehydropantoate 2-reductase | panE |
| pyruvate kinase | pykA |
| hypothetical protein | ctsR |
| sensory box histidine kinase VicK | walK |
| threonyl-tRNA synthetase | thrS |
| cyclophilin-type peptidyl-prolyl cis-trans isomerase | ppiB |
| hypothetical protein | thrR |
| hypothetical protein | hypothetical protein |
| hypothetical protein | hypothetical protein |
| ABC transporter ATP-binding protein | vga |
| recombination protein RecR | recR |
| succinate dehydrogenase flavoprotein subunit | sdhA |
| cation transporter E1-E2 family ATPase | copZ |
| hypothetical protein | ftsK |
| PTS system glucose-specific protein | glcA |
| 1,4-dihydroxy-2-naphthoate octaprenyltransferase | menA |
| hypothetical protein | walH |
| hypothetical protein | hypothetical protein |
| trigger factor | tig |
| hypothetical protein | hypothetical protein |
| hypothetical protein | hypothetical protein |
| hypothetical protein | hypothetical protein |
| hypothetical protein | gmpA |
| tRNA pseudouridine synthase A | truA |
| hypothetical protein | traP |
| glutathione peroxidase | bsaA |
| high affinity nickel transporter | nixA |
| hypothetical protein | esxC |
| undecaprenyl pyrophosphate phosphatase | uppP |

Up

| | |
|---|---|
| hypothetical protein | hypothetical protein |
| monovalent cation/H+ antiporter subunit D | mnhD |
| hypothetical protein | pde2 |
| hypothetical protein | isdH |
| hypothetical protein | hypothetical protein |
| hypothetical protein | hypothetical protein |
| chorismate synthase | aroC |
| hypothetical protein | hypothetical protein |
| replicative DNA helicase | dnaC |
| hypothetical protein | sbnI |
| hypothetical protein | hypothetical protein |
| penicillin-binding protein 3 | pbp3 |
| phosphoribosylamine--glycine ligase | purD |
| hypothetical protein | hypothetical protein |
| hypothetical protein | hypothetical protein |
| molybdopterin converting factor subunit 1 | moaD |
| 2,3,4,5-tetrahydropyridine-2,6-dicarboxylate N-acetyltrar | dapD |
| hypothetical protein | polX |
| hypothetical protein | hypothetical protein |
| queuine tRNA-ribosyltransferase | tgt |
| hypothetical protein | ftsH |
| accessory regulator A | sarA |
| transcriptional regulator Spx | spxA |
| extramembranal protein | dltD |
| hypothetical protein | hypothetical protein |
| hypothetical protein | hypothetical protein |
| hypothetical protein | hypothetical protein |
| ABC transporter | znuC |
| 4-hydroxy-tetrahydrodipicolinate synthase | dapA |
| hypothetical protein | mhqR |
| phenylalanyl-tRNA synthetase subunit beta | pheT |
| hypothetical protein | cozEa |
| hypothetical protein | hypothetical protein |
| glutamate racemase | murI |
| 23S rRNA (adenine(2503)-C(2))-methyltransferase RlmN | rlmN |
| molybdopterin-guanine dinucleotide biosynthesis protein | mobA |
| hypothetical protein | hypothetical protein |
| hypothetical protein | hypothetical protein |
| quinol oxidase subunit IV | qoxD |
| methionyl-tRNA synthetase | metS |
| hypothetical protein | ktrD |
| hypothetical protein | hypothetical protein |
| hypothetical protein | hypothetical protein |
| Na+/H+ antiporter NhaC | nhaC |
| ABC transporter ATP-binding protein | sufC |
| hypothetical protein | hypothetical protein |
| glyceraldehyde-3-phosphate dehydrogenase | gapA |
| RNasaY | rny |
| ica operon transcriptional regulator IcaR | *icaR* |

| gene | locus | sample_1 | sample_2 | status | value_1 | value_2 |
|---|---|---|---|---|---|---|
| SAOUHSC_00069 | NC_007795.1 | wt_R | mut_A | OK | 195.696 | 3255.41 |
| SAOUHSC_02416 | NC_007795.1 | mut_A | wt_R | OK | 22.9439 | 276.812 |
| SAOUHSC_00093 | NC_007795.1 | mut_A | wt_R | OK | 306.813 | 2403.5 |
| SAOUHSC_02820 | NC_007795.1 | mut_A | wt_R | OK | 9.08806 | 68.5361 |
| SAOUHSC_02504 | NC_007795.1 | mut_A | wt_R | OK | 2524.7 | 15010.1 |
| SAOUHSC_03024 | NC_007795.1 | mut_A | wt_R | OK | 72.3003 | 388.027 |
| SAOUHSC_02571 | NC_007795.1 | mut_A | wt_R | OK | 49.9851 | 261.399 |
| SAOUHSC_02485 | NC_007795.1 | mut_A | wt_R | OK | 1721.24 | 8808.25 |
| SAOUHSC_00499 | NC_007795.1 | mut_A | wt_R | OK | 211.35 | 1043.11 |
| SAOUHSC_02486 | NC_007795.1 | mut_A | wt_R | OK | 1964.09 | 9670.1 |
| SAOUHSC_02821 | NC_007795.1 | mut_A | wt_R | OK | 8.61057 | 41.9795 |
| SAOUHSC_02887 | NC_007795.1 | mut_A | wt_R | OK | 117.92 | 560.681 |
| SAOUHSC_00070 | NC_007795.1 | mut_A | wt_R | OK | 18.3886 | 85.5803 |
| SAOUHSC_00500 | NC_007795.1 | mut_A | wt_R | OK | 271.89 | 1259.81 |
| SAOUHSC_02830 | NC_007795.1 | mut_A | wt_R | OK | 171.833 | 789.217 |
| SAOUHSC_00246 | NC_007795.1 | mut_A | wt_R | OK | 19.9601 | 91.3027 |
| SAOUHSC_02512a | NC_007795.1 | mut_A | wt_R | OK | 2328.51 | 10303.2 |
| SAOUHSC_02580 | NC_007795.1 | mut_A | wt_R | OK | 72.9631 | 314.806 |
| SAOUHSC_00007 | NC_007795.1 | mut_A | wt_R | OK | 331.456 | 1345.21 |
| SAOUHSC_00349 | NC_007795.1 | mut_A | wt_R | OK | 2209.01 | 8672.14 |
| SAOUHSC_01887 | NC_007795.1 | mut_A | wt_R | OK | 20.7329 | 80.0338 |
| SAOUHSC_01888 | NC_007795.1 | mut_A | wt_R | OK | 25.802 | 97.7484 |
| SAOUHSC_01504 | NC_007795.1 | mut_A | wt_R | OK | 135.149 | 509.517 |
| SAOUHSC_00253 | NC_007795.1 | mut_A | wt_R | OK | 47.37 | 166.317 |
| SAOUHSC_00995 | NC_007795.1 | mut_A | wt_R | OK | 27.0405 | 91.773 |
| SAOUHSC_02725 | NC_007795.1 | mut_A | wt_R | OK | 200.843 | 674.284 |
| SAOUHSC_02851 | NC_007795.1 | mut_A | wt_R | OK | 139.742 | 461.794 |
| SAOUHSC_02849 | NC_007795.1 | mut_A | wt_R | OK | 247.537 | 816.494 |
| SAOUHSC_02815 | NC_007795.1 | mut_A | wt_R | OK | 42.5985 | 140.345 |
| SAOUHSC_01058 | NC_007795.1 | mut_A | wt_R | OK | 420.362 | 1368.01 |
| SAOUHSC_01031 | NC_007795.1 | mut_A | wt_R | OK | 113.763 | 366.494 |
| SAOUHSC_00581 | NC_007795.1 | mut_A | wt_R | OK | 29.0582 | 93.2719 |
| SAOUHSC_02773 | NC_007795.1 | mut_A | wt_R | OK | 116.335 | 367.75 |
| SAOUHSC_01877 | NC_007795.1 | mut_A | wt_R | OK | 64.3811 | 199.233 |
| SAOUHSC_02850 | NC_007795.1 | mut_A | wt_R | OK | 177.466 | 541.001 |
| SAOUHSC_00582 | NC_007795.1 | mut_A | wt_R | OK | 19.919 | 60.3984 |
| SAOUHSC_01889 | NC_007795.1 | mut_A | wt_R | OK | 40.9755 | 123.797 |
| SAOUHSC_00920 | NC_007795.1 | mut_A | wt_R | OK | 176.008 | 531.612 |
| SAOUHSC_00880 | NC_007795.1 | mut_A | wt_R | OK | 33.6507 | 101.186 |
| SAOUHSC_00525 | NC_007795.1 | mut_A | wt_R | OK | 1648.22 | 4927.18 |
| SAOUHSC_02866 | NC_007795.1 | mut_A | wt_R | OK | 10.594 | 30.9026 |
| SAOUHSC_02923 | NC_007795.1 | mut_A | wt_R | OK | 14.0127 | 40.6753 |
| SAOUHSC_00255 | NC_007795.1 | mut_A | wt_R | OK | 18.6761 | 54.0742 |
| SAOUHSC_00435 | NC_007795.1 | mut_A | wt_R | OK | 21.9898 | 63.0382 |
| SAOUHSC_03042 | NC_007795.1 | mut_A | wt_R | OK | 5.21324 | 14.8756 |
| SAOUHSC_01993 | NC_007795.1 | mut_A | wt_R | OK | 0.427284 | 1.2124 |
| SAOUHSC_02388 | NC_007795.1 | mut_A | wt_R | OK | 40.3193 | 113.89 |
| SAOUHSC_01451 | NC_007795.1 | mut_A | wt_R | OK | 290.148 | 818.001 |
| trmD | NC_007795.1 | mut_A | wt_R | OK | 129.04 | 355.249 |
| SAOUHSC_02374 | NC_007795.1 | mut_A | wt_R | OK | 55.2502 | 151.429 |

| | | | | | | |
|---|---|---|---|---|---|---|
| SAOUHSC_01893 | NC_007795.1 | mut_A | wt_R | OK | 20.7918 | 56.9135 |
| SAOUHSC_02823 | NC_007795.1 | mut_A | wt_R | OK | 33.3822 | 91.0264 |
| SAOUHSC_02554 | NC_007795.1 | mut_A | wt_R | OK | 140.186 | 382.165 |
| SAOUHSC_02937 | NC_007795.1 | mut_A | wt_R | OK | 15.1112 | 41.0593 |
| SAOUHSC_02963 | NC_007795.1 | mut_A | wt_R | OK | 140.392 | 380.615 |
| SAOUHSC_00373 | NC_007795.1 | mut_A | wt_R | OK | 193.527 | 523.011 |
| SAOUHSC_01452 | NC_007795.1 | mut_A | wt_R | OK | 190.67 | 512.653 |
| SAOUHSC_01825 | NC_007795.1 | mut_A | wt_R | OK | 135.332 | 361.713 |
| SAOUHSC_01392 | NC_007795.1 | mut_A | wt_R | OK | 400.841 | 1067.42 |
| rimM | NC_007795.1 | mut_A | wt_R | OK | 95.8352 | 253.995 |
| SAOUHSC_02369 | NC_007795.1 | mut_A | wt_R | OK | 297.155 | 785.794 |
| SAOUHSC_01385 | NC_007795.1 | mut_A | wt_R | OK | 12.7608 | 33.6802 |
| SAOUHSC_03002 | NC_007795.1 | mut_A | wt_R | OK | 2.34643 | 6.1756 |
| SAOUHSC_00372 | NC_007795.1 | mut_A | wt_R | OK | 162.549 | 427.487 |
| SAOUHSC_02410 | NC_007795.1 | mut_A | wt_R | OK | 0.415289 | 1.08876 |
| SAOUHSC_02389 | NC_007795.1 | mut_A | wt_R | OK | 31.118 | 81.4942 |
| SAOUHSC_00427 | NC_007795.1 | mut_A | wt_R | OK | 184.714 | 480.459 |
| SAOUHSC_02630 | NC_007795.1 | mut_A | wt_R | OK | 413.512 | 1062.85 |
| SAOUHSC_01878 | NC_007795.1 | mut_A | wt_R | OK | 63.0665 | 161.614 |
| SAOUHSC_00528 | NC_007795.1 | mut_A | wt_R | OK | 6241.96 | 15978.5 |
| SAOUHSC_01062 | NC_007795.1 | mut_A | wt_R | OK | 992.115 | 2531.95 |
| SAOUHSC_01991 | NC_007795.1 | mut_A | wt_R | OK | 44.1098 | 112.543 |
| SAOUHSC_00483 | NC_007795.1 | mut_A | wt_R | OK | 1422.37 | 3618.92 |
| rplA | NC_007795.1 | mut_A | wt_R | OK | 1392.27 | 3529.46 |
| SAOUHSC_01330 | NC_007795.1 | mut_A | wt_R | OK | 82.629 | 209.451 |
| SAOUHSC_A02450 | NC_007795.1 | mut_A | wt_R | OK | 36.5843 | 92.3736 |
| SAOUHSC_01909 | NC_007795.1 | mut_A | wt_R | OK | 230.815 | 582.097 |
| SAOUHSC_00580 | NC_007795.1 | mut_A | wt_R | OK | 95.7709 | 240.69 |
| SAOUHSC_02373 | NC_007795.1 | mut_A | wt_R | OK | 71.26 | 177.957 |
| SAOUHSC_02750 | NC_007795.1 | mut_A | wt_R | OK | 79.8607 | 199.055 |
| SAOUHSC_00748 | NC_007795.1 | mut_A | wt_R | OK | 12.4981 | 31.0813 |
| SAOUHSC_00545 | NC_007795.1 | mut_A | wt_R | OK | 242.968 | 603.911 |
| SAOUHSC_01032 | NC_007795.1 | mut_A | wt_R | OK | 120.534 | 298.055 |
| SAOUHSC_02093 | NC_007795.1 | mut_A | wt_R | OK | 71.2455 | 176.151 |
| SAOUHSC_02590 | NC_007795.1 | mut_A | wt_R | OK | 128.891 | 317.859 |
| SAOUHSC_00244 | NC_007795.1 | mut_A | wt_R | OK | 79.7163 | 195.958 |
| SAOUHSC_01386 | NC_007795.1 | mut_A | wt_R | OK | 12.071 | 29.6696 |
| hslO | NC_007795.1 | mut_A | wt_R | OK | 140.546 | 344.98 |
| SAOUHSC_01824 | NC_007795.1 | mut_A | wt_R | OK | 188.915 | 463.593 |
| SAOUHSC_01440 | NC_007795.1 | mut_A | wt_R | OK | 83.8684 | 205.618 |
| SAOUHSC_01354 | NC_007795.1 | mut_A | wt_R | OK | 331.867 | 812.279 |
| SAOUHSC_01387 | NC_007795.1 | mut_A | wt_R | OK | 10.0586 | 24.6066 |
| SAOUHSC_03009 | NC_007795.1 | mut_A | wt_R | OK | 4.7706 | 11.6516 |
| SAOUHSC_01049 | NC_007795.1 | mut_A | wt_R | OK | 191.984 | 468.678 |
| SAOUHSC_00039 | NC_007795.1 | mut_A | wt_R | OK | 245.149 | 598.227 |
| SAOUHSC_01008 | NC_007795.1 | mut_A | wt_R | OK | 28.9332 | 70.4725 |
| SAOUHSC_02825 | NC_007795.1 | mut_A | wt_R | OK | 2.39355 | 5.81107 |
| SAOUHSC_00524 | NC_007795.1 | mut_A | wt_R | OK | 1378.76 | 3344.38 |
| SAOUHSC_00853 | NC_007795.1 | mut_A | wt_R | OK | 11.3355 | 27.3865 |
| SAOUHSC_02695 | NC_007795.1 | mut_A | wt_R | OK | 159.299 | 383.103 |
| SAOUHSC_01891 | NC_007795.1 | mut_A | wt_R | OK | 47.253 | 113.327 |

| | | | | | |
|---|---|---|---|---|---|
| SAOUHSC_01222 | NC_007795.1 mut_A | wt_R | OK | 303.054 | 725.047 |
| SAOUHSC_01894 | NC_007795.1 mut_A | wt_R | OK | 39.0062 | 92.8432 |
| SAOUHSC_00391 | NC_007795.1 mut_A | wt_R | OK | 3.88072 | 9.20243 |
| SAOUHSC_01045 | NC_007795.1 mut_A | wt_R | OK | 69.0998 | 163.694 |
| SAOUHSC_02417 | NC_007795.1 mut_A | wt_R | OK | 318.78 | 752.553 |
| SAOUHSC_00523 | NC_007795.1 mut_A | wt_R | OK | 285.045 | 672.192 |
| SAOUHSC_01651 | NC_007795.1 mut_A | wt_R | OK | 5662.9 | 13327 |
| SAOUHSC_00863 | NC_007795.1 mut_A | wt_R | OK | 26.4847 | 62.3263 |
| SAOUHSC_00701 | NC_007795.1 mut_A | wt_R | OK | 49.4668 | 115.661 |
| SAOUHSC_00910 | NC_007795.1 mut_A | wt_R | OK | 67.7719 | 157.69 |
| SAOUHSC_03010 | NC_007795.1 mut_A | wt_R | OK | 5.13398 | 11.9226 |
| SAOUHSC_01727 | NC_007795.1 mut_A | wt_R | OK | 179.884 | 416.506 |
| SAOUHSC_02629 | NC_007795.1 mut_A | wt_R | OK | 694.918 | 1608.27 |
| SAOUHSC_00556 | NC_007795.1 mut_A | wt_R | OK | 49.3198 | 114.136 |
| SAOUHSC_00954 | NC_007795.1 mut_A | wt_R | OK | 77.6226 | 179.55 |
| rpsL | NC_007795.1 mut_A | wt_R | OK | 6893.2 | 15938 |
| rpsD | NC_007795.1 mut_A | wt_R | OK | 2199.03 | 5084 |
| rplK | NC_007795.1 mut_A | wt_R | OK | 4542 | 10443.1 |
| SAOUHSC_02316 | NC_007795.1 mut_A | wt_R | OK | 830.41 | 1906.92 |
| SAOUHSC_02272 | NC_007795.1 mut_A | wt_R | OK | 43.0341 | 98.7177 |
| SAOUHSC_01384 | NC_007795.1 mut_A | wt_R | OK | 26.0865 | 59.8097 |
| rplS | NC_007795.1 mut_A | wt_R | OK | 3803.7 | 8696.28 |
| SAOUHSC_01756 | NC_007795.1 mut_A | wt_R | OK | 2439.14 | 5552.85 |
| rpsO | NC_007795.1 mut_A | wt_R | OK | 3340.38 | 7582.34 |
| SAOUHSC_01835 | NC_007795.1 mut_A | wt_R | OK | 50.2585 | 113.496 |
| SAOUHSC_02740 | NC_007795.1 mut_A | wt_R | OK | 85.7202 | 193.28 |
| SAOUHSC_01030 | NC_007795.1 mut_A | wt_R | OK | 188.173 | 423.678 |
| SAOUHSC_02829 | NC_007795.1 mut_A | wt_R | OK | 163.584 | 367.362 |
| rnpA | NC_007795.1 mut_A | wt_R | OK | 47.225 | 105.674 |
| SAOUHSC_02694 | NC_007795.1 mut_A | wt_R | OK | 142.877 | 319.124 |
| SAOUHSC_01285 | NC_007795.1 mut_A | wt_R | OK | 404.393 | 902.39 |
| SAOUHSC_02448 | NC_007795.1 mut_A | wt_R | OK | 46.2121 | 103.086 |
| SAOUHSC_00474 | NC_007795.1 mut_A | wt_R | OK | 2047.36 | 4560.05 |
| SAOUHSC_02516 | NC_007795.1 mut_A | wt_R | OK | 270.378 | 601.18 |
| SAOUHSC_00700 | NC_007795.1 mut_A | wt_R | OK | 55.6998 | 123.683 |
| SAOUHSC_02731 | NC_007795.1 mut_A | wt_R | OK | 44.4841 | 98.4773 |
| SAOUHSC_02119 | NC_007795.1 mut_A | wt_R | OK | 113.98 | 251.429 |
| SAOUHSC_01316 | NC_007795.1 mut_A | wt_R | OK | 136.996 | 301.138 |
| SAOUHSC_01823 | NC_007795.1 mut_A | wt_R | OK | 137.482 | 301.961 |
| SAOUHSC_00749 | NC_007795.1 mut_A | wt_R | OK | 29.5817 | 64.6888 |
| SAOUHSC_02886 | NC_007795.1 mut_A | wt_R | OK | 1.00592 | 2.19785 |
| gid | NC_007795.1 mut_A | wt_R | OK | 352.794 | 769.223 |
| SAOUHSC_02271 | NC_007795.1 mut_A | wt_R | OK | 123.551 | 268.793 |
| SAOUHSC_02611 | NC_007795.1 mut_A | wt_R | OK | 102.25 | 222.076 |
| SAOUHSC_01702 | NC_007795.1 mut_A | wt_R | OK | 145.567 | 314.705 |
| SAOUHSC_02622 | NC_007795.1 mut_A | wt_R | OK | 82.8031 | 178.781 |
| SAOUHSC_00342 | NC_007795.1 mut_A | wt_R | OK | 55.8969 | 120.475 |
| SAOUHSC_02824 | NC_007795.1 mut_A | wt_R | OK | 6.24652 | 13.419 |
| SAOUHSC_00844 | NC_007795.1 mut_A | wt_R | OK | 57.9129 | 124.203 |
| SAOUHSC_00842 | NC_007795.1 mut_A | wt_R | OK | 94.3441 | 202.331 |
| SAOUHSC_00881 | NC_007795.1 mut_A | wt_R | OK | 213.917 | 456.829 |

| | | | | | | |
|---|---|---|---|---|---|---|
| rpmH | NC_007795.1 | mut_A | wt_R | OK | 4237.84 | 9048.77 |
| prfC | NC_007795.1 | mut_A | wt_R | OK | 209.272 | 445.847 |
| SAOUHSC_01046 | NC_007795.1 | mut_A | wt_R | OK | 56.9923 | 121.242 |
| SAOUHSC_00721 | NC_007795.1 | mut_A | wt_R | OK | 64.9092 | 137.75 |
| SAOUHSC_01034 | NC_007795.1 | mut_A | wt_R | OK | 189.932 | 402.136 |
| SAOUHSC_02658 | NC_007795.1 | mut_A | wt_R | OK | 22.7 | 47.9568 |
| hisS | NC_007795.1 | mut_A | wt_R | OK | 131.913 | 277.515 |
| SAOUHSC_01984 | NC_007795.1 | mut_A | wt_R | OK | 57.1979 | 119.271 |
| SAOUHSC_02321 | NC_007795.1 | mut_A | wt_R | OK | 264.807 | 551.962 |
| SAOUHSC_02789 | NC_007795.1 | mut_A | wt_R | OK | 23.7469 | 49.4696 |
| SAOUHSC_01266 | NC_007795.1 | mut_A | wt_R | OK | 168.065 | 348.833 |
| SAOUHSC_01757 | NC_007795.1 | mut_A | wt_R | OK | 4016.12 | 8333.43 |
| SAOUHSC_03015 | NC_007795.1 | mut_A | wt_R | OK | 1.77851 | 3.68972 |
| SAOUHSC_01021 | NC_007795.1 | mut_A | wt_R | OK | 1715.81 | 3557.57 |
| SAOUHSC_02530 | NC_007795.1 | mut_A | wt_R | OK | 19.8511 | 41.0909 |
| SAOUHSC_01700 | NC_007795.1 | mut_A | wt_R | OK | 190.412 | 393.842 |
| SAOUHSC_01019 | NC_007795.1 | mut_A | wt_R | OK | 981.971 | 2028.21 |
| SAOUHSC_03053 | NC_007795.1 | mut_A | wt_R | OK | 112.671 | 232.5 |
| SAOUHSC_03012 | NC_007795.1 | mut_A | wt_R | OK | 4.44946 | 9.17124 |
| SAOUHSC_01726 | NC_007795.1 | mut_A | wt_R | OK | 297.599 | 612.544 |
| SAOUHSC_01048 | NC_007795.1 | mut_A | wt_R | OK | 151.952 | 312.408 |
| SAOUHSC_00720 | NC_007795.1 | mut_A | wt_R | OK | 139.194 | 284.746 |
| SAOUHSC_03052 | NC_007795.1 | mut_A | wt_R | OK | 151.38 | 308.873 |
| SAOUHSC_02576 | NC_007795.1 | mut_A | wt_R | OK | 28.3524 | 57.6284 |
| SAOUHSC_01265 | NC_007795.1 | mut_A | wt_R | OK | 117.838 | 239.499 |
| SAOUHSC_00660 | NC_007795.1 | mut_A | wt_R | OK | 198.326 | 402.625 |
| SAOUHSC_01876 | NC_007795.1 | mut_A | wt_R | OK | 72.7331 | 147.58 |
| SAOUHSC_00771 | NC_007795.1 | mut_A | wt_R | OK | 200.723 | 407.203 |
| SAOUHSC_02250 | NC_007795.1 | mut_A | wt_R | OK | 6.20687 | 12.5805 |
| SAOUHSC_01661 | NC_007795.1 | mut_A | wt_R | OK | 73.9267 | 149.407 |
| SAOUHSC_00965 | NC_007795.1 | mut_A | wt_R | OK | 224.682 | 454.033 |
| SAOUHSC_00027 | NC_007795.1 | mut_A | wt_R | OK | 115.128 | 232.272 |
| SAOUHSC_02341 | NC_007795.1 | mut_A | wt_R | OK | 2564.88 | 5171.67 |
| SAOUHSC_01701 | NC_007795.1 | mut_A | wt_R | OK | 214.988 | 432.497 |
| SAOUHSC_02005 | NC_007795.1 | mut_A | wt_R | OK | 25.075 | 50.3813 |
| SAOUHSC_02139 | NC_007795.1 | mut_A | wt_R | OK | 647.126 | 1297.52 |
| SAOUHSC_01716 | NC_007795.1 | mut_A | wt_R | OK | 257.921 | 516.681 |
| SAOUHSC_00928 | NC_007795.1 | mut_A | wt_R | OK | 1.69668 | 3.39649 |
| SAOUHSC_01078 | NC_007795.1 | mut_A | wt_R | OK | 2107.09 | 4216.9 |
| SAOUHSC_00719 | NC_007795.1 | mut_A | wt_R | OK | 126.211 | 252.241 |
| SAOUHSC_02831 | NC_007795.1 | mut_A | wt_R | OK | 68.3411 | 136.4 |
| SAOUHSC_02372 | NC_007795.1 | mut_A | wt_R | OK | 170.56 | 339.962 |
| SAOUHSC_02779 | NC_007795.1 | mut_A | wt_R | OK | 572.169 | 1139.59 |
| SAOUHSC_00248 | NC_007795.1 | mut_A | wt_R | OK | 48.5688 | 96.6277 |
| SAOUHSC_01717 | NC_007795.1 | mut_A | wt_R | OK | 149.233 | 296.729 |
| SAOUHSC_03008 | NC_007795.1 | mut_A | wt_R | OK | 8.24197 | 16.3593 |
| SAOUHSC_02978 | NC_007795.1 | mut_A | wt_R | OK | 223.552 | 442.03 |
| SAOUHSC_01425 | NC_007795.1 | mut_A | wt_R | OK | 101.376 | 200.177 |
| SAOUHSC_03051 | NC_007795.1 | mut_A | wt_R | OK | 207.252 | 408.805 |
| SAOUHSC_01176 | NC_007795.1 | mut_A | wt_R | OK | 593.177 | 1169.28 |
| SAOUHSC_00964 | NC_007795.1 | mut_A | wt_R | OK | 190.171 | 374.034 |

| | | | | | |
|---|---|---|---|---|---|
| SAOUHSC_02896 | NC_007795.1 mut_A | wt_R | OK | 35.934 | 70.612 |
| SAOUHSC_00646 | NC_007795.1 mut_A | wt_R | OK | 45.8099 | 89.8873 |
| SAOUHSC_02322 | NC_007795.1 mut_A | wt_R | OK | 159.774 | 313.186 |
| SAOUHSC_02340 | NC_007795.1 mut_A | wt_R | OK | 2585.62 | 5052.21 |
| SAOUHSC_00550 | NC_007795.1 mut_A | wt_R | OK | 503.318 | 982.697 |
| SAOUHSC_00572 | NC_007795.1 mut_A | wt_R | OK | 66.2268 | 129.23 |
| SAOUHSC_02003 | NC_007795.1 mut_A | wt_R | OK | 129.889 | 253.216 |
| SAOUHSC_02362 | NC_007795.1 mut_A | wt_R | OK | 304.004 | 591.384 |
| SAOUHSC_02273 | NC_007795.1 mut_A | wt_R | OK | 522.829 | 1015.6 |
| SAOUHSC_01978 | NC_007795.1 mut_A | wt_R | OK | 32.103 | 62.3557 |
| SAOUHSC_00873 | NC_007795.1 mut_A | wt_R | OK | 404.844 | 782.682 |
| SAOUHSC_01737 | NC_007795.1 mut_A | wt_R | OK | 246.876 | 476.248 |
| SAOUHSC_00794 | NC_007795.1 mut_A | wt_R | OK | 3632.07 | 6980.39 |
| SAOUHSC_01990 | NC_007795.1 mut_A | wt_R | OK | 67.6997 | 130.078 |
| SAOUHSC_01496 | NC_007795.1 mut_A | wt_R | OK | 583.182 | 1118.96 |
| SAOUHSC_02933 | NC_007795.1 mut_A | wt_R | OK | 9.68142 | 18.563 |
| SAOUHSC_01653 | NC_007795.1 mut_A | wt_R | OK | 2018.48 | 3869.42 |
| SAOUHSC_01680 | NC_007795.1 mut_A | wt_R | OK | 177.018 | 338.152 |
| SAOUHSC_01047 | NC_007795.1 mut_A | wt_R | OK | 100.093 | 191.042 |
| SAOUHSC_01992 | NC_007795.1 mut_A | wt_R | OK | 82.6518 | 157.434 |
| SAOUHSC_01715 | NC_007795.1 mut_A | wt_R | OK | 428.214 | 815.211 |
| SAOUHSC_01898 | NC_007795.1 mut_A | wt_R | OK | 49.353 | 93.6425 |
| SAOUHSC_00753 | NC_007795.1 mut_A | wt_R | OK | 45.1968 | 85.7555 |
| SAOUHSC_00843 | NC_007795.1 mut_A | wt_R | OK | 55.8504 | 105.818 |
| SAOUHSC_00394 | NC_007795.1 mut_A | wt_R | OK | 5.51258 | 10.4436 |
| SAOUHSC_02605 | NC_007795.1 mut_A | wt_R | OK | 14.8155 | 28.0172 |
| SAOUHSC_00341 | NC_007795.1 mut_A | wt_R | OK | 8.28645 | 15.6601 |
| SAOUHSC_00374 | NC_007795.1 mut_A | wt_R | OK | 659.137 | 1245.03 |
| SAOUHSC_01807 | NC_007795.1 mut_A | wt_R | OK | 959.726 | 1811.2 |
| SAOUHSC_02461 | NC_007795.1 mut_A | wt_R | OK | 33.3982 | 63.0001 |
| SAOUHSC_02816 | NC_007795.1 mut_A | wt_R | OK | 51.5002 | 97.0121 |
| SAOUHSC_01295 | NC_007795.1 mut_A | wt_R | OK | 23.2574 | 43.804 |
| SAOUHSC_00375 | NC_007795.1 mut_A | wt_R | OK | 688.673 | 1296.93 |
| SAOUHSC_02919 | NC_007795.1 mut_A | wt_R | OK | 372.093 | 700.154 |
| SAOUHSC_02327 | NC_007795.1 mut_A | wt_R | OK | 638.141 | 1198.16 |
| SAOUHSC_02468 | NC_007795.1 mut_A | wt_R | OK | 1370.13 | 2565.33 |
| SAOUHSC_01755 | NC_007795.1 mut_A | wt_R | OK | 7005.75 | 13115.1 |
| SAOUHSC_00747 | NC_007795.1 mut_A | wt_R | OK | 16.4633 | 30.7908 |
| SAOUHSC_00060 | NC_007795.1 mut_A | wt_R | OK | 106.178 | 198.33 |
| SAOUHSC_00551 | NC_007795.1 mut_A | wt_R | OK | 499.295 | 928.961 |
| SAOUHSC_02582 | NC_007795.1 mut_A | wt_R | OK | 401.951 | 747.246 |
| SAOUHSC_01214 | NC_007795.1 mut_A | wt_R | OK | 167.901 | 310.708 |
| SAOUHSC_02579 | NC_007795.1 mut_A | wt_R | OK | 162.216 | 299.874 |
| SAOUHSC_01714 | NC_007795.1 mut_A | wt_R | OK | 524.437 | 968.196 |
| SAOUHSC_00994 | NC_007795.1 mut_A | wt_R | OK | 168.842 | 311.535 |
| SAOUHSC_00413 | NC_007795.1 mut_A | wt_R | OK | 184.313 | 339.742 |
| SAOUHSC_00549 | NC_007795.1 mut_A | wt_R | OK | 625.657 | 1152.93 |
| SAOUHSC_01662 | NC_007795.1 mut_A | wt_R | OK | 581.596 | 1070.45 |
| SAOUHSC_00531 | NC_007795.1 mut_A | wt_R | OK | 93.0156 | 170.559 |
| SAOUHSC_01438 | NC_007795.1 mut_A | wt_R | OK | 78.703 | 143.886 |
| SAOUHSC_01679 | NC_007795.1 mut_A | wt_R | OK | 347.154 | 633.699 |

| | | | | | |
|---|---|---|---|---|---|
| SAOUHSC_01352 | NC_007795.1 mut_A | wt_R | OK | 346.861 | 632.1 |
| SAOUHSC_02349 | NC_007795.1 mut_A | wt_R | OK | 1133.74 | 2054.51 |
| SAOUHSC_01040 | NC_007795.1 mut_A | wt_R | OK | 2928.94 | 5304.93 |
| SAOUHSC_00912 | NC_007795.1 mut_A | wt_R | OK | 290.641 | 525.705 |
| SAOUHSC_01988 | NC_007795.1 mut_A | wt_R | OK | 87.6462 | 158.532 |
| SAOUHSC_02014 | NC_007795.1 mut_A | wt_R | OK | 312.641 | 563.926 |
| SAOUHSC_00330 | NC_007795.1 mut_A | wt_R | OK | 107.518 | 193.781 |
| SAOUHSC_01496 | NC_007795.1 mut_A | wt_R | OK | 153.515 | 276.557 |
| SAOUHSC_00010 | NC_007795.1 mut_A | wt_R | OK | 16.6233 | 29.845 |
| SAOUHSC_00428 | NC_007795.1 mut_A | wt_R | OK | 218.317 | 391.876 |
| SAOUHSC_1342a | NC_007795.1 mut_A | wt_R | OK | 355.206 | 636.837 |
| SAOUHSC_02092 | NC_007795.1 mut_A | wt_R | OK | 266.581 | 477.355 |
| SAOUHSC_01077 | NC_007795.1 mut_A | wt_R | OK | 2312.71 | 4140.86 |
| SAOUHSC_00267 | NC_007795.1 mut_A | wt_R | OK | 112.185 | 200.777 |
| SAOUHSC_01197 | NC_007795.1 mut_A | wt_R | OK | 570.727 | 1016.98 |
| SAOUHSC_01022 | NC_007795.1 mut_A | wt_R | OK | 1205.52 | 2142.87 |
| SAOUHSC_01199 | NC_007795.1 mut_A | wt_R | OK | 821.375 | 1455.91 |
| SAOUHSC_01989 | NC_007795.1 mut_A | wt_R | OK | 116.646 | 206.74 |
| SAOUHSC_00670 | NC_007795.1 mut_A | wt_R | OK | 522.989 | 926.324 |
| SAOUHSC_02343 | NC_007795.1 mut_A | wt_R | OK | 2460.47 | 4350.97 |
| SAOUHSC_01697 | NC_007795.1 mut_A | wt_R | OK | 260.016 | 459.348 |
| SAOUHSC_01009 | NC_007795.1 mut_A | wt_R | OK | 83.9636 | 148.157 |
| SAOUHSC_02255 | NC_007795.1 mut_A | wt_R | OK | 200.418 | 353.462 |
| SAOUHSC_00662 | NC_007795.1 mut_A | wt_R | OK | 87.3649 | 153.744 |
| SAOUHSC_02690 | NC_007795.1 mut_A | wt_R | OK | 70.2115 | 123.43 |
| SAOUHSC_00344 | NC_007795.1 mut_A | wt_R | OK | 129.313 | 227.176 |
| SAOUHSC_00571 | NC_007795.1 mut_A | wt_R | OK | 96.8815 | 169.959 |
| SAOUHSC_00324 | NC_007795.1 mut_A | wt_R | OK | 74.6881 | 130.933 |
| SAOUHSC_00493 | NC_007795.1 mut_A | wt_R | OK | 607.495 | 1064.17 |
| SAOUHSC_01439 | NC_007795.1 mut_A | wt_R | OK | 84.7528 | 148.373 |
| SAOUHSC_01215 | NC_007795.1 mut_A | wt_R | OK | 271.329 | 474.782 |
| SAOUHSC_02897 | NC_007795.1 mut_A | wt_R | OK | 17.221 | 30.0911 |
| SAOUHSC_01492 | NC_007795.1 mut_A | wt_R | OK | 343.007 | 598.112 |
| SAOUHSC_01163 | NC_007795.1 mut_A | wt_R | OK | 407.401 | 709.067 |
| SAOUHSC_00265 | NC_007795.1 mut_A | wt_R | OK | 105.975 | 184.296 |
| SAOUHSC_02680 | NC_007795.1 mut_A | wt_R | OK | 669.726 | 1163.74 |
| SAOUHSC_00669 | NC_007795.1 mut_A | wt_R | OK | 376.802 | 654.716 |
| SAOUHSC_01666 | NC_007795.1 mut_A | wt_R | OK | 1085.39 | 1885.73 |
| SAOUHSC_02628 | NC_007795.1 mut_A | wt_R | OK | 87.7174 | 152.165 |
| SAOUHSC_01875 | NC_007795.1 mut_A | wt_R | OK | 287.755 | 498.993 |
| SAOUHSC_00005 | NC_007795.1 mut_A | wt_R | OK | 529.858 | 918.053 |
| SAOUHSC_02335 | NC_007795.1 mut_A | wt_R | OK | 44.889 | 77.4858 |
| SAOUHSC_00989 | NC_007795.1 mut_A | wt_R | OK | 57.1517 | 98.5299 |
| SAOUHSC_01351 | NC_007795.1 mut_A | wt_R | OK | 190.341 | 327.929 |
| SAOUHSC_01684 | NC_007795.1 mut_A | wt_R | OK | 827.907 | 1424.55 |
| SAOUHSC_00921 | NC_007795.1 mut_A | wt_R | OK | 635.529 | 1092.7 |
| SAOUHSC_01346 | NC_007795.1 mut_A | wt_R | OK | 228.568 | 392.579 |
| SAOUHSC_01491 | NC_007795.1 mut_A | wt_R | OK | 556.921 | 955.717 |
| SAOUHSC_02921 | NC_007795.1 mut_A | wt_R | OK | 138.781 | 237.476 |
| SAOUHSC_01777 | NC_007795.1 mut_A | wt_R | OK | 206.831 | 352.936 |
| SAOUHSC_00773 | NC_007795.1 mut_A | wt_R | OK | 17.4077 | 29.6705 |

| | | | | | |
|---|---|---|---|---|---|
| SAOUHSC_00733 | NC_007795.1 mut_A | wt_R | OK | 14.8221 | 25.235 |
| SAOUHSC_01267 | NC_007795.1 mut_A | wt_R | OK | 233.308 | 397.075 |
| SAOUHSC_02004 | NC_007795.1 mut_A | wt_R | OK | 208.601 | 354.948 |
| SAOUHSC_01710 | NC_007795.1 mut_A | wt_R | OK | 37.2478 | 63.1667 |
| SAOUHSC_01635 | NC_007795.1 mut_A | wt_R | OK | 99.4027 | 168.282 |
| SAOUHSC_01091 | NC_007795.1 mut_A | wt_R | OK | 188.894 | 319.294 |
| SAOUHSC_01198 | NC_007795.1 mut_A | wt_R | OK | 624.253 | 1054.24 |
| SAOUHSC_02802 | NC_007795.1 mut_A | wt_R | OK | 14.66 | 24.751 |
| SAOUHSC_02254 | NC_007795.1 mut_A | wt_R | OK | 308.745 | 521.23 |
| SAOUHSC_02686 | NC_007795.1 mut_A | wt_R | OK | 65.029 | 109.682 |
| SAOUHSC_00562 | NC_007795.1 mut_A | wt_R | OK | 212.299 | 358.014 |
| SAOUHSC_01856 | NC_007795.1 mut_A | wt_R | OK | 305.24 | 514.435 |
| SAOUHSC_02359 | NC_007795.1 mut_A | wt_R | OK | 183.523 | 309.171 |
| SAOUHSC_01689 | NC_007795.1 mut_A | wt_R | OK | 6237.36 | 10485.7 |
| SAOUHSC_A00992 | NC_007795.1 mut_A | wt_R | OK | 405.284 | 681.086 |
| SAOUHSC_00933 | NC_007795.1 mut_A | wt_R | OK | 430.734 | 723.846 |
| SAOUHSC_02520 | NC_007795.1 mut_A | wt_R | OK | 236.024 | 396.346 |
| SAOUHSC_00802 | NC_007795.1 mut_A | wt_R | OK | 188.777 | 316.71 |
| SAOUHSC_01735 | NC_007795.1 mut_A | wt_R | OK | 226.849 | 380.464 |
| SAOUHSC_01041 | NC_007795.1 mut_A | wt_R | OK | 3654.93 | 6128.65 |
| SAOUHSC_01437 | NC_007795.1 mut_A | wt_R | OK | 178.166 | 298.737 |
| SAOUHSC_02640 | NC_007795.1 mut_A | wt_R | OK | 8.61158 | 14.4379 |
| SAOUHSC_02932 | NC_007795.1 mut_A | wt_R | OK | 7.06896 | 11.8508 |
| SAOUHSC_01683 | NC_007795.1 mut_A | wt_R | OK | 1198.69 | 2006.33 |
| SAOUHSC_00991 | NC_007795.1 mut_A | wt_R | OK | 25.7221 | 42.9384 |
| SAOUHSC_00290 | NC_007795.1 mut_A | wt_R | OK | 185.775 | 310.106 |
| SAOUHSC_01732 | NC_007795.1 mut_A | wt_R | OK | 236.284 | 393.713 |
| SAOUHSC_01241 | NC_007795.1 mut_A | wt_R | OK | 88.9779 | 147.932 |
| SAOUHSC_01287 | NC_007795.1 mut_A | wt_R | OK | 988.274 | 1642.95 |
| SAOUHSC_00441 | NC_007795.1 mut_A | wt_R | OK | 95.1566 | 158.185 |
| SAOUHSC_00012 | NC_007795.1 mut_A | wt_R | OK | 24.8408 | 41.2529 |
| SAOUHSC_03049 | NC_007795.1 mut_A | wt_R | OK | 467.758 | 775.66 |
| SAOUHSC_02848 | NC_007795.1 mut_A | wt_R | OK | 1239.42 | 2054.38 |
| SAOUHSC_02696 | NC_007795.1 mut_A | wt_R | OK | 13.3134 | 22.0343 |
| SAOUHSC_01908 | NC_007795.1 mut_A | wt_R | OK | 415.661 | 687.885 |
| SAOUHSC_00480 | NC_007795.1 mut_A | wt_R | OK | 252.975 | 418.593 |
| SAOUHSC_02550 | NC_007795.1 mut_A | wt_R | OK | 53.531 | 88.5028 |
| SAOUHSC_02467 | NC_007795.1 mut_A | wt_R | OK | 1658.93 | 2742.37 |
| SAOUHSC_02935 | NC_007795.1 mut_A | wt_R | OK | 159.081 | 262.335 |
| SAOUHSC_01699 | NC_007795.1 mut_A | wt_R | OK | 140.772 | 231.155 |
| SAOUHSC_00452 | NC_007795.1 mut_A | wt_R | OK | 780.108 | 1279.82 |
| SAOUHSC_02733 | NC_007795.1 mut_A | wt_R | OK | 1108.71 | 1818.69 |
| SAOUHSC_00299 | NC_007795.1 mut_A | wt_R | OK | 1.31101 | 2.15046 |
| SAOUHSC_01659 | NC_007795.1 mut_A | wt_R | OK | 271.602 | 445.298 |
| SAOUHSC_00345 | NC_007795.1 mut_A | wt_R | OK | 170.946 | 280.122 |
| SAOUHSC_01234 | NC_007795.1 mut_A | wt_R | OK | 2004.72 | 3281.65 |
| SAOUHSC_00746 | NC_007795.1 mut_A | wt_R | OK | 15.6939 | 25.6409 |
| SAOUHSC_00004 | NC_007795.1 mut_A | wt_R | OK | 472.055 | 770.681 |
| SAOUHSC_01682 | NC_007795.1 mut_A | wt_R | OK | 336.388 | 548.119 |
| SAOUHSC_02918 | NC_007795.1 mut_A | wt_R | OK | 384.751 | 626.2 |
| SAOUHSC_00009 | NC_007795.1 mut_A | wt_R | OK | 580.75 | 944.416 |

| | | | | | |
|---|---|---|---|---|---|
| SAOUHSC_00392 | NC_007795.1 mut_A | wt_R | OK | 11.0552 | 17.9771 |
| SAOUHSC_01424 | NC_007795.1 mut_A | wt_R | OK | 284.214 | 462.072 |
| SAOUHSC_01696 | NC_007795.1 mut_A | wt_R | OK | 292.215 | 474.963 |
| SAOUHSC_01718 | NC_007795.1 mut_A | wt_R | OK | 102.36 | 166.02 |
| SAOUHSC_02368 | NC_007795.1 mut_A | wt_R | OK | 508.114 | 824.042 |
| SAOUHSC_02347 | NC_007795.1 mut_A | wt_R | OK | 1755.89 | 2843.29 |
| SAOUHSC_02685 | NC_007795.1 mut_A | wt_R | OK | 394.489 | 638.461 |
| SAOUHSC_00442 | NC_007795.1 mut_A | wt_R | OK | 224.997 | 364.118 |
| SAOUHSC_01207 | NC_007795.1 mut_A | wt_R | OK | 400.072 | 646.789 |
| SAOUHSC_03013 | NC_007795.1 mut_A | wt_R | OK | 3.96872 | 6.41346 |
| SAOUHSC_00482 | NC_007795.1 mut_A | wt_R | OK | 464.968 | 749.637 |
| SAOUHSC_00268 | NC_007795.1 mut_A | wt_R | OK | 115.404 | 185.991 |
| SAOUHSC_00764 | NC_007795.1 mut_A | wt_R | OK | 460.14 | 741.551 |
| SAOUHSC_00254 | NC_007795.1 mut_A | wt_R | OK | 5.49939 | 8.85272 |
| SAOUHSC_02337 | NC_007795.1 mut_A | wt_R | OK | 446.517 | 717.425 |
| SAOUHSC_02158 | NC_007795.1 mut_A | wt_R | OK | 389.163 | 623.784 |
| SAOUHSC_00702 | NC_007795.1 mut_A | wt_R | OK | 55.4272 | 88.7287 |
| SAOUHSC_01332 | NC_007795.1 mut_A | wt_R | OK | 258.55 | 413.415 |
| SAOUHSC_01778 | NC_007795.1 mut_A | wt_R | OK | 949.374 | 1516.03 |
| SAOUHSC_02855 | NC_007795.1 mut_A | wt_R | OK | 35.2509 | 56.2418 |
| SAOUHSC_00861 | NC_007795.1 mut_A | wt_R | OK | 686.7 | 1095.22 |
| SAOUHSC_02860 | NC_007795.1 mut_A | wt_R | OK | 557.74 | 886.436 |
| SAOUHSC_03025 | NC_007795.1 mut_A | wt_R | OK | 32.7309 | 52.0014 |
| SAOUHSC_00927 | NC_007795.1 mut_A | wt_R | OK | 60.6698 | 96.3214 |
| SAOUHSC_00383 | NC_007795.1 mut_A | wt_R | OK | 10.8081 | 17.1589 |
| SAOUHSC_00174 | NC_007795.1 mut_A | wt_R | OK | 34.966 | 55.4829 |
| SAOUHSC_02973 | NC_007795.1 mut_A | wt_R | OK | 100.945 | 160.147 |
| SAOUHSC_01191 | NC_007795.1 mut_A | wt_R | OK | 21172.5 | 33583.2 |
| SAOUHSC_02360 | NC_007795.1 mut_A | wt_R | OK | 124.259 | 197.04 |
| SAOUHSC_02338 | NC_007795.1 mut_A | wt_R | OK | 245.064 | 388.336 |
| SAOUHSC_00274 | NC_007795.1 mut_A | wt_R | OK | 55.8264 | 88.3769 |
| SAOUHSC_02345 | NC_007795.1 mut_A | wt_R | OK | 1652.82 | 2616.32 |
| SAOUHSC_02091 | NC_007795.1 mut_A | wt_R | OK | 369.414 | 584.56 |
| SAOUHSC_01066 | NC_007795.1 mut_A | wt_R | OK | 780.183 | 1233.42 |
| SAOUHSC_01368 | NC_007795.1 mut_A | wt_R | OK | 7.85081 | 12.3994 |
| SAOUHSC_02122 | NC_007795.1 mut_A | wt_R | OK | 138.857 | 218.823 |
| SAOUHSC_02123 | NC_007795.1 mut_A | wt_R | OK | 101.342 | 159.613 |
| SAOUHSC_01413 | NC_007795.1 mut_A | wt_R | OK | 132.661 | 207.876 |
| SAOUHSC_01284 | NC_007795.1 mut_A | wt_R | OK | 149.002 | 233.354 |
| SAOUHSC_02351 | NC_007795.1 mut_A | wt_R | OK | 1585.8 | 2482.35 |
| SAOUHSC_02346 | NC_007795.1 mut_A | wt_R | OK | 2139.75 | 3345.7 |
| SAOUHSC_02270 | NC_007795.1 mut_A | wt_R | OK | 4.95394 | 7.74456 |
| SAOUHSC_00503 | NC_007795.1 mut_A | wt_R | OK | 397.396 | 620.897 |
| SAOUHSC_00707 | NC_007795.1 mut_A | wt_R | OK | 353.819 | 552.673 |
| SAOUHSC_02623 | NC_007795.1 mut_A | wt_R | OK | 171.052 | 267.023 |
| SAOUHSC_00526 | NC_007795.1 mut_A | wt_R | OK | 1585.48 | 2473.06 |
| SAOUHSC_02684 | NC_007795.1 mut_A | wt_R | OK | 898.622 | 1400.5 |
| SAOUHSC_01794 | NC_007795.1 mut_A | wt_R | OK | 1422.38 | 2215.56 |
| SAOUHSC_02702 | NC_007795.1 mut_A | wt_R | OK | 7610.1 | 11832.1 |
| SAOUHSC_02883 | NC_007795.1 mut_A | wt_R | OK | 6.95517 | 10.8025 |
| SAOUHSC_01251 | NC_007795.1 mut_A | wt_R | OK | 798.248 | 1239.25 |

| | | | | | |
|---|---|---|---|---|---|
| SAOUHSC_02427 | NC_007795.1 mut_A | wt_R | OK | 53.9464 | 83.611 |
| SAOUHSC_01585 | NC_007795.1 mut_A | wt_R | OK | 610.651 | 946.409 |
| SAUSA300_RS08385 | NC_007795.1 mut_A | wt_R | OK | 412.185 | 638.715 |
| SAOUHSC_03021 | NC_007795.1 mut_A | wt_R | OK | 3.49547 | 5.41549 |
| SAOUHSC_00729 | NC_007795.1 mut_A | wt_R | OK | 85.2317 | 132.048 |
| SAOUHSC_01658 | NC_007795.1 mut_A | wt_R | OK | 348.209 | 539.208 |
| SAOUHSC_02615 | NC_007795.1 mut_A | wt_R | OK | 291.263 | 450.989 |
| SAOUHSC_02641 | NC_007795.1 mut_A | wt_R | OK | 3.95222 | 6.11878 |
| SAOUHSC_02739 | NC_007795.1 mut_A | wt_R | OK | 196.136 | 303.634 |
| SAOUHSC_02700 | NC_007795.1 mut_A | wt_R | OK | 115.295 | 178.452 |
| SAOUHSC_01685 | NC_007795.1 mut_A | wt_R | OK | 440.186 | 681.024 |
| SAOUHSC_00235 | NC_007795.1 mut_A | wt_R | OK | 41.7352 | 64.5038 |
| SAOUHSC_00992 | NC_007795.1 mut_A | wt_R | OK | 11.6798 | 18.0379 |
| SAOUHSC_00050 | NC_007795.1 mut_A | wt_R | OK | 70.7283 | 109.015 |
| SAOUHSC_01042 | NC_007795.1 mut_A | wt_R | OK | 3611.01 | 5560.26 |
| SAOUHSC_02483 | NC_007795.1 mut_A | wt_R | OK | 148.271 | 228.161 |
| SAOUHSC_01897 | NC_007795.1 mut_A | wt_R | OK | 47.2134 | 72.6405 |
| SAOUHSC_00282 | NC_007795.1 mut_A | wt_R | OK | 38.4807 | 59.1468 |
| SAOUHSC_02286 | NC_007795.1 mut_A | wt_R | OK | 14.6579 | 22.5274 |
| SAOUHSC_02529 | NC_007795.1 mut_A | wt_R | OK | 44.3897 | 68.1761 |
| SAOUHSC_01981 | NC_007795.1 mut_A | wt_R | OK | 37.0546 | 56.9101 |
| SAOUHSC_02350 | NC_007795.1 mut_A | wt_R | OK | 2317.87 | 3558.17 |
| SAOUHSC_03041 | NC_007795.1 mut_A | wt_R | OK | 6.40719 | 9.82233 |
| SAOUHSC_01930 | NC_007795.1 mut_A | wt_R | OK | 63.7969 | 97.7 |
| SAOUHSC_01436 | NC_007795.1 mut_A | wt_R | OK | 183.96 | 281.647 |
| SAOUHSC_00730 | NC_007795.1 mut_A | wt_R | OK | 140.096 | 214.347 |
| SAOUHSC_00463 | NC_007795.1 mut_A | wt_R | OK | 145.786 | 222.732 |
| SAOUHSC_02924 | NC_007795.1 mut_A | wt_R | OK | 10.0952 | 15.4068 |
| SAOUHSC_01479 | NC_007795.1 mut_A | wt_R | OK | 122.175 | 186.3 |
| SAOUHSC_02531 | NC_007795.1 mut_A | wt_R | OK | 18.401 | 28.015 |
| SAOUHSC_01874 | NC_007795.1 mut_A | wt_R | OK | 403.709 | 613.468 |
| SAOUHSC_02284 | NC_007795.1 mut_A | wt_R | OK | 11.1596 | 16.9521 |
| SAOUHSC_01389 | NC_007795.1 mut_A | wt_R | OK | 23.1832 | 35.1524 |
| SAOUHSC_01467 | NC_007795.1 mut_A | wt_R | OK | 1132.53 | 1715.11 |
| SAOUHSC_01743 | NC_007795.1 mut_A | wt_R | OK | 360.034 | 545.071 |
| SAOUHSC_01454 | NC_007795.1 mut_A | wt_R | OK | 175.734 | 266.009 |
| SAOUHSC_00661 | NC_007795.1 mut_A | wt_R | OK | 40.2726 | 60.9207 |
| SAOUHSC_02012 | NC_007795.1 mut_A | wt_R | OK | 205.835 | 311.288 |
| SAOUHSC_00451 | NC_007795.1 mut_A | wt_R | OK | 45.5309 | 68.8129 |
| SAOUHSC_00129 | NC_007795.1 mut_A | wt_R | OK | 58.9118 | 88.9062 |
| SAOUHSC_02697 | NC_007795.1 mut_A | wt_R | OK | 1952.5 | 2938.2 |
| SAOUHSC_00014 | NC_007795.1 mut_A | wt_R | OK | 118.834 | 178.791 |
| SAOUHSC_01471 | NC_007795.1 mut_A | wt_R | OK | 1154.35 | 1734.05 |
| SAOUHSC_02894 | NC_007795.1 mut_A | wt_R | OK | 61.9465 | 93.0057 |
| SAOUHSC_02285 | NC_007795.1 mut_A | wt_R | OK | 11.8964 | 17.8378 |
| SAOUHSC_01326 | NC_007795.1 mut_A | wt_R | OK | 93.1811 | 139.635 |
| SAOUHSC_02898 | NC_007795.1 mut_A | wt_R | OK | 28.9455 | 43.372 |
| SAOUHSC_00803 | NC_007795.1 mut_A | wt_R | OK | 381.283 | 571.291 |
| SAOUHSC_01338 | NC_007795.1 mut_A | wt_R | OK | 1282.55 | 1919.54 |
| SAOUHSC_01043 | NC_007795.1 mut_A | wt_R | OK | 5119.66 | 7647.6 |
| SAOUHSC_02752 | NC_007795.1 mut_A | wt_R | OK | 70.9169 | 105.859 |

| | | | | | |
|---|---|---|---|---|---|
| SAOUHSC_01448 | NC_007795.1 mut_A | wt_R | OK | 89.504 | 133.383 |
| SAOUHSC_01395 | NC_007795.1 mut_A | wt_R | OK | 45.3643 | 67.5683 |
| SAOUHSC_01751 | NC_007795.1 mut_A | wt_R | OK | 360.569 | 536.907 |
| SAOUHSC_01753 | NC_007795.1 mut_A | wt_R | OK | 436.329 | 649.492 |
| SAOUHSC_01871 | NC_007795.1 mut_A | wt_R | OK | 218.469 | 324.887 |
| SAOUHSC_01839 | NC_007795.1 mut_A | wt_R | OK | 144.742 | 215.12 |
| SAOUHSC_02836 | NC_007795.1 mut_A | wt_R | OK | 78.948 | 117.271 |
| SAOUHSC_00985 | NC_007795.1 mut_A | wt_R | OK | 1063.38 | 1579.21 |
| SAOUHSC_02867 | NC_007795.1 mut_A | wt_R | OK | 63.3891 | 93.989 |
| SAOUHSC_01767 | NC_007795.1 mut_A | wt_R | OK | 415.449 | 615.807 |
| SAOUHSC_02895 | NC_007795.1 mut_A | wt_R | OK | 30.66 | 45.44 |
| SAOUHSC_01261 | NC_007795.1 mut_A | wt_R | OK | 80.9039 | 119.831 |
| SAOUHSC_01205 | NC_007795.1 mut_A | wt_R | OK | 200.823 | 297.402 |
| SAOUHSC_02983 | NC_007795.1 mut_A | wt_R | OK | 131.084 | 193.936 |
| SAOUHSC_02652 | NC_007795.1 mut_A | wt_R | OK | 235.965 | 349.094 |
| SAOUHSC_01060 | NC_007795.1 mut_A | wt_R | OK | 249.789 | 369.278 |
| SAOUHSC_01742 | NC_007795.1 mut_A | wt_R | OK | 243.818 | 359.423 |
| SAOUHSC_01283 | NC_007795.1 mut_A | wt_R | OK | 76.9871 | 113.412 |
| SAOUHSC_00266 | NC_007795.1 mut_A | wt_R | OK | 104.798 | 154.361 |
| SAOUHSC_00430 | NC_007795.1 mut_A | wt_R | OK | 154.611 | 227.683 |
| SAOUHSC_00671 | NC_007795.1 mut_A | wt_R | OK | 377.117 | 555.263 |
| SAOUHSC_02428 | NC_007795.1 mut_A | wt_R | OK | 36.172 | 53.2392 |
| SAOUHSC_02121 | NC_007795.1 mut_A | wt_R | OK | 201.013 | 295.546 |
| SAOUHSC_01076 | NC_007795.1 mut_A | wt_R | OK | 55.4916 | 81.552 |
| SAOUHSC_00703 | NC_007795.1 mut_A | wt_R | OK | 46.9662 | 68.9919 |
| SAOUHSC_02288 | NC_007795.1 mut_A | wt_R | OK | 11.0556 | 16.2006 |
| SAOUHSC_02524 | NC_007795.1 mut_A | wt_R | OK | 128.732 | 188.63 |
| SAOUHSC_01899 | NC_007795.1 mut_A | wt_R | OK | 52.259 | 76.5211 |
| SAOUHSC_01374 | NC_007795.1 mut_A | wt_R | OK | 470.373 | 688.278 |
| SAOUHSC_01412 | NC_007795.1 mut_A | wt_R | OK | 115.35 | 168.501 |
| SAOUHSC_01196 | NC_007795.1 mut_A | wt_R | OK | 439.134 | 641.196 |
| SAOUHSC_01156 | NC_007795.1 mut_A | wt_R | OK | 114.577 | 167.26 |
| SAOUHSC_02287 | NC_007795.1 mut_A | wt_R | OK | 13.3308 | 19.4377 |
| SAOUHSC_01695 | NC_007795.1 mut_A | wt_R | OK | 155.436 | 226.578 |
| SAOUHSC_01630 | NC_007795.1 mut_A | wt_R | OK | 1428.94 | 2081.06 |
| SAOUHSC_02682 | NC_007795.1 mut_A | wt_R | OK | 1092.07 | 1588.38 |
| SAOUHSC_00346 | NC_007795.1 mut_A | wt_R | OK | 359.494 | 522.703 |
| SAOUHSC_00811 | NC_007795.1 mut_A | wt_R | OK | 44.9333 | 65.2782 |
| SAOUHSC_01050 | NC_007795.1 mut_A | wt_R | OK | 695.405 | 1009.14 |
| SAOUHSC_00106 | NC_007795.1 mut_A | wt_R | OK | 10.8561 | 15.7507 |
| SAOUHSC_01193 | NC_007795.1 mut_A | wt_R | OK | 563.808 | 817.676 |
| SAOUHSC_00317 | NC_007795.1 mut_A | wt_R | OK | 128.698 | 186.409 |
| SAOUHSC_02916 | NC_007795.1 mut_A | wt_R | OK | 460.719 | 666.161 |
| SAOUHSC_00195 | NC_007795.1 mut_A | wt_R | OK | 11.6027 | 16.7678 |
| SAOUHSC_01067 | NC_007795.1 mut_A | wt_R | OK | 623.864 | 901.078 |
| SAOUHSC_02698 | NC_007795.1 mut_A | wt_R | OK | 1731.91 | 2499.39 |
| SAOUHSC_01055a | NC_007795.1 mut_A | wt_R | OK | 207.859 | 299.773 |
| SAOUHSC_01423 | NC_007795.1 mut_A | wt_R | OK | 267.746 | 383.889 |
| SAOUHSC_01188 | NC_007795.1 mut_A | wt_R | OK | 195.556 | 280.363 |
| SAOUHSC_01075 | NC_007795.1 mut_A | wt_R | OK | 279.863 | 401.065 |
| SAOUHSC_00491 | NC_007795.1 mut_A | wt_R | OK | 239.503 | 342.816 |

| | | | | | |
|---|---|---|---|---|---|
| SAOUHSC_01895 | NC_007795.1 mut_A | wt_R | OK | 610.247 | 873.411 |
| SAOUHSC_01698 | NC_007795.1 mut_A | wt_R | OK | 148.519 | 212.538 |
| SAOUHSC_00740 | NC_007795.1 mut_A | wt_R | OK | 219.634 | 313.915 |
| SAOUHSC_02618 | NC_007795.1 mut_A | wt_R | OK | 66.333 | 94.7516 |
| SAOUHSC_01505 | NC_007795.1 mut_A | wt_R | OK | 365.124 | 521.459 |
| SAOUHSC_02936 | NC_007795.1 mut_A | wt_R | OK | 7.7327 | 11.0394 |
| SAOUHSC_00020 | NC_007795.1 mut_A | wt_R | OK | 238.324 | 340.226 |
| SAOUHSC_00725 | NC_007795.1 mut_A | wt_R | OK | 268.487 | 382.818 |
| SAOUHSC_02318 | NC_007795.1 mut_A | wt_R | OK | 428.981 | 610.831 |
| SAOUHSC_00404 | NC_007795.1 mut_A | wt_R | OK | 73.7193 | 104.942 |
| SAOUHSC_00412 | NC_007795.1 mut_A | wt_R | OK | 117.681 | 167.495 |
| SAOUHSC_02614 | NC_007795.1 mut_A | wt_R | OK | 639.528 | 910.043 |
| SAOUHSC_00663 | NC_007795.1 mut_A | wt_R | OK | 185.175 | 263.208 |
| SAOUHSC_01768 | NC_007795.1 mut_A | wt_R | OK | 263.087 | 373.515 |
| SAOUHSC_01158 | NC_007795.1 mut_A | wt_R | OK | 1246.09 | 1767.89 |
| SAOUHSC_02358 | NC_007795.1 mut_A | wt_R | OK | 300.989 | 426.591 |
| SAOUHSC_00393 | NC_007795.1 mut_A | wt_R | OK | 7.51455 | 10.6462 |
| SAOUHSC_01793 | NC_007795.1 mut_A | wt_R | OK | 745.396 | 1055.39 |
| SAOUHSC_00167 | NC_007795.1 mut_A | wt_R | OK | 57.5928 | 81.4775 |
| SAOUHSC_00925 | NC_007795.1 mut_A | wt_R | OK | 57.3863 | 81.0637 |
| SAOUHSC_02336 | NC_007795.1 mut_A | wt_R | OK | 707.472 | 999.113 |
| SAOUHSC_02798 | NC_007795.1 mut_A | wt_R | OK | 30.57 | 43.0796 |
| SAOUHSC_02668 | NC_007795.1 mut_A | wt_R | OK | 235.891 | 332.391 |
| SAOUHSC_02426 | NC_007795.1 mut_A | wt_R | OK | 66.631 | 93.7901 |
| SAOUHSC_00801 | NC_007795.1 mut_A | wt_R | OK | 671.56 | 944.683 |
| SAOUHSC_01103 | NC_007795.1 mut_A | wt_R | OK | 763.34 | 1073.33 |
| SAOUHSC_00275 | NC_007795.1 mut_A | wt_R | OK | 22.4265 | 31.5134 |
| SAOUHSC_02915 | NC_007795.1 mut_A | wt_R | OK | 31.5466 | 44.3183 |
| SAOUHSC_02597 | NC_007795.1 mut_A | wt_R | OK | 90.0265 | 126.373 |
| SAOUHSC_00019 | NC_007795.1 mut_A | wt_R | OK | 200.317 | 281.189 |
| SAOUHSC_01028 | NC_007795.1 mut_A | wt_R | OK | 6255.19 | 8765.62 |
| SAOUHSC_01766 | NC_007795.1 mut_A | wt_R | OK | 506.017 | 709.09 |
| SAOUHSC_02140 | NC_007795.1 mut_A | wt_R | OK | 1612.37 | 2258.68 |
| SAOUHSC_02517 | NC_007795.1 mut_A | wt_R | OK | 93.8195 | 131.359 |
| SAOUHSC_00876 | NC_007795.1 mut_A | wt_R | OK | 763.212 | 1068.49 |
| SAOUHSC_02912 | NC_007795.1 mut_A | wt_R | OK | 277.337 | 388.177 |
| SAOUHSC_01149 | NC_007795.1 mut_A | wt_R | OK | 1565.63 | 2189.85 |
| SAOUHSC_00395 | NC_007795.1 mut_A | wt_R | OK | 14.3614 | 20.0774 |
| SAOUHSC_00738 | NC_007795.1 mut_A | wt_R | OK | 897.594 | 1253.42 |
| SAOUHSC_00462 | NC_007795.1 mut_A | wt_R | OK | 813.78 | 1136.24 |
| SAOUHSC_01634 | NC_007795.1 mut_A | wt_R | OK | 154.561 | 215.735 |
| SAOUHSC_02982 | NC_007795.1 mut_A | wt_R | OK | 537.734 | 750.412 |
| SAOUHSC_02625 | NC_007795.1 mut_A | wt_R | OK | 148.988 | 207.794 |
| SAOUHSC_02984 | NC_007795.1 mut_A | wt_R | OK | 92.2427 | 128.622 |
| SAOUHSC_00335 | NC_007795.1 mut_A | wt_R | OK | 12.9943 | 18.1079 |
| SAOUHSC_02113 | NC_007795.1 mut_A | wt_R | OK | 79.2689 | 110.414 |
| SAOUHSC_02527 | NC_007795.1 mut_A | wt_R | OK | 228.244 | 317.676 |
| SAOUHSC_00926 | NC_007795.1 mut_A | wt_R | OK | 74.0487 | 103.055 |
| SAOUHSC_02911 | NC_007795.1 mut_A | wt_R | OK | 51.0019 | 70.836 |
| SAOUHSC_02013 | NC_007795.1 mut_A | wt_R | OK | 488.548 | 677.679 |
| SAOUHSC_00301 | NC_007795.1 mut_A | wt_R | OK | 62.3701 | 86.4718 |

| | | | | | |
|---|---|---|---|---|---|
| SAOUHSC_01980 | NC_007795.1 mut_A | wt_R | OK | 68.9859 | 95.584 |
| SAOUHSC_01394 | NC_007795.1 mut_A | wt_R | OK | 53.1221 | 73.5696 |
| SAOUHSC_02536 | NC_007795.1 mut_A | wt_R | OK | 287.093 | 397.549 |
| SAOUHSC_02669 | NC_007795.1 mut_A | wt_R | OK | 1037.71 | 1436.85 |
| SAOUHSC_00471 | NC_007795.1 mut_A | wt_R | OK | 506.091 | 700.733 |
| SAOUHSC_01061 | NC_007795.1 mut_A | wt_R | OK | 180.535 | 249.707 |
| SAOUHSC_02110 | NC_007795.1 mut_A | wt_R | OK | 207.891 | 287.462 |
| SAOUHSC_01660 | NC_007795.1 mut_A | wt_R | OK | 117.259 | 161.926 |
| SAOUHSC_00666 | NC_007795.1 mut_A | wt_R | OK | 63.1048 | 87.0949 |
| SAOUHSC_00504 | NC_007795.1 mut_A | wt_R | OK | 365.758 | 504.313 |
| SAOUHSC_00402 | NC_007795.1 mut_A | wt_R | OK | 20.8367 | 28.7175 |
| SAOUHSC_01029 | NC_007795.1 mut_A | wt_R | OK | 3756.49 | 5169.1 |
| SAOUHSC_01480 | NC_007795.1 mut_A | wt_R | OK | 290.407 | 399.575 |
| SAOUHSC_03020 | NC_007795.1 mut_A | wt_R | OK | 13.3297 | 18.34 |
| SAOUHSC_01044 | NC_007795.1 mut_A | wt_R | OK | 144.911 | 199.147 |
| SAOUHSC_02000 | NC_007795.1 mut_A | wt_R | OK | 200.249 | 275.162 |
| SAOUHSC_02449 | NC_007795.1 mut_A | wt_R | OK | 11.0466 | 15.1651 |
| SAOUHSC_00006 | NC_007795.1 mut_A | wt_R | OK | 592.926 | 813.88 |
| SAOUHSC_01350 | NC_007795.1 mut_A | wt_R | OK | 484.452 | 664.954 |
| SAOUHSC_00533 | NC_007795.1 mut_A | wt_R | OK | 1634.7 | 2243.03 |
| SAOUHSC_00024 | NC_007795.1 mut_A | wt_R | OK | 53.862 | 73.8732 |
| SAOUHSC_02281 | NC_007795.1 mut_A | wt_R | OK | 11.6113 | 15.8965 |
| SAOUHSC_02114 | NC_007795.1 mut_A | wt_R | OK | 324.916 | 444.319 |
| SAOUHSC_00013 | NC_007795.1 mut_A | wt_R | OK | 31.3026 | 42.7988 |
| SAOUHSC_01269 | NC_007795.1 mut_A | wt_R | OK | 257.046 | 351.237 |
| SAOUHSC_00464 | NC_007795.1 mut_A | wt_R | OK | 90.5125 | 123.637 |
| SAOUHSC_01709 | NC_007795.1 mut_A | wt_R | OK | 44.8171 | 61.0936 |
| SAOUHSC_00708 | NC_007795.1 mut_A | wt_R | OK | 691.041 | 941.785 |
| SAOUHSC_00334 | NC_007795.1 mut_A | wt_R | OK | 184.268 | 251.021 |
| SAOUHSC_01010 | NC_007795.1 mut_A | wt_R | OK | 56.6393 | 77.1546 |
| SAOUHSC_00755 | NC_007795.1 mut_A | wt_R | OK | 326.168 | 443.786 |
| SAOUHSC_00414 | NC_007795.1 mut_A | wt_R | OK | 492.599 | 669.5 |
| SAOUHSC_01750 | NC_007795.1 mut_A | wt_R | OK | 483.475 | 655.857 |
| SAOUHSC_00229 | NC_007795.1 mut_A | wt_R | OK | 470.154 | 637.213 |
| SAOUHSC_00804 | NC_007795.1 mut_A | wt_R | OK | 427.383 | 579.055 |
| SAOUHSC_01150 | NC_007795.1 mut_A | wt_R | OK | 2572.3 | 3482.97 |
| SAOUHSC_01248 | NC_007795.1 mut_A | wt_R | OK | 65.5783 | 88.7761 |
| SAOUHSC_01363 | NC_007795.1 mut_A | wt_R | OK | 41.7913 | 56.5617 |
| SAOUHSC_02499 | NC_007795.1 mut_A | wt_R | OK | 158.058 | 213.825 |
| SAOUHSC_00457 | NC_007795.1 mut_A | wt_R | OK | 46.972 | 63.5155 |
| SAOUHSC_01747 | NC_007795.1 mut_A | wt_R | OK | 903.434 | 1221.35 |
| SAOUHSC_02525 | NC_007795.1 mut_A | wt_R | OK | 119.695 | 161.678 |
| SAOUHSC_00885 | NC_007795.1 mut_A | wt_R | OK | 235.626 | 317.759 |
| SAOUHSC_02737 | NC_007795.1 mut_A | wt_R | OK | 284.9 | 383.834 |
| SAOUHSC_01072 | NC_007795.1 mut_A | wt_R | OK | 99.7483 | 134.352 |
| SAOUHSC_02861 | NC_007795.1 mut_A | wt_R | OK | 301.966 | 406.706 |
| SAOUHSC_01110 | NC_007795.1 mut_A | wt_R | OK | 271.022 | 364.838 |
| SAOUHSC_01201 | NC_007795.1 mut_A | wt_R | OK | 5727.73 | 7680.41 |
| SAOUHSC_02660 | NC_007795.1 mut_A | wt_R | OK | 145.491 | 194.901 |
| SAOUHSC_03001 | NC_007795.1 mut_A | wt_R | OK | 211.512 | 283.313 |
| SAOUHSC_00739 | NC_007795.1 mut_A | wt_R | OK | 313.997 | 420.293 |

| | | | | | |
|---|---|---|---|---|---|
| SAOUHSC_01192 | NC_007795.1 mut_A | wt_R | OK | 345.789 | 462.762 |
| SAOUHSC_01497 | NC_007795.1 mut_A | wt_R | OK | 192.411 | 257.49 |
| SAOUHSC_00903 | NC_007795.1 mut_A | wt_R | OK | 340.588 | 455.786 |
| SAOUHSC_01361 | NC_007795.1 mut_A | wt_R | OK | 296.502 | 396.721 |
| SAOUHSC_00536 | NC_007795.1 mut_A | wt_R | OK | 317.142 | 424.245 |
| SAOUHSC_01035 | NC_007795.1 mut_A | wt_R | OK | 738.705 | 988.019 |
| SAOUHSC_02382 | NC_007795.1 mut_A | wt_R | OK | 403.029 | 538.467 |
| SAOUHSC_02927 | NC_007795.1 mut_A | wt_R | OK | 6286.87 | 8390.23 |
| SAOUHSC_01038 | NC_007795.1 mut_A | wt_R | OK | 449.141 | 599.303 |
| SAOUHSC_01373 | NC_007795.1 mut_A | wt_R | OK | 185.37 | 247.31 |
| SAOUHSC_02460 | NC_007795.1 mut_A | wt_R | OK | 117.901 | 157.209 |
| SAOUHSC_00276 | NC_007795.1 mut_A | wt_R | OK | 19.1363 | 25.5018 |
| SAOUHSC_00878 | NC_007795.1 mut_A | wt_R | OK | 2009.98 | 2674.15 |
| SAOUHSC_01749 | NC_007795.1 mut_A | wt_R | OK | 355.911 | 473.461 |
| SAOUHSC_01249 | NC_007795.1 mut_A | wt_R | OK | 232.476 | 308.743 |
| SAOUHSC_01450 | NC_007795.1 mut_A | wt_R | OK | 215.352 | 285.546 |
| SAOUHSC_02901 | NC_007795.1 mut_A | wt_R | OK | 11.8271 | 15.6712 |
| SAOUHSC_01320 | NC_007795.1 mut_A | wt_R | OK | 29.0142 | 38.4324 |
| SAOUHSC_00683 | NC_007795.1 mut_A | wt_R | OK | 23.6255 | 31.2903 |
| SAOUHSC_02624 | NC_007795.1 mut_A | wt_R | OK | 70.2471 | 92.8894 |
| SAOUHSC_00150 | NC_007795.1 mut_A | wt_R | OK | 12.957 | 17.1302 |
| SAOUHSC_00854 | NC_007795.1 mut_A | wt_R | OK | 44.4296 | 58.6888 |
| SAOUHSC_02920 | NC_007795.1 mut_A | wt_R | OK | 144.111 | 190.238 |
| SAOUHSC_01806 | NC_007795.1 mut_A | wt_R | OK | 2825.24 | 3721.62 |
| SAOUHSC_00502 | NC_007795.1 mut_A | wt_R | OK | 510.624 | 672.441 |
| SAOUHSC_00021 | NC_007795.1 mut_A | wt_R | OK | 373.587 | 491.388 |
| SAOUHSC_01788 | NC_007795.1 mut_A | wt_R | OK | 1011.64 | 1329.86 |
| SAOUHSC_00891 | NC_007795.1 mut_A | wt_R | OK | 333.947 | 438.915 |
| SAOUHSC_01752 | NC_007795.1 mut_A | wt_R | OK | 487.192 | 639.591 |
| SAOUHSC_00874 | NC_007795.1 mut_A | wt_R | OK | 207.325 | 272.092 |
| SAOUHSC_00828 | NC_007795.1 mut_A | wt_R | OK | 21.4352 | 28.1047 |
| SAOUHSC_02274 | NC_007795.1 mut_A | wt_R | OK | 64.8073 | 84.9221 |
| SAOUHSC_00445 | NC_007795.1 mut_A | wt_R | OK | 852.871 | 1116.37 |
| SAOUHSC_01104 | NC_007795.1 mut_A | wt_R | OK | 1065.88 | 1394.98 |
| SAOUHSC_02874 | NC_007795.1 mut_A | wt_R | OK | 207.793 | 271.388 |
| SAOUHSC_01857 | NC_007795.1 mut_A | wt_R | OK | 227.316 | 296.827 |
| SAOUHSC_00155 | NC_007795.1 mut_A | wt_R | OK | 616.48 | 803.868 |
| SAOUHSC_00980 | NC_007795.1 mut_A | wt_R | OK | 267.243 | 348.168 |
| SAOUHSC_00022 | NC_007795.1 mut_A | wt_R | OK | 456.028 | 593.935 |
| SAOUHSC_00302 | NC_007795.1 mut_A | wt_R | OK | 18.0779 | 23.5424 |
| SAOUHSC_01779 | NC_007795.1 mut_A | wt_R | OK | 2683.31 | 3493.18 |
| SAOUHSC_01813 | NC_007795.1 mut_A | wt_R | OK | 92.6012 | 120.399 |
| SAOUHSC_01762 | NC_007795.1 mut_A | wt_R | OK | 650.155 | 844.661 |
| SAOUHSC_00962 | NC_007795.1 mut_A | wt_R | OK | 1590.55 | 2065.35 |
| SAOUHSC_00423 | NC_007795.1 mut_A | wt_R | OK | 34.2614 | 44.4777 |
| SAOUHSC_02480 | NC_007795.1 mut_A | wt_R | OK | 191.27 | 247.996 |
| SAOUHSC_01964 | NC_007795.1 mut_A | wt_R | OK | 2419.88 | 3136.95 |
| SAOUHSC_01282 | NC_007795.1 mut_A | wt_R | OK | 467.559 | 606.012 |
| SAOUHSC_03033 | NC_007795.1 mut_A | wt_R | OK | 15.5905 | 20.2062 |
| SAOUHSC_00264 | NC_007795.1 mut_A | wt_R | OK | 53.5157 | 69.3202 |
| SAOUHSC_00691 | NC_007795.1 mut_A | wt_R | OK | 151.025 | 195.574 |

| | | | | | | |
|---|---|---|---|---|---|---|
| SAOUHSC_02131 | NC_007795.1 | mut_A | wt_R | OK | 513.572 | 664.925 |
| SAOUHSC_00886 | NC_007795.1 | mut_A | wt_R | OK | 174.677 | 225.861 |
| SAOUHSC_01812 | NC_007795.1 | mut_A | wt_R | OK | 231.072 | 298.458 |
| SAOUHSC_01843 | NC_007795.1 | mut_A | wt_R | OK | 7.90036 | 10.198 |
| SAOUHSC_02370 | NC_007795.1 | mut_A | wt_R | OK | 58.827 | 75.9282 |
| SAOUHSC_00949 | NC_007795.1 | mut_A | wt_R | OK | 10.1033 | 13.0363 |
| SAOUHSC_01483 | NC_007795.1 | mut_A | wt_R | OK | 108.567 | 140.059 |
| SAOUHSC_03016 | NC_007795.1 | mut_A | wt_R | OK | 22.3819 | 28.8126 |
| SAOUHSC_00018 | NC_007795.1 | mut_A | wt_R | OK | 443.236 | 569.261 |
| SAOUHSC_00083 | NC_007795.1 | mut_A | wt_R | OK | 21.6585 | 27.8003 |
| SAOUHSC_02591 | NC_007795.1 | mut_A | wt_R | OK | 167.349 | 214.784 |
| SAOUHSC_01652 | NC_007795.1 | mut_A | wt_R | OK | 265.69 | 340.51 |
| SAOUHSC_01018 | NC_007795.1 | mut_A | wt_R | OK | 439.69 | 563.455 |
| SAOUHSC_00734 | NC_007795.1 | mut_A | wt_R | OK | 47.4168 | 60.6735 |
| SAOUHSC_02383 | NC_007795.1 | mut_A | wt_R | OK | 284.855 | 364.235 |
| SAOUHSC_02538 | NC_007795.1 | mut_A | wt_R | OK | 373.385 | 477.088 |
| SAOUHSC_01398 | NC_007795.1 | mut_A | wt_R | OK | 58.7058 | 74.9821 |
| SAOUHSC_01098 | NC_007795.1 | mut_A | wt_R | OK | 172.278 | 219.965 |
| SAOUHSC_02627 | NC_007795.1 | mut_A | wt_R | OK | 439.257 | 559.947 |
| SAOUHSC_01748 | NC_007795.1 | mut_A | wt_R | OK | 374.79 | 477.72 |
| SAOUHSC_00486 | NC_007795.1 | mut_A | wt_R | OK | 929.831 | 1184.94 |
| SAOUHSC_00620 | NC_007795.1 | mut_A | wt_R | OK | 1538.51 | 1960.58 |
| SAOUHSC_00934 | NC_007795.1 | mut_A | wt_R | OK | 11914.5 | 15181 |
| SAOUHSC_00872 | NC_007795.1 | mut_A | wt_R | OK | 1086.31 | 1383.65 |
| SAOUHSC_02613 | NC_007795.1 | mut_A | wt_R | OK | 93.8043 | 118.827 |
| SAOUHSC_00718 | NC_007795.1 | mut_A | wt_R | OK | 130.626 | 165.268 |
| SAOUHSC_00281 | NC_007795.1 | mut_A | wt_R | OK | 614.9 | 776.442 |
| SAOUHSC_01657 | NC_007795.1 | mut_A | wt_R | OK | 223.621 | 282.104 |
| SAOUHSC_01396 | NC_007795.1 | mut_A | wt_R | OK | 60.2317 | 75.9643 |
| SAOUHSC_02826 | NC_007795.1 | mut_A | wt_R | OK | 97.2011 | 122.483 |
| SAOUHSC_01093 | NC_007795.1 | mut_A | wt_R | OK | 361.857 | 455.865 |
| SAOUHSC_00948 | NC_007795.1 | mut_A | wt_R | OK | 343.565 | 432.652 |
| SAOUHSC_01896 | NC_007795.1 | mut_A | wt_R | OK | 553.676 | 696.4 |
| SAOUHSC_01106 | NC_007795.1 | mut_A | wt_R | OK | 206.682 | 259.224 |
| SAOUHSC_01185 | NC_007795.1 | mut_A | wt_R | OK | 755.526 | 947.125 |
| SAOUHSC_02537 | NC_007795.1 | mut_A | wt_R | OK | 243.858 | 305.086 |
| SAOUHSC_01073 | NC_007795.1 | mut_A | wt_R | OK | 145.558 | 182.044 |
| SAOUHSC_00909 | NC_007795.1 | mut_A | wt_R | OK | 148.366 | 185.55 |
| SAOUHSC_00999 | NC_007795.1 | mut_A | wt_R | OK | 7622.38 | 9531.64 |
| SAOUHSC_00461 | NC_007795.1 | mut_A | wt_R | OK | 830.306 | 1034.83 |
| SAOUHSC_00959 | NC_007795.1 | mut_A | wt_R | OK | 220.818 | 275.15 |
| SAOUHSC_00049 | NC_007795.1 | mut_A | wt_R | OK | 36.1584 | 44.9779 |
| SAOUHSC_00569 | NC_007795.1 | mut_A | wt_R | OK | 292.691 | 363.723 |
| SAOUHSC_02573 | NC_007795.1 | mut_A | wt_R | OK | 94.1177 | 116.816 |
| SAOUHSC_00847 | NC_007795.1 | mut_A | wt_R | OK | 356.16 | 441.141 |
| SAOUHSC_01271 | NC_007795.1 | mut_A | wt_R | OK | 574.937 | 710.558 |
| SAOUHSC_00795 | NC_007795.1 | mut_A | wt_R | OK | 19599.1 | 20071.8 |
| SAOUHSC_01263 | NC_007795.1 | mut_A | wt_R | OK | 913.108 | 932.541 |
| SAOUHSC_03001 | '95.1:2774449 | mut_A | wt_R | OK | 211.512 | 283.313 |

| log2(fold_cha | test_stat | p_value | q_value | significant | FC | Log2fC |
|---|---|---|---|---|---|---|
| 4.05615 | 16.7546 | 5.00E-05 | 0.00013743 | yes | 16.635036 | 4.05615308 |
| 3.59272 | 6.52882 | 5.00E-05 | 0.00013743 | yes | | |
| 2.96971 | 12.1867 | 5.00E-05 | 0.00013743 | yes | | |
| 2.91482 | 9.4386 | 5.00E-05 | 0.00013743 | yes | | |
| 2.57175 | 3.83527 | 5.00E-05 | 0.00013743 | yes | | |
| 2.42408 | 9.88015 | 5.00E-05 | 0.00013743 | yes | | |
| 2.38668 | 9.51847 | 5.00E-05 | 0.00013743 | yes | | |
| 2.35541 | 9.65368 | 5.00E-05 | 0.00013743 | yes | | |
| 2.30318 | 9.48112 | 5.00E-05 | 0.00013743 | yes | | |
| 2.29967 | 9.41993 | 5.00E-05 | 0.00013743 | yes | | |
| 2.28551 | 7.39334 | 5.00E-05 | 0.00013743 | yes | | |
| 2.24937 | 9.18681 | 5.00E-05 | 0.00013743 | yes | | |
| 2.21847 | 8.0579 | 5.00E-05 | 0.00013743 | yes | | |
| 2.21211 | 9.06692 | 5.00E-05 | 0.00013743 | yes | | |
| 2.19942 | 9.07467 | 5.00E-05 | 0.00013743 | yes | | |
| 2.19354 | 8.63719 | 5.00E-05 | 0.00013743 | yes | | |
| 2.14562 | 8.81164 | 5.00E-05 | 0.00013743 | yes | | |
| 2.10923 | 8.42071 | 5.00E-05 | 0.00013743 | yes | | |
| 2.02095 | 8.28988 | 5.00E-05 | 0.00013743 | yes | | |
| 1.97299 | 8.07524 | 5.00E-05 | 0.00013743 | yes | | |
| 1.94869 | 7.60235 | 5.00E-05 | 0.00013743 | yes | | |
| 1.92159 | 7.11941 | 5.00E-05 | 0.00013743 | yes | | |
| 1.91458 | 6.60275 | 5.00E-05 | 0.00013743 | yes | | |
| 1.81189 | 7.39008 | 5.00E-05 | 0.00013743 | yes | | |
| 1.76295 | 6.00004 | 5.00E-05 | 0.00013743 | yes | | |
| 1.74729 | 7.21221 | 5.00E-05 | 0.00013743 | yes | | |
| 1.72449 | 3.49157 | 5.00E-05 | 0.00013743 | yes | | |
| 1.7218 | 7.06309 | 5.00E-05 | 0.00013743 | yes | | |
| 1.7201 | 6.95518 | 5.00E-05 | 0.00013743 | yes | | |
| 1.70237 | 7.01478 | 5.00E-05 | 0.00013743 | yes | | |
| 1.68776 | 5.3864 | 5.00E-05 | 0.00013743 | yes | | |
| 1.6825 | 6.26348 | 5.00E-05 | 0.00013743 | yes | | |
| 1.66044 | 6.83752 | 5.00E-05 | 0.00013743 | yes | | |
| 1.62975 | 5.52191 | 5.00E-05 | 0.00013743 | yes | | |
| 1.60808 | 5.67082 | 5.00E-05 | 0.00013743 | yes | | |
| 1.60037 | 2.19515 | 0.0001 | 0.00026595 | yes | | |
| 1.59514 | 6.39026 | 5.00E-05 | 0.00013743 | yes | | |
| 1.59474 | 6.53893 | 5.00E-05 | 0.00013743 | yes | | |
| 1.5883 | 6.38343 | 5.00E-05 | 0.00013743 | yes | | |
| 1.57985 | 6.48266 | 5.00E-05 | 0.00013743 | yes | | |
| 1.54449 | 6.07791 | 5.00E-05 | 0.00013743 | yes | | |
| 1.53742 | 5.86864 | 5.00E-05 | 0.00013743 | yes | | |
| 1.53375 | 5.29532 | 5.00E-05 | 0.00013743 | yes | | |
| 1.51939 | 6.23198 | 5.00E-05 | 0.00013743 | yes | | |
| 1.51269 | 2.89536 | 5.00E-05 | 0.00013743 | yes | | |
| 1.5046 | 2.26334 | 0.0013 | 0.00286816 | yes | | |
| 1.4981 | 4.85983 | 5.00E-05 | 0.00013743 | yes | | |
| 1.49531 | 6.16067 | 5.00E-05 | 0.00013743 | yes | | |
| 1.46101 | 5.01725 | 5.00E-05 | 0.00013743 | yes | | |
| 1.45459 | 3.94188 | 5.00E-05 | 0.00013743 | yes | | |

| | | | | |
|---|---|---|---|---|
| 1.45276 | 5.14843 | 5.00E-05 | 0.00013743 | yes |
| 1.44721 | 5.67875 | 5.00E-05 | 0.00013743 | yes |
| 1.44685 | 5.94418 | 5.00E-05 | 0.00013743 | yes |
| 1.44209 | 5.62358 | 5.00E-05 | 0.00013743 | yes |
| 1.43887 | 5.89184 | 5.00E-05 | 0.00013743 | yes |
| 1.4343 | 5.18371 | 5.00E-05 | 0.00013743 | yes |
| 1.4269 | 5.83874 | 5.00E-05 | 0.00013743 | yes |
| 1.41834 | 3.66083 | 5.00E-05 | 0.00013743 | yes |
| 1.41303 | 5.84099 | 5.00E-05 | 0.00013743 | yes |
| 1.40617 | 3.03866 | 5.00E-05 | 0.00013743 | yes |
| 1.40294 | 5.73269 | 5.00E-05 | 0.00013743 | yes |
| 1.40019 | 4.91863 | 5.00E-05 | 0.00013743 | yes |
| 1.39612 | 2.48466 | 0.0001 | 0.00026595 | yes |
| 1.39501 | 2.73029 | 5.00E-05 | 0.00013743 | yes |
| 1.3905 | 2.02644 | 0.00225 | 0.00474293 | yes |
| 1.38895 | 5.46694 | 5.00E-05 | 0.00013743 | yes |
| 1.37912 | 5.63973 | 5.00E-05 | 0.00013743 | yes |
| 1.36194 | 5.60737 | 5.00E-05 | 0.00013743 | yes |
| 1.3576 | 3.02213 | 5.00E-05 | 0.00013743 | yes |
| 1.35606 | 5.54732 | 5.00E-05 | 0.00013743 | yes |
| 1.35167 | 5.55738 | 5.00E-05 | 0.00013743 | yes |
| 1.35131 | 4.36212 | 5.00E-05 | 0.00013743 | yes |
| 1.34726 | 5.53577 | 5.00E-05 | 0.00013743 | yes |
| 1.34201 | 5.52181 | 5.00E-05 | 0.00013743 | yes |
| 1.3419 | 5.44503 | 5.00E-05 | 0.00013743 | yes |
| 1.33625 | 3.39921 | 5.00E-05 | 0.00013743 | yes |
| 1.33452 | 5.50516 | 5.00E-05 | 0.00013743 | yes |
| 1.32952 | 5.01976 | 5.00E-05 | 0.00013743 | yes |
| 1.32036 | 4.00563 | 5.00E-05 | 0.00013743 | yes |
| 1.31761 | 5.40751 | 5.00E-05 | 0.00013743 | yes |
| 1.31434 | 2.51246 | 5.00E-05 | 0.00013743 | yes |
| 1.31357 | 5.41649 | 5.00E-05 | 0.00013743 | yes |
| 1.30614 | 3.29071 | 5.00E-05 | 0.00013743 | yes |
| 1.30595 | 3.48398 | 5.00E-05 | 0.00013743 | yes |
| 1.30224 | 5.34644 | 5.00E-05 | 0.00013743 | yes |
| 1.2976 | 4.91415 | 5.00E-05 | 0.00013743 | yes |
| 1.29744 | 4.56728 | 5.00E-05 | 0.00013743 | yes |
| 1.29547 | 5.31506 | 5.00E-05 | 0.00013743 | yes |
| 1.29512 | 4.06119 | 5.00E-05 | 0.00013743 | yes |
| 1.29377 | 5.2142 | 5.00E-05 | 0.00013743 | yes |
| 1.29137 | 5.32807 | 5.00E-05 | 0.00013743 | yes |
| 1.29062 | 4.40669 | 5.00E-05 | 0.00013743 | yes |
| 1.28829 | 1.79576 | 0.0027 | 0.00560506 | yes |
| 1.28761 | 4.3275 | 5.00E-05 | 0.00013743 | yes |
| 1.28703 | 5.26737 | 5.00E-05 | 0.00013743 | yes |
| 1.28433 | 1.48192 | 0.0125 | 0.0229993 | yes |
| 1.27965 | 2.95666 | 5.00E-05 | 0.00013743 | yes |
| 1.27837 | 5.24631 | 5.00E-05 | 0.00013743 | yes |
| 1.27262 | 2.83687 | 0.0001 | 0.00026595 | yes |
| 1.26599 | 5.00924 | 5.00E-05 | 0.00013743 | yes |
| 1.26201 | 2.26389 | 5.00E-05 | 0.00013743 | yes |

| | | | | |
|---|---|---|---|---|
| 1.2585 | 5.20265 | 5.00E-05 | 0.00013743 | yes |
| 1.25109 | 4.35928 | 5.00E-05 | 0.00013743 | yes |
| 1.24569 | 3.05504 | 5.00E-05 | 0.00013743 | yes |
| 1.24425 | 4.88304 | 5.00E-05 | 0.00013743 | yes |
| 1.23923 | 5.12396 | 5.00E-05 | 0.00013743 | yes |
| 1.23768 | 5.08224 | 5.00E-05 | 0.00013743 | yes |
| 1.23474 | 4.99756 | 5.00E-05 | 0.00013743 | yes |
| 1.23468 | 3.16437 | 5.00E-05 | 0.00013743 | yes |
| 1.22537 | 4.00708 | 5.00E-05 | 0.00013743 | yes |
| 1.21833 | 4.00362 | 5.00E-05 | 0.00013743 | yes |
| 1.21555 | 1.46967 | 0.0139 | 0.0253101 | yes |
| 1.21127 | 4.95389 | 5.00E-05 | 0.00013743 | yes |
| 1.21059 | 4.99941 | 5.00E-05 | 0.00013743 | yes |
| 1.21052 | 4.93462 | 5.00E-05 | 0.00013743 | yes |
| 1.20983 | 2.56394 | 5.00E-05 | 0.00013743 | yes |
| 1.20923 | 4.95004 | 5.00E-05 | 0.00013743 | yes |
| 1.2091 | 4.99665 | 5.00E-05 | 0.00013743 | yes |
| 1.20115 | 4.9439 | 5.00E-05 | 0.00013743 | yes |
| 1.19935 | 4.93328 | 5.00E-05 | 0.00013743 | yes |
| 1.19783 | 4.79421 | 5.00E-05 | 0.00013743 | yes |
| 1.19708 | 4.38229 | 5.00E-05 | 0.00013743 | yes |
| 1.19299 | 4.91142 | 5.00E-05 | 0.00013743 | yes |
| 1.18686 | 4.81427 | 5.00E-05 | 0.00013743 | yes |
| 1.18263 | 4.83438 | 5.00E-05 | 0.00013743 | yes |
| 1.1752 | 4.74813 | 5.00E-05 | 0.00013743 | yes |
| 1.17298 | 4.81642 | 5.00E-05 | 0.00013743 | yes |
| 1.17091 | 4.26097 | 5.00E-05 | 0.00013743 | yes |
| 1.16717 | 4.74383 | 5.00E-05 | 0.00013743 | yes |
| 1.162 | 4.0161 | 5.00E-05 | 0.00013743 | yes |
| 1.15935 | 4.72574 | 5.00E-05 | 0.00013743 | yes |
| 1.15799 | 4.72632 | 5.00E-05 | 0.00013743 | yes |
| 1.15751 | 4.61487 | 5.00E-05 | 0.00013743 | yes |
| 1.15529 | 4.75366 | 5.00E-05 | 0.00013743 | yes |
| 1.15282 | 4.76996 | 5.00E-05 | 0.00013743 | yes |
| 1.15091 | 2.32886 | 0.0001 | 0.00026595 | yes |
| 1.1465 | 4.68488 | 5.00E-05 | 0.00013743 | yes |
| 1.14137 | 4.68639 | 5.00E-05 | 0.00013743 | yes |
| 1.1363 | 4.59347 | 5.00E-05 | 0.00013743 | yes |
| 1.13512 | 4.63646 | 5.00E-05 | 0.00013743 | yes |
| 1.12881 | 4.43412 | 5.00E-05 | 0.00013743 | yes |
| 1.12758 | 1.33692 | 0.02935 | 0.049897 | yes |
| 1.12458 | 4.64414 | 5.00E-05 | 0.00013743 | yes |
| 1.12139 | 3.69585 | 5.00E-05 | 0.00013743 | yes |
| 1.11895 | 4.59475 | 5.00E-05 | 0.00013743 | yes |
| 1.11231 | 4.54436 | 5.00E-05 | 0.00013743 | yes |
| 1.11043 | 4.52356 | 5.00E-05 | 0.00013743 | yes |
| 1.10789 | 4.43283 | 5.00E-05 | 0.00013743 | yes |
| 1.10316 | 2.89498 | 5.00E-05 | 0.00013743 | yes |
| 1.10074 | 4.34642 | 5.00E-05 | 0.00013743 | yes |
| 1.10071 | 3.85432 | 5.00E-05 | 0.00013743 | yes |
| 1.0946 | 4.41287 | 5.00E-05 | 0.00013743 | yes |

| | | | | |
|---|---|---|---|---|
| 1.09439 | 4.2293 | 5.00E-05 | 0.00013743 | yes |
| 1.09117 | 3.80071 | 5.00E-05 | 0.00013743 | yes |
| 1.08905 | 3.01221 | 5.00E-05 | 0.00013743 | yes |
| 1.08555 | 4.31089 | 5.00E-05 | 0.00013743 | yes |
| 1.0822 | 4.42836 | 5.00E-05 | 0.00013743 | yes |
| 1.07904 | 4.23281 | 5.00E-05 | 0.00013743 | yes |
| 1.07298 | 4.40661 | 5.00E-05 | 0.00013743 | yes |
| 1.06021 | 3.87506 | 5.00E-05 | 0.00013743 | yes |
| 1.05963 | 3.85061 | 5.00E-05 | 0.00013743 | yes |
| 1.0588 | 3.87219 | 5.00E-05 | 0.00013743 | yes |
| 1.05352 | 4.35876 | 5.00E-05 | 0.00013743 | yes |
| 1.05311 | 4.31436 | 5.00E-05 | 0.00013743 | yes |
| 1.05284 | 2.11415 | 0.00115 | 0.00256018 | yes |
| 1.052 | 3.81277 | 5.00E-05 | 0.00013743 | yes |
| 1.0496 | 1.90525 | 0.0015 | 0.00327128 | yes |
| 1.0485 | 4.30246 | 5.00E-05 | 0.00013743 | yes |
| 1.04645 | 2.01343 | 0.00065 | 0.00151996 | yes |
| 1.04511 | 4.3043 | 5.00E-05 | 0.00013743 | yes |
| 1.04349 | 1.71943 | 0.0026 | 0.00542494 | yes |
| 1.04144 | 4.29528 | 5.00E-05 | 0.00013743 | yes |
| 1.03981 | 2.45198 | 5.00E-05 | 0.00013743 | yes |
| 1.03258 | 4.09223 | 5.00E-05 | 0.00013743 | yes |
| 1.02883 | 3.5548 | 5.00E-05 | 0.00013743 | yes |
| 1.02331 | 3.56648 | 5.00E-05 | 0.00013743 | yes |
| 1.02321 | 4.16348 | 5.00E-05 | 0.00013743 | yes |
| 1.02157 | 4.18818 | 5.00E-05 | 0.00013743 | yes |
| 1.02081 | 4.1696 | 5.00E-05 | 0.00013743 | yes |
| 1.02054 | 4.20388 | 5.00E-05 | 0.00013743 | yes |
| 1.01925 | 2.53539 | 0.00015 | 0.00038924 | yes |
| 1.01508 | 4.07009 | 5.00E-05 | 0.00013743 | yes |
| 1.01491 | 4.13644 | 5.00E-05 | 0.00013743 | yes |
| 1.01258 | 4.01572 | 5.00E-05 | 0.00013743 | yes |
| 1.01174 | 4.15692 | 5.00E-05 | 0.00013743 | yes |
| 1.00844 | 4.12436 | 5.00E-05 | 0.00013743 | yes |
| 1.00664 | 3.89819 | 5.00E-05 | 0.00013743 | yes |
| 1.00364 | 4.14328 | 5.00E-05 | 0.00013743 | yes |
| 1.00234 | 3.20639 | 5.00E-05 | 0.00013743 | yes |
| 1.00133 | 2.59518 | 5.00E-05 | 0.00013743 | yes |
| 1.00093 | 3.99754 | 5.00E-05 | 0.00013743 | yes |
| 0.998972 | 4.07464 | 5.00E-05 | 0.00013743 | yes |
| 0.997017 | 4.01666 | 5.00E-05 | 0.00013743 | yes |
| 0.995095 | 4.05521 | 5.00E-05 | 0.00013743 | yes |
| 0.994 | 4.02082 | 5.00E-05 | 0.00013743 | yes |
| 0.992408 | 3.97816 | 5.00E-05 | 0.00013743 | yes |
| 0.991578 | 4.06653 | 5.00E-05 | 0.00013743 | yes |
| 0.989053 | 2.57733 | 5.00E-05 | 0.00013743 | yes |
| 0.983535 | 4.06894 | 5.00E-05 | 0.00013743 | yes |
| 0.981555 | 3.91432 | 5.00E-05 | 0.00013743 | yes |
| 0.980028 | 2.20627 | 0.0001 | 0.00026595 | yes |
| 0.979084 | 3.61243 | 5.00E-05 | 0.00013743 | yes |
| 0.975872 | 2.86 | 5.00E-05 | 0.00013743 | yes |

| | | | | |
|---|---|---|---|---|
| 0.974562 | 2.9258 | 5.00E-05 | 0.00013743 | yes |
| 0.972458 | 3.92572 | 5.00E-05 | 0.00013743 | yes |
| 0.970988 | 3.7142 | 5.00E-05 | 0.00013743 | yes |
| 0.966405 | 3.9831 | 5.00E-05 | 0.00013743 | yes |
| 0.965277 | 3.97795 | 5.00E-05 | 0.00013743 | yes |
| 0.964458 | 3.82853 | 5.00E-05 | 0.00013743 | yes |
| 0.96309 | 3.95815 | 5.00E-05 | 0.00013743 | yes |
| 0.960005 | 3.95232 | 5.00E-05 | 0.00013743 | yes |
| 0.957923 | 3.95767 | 5.00E-05 | 0.00013743 | yes |
| 0.957815 | 3.78902 | 5.00E-05 | 0.00013743 | yes |
| 0.951061 | 3.74688 | 5.00E-05 | 0.00013743 | yes |
| 0.947929 | 3.91008 | 5.00E-05 | 0.00013743 | yes |
| 0.942514 | 3.87057 | 5.00E-05 | 0.00013743 | yes |
| 0.94215 | 2.27607 | 5.00E-05 | 0.00013743 | yes |
| 0.940146 | 3.79831 | 5.00E-05 | 0.00013743 | yes |
| 0.939139 | 3.4349 | 5.00E-05 | 0.00013743 | yes |
| 0.938849 | 3.88517 | 5.00E-05 | 0.00013743 | yes |
| 0.933778 | 3.81763 | 5.00E-05 | 0.00013743 | yes |
| 0.932547 | 2.77279 | 5.00E-05 | 0.00013743 | yes |
| 0.92963 | 3.80201 | 5.00E-05 | 0.00013743 | yes |
| 0.928843 | 2.37912 | 5.00E-05 | 0.00013743 | yes |
| 0.924028 | 2.44999 | 5.00E-05 | 0.00013743 | yes |
| 0.924008 | 3.44645 | 5.00E-05 | 0.00013743 | yes |
| 0.921944 | 1.8751 | 0.00105 | 0.0023589 | yes |
| 0.921814 | 2.45189 | 5.00E-05 | 0.00013743 | yes |
| 0.919204 | 3.40245 | 5.00E-05 | 0.00013743 | yes |
| 0.918268 | 1.38334 | 0.0206 | 0.036223 | yes |
| 0.91753 | 3.79484 | 5.00E-05 | 0.00013743 | yes |
| 0.91625 | 3.76559 | 5.00E-05 | 0.00013743 | yes |
| 0.915583 | 3.08524 | 5.00E-05 | 0.00013743 | yes |
| 0.913588 | 3.54037 | 5.00E-05 | 0.00013743 | yes |
| 0.913373 | 2.10236 | 0.00055 | 0.00130598 | yes |
| 0.913212 | 3.77358 | 5.00E-05 | 0.00013743 | yes |
| 0.91201 | 2.59412 | 5.00E-05 | 0.00013743 | yes |
| 0.908869 | 3.75004 | 5.00E-05 | 0.00013743 | yes |
| 0.904838 | 3.73191 | 5.00E-05 | 0.00013743 | yes |
| 0.904617 | 3.7132 | 5.00E-05 | 0.00013743 | yes |
| 0.903242 | 2.17908 | 5.00E-05 | 0.00013743 | yes |
| 0.901413 | 3.68551 | 5.00E-05 | 0.00013743 | yes |
| 0.895726 | 3.64607 | 5.00E-05 | 0.00013743 | yes |
| 0.894564 | 3.51598 | 5.00E-05 | 0.00013743 | yes |
| 0.887949 | 2.40068 | 5.00E-05 | 0.00013743 | yes |
| 0.886445 | 3.64901 | 5.00E-05 | 0.00013743 | yes |
| 0.884529 | 3.64537 | 5.00E-05 | 0.00013743 | yes |
| 0.88372 | 3.65452 | 5.00E-05 | 0.00013743 | yes |
| 0.882279 | 3.64734 | 5.00E-05 | 0.00013743 | yes |
| 0.881863 | 3.64282 | 5.00E-05 | 0.00013743 | yes |
| 0.880127 | 3.6371 | 5.00E-05 | 0.00013743 | yes |
| 0.874727 | 3.57264 | 5.00E-05 | 0.00013743 | yes |
| 0.870438 | 2.79617 | 5.00E-05 | 0.00013743 | yes |
| 0.868224 | 3.58898 | 5.00E-05 | 0.00013743 | yes |

| | | | | |
|---|---|---|---|---|
| 0.865798 | 2.96585 | 5.00E-05 | 0.00013743 | yes |
| 0.857702 | 3.45085 | 5.00E-05 | 0.00013743 | yes |
| 0.856956 | 3.5246 | 5.00E-05 | 0.00013743 | yes |
| 0.855018 | 3.52406 | 5.00E-05 | 0.00013743 | yes |
| 0.855012 | 3.34774 | 5.00E-05 | 0.00013743 | yes |
| 0.850999 | 2.52001 | 5.00E-05 | 0.00013743 | yes |
| 0.849853 | 3.27687 | 5.00E-05 | 0.00013743 | yes |
| 0.849202 | 3.46399 | 5.00E-05 | 0.00013743 | yes |
| 0.844281 | 2.49389 | 5.00E-05 | 0.00013743 | yes |
| 0.843974 | 3.25303 | 5.00E-05 | 0.00013743 | yes |
| 0.842268 | 3.41935 | 5.00E-05 | 0.00013743 | yes |
| 0.840488 | 3.46931 | 5.00E-05 | 0.00013743 | yes |
| 0.84035 | 3.45064 | 5.00E-05 | 0.00013743 | yes |
| 0.839709 | 1.50239 | 0.00995 | 0.0186705 | yes |
| 0.833419 | 1.97873 | 0.00055 | 0.00130598 | yes |
| 0.82989 | 3.42769 | 5.00E-05 | 0.00013743 | yes |
| 0.825809 | 1.93817 | 0.0008 | 0.00183411 | yes |
| 0.825687 | 3.37324 | 5.00E-05 | 0.00013743 | yes |
| 0.824737 | 3.40888 | 5.00E-05 | 0.00013743 | yes |
| 0.822403 | 3.38325 | 5.00E-05 | 0.00013743 | yes |
| 0.820986 | 2.28519 | 0.0002 | 0.00051037 | yes |
| 0.819288 | 3.17721 | 5.00E-05 | 0.00013743 | yes |
| 0.818542 | 3.16465 | 5.00E-05 | 0.00013743 | yes |
| 0.815404 | 3.14783 | 5.00E-05 | 0.00013743 | yes |
| 0.813916 | 3.32379 | 5.00E-05 | 0.00013743 | yes |
| 0.812946 | 3.32108 | 5.00E-05 | 0.00013743 | yes |
| 0.810893 | 3.18283 | 5.00E-05 | 0.00013743 | yes |
| 0.809879 | 3.17 | 5.00E-05 | 0.00013743 | yes |
| 0.808784 | 3.34951 | 5.00E-05 | 0.00013743 | yes |
| 0.807895 | 2.37482 | 5.00E-05 | 0.00013743 | yes |
| 0.807223 | 2.51353 | 5.00E-05 | 0.00013743 | yes |
| 0.805172 | 1.68481 | 0.00305 | 0.0062525 | yes |
| 0.802178 | 3.31292 | 5.00E-05 | 0.00013743 | yes |
| 0.799474 | 3.22991 | 5.00E-05 | 0.00013743 | yes |
| 0.798302 | 1.36652 | 0.01875 | 0.0333514 | yes |
| 0.797121 | 2.58841 | 5.00E-05 | 0.00013743 | yes |
| 0.797063 | 3.27494 | 5.00E-05 | 0.00013743 | yes |
| 0.796918 | 3.29109 | 5.00E-05 | 0.00013743 | yes |
| 0.794702 | 3.18664 | 5.00E-05 | 0.00013743 | yes |
| 0.79418 | 3.27391 | 5.00E-05 | 0.00013743 | yes |
| 0.792971 | 3.28015 | 5.00E-05 | 0.00013743 | yes |
| 0.787569 | 2.91687 | 5.00E-05 | 0.00013743 | yes |
| 0.785765 | 3.17017 | 5.00E-05 | 0.00013743 | yes |
| 0.784797 | 1.78273 | 0.0021 | 0.00446114 | yes |
| 0.782969 | 3.23299 | 5.00E-05 | 0.00013743 | yes |
| 0.781869 | 3.2072 | 5.00E-05 | 0.00013743 | yes |
| 0.780358 | 3.21501 | 5.00E-05 | 0.00013743 | yes |
| 0.779111 | 3.22564 | 5.00E-05 | 0.00013743 | yes |
| 0.774973 | 3.16348 | 5.00E-05 | 0.00013743 | yes |
| 0.770956 | 3.15571 | 5.00E-05 | 0.00013743 | yes |
| 0.769307 | 2.76306 | 5.00E-05 | 0.00013743 | yes |

| | | | | |
|---|---|---|---|---|
| 0.767679 | 2.80576 | 5.00E-05 | 0.00013743 | yes |
| 0.767177 | 3.16143 | 5.00E-05 | 0.00013743 | yes |
| 0.766864 | 3.13477 | 5.00E-05 | 0.00013743 | yes |
| 0.762009 | 2.63842 | 5.00E-05 | 0.00013743 | yes |
| 0.759525 | 3.02945 | 5.00E-05 | 0.00013743 | yes |
| 0.757313 | 3.09945 | 5.00E-05 | 0.00013743 | yes |
| 0.756003 | 1.78991 | 0.0021 | 0.00446114 | yes |
| 0.755598 | 3.01077 | 5.00E-05 | 0.00013743 | yes |
| 0.755506 | 3.11919 | 5.00E-05 | 0.00013743 | yes |
| 0.754173 | 2.92663 | 5.00E-05 | 0.00013743 | yes |
| 0.753919 | 3.08107 | 5.00E-05 | 0.00013743 | yes |
| 0.753043 | 3.10655 | 5.00E-05 | 0.00013743 | yes |
| 0.752446 | 3.09509 | 5.00E-05 | 0.00013743 | yes |
| 0.749421 | 3.09422 | 5.00E-05 | 0.00013743 | yes |
| 0.748904 | 2.18836 | 0.00025 | 0.00063012 | yes |
| 0.748887 | 3.08442 | 5.00E-05 | 0.00013743 | yes |
| 0.747823 | 3.08295 | 5.00E-05 | 0.00013743 | yes |
| 0.746479 | 3.05242 | 5.00E-05 | 0.00013743 | yes |
| 0.746027 | 3.07294 | 5.00E-05 | 0.00013743 | yes |
| 0.745726 | 3.06131 | 5.00E-05 | 0.00013743 | yes |
| 0.745655 | 3.0626 | 5.00E-05 | 0.00013743 | yes |
| 0.745514 | 1.7953 | 0.00225 | 0.00474293 | yes |
| 0.745417 | 2.64919 | 5.00E-05 | 0.00013743 | yes |
| 0.743098 | 3.06508 | 5.00E-05 | 0.00013743 | yes |
| 0.739259 | 2.80397 | 5.00E-05 | 0.00013743 | yes |
| 0.739207 | 3.04539 | 5.00E-05 | 0.00013743 | yes |
| 0.73662 | 2.95408 | 5.00E-05 | 0.00013743 | yes |
| 0.733414 | 3.01972 | 5.00E-05 | 0.00013743 | yes |
| 0.733303 | 3.02683 | 5.00E-05 | 0.00013743 | yes |
| 0.733236 | 2.91419 | 5.00E-05 | 0.00013743 | yes |
| 0.731783 | 1.34508 | 0.02205 | 0.0385522 | yes |
| 0.72966 | 3.00214 | 5.00E-05 | 0.00013743 | yes |
| 0.729038 | 2.95896 | 5.00E-05 | 0.00013743 | yes |
| 0.726868 | 2.68562 | 5.00E-05 | 0.00013743 | yes |
| 0.72676 | 2.98362 | 5.00E-05 | 0.00013743 | yes |
| 0.726556 | 1.42715 | 0.01405 | 0.0255643 | yes |
| 0.725349 | 2.87985 | 5.00E-05 | 0.00013743 | yes |
| 0.725172 | 3.00236 | 5.00E-05 | 0.00013743 | yes |
| 0.721649 | 2.92659 | 5.00E-05 | 0.00013743 | yes |
| 0.715501 | 2.92565 | 5.00E-05 | 0.00013743 | yes |
| 0.714192 | 2.91303 | 5.00E-05 | 0.00013743 | yes |
| 0.714015 | 2.94107 | 5.00E-05 | 0.00013743 | yes |
| 0.713971 | 1.51837 | 0.0097 | 0.0182431 | yes |
| 0.713277 | 2.94068 | 5.00E-05 | 0.00013743 | yes |
| 0.71252 | 2.31984 | 0.00015 | 0.00038924 | yes |
| 0.711022 | 2.93523 | 5.00E-05 | 0.00013743 | yes |
| 0.708247 | 1.61439 | 0.0056 | 0.0109594 | yes |
| 0.707181 | 2.73776 | 5.00E-05 | 0.00013743 | yes |
| 0.704365 | 2.905 | 5.00E-05 | 0.00013743 | yes |
| 0.702698 | 2.05657 | 0.00035 | 0.00086186 | yes |
| 0.701506 | 2.90195 | 5.00E-05 | 0.00013743 | yes |

| | | | | |
|---|---|---|---|---|
| 0.701429 | 2.20727 | 0.0002 | 0.00051037 | yes |
| 0.701139 | 2.89344 | 5.00E-05 | 0.00013743 | yes |
| 0.700787 | 2.0706 | 0.00015 | 0.00038924 | yes |
| 0.697703 | 2.80801 | 5.00E-05 | 0.00013743 | yes |
| 0.697565 | 2.88703 | 5.00E-05 | 0.00013743 | yes |
| 0.695358 | 1.90183 | 0.0009 | 0.00204621 | yes |
| 0.694615 | 2.81267 | 5.00E-05 | 0.00013743 | yes |
| 0.694498 | 2.86433 | 5.00E-05 | 0.00013743 | yes |
| 0.693034 | 2.8655 | 5.00E-05 | 0.00013743 | yes |
| 0.692428 | 1.78508 | 0.00325 | 0.00664037 | yes |
| 0.68906 | 2.81513 | 5.00E-05 | 0.00013743 | yes |
| 0.68853 | 2.82849 | 5.00E-05 | 0.00013743 | yes |
| 0.688472 | 2.81387 | 5.00E-05 | 0.00013743 | yes |
| 0.686849 | 1.74439 | 0.0035 | 0.0071157 | yes |
| 0.684114 | 2.82812 | 5.00E-05 | 0.00013743 | yes |
| 0.680673 | 2.81422 | 5.00E-05 | 0.00013743 | yes |
| 0.678807 | 2.48078 | 5.00E-05 | 0.00013743 | yes |
| 0.677144 | 2.78205 | 5.00E-05 | 0.00013743 | yes |
| 0.675248 | 2.78865 | 5.00E-05 | 0.00013743 | yes |
| 0.673984 | 2.31669 | 5.00E-05 | 0.00013743 | yes |
| 0.673465 | 2.78856 | 5.00E-05 | 0.00013743 | yes |
| 0.668424 | 2.76352 | 5.00E-05 | 0.00013743 | yes |
| 0.667899 | 2.47177 | 5.00E-05 | 0.00013743 | yes |
| 0.666878 | 2.69182 | 5.00E-05 | 0.00013743 | yes |
| 0.666846 | 2.06623 | 0.0006 | 0.00141244 | yes |
| 0.66609 | 2.42998 | 5.00E-05 | 0.00013743 | yes |
| 0.66583 | 2.61395 | 5.00E-05 | 0.00013743 | yes |
| 0.665547 | 2.72118 | 5.00E-05 | 0.00013743 | yes |
| 0.665133 | 2.68637 | 5.00E-05 | 0.00013743 | yes |
| 0.664146 | 2.46479 | 5.00E-05 | 0.00013743 | yes |
| 0.662722 | 2.5055 | 5.00E-05 | 0.00013743 | yes |
| 0.662612 | 2.72115 | 5.00E-05 | 0.00013743 | yes |
| 0.662111 | 2.71857 | 5.00E-05 | 0.00013743 | yes |
| 0.660784 | 2.73306 | 5.00E-05 | 0.00013743 | yes |
| 0.659359 | 2.11902 | 0.0005 | 0.00119766 | yes |
| 0.656159 | 2.7003 | 5.00E-05 | 0.00013743 | yes |
| 0.655351 | 2.48897 | 5.00E-05 | 0.00013743 | yes |
| 0.647983 | 2.64989 | 5.00E-05 | 0.00013743 | yes |
| 0.647189 | 2.66662 | 5.00E-05 | 0.00013743 | yes |
| 0.646502 | 2.67229 | 5.00E-05 | 0.00013743 | yes |
| 0.644867 | 1.98239 | 0.00055 | 0.00130598 | yes |
| 0.644607 | 1.98263 | 0.0008 | 0.00183411 | yes |
| 0.643778 | 1.43646 | 0.0141 | 0.0256364 | yes |
| 0.643415 | 2.01194 | 0.0003 | 0.00074621 | yes |
| 0.642532 | 2.64679 | 5.00E-05 | 0.00013743 | yes |
| 0.641373 | 2.60557 | 5.00E-05 | 0.00013743 | yes |
| 0.640152 | 2.59117 | 5.00E-05 | 0.00013743 | yes |
| 0.639364 | 2.61595 | 5.00E-05 | 0.00013743 | yes |
| 0.636719 | 1.34344 | 0.02635 | 0.0452345 | yes |
| 0.635204 | 1.82873 | 0.0012 | 0.00266186 | yes |
| 0.634561 | 2.6172 | 5.00E-05 | 0.00013743 | yes |

| | | | | |
|---|---|---|---|---|
| 0.632166 | 1.97051 | 0.0005 | 0.00119766 | yes |
| 0.632115 | 1.66245 | 0.00445 | 0.00886397 | yes |
| 0.631881 | 2.60652 | 5.00E-05 | 0.00013743 | yes |
| 0.631605 | 1.53068 | 0.00925 | 0.0174368 | yes |
| 0.631597 | 2.59885 | 5.00E-05 | 0.00013743 | yes |
| 0.63089 | 2.60212 | 5.00E-05 | 0.00013743 | yes |
| 0.630767 | 2.53271 | 5.00E-05 | 0.00013743 | yes |
| 0.630579 | 1.34496 | 0.021 | 0.0368474 | yes |
| 0.630477 | 2.59309 | 5.00E-05 | 0.00013743 | yes |
| 0.630208 | 2.57932 | 5.00E-05 | 0.00013743 | yes |
| 0.629594 | 2.56823 | 5.00E-05 | 0.00013743 | yes |
| 0.628118 | 2.44688 | 0.0001 | 0.00026595 | yes |
| 0.627009 | 1.62512 | 0.0065 | 0.0125806 | yes |
| 0.624166 | 2.39978 | 5.00E-05 | 0.00013743 | yes |
| 0.622749 | 2.55436 | 5.00E-05 | 0.00013743 | yes |
| 0.621821 | 1.32353 | 0.0224 | 0.0390809 | yes |
| 0.621576 | 2.27409 | 5.00E-05 | 0.00013743 | yes |
| 0.620166 | 2.48872 | 5.00E-05 | 0.00013743 | yes |
| 0.620002 | 2.23785 | 5.00E-05 | 0.00013743 | yes |
| 0.619042 | 2.16885 | 5.00E-05 | 0.00013743 | yes |
| 0.619033 | 2.45713 | 5.00E-05 | 0.00013743 | yes |
| 0.61834 | 2.55227 | 5.00E-05 | 0.00013743 | yes |
| 0.616374 | 1.42287 | 0.0153 | 0.0276547 | yes |
| 0.614871 | 2.17648 | 0.0002 | 0.00051037 | yes |
| 0.614491 | 2.47313 | 5.00E-05 | 0.00013743 | yes |
| 0.613531 | 2.52355 | 5.00E-05 | 0.00013743 | yes |
| 0.611457 | 2.4742 | 5.00E-05 | 0.00013743 | yes |
| 0.609895 | 2.16141 | 0.0001 | 0.00026595 | yes |
| 0.608677 | 1.91011 | 0.001 | 0.00225275 | yes |
| 0.606416 | 2.02998 | 0.0005 | 0.00119766 | yes |
| 0.603671 | 2.44013 | 5.00E-05 | 0.00013743 | yes |
| 0.603177 | 2.06495 | 0.00055 | 0.00130598 | yes |
| 0.600539 | 2.26978 | 0.0001 | 0.00026595 | yes |
| 0.598758 | 2.3143 | 0.0001 | 0.00026595 | yes |
| 0.59831 | 2.44864 | 5.00E-05 | 0.00013743 | yes |
| 0.598085 | 2.45159 | 5.00E-05 | 0.00013743 | yes |
| 0.597132 | 2.37224 | 0.0001 | 0.00026595 | yes |
| 0.596759 | 2.4484 | 5.00E-05 | 0.00013743 | yes |
| 0.595833 | 2.27452 | 0.00015 | 0.00038924 | yes |
| 0.593727 | 2.40605 | 5.00E-05 | 0.00013743 | yes |
| 0.589605 | 1.52512 | 0.00805 | 0.0153155 | yes |
| 0.589331 | 2.41258 | 5.00E-05 | 0.00013743 | yes |
| 0.587071 | 2.42131 | 5.00E-05 | 0.00013743 | yes |
| 0.586297 | 2.1052 | 0.0001 | 0.00026595 | yes |
| 0.58442 | 2.15394 | 0.0002 | 0.00051037 | yes |
| 0.583549 | 2.3891 | 0.00015 | 0.00038924 | yes |
| 0.583423 | 1.83537 | 0.0015 | 0.00327128 | yes |
| 0.583364 | 2.41258 | 5.00E-05 | 0.00013743 | yes |
| 0.581748 | 2.34517 | 5.00E-05 | 0.00013743 | yes |
| 0.578959 | 2.37465 | 5.00E-05 | 0.00013743 | yes |
| 0.577949 | 2.35268 | 5.00E-05 | 0.00013743 | yes |

| | | | | |
|---|---|---|---|---|
| 0.57555 | 2.336 | 5.00E-05 | 0.00013743 | yes |
| 0.574789 | 2.15247 | 0.00035 | 0.00086186 | yes |
| 0.574399 | 2.36371 | 5.00E-05 | 0.00013743 | yes |
| 0.573893 | 2.37538 | 5.00E-05 | 0.00013743 | yes |
| 0.572507 | 1.94528 | 0.00065 | 0.00151996 | yes |
| 0.571661 | 2.35352 | 5.00E-05 | 0.00013743 | yes |
| 0.570868 | 2.21787 | 0.00015 | 0.00038924 | yes |
| 0.570551 | 1.93352 | 0.001 | 0.00225275 | yes |
| 0.568257 | 2.19694 | 0.0002 | 0.00051037 | yes |
| 0.567809 | 2.34515 | 5.00E-05 | 0.00013743 | yes |
| 0.567603 | 2.16006 | 0.00015 | 0.00038924 | yes |
| 0.566714 | 2.31499 | 0.00015 | 0.00038924 | yes |
| 0.566491 | 1.45741 | 0.01115 | 0.0207795 | yes |
| 0.565088 | 1.74365 | 0.0023 | 0.00483177 | yes |
| 0.56504 | 2.32368 | 5.00E-05 | 0.00013743 | yes |
| 0.563999 | 1.60901 | 0.00575 | 0.0112351 | yes |
| 0.559878 | 2.31022 | 0.0001 | 0.00026595 | yes |
| 0.558878 | 2.28246 | 0.0001 | 0.00026595 | yes |
| 0.558699 | 1.72779 | 0.0029 | 0.00598992 | yes |
| 0.558388 | 1.90126 | 0.0007 | 0.00162914 | yes |
| 0.558158 | 2.29517 | 5.00E-05 | 0.00013743 | yes |
| 0.557615 | 1.45257 | 0.0116 | 0.0215366 | yes |
| 0.556093 | 2.28877 | 5.00E-05 | 0.00013743 | yes |
| 0.555453 | 2.24987 | 0.00015 | 0.00038924 | yes |
| 0.554803 | 2.22993 | 0.00015 | 0.00038924 | yes |
| 0.551266 | 1.59363 | 0.00685 | 0.0131958 | yes |
| 0.551191 | 2.07607 | 0.0003 | 0.00074621 | yes |
| 0.550179 | 1.38621 | 0.01755 | 0.0314672 | yes |
| 0.549188 | 2.27166 | 5.00E-05 | 0.00013743 | yes |
| 0.546735 | 2.25018 | 5.00E-05 | 0.00013743 | yes |
| 0.546105 | 2.24961 | 0.00015 | 0.00038924 | yes |
| 0.545772 | 2.22369 | 5.00E-05 | 0.00013743 | yes |
| 0.544096 | 2.01737 | 0.0006 | 0.00141244 | yes |
| 0.543694 | 2.12204 | 0.00015 | 0.00038924 | yes |
| 0.542373 | 2.23172 | 0.0001 | 0.00026595 | yes |
| 0.540483 | 2.00738 | 0.0004 | 0.00097813 | yes |
| 0.540024 | 2.22492 | 5.00E-05 | 0.00013743 | yes |
| 0.538815 | 1.99635 | 0.00075 | 0.00173402 | yes |
| 0.537204 | 2.22419 | 0.00025 | 0.00063012 | yes |
| 0.536907 | 1.98033 | 0.0004 | 0.00097813 | yes |
| 0.536326 | 2.21919 | 0.00015 | 0.00038924 | yes |
| 0.534485 | 2.14845 | 0.0001 | 0.00026595 | yes |
| 0.531985 | 2.16954 | 0.00015 | 0.00038924 | yes |
| 0.531236 | 1.89226 | 0.001 | 0.00225275 | yes |
| 0.530421 | 2.18417 | 5.00E-05 | 0.00013743 | yes |
| 0.529208 | 1.26723 | 0.02795 | 0.0477811 | yes |
| 0.528265 | 1.63555 | 0.0055 | 0.0107809 | yes |
| 0.519824 | 2.13023 | 5.00E-05 | 0.00013743 | yes |
| 0.519713 | 2.13508 | 0.0003 | 0.00074621 | yes |
| 0.519116 | 2.12415 | 0.0002 | 0.00051037 | yes |
| 0.517388 | 1.48088 | 0.01105 | 0.0206088 | yes |

| | | | | |
|---|---|---|---|---|
| 0.517269 | 2.13486 | 0.00015 | 0.00038924 | yes |
| 0.517069 | 1.93415 | 0.0013 | 0.00286816 | yes |
| 0.51527 | 2.1093 | 0.00025 | 0.00063012 | yes |
| 0.514424 | 1.80004 | 0.00195 | 0.00417493 | yes |
| 0.514169 | 2.10914 | 0.00035 | 0.00086186 | yes |
| 0.513622 | 1.44548 | 0.01325 | 0.0241982 | yes |
| 0.513569 | 2.10224 | 0.0002 | 0.00051037 | yes |
| 0.511806 | 1.81565 | 0.00175 | 0.00376311 | yes |
| 0.509859 | 2.10143 | 0.0002 | 0.00051037 | yes |
| 0.509482 | 2.04389 | 0.00035 | 0.00086186 | yes |
| 0.509229 | 2.08705 | 0.0004 | 0.00097813 | yes |
| 0.508926 | 2.1072 | 0.00025 | 0.00063012 | yes |
| 0.50731 | 2.05103 | 0.0003 | 0.00074621 | yes |
| 0.505629 | 2.06797 | 0.00035 | 0.00086186 | yes |
| 0.504624 | 2.08815 | 5.00E-05 | 0.00013743 | yes |
| 0.503143 | 2.06855 | 0.0003 | 0.00074621 | yes |
| 0.502585 | 1.40941 | 0.01565 | 0.0282665 | yes |
| 0.501699 | 2.06349 | 0.0004 | 0.00097813 | yes |
| 0.500514 | 1.91259 | 0.00075 | 0.00173402 | yes |
| 0.49835 | 1.34992 | 0.0199 | 0.0351176 | yes |
| 0.497976 | 2.05162 | 0.00015 | 0.00038924 | yes |
| 0.494887 | 2.03085 | 0.0003 | 0.00074621 | yes |
| 0.494762 | 1.98446 | 0.0005 | 0.00119766 | yes |
| 0.493241 | 1.99176 | 0.00075 | 0.00173402 | yes |
| 0.492314 | 1.97031 | 0.00065 | 0.00151996 | yes |
| 0.4917 | 2.01841 | 0.0003 | 0.00074621 | yes |
| 0.490763 | 1.59155 | 0.0067 | 0.0129474 | yes |
| 0.490415 | 1.97573 | 0.00045 | 0.0010885 | yes |
| 0.489264 | 1.98974 | 0.0006 | 0.00141244 | yes |
| 0.489252 | 2.00941 | 0.00045 | 0.0010885 | yes |
| 0.486802 | 2.01768 | 0.00035 | 0.00086186 | yes |
| 0.486783 | 2.01715 | 0.00045 | 0.0010885 | yes |
| 0.486303 | 2.00834 | 0.0006 | 0.00141244 | yes |
| 0.485555 | 2.00038 | 0.0005 | 0.00119766 | yes |
| 0.48542 | 1.95627 | 0.0006 | 0.00141244 | yes |
| 0.485076 | 1.96749 | 0.0007 | 0.00162914 | yes |
| 0.48409 | 1.99945 | 0.0005 | 0.00119766 | yes |
| 0.48338 | 1.57443 | 0.00795 | 0.0151605 | yes |
| 0.481737 | 1.9902 | 0.00065 | 0.00151996 | yes |
| 0.481553 | 1.99078 | 0.00055 | 0.00130598 | yes |
| 0.481085 | 1.97134 | 0.0007 | 0.00162914 | yes |
| 0.480791 | 1.98836 | 0.00065 | 0.00151996 | yes |
| 0.479957 | 1.29181 | 0.0232 | 0.0402482 | yes |
| 0.479628 | 1.28024 | 0.0273 | 0.0467674 | yes |
| 0.478744 | 1.71798 | 0.0024 | 0.00502896 | yes |
| 0.478102 | 1.95166 | 0.00075 | 0.00173402 | yes |
| 0.47698 | 1.96593 | 0.00085 | 0.0019415 | yes |
| 0.476866 | 1.35807 | 0.01805 | 0.0322228 | yes |
| 0.473932 | 1.85453 | 0.0013 | 0.00286816 | yes |
| 0.472103 | 1.93735 | 0.001 | 0.00225275 | yes |
| 0.471375 | 1.88263 | 0.0014 | 0.003075 | yes |

| | | | | |
|---|---|---|---|---|
| 0.470468 | 1.85328 | 0.0011 | 0.00246224 | yes |
| 0.469798 | 1.78266 | 0.00265 | 0.0055199 | yes |
| 0.469612 | 1.93615 | 0.0009 | 0.00204621 | yes |
| 0.469507 | 1.93323 | 0.0008 | 0.00183411 | yes |
| 0.469468 | 1.94486 | 0.00095 | 0.0021559 | yes |
| 0.467959 | 1.89443 | 0.00135 | 0.00297048 | yes |
| 0.467543 | 1.90778 | 0.001 | 0.00225275 | yes |
| 0.46563 | 1.907 | 0.0006 | 0.00141244 | yes |
| 0.464839 | 1.54931 | 0.0059 | 0.0115099 | yes |
| 0.463431 | 1.59433 | 0.0055 | 0.0107809 | yes |
| 0.462804 | 1.67305 | 0.0034 | 0.00692958 | yes |
| 0.460527 | 1.89938 | 0.001 | 0.00225275 | yes |
| 0.460388 | 1.90037 | 0.001 | 0.00225275 | yes |
| 0.460343 | 1.36509 | 0.0189 | 0.0335697 | yes |
| 0.458669 | 1.67981 | 0.00405 | 0.00810659 | yes |
| 0.458491 | 1.8888 | 0.00085 | 0.0019415 | yes |
| 0.457142 | 1.662 | 0.0044 | 0.00877859 | yes |
| 0.456962 | 1.88831 | 0.0011 | 0.00246224 | yes |
| 0.4569 | 1.88183 | 0.0008 | 0.00183411 | yes |
| 0.456421 | 1.88377 | 0.00155 | 0.00336243 | yes |
| 0.455784 | 1.80273 | 0.00215 | 0.00455164 | yes |
| 0.45318 | 1.65804 | 0.0037 | 0.0074852 | yes |
| 0.451532 | 1.85858 | 0.00125 | 0.00277027 | yes |
| 0.451287 | 1.73862 | 0.0026 | 0.00542494 | yes |
| 0.450418 | 1.85414 | 0.0014 | 0.003075 | yes |
| 0.449919 | 1.8299 | 0.00175 | 0.00376311 | yes |
| 0.44697 | 1.4134 | 0.0139 | 0.0253101 | yes |
| 0.446627 | 1.83755 | 0.0008 | 0.00183411 | yes |
| 0.446004 | 1.41555 | 0.01285 | 0.0235727 | yes |
| 0.445948 | 1.62135 | 0.00455 | 0.00905583 | yes |
| 0.444248 | 1.78659 | 0.0027 | 0.00560506 | yes |
| 0.442672 | 1.80988 | 0.00175 | 0.00376311 | yes |
| 0.439939 | 1.81537 | 0.00145 | 0.00317914 | yes |
| 0.438642 | 1.80714 | 0.0016 | 0.00346174 | yes |
| 0.438169 | 1.79153 | 0.0013 | 0.00286816 | yes |
| 0.437262 | 1.80228 | 0.002 | 0.00427826 | yes |
| 0.436954 | 1.75937 | 0.0021 | 0.00446114 | yes |
| 0.436623 | 1.75515 | 0.00215 | 0.00455164 | yes |
| 0.43598 | 1.5923 | 0.0056 | 0.0109594 | yes |
| 0.435308 | 1.50464 | 0.0083 | 0.0157546 | yes |
| 0.434981 | 1.77611 | 0.00205 | 0.00437002 | yes |
| 0.43376 | 1.78681 | 0.00155 | 0.00336243 | yes |
| 0.431435 | 1.30336 | 0.02265 | 0.039461 | yes |
| 0.430027 | 1.76806 | 0.0019 | 0.00407498 | yes |
| 0.42965 | 1.4259 | 0.01415 | 0.0256893 | yes |
| 0.429598 | 1.75695 | 0.0034 | 0.00692958 | yes |
| 0.428845 | 1.71786 | 0.00275 | 0.00569444 | yes |
| 0.42322 | 1.74409 | 0.00245 | 0.00512936 | yes |
| 0.421809 | 1.72761 | 0.00235 | 0.00493259 | yes |
| 0.421655 | 1.71381 | 0.0028 | 0.0057931 | yes |
| 0.420644 | 1.72011 | 0.0036 | 0.00730091 | yes |

| | | | | |
|---|---|---|---|---|
| 0.420378 | 1.71154 | 0.0023 | 0.00483177 | yes |
| 0.420326 | 1.72741 | 0.00275 | 0.00569444 | yes |
| 0.420326 | 1.72172 | 0.0027 | 0.00560506 | yes |
| 0.420085 | 1.72301 | 0.0031 | 0.00634971 | yes |
| 0.419767 | 1.72644 | 0.00305 | 0.0062525 | yes |
| 0.419541 | 1.71939 | 0.0025 | 0.00522959 | yes |
| 0.417974 | 1.70087 | 0.00275 | 0.00569444 | yes |
| 0.416368 | 1.7091 | 0.0037 | 0.0074852 | yes |
| 0.416116 | 1.70975 | 0.00305 | 0.0062525 | yes |
| 0.415916 | 1.70261 | 0.003 | 0.00617057 | yes |
| 0.415102 | 1.69162 | 0.00345 | 0.00701985 | yes |
| 0.414286 | 1.30171 | 0.02505 | 0.0432746 | yes |
| 0.411897 | 1.69086 | 0.0027 | 0.00560506 | yes |
| 0.411729 | 1.69643 | 0.00355 | 0.0072114 | yes |
| 0.409324 | 1.68506 | 0.00345 | 0.00701985 | yes |
| 0.407024 | 1.65674 | 0.00385 | 0.00775676 | yes |
| 0.406014 | 1.36123 | 0.018 | 0.0321569 | yes |
| 0.405566 | 1.59019 | 0.006 | 0.0116865 | yes |
| 0.405373 | 1.44123 | 0.01305 | 0.0238684 | yes |
| 0.403076 | 1.62562 | 0.004 | 0.0080261 | yes |
| 0.402808 | 1.45157 | 0.01185 | 0.021918 | yes |
| 0.401565 | 1.59237 | 0.0055 | 0.0107809 | yes |
| 0.400627 | 1.64139 | 0.00445 | 0.00886397 | yes |
| 0.397555 | 1.63527 | 0.004 | 0.0080261 | yes |
| 0.397145 | 1.62501 | 0.00405 | 0.00810659 | yes |
| 0.395421 | 1.63737 | 0.0059 | 0.0115099 | yes |
| 0.394575 | 1.62953 | 0.00535 | 0.0105204 | yes |
| 0.394325 | 1.61753 | 0.00505 | 0.0100024 | yes |
| 0.39266 | 1.61214 | 0.00485 | 0.00962177 | yes |
| 0.392205 | 1.52277 | 0.00905 | 0.0170729 | yes |
| 0.390825 | 1.33607 | 0.0198 | 0.0350165 | yes |
| 0.389984 | 1.59764 | 0.00495 | 0.00981225 | yes |
| 0.388418 | 1.5991 | 0.0053 | 0.0104304 | yes |
| 0.388197 | 1.28583 | 0.0274 | 0.0468734 | yes |
| 0.385208 | 1.28401 | 0.02295 | 0.0399272 | yes |
| 0.384922 | 1.59273 | 0.00605 | 0.0117652 | yes |
| 0.382904 | 1.56095 | 0.00705 | 0.0135598 | yes |
| 0.381632 | 1.57266 | 0.00645 | 0.0124937 | yes |
| 0.381181 | 1.57533 | 0.0063 | 0.0122128 | yes |
| 0.381035 | 1.43142 | 0.01275 | 0.0234242 | yes |
| 0.380531 | 1.56978 | 0.0068 | 0.01312 | yes |
| 0.378714 | 1.55184 | 0.0069 | 0.0132817 | yes |
| 0.377589 | 1.53447 | 0.00685 | 0.0131958 | yes |
| 0.376854 | 1.4559 | 0.0125 | 0.0229993 | yes |
| 0.376499 | 1.44339 | 0.01305 | 0.0238684 | yes |
| 0.374709 | 1.5359 | 0.00755 | 0.0144312 | yes |
| 0.374427 | 1.55002 | 0.0084 | 0.0159322 | yes |
| 0.374199 | 1.53429 | 0.008 | 0.0152322 | yes |
| 0.37413 | 1.34545 | 0.01915 | 0.0339892 | yes |
| 0.373313 | 1.31802 | 0.02145 | 0.0375833 | yes |
| 0.372926 | 1.52651 | 0.0079 | 0.0150885 | yes |

| | | | | |
|---|---|---|---|---|
| 0.372625 | 1.50356 | 0.0071 | 0.0136453 | yes |
| 0.37075 | 1.36378 | 0.0166 | 0.0298509 | yes |
| 0.369186 | 1.51631 | 0.0073 | 0.0139642 | yes |
| 0.368294 | 1.39328 | 0.01635 | 0.0294228 | yes |
| 0.368158 | 1.46811 | 0.011 | 0.0205311 | yes |
| 0.367698 | 1.33328 | 0.0215 | 0.0376441 | yes |
| 0.367445 | 1.50649 | 0.0083 | 0.0157546 | yes |
| 0.364368 | 1.38628 | 0.0158 | 0.0284747 | yes |
| 0.361016 | 1.49153 | 0.0089 | 0.0168157 | yes |
| 0.360168 | 1.29093 | 0.02405 | 0.0416055 | yes |
| 0.360023 | 1.29863 | 0.0221 | 0.0386122 | yes |
| 0.357955 | 1.48137 | 0.0096 | 0.0180689 | yes |
| 0.357815 | 1.47745 | 0.0098 | 0.0184031 | yes |
| 0.355668 | 1.32414 | 0.02 | 0.0352688 | yes |
| 0.354641 | 1.45919 | 0.01155 | 0.02146 | yes |
| 0.353592 | 1.35711 | 0.01675 | 0.0300986 | yes |
| 0.353043 | 1.35435 | 0.0196 | 0.0347378 | yes |
| 0.35253 | 1.455 | 0.0115 | 0.0213832 | yes |
| 0.350225 | 1.44098 | 0.0112 | 0.0208569 | yes |
| 0.350081 | 1.43855 | 0.01235 | 0.0227573 | yes |
| 0.349777 | 1.44483 | 0.0109 | 0.0203599 | yes |
| 0.349746 | 1.4324 | 0.01285 | 0.0235727 | yes |
| 0.349547 | 1.43982 | 0.0123 | 0.0226822 | yes |
| 0.34904 | 1.40295 | 0.01415 | 0.0256893 | yes |
| 0.34114 | 1.36598 | 0.0179 | 0.0320247 | yes |
| 0.339364 | 1.34704 | 0.01755 | 0.0314672 | yes |
| 0.336528 | 1.37354 | 0.01765 | 0.0316235 | yes |
| 0.335169 | 1.37256 | 0.01825 | 0.0325091 | yes |
| 0.334799 | 1.2637 | 0.02935 | 0.049897 | yes |
| 0.333532 | 1.28861 | 0.02635 | 0.0452345 | yes |
| 0.333186 | 1.22999 | 0.0291 | 0.0495748 | yes |
| 0.332624 | 1.36827 | 0.0182 | 0.0324435 | yes |
| 0.330874 | 1.30073 | 0.0256 | 0.04407 | yes |
| 0.326785 | 1.33856 | 0.021 | 0.0368474 | yes |
| 0.326074 | 1.35211 | 0.01815 | 0.0323778 | yes |
| 0.323172 | 1.32279 | 0.02135 | 0.0374348 | yes |
| 0.322693 | 1.26781 | 0.02625 | 0.0451258 | yes |
| 0.322648 | 1.31764 | 0.0232 | 0.0402482 | yes |
| 0.322483 | 1.3212 | 0.02155 | 0.0377048 | yes |
| 0.317678 | 1.31295 | 0.0197 | 0.0348898 | yes |
| 0.31736 | 1.3091 | 0.023 | 0.0399859 | yes |
| 0.314883 | 1.28982 | 0.02305 | 0.0400445 | yes |
| 0.313464 | 1.29192 | 0.0255 | 0.0439594 | yes |
| 0.311695 | 1.27879 | 0.02515 | 0.0434168 | yes |
| 0.308718 | 1.27137 | 0.02595 | 0.0446413 | yes |
| 0.305549 | 1.25429 | 0.02885 | 0.049183 | yes |
| 0.0343834 | 0.141734 | 0.8051 | 0.849981 | no |
| 0.0303819 | 0.125415 | 0.8262 | 0.865241 | no |
| 0.421655 | 1.71381 | 0.0028 | 0.0057931 | yes |

Up

| V1 | V2 | | |
|---|---|---|---|
| 1 | 3 | 3 | 1.5849625 |

| | gene |
|---|---|
| acetyl-CoA synthetase | acetyl-CoA synthetase |
| carbamate kinase | arcC |
| hypothetical protein | hypothetical protein |
| hypothetical protein | hypothetical protein |
| hypothetical protein | hypothetical protein |
| hypothetical protein | hypothetical protein |
| hypothetical protein | hypothetical protein |
| catabolite control protein A | ccpA |
| hypothetical protein | hypothetical protein |
| hypothetical protein | hypothetical protein |
| manganese transport protein MntH | mntH |
| hypothetical protein | hypothetical protein |
| L-lactate dehydrogenase | ldh1 |
| hypothetical protein | hypothetical protein |
| serine protease HtrA | htrA2 |
| glycerol kinase | glpK |
| monovalent cation/H+ antiporter subunit G | mnhG2 |
| 1-pyrroline-5-carboxylate dehydrogenase | rocA |
| D-lactate dehydrogenase | D-lactate dehydrogenase |
| | sarX |
| ribonucleotide-diphosphate reductase subunit beta | nrdF |
| ribonuclease III | rnc |
| hypothetical protein | bmrU |
| molybdenum ABC transporter permease | modB |
| PTS system transporter subunit IIA | crr |
| hypothetical protein | hypothetical protein |
| hypothetical protein | tcaA |
| phosphopyruvate hydratase | eno |
| hypothetical protein | hypothetical protein |
| DNA polymerase I | polA |
| hypothetical protein | hypothetical protein |
| recombination regulator RecX | recX |
| hypothetical protein | hypothetical protein |
| hypothetical protein | hypothetical protein |
| hypothetical protein | mtlR |
| nisin susceptibility-associated sensor histidine kinase | nsaS |
| hypothetical protein | hypothetical protein |
| hypothetical protein | hypothetical protein |
| inorganic polyphosphate/ATP-NAD kinase | ppnK |
| cardiolipin synthetase | cls1 |
| hypothetical protein | hypothetical protein |
| ATP-dependent protease ATP-binding subunit HslU | hslU |
| hypothetical protein | hypothetical protein |
| inositol monophosphatase family protein | imp |
| aldehyde dehydrogenase | aldH |
| hypothetical protein | hypothetical protein |
| hypothetical protein | sfnaB |
| hypothetical protein | hypothetical protein |
| hypothetical protein | tatC |
| hypothetical protein | hypothetical protein |
| hypothetical protein | hemX |
| formate--tetrahydrofolate ligase | fhs |
| regulatory protein SpoVG | spoVG |
| D-isomer specific 2-hydroxyacid dehydrogenase NAD binding | oxyacid dehydrogenase NAD binding d |

| | |
|---|---|
| 16S Ribosomal RNA | rrsD |
| sodium, sulfate symporter | sodium, sulfate symporter |
| peptidase T | pepT |
| hypothetical protein | hypothetical protein |
| excinuclease ABC subunit B | uvrB |
| hypothetical protein | hypothetical protein |
| hypothetical protein | sufB |
| hypothetical protein | hypothetical protein |
| heme-degrading monooxygenase IsdI | isdI |
| dihydroorotate dehydrogenase 2 | pyrD |
| arginase | rocF |
| excinuclease ABC subunit A | uvrA |
| hypothetical protein | prsS |
| hypothetical protein | yabJ |
| hypothetical protein | hypothetical protein |
| hypothetical protein | hypothetical protein |
| hypothetical protein | hypothetical protein |
| formamidopyrimidine-DNA glycosylase | mutM |
| hypothetical protein | hypothetical protein |
| hypothetical protein | flp |
| D-isomer specific 2-hydroxyacid dehydrogenase NAD binding | oxyacid dehydrogenase NAD binding d |
| teichoic acid biosynthesis protein F | tarF |
| 2-oxoglutarate dehydrogenase E1 component | sucA |
| hypothetical protein | hypothetical protein |
| acetate--CoA ligase | acsA |
| 2-deoxyribose-5-phosphate aldolase | deoC1 |
| hypothetical protein | eta |
| D-alanine aminotransferase | dat |
| hypothetical protein | spdA |
| hypothetical protein | uspA1 |
| hypothetical protein | hypothetical protein |
| haloacid dehalogenase-like hydrolase | haloacid dehalogenase-like hydrolase |
| endonuclease III | endonuclease III |
| hypothetical protein | csa1A |
| hypothetical protein | hypothetical protein |
| phosphoenolpyruvate carboxykinase | pckA |
| hypothetical protein | hypothetical protein |
| hypothetical protein | hypothetical protein |
| tRNA delta(2)-isopentenylpyrophosphate transferase | miaA |
| hypothetical protein | hypothetical protein |
| hypothetical protein | ydbM |
| hypothetical protein | tet38 |
| hypothetical protein | hypothetical protein |
| Glycerol uptake facilitator | glpF |
| hypothetical protein | esaA |
| aspartate kinase | thrD |
| hypothetical protein | glpQ |
| ornithine carbamoyltransferase | arcB |
| hypothetical protein | hypothetical protein |
| hypothetical protein | kapB |
| 2-oxoisovalerate dehydrogenase, E1 component subunit beta | bfmBAB |
| hypothetical protein | hypothetical protein |
| hypothetical protein | hypothetical protein |
| hypothetical protein | rarD |
| hypothetical protein | fakB2 |

| | |
|---|---|
| hypothetical protein | hypothetical protein |
| hypothetical protein | sasD |
| 2-oxoisovalerate dehydrogenase, E2 component, dihydrolipoa | bmfBB |
| hypothetical protein | selX |
| prolipoprotein diacylglyceryl transferase | lgt |
| zinc metalloproteinase aureolysin | aur |
| hydroxyethylthiazole kinase | thiM |
| hypothetical protein | hypothetical protein |
| hypothetical protein | vraX |
| transcriptional repressor CodY | codY |
| hypothetical protein | hypothetical protein |
| hypothetical protein | hypothetical protein |
| dihydrolipoamide succinyltransferase | sucB |
| peptide ABC transporter substrate-binding protein | cntA |
| hypothetical protein | hypothetical protein |
| hypothetical protein | norG |
| hypothetical protein | sqr |
| hypothetical protein | hypothetical protein |
| antiholin-like protein LrgB | lrgB |
| hypothetical protein | hypothetical protein |
| hypothetical protein | vraA |
| sucrose operon repressor | scrR |
| hypothetical protein | hypothetical protein |
| hypothetical protein | hypothetical protein |
| hypothetical protein | recD2 |
| hypothetical protein | hypothetical protein |
| arginine deiminase | arcA |
| carbamoyl phosphate synthase large subunit | carB |
| hypothetical protein | essB |
| hypothetical protein | hypothetical protein |
| hypothetical protein | hypothetical protein |
| hypothetical protein | hypothetical protein |
| hypothetical protein | hypothetical protein |
| hypothetical protein | hypothetical protein |
| hypothetical protein | hypothetical protein |
| 2-Cys peroxiredoxin | tpx |
| alpha amylase family protein | treC |
| phospho-N-acetylmuramoyl-pentapeptide- transferase | mraY |
| hypothetical protein | menC |
| hypothetical protein | hypothetical protein |
| DnaQ family exonuclease/DinG family helicase | dinG |
| preprotein translocase subunit SecY | secY |
| hypothetical protein | hypothetical protein |
| porphobilinogen deaminase | hemC |
| superantigen-like protein | ssl4 |
| permease domain-containing protein | nsaB |
| permease | vraE |
| hypothetical protein | hypothetical protein |
| hypothetical protein | hypothetical protein |
| hypothetical protein | hypothetical protein |
| hydrolase alpha/beta fold domain-containing protein | ase alpha/beta fold domain-containing |
| molybdenum ABC transporter substrate-binding protein | modA |
| transcriptional regulator | transcriptional regulator |
| hypothetical protein | hypothetical protein |
| hypothetical protein | hypothetical protein |

| | |
|---|---|
| hypothetical protein | hypothetical protein |
| DNA polymerase IV | dinB |
| dihydrolipoamide dehydrogenase | lpdA |
| ribonuclease HIII | rnhC |
| hypothetical protein | hypothetical protein |
| hypothetical protein | nupC |
| ABC transporter ATP-binding protein | ecsA |
| hypothetical protein | hypothetical protein |
| histidine kinase | vraS |
| hypothetical protein | hypothetical protein |
| hypothetical protein | adsA |
| hypothetical protein | mpfA |
| hypothetical protein | hypothetical protein |
| hypothetical protein | hypothetical protein |
| hypothetical protein | hypothetical protein |
| alkaline phosphatase synthesis transcriptional regulatory prot | phoP |
| hypothetical protein | tcyP |
| hypothetical protein | hypothetical protein |
| hypothetical protein | hypothetical protein |
| hypothetical protein | hypothetical protein |
| phosphonate ABC transporter substrate-binding protein | phnD |
| arginine/ornithine antiporter | arcD |
| 2,3-bisphosphoglycerate-dependent phosphoglycerate mutas | gpmA |
| hypothetical protein | hypothetical protein |
| hypothetical protein | hypothetical protein |
| hypothetical protein | hypothetical protein |
| hypothetical protein | hypothetical protein |
| methionine sulfoxide reductase A | msrA3 |
| hypothetical protein | hypothetical protein |
| Trehalose operon repressor | treR |
| hypothetical protein | hypothetical protein |
| protein export protein PrsA | prsA |
| | saeS |
| cap5L protein/glycosyltransferase | capL |
| 2-deoxyribose-5-phosphate aldolase | deoC2 |
| extracellular matrix and plasma binding protein | emp |
| hypothetical protein | putA |
| hypothetical protein | hypothetical protein |
| monovalent cation/H+ antiporter subunit D | mnhD2 |
| hypothetical protein | hypothetical protein |
| RNA polymerase sigma factor SigB | sigB |
| hypothetical protein | hypothetical protein |
| hypothetical protein | hypothetical protein |
| virulence factor regulator protein | rot |
| BCCT family osmoprotectant transporter | opuD2 |
| oligopeptide ABC transporter permease | opp-4B |
| hypothetical protein | hypothetical protein |
| hypothetical protein | yjbI |
| hypothetical protein | sfnaC |
| hypothetical protein | hypothetical protein |
| response regulator | saeR |
| hypothetical protein | hypothetical protein |
| hypothetical protein | hypothetical protein |
| isocitrate dehydrogenase | citC |
| pyruvate formate-lyase 1 activating enzyme | pflA |

| | |
|---|---|
| dihydropteroate synthase | folP |
| hypothetical protein | hypothetical protein |
| hypothetical protein | hypothetical protein |
| pyrimidine-nucleoside phosphorylase | pdp |
| hypothetical protein | hypothetical protein |
| excinuclease ABC subunit B | uvrB2 |
| teichoic acid biosynthesis protein TagB | tarL |
| HTH-type transcriptional regulator SarV | sarV |
| histidine kinase | phoR |
| teichoic acid biosynthesis protein TagB | tagB |
| hypothetical protein | hypothetical protein |
| lipoprotein SirC | sirC |
| hypothetical protein | hypothetical protein |
| hypothetical protein | hypothetical protein |
| *cntM* | cntM |
| hypothetical protein | hypothetical protein |
| hypothetical protein | bglR |
| hypothetical protein | bioX |
| hypothetical protein | nrdG |
| hypothetical protein | hypothetical protein |
| alkyl hydroperoxide reductase subunit C | ahpC |
| hypothetical protein | hypothetical protein |
| hypothetical protein | hypothetical protein |
| myosin-cross-reactive antigen | myosin-cross-reactive antigen |
| lipoyltransferase and lipoate-protein ligase | lplA1 |
| PTS system glucose-specific transporter subunit IIBC | glcC |
| hypothetical protein | sbcD |
| hypothetical protein | ptpB |
| MutS domain-containing protein | MutS domain-containing protein |
| hypothetical protein | hypothetical protein |
| hypothetical protein | hypothetical protein |
| bifunctional biotin operon repressor/biotin--[acetyl-CoA-carbc | birA |
| gluconate operon transcriptional repressor | gntR |
| Multidrug efflux pump SdrM | sdrM |
| hypothetical protein | hypothetical protein |
| arginine repressor | argR |
| 16S Ribosomal RNA | rrsE |
| hypothetical protein | hypothetical protein |
| hypothetical protein | hypothetical protein |
| acetoin reductase | butA |
| hypothetical protein | hypothetical protein |
| hypothetical protein | hypothetical protein |
| hypothetical protein | hypothetical protein |
| teichoic acid biosynthesis protein | tagA |
| hypothetical protein | hypothetical protein |
| N-acetylmuramoyl-L-alanine amidase | N-acetylmuramoyl-L-alanine amidase |
| hypothetical protein | iruO |
| hypothetical protein | acuA |
| hypothetical protein | hypothetical protein |
| hypothetical protein | hypothetical protein |
| clumping factor | clfA |
| hypothetical protein | hypothetical protein |
| phosphatidate cytidylyltransferase | cdsA |
| RGD-containing lipoprotein | RGD-containing lipoprotein |
| hypothetical protein | ssuC |

| | |
|---|---|
| methionine aminopeptidase | map |
| hypothetical protein | hypothetical protein |
| DNA-binding response regulator | arlR |
| hypothetical protein | hypothetical protein |
| dihydroorotase | pyrC |
| alkyl hydroperoxide reductase subunit F | ahpF |
| hypothetical protein | mepA |
| bacteriophage L54a single-stranded DNA binding protein | ssbB |
| sodium-dependent transporter | sdcS |
| hypothetical protein | csbB |
| hypothetical protein | hypothetical protein |
| hypothetical protein | hypothetical protein |
| DNA polymerase III subunit delta' | holB |
| hypothetical protein | oatA |
| hypothetical protein | nagD |
| | comFA |
| hypothetical protein | hypothetical protein |
| hypothetical protein | hypothetical protein |
| aspartyl/glutamyl-tRNA amidotransferase subunit C | gatC |
| hypothetical protein | hypothetical protein |
| dipeptidase PepV | dipeptidase PepV |
| hypothetical protein | rbsR |
| ribulokinase | araB |
| hypothetical protein | hypothetical protein |
| ATP-dependent Clp protease ATP-binding subunit ClpC | clpL |
| hypothetical protein | gcvH-L |
| hypothetical protein | tenA |
| hypothetical protein | hypothetical protein |
| hypothetical protein | fadD |
| hypothetical protein | hypothetical protein |
| hyaluronate lyase | hysA |
| cold shock protein | cspB |
| malate:quinone oxidoreductase | mqo |
| potassium-transporting ATPase subunit A | kdpA |
| transglycosylase domain-containing protein | sgtA |
| hypothetical protein | isdD |
| LysR family transcriptional regulator | gltC |
| hypothetical protein | isdC |
| hypothetical protein | hypothetical protein |
| hypothetical protein | hypothetical protein |
| hypothetical protein | hypothetical protein |
| L-lactate dehydrogenase | ldh2 |
| proline dipeptidase | pepQ2 |
| hypothetical protein | hypothetical protein |
| hypothetical protein | hypothetical protein |
| hypothetical protein | sirTM |
| nucleoside diphosphate kinase | ndk |
| phosphopentomutase | deoB |
| oligoendopeptidase F | pepF |
| hypothetical protein | argR3 |
| peptide ABC transporter permease | cntB |
| hypothetical protein | fmhC |
| response regulator receiver domain-containing protein | hptR |
| peptidase T | peptidase T |
| hypothetical protein | hypothetical protein |

| | |
|---|---|
| immunoglobulin G-binding protein Sbi | sbi |
| hypothetical protein | hypothetical protein |
| hypothetical protein | hypothetical protein |
| hypothetical protein | hypothetical protein |
| histidine ammonia-lyase | hutH |
| 30S ribosomal protein S1 | rpsA |
| 2-succinylbenzoate-CoA ligase | menE |
| lipoprotein SirB | sirB |
| hypothetical protein | hypothetical protein |
| hypothetical protein | hypothetical protein |
| capsular polysaccharide biosynthesis protein Cap5C | cap1C |
| hypothetical protein | hypothetical protein |
| hypothetical protein | vraU |
| camphor resistance protein CrcB | crcB1 |
| acyl CoA:acetate/3-ketoacid CoA transferase | fadX |
| | agrB |
| hypothetical protein | cntE |
| superantigen-like protein | ssl11 |
| immunodominant antigen B | isaB |
| hypothetical protein | fadE |
| hypothetical protein | hypothetical protein |
| hypothetical protein | hypothetical protein |
| hypothetical protein | hypothetical protein |
| AtsA/ElaC family protein | rnz |
| hypothetical protein | hypothetical protein |
| hypothetical protein | hypothetical protein |
| hypothetical protein | hypothetical protein |
| hypothetical protein | hypothetical protein |
| two-component response regulator | lytR |
| hypothetical protein | hypothetical protein |
| ferric uptake regulator-like protein | perR |
| 5-formyltetrahydrofolate cyclo-ligase | 5-formyltetrahydrofolate cyclo-ligase |
| hypothetical protein | lnsB |
| hypothetical protein | hypothetical protein |
| superantigen-like protein | ssl12 |
| cell division protein MraZ | mraZ |
| hypothetical protein | hypothetical protein |
| hypothetical protein | hypothetical protein |
| hypothetical protein | hypothetical protein |
| anaerobic ribonucleoside triphosphate reductase | nrdD |
| N-acetylglucosamine-6-phosphate deacetylase | nagA |
| hypothetical protein | rbf |
| hypothetical protein | isdE |
| hypothetical protein | hypothetical protein |
| hypothetical protein | hypothetical protein |
| ATP-dependent DNA helicase RecQ | recQ2 |
| hypothetical protein | hypothetical protein |
| succinyl-diaminopimelate desuccinylase | succinyl-diaminopimelate desuccinylase |
| hypothetical protein | hypothetical protein |
| hypothetical protein | hypothetical protein |
| hypothetical protein | hypothetical protein |
| hypothetical protein | hypothetical protein |
| membrane protein | membrane protein |
| glutamate synthase subunit alpha | glutamate synthase subunit alpha |
| hypothetical protein | sepA |

| | |
|---|---|
| mannitol-1-phosphate 5-dehydrogenase | mtlD |
| hypothetical protein | hypothetical protein |
| oligopeptide ABC transporter ATP-binding protein | opp-2F |
| hypothetical protein | tarM |
| methicillin resistance protein FmtA | fmtA |
| Signal transduction histidine-protein kinase ArlS | arlS |
| hypothetical protein | hypothetical protein |
| hypothetical protein | fepC |
| hypothetical protein | cidR |
| glycerophosphoryl diester phosphodiesterase | cerophosphoryl diester phosphodiester |
| hypothetical protein | hypothetical protein |
| hypothetical protein | hypothetical protein |
| Alcohol dehydrogenase (quinone) | adhA |
| hypothetical protein | hypothetical protein |
| hypothetical protein | dps |
| iron compound ABC transporter permease | isdF |
| hypothetical protein | hypothetical protein |
| hypothetical protein | hypothetical protein |
| hypothetical protein | hypothetical protein |
| hypothetical protein | hypothetical protein |
| hypothetical protein | hypothetical protein |
| iron compound ABC transporter permease | on compound ABC transporter permeas |
| hypothetical protein | hypothetical protein |
| hypothetical protein | rodA |
| superantigen-like protein | ssl14 |
| hypothetical protein | hypothetical protein |
| hypothetical protein | hypothetical protein |
| hypothetical protein | hypothetical protein |
| ferritin | ftnA |
| superantigen-like protein | ssl13 |
| hypothetical protein | hypothetical protein |
| hypothetical protein | hypothetical protein |
| NADPH-dependent FMN reductase | NADPH-dependent FMN reductase |
| hypothetical protein | hypothetical protein |
| hypothetical protein | ohr |
| hypothetical protein | hypothetical protein |
| dihydroxyacetone kinase subunit DhaK | dhaK |
| hypothetical protein | veg |
| hypothetical protein | hypothetical protein |
| hypothetical protein | hypothetical protein |
| hypothetical protein | hypothetical protein |
| hypothetical protein | hypothetical protein |
| N-acetylneuraminate lyase | nanA |
| hypothetical protein | hypothetical protein |
| hypothetical protein | yjbH |
| hypothetical protein | hypothetical protein |
| | sarR |
| hex regulon repressor | glvR |
| hypothetical protein | hypothetical protein |
| hypothetical protein | hypothetical protein |
| aerobic glycerol-3-phosphate dehydrogenase | glpD |
| fosfomycin resistance protein FosB | fosB |
| hypothetical protein | hypothetical protein |
| capsular polysaccharide biosynthesis protein O-acetyl transfe | capH |
| hypothetical protein | hypothetical protein |

| | |
|---|---|
| | agrD |
| capsular polysaccharide biosynthesis protein Cap5I | capI |
| hypothetical protein | cwrA |
| urea transporter | urea transporter |
| hypothetical protein | hypothetical protein |
| hypothetical protein | relP |
| hypothetical protein | hypothetical protein |
| hypothetical protein | fhuD1 |
| hypothetical protein | hypothetical protein |
| accessory gene regulator protein C | agrC |
| hypothetical protein | nanT |
| hypothetical protein | hypothetical protein |
| 16S Ribosomal RNA | rrsF |
| hypothetical protein | hypothetical protein |
| hypothetical protein | hypothetical protein |
| hypothetical protein | hypothetical protein |
| hypothetical protein | hypothetical protein |
| capsular polysaccharide biosynthesis protein Cap5K | capK |
| hypothetical protein | hypothetical protein |
| lipoprotein | hptA |
| hypothetical protein | rnhA |
| hypothetical protein | hypothetical protein |
| UDP-N-acetylglucosamine 2-epimerase | capG |
| hypothetical protein | hypothetical protein |
| hypothetical protein | dprA |
| DNA internalization-related competence protein ComEC/Rec | comEC |
| hypothetical protein | fepB |
| hypothetical protein | hypothetical protein |
| hypothetical protein | hypothetical protein |
| hypothetical protein | hypothetical protein |
| hypothetical protein | hypothetical protein |
| DNA repair protein RecN | recN |
| hypothetical protein | htrA1 |
| hypothetical protein | hypothetical protein |
| hypothetical protein | hypothetical protein |
| hypothetical protein | hypothetical protein |
| hypothetical protein | hypothetical protein |
| peptide ABC transporter permease | peptide ABC transporter permease |
| hypothetical protein | hypothetical protein |
| aspartate carbamoyltransferase catalytic subunit | pyrB |
| hypothetical protein | dhaL |
| 5-nucleotidase family protein | 5-nucleotidase family protein |
| hypothetical protein | hypothetical protein |
| ABC transporter substrate-binding protein | mntC |
| hypothetical protein | hypothetical protein |
| hypothetical protein | hypothetical protein |
| | saeP |
| hypothetical protein | vraC |
| hypothetical protein | hypothetical protein |
| catalase | katA |
| oligopeptide transporter permease | opp-2C |
| hypothetical protein | hypothetical protein |
| hypothetical protein | pmtR |
| hypothetical protein | hypothetical protein |
| putative fluoride ion transporter crcB2 | crcB2 |

| | |
|---|---|
| hypothetical protein | comGB |
| acetoin utilization protein AcuC | acuC |
| hypothetical protein | hptS |
| cytosol aminopeptidase | pepZ |
| hypothetical protein | pgcA |
| urocanate hydratase | hutU |
| hypothetical protein | hutR |
| imidazolonepropionase | hutI |
| bifunctional pyrimidine regulatory protein PyrR/uracil phospho | pyrR |
| hypothetical protein | hypothetical protein |
| periplasmic binding protein | sirA |
| hypothetical protein | hypothetical protein |
| peptide ABC transporter ATP-binding protein | opp-2D |
| cell wall hydrolase | lytN |
| hypothetical protein | hypothetical protein |
| monovalent cation/H+ antiporter subunit A | mnhA2 |
| alpha-D-1,4-glucosidase | malA |
| hypothetical protein | hypothetical protein |
| a positive activator of SigB | rbsU |
| hypothetical protein | hypothetical protein |
| hypothetical protein | hypothetical protein |
| hypothetical protein | mroQ |
| capsular polysaccharide biosynthesis protein Cap5J | capJ |
| heme-degrading monooxygenase IsdG | isdG |
| hypothetical protein | fepA |
| capsular polysaccharide biosynthesis protein Cap8F | capF |
| amino acid ABC transporter ATP-binding protein | opuCA |
| hypothetical protein | hypothetical protein |
| hypothetical protein | epiG |
| hypothetical protein | spdB |
| oligopeptide transporter permease | opp-2B |
| hypothetical protein | map_2 |
| hypothetical protein | hypothetical protein |
| hypothetical protein | opp-5A |
| hypothetical protein | bstA |
| formate dehydrogenase | fdh |
| hypothetical protein | hypothetical protein |
| hypothetical protein | csbA |
| hypothetical protein | hypothetical protein |
| squalene synthase | crtN |
| PTS system mannnose-specific family transporter subunit IIA | dhaM |
| amino acid ABC transporter permease | opuCD |
| hypothetical protein | hypothetical protein |
| uracil permease | pyrP |
| hypothetical protein | hypothetical protein |
| hypothetical protein | hypothetical protein |
| peptide ABC transporter ATP-binding protein | cntF |
| accessory gene regulator protein A | agrA |
| hypothetical protein | hypothetical protein |
| hypothetical protein | hypothetical protein |
| capsular polysaccharide biosynthesis protein Cap5E | capE |
| hypothetical protein | abcA |
| hypothetical protein | hypothetical protein |
| ABC transporter ATP-binding protein | pmtC |
| D-ribose pyranase | rbsD |

| | |
|---|---|
| hypothetical protein | hypothetical protein |
| formyl peptide receptor-like 1 inhibitory protein | flr |
| hypothetical protein | hypothetical protein |
| PTS system ascorbate-specific transporter subunit IIC | ulaA |
| amino acid ABC transporter permease | opuCB |
| nitric oxide synthase oxygenase subunit | nos |
| hypothetical protein | hypothetical protein |
| Sortase B | srtB |
| ABC transporter ATP-binding protein | pmtA |
| para-nitrobenzyl esterase | pnbA |
| extracellular solute-binding protein | malE |
| lipase | gehA |
| MHC class II analog protein | eap/map |
| amino acid transporter | opuCC |
| nickel ABC transporter permease | cnzC |
| cell division protein | ftsL |
| hypothetical protein | hypothetical protein |
| gamma-hemolysin h-gamma-II subunit | hlgA |
| hypothetical protein | epiE |
| hypothetical protein | hypothetical protein |
| hypothetical protein | hypothetical protein |
| carbamate kinase | arcC1 |
| urease accessory protein UreD | ureD |
| hypothetical protein | csbD |
| peptide ABC transporter ATP-binding protein | cntD |
| hypothetical protein | scpB |
| hypothetical protein | hypothetical protein |
| gallidermin superfamily epiA protein | epiA |
| hypothetical protein | radC |
| hypothetical protein | hypothetical protein |
| hypothetical protein | hypothetical protein |
| hypothetical protein | hypothetical protein |
| | saeQ |
| urease accessory protein UreG | ureG |
| hypothetical protein | sfnaD |
| ABC transporter ATP-binding protein | ABC transporter ATP-binding protein |
| maltose ABC transporter permease | malF |
| hypothetical protein | hypothetical protein |
| hypothetical protein | hypothetical protein |
| hypothetical protein | pmtD |
| lactonase Drp35 | drp35 |
| hypothetical protein | comGA |
| ribokinase | rbsK |
| hypothetical protein | pheA |
| hypothetical protein | pmtB |
| hypothetical protein | hypothetical protein |
| gluconate permease | gntP |
| iron (chelated) ABC transporter permease | mntB |
| hypothetical protein | hypothetical protein |
| hypothetical protein | hypothetical protein |
| hypothetical protein | hypothetical protein |
| ABC transporter | epiF |
| intracellular serine protease | epiP |
| hypothetical protein | hypothetical protein |
| hypothetical protein | hypothetical protein |

| | |
|---|---|
| hypothetical protein | hypothetical protein |
| hypothetical protein | feoA |
| fibrinogen-binding protein SdrC | sdrC |
| hypothetical protein | mntA |
| hypothetical protein | isdA |
| capsular polysaccharide biosynthesis protein | capA |
| hypothetical protein | comC |
| capsular polysaccharide biosynthesis protein Cap5D | capD |
| | hypothetical protein |
| hypothetical protein | zagA |
| | hypothetical protein |
| | hypothetical protein |
| | hypothetical protein |
| ferrous iron transport protein B | feoB |
| multiple sugar-binding transport ATP-binding protein | malK |
| | hypothetical protein |
| sensor protein KdpD | kdpD |
| | hypothetical protein |
| hypothetical protein | rppH |
| capsular polysaccharide biosynthesis protein Cap5B | capB |
| hypothetical protein | ausA |
| squalene desaturase | crtM |
| | hypothetical protein |
| | maltose ABC transporter permease |
| urease accessory protein UreF | ureF |
| | hypothetical protein |
| gluconate kinase | gntK |
| | hypothetical protein |
| maltose operon transcriptional repressor | malR |
| capsular polysaccharide biosynthesis protein Cap8C | capC |
| mannose-6-phosphate isomerase | pmi |
| L-serine dehydratase, iron-sulfur-dependent subunit alpha | sdaAA |
| | hypothetical protein |
| | 5'-nucleotidase |
| PTS system fructose-specific transporter subunit IIABC | manP |
| ornithine carbamoyltransferase | argF |
| lipase | gehB |
| hypothetical protein | crtO |
| hypothetical protein | hypothetical protein |
| hypothetical protein | crtQ |
| | integrase/recombinase |
| urease accessory protein UreE | ureE |
| thermonuclease | nuc |
| | hypothetical protein |
| flavoprotein EpiD | epiD |
| | hypothetical protein |
| DNA-binding response regulator | kdpE |
| alpha-hemolysin | hla |
| epidermin biosynthesis protein EpiC | epiC |
| | hypothetical protein |
| | hypothetical protein |
| | hypothetical protein |

| | |
|---|---|
| staphopain thiol proteinase | scpA |
| L-serine dehydratase iron-sulfur-dependent subunit beta | sdaAB |
| Phenol-soluble modulins (PSMs) | psmβ1 |
| hypothetical protein | crtP |
| hypothetical protein | lukG |
| PTS system transporter | murP |
| Phenol-soluble modulins (PSMs) | psmβ2 |
| hypothetical protein | hypothetical protein |
| alkaline shock protein 23 | asp23 |
| lantibiotic epidermin biosynthesis protein EpiB | epiB |
| hypothetical protein | mupG |
| hypothetical protein | murR |
| Leukocidin; Leukocidin/Hemolysin toxin family | lukH |
| leukotoxin LukD | lukD |
| hypothetical protein | amaP |
| hypothetical protein | hypothetical protein |
| hypothetical protein | hypothetical protein |
| delta-hemolysin | hld |
| bifunctional acetaldehyde-CoA/alcohol dehydrogenase | adhE |
| leukotoxin LukE | lukE |
| Gamma-hemolysin component B | hlgB |
| Gamma-hemolysin component C | hlgC |
| hypothetical protein | hypothetical protein |
| hypothetical protein | hypothetical protein |
| glutamyl endopeptidase | sspA |
| cysteine protease | sspC |
| 1-phosphatidylinositol phosphodiesterase | plc |
| cysteine protease | sspB |
| Competence transcription factor | comK1 |
| serine protease SplC | splC |
| serine protease SplB | splB |
| serine protease SplA | splA |
| serine protease SplF | splF |
| serine protease SplD | splD |
| serine protease SplE | splE |

| | locus | sample_1 | sample_2 | status | value_1 | value_2 |
|---|---|---|---|---|---|---|
| SAOUHSC_02929 | NC_007795.1 | mut_A | wt_R | OK | 220.294 | 178.544 |
| SAOUHSC_02965 | NC_007795.1 | mut_A | wt_R | OK | 138.815 | 112.242 |
| SAOUHSC_01254 | NC_007795.1 | mut_A | wt_R | OK | 133.905 | 108.132 |
| SAOUHSC_00163 | NC_007795.1 | mut_A | wt_R | OK | 448.143 | 361.616 |
| SAOUHSC_00309 | NC_007795.1 | mut_A | wt_R | OK | 155.48 | 125.093 |
| SAOUHSC_00618 | NC_007795.1 | mut_A | wt_R | OK | 349.357 | 280.081 |
| SAOUHSC_00055 | NC_007795.1 | mut_A | wt_R | OK | 75.0427 | 59.9552 |
| SAOUHSC_01850 | NC_007795.1 | mut_A | wt_R | OK | 1204 | 961.623 |
| SAOUHSC_02424 | NC_007795.1 | mut_A | wt_R | OK | 363.343 | 290.196 |
| SAOUHSC_00056 | NC_007795.1 | mut_A | wt_R | OK | 97.8497 | 78.1274 |
| SAOUHSC_01053 | NC_007795.1 | mut_A | wt_R | OK | 50.6654 | 40.4442 |
| SAOUHSC_00996 | NC_007795.1 | mut_A | wt_R | OK | 1113.05 | 888.31 |
| SAOUHSC_00206 | NC_007795.1 | mut_A | wt_R | OK | 14435 | 11508.8 |
| SAOUHSC_01998 | NC_007795.1 | mut_A | wt_R | OK | 471.72 | 375.982 |
| SAOUHSC_00958 | NC_007795.1 | mut_A | wt_R | OK | 52.5419 | 41.7408 |
| SAOUHSC_01276 | NC_007795.1 | mut_A | wt_R | OK | 544.864 | 431.976 |
| SAOUHSC_00632 | NC_007795.1 | mut_A | wt_R | OK | 80.283 | 63.6391 |
| SAOUHSC_02869 | NC_007795.1 | mut_A | wt_R | OK | 323.3 | 255.572 |
| SAOUHSC_02875 | NC_007795.1 | mut_A | wt_R | OK | 40.7164 | 32.1817 |
| SAOUHSC_00674 | NC_007795.1 | mut_A | wt_R | OK | 1660.42 | 1309.85 |
| SAOUHSC_00743 | NC_007795.1 | mut_A | wt_R | OK | 1267.84 | 998.673 |
| SAOUHSC_01203 | NC_007795.1 | mut_A | wt_R | OK | 82.1663 | 64.6885 |
| SAOUHSC_00736 | NC_007795.1 | mut_A | wt_R | OK | 48.6494 | 38.2956 |
| SAOUHSC_02547 | NC_007795.1 | mut_A | wt_R | OK | 494.811 | 388.156 |
| SAOUHSC_01430 | NC_007795.1 | mut_A | wt_R | OK | 2475.72 | 1940.94 |
| SAOUHSC_02150 | NC_007795.1 | mut_A | wt_R | OK | 1513.96 | 1186.92 |
| SAOUHSC_02635 | NC_007795.1 | mut_A | wt_R | OK | 132.183 | 103.577 |
| SAOUHSC_00799 | NC_007795.1 | mut_A | wt_R | OK | 11812.8 | 9232.18 |
| SAOUHSC_00659 | NC_007795.1 | mut_A | wt_R | OK | 1295.78 | 1010.61 |
| SAOUHSC_01797 | NC_007795.1 | mut_A | wt_R | OK | 520.719 | 405.735 |
| SAOUHSC_01512 | NC_007795.1 | mut_A | wt_R | OK | 61.9178 | 48.2251 |
| SAOUHSC_02011 | NC_007795.1 | mut_A | wt_R | OK | 138.02 | 107.494 |
| SAOUHSC_00203 | NC_007795.1 | mut_A | wt_R | OK | 2953.23 | 2292.59 |
| SAOUHSC_01408 | NC_007795.1 | mut_A | wt_R | OK | 657.745 | 509.528 |
| SAOUHSC_02401 | NC_007795.1 | mut_A | wt_R | OK | 142.225 | 109.998 |
| SAOUHSC_02955 | NC_007795.1 | mut_A | wt_R | OK | 464.29 | 358.981 |
| SAOUHSC_00535 | NC_007795.1 | mut_A | wt_R | OK | 608.15 | 469.049 |
| SAOUHSC_02138 | NC_007795.1 | mut_A | wt_R | OK | 33.6011 | 25.87 |
| SAOUHSC_00943 | NC_007795.1 | mut_A | wt_R | OK | 977.874 | 752.799 |
| SAOUHSC_01310 | NC_007795.1 | mut_A | wt_R | OK | 466.195 | 358.291 |
| SAOUHSC_00406 | NC_007795.1 | mut_A | wt_R | OK | 413.1 | 317.161 |
| SAOUHSC_01226 | NC_007795.1 | mut_A | wt_R | OK | 1142.33 | 875.315 |
| SAOUHSC_00604 | NC_007795.1 | mut_A | wt_R | OK | 567.773 | 434.831 |
| SAOUHSC_01055 | NC_007795.1 | mut_A | wt_R | OK | 1740.66 | 1329.25 |
| SAOUHSC_02142 | NC_007795.1 | mut_A | wt_R | OK | 272.832 | 208.298 |
| SAOUHSC_02570 | NC_007795.1 | mut_A | wt_R | OK | 7.83939 | 5.98344 |
| SAOUHSC_02434 | NC_007795.1 | mut_A | wt_R | OK | 81.5192 | 62.1912 |
| SAOUHSC_01488 | NC_007795.1 | mut_A | wt_R | OK | 79.2138 | 60.2976 |
| SAOUHSC_00328 | NC_007795.1 | mut_A | wt_R | OK | 65.1649 | 49.3832 |
| SAOUHSC_01317 | NC_007795.1 | mut_A | wt_R | OK | 1044.99 | 791.723 |
| SAOUHSC_01775 | NC_007795.1 | mut_A | wt_R | OK | 371.115 | 280.974 |
| SAOUHSC_01845 | NC_007795.1 | mut_A | wt_R | OK | 739.295 | 559.431 |
| SAOUHSC_00469 | NC_007795.1 | mut_A | wt_R | OK | 2887.03 | 2184.02 |
| SAOUHSC_00866 | NC_007795.1 | mut_A | wt_R | OK | 488.402 | 369.31 |

| | | | | | |
|---|---|---|---|---|---|
| SAOUHSC_R0003 | NC_007795.1 mut_A | wt_R | OK | 30.0759 | 22.6932 |
| SAOUHSC_03030 | NC_007795.1 mut_A | wt_R | OK | 909.003 | 684.474 |
| SAOUHSC_00757 | NC_007795.1 mut_A | wt_R | OK | 563.178 | 423.958 |
| SAOUHSC_01958 | NC_007795.1 mut_A | wt_R | OK | 89.6576 | 67.4115 |
| SAOUHSC_00776 | NC_007795.1 mut_A | wt_R | OK | 430.956 | 321.858 |
| SAOUHSC_02367 | NC_007795.1 mut_A | wt_R | OK | 78.0026 | 58.1478 |
| SAOUHSC_00851 | NC_007795.1 mut_A | wt_R | OK | 1688.96 | 1258.63 |
| SAOUHSC_01724 | NC_007795.1 mut_A | wt_R | OK | 93.0829 | 69.3203 |
| SAOUHSC_00130 | NC_007795.1 mut_A | wt_R | OK | 148.065 | 110.185 |
| SAOUHSC_02909 | NC_007795.1 mut_A | wt_R | OK | 158.459 | 117.476 |
| SAOUHSC_02409 | NC_007795.1 mut_A | wt_R | OK | 203.63 | 150.923 |
| SAOUHSC_00780 | NC_007795.1 mut_A | wt_R | OK | 746.23 | 552.471 |
| SAOUHSC_00200 | NC_007795.1 mut_A | wt_R | OK | 56.8579 | 42.015 |
| SAOUHSC_00468 | NC_007795.1 mut_A | wt_R | OK | 1273.58 | 940.874 |
| SAOUHSC_00133 | NC_007795.1 mut_A | wt_R | OK | 38.186 | 28.2099 |
| SAOUHSC_02568 | NC_007795.1 mut_A | wt_R | OK | 380.447 | 280.865 |
| SAOUHSC_02324 | NC_007795.1 mut_A | wt_R | OK | 155.431 | 114.723 |
| SAOUHSC_01796 | NC_007795.1 mut_A | wt_R | OK | 399.733 | 294.944 |
| SAOUHSC_01781 | NC_007795.1 mut_A | wt_R | OK | 349.147 | 256.89 |
| SAOUHSC_02736 | NC_007795.1 mut_A | wt_R | OK | 93.2127 | 68.5714 |
| SAOUHSC_02577 | NC_007795.1 mut_A | wt_R | OK | 1189.36 | 874.597 |
| SAOUHSC_00223 | NC_007795.1 mut_A | wt_R | OK | 161.956 | 119.087 |
| SAOUHSC_01418 | NC_007795.1 mut_A | wt_R | OK | 976.761 | 717.738 |
| SAOUHSC_00919 | NC_007795.1 mut_A | wt_R | OK | 3399.19 | 2497.27 |
| SAOUHSC_01846 | NC_007795.1 mut_A | wt_R | OK | 45.7836 | 33.6293 |
| SAOUHSC_00100 | NC_007795.1 mut_A | wt_R | OK | 5604.5 | 4111.65 |
| SAOUHSC_01133 | NC_007795.1 mut_A | wt_R | OK | 149.206 | 109.435 |
| SAOUHSC_01867 | NC_007795.1 mut_A | wt_R | OK | 680.257 | 498.07 |
| SAOUHSC_01900 | NC_007795.1 mut_A | wt_R | OK | 22.9862 | 16.8269 |
| SAOUHSC_01819 | NC_007795.1 mut_A | wt_R | OK | 7762.33 | 5675.99 |
| SAOUHSC_02980 | NC_007795.1 mut_A | wt_R | OK | 1020.88 | 746.188 |
| SAOUHSC_00538 | NC_007795.1 mut_A | wt_R | OK | 128.09 | 93.371 |
| SAOUHSC_00612 | NC_007795.1 mut_A | wt_R | OK | 13.8187 | 10.0671 |
| SAOUHSC_00052 | NC_007795.1 mut_A | wt_R | OK | 31.4759 | 22.9036 |
| SAOUHSC_02842 | NC_007795.1 mut_A | wt_R | OK | 425.492 | 309.139 |
| SAOUHSC_01910 | NC_007795.1 mut_A | wt_R | OK | 798.405 | 579.444 |
| SAOUHSC_02147 | NC_007795.1 mut_A | wt_R | OK | 164.505 | 119.317 |
| SAOUHSC_01861 | NC_007795.1 mut_A | wt_R | OK | 227.232 | 164.626 |
| SAOUHSC_01280 | NC_007795.1 mut_A | wt_R | OK | 152.706 | 110.557 |
| SAOUHSC_02472 | NC_007795.1 mut_A | wt_R | OK | 256.737 | 185.494 |
| SAOUHSC_02555 | NC_007795.1 mut_A | wt_R | OK | 198.788 | 143.452 |
| SAOUHSC_00099 | NC_007795.1 mut_A | wt_R | OK | 4346.75 | 3135.39 |
| SAOUHSC_02515 | NC_007795.1 mut_A | wt_R | OK | 304.402 | 219.499 |
| SAOUHSC_01275 | NC_007795.1 mut_A | wt_R | OK | 2143.56 | 1543.33 |
| SAOUHSC_00258 | NC_007795.1 mut_A | wt_R | OK | 168.787 | 121.449 |
| SAOUHSC_01319 | NC_007795.1 mut_A | wt_R | OK | 6.06517 | 4.36343 |
| SAOUHSC_00897 | NC_007795.1 mut_A | wt_R | OK | 113.481 | 81.5389 |
| SAOUHSC_02968 | NC_007795.1 mut_A | wt_R | OK | 96.1721 | 69.0968 |
| SAOUHSC_00401 | NC_007795.1 mut_A | wt_R | OK | 182.576 | 131.036 |
| SAOUHSC_00890 | NC_007795.1 mut_A | wt_R | OK | 50.297 | 36.0806 |
| SAOUHSC_01612 | NC_007795.1 mut_A | wt_R | OK | 759.963 | 544.096 |
| SAOUHSC_01405 | NC_007795.1 mut_A | wt_R | OK | 318.349 | 227.875 |
| SAOUHSC_01069 | NC_007795.1 mut_A | wt_R | OK | 175.896 | 125.811 |
| SAOUHSC_03031 | NC_007795.1 mut_A | wt_R | OK | 70.2103 | 50.1904 |
| SAOUHSC_01433 | NC_007795.1 mut_A | wt_R | OK | 412.93 | 294.952 |

| | | | | | |
|---|---|---|---|---|---|
| SAOUHSC_02112 | NC_007795.1 mut_A | wt_R | OK | 211.762 | 151.217 |
| SAOUHSC_00094 | NC_007795.1 mut_A | wt_R | OK | 15767 | 11241.9 |
| SAOUHSC_01611 | NC_007795.1 mut_A | wt_R | OK | 854.337 | 608.998 |
| SAOUHSC_00354 | NC_007795.1 mut_A | wt_R | OK | 51.0256 | 36.3213 |
| SAOUHSC_00782 | NC_007795.1 mut_A | wt_R | OK | 242.123 | 172.198 |
| SAOUHSC_02971 | NC_007795.1 mut_A | wt_R | OK | 76.7144 | 54.5085 |
| SAOUHSC_02329 | NC_007795.1 mut_A | wt_R | OK | 118.552 | 84.1393 |
| SAOUHSC_00030 | NC_007795.1 mut_A | wt_R | OK | 38.0609 | 27.0113 |
| SAOUHSC_00561 | NC_007795.1 mut_A | wt_R | OK | 10296.3 | 7267.74 |
| SAOUHSC_01228 | NC_007795.1 mut_A | wt_R | OK | 1408.68 | 993.279 |
| SAOUHSC_02853 | NC_007795.1 mut_A | wt_R | OK | 35703.4 | 25166.2 |
| SAOUHSC_00172 | NC_007795.1 mut_A | wt_R | OK | 89.8112 | 63.2665 |
| SAOUHSC_01416 | NC_007795.1 mut_A | wt_R | OK | 755.39 | 531.944 |
| SAOUHSC_02767 | NC_007795.1 mut_A | wt_R | OK | 71.5204 | 50.3555 |
| SAOUHSC_00141 | NC_007795.1 mut_A | wt_R | OK | 1418.42 | 998.65 |
| SAOUHSC_00064 | NC_007795.1 mut_A | wt_R | OK | 7.19525 | 5.05499 |
| SAOUHSC_00037 | NC_007795.1 mut_A | wt_R | OK | 54.2094 | 38.0686 |
| SAOUHSC_02006 | NC_007795.1 mut_A | wt_R | OK | 418.046 | 293.557 |
| SAOUHSC_00233 | NC_007795.1 mut_A | wt_R | OK | 520.403 | 364.855 |
| SAOUHSC_00774 | NC_007795.1 mut_A | wt_R | OK | 193.768 | 135.747 |
| SAOUHSC_00557 | NC_007795.1 mut_A | wt_R | OK | 6.16447 | 4.31839 |
| SAOUHSC_02269 | NC_007795.1 mut_A | wt_R | OK | 27.8011 | 19.4723 |
| SAOUHSC_01414 | NC_007795.1 mut_A | wt_R | OK | 1776.23 | 1241.82 |
| SAOUHSC_01761 | NC_007795.1 mut_A | wt_R | OK | 325.41 | 227.12 |
| SAOUHSC_01723 | NC_007795.1 mut_A | wt_R | OK | 115.283 | 80.4485 |
| SAOUHSC_00830 | NC_007795.1 mut_A | wt_R | OK | 1439.66 | 1004.51 |
| SAOUHSC_02969 | NC_007795.1 mut_A | wt_R | OK | 96.6171 | 67.325 |
| SAOUHSC_01170 | NC_007795.1 mut_A | wt_R | OK | 347.775 | 242.088 |
| SAOUHSC_00261 | NC_007795.1 mut_A | wt_R | OK | 60.9166 | 42.3208 |
| SAOUHSC_00322 | NC_007795.1 mut_A | wt_R | OK | 67.6231 | 46.8711 |
| SAOUHSC_02585 | NC_007795.1 mut_A | wt_R | OK | 574.526 | 398.065 |
| SAOUHSC_02900 | NC_007795.1 mut_A | wt_R | OK | 113.992 | 78.9717 |
| SAOUHSC_01957 | NC_007795.1 mut_A | wt_R | OK | 305.425 | 211.081 |
| SAOUHSC_01831 | NC_007795.1 mut_A | wt_R | OK | 287.833 | 198.816 |
| SAOUHSC_01324 | NC_007795.1 mut_A | wt_R | OK | 1742.12 | 1201.87 |
| SAOUHSC_01822 | NC_007795.1 mut_A | wt_R | OK | 2109.87 | 1449.77 |
| SAOUHSC_00438 | NC_007795.1 mut_A | wt_R | OK | 410.259 | 280.959 |
| SAOUHSC_01146 | NC_007795.1 mut_A | wt_R | OK | 132.711 | 90.6666 |
| SAOUHSC_01915 | NC_007795.1 mut_A | wt_R | OK | 146.664 | 100.164 |
| SAOUHSC_00565 | NC_007795.1 mut_A | wt_R | OK | 116.381 | 79.3737 |
| SAOUHSC_01472 | NC_007795.1 mut_A | wt_R | OK | 172.977 | 117.401 |
| SAOUHSC_02491 | NC_007795.1 mut_A | wt_R | OK | 22.5765 | 15.3118 |
| SAOUHSC_01005 | NC_007795.1 mut_A | wt_R | OK | 1678.95 | 1137.17 |
| SAOUHSC_01774 | NC_007795.1 mut_A | wt_R | OK | 363.133 | 245.72 |
| SAOUHSC_00389 | NC_007795.1 mut_A | wt_R | OK | 5.02466 | 3.39935 |
| SAOUHSC_02953 | NC_007795.1 mut_A | wt_R | OK | 128.841 | 87.1478 |
| SAOUHSC_03037 | NC_007795.1 mut_A | wt_R | OK | 10.1603 | 6.86217 |
| SAOUHSC_02143 | NC_007795.1 mut_A | wt_R | OK | 299.705 | 202.408 |
| SAOUHSC_02396 | NC_007795.1 mut_A | wt_R | OK | 470.148 | 317.38 |
| SAOUHSC_01349 | NC_007795.1 mut_A | wt_R | OK | 228.351 | 154.107 |
| SAOUHSC_01279 | NC_007795.1 mut_A | wt_R | OK | 129.2 | 87.1398 |
| SAOUHSC_02549 | NC_007795.1 mut_A | wt_R | OK | 457.222 | 307.467 |
| SAOUHSC_02583 | NC_007795.1 mut_A | wt_R | OK | 994.89 | 668.863 |
| SAOUHSC_00280 | NC_007795.1 mut_A | wt_R | OK | 741.206 | 497.607 |
| SAOUHSC_00599 | NC_007795.1 mut_A | wt_R | OK | 41.2007 | 27.599 |

| | | | | | |
|---|---|---|---|---|---|
| SAOUHSC_02892 | NC_007795.1 mut_A | wt_R | OK | 83.6455 | 55.9612 |
| SAOUHSC_02111 | NC_007795.1 mut_A | wt_R | OK | 123.211 | 82.4042 |
| SAOUHSC_01614 | NC_007795.1 mut_A | wt_R | OK | 577.793 | 385.579 |
| SAOUHSC_01095 | NC_007795.1 mut_A | wt_R | OK | 71.3502 | 47.6046 |
| SAOUHSC_02995 | NC_007795.1 mut_A | wt_R | OK | 53.4669 | 35.6494 |
| SAOUHSC_00501 | NC_007795.1 mut_A | wt_R | OK | 290.244 | 193.349 |
| SAOUHSC_01967 | NC_007795.1 mut_A | wt_R | OK | 216.855 | 144.437 |
| SAOUHSC_02908 | NC_007795.1 mut_A | wt_R | OK | 50.8234 | 33.8427 |
| SAOUHSC_02099 | NC_007795.1 mut_A | wt_R | OK | 309.883 | 206.216 |
| SAOUHSC_00508 | NC_007795.1 mut_A | wt_R | OK | 554.863 | 368.998 |
| SAOUHSC_00025 | NC_007795.1 mut_A | wt_R | OK | 46.8628 | 31.1176 |
| SAOUHSC_00711 | NC_007795.1 mut_A | wt_R | OK | 338.182 | 224.468 |
| SAOUHSC_00084 | NC_007795.1 mut_A | wt_R | OK | 58.4711 | 38.7793 |
| SAOUHSC_02727 | NC_007795.1 mut_A | wt_R | OK | 543.177 | 359.757 |
| SAOUHSC_01304 | NC_007795.1 mut_A | wt_R | OK | 385.732 | 255.392 |
| SAOUHSC_01800 | NC_007795.1 mut_A | wt_R | OK | 313.19 | 206.641 |
| SAOUHSC_00367 | NC_007795.1 mut_A | wt_R | OK | 1921.59 | 1266.53 |
| SAOUHSC_01859 | NC_007795.1 mut_A | wt_R | OK | 219.912 | 144.812 |
| SAOUHSC_00633 | NC_007795.1 mut_A | wt_R | OK | 298.183 | 196.127 |
| SAOUHSC_02650 | NC_007795.1 mut_A | wt_R | OK | 949.067 | 624.179 |
| SAOUHSC_00105 | NC_007795.1 mut_A | wt_R | OK | 11.9908 | 7.88585 |
| SAOUHSC_02967 | NC_007795.1 mut_A | wt_R | OK | 90.9532 | 59.806 |
| SAOUHSC_02703 | NC_007795.1 mut_A | wt_R | OK | 2201.17 | 1439.85 |
| SAOUHSC_01256 | NC_007795.1 mut_A | wt_R | OK | 135.45 | 88.3543 |
| SAOUHSC_00922 | NC_007795.1 mut_A | wt_R | OK | 35.2933 | 23.0057 |
| SAOUHSC_02913 | NC_007795.1 mut_A | wt_R | OK | 67.6561 | 43.973 |
| SAOUHSC_02845 | NC_007795.1 mut_A | wt_R | OK | 821.478 | 533.415 |
| SAOUHSC_02996 | NC_007795.1 mut_A | wt_R | OK | 17.254 | 11.2015 |
| SAOUHSC_00976 | NC_007795.1 mut_A | wt_R | OK | 15.3432 | 9.95458 |
| SAOUHSC_00439 | NC_007795.1 mut_A | wt_R | OK | 480.996 | 311.309 |
| SAOUHSC_00270 | NC_007795.1 mut_A | wt_R | OK | 91.3413 | 59.1161 |
| SAOUHSC_01972 | NC_007795.1 mut_A | wt_R | OK | 1055.15 | 680.246 |
| SAOUHSC_00714 | NC_007795.1 mut_A | wt_R | OK | 1142.87 | 736.38 |
| SAOUHSC_00125 | NC_007795.1 mut_A | wt_R | OK | 31.9509 | 20.5471 |
| SAOUHSC_02379 | NC_007795.1 mut_A | wt_R | OK | 944.92 | 607.077 |
| SAOUHSC_00816 | NC_007795.1 mut_A | wt_R | OK | 17.5372 | 11.2589 |
| SAOUHSC_01884 | NC_007795.1 mut_A | wt_R | OK | 320.689 | 205.822 |
| SAOUHSC_01971 | NC_007795.1 mut_A | wt_R | OK | 11.8283 | 7.5907 |
| SAOUHSC_00628 | NC_007795.1 mut_A | wt_R | OK | 34.845 | 22.3549 |
| SAOUHSC_01463 | NC_007795.1 mut_A | wt_R | OK | 841.974 | 539.985 |
| SAOUHSC_02298 | NC_007795.1 mut_A | wt_R | OK | 532.967 | 341.494 |
| SAOUHSC_02393 | NC_007795.1 mut_A | wt_R | OK | 90.1269 | 57.6563 |
| SAOUHSC_00621 | NC_007795.1 mut_A | wt_R | OK | 40.8536 | 26.1046 |
| SAOUHSC_01879 | NC_007795.1 mut_A | wt_R | OK | 1186.04 | 756.195 |
| SAOUHSC_02444 | NC_007795.1 mut_A | wt_R | OK | 9.52316 | 6.06528 |
| SAOUHSC_00931 | NC_007795.1 mut_A | wt_R | OK | 11.4891 | 7.30772 |
| SAOUHSC_00602 | NC_007795.1 mut_A | wt_R | OK | 35.5185 | 22.5375 |
| SAOUHSC_00939 | NC_007795.1 mut_A | wt_R | OK | 513.047 | 324.793 |
| SAOUHSC_02433 | NC_007795.1 mut_A | wt_R | OK | 57.7768 | 36.4572 |
| SAOUHSC_02827 | NC_007795.1 mut_A | wt_R | OK | 732.822 | 461.59 |
| SAOUHSC_00715 | NC_007795.1 mut_A | wt_R | OK | 1471.9 | 926.126 |
| SAOUHSC_01649 | NC_007795.1 mut_A | wt_R | OK | 386.101 | 242.824 |
| SAOUHSC_01508 | NC_007795.1 mut_A | wt_R | OK | 73.8166 | 46.3934 |
| SAOUHSC_01801 | NC_007795.1 mut_A | wt_R | OK | 462.002 | 289.82 |
| SAOUHSC_00188 | NC_007795.1 mut_A | wt_R | OK | 10483.3 | 6573.27 |

| | | | | | |
|---|---|---|---|---|---|
| SAOUHSC_00489 | NC_007795.1 mut_A | wt_R | OK | 117.437 | 73.5576 |
| SAOUHSC_00672 | NC_007795.1 mut_A | wt_R | OK | 233.151 | 146.016 |
| SAOUHSC_01464 | NC_007795.1 mut_A | wt_R | OK | 1207.65 | 756.045 |
| SAOUHSC_02377 | NC_007795.1 mut_A | wt_R | OK | 709.67 | 443.02 |
| SAOUHSC_01296 | NC_007795.1 mut_A | wt_R | OK | 6.58423 | 4.10713 |
| SAOUHSC_00779 | NC_007795.1 mut_A | wt_R | OK | 527.088 | 327.831 |
| SAOUHSC_00222 | NC_007795.1 mut_A | wt_R | OK | 111.98 | 69.6414 |
| SAOUHSC_02532 | NC_007795.1 mut_A | wt_R | OK | 283.472 | 176.195 |
| SAOUHSC_01799 | NC_007795.1 mut_A | wt_R | OK | 282.199 | 175.2 |
| SAOUHSC_00643 | NC_007795.1 mut_A | wt_R | OK | 88.4507 | 54.817 |
| SAOUHSC_00598 | NC_007795.1 mut_A | wt_R | OK | 14.9193 | 9.16816 |
| SAOUHSC_00071 | NC_007795.1 mut_A | wt_R | OK | 40.4676 | 24.8174 |
| SAOUHSC_03047 | NC_007795.1 mut_A | wt_R | OK | 253.635 | 154.604 |
| SAOUHSC_02775 | NC_007795.1 mut_A | wt_R | OK | 112.907 | 68.8169 |
| SAOUHSC_02768 | NC_007795.1 mut_A | wt_R | OK | 7.55563 | 4.60333 |
| SAOUHSC_02797 | NC_007795.1 mut_A | wt_R | OK | 40.0775 | 24.4146 |
| SAOUHSC_00234 | NC_007795.1 mut_A | wt_R | OK | 103.564 | 63.0015 |
| SAOUHSC_02711 | NC_007795.1 mut_A | wt_R | OK | 9.04618 | 5.48533 |
| SAOUHSC_02941 | NC_007795.1 mut_A | wt_R | OK | 337.637 | 204.462 |
| SAOUHSC_01134 | NC_007795.1 mut_A | wt_R | OK | 62.7518 | 37.9773 |
| SAOUHSC_00365 | NC_007795.1 mut_A | wt_R | OK | 7823.45 | 4733.4 |
| SAOUHSC_01798 | NC_007795.1 mut_A | wt_R | OK | 22.0974 | 13.3365 |
| SAOUHSC_00139 | NC_007795.1 mut_A | wt_R | OK | 419.912 | 253.307 |
| SAOUHSC_00061 | NC_007795.1 mut_A | wt_R | OK | 33.1365 | 19.9673 |
| SAOUHSC_00963 | NC_007795.1 mut_A | wt_R | OK | 325.377 | 196.009 |
| SAOUHSC_00209 | NC_007795.1 mut_A | wt_R | OK | 90.9995 | 54.7864 |
| SAOUHSC_01341 | NC_007795.1 mut_A | wt_R | OK | 70.651 | 42.4644 |
| SAOUHSC_02356 | NC_007795.1 mut_A | wt_R | OK | 101.253 | 60.8546 |
| SAOUHSC_02276 | NC_007795.1 mut_A | wt_R | OK | 31.7785 | 19.0979 |
| SAOUHSC_00042 | NC_007795.1 mut_A | wt_R | OK | 26.7754 | 16.0695 |
| SAOUHSC_02781 | NC_007795.1 mut_A | wt_R | OK | 21887 | 13112.7 |
| SAOUHSC_01473 | NC_007795.1 mut_A | wt_R | OK | 169.116 | 101.276 |
| SAOUHSC_02809 | NC_007795.1 mut_A | wt_R | OK | 561.174 | 335.886 |
| SAOUHSC_02420 | NC_007795.1 mut_A | wt_R | OK | 189.628 | 113.227 |
| SAOUHSC_01511 | NC_007795.1 mut_A | wt_R | OK | 23.0061 | 13.7305 |
| SAOUHSC_01617 | NC_007795.1 mut_A | wt_R | OK | 406.966 | 241.692 |
| SAOUHSC_R0004 | NC_007795.1 mut_A | wt_R | OK | 3.56432 | 2.11223 |
| SAOUHSC_02471 | NC_007795.1 mut_A | wt_R | OK | 123.467 | 73.1449 |
| SAOUHSC_02008 | NC_007795.1 mut_A | wt_R | OK | 129.138 | 76.4356 |
| SAOUHSC_00086 | NC_007795.1 mut_A | wt_R | OK | 1375.03 | 813.666 |
| SAOUHSC_01860 | NC_007795.1 mut_A | wt_R | OK | 726.291 | 429.359 |
| SAOUHSC_01830 | NC_007795.1 mut_A | wt_R | OK | 77.8376 | 46.0048 |
| SAOUHSC_00696 | NC_007795.1 mut_A | wt_R | OK | 183.807 | 108.62 |
| SAOUHSC_00640 | NC_007795.1 mut_A | wt_R | OK | 81.9318 | 48.4058 |
| SAOUHSC_00913 | NC_007795.1 mut_A | wt_R | OK | 5.0355 | 2.97029 |
| SAOUHSC_02979 | NC_007795.1 mut_A | wt_R | OK | 931.268 | 549.116 |
| SAOUHSC_02654 | NC_007795.1 mut_A | wt_R | OK | 53.7352 | 31.6447 |
| SAOUHSC_01847 | NC_007795.1 mut_A | wt_R | OK | 170.394 | 100.321 |
| SAOUHSC_01966 | NC_007795.1 mut_A | wt_R | OK | 242.34 | 142.387 |
| SAOUHSC_00810 | NC_007795.1 mut_A | wt_R | OK | 9.57105 | 5.61409 |
| SAOUHSC_00812 | NC_007795.1 mut_A | wt_R | OK | 179.72 | 105.414 |
| SAOUHSC_01376 | NC_007795.1 mut_A | wt_R | OK | 71.4447 | 41.775 |
| SAOUHSC_01238 | NC_007795.1 mut_A | wt_R | OK | 153.663 | 89.7839 |
| SAOUHSC_00170 | NC_007795.1 mut_A | wt_R | OK | 26.174 | 15.2896 |
| SAOUHSC_00138 | NC_007795.1 mut_A | wt_R | OK | 298.948 | 174.456 |

| | | | | | |
|---|---|---|---|---|---|
| SAOUHSC_02102 | NC_007795.1 mut_A | wt_R | OK | 417.465 | 243.286 |
| SAOUHSC_01113 | NC_007795.1 mut_A | wt_R | OK | 145.52 | 84.6779 |
| SAOUHSC_01420 | NC_007795.1 mut_A | wt_R | OK | 690.191 | 400.855 |
| SAOUHSC_00697 | NC_007795.1 mut_A | wt_R | OK | 75.1285 | 43.6145 |
| SAOUHSC_01168 | NC_007795.1 mut_A | wt_R | OK | 279.838 | 162.217 |
| SAOUHSC_00364 | NC_007795.1 mut_A | wt_R | OK | 6207.64 | 3595.38 |
| SAOUHSC_00315 | NC_007795.1 mut_A | wt_R | OK | 35.0472 | 20.2829 |
| SAOUHSC_02334 | NC_007795.1 mut_A | wt_R | OK | 75.3873 | 43.5546 |
| SAOUHSC_02137 | NC_007795.1 mut_A | wt_R | OK | 36.9902 | 21.3668 |
| SAOUHSC_00713 | NC_007795.1 mut_A | wt_R | OK | 215.898 | 124.539 |
| SAOUHSC_02001 | NC_007795.1 mut_A | wt_R | OK | 637.454 | 367.518 |
| SAOUHSC_03017 | NC_007795.1 mut_A | wt_R | OK | 18.0792 | 10.4086 |
| SAOUHSC_00454 | NC_007795.1 mut_A | wt_R | OK | 82.9905 | 47.7384 |
| SAOUHSC_02885 | NC_007795.1 mut_A | wt_R | OK | 390.704 | 224.331 |
| SAOUHSC_00865 | NC_007795.1 mut_A | wt_R | OK | 323.343 | 185.532 |
| SAOUHSC_00765 | NC_007795.1 mut_A | wt_R | OK | 5.40992 | 3.1039 |
| SAOUHSC_00431 | NC_007795.1 mut_A | wt_R | OK | 159.838 | 91.2587 |
| SAOUHSC_02464 | NC_007795.1 mut_A | wt_R | OK | 222.746 | 127.123 |
| SAOUHSC_02118 | NC_007795.1 mut_A | wt_R | OK | 178.994 | 102.147 |
| SAOUHSC_1307a | NC_007795.1 mut_A | wt_R | OK | 58.8179 | 33.5546 |
| SAOUHSC_01868 | NC_007795.1 mut_A | wt_R | OK | 341.699 | 194.79 |
| SAOUHSC_00242 | NC_007795.1 mut_A | wt_R | OK | 30.7607 | 17.4827 |
| SAOUHSC_00534 | NC_007795.1 mut_A | wt_R | OK | 47.3229 | 26.7724 |
| SAOUHSC_02376 | NC_007795.1 mut_A | wt_R | OK | 1774.11 | 1002.74 |
| SAOUHSC_02862 | NC_007795.1 mut_A | wt_R | OK | 149.435 | 84.2789 |
| SAOUHSC_00305 | NC_007795.1 mut_A | wt_R | OK | 76.1623 | 42.9191 |
| SAOUHSC_02331 | NC_007795.1 mut_A | wt_R | OK | 69.136 | 38.9079 |
| SAOUHSC_00272 | NC_007795.1 mut_A | wt_R | OK | 233.107 | 131.153 |
| SAOUHSC_00197 | NC_007795.1 mut_A | wt_R | OK | 9.86031 | 5.54682 |
| SAOUHSC_02726 | NC_007795.1 mut_A | wt_R | OK | 15.8895 | 8.9344 |
| SAOUHSC_02463 | NC_007795.1 mut_A | wt_R | OK | 56.0332 | 31.479 |
| SAOUHSC_03045 | NC_007795.1 mut_A | wt_R | OK | 4808.41 | 2700.14 |
| SAOUHSC_02647 | NC_007795.1 mut_A | wt_R | OK | 401.833 | 225.533 |
| SAOUHSC_02312 | NC_007795.1 mut_A | wt_R | OK | 6.38584 | 3.58305 |
| SAOUHSC_01840 | NC_007795.1 mut_A | wt_R | OK | 46.2909 | 25.9718 |
| SAOUHSC_01084 | NC_007795.1 mut_A | wt_R | OK | 16.0519 | 9.003 |
| SAOUHSC_00434 | NC_007795.1 mut_A | wt_R | OK | 16.4003 | 9.19698 |
| SAOUHSC_01082 | NC_007795.1 mut_A | wt_R | OK | 20.4195 | 11.4356 |
| SAOUHSC_01926 | NC_007795.1 mut_A | wt_R | OK | 158.458 | 88.4075 |
| SAOUHSC_02144 | NC_007795.1 mut_A | wt_R | OK | 124.801 | 69.6079 |
| SAOUHSC_02596 | NC_007795.1 mut_A | wt_R | OK | 35.4462 | 19.6785 |
| SAOUHSC_02922 | NC_007795.1 mut_A | wt_R | OK | 1009.82 | 560.18 |
| SAOUHSC_01626 | NC_007795.1 mut_A | wt_R | OK | 541.766 | 299.668 |
| SAOUHSC_00408 | NC_007795.1 mut_A | wt_R | OK | 253.31 | 139.951 |
| SAOUHSC_00825 | NC_007795.1 mut_A | wt_R | OK | 208.871 | 115.391 |
| SAOUHSC_00307 | NC_007795.1 mut_A | wt_R | OK | 39.375 | 21.7229 |
| SAOUHSC_01485 | NC_007795.1 mut_A | wt_R | OK | 562.019 | 309.57 |
| SAOUHSC_00101 | NC_007795.1 mut_A | wt_R | OK | 8832.23 | 4853.8 |
| SAOUHSC_00937 | NC_007795.1 mut_A | wt_R | OK | 133.521 | 73.3333 |
| SAOUHSC_02970 | NC_007795.1 mut_A | wt_R | OK | 262.885 | 143.657 |
| SAOUHSC_02766 | NC_007795.1 mut_A | wt_R | OK | 32.6084 | 17.8033 |
| SAOUHSC_01220 | NC_007795.1 mut_A | wt_R | OK | 28.3659 | 15.4588 |
| SAOUHSC_00184 | NC_007795.1 mut_A | wt_R | OK | 133.192 | 72.4591 |
| SAOUHSC_01606 | NC_007795.1 mut_A | wt_R | OK | 324.738 | 176.514 |
| SAOUHSC_00146 | NC_007795.1 mut_A | wt_R | OK | 5045.5 | 2738.73 |

| | | | | | | |
|---|---|---|---|---|---|---|
| SAOUHSC_02706 | NC_007795.1 | mut_A | wt_R | OK | 439.12 | 238.135 |
| SAOUHSC_00271 | NC_007795.1 | mut_A | wt_R | OK | 65.5888 | 35.5161 |
| SAOUHSC_01292 | NC_007795.1 | mut_A | wt_R | OK | 53.7639 | 29.1077 |
| SAOUHSC_02009 | NC_007795.1 | mut_A | wt_R | OK | 86.0501 | 46.5725 |
| SAOUHSC_00008 | NC_007795.1 | mut_A | wt_R | OK | 67.0104 | 36.1763 |
| SAOUHSC_01493 | NC_007795.1 | mut_A | wt_R | OK | 3883.19 | 2094.87 |
| SAOUHSC_01916 | NC_007795.1 | mut_A | wt_R | OK | 82.7153 | 44.2704 |
| SAOUHSC_00072 | NC_007795.1 | mut_A | wt_R | OK | 26.1406 | 13.9831 |
| SAOUHSC_00415 | NC_007795.1 | mut_A | wt_R | OK | 17.9073 | 9.57577 |
| SAOUHSC_00823 | NC_007795.1 | mut_A | wt_R | OK | 442.314 | 236.156 |
| SAOUHSC_02998 | NC_007795.1 | mut_A | wt_R | OK | 17.9892 | 9.57611 |
| SAOUHSC_00587 | NC_007795.1 | mut_A | wt_R | OK | 47.714 | 25.3777 |
| SAOUHSC_02101 | NC_007795.1 | mut_A | wt_R | OK | 96.7924 | 51.4693 |
| SAOUHSC_01903 | NC_007795.1 | mut_A | wt_R | OK | 22.5062 | 11.9559 |
| SAOUHSC_00199 | NC_007795.1 | mut_A | wt_R | OK | 36.9758 | 19.631 |
| SAOUHSC_02261 | NC_007795.1 | mut_A | wt_R | OK | 9880.19 | 5228.28 |
| SAOUHSC_02762 | NC_007795.1 | mut_A | wt_R | OK | 57.5488 | 30.4256 |
| SAOUHSC_00399 | NC_007795.1 | mut_A | wt_R | OK | 13.2213 | 6.98001 |
| SAOUHSC_02972 | NC_007795.1 | mut_A | wt_R | OK | 985.006 | 519.684 |
| SAOUHSC_00198 | NC_007795.1 | mut_A | wt_R | OK | 25.1644 | 13.2632 |
| SAOUHSC_02653 | NC_007795.1 | mut_A | wt_R | OK | 22.4125 | 11.7893 |
| SAOUHSC_00917 | NC_007795.1 | mut_A | wt_R | OK | 27.0291 | 14.2043 |
| SAOUHSC_01305 | NC_007795.1 | mut_A | wt_R | OK | 86.9647 | 45.6665 |
| SAOUHSC_01598 | NC_007795.1 | mut_A | wt_R | OK | 124.841 | 65.5092 |
| SAOUHSC_02772 | NC_007795.1 | mut_A | wt_R | OK | 19.9469 | 10.4515 |
| SAOUHSC_02149 | NC_007795.1 | mut_A | wt_R | OK | 49.9183 | 26.0976 |
| SAOUHSC_A01081 | NC_007795.1 | mut_A | wt_R | OK | 77.2697 | 40.3349 |
| SAOUHSC_02104 | NC_007795.1 | mut_A | wt_R | OK | 17.5075 | 9.13818 |
| SAOUHSC_00231 | NC_007795.1 | mut_A | wt_R | OK | 53.9695 | 28.1132 |
| SAOUHSC_00758 | NC_007795.1 | mut_A | wt_R | OK | 262.528 | 136.317 |
| SAOUHSC_01997 | NC_007795.1 | mut_A | wt_R | OK | 3998.03 | 2073.04 |
| SAOUHSC_01650 | NC_007795.1 | mut_A | wt_R | OK | 421.214 | 218.077 |
| SAOUHSC_02761 | NC_007795.1 | mut_A | wt_R | OK | 49.0531 | 25.3778 |
| SAOUHSC_00318 | NC_007795.1 | mut_A | wt_R | OK | 53.5646 | 27.6739 |
| SAOUHSC_01124 | NC_007795.1 | mut_A | wt_R | OK | 14.5978 | 7.53547 |
| SAOUHSC_01142 | NC_007795.1 | mut_A | wt_R | OK | 6753.64 | 3484.32 |
| SAOUHSC_00182 | NC_007795.1 | mut_A | wt_R | OK | 124.849 | 64.4074 |
| SAOUHSC_01137 | NC_007795.1 | mut_A | wt_R | OK | 280.745 | 144.798 |
| SAOUHSC_01805 | NC_007795.1 | mut_A | wt_R | OK | 1.66117 | 0.856748 |
| SAOUHSC_02942 | NC_007795.1 | mut_A | wt_R | OK | 250.813 | 128.868 |
| SAOUHSC_00710 | NC_007795.1 | mut_A | wt_R | OK | 282.085 | 144.803 |
| SAOUHSC_00673 | NC_007795.1 | mut_A | wt_R | OK | 18.5707 | 9.5314 |
| SAOUHSC_01085 | NC_007795.1 | mut_A | wt_R | OK | 27.0902 | 13.8891 |
| SAOUHSC_02465 | NC_007795.1 | mut_A | wt_R | OK | 242.88 | 124.439 |
| SAOUHSC_02844 | NC_007795.1 | mut_A | wt_R | OK | 870.125 | 445.688 |
| SAOUHSC_01502 | NC_007795.1 | mut_A | wt_R | OK | 122.024 | 62.4893 |
| SAOUHSC_00202 | NC_007795.1 | mut_A | wt_R | OK | 145.541 | 74.5033 |
| SAOUHSC_02244 | NC_007795.1 | mut_A | wt_R | OK | 72.7905 | 37.1005 |
| SAOUHSC_00940 | NC_007795.1 | mut_A | wt_R | OK | 354.503 | 180.304 |
| SAOUHSC_00409 | NC_007795.1 | mut_A | wt_R | OK | 325.836 | 164.959 |
| SAOUHSC_03022 | NC_007795.1 | mut_A | wt_R | OK | 473.979 | 239.87 |
| SAOUHSC_02422 | NC_007795.1 | mut_A | wt_R | OK | 253.65 | 127.629 |
| SAOUHSC_01761a | NC_007795.1 | mut_A | wt_R | OK | 50.2203 | 25.2604 |
| SAOUHSC_02760 | NC_007795.1 | mut_A | wt_R | OK | 159.282 | 79.9994 |
| SAOUHSC_02419 | NC_007795.1 | mut_A | wt_R | OK | 249.618 | 125.243 |

| | | | | | | |
|---|---|---|---|---|---|---|
| SAOUHSC_02403 | NC_007795.1 | mut_A | wt_R | OK | 176.73 | 88.5725 |
| SAOUHSC_02832 | NC_007795.1 | mut_A | wt_R | OK | 6.82155 | 3.4137 |
| SAOUHSC_01377 | NC_007795.1 | mut_A | wt_R | OK | 156.83 | 78.2684 |
| SAOUHSC_00973 | NC_007795.1 | mut_A | wt_R | OK | 320.405 | 159.722 |
| SAOUHSC_00998 | NC_007795.1 | mut_A | wt_R | OK | 350.529 | 174.25 |
| SAOUHSC_01419 | NC_007795.1 | mut_A | wt_R | OK | 541.949 | 268.38 |
| SAOUHSC_02663 | NC_007795.1 | mut_A | wt_R | OK | 100.886 | 49.935 |
| SAOUHSC_00327 | NC_007795.1 | mut_A | wt_R | OK | 33.0528 | 16.3341 |
| SAOUHSC_02852 | NC_007795.1 | mut_A | wt_R | OK | 81.4465 | 40.2109 |
| SAOUHSC_01071 | NC_007795.1 | mut_A | wt_R | OK | 158.472 | 78.1939 |
| SAOUHSC_00591 | NC_007795.1 | mut_A | wt_R | OK | 32.2366 | 15.895 |
| SAOUHSC_02718 | NC_007795.1 | mut_A | wt_R | OK | 13.6716 | 6.73355 |
| adhA | NC_007795.1 | mut_A | wt_R | OK | 7984.49 | 3932.26 |
| SAOUHSC_00433 | NC_007795.1 | mut_A | wt_R | OK | 62.5871 | 30.8078 |
| SAOUHSC_02381 | NC_007795.1 | mut_A | wt_R | OK | 2615.34 | 1287.28 |
| SAOUHSC_01086 | NC_007795.1 | mut_A | wt_R | OK | 16.4056 | 8.07248 |
| SAOUHSC_00360 | NC_007795.1 | mut_A | wt_R | OK | 108.615 | 53.3785 |
| SAOUHSC_00617 | NC_007795.1 | mut_A | wt_R | OK | 10620.2 | 5213.46 |
| SAOUHSC_02097 | NC_007795.1 | mut_A | wt_R | OK | 95.0237 | 46.6193 |
| SAOUHSC_00826 | NC_007795.1 | mut_A | wt_R | OK | 141.544 | 69.4262 |
| SAOUHSC_01213 | NC_007795.1 | mut_A | wt_R | OK | 62.7769 | 30.7652 |
| SAOUHSC_01087 | NC_007795.1 | mut_A | wt_R | OK | 23.047 | 11.2517 |
| SAOUHSC_02776 | NC_007795.1 | mut_A | wt_R | OK | 46.786 | 22.8141 |
| SAOUHSC_02319 | NC_007795.1 | mut_A | wt_R | OK | 138.441 | 67.3258 |
| SAOUHSC_01127 | NC_007795.1 | mut_A | wt_R | OK | 9.06051 | 4.39188 |
| SAOUHSC_00256 | NC_007795.1 | mut_A | wt_R | OK | 133.125 | 64.1592 |
| SAOUHSC_00306 | NC_007795.1 | mut_A | wt_R | OK | 91.3297 | 44.0145 |
| SAOUHSC_01924 | NC_007795.1 | mut_A | wt_R | OK | 43.1161 | 20.5686 |
| SAOUHSC_02108 | NC_007795.1 | mut_A | wt_R | OK | 6153.27 | 2931.73 |
| SAOUHSC_01125 | NC_007795.1 | mut_A | wt_R | OK | 7.94434 | 3.75008 |
| SAOUHSC_00975 | NC_007795.1 | mut_A | wt_R | OK | 163.391 | 77.0966 |
| SAOUHSC_00686 | NC_007795.1 | mut_A | wt_R | OK | 152.431 | 71.9068 |
| SAOUHSC_00320 | NC_007795.1 | mut_A | wt_R | OK | 95.2683 | 44.9011 |
| SAOUHSC_00319 | NC_007795.1 | mut_A | wt_R | OK | 100.032 | 47.1426 |
| SAOUHSC_00831 | NC_007795.1 | mut_A | wt_R | OK | 68.4067 | 32.2347 |
| SAOUHSC_00312 | NC_007795.1 | mut_A | wt_R | OK | 159.658 | 75.1381 |
| SAOUHSC_00655 | NC_007795.1 | mut_A | wt_R | OK | 513.119 | 240.723 |
| SAOUHSC_00465 | NC_007795.1 | mut_A | wt_R | OK | 1603.01 | 750.274 |
| SAOUHSC_00712 | NC_007795.1 | mut_A | wt_R | OK | 310.753 | 145.142 |
| SAOUHSC_02994 | NC_007795.1 | mut_A | wt_R | OK | 166.495 | 77.6148 |
| SAOUHSC_01301 | NC_007795.1 | mut_A | wt_R | OK | 534.028 | 248.128 |
| SAOUHSC_02759 | NC_007795.1 | mut_A | wt_R | OK | 4021.01 | 1868.07 |
| SAOUHSC_00295 | NC_007795.1 | mut_A | wt_R | OK | 25.8372 | 12.0032 |
| SAOUHSC_00972 | NC_007795.1 | mut_A | wt_R | OK | 157.263 | 72.9097 |
| SAOUHSC_00938 | NC_007795.1 | mut_A | wt_R | OK | 674.711 | 311.253 |
| SAOUHSC_02646 | NC_007795.1 | mut_A | wt_R | OK | 104.561 | 48.0677 |
| SAOUHSC_02566 | NC_007795.1 | mut_A | wt_R | OK | 2903.01 | 1330.16 |
| SAOUHSC_02599 | NC_007795.1 | mut_A | wt_R | OK | 22.9331 | 10.5028 |
| SAOUHSC_02753 | NC_007795.1 | mut_A | wt_R | OK | 34.1413 | 15.6175 |
| SAOUHSC_00311 | NC_007795.1 | mut_A | wt_R | OK | 165.647 | 75.6399 |
| SAOUHSC_01278 | NC_007795.1 | mut_A | wt_R | OK | 3314.07 | 1505.36 |
| SAOUHSC_02609 | NC_007795.1 | mut_A | wt_R | OK | 7.86152 | 3.56784 |
| SAOUHSC_03034 | NC_007795.1 | mut_A | wt_R | OK | 256.736 | 116.292 |
| SAOUHSC_00121 | NC_007795.1 | mut_A | wt_R | OK | 52.3567 | 23.6951 |
| SAOUHSC_01922 | NC_007795.1 | mut_A | wt_R | OK | 96.649 | 43.6828 |

| | | | | | |
|---|---|---|---|---|---|
| SAOUHSC_02262 | NC_007795.1 mut_A | wt_R | OK | 13041.5 | 5889.39 |
| SAOUHSC_00122 | NC_007795.1 mut_A | wt_R | OK | 41.3879 | 18.651 |
| SAOUHSC_02872 | NC_007795.1 mut_A | wt_R | OK | 914.525 | 410.939 |
| SAOUHSC_02557 | NC_007795.1 mut_A | wt_R | OK | 48.4266 | 21.75 |
| SAOUHSC_00806 | NC_007795.1 mut_A | wt_R | OK | 547.825 | 245.925 |
| SAOUHSC_02811 | NC_007795.1 mut_A | wt_R | OK | 108.376 | 48.6334 |
| SAOUHSC_01596 | NC_007795.1 mut_A | wt_R | OK | 61.0562 | 27.3655 |
| SAOUHSC_02246 | NC_007795.1 mut_A | wt_R | OK | 5.68685 | 2.54544 |
| SAOUHSC_00593 | NC_007795.1 mut_A | wt_R | OK | 26.6036 | 11.9009 |
| SAOUHSC_02264 | NC_007795.1 mut_A | wt_R | OK | 5157.81 | 2300.9 |
| SAOUHSC_00294 | NC_007795.1 mut_A | wt_R | OK | 23.093 | 10.2972 |
| SAOUHSC_00377 | NC_007795.1 mut_A | wt_R | OK | 6.18431 | 2.75491 |
| SAOUHSC_R0005 | NC_007795.1 mut_A | wt_R | OK | 1.69671 | 0.755567 |
| SAOUHSC_02604 | NC_007795.1 mut_A | wt_R | OK | 110.922 | 49.3462 |
| SAOUHSC_02245 | NC_007795.1 mut_A | wt_R | OK | 21.8643 | 9.68438 |
| SAOUHSC_02103 | NC_007795.1 mut_A | wt_R | OK | 22.5844 | 9.99092 |
| SAOUHSC_02810 | NC_007795.1 mut_A | wt_R | OK | 326.686 | 143.889 |
| SAOUHSC_00124 | NC_007795.1 mut_A | wt_R | OK | 17.7542 | 7.81313 |
| SAOUHSC_00237 | NC_007795.1 mut_A | wt_R | OK | 48.7174 | 21.3152 |
| SAOUHSC_00186 | NC_007795.1 mut_A | wt_R | OK | 52.9736 | 23.1368 |
| SAOUHSC_01443 | NC_007795.1 mut_A | wt_R | OK | 9.03825 | 3.93639 |
| SAOUHSC_00864 | NC_007795.1 mut_A | wt_R | OK | 272.669 | 117.804 |
| SAOUHSC_00120 | NC_007795.1 mut_A | wt_R | OK | 53.5832 | 23.1202 |
| SAOUHSC_01912 | NC_007795.1 mut_A | wt_R | OK | 50.008 | 21.5755 |
| SAOUHSC_01221 | NC_007795.1 mut_A | wt_R | OK | 10.2443 | 4.41763 |
| SAOUHSC_01691 | NC_007795.1 mut_A | wt_R | OK | 15.7088 | 6.77086 |
| SAOUHSC_00326 | NC_007795.1 mut_A | wt_R | OK | 46.8995 | 20.2035 |
| SAOUHSC_00759 | NC_007795.1 mut_A | wt_R | OK | 95.5555 | 41.1394 |
| SAOUHSC_01303 | NC_007795.1 mut_A | wt_R | OK | 38.4013 | 16.4553 |
| SAOUHSC_00858 | NC_007795.1 mut_A | wt_R | OK | 96.7659 | 41.4165 |
| SAOUHSC_00808 | NC_007795.1 mut_A | wt_R | OK | 425.309 | 181.92 |
| SAOUHSC_01615 | NC_007795.1 mut_A | wt_R | OK | 431.617 | 183.928 |
| SAOUHSC_01838 | NC_007795.1 mut_A | wt_R | OK | 461.618 | 194.336 |
| SAOUHSC_00807 | NC_007795.1 mut_A | wt_R | OK | 189.222 | 79.6152 |
| SAOUHSC_01890 | NC_007795.1 mut_A | wt_R | OK | 12.3276 | 5.185 |
| SAOUHSC_00313 | NC_007795.1 mut_A | wt_R | OK | 97.5483 | 40.994 |
| SAOUHSC_01181 | NC_007795.1 mut_A | wt_R | OK | 1233.89 | 517.422 |
| SAOUHSC_00169 | NC_007795.1 mut_A | wt_R | OK | 18.3963 | 7.63483 |
| SAOUHSC_00065 | NC_007795.1 mut_A | wt_R | OK | 77.9845 | 32.1927 |
| SAOUHSC_01166 | NC_007795.1 mut_A | wt_R | OK | 308.815 | 127.429 |
| SAOUHSC_00656 | NC_007795.1 mut_A | wt_R | OK | 359.913 | 148.364 |
| SAOUHSC_00860 | NC_007795.1 mut_A | wt_R | OK | 234.771 | 96.7587 |
| SAOUHSC_00592 | NC_007795.1 mut_A | wt_R | OK | 37.7384 | 15.53 |
| SAOUHSC_00634 | NC_007795.1 mut_A | wt_R | OK | 2057.57 | 840.169 |
| SAOUHSC_00043 | NC_007795.1 mut_A | wt_R | OK | 16.1431 | 6.57021 |
| SAOUHSC_00596 | NC_007795.1 mut_A | wt_R | OK | 5.64938 | 2.28841 |
| SAOUHSC_00717 | NC_007795.1 mut_A | wt_R | OK | 913.813 | 368.926 |
| SAOUHSC_00559 | NC_007795.1 mut_A | wt_R | OK | 43.1828 | 17.4275 |
| SAOUHSC_01595 | NC_007795.1 mut_A | wt_R | OK | 108.515 | 43.7895 |
| SAOUHSC_01327 | NC_007795.1 mut_A | wt_R | OK | 1353.6 | 544.973 |
| SAOUHSC_01379 | NC_007795.1 mut_A | wt_R | OK | 127.704 | 51.3783 |
| SAOUHSC_00304 | NC_007795.1 mut_A | wt_R | OK | 48.079 | 19.3398 |
| SAOUHSC_02155 | NC_007795.1 mut_A | wt_R | OK | 204.826 | 81.6408 |
| SAOUHSC_03046 | NC_007795.1 mut_A | wt_R | OK | 109.374 | 43.5858 |
| SAOUHSC_01904 | NC_007795.1 mut_A | wt_R | OK | 45.9792 | 18.1916 |

| | | | | | | |
|---|---|---|---|---|---|---|
| SAOUHSC_01641 | NC_007795.1 | mut_A | wt_R | OK | 4.01017 | 1.58556 |
| SAOUHSC_01849 | NC_007795.1 | mut_A | wt_R | OK | 135.193 | 53.3767 |
| SAOUHSC_00185 | NC_007795.1 | mut_A | wt_R | OK | 62.6209 | 24.72 |
| SAOUHSC_00879 | NC_007795.1 | mut_A | wt_R | OK | 295.615 | 116.41 |
| SAOUHSC_02793 | NC_007795.1 | mut_A | wt_R | OK | 83.8589 | 32.9859 |
| SAOUHSC_02607 | NC_007795.1 | mut_A | wt_R | OK | 90.0776 | 35.4286 |
| SAOUHSC_02608 | NC_007795.1 | mut_A | wt_R | OK | 48.8829 | 19.1926 |
| SAOUHSC_02606 | NC_007795.1 | mut_A | wt_R | OK | 124.171 | 48.5198 |
| SAOUHSC_01164 | NC_007795.1 | mut_A | wt_R | OK | 314.776 | 122.874 |
| SAOUHSC_00363 | NC_007795.1 | mut_A | wt_R | OK | 76.8815 | 29.7569 |
| SAOUHSC_00074 | NC_007795.1 | mut_A | wt_R | OK | 52.513 | 20.3179 |
| SAOUHSC_02974 | NC_007795.1 | mut_A | wt_R | OK | 61.8764 | 23.8758 |
| SAOUHSC_01378 | NC_007795.1 | mut_A | wt_R | OK | 125.465 | 48.344 |
| SAOUHSC_01219 | NC_007795.1 | mut_A | wt_R | OK | 43.2946 | 16.6114 |
| SAOUHSC_01255 | NC_007795.1 | mut_A | wt_R | OK | 83.3386 | 31.9751 |
| SAOUHSC_00625 | NC_007795.1 | mut_A | wt_R | OK | 39.7652 | 15.1666 |
| SAOUHSC_01601 | NC_007795.1 | mut_A | wt_R | OK | 509.012 | 193.654 |
| SAOUHSC_00859 | NC_007795.1 | mut_A | wt_R | OK | 165.324 | 62.7054 |
| SAOUHSC_00241 | NC_007795.1 | mut_A | wt_R | OK | 310.107 | 116.926 |
| SAOUHSC_02645 | NC_007795.1 | mut_A | wt_R | OK | 80.6859 | 30.2699 |
| SAOUHSC_00371 | NC_007795.1 | mut_A | wt_R | OK | 1259.19 | 471.055 |
| SAOUHSC_02256 | NC_007795.1 | mut_A | wt_R | OK | 447.732 | 167.154 |
| SAOUHSC_00123 | NC_007795.1 | mut_A | wt_R | OK | 18.8518 | 7.00894 |
| SAOUHSC_01089 | NC_007795.1 | mut_A | wt_R | OK | 20.7374 | 7.68765 |
| SAOUHSC_00325 | NC_007795.1 | mut_A | wt_R | OK | 47.9558 | 17.7688 |
| SAOUHSC_00119 | NC_007795.1 | mut_A | wt_R | OK | 65.7044 | 24.3179 |
| SAOUHSC_02744 | NC_007795.1 | mut_A | wt_R | OK | 257.426 | 95.1491 |
| SAOUHSC_01821 | NC_007795.1 | mut_A | wt_R | OK | 181.825 | 67.164 |
| SAOUHSC_01945 | NC_007795.1 | mut_A | wt_R | OK | 43.0533 | 15.8967 |
| SAOUHSC_02587 | NC_007795.1 | mut_A | wt_R | OK | 89.2256 | 32.9127 |
| SAOUHSC_01380 | NC_007795.1 | mut_A | wt_R | OK | 101.863 | 37.4545 |
| SAOUHSC_02160 | NC_007795.1 | mut_A | wt_R | OK | 112.184 | 41.1503 |
| SAOUHSC_02771 | NC_007795.1 | mut_A | wt_R | OK | 19.2869 | 7.06981 |
| SAOUHSC_00201 | NC_007795.1 | mut_A | wt_R | OK | 189.276 | 69.2027 |
| SAOUHSC_03028 | NC_007795.1 | mut_A | wt_R | OK | 36.6991 | 13.3974 |
| SAOUHSC_00142 | NC_007795.1 | mut_A | wt_R | OK | 81.781 | 29.6092 |
| SAOUHSC_00586 | NC_007795.1 | mut_A | wt_R | OK | 44.0575 | 15.9168 |
| SAOUHSC_00775 | NC_007795.1 | mut_A | wt_R | OK | 202.598 | 73.0233 |
| SAOUHSC_02794 | NC_007795.1 | mut_A | wt_R | OK | 95.0469 | 34.2038 |
| SAOUHSC_02877 | NC_007795.1 | mut_A | wt_R | OK | 54.0366 | 19.3139 |
| SAOUHSC_00658 | NC_007795.1 | mut_A | wt_R | OK | 835.765 | 298.628 |
| SAOUHSC_02741 | NC_007795.1 | mut_A | wt_R | OK | 380.787 | 135.985 |
| SAOUHSC_03035 | NC_007795.1 | mut_A | wt_R | OK | 94.5716 | 33.6803 |
| SAOUHSC_01165 | NC_007795.1 | mut_A | wt_R | OK | 184.164 | 65.5239 |
| SAOUHSC_00588 | NC_007795.1 | mut_A | wt_R | OK | 28.5917 | 10.142 |
| SAOUHSC_01921 | NC_007795.1 | mut_A | wt_R | OK | 64.8022 | 22.9793 |
| SAOUHSC_02763 | NC_007795.1 | mut_A | wt_R | OK | 41.3378 | 14.5978 |
| SAOUHSC_02265 | NC_007795.1 | mut_A | wt_R | OK | 8380.57 | 2901.37 |
| SAOUHSC_01054 | NC_007795.1 | mut_A | wt_R | OK | 455.747 | 157.703 |
| SAOUHSC_00085 | NC_007795.1 | mut_A | wt_R | OK | 59.4873 | 20.5458 |
| SAOUHSC_00118 | NC_007795.1 | mut_A | wt_R | OK | 44.5584 | 15.3751 |
| SAOUHSC_00647 | NC_007795.1 | mut_A | wt_R | OK | 2396.66 | 822.244 |
| SAOUHSC_01923 | NC_007795.1 | mut_A | wt_R | OK | 33.6677 | 11.5182 |
| SAOUHSC_02152 | NC_007795.1 | mut_A | wt_R | OK | 284.404 | 97.0149 |
| SAOUHSC_00240 | NC_007795.1 | mut_A | wt_R | OK | 471.561 | 160.622 |

| | | | | | | |
|---|---|---|---|---|---|---|
| SAOUHSC_02719 | NC_007795.1 | mut_A | wt_R | OK | 9.90979 | 3.34984 |
| SAOUHSC_01112 | NC_007795.1 | mut_A | wt_R | OK | 195.72 | 66.1195 |
| SAOUHSC_00687 | NC_007795.1 | mut_A | wt_R | OK | 31.5383 | 10.5457 |
| SAOUHSC_00310 | NC_007795.1 | mut_A | wt_R | OK | 231.248 | 76.9547 |
| SAOUHSC_02743 | NC_007795.1 | mut_A | wt_R | OK | 275.237 | 91.2561 |
| SAOUHSC_02134 | NC_007795.1 | mut_A | wt_R | OK | 572.75 | 189.893 |
| SAOUHSC_01920 | NC_007795.1 | mut_A | wt_R | OK | 36.0435 | 11.9403 |
| SAOUHSC_01088 | NC_007795.1 | mut_A | wt_R | OK | 9.59275 | 3.1561 |
| SAOUHSC_02154 | NC_007795.1 | mut_A | wt_R | OK | 187.272 | 61.6124 |
| SAOUHSC_02751 | NC_007795.1 | mut_A | wt_R | OK | 656.262 | 215.303 |
| SAOUHSC_00176 | NC_007795.1 | mut_A | wt_R | OK | 126.547 | 41.3088 |
| SAOUHSC_03006 | NC_007795.1 | mut_A | wt_R | OK | 388.644 | 126.48 |
| SAOUHSC_02161 | NC_007795.1 | mut_A | wt_R | OK | 96.9731 | 31.4053 |
| SAOUHSC_02742 | NC_007795.1 | mut_A | wt_R | OK | 321.686 | 103.54 |
| SAOUHSC_02765 | NC_007795.1 | mut_A | wt_R | OK | 19.4322 | 6.212 |
| SAOUHSC_01144 | NC_007795.1 | mut_A | wt_R | OK | 1015.65 | 323.841 |
| SAOUHSC_02778 | NC_007795.1 | mut_A | wt_R | OK | 270.004 | 84.9334 |
| SAOUHSC_02708 | NC_007795.1 | mut_A | wt_R | OK | 38.405 | 11.6909 |
| SAOUHSC_01947 | NC_007795.1 | mut_A | wt_R | OK | 57.4189 | 17.2622 |
| SAOUHSC_01130 | NC_007795.1 | mut_A | wt_R | OK | 162.038 | 48.6688 |
| SAOUHSC_02822 | NC_007795.1 | mut_A | wt_R | OK | 188.679 | 56.658 |
| SAOUHSC_01129 | NC_007795.1 | mut_A | wt_R | OK | 441.569 | 132.509 |
| SAOUHSC_02565 | NC_007795.1 | mut_A | wt_R | OK | 310.997 | 93.1889 |
| SAOUHSC_01730 | NC_007795.1 | mut_A | wt_R | OK | 48.4644 | 14.2989 |
| SAOUHSC_02764 | NC_007795.1 | mut_A | wt_R | OK | 16.7358 | 4.90859 |
| SAOUHSC_02129 | NC_007795.1 | mut_A | wt_R | OK | 117.197 | 34.3422 |
| SAOUHSC_00180 | NC_007795.1 | mut_A | wt_R | OK | 97.2152 | 28.4286 |
| SAOUHSC_01953 | NC_007795.1 | mut_A | wt_R | OK | 896.145 | 262.058 |
| SAOUHSC_01763 | NC_007795.1 | mut_A | wt_R | OK | 18.0296 | 5.26878 |
| SAOUHSC_00145 | NC_007795.1 | mut_A | wt_R | OK | 445.927 | 130.261 |
| SAOUHSC_00181 | NC_007795.1 | mut_A | wt_R | OK | 117.877 | 34.2728 |
| SAOUHSC_02812 | NC_007795.1 | mut_A | wt_R | OK | 138.487 | 39.8362 |
| SAOUHSC_00716 | NC_007795.1 | mut_A | wt_R | OK | 842.16 | 241.589 |
| SAOUHSC_02564 | NC_007795.1 | mut_A | wt_R | OK | 278.427 | 79.1578 |
| SAOUHSC_02436 | NC_007795.1 | mut_A | wt_R | OK | 203.122 | 57.448 |
| SAOUHSC_02754 | NC_007795.1 | mut_A | wt_R | OK | 23.6721 | 6.69302 |
| SAOUHSC_00177 | NC_007795.1 | mut_A | wt_R | OK | 186.114 | 52.5924 |
| SAOUHSC_01318 | NC_007795.1 | mut_A | wt_R | OK | 160.068 | 45.0865 |
| SAOUHSC_01729 | NC_007795.1 | mut_A | wt_R | OK | 837.736 | 235.939 |
| SAOUHSC_02151 | NC_007795.1 | mut_A | wt_R | OK | 157.273 | 44.1139 |
| SAOUHSC_03023 | NC_007795.1 | mut_A | wt_R | OK | 188.55 | 52.5708 |
| SAOUHSC_01643 | NC_007795.1 | mut_A | wt_R | OK | 21.7432 | 6.00014 |
| SAOUHSC_00239 | NC_007795.1 | mut_A | wt_R | OK | 407.912 | 112.305 |
| SAOUHSC_02135 | NC_007795.1 | mut_A | wt_R | OK | 374.296 | 100.02 |
| SAOUHSC_02153 | NC_007795.1 | mut_A | wt_R | OK | 88.7519 | 23.7104 |
| SAOUHSC_02777 | NC_007795.1 | mut_A | wt_R | OK | 217.411 | 57.9126 |
| SAOUHSC_02806 | NC_007795.1 | mut_A | wt_R | OK | 837.862 | 223.044 |
| SAOUHSC_00636 | NC_007795.1 | mut_A | wt_R | OK | 1373.52 | 365.066 |
| SAOUHSC_00585 | NC_007795.1 | mut_A | wt_R | OK | 8.76834 | 2.32928 |
| SAOUHSC_02813 | NC_007795.1 | mut_A | wt_R | OK | 102.246 | 27.083 |
| SAOUHSC_02774 | NC_007795.1 | mut_A | wt_R | OK | 105.902 | 27.8012 |
| SAOUHSC_01948 | NC_007795.1 | mut_A | wt_R | OK | 106.411 | 27.8535 |
| SAOUHSC_01949 | NC_007795.1 | mut_A | wt_R | OK | 100.229 | 26.1285 |
| SAOUHSC_01297 | NC_007795.1 | mut_A | wt_R | OK | 20.3213 | 5.28775 |
| SAOUHSC_00179 | NC_007795.1 | mut_A | wt_R | OK | 122.306 | 31.4392 |

| | | | | | | |
|---|---|---|---|---|---|---|
| SAOUHSC_02688 | NC_007795.1 | mut_A | wt_R | OK | 205.351 | 52.1151 |
| SAOUHSC_02865 | NC_007795.1 | mut_A | wt_R | OK | 251.291 | 63.6213 |
| SAOUHSC_00544 | NC_007795.1 | mut_A | wt_R | OK | 2896.54 | 728.422 |
| SAOUHSC_00637 | NC_007795.1 | mut_A | wt_R | OK | 1506.62 | 378.337 |
| SAOUHSC_01081 | NC_007795.1 | mut_A | wt_R | OK | 117.118 | 29.2876 |
| SAOUHSC_00114 | NC_007795.1 | mut_A | wt_R | OK | 53.9104 | 13.4363 |
| SAOUHSC_01764 | NC_007795.1 | mut_A | wt_R | OK | 9.51841 | 2.36934 |
| SAOUHSC_00117 | NC_007795.1 | mut_A | wt_R | OK | 52.2189 | 12.9246 |
| SAOUHSC_01383 | NC_007795.1 | mut_A | wt_R | OK | 239.922 | 58.8581 |
| SAOUHSC_00410 | NC_007795.1 | mut_A | wt_R | OK | 140.074 | 34.0462 |
| SAOUHSC_00827 | NC_007795.1 | mut_A | wt_R | OK | 468.842 | 113.265 |
| SAOUHSC_01307 | NC_007795.1 | mut_A | wt_R | OK | 92.4814 | 22.2754 |
| SAOUHSC_02387 | NC_007795.1 | mut_A | wt_R | OK | 70.401 | 16.8014 |
| SAOUHSC_02864 | NC_007795.1 | mut_A | wt_R | OK | 126.567 | 30.0282 |
| SAOUHSC_00175 | NC_007795.1 | mut_A | wt_R | OK | 30.4026 | 7.07907 |
| SAOUHSC_02519 | NC_007795.1 | mut_A | wt_R | OK | 377.88 | 87.7048 |
| SAOUHSC_02314 | NC_007795.1 | mut_A | wt_R | OK | 77.6756 | 17.9153 |
| SAOUHSC_00356 | NC_007795.1 | mut_A | wt_R | OK | 374.343 | 85.9568 |
| SAOUHSC_01913 | NC_007795.1 | mut_A | wt_R | OK | 36.8241 | 8.44903 |
| SAOUHSC_00115 | NC_007795.1 | mut_A | wt_R | OK | 89.0232 | 20.2152 |
| SAOUHSC_00144 | NC_007795.1 | mut_A | wt_R | OK | 530.855 | 120.336 |
| SAOUHSC_02879 | NC_007795.1 | mut_A | wt_R | OK | 40.7822 | 9.18246 |
| SAOUHSC_02666 | NC_007795.1 | mut_A | wt_R | OK | 621.258 | 138.969 |
| SAOUHSC_00178 | NC_007795.1 | mut_A | wt_R | OK | 168.278 | 37.502 |
| SAOUHSC_02563 | NC_007795.1 | mut_A | wt_R | OK | 319.133 | 70.5952 |
| SAOUHSC_02863 | NC_007795.1 | mut_A | wt_R | OK | 473.95 | 102.726 |
| SAOUHSC_02808 | NC_007795.1 | mut_A | wt_R | OK | 320.433 | 69.1463 |
| SAOUHSC_00378 | NC_007795.1 | mut_A | wt_R | OK | 7.02085 | 1.50354 |
| SAOUHSC_01602 | NC_007795.1 | mut_A | wt_R | OK | 285.02 | 60.5031 |
| SAOUHSC_00116 | NC_007795.1 | mut_A | wt_R | OK | 85.2354 | 18.0643 |
| SAOUHSC_02976 | NC_007795.1 | mut_A | wt_R | OK | 2008.49 | 423.23 |
| SAOUHSC_02839 | NC_007795.1 | mut_A | wt_R | OK | 823.667 | 172.759 |
| SAOUHSC_00358 | NC_007795.1 | mut_A | wt_R | OK | 704.656 | 146.669 |
| SAOUHSC_00284 | NC_007795.1 | mut_A | wt_R | OK | 592.66 | 122.475 |
| SAOUHSC_02975 | NC_007795.1 | mut_A | wt_R | OK | 1776.6 | 365.745 |
| SAOUHSC_01128 | NC_007795.1 | mut_A | wt_R | OK | 338.884 | 69.4889 |
| SAOUHSC_00300 | NC_007795.1 | mut_A | wt_R | OK | 845.431 | 169.643 |
| SAOUHSC_02882 | NC_007795.1 | mut_A | wt_R | OK | 14.2038 | 2.84258 |
| SAOUHSC_00211 | NC_007795.1 | mut_A | wt_R | OK | 96.0708 | 19.062 |
| SAOUHSC_02880 | NC_007795.1 | mut_A | wt_R | OK | 35.3579 | 6.93459 |
| SAOUHSC_00624 | NC_007795.1 | mut_A | wt_R | OK | 25.6712 | 5.00664 |
| SAOUHSC_02562 | NC_007795.1 | mut_A | wt_R | OK | 398.416 | 76.1546 |
| SAOUHSC_00818 | NC_007795.1 | mut_A | wt_R | OK | 52.6275 | 10.0555 |
| ureA | NC_007795.1 | mut_A | wt_R | OK | 176.711 | 33.7378 |
| SAOUHSC_00143 | NC_007795.1 | mut_A | wt_R | OK | 898.957 | 171.459 |
| murQ | NC_007795.1 | mut_A | wt_R | OK | 185.994 | 34.8683 |
| SAOUHSC_01950 | NC_007795.1 | mut_A | wt_R | OK | 51.8544 | 9.56723 |
| SAOUHSC_01456 | NC_007795.1 | mut_A | wt_R | OK | 705.476 | 128.614 |
| SAOUHSC_02315 | NC_007795.1 | mut_A | wt_R | OK | 89.0358 | 16.1219 |
| SAOUHSC_01121 | NC_007795.1 | mut_A | wt_R | OK | 10389.3 | 1855.62 |
| SAOUHSC_01951 | NC_007795.1 | mut_A | wt_R | OK | 64.8333 | 11.4722 |
| ureB | NC_007795.1 | mut_A | wt_R | OK | 375.159 | 65.1554 |
| SAOUHSC_02795 | NC_007795.1 | mut_A | wt_R | OK | 15.6119 | 2.68344 |
| SAOUHSC_00560 | NC_007795.1 | mut_A | wt_R | OK | 113.52 | 19.278 |
| SAOUHSC_02841 | NC_007795.1 | mut_A | wt_R | OK | 850.652 | 142.767 |

| | | | | | | |
|---|---|---|---|---|---|---|
| SAOUHSC_02127 | NC_007795.1 | mut_A | wt_R | OK | 116.919 | 18.9626 |
| ureC | NC_007795.1 | mut_A | wt_R | OK | 347.626 | 55.811 |
| SAOUHSC_02840 | NC_007795.1 | mut_A | wt_R | OK | 648.149 | 104.005 |
| SAOUHSC_01135 | NC_007795.1 | mut_A | wt_R | OK | 48108.7 | 7713.6 |
| SAOUHSC_02881 | NC_007795.1 | mut_A | wt_R | OK | 30.2576 | 4.78792 |
| SAOUHSC_02241 | NC_007795.1 | mut_A | wt_R | OK | 1961.34 | 302.378 |
| SAOUHSC_00158 | NC_007795.1 | mut_A | wt_R | OK | 174.534 | 26.8247 |
| SAOUHSC_01136 | NC_007795.1 | mut_A | wt_R | OK | 33635.4 | 5124.59 |
| SAOUHSC_01306 | NC_007795.1 | mut_A | wt_R | OK | 42.372 | 6.39563 |
| SAOUHSC_02441 | NC_007795.1 | mut_A | wt_R | OK | 973.21 | 143.508 |
| SAOUHSC_01952 | NC_007795.1 | mut_A | wt_R | OK | 94.6178 | 13.9116 |
| SAOUHSC_00156 | NC_007795.1 | mut_A | wt_R | OK | 331.261 | 46.2714 |
| SAOUHSC_00160 | NC_007795.1 | mut_A | wt_R | OK | 226.63 | 27.7473 |
| SAOUHSC_02243 | NC_007795.1 | mut_A | wt_R | OK | 1516.88 | 175.37 |
| SAOUHSC_01954 | NC_007795.1 | mut_A | wt_R | OK | 56.1343 | 6.30958 |
| SAOUHSC_02443 | NC_007795.1 | mut_A | wt_R | OK | 703.533 | 78.2347 |
| SAOUHSC_02442 | NC_007795.1 | mut_A | wt_R | OK | 700.348 | 75.642 |
| SAOUHSC_00845 | NC_007795.1 | mut_A | wt_R | OK | 3399.74 | 345.716 |
| SAOUHSC_02260 | NC_007795.1 | mut_A | wt_R | OK | 202532 | 17173.3 |
| SAOUHSC_00113 | NC_007795.1 | mut_A | wt_R | OK | 2039.5 | 171.308 |
| SAOUHSC_01955 | NC_007795.1 | mut_A | wt_R | OK | 40.0359 | 2.92741 |
| SAOUHSC_02710 | NC_007795.1 | mut_A | wt_R | OK | 710.44 | 49.6686 |
| SAOUHSC_02709 | NC_007795.1 | mut_A | wt_R | OK | 640.818 | 42.8504 |
| SAOUHSC_01944 | NC_007795.1 | mut_A | wt_R | OK | 105.347 | 6.80098 |
| SAOUHSC_00622 | NC_007795.1 | mut_A | wt_R | OK | 133.018 | 7.26972 |
| SAOUHSC_00988 | NC_007795.1 | mut_A | wt_R | OK | 2231.8 | 81.5208 |
| SAOUHSC_00986 | NC_007795.1 | mut_A | wt_R | OK | 2466.3 | 86.1855 |
| SAOUHSC_00051 | NC_007795.1 | mut_A | wt_R | OK | 697.944 | 24.2409 |
| SAOUHSC_00987 | NC_007795.1 | mut_A | wt_R | OK | 2210.92 | 76.6098 |
| SAOUHSC_00961 | NC_007795.1 | mut_A | wt_R | OK | 563.03 | 11.1716 |
| SAOUHSC_01939 | NC_007795.1 | mut_A | wt_R | OK | 3823.21 | 17.5841 |
| SAOUHSC_01941 | NC_007795.1 | mut_A | wt_R | OK | 2918.41 | 12.9372 |
| SAOUHSC_01942 | NC_007795.1 | mut_A | wt_R | OK | 3160.45 | 13.525 |
| SAOUHSC_01935 | NC_007795.1 | mut_A | wt_R | OK | 3836.11 | 11.4284 |
| SAOUHSC_01938 | NC_007795.1 | mut_A | wt_R | OK | 4177.95 | 11.9149 |
| SAOUHSC_01936 | NC_007795.1 | mut_A | wt_R | OK | 7015.9 | 11.4073 |

| g2(fold_chang | test_stat | p_value | q_value | significant |
|---|---|---|---|---|
| -0.303152 | -1.25214 | 0.0285 | 0.0486537 | yes |
| -0.30655 | -1.2513 | 0.0283 | 0.0483458 | yes |
| -0.308426 | -1.24017 | 0.02915 | 0.0496256 | yes |
| -0.309504 | -1.25654 | 0.02735 | 0.0468205 | yes |
| -0.313731 | -1.28296 | 0.0233 | 0.0403932 | yes |
| -0.318861 | -1.29116 | 0.02535 | 0.0437314 | yes |
| -0.323827 | -1.28022 | 0.0256 | 0.04407 | yes |
| -0.324286 | -1.34173 | 0.01975 | 0.0349532 | yes |
| -0.324304 | -1.32718 | 0.0203 | 0.035721 | yes |
| -0.32474 | -1.33255 | 0.0203 | 0.035721 | yes |
| -0.325067 | -1.3068 | 0.02335 | 0.0404514 | yes |
| -0.325378 | -1.3434 | 0.01855 | 0.0330195 | yes |
| -0.326831 | -1.34439 | 0.02095 | 0.0368121 | yes |
| -0.327268 | -1.34737 | 0.01795 | 0.0320908 | yes |
| -0.332011 | -1.35382 | 0.0172 | 0.0308847 | yes |
| -0.334945 | -1.36387 | 0.0177 | 0.03169 | yes |
| -0.33518 | -1.24923 | 0.0288 | 0.0491318 | yes |
| -0.339143 | -1.37378 | 0.01465 | 0.0265383 | yes |
| -0.339368 | -1.31395 | 0.02245 | 0.0391403 | yes |
| -0.342147 | -1.40478 | 0.0146 | 0.0264672 | yes |
| -0.344284 | -1.42601 | 0.0127 | 0.0233498 | yes |
| -0.345038 | -1.36802 | 0.0161 | 0.0289941 | yes |
| -0.345244 | -1.34244 | 0.0199 | 0.0351176 | yes |
| -0.350242 | -1.43946 | 0.0105 | 0.0196426 | yes |
| -0.351095 | -1.34425 | 0.01935 | 0.0343194 | yes |
| -0.351102 | -1.45177 | 0.0138 | 0.0251653 | yes |
| -0.351836 | -1.44409 | 0.01335 | 0.0243628 | yes |
| -0.355607 | -1.45597 | 0.01065 | 0.0199081 | yes |
| -0.358591 | -1.4668 | 0.01025 | 0.0191895 | yes |
| -0.359966 | -1.48753 | 0.0086 | 0.0162738 | yes |
| -0.360569 | -1.42937 | 0.0132 | 0.0241248 | yes |
| -0.360618 | -1.46638 | 0.00975 | 0.0183231 | yes |
| -0.365314 | -1.47917 | 0.0101 | 0.0189231 | yes |
| -0.368368 | -1.52035 | 0.00825 | 0.0156839 | yes |
| -0.370697 | -1.49835 | 0.00895 | 0.0168972 | yes |
| -0.371121 | -1.52724 | 0.0073 | 0.0139642 | yes |
| -0.37469 | -1.5288 | 0.0067 | 0.0129474 | yes |
| -0.377225 | -1.45423 | 0.01175 | 0.0217822 | yes |
| -0.377383 | -1.38479 | 0.0157 | 0.028336 | yes |
| -0.379802 | -1.56742 | 0.0063 | 0.0122128 | yes |
| -0.381274 | -1.38931 | 0.0145 | 0.0263053 | yes |
| -0.384099 | -1.57765 | 0.0062 | 0.0120379 | yes |
| -0.38486 | -1.57748 | 0.00525 | 0.0103403 | yes |
| -0.389023 | -1.60094 | 0.00525 | 0.0103403 | yes |
| -0.389361 | -1.60248 | 0.00525 | 0.0103403 | yes |
| -0.389765 | -1.35989 | 0.0189 | 0.0335697 | yes |
| -0.390429 | -1.47978 | 0.0088 | 0.0166395 | yes |
| -0.393651 | -1.51213 | 0.0072 | 0.0138159 | yes |
| -0.400075 | -1.53773 | 0.00795 | 0.0151605 | yes |
| -0.400421 | -1.65212 | 0.00465 | 0.00923245 | yes |
| -0.401433 | -1.6503 | 0.00365 | 0.00739621 | yes |
| -0.402191 | -1.66411 | 0.00395 | 0.00793873 | yes |
| -0.402602 | -1.66168 | 0.00465 | 0.00923245 | yes |
| -0.403236 | -1.66123 | 0.00375 | 0.00757389 | yes |

| | | | | |
|---|---|---|---|---|
| -0.406344 | -1.5132 | 0.008 | 0.0152322 | yes |
| -0.409289 | -1.69046 | 0.0029 | 0.00598992 | yes |
| -0.40967 | -1.69315 | 0.0036 | 0.00730091 | yes |
| -0.411431 | -1.59395 | 0.00565 | 0.0110485 | yes |
| -0.421117 | -1.73658 | 0.00305 | 0.0062525 | yes |
| -0.423797 | -1.60591 | 0.0052 | 0.0102747 | yes |
| -0.424279 | -1.75268 | 0.0022 | 0.00464948 | yes |
| -0.425238 | -1.68659 | 0.003 | 0.00617057 | yes |
| -0.426303 | -1.58455 | 0.00615 | 0.0119502 | yes |
| -0.431749 | -1.74509 | 0.003 | 0.00617057 | yes |
| -0.432138 | -1.76957 | 0.00205 | 0.00437002 | yes |
| -0.433721 | -1.79256 | 0.00155 | 0.00336243 | yes |
| -0.436458 | -1.74516 | 0.00255 | 0.00532965 | yes |
| -0.436813 | -1.80021 | 0.0012 | 0.00266186 | yes |
| -0.436842 | -1.67488 | 0.0041 | 0.0082 | yes |
| -0.43782 | -1.7403 | 0.0024 | 0.00502896 | yes |
| -0.438126 | -1.77251 | 0.0019 | 0.00407498 | yes |
| -0.438596 | -1.80446 | 0.0022 | 0.00464948 | yes |
| -0.442687 | -1.82494 | 0.00115 | 0.00256018 | yes |
| -0.442919 | -1.80854 | 0.0011 | 0.00246224 | yes |
| -0.443492 | -1.83515 | 0.00115 | 0.00256018 | yes |
| -0.443579 | -1.81328 | 0.0015 | 0.00327128 | yes |
| -0.444548 | -1.83604 | 0.00135 | 0.00297048 | yes |
| -0.444839 | -1.81548 | 0.0021 | 0.00446114 | yes |
| -0.445113 | -1.79457 | 0.00205 | 0.00437002 | yes |
| -0.446869 | -1.8371 | 0.00145 | 0.00317914 | yes |
| -0.447236 | -1.82609 | 0.0017 | 0.00366842 | yes |
| -0.44973 | -1.85072 | 0.00075 | 0.00173402 | yes |
| -0.449996 | -1.47728 | 0.01125 | 0.0209342 | yes |
| -0.451617 | -1.85037 | 0.00135 | 0.00297048 | yes |
| -0.452207 | -1.86408 | 0.0008 | 0.00183411 | yes |
| -0.456107 | -1.83563 | 0.0017 | 0.00366842 | yes |
| -0.456969 | -1.33185 | 0.0245 | 0.0423542 | yes |
| -0.458675 | -1.66494 | 0.00385 | 0.00775676 | yes |
| -0.460877 | -1.63246 | 0.00395 | 0.00793873 | yes |
| -0.46245 | -1.88777 | 0.0009 | 0.00204621 | yes |
| -0.463328 | -1.75886 | 0.0023 | 0.00483177 | yes |
| -0.464979 | -1.90687 | 0.00085 | 0.0019415 | yes |
| -0.465969 | -1.90333 | 0.0012 | 0.00266186 | yes |
| -0.468917 | -1.91064 | 0.00115 | 0.00256018 | yes |
| -0.47067 | -1.93297 | 0.0006 | 0.00141244 | yes |
| -0.471291 | -1.92873 | 0.0008 | 0.00183411 | yes |
| -0.471762 | -1.87856 | 0.00115 | 0.00256018 | yes |
| -0.473964 | -1.93903 | 0.00045 | 0.0010885 | yes |
| -0.474856 | -1.85031 | 0.00105 | 0.0023589 | yes |
| -0.475085 | -1.43838 | 0.0151 | 0.0273132 | yes |
| -0.476894 | -1.88515 | 0.0005 | 0.00119766 | yes |
| -0.476999 | -1.93054 | 0.0007 | 0.00162914 | yes |
| -0.47854 | -1.79172 | 0.0016 | 0.00346174 | yes |
| -0.479248 | -1.56719 | 0.0073 | 0.0139642 | yes |
| -0.482068 | -1.42219 | 0.01185 | 0.021918 | yes |
| -0.482367 | -1.89138 | 0.00095 | 0.0021559 | yes |
| -0.483465 | -1.97036 | 0.00055 | 0.00130598 | yes |
| -0.48427 | -1.92531 | 0.0008 | 0.00183411 | yes |
| -0.485418 | -1.99701 | 0.00045 | 0.0010885 | yes |

| | | | | |
|---|---|---|---|---|
| -0.485821 | -1.9491 | 0.00085 | 0.0019415 | yes |
| -0.488016 | -2.0039 | 0.00055 | 0.00130598 | yes |
| -0.488368 | -2.02349 | 0.00045 | 0.0010885 | yes |
| -0.490408 | -1.8198 | 0.00155 | 0.00336243 | yes |
| -0.491671 | -2.0126 | 0.0008 | 0.00183411 | yes |
| -0.493016 | -2.00452 | 0.0005 | 0.00119766 | yes |
| -0.494667 | -1.27212 | 0.027 | 0.046318 | yes |
| -0.494745 | -1.95887 | 0.00065 | 0.00151996 | yes |
| -0.502552 | -1.99636 | 0.0009 | 0.00204621 | yes |
| -0.504077 | -2.08178 | 0.0003 | 0.00074621 | yes |
| -0.504574 | -2.0462 | 0.00025 | 0.00063012 | yes |
| -0.505455 | -2.01151 | 0.00075 | 0.00173402 | yes |
| -0.505948 | -2.0901 | 0.0003 | 0.00074621 | yes |
| -0.506206 | -2.0573 | 0.0004 | 0.00097813 | yes |
| -0.50623 | -2.05369 | 0.00025 | 0.00063012 | yes |
| -0.509339 | -1.59938 | 0.00605 | 0.0117652 | yes |
| -0.50994 | -2.02595 | 0.0007 | 0.00162914 | yes |
| -0.510019 | -2.09041 | 0.0002 | 0.00051037 | yes |
| -0.512305 | -1.81401 | 0.0015 | 0.00327128 | yes |
| -0.513408 | -1.98116 | 0.0003 | 0.00074621 | yes |
| -0.513482 | -1.55735 | 0.00715 | 0.0137307 | yes |
| -0.513723 | -1.8831 | 0.0015 | 0.00327128 | yes |
| -0.516364 | -2.08231 | 0.0002 | 0.00051037 | yes |
| -0.518807 | -2.1113 | 0.00025 | 0.00063012 | yes |
| -0.519036 | -2.13735 | 5.00E-05 | 0.00013743 | yes |
| -0.519239 | -2.1374 | 0.00015 | 0.00038924 | yes |
| -0.521135 | -2.11366 | 0.00015 | 0.00038924 | yes |
| -0.522621 | -1.53012 | 0.0118 | 0.0218584 | yes |
| -0.525469 | -2.11775 | 0.00015 | 0.00038924 | yes |
| -0.528818 | -2.09497 | 0.00035 | 0.00086186 | yes |
| -0.529369 | -2.17852 | 5.00E-05 | 0.00013743 | yes |
| -0.529526 | -2.12703 | 0.00015 | 0.00038924 | yes |
| -0.53302 | -2.13816 | 0.00025 | 0.00063012 | yes |
| -0.533801 | -2.16089 | 0.0002 | 0.00051037 | yes |
| -0.535561 | -2.18718 | 0.0001 | 0.00026595 | yes |
| -0.541325 | -2.24008 | 0.0002 | 0.00051037 | yes |
| -0.546176 | -2.22058 | 0.0001 | 0.00026595 | yes |
| -0.549645 | -2.23628 | 0.00025 | 0.00063012 | yes |
| -0.550145 | -1.84655 | 0.00225 | 0.00474293 | yes |
| -0.552126 | -2.05254 | 0.00055 | 0.00130598 | yes |
| -0.559134 | -2.30646 | 0.0001 | 0.00026595 | yes |
| -0.560174 | -2.0845 | 0.00045 | 0.0010885 | yes |
| -0.562107 | -2.31264 | 5.00E-05 | 0.00013743 | yes |
| -0.563485 | -2.31404 | 0.0001 | 0.00026595 | yes |
| -0.563766 | -1.41013 | 0.0148 | 0.0267903 | yes |
| -0.564057 | -1.95464 | 0.00075 | 0.00173402 | yes |
| -0.566203 | -1.73836 | 0.00315 | 0.00644676 | yes |
| -0.566274 | -2.33211 | 5.00E-05 | 0.00013743 | yes |
| -0.566904 | -2.31374 | 5.00E-05 | 0.00013743 | yes |
| -0.567316 | -2.11888 | 0.00045 | 0.0010885 | yes |
| -0.568201 | -2.30758 | 5.00E-05 | 0.00013743 | yes |
| -0.572464 | -2.35697 | 5.00E-05 | 0.00013743 | yes |
| -0.572826 | -2.37204 | 5.00E-05 | 0.00013743 | yes |
| -0.574867 | -2.35362 | 0.0001 | 0.00026595 | yes |
| -0.578054 | -1.55397 | 0.0086 | 0.0162738 | yes |

| | | | | | |
|---|---|---|---|---|---|
| -0.579859 | -2.18341 | 0.0002 | 0.00051037 | yes | |
| -0.580343 | -2.3543 | 5.00E-05 | 0.00013743 | yes | |
| -0.583528 | -2.41452 | 5.00E-05 | 0.00013743 | yes | |
| -0.583817 | -2.32763 | 0.0001 | 0.00026595 | yes | |
| -0.584767 | -2.16935 | 5.00E-05 | 0.00013743 | yes | |
| -0.586059 | -2.40918 | 5.00E-05 | 0.00013743 | yes | |
| -0.586291 | -1.38586 | 0.01575 | 0.0284054 | yes | |
| -0.586648 | -2.26663 | 5.00E-05 | 0.00013743 | yes | |
| -0.587569 | -1.78139 | 0.0023 | 0.00483177 | yes | |
| -0.58852 | -2.42094 | 5.00E-05 | 0.00013743 | yes | |
| -0.590714 | -2.40411 | 5.00E-05 | 0.00013743 | yes | |
| -0.59129 | -2.43852 | 5.00E-05 | 0.00013743 | yes | |
| -0.592436 | -2.01986 | 0.00065 | 0.00151996 | yes | |
| -0.594401 | -2.44204 | 5.00E-05 | 0.00013743 | yes | |
| -0.594887 | -1.9848 | 0.00045 | 0.0010885 | yes | |
| -0.599915 | -1.3156 | 0.0224 | 0.0390809 | yes | |
| -0.60142 | -2.48356 | 5.00E-05 | 0.00013743 | yes | |
| -0.602744 | -2.46498 | 5.00E-05 | 0.00013743 | yes | |
| -0.604405 | -2.49226 | 5.00E-05 | 0.00013743 | yes | |
| -0.60455 | -2.48732 | 5.00E-05 | 0.00013743 | yes | |
| -0.604583 | -1.92069 | 0.00105 | 0.0023589 | yes | |
| -0.604834 | -2.46668 | 5.00E-05 | 0.00013743 | yes | |
| -0.612353 | -2.5218 | 0.0001 | 0.00026595 | yes | |
| -0.616393 | -1.65669 | 0.003 | 0.00617057 | yes | |
| -0.617399 | -1.81343 | 0.00205 | 0.00437002 | yes | |
| -0.621603 | -2.34818 | 5.00E-05 | 0.00013743 | yes | |
| -0.622965 | -2.5342 | 5.00E-05 | 0.00013743 | yes | |
| -0.623243 | -1.68749 | 0.0042 | 0.00839318 | yes | |
| -0.624172 | -2.02714 | 0.00025 | 0.00063012 | yes | |
| -0.627679 | -2.54085 | 5.00E-05 | 0.00013743 | yes | |
| -0.627717 | -2.44886 | 5.00E-05 | 0.00013743 | yes | |
| -0.633317 | -2.61609 | 5.00E-05 | 0.00013743 | yes | |
| -0.634144 | -1.82973 | 0.0017 | 0.00366842 | | yes |
| -0.636922 | -2.44638 | 0.0001 | 0.00026595 | yes | |
| -0.638312 | -2.63096 | 5.00E-05 | 0.00013743 | yes | |
| -0.639343 | -2.21842 | 0.0002 | 0.00051037 | yes | |
| -0.639779 | -2.59245 | 5.00E-05 | 0.00013743 | yes | |
| -0.639939 | -1.66146 | 0.0038 | 0.00766858 | yes | |
| -0.64036 | -1.4639 | 0.01225 | 0.0226069 | yes | |
| -0.640857 | -2.03802 | 0.0006 | 0.00141244 | yes | |
| -0.642188 | -2.22854 | 0.0001 | 0.00026595 | yes | |
| -0.64448 | -2.59127 | 5.00E-05 | 0.00013743 | yes | |
| -0.646161 | -2.44521 | 5.00E-05 | 0.00013743 | yes | |
| -0.649318 | -2.67711 | 5.00E-05 | 0.00013743 | yes | |
| -0.650865 | -2.19946 | 0.0001 | 0.00026595 | yes | |
| -0.652775 | -1.79861 | 0.00175 | 0.00376311 | yes | |
| -0.656243 | -2.3027 | 5.00E-05 | 0.00013743 | yes | |
| -0.659571 | -2.64879 | 5.00E-05 | 0.00013743 | yes | |
| -0.664285 | -2.63092 | 5.00E-05 | 0.00013743 | yes | |
| -0.666849 | -2.67571 | 5.00E-05 | 0.00013743 | yes | |
| -0.668403 | -1.6609 | 0.00405 | 0.00810659 | | yes |
| -0.669064 | -1.52404 | 0.01005 | 0.0188437 | yes | |
| -0.670026 | -2.66201 | 5.00E-05 | 0.00013743 | yes | |
| -0.672744 | -2.7692 | 5.00E-05 | 0.00013743 | yes | |
| -0.673411 | -2.75973 | 5.00E-05 | 0.00013743 | yes | |

| | | | | |
|---|---|---|---|---|
| -0.674936 | -1.58768 | 0.00675 | 0.0130338 | yes |
| -0.675138 | -2.74907 | 5.00E-05 | 0.00013743 | yes |
| -0.675654 | -1.69634 | 0.00335 | 0.006839 | yes |
| -0.679776 | -2.80944 | 5.00E-05 | 0.00013743 | yes |
| -0.680885 | -1.49826 | 0.0117 | 0.0217059 | yes |
| -0.685092 | -2.83035 | 5.00E-05 | 0.00013743 | yes |
| -0.685226 | -2.7936 | 5.00E-05 | 0.00013743 | yes |
| -0.686038 | -2.69238 | 5.00E-05 | 0.00013743 | yes |
| -0.687714 | -2.38934 | 5.00E-05 | 0.00013743 | yes |
| -0.69025 | -1.57413 | 0.00725 | 0.013901 | yes |
| -0.702476 | -1.96038 | 0.0016 | 0.00346174 | yes |
| -0.705416 | -2.67613 | 5.00E-05 | 0.00013743 | yes |
| -0.714181 | -2.84126 | 5.00E-05 | 0.00013743 | yes |
| -0.714299 | -2.68445 | 5.00E-05 | 0.00013743 | yes |
| -0.714874 | -2.14011 | 0.00025 | 0.00063012 | yes |
| -0.71505 | -2.77206 | 5.00E-05 | 0.00013743 | yes |
| -0.717065 | -2.84291 | 5.00E-05 | 0.00013743 | yes |
| -0.72173 | -1.55652 | 0.0095 | 0.0178943 | yes |
| -0.72364 | -1.44816 | 0.0129 | 0.0236468 | yes |
| -0.724521 | -1.36718 | 0.02015 | 0.0355079 | yes |
| -0.724927 | -2.99491 | 5.00E-05 | 0.00013743 | yes |
| -0.728491 | -2.72954 | 5.00E-05 | 0.00013743 | yes |
| -0.729197 | -2.97494 | 5.00E-05 | 0.00013743 | yes |
| -0.730784 | -2.89067 | 5.00E-05 | 0.00013743 | yes |
| -0.731191 | -3.01309 | 5.00E-05 | 0.00013743 | yes |
| -0.73204 | -2.96776 | 5.00E-05 | 0.00013743 | yes |
| -0.734457 | -2.94451 | 5.00E-05 | 0.00013743 | yes |
| -0.734528 | -1.31816 | 0.0228 | 0.0396943 | yes |
| -0.73464 | -2.8821 | 5.00E-05 | 0.00013743 | yes |
| -0.736581 | -2.57035 | 0.0001 | 0.00026595 | yes |
| -0.739112 | -2.99904 | 5.00E-05 | 0.00013743 | yes |
| -0.739721 | -1.73912 | 0.00265 | 0.0055199 | yes |
| -0.740475 | -2.9748 | 5.00E-05 | 0.00013743 | yes |
| -0.743955 | -3.05948 | 5.00E-05 | 0.00013743 | yes |
| -0.744637 | -2.52059 | 5.00E-05 | 0.00013743 | yes |
| -0.751739 | -3.04792 | 5.00E-05 | 0.00013743 | yes |
| -0.754856 | -1.94183 | 0.001 | 0.00225275 | yes |
| -0.755301 | -3.01253 | 5.00E-05 | 0.00013743 | yes |
| -0.7566 | -3.02797 | 5.00E-05 | 0.00013743 | yes |
| -0.756958 | -3.07038 | 5.00E-05 | 0.00013743 | yes |
| -0.758365 | -3.08066 | 5.00E-05 | 0.00013743 | yes |
| -0.758684 | -2.95428 | 5.00E-05 | 0.00013743 | yes |
| -0.75891 | -3.09842 | 5.00E-05 | 0.00013743 | yes |
| -0.759243 | -2.98807 | 5.00E-05 | 0.00013743 | yes |
| -0.76153 | -1.8153 | 0.00195 | 0.00417493 | yes |
| -0.762085 | -3.15194 | 5.00E-05 | 0.00013743 | yes |
| -0.763904 | -2.99838 | 5.00E-05 | 0.00013743 | yes |
| -0.76425 | -3.06902 | 5.00E-05 | 0.00013743 | yes |
| -0.767222 | -2.56809 | 5.00E-05 | 0.00013743 | yes |
| -0.769624 | -1.77585 | 0.00295 | 0.00608809 | yes |
| -0.769688 | -3.17436 | 5.00E-05 | 0.00013743 | yes |
| -0.774189 | -2.80536 | 5.00E-05 | 0.00013743 | yes |
| -0.775237 | -3.14149 | 5.00E-05 | 0.00013743 | yes |
| -0.77559 | -3.01463 | 5.00E-05 | 0.00013743 | yes |
| -0.777033 | -2.24987 | 0.0001 | 0.00026595 | yes |

| | | | | |
|---|---|---|---|---|
| -0.779005 | -3.20823 | 5.00E-05 | 0.00013743 | yes |
| -0.781155 | -2.98868 | 5.00E-05 | 0.00013743 | yes |
| -0.783915 | -1.927 | 0.0005 | 0.00119766 | yes |
| -0.784553 | -2.33182 | 5.00E-05 | 0.00013743 | yes |
| -0.786665 | -2.79066 | 5.00E-05 | 0.00013743 | yes |
| -0.787901 | -3.24646 | 5.00E-05 | 0.00013743 | yes |
| -0.789034 | -3.06757 | 5.00E-05 | 0.00013743 | yes |
| -0.791496 | -2.75684 | 5.00E-05 | 0.00013743 | yes |
| -0.791772 | -3.12138 | 5.00E-05 | 0.00013743 | yes |
| -0.793748 | -3.24007 | 5.00E-05 | 0.00013743 | yes |
| -0.794504 | -3.26572 | 5.00E-05 | 0.00013743 | yes |
| -0.796555 | -2.17491 | 0.0005 | 0.00119766 | yes |
| -0.797797 | -3.16643 | 5.00E-05 | 0.00013743 | yes |
| -0.800448 | -3.30213 | 5.00E-05 | 0.00013743 | yes |
| -0.801402 | -2.87801 | 5.00E-05 | 0.00013743 | yes |
| -0.801525 | -1.70583 | 0.00395 | 0.00793873 | yes |
| -0.808575 | -2.62736 | 5.00E-05 | 0.00013743 | yes |
| -0.809176 | -3.285 | 5.00E-05 | 0.00013743 | yes |
| -0.809271 | -2.95125 | 5.00E-05 | 0.00013743 | yes |
| -0.809746 | -1.56115 | 0.01215 | 0.0224392 | yes |
| -0.810807 | -3.3392 | 5.00E-05 | 0.00013743 | yes |
| -0.815163 | -3.01618 | 5.00E-05 | 0.00013743 | yes |
| -0.821792 | -3.25445 | 5.00E-05 | 0.00013743 | yes |
| -0.823142 | -3.38064 | 5.00E-05 | 0.00013743 | yes |
| -0.826272 | -3.3994 | 5.00E-05 | 0.00013743 | yes |
| -0.827457 | -2.70737 | 5.00E-05 | 0.00013743 | yes |
| -0.829372 | -1.37537 | 0.0199 | 0.0351176 | yes |
| -0.82974 | -2.70206 | 5.00E-05 | 0.00013743 | yes |
| -0.829972 | -2.62124 | 5.00E-05 | 0.00013743 | yes |
| -0.830632 | -2.12782 | 0.00035 | 0.00086186 | yes |
| -0.831891 | -3.36979 | 5.00E-05 | 0.00013743 | yes |
| -0.832523 | -3.37167 | 5.00E-05 | 0.00013743 | yes |
| -0.83326 | -3.44172 | 5.00E-05 | 0.00013743 | yes |
| -0.833686 | -2.60713 | 5.00E-05 | 0.00013743 | yes |
| -0.83378 | -3.18208 | 5.00E-05 | 0.00013743 | yes |
| -0.834262 | -2.23367 | 0.0001 | 0.00026595 | yes |
| -0.83449 | -2.69815 | 5.00E-05 | 0.00013743 | yes |
| -0.836414 | -1.8575 | 0.0013 | 0.00286816 | yes |
| -0.841856 | -3.18613 | 5.00E-05 | 0.00013743 | yes |
| -0.842308 | -3.15553 | 5.00E-05 | 0.00013743 | yes |
| -0.849015 | -2.79294 | 5.00E-05 | 0.00013743 | yes |
| -0.850139 | -3.49872 | 5.00E-05 | 0.00013743 | yes |
| -0.854307 | -3.52904 | 5.00E-05 | 0.00013743 | yes |
| -0.85598 | -3.27495 | 5.00E-05 | 0.00013743 | yes |
| -0.856085 | -1.90117 | 0.0019 | 0.00407498 | yes |
| -0.858064 | -1.63297 | 0.0052 | 0.0102747 | yes |
| -0.860354 | -3.49552 | 5.00E-05 | 0.00013743 | yes |
| -0.863663 | -3.53974 | 5.00E-05 | 0.00013743 | yes |
| -0.864528 | -3.54459 | 5.00E-05 | 0.00013743 | yes |
| -0.871808 | -3.47933 | 5.00E-05 | 0.00013743 | yes |
| -0.873101 | -2.08804 | 0.00045 | 0.0010885 | yes |
| -0.875733 | -3.29367 | 5.00E-05 | 0.00013743 | yes |
| -0.878266 | -2.21038 | 0.00025 | 0.00063012 | yes |
| -0.879494 | -3.60647 | 5.00E-05 | 0.00013743 | yes |
| -0.881491 | -3.59297 | 5.00E-05 | 0.00013743 | yes |

| | | | | | |
|---|---|---|---|---|---|
| -0.882835 | -3.6403 | 5.00E-05 | 0.00013743 | yes | |
| -0.884977 | -3.30903 | 5.00E-05 | 0.00013743 | yes | |
| -0.885238 | -1.49973 | 0.0121 | 0.0223636 | yes | |
| -0.885698 | -3.6044 | 5.00E-05 | 0.00013743 | yes | |
| -0.889339 | -3.58258 | 5.00E-05 | 0.00013743 | yes | |
| -0.890382 | -3.67399 | 5.00E-05 | 0.00013743 | yes | |
| -0.901812 | -3.6566 | 5.00E-05 | 0.00013743 | yes | |
| -0.902614 | -3.24993 | 5.00E-05 | 0.00013743 | yes | |
| -0.903088 | -2.23632 | 0.0002 | 0.00051037 | yes | |
| -0.90533 | -2.94673 | 5.00E-05 | 0.00013743 | yes | |
| -0.909617 | -2.33006 | 0.00015 | 0.00038924 | yes | |
| -0.910851 | -3.18323 | 5.00E-05 | 0.00013743 | yes | |
| -0.911181 | -3.2364 | 5.00E-05 | 0.00013743 | yes | |
| -0.9126 | -1.66666 | 0.00595 | 0.0115983 | yes | |
| -0.913447 | -3.58581 | 5.00E-05 | 0.00013743 | yes | |
| -0.918203 | -3.75397 | 5.00E-05 | 0.00013743 | | yes |
| -0.9195 | -3.63277 | 5.00E-05 | 0.00013743 | yes | |
| -0.921561 | -2.59532 | 5.00E-05 | 0.00013743 | yes | |
| -0.922498 | -3.74225 | 5.00E-05 | 0.00013743 | yes | |
| -0.923951 | -3.50175 | 5.00E-05 | 0.00013743 | yes | |
| -0.926824 | -2.14803 | 0.0004 | 0.00097813 | yes | |
| -0.928191 | -1.77845 | 0.00215 | 0.00455164 | yes | |
| -0.929296 | -3.01933 | 5.00E-05 | 0.00013743 | yes | |
| -0.930317 | -3.75972 | 5.00E-05 | 0.00013743 | yes | |
| -0.932452 | -2.9591 | 5.00E-05 | 0.00013743 | yes | |
| -0.935652 | -3.25702 | 5.00E-05 | 0.00013743 | yes | |
| -0.937875 | -2.14912 | 0.0006 | 0.00141244 | yes | |
| -0.937997 | -2.27247 | 0.00035 | 0.00086186 | yes | |
| -0.940895 | -3.51724 | 5.00E-05 | 0.00013743 | yes | |
| -0.945501 | -3.81255 | 5.00E-05 | 0.00013743 | yes | |
| -0.947541 | -3.90884 | 5.00E-05 | 0.00013743 | yes | |
| -0.949716 | -3.82434 | 5.00E-05 | 0.00013743 | yes | |
| -0.950777 | -3.52363 | 5.00E-05 | 0.00013743 | yes | |
| -0.952753 | -3.61782 | 5.00E-05 | 0.00013743 | yes | |
| -0.953978 | -2.7891 | 5.00E-05 | 0.00013743 | yes | |
| -0.954787 | -3.95174 | 5.00E-05 | 0.00013743 | yes | |
| -0.954886 | -3.69446 | 5.00E-05 | 0.00013743 | yes | |
| -0.955212 | -3.90266 | 5.00E-05 | 0.00013743 | yes | |
| -0.955259 | -1.84609 | 0.00325 | 0.00664037 | yes | |
| -0.960726 | -3.46405 | 5.00E-05 | 0.00013743 | yes | |
| -0.962039 | -3.95353 | 5.00E-05 | 0.00013743 | yes | |
| -0.962268 | -3.67348 | 5.00E-05 | 0.00013743 | yes | |
| -0.963816 | -3.37351 | 5.00E-05 | 0.00013743 | yes | |
| -0.964805 | -3.92425 | 5.00E-05 | 0.00013743 | yes | |
| -0.965188 | -3.98506 | 5.00E-05 | 0.00013743 | yes | |
| -0.965486 | -3.39786 | 5.00E-05 | 0.00013743 | yes | |
| -0.966048 | -3.53725 | 5.00E-05 | 0.00013743 | yes | |
| -0.97231 | -3.88655 | 5.00E-05 | 0.00013743 | yes | |
| -0.975366 | -3.95863 | 5.00E-05 | 0.00013743 | yes | |
| -0.982044 | -3.57543 | 5.00E-05 | 0.00013743 | yes | |
| -0.982568 | -3.86004 | 5.00E-05 | 0.00013743 | yes | |
| -0.990882 | -4.03688 | 5.00E-05 | 0.00013743 | yes | |
| -0.991391 | -1.76153 | 0.00525 | 0.0103403 | yes | |
| -0.993524 | -4.07947 | 5.00E-05 | 0.00013743 | yes | |
| -0.994997 | -3.95621 | 5.00E-05 | 0.00013743 | yes | |

| | | | | |
|---|---|---|---|---|
| -0.996615 | -3.5134 | 5.00E-05 | 0.00013743 | yes |
| -0.998762 | -2.20099 | 0.00045 | 0.0010885 | yes |
| -1.0027 | -2.14334 | 0.0003 | 0.00074621 | yes |
| -1.00434 | -4.12587 | 5.00E-05 | 0.00013743 | yes |
| -1.00838 | -4.13425 | 5.00E-05 | 0.00013743 | yes |
| -1.01388 | -3.3407 | 5.00E-05 | 0.00013743 | yes |
| -1.0146 | -3.60681 | 5.00E-05 | 0.00013743 | yes |
| -1.01688 | -3.96566 | 5.00E-05 | 0.00013743 | yes |
| -1.01827 | -3.99694 | 5.00E-05 | 0.00013743 | yes |
| -1.0191 | -4.11584 | 5.00E-05 | 0.00013743 | yes |
| -1.02013 | -3.34151 | 5.00E-05 | 0.00013743 | yes |
| -1.02174 | -3.64866 | 5.00E-05 | 0.00013743 | yes |
| -1.02184 | -4.20523 | 5.00E-05 | 0.00013743 | yes |
| -1.02257 | -2.45899 | 5.00E-05 | 0.00013743 | yes |
| -1.02267 | -4.21744 | 5.00E-05 | 0.00013743 | yes |
| -1.02311 | -2.92591 | 5.00E-05 | 0.00013743 | yes |
| -1.02489 | -3.68063 | 5.00E-05 | 0.00013743 | yes |
| -1.0265 | -4.21376 | 5.00E-05 | 0.00013743 | yes |
| -1.02736 | -4.1559 | 5.00E-05 | 0.00013743 | yes |
| -1.0277 | -2.36918 | 0.0002 | 0.00051037 | yes |
| -1.02893 | -4.1793 | 5.00E-05 | 0.00013743 | yes |
| -1.03444 | -1.4793 | 0.01305 | 0.0238684 | yes |
| -1.03615 | -3.27509 | 5.00E-05 | 0.00013743 | yes |
| -1.04004 | -4.23401 | 5.00E-05 | 0.00013743 | yes |
| -1.04475 | -2.64338 | 5.00E-05 | 0.00013743 | yes |
| -1.05305 | -4.24719 | 5.00E-05 | 0.00013743 | yes |
| -1.05311 | -2.60838 | 5.00E-05 | 0.00013743 | yes |
| -1.06778 | -1.64096 | 0.0085 | 0.0161094 | yes |
| -1.0696 | -4.41921 | 5.00E-05 | 0.00013743 | yes |
| -1.083 | -2.64605 | 5.00E-05 | 0.00013743 | yes |
| -1.08359 | -4.02406 | 5.00E-05 | 0.00013743 | yes |
| -1.08396 | -3.779 | 5.00E-05 | 0.00013743 | yes |
| -1.08525 | -4.10297 | 5.00E-05 | 0.00013743 | yes |
| -1.08535 | -4.32084 | 5.00E-05 | 0.00013743 | yes |
| -1.08552 | -3.68905 | 5.00E-05 | 0.00013743 | yes |
| -1.08736 | -4.14919 | 5.00E-05 | 0.00013743 | yes |
| -1.09192 | -4.45896 | 5.00E-05 | 0.00013743 | yes |
| -1.09529 | -4.41501 | 5.00E-05 | 0.00013743 | yes |
| -1.0983 | -4.48975 | 5.00E-05 | 0.00013743 | yes |
| -1.10107 | -4.06317 | 5.00E-05 | 0.00013743 | yes |
| -1.10583 | -2.2421 | 0.00055 | 0.00130598 | yes |
| -1.10601 | -4.54276 | 5.00E-05 | 0.00013743 | yes |
| -1.10602 | -3.78391 | 5.00E-05 | 0.00013743 | yes |
| -1.10899 | -3.80823 | 5.00E-05 | 0.00013743 | yes |
| -1.11618 | -4.56811 | 5.00E-05 | 0.00013743 | yes |
| -1.1212 | -3.15083 | 5.00E-05 | 0.00013743 | yes |
| -1.12595 | -4.63474 | 5.00E-05 | 0.00013743 | yes |
| -1.12666 | -3.68068 | 5.00E-05 | 0.00013743 | yes |
| -1.12836 | -3.31996 | 5.00E-05 | 0.00013743 | yes |
| -1.13089 | -3.86209 | 5.00E-05 | 0.00013743 | yes |
| -1.1385 | -4.60465 | 5.00E-05 | 0.00013743 | yes |
| -1.13976 | -2.04969 | 0.0015 | 0.00327128 | yes |
| -1.14253 | -4.62653 | 5.00E-05 | 0.00013743 | yes |
| -1.14378 | -4.10619 | 5.00E-05 | 0.00013743 | yes |
| -1.14569 | -4.33771 | 5.00E-05 | 0.00013743 | yes |

| | | | | | |
|---|---|---|---|---|---|
| -1.14692 | -0.632922 | 0.2397 | 0.314151 | | no |
| -1.14995 | -4.38211 | 5.00E-05 | 0.00013743 | yes | |
| -1.1541 | -4.21229 | 5.00E-05 | 0.00013743 | yes | |
| -1.15478 | -4.35547 | 5.00E-05 | 0.00013743 | yes | |
| -1.1555 | -4.60665 | 5.00E-05 | 0.00013743 | yes | |
| -1.15603 | -4.53055 | 5.00E-05 | 0.00013743 | yes | |
| -1.15778 | -4.3732 | 5.00E-05 | 0.00013743 | yes | |
| -1.15971 | -2.7557 | 0.0001 | 0.00026595 | yes | |
| -1.16055 | -3.67088 | 5.00E-05 | 0.00013743 | yes | |
| -1.16456 | -4.80613 | 5.00E-05 | 0.00013743 | | yes |
| -1.1652 | -4.29858 | 5.00E-05 | 0.00013743 | yes | |
| -1.16661 | -2.5398 | 0.00015 | 0.00038924 | yes | |
| -1.16711 | -2.20846 | 0.0009 | 0.00204621 | yes | |
| -1.16854 | -4.66048 | 5.00E-05 | 0.00013743 | yes | |
| -1.17485 | -3.85852 | 5.00E-05 | 0.00013743 | yes | |
| -1.17664 | -4.04152 | 5.00E-05 | 0.00013743 | yes | |
| -1.18294 | -4.83167 | 5.00E-05 | 0.00013743 | yes | |
| -1.18419 | -3.07364 | 5.00E-05 | 0.00013743 | yes | |
| -1.19256 | -4.37227 | 5.00E-05 | 0.00013743 | yes | |
| -1.19509 | -2.11517 | 0.0003 | 0.00074621 | yes | |
| -1.19917 | -2.20161 | 0.00035 | 0.00086186 | yes | |
| -1.21076 | -2.40856 | 5.00E-05 | 0.00013743 | yes | |
| -1.21263 | -4.71969 | 5.00E-05 | 0.00013743 | yes | |
| -1.21277 | -4.45507 | 5.00E-05 | 0.00013743 | yes | |
| -1.21348 | -3.15755 | 5.00E-05 | 0.00013743 | yes | |
| -1.21416 | -4.50016 | 5.00E-05 | 0.00013743 | yes | |
| -1.21497 | -3.43772 | 5.00E-05 | 0.00013743 | yes | |
| -1.21582 | -4.76531 | 5.00E-05 | 0.00013743 | yes | |
| -1.2226 | -3.71114 | 5.00E-05 | 0.00013743 | yes | |
| -1.22429 | -4.84255 | 5.00E-05 | 0.00013743 | yes | |
| -1.22521 | -5.03003 | 5.00E-05 | 0.00013743 | yes | |
| -1.23061 | -5.07805 | 5.00E-05 | 0.00013743 | yes | |
| -1.24815 | -5.14627 | 5.00E-05 | 0.00013743 | yes | |
| -1.24896 | -2.03679 | 0.00155 | 0.00336243 | yes | |
| -1.24948 | -4.1993 | 5.00E-05 | 0.00013743 | yes | |
| -1.2507 | -5.0472 | 5.00E-05 | 0.00013743 | yes | |
| -1.2538 | -5.06745 | 5.00E-05 | 0.00013743 | yes | |
| -1.26875 | -4.29743 | 5.00E-05 | 0.00013743 | yes | |
| -1.27645 | -4.51802 | 5.00E-05 | 0.00013743 | yes | |
| -1.27705 | -4.61925 | 5.00E-05 | 0.00013743 | yes | |
| -1.27851 | -3.70343 | 5.00E-05 | 0.00013743 | yes | |
| -1.27879 | -5.23918 | 5.00E-05 | 0.00013743 | yes | |
| -1.28097 | -4.3267 | 5.00E-05 | 0.00013743 | yes | |
| -1.29219 | -3.79946 | 5.00E-05 | 0.00013743 | yes | |
| -1.2969 | -1.73908 | 0.00425 | 0.0084862 | yes | |
| -1.30375 | -2.53126 | 0.00015 | 0.00038924 | yes | |
| -1.30857 | -5.34753 | 5.00E-05 | 0.00013743 | | yes |
| -1.30909 | -1.71259 | 0.0055 | 0.0107809 | yes | |
| -1.30924 | -3.67765 | 5.00E-05 | 0.00013743 | yes | |
| -1.31254 | -5.43155 | 5.00E-05 | 0.00013743 | yes | |
| -1.31357 | -2.70265 | 5.00E-05 | 0.00013743 | yes | |
| -1.31383 | -4.98275 | 5.00E-05 | 0.00013743 | yes | |
| -1.32704 | -2.58165 | 5.00E-05 | 0.00013743 | yes | |
| -1.32734 | -5.04865 | 5.00E-05 | 0.00013743 | yes | |
| -1.33771 | -3.13906 | 5.00E-05 | 0.00013743 | yes | |

| | | | | | |
|---|---|---|---|---|---|
| -1.33867 | -2.76454 | 5.00E-05 | 0.00013743 | yes | |
| -1.34074 | -5.42854 | 5.00E-05 | 0.00013743 | yes | |
| -1.34097 | -3.33355 | 5.00E-05 | 0.00013743 | yes | |
| -1.34451 | -5.52511 | 5.00E-05 | 0.00013743 | yes | |
| -1.34612 | -5.46668 | 5.00E-05 | 0.00013743 | yes | |
| -1.34625 | -3.86921 | 5.00E-05 | 0.00013743 | yes | |
| -1.34878 | -4.98707 | 5.00E-05 | 0.00013743 | yes | |
| -1.35568 | -3.83986 | 5.00E-05 | 0.00013743 | yes | |
| -1.35714 | -5.08757 | 5.00E-05 | 0.00013743 | yes | |
| -1.36941 | -3.96186 | 5.00E-05 | 0.00013743 | yes | |
| -1.36993 | -5.21258 | 5.00E-05 | 0.00013743 | yes | |
| -1.37384 | -5.48564 | 5.00E-05 | 0.00013743 | yes | |
| -1.37587 | -2.64573 | 5.00E-05 | 0.00013743 | yes | |
| -1.38202 | -5.20963 | 5.00E-05 | 0.00013743 | yes | |
| -1.38204 | -2.46372 | 0.0001 | 0.00026595 | yes | |
| -1.39061 | -3.95645 | 5.00E-05 | 0.00013743 | yes | |
| -1.39422 | -5.68658 | 5.00E-05 | 0.00013743 | yes | |
| -1.39863 | -5.60806 | 5.00E-05 | 0.00013743 | yes | |
| -1.40717 | -5.75407 | 5.00E-05 | 0.00013743 | yes | |
| -1.41443 | -3.10259 | 5.00E-05 | 0.00013743 | yes | |
| -1.41853 | -5.82551 | 5.00E-05 | 0.00013743 | yes | |
| -1.42145 | -5.71041 | 5.00E-05 | 0.00013743 | yes | |
| -1.42743 | -3.5651 | 5.00E-05 | 0.00013743 | yes | |
| -1.43162 | -2.9424 | 5.00E-05 | 0.00013743 | yes | |
| -1.43236 | -2.95626 | 5.00E-05 | 0.00013743 | yes | |
| -1.43397 | -5.60503 | 5.00E-05 | 0.00013743 | yes | |
| -1.4359 | -4.88644 | 5.00E-05 | 0.00013743 | yes | |
| -1.43679 | -5.83508 | 5.00E-05 | 0.00013743 | yes | |
| -1.43739 | -2.50352 | 5.00E-05 | 0.00013743 | yes | |
| -1.43881 | -5.51625 | 5.00E-05 | 0.00013743 | yes | |
| -1.44342 | -2.86842 | 5.00E-05 | 0.00013743 | yes | |
| -1.4469 | -4.51727 | 5.00E-05 | 0.00013743 | yes | |
| -1.44788 | -3.12843 | 5.00E-05 | 0.00013743 | yes | |
| -1.45159 | -5.94321 | 5.00E-05 | 0.00013743 | yes | |
| -1.45379 | -4.27912 | 5.00E-05 | 0.00013743 | yes | |
| -1.46572 | -5.68934 | 5.00E-05 | 0.00013743 | yes | |
| -1.46884 | -4.92745 | 5.00E-05 | 0.00013743 | yes | |
| -1.47219 | -1.74648 | 0.0051 | 0.0100933 | yes | |
| -1.47448 | -4.16545 | 5.00E-05 | 0.00013743 | yes | |
| -1.48429 | -5.84502 | 5.00E-05 | 0.00013743 | yes | |
| -1.48475 | -4.26845 | 5.00E-05 | 0.00013743 | yes | |
| -1.48554 | -4.02423 | 5.00E-05 | 0.00013743 | yes | |
| -1.4895 | -5.40228 | 5.00E-05 | 0.00013743 | yes | |
| -1.4909 | -5.44478 | 5.00E-05 | 0.00013743 | yes | |
| -1.49526 | -4.63544 | 5.00E-05 | 0.00013743 | yes | |
| -1.49571 | -4.46163 | 5.00E-05 | 0.00013743 | yes | |
| -1.50172 | -2.95593 | 5.00E-05 | 0.00013743 | yes | |
| -1.53032 | -6.30266 | 5.00E-05 | 0.00013743 | | yes |
| -1.53102 | -6.23934 | 5.00E-05 | 0.00013743 | yes | |
| -1.53374 | -5.45054 | 5.00E-05 | 0.00013743 | yes | |
| -1.5351 | -3.54585 | 5.00E-05 | 0.00013743 | yes | |
| -1.54339 | -6.36301 | 5.00E-05 | 0.00013743 | yes | |
| -1.54744 | -4.92142 | 5.00E-05 | 0.00013743 | yes | |
| -1.55166 | -4.45664 | 5.00E-05 | 0.00013743 | yes | |
| -1.55378 | -6.24375 | 5.00E-05 | 0.00013743 | yes | |

| | | | | | |
|---|---|---|---|---|---|
| -1.56476 | -5.03224 | 5.00E-05 | 0.00013743 | yes | |
| -1.56564 | -5.90077 | 5.00E-05 | 0.00013743 | yes | |
| -1.58045 | -4.73484 | 5.00E-05 | 0.00013743 | yes | |
| -1.58736 | -6.40231 | 5.00E-05 | 0.00013743 | yes | |
| -1.59268 | -3.57639 | 5.00E-05 | 0.00013743 | yes | |
| -1.59272 | -6.51463 | 5.00E-05 | 0.00013743 | yes | |
| -1.5939 | -5.13921 | 5.00E-05 | 0.00013743 | yes | |
| -1.6038 | -3.84679 | 5.00E-05 | 0.00013743 | yes | |
| -1.60384 | -5.55114 | 5.00E-05 | 0.00013743 | yes | |
| -1.6079 | -6.6617 | 5.00E-05 | 0.00013743 | yes | |
| -1.61515 | -6.3535 | 5.00E-05 | 0.00013743 | yes | |
| -1.61954 | -6.54677 | 5.00E-05 | 0.00013743 | yes | |
| -1.62658 | -6.57783 | 5.00E-05 | 0.00013743 | yes | |
| -1.63547 | -4.8598 | 5.00E-05 | 0.00013743 | yes | |
| -1.64532 | -2.59701 | 5.00E-05 | 0.00013743 | yes | |
| -1.64904 | -1.93778 | 0.00375 | 0.00757389 | yes | |
| -1.66858 | -6.72227 | 5.00E-05 | 0.00013743 | yes | |
| -1.71591 | -6.00114 | 5.00E-05 | 0.00013743 | yes | |
| -1.73391 | -3.63359 | 5.00E-05 | 0.00013743 | yes | |
| -1.73526 | -6.75093 | 5.00E-05 | 0.00013743 | yes | |
| -1.73558 | -6.98342 | 5.00E-05 | 0.00013743 | yes | |
| -1.73654 | -6.73361 | 5.00E-05 | 0.00013743 | yes | |
| -1.73867 | -5.60794 | 5.00E-05 | 0.00013743 | yes | |
| -1.76102 | -2.30625 | 0.0012 | 0.00266186 | yes | |
| -1.76956 | -2.32972 | 0.0004 | 0.00097813 | yes | |
| -1.77088 | -5.6467 | 5.00E-05 | 0.00013743 | yes | |
| -1.77384 | -6.78175 | 5.00E-05 | 0.00013743 | yes | |
| -1.77385 | -4.3008 | 5.00E-05 | 0.00013743 | yes | |
| -1.77483 | -4.78362 | 5.00E-05 | 0.00013743 | yes | |
| -1.7754 | -7.22818 | 5.00E-05 | 0.00013743 | yes | |
| -1.78214 | -6.88406 | 5.00E-05 | 0.00013743 | yes | |
| -1.7976 | -6.46646 | 5.00E-05 | 0.00013743 | yes | |
| -1.80154 | -2.06979 | 0.00115 | 0.00256018 | | yes |
| -1.8145 | -4.36074 | 5.00E-05 | 0.00013743 | yes | |
| -1.82202 | -7.46095 | 5.00E-05 | 0.00013743 | yes | |
| -1.82246 | -3.28093 | 5.00E-05 | 0.00013743 | yes | |
| -1.82326 | -7.22201 | 5.00E-05 | 0.00013743 | yes | |
| -1.82792 | -6.98083 | 5.00E-05 | 0.00013743 | yes | |
| -1.82808 | -4.52677 | 5.00E-05 | 0.00013743 | yes | |
| -1.83397 | -3.36114 | 5.00E-05 | 0.00013743 | yes | |
| -1.84261 | -7.39319 | 5.00E-05 | 0.00013743 | yes | |
| -1.85749 | -5.90366 | 5.00E-05 | 0.00013743 | yes | |
| -1.86084 | -7.62987 | 5.00E-05 | 0.00013743 | yes | |
| -1.9039 | -7.71247 | 5.00E-05 | 0.00013743 | yes | |
| -1.90426 | -2.57889 | 5.00E-05 | 0.00013743 | yes | |
| -1.90848 | -7.19225 | 5.00E-05 | 0.00013743 | yes | |
| -1.90938 | -7.68918 | 5.00E-05 | 0.00013743 | yes | |
| -1.91165 | -3.78137 | 5.00E-05 | 0.00013743 | yes | |
| -1.91242 | -3.62785 | 5.00E-05 | 0.00013743 | yes | |
| -1.91659 | -7.73661 | 5.00E-05 | 0.00013743 | yes | |
| -1.92952 | -6.49653 | 5.00E-05 | 0.00013743 | yes | |
| -1.93371 | -5.11772 | 5.00E-05 | 0.00013743 | yes | |
| -1.9396 | -7.71499 | 5.00E-05 | 0.00013743 | yes | |
| -1.94226 | -2.82906 | 0.0003 | 0.00074621 | yes | |
| -1.95986 | -7.58388 | 5.00E-05 | 0.00013743 | yes | |

| | | | | | |
|---|---|---|---|---|---|
| -1.97832 | -4.43851 | 5.00E-05 | 0.00013743 | yes | |
| -1.98178 | -5.9667 | 5.00E-05 | 0.00013743 | yes | |
| -1.99149 | -8.16065 | 5.00E-05 | 0.00013743 | yes | |
| -1.99357 | -3.80816 | 5.00E-05 | 0.00013743 | yes | |
| -1.9996 | -7.90126 | 5.00E-05 | 0.00013743 | yes | |
| -2.00443 | -6.80575 | 0.00005 | 0.00013743 | yes | |
| -2.00624 | -4.43282 | 5.00E-05 | 0.00013743 | | yes |
| -2.01445 | -6.53562 | 5.00E-05 | 0.00013743 | | yes |
| -2.02725 | -8.28772 | 5.00E-05 | 0.00013743 | | yes |
| -2.04062 | -8.19651 | 5.00E-05 | 0.00013743 | | yes |
| -2.0494 | -8.2713 | 5.00E-05 | 0.00013743 | | yes |
| -2.05371 | -7.94591 | 5.00E-05 | 0.00013743 | | yes |
| -2.06701 | -7.2345 | 5.00E-05 | 0.00013743 | | yes |
| -2.07551 | -8.43841 | 5.00E-05 | 0.00013743 | | yes |
| -2.10256 | -6.98117 | 5.00E-05 | 0.00013743 | | yes |
| -2.1072 | -8.58673 | 5.00E-05 | 0.00013743 | | yes |
| -2.11627 | -7.80104 | 5.00E-05 | 0.00013743 | | yes |
| -2.12268 | -8.36942 | 5.00E-05 | 0.00013743 | | yes |
| -2.12379 | -5.75652 | 5.00E-05 | 0.00013743 | | yes |
| -2.13874 | -7.82673 | 5.00E-05 | 0.00013743 | | yes |
| -2.14125 | -8.83247 | 5.00E-05 | 0.00013743 | | yes |
| -2.15099 | -7.28821 | 5.00E-05 | 0.00013743 | | yes |
| -2.16044 | -8.42995 | 5.00E-05 | 0.00013743 | | yes |
| -2.1658 | -8.32054 | 5.00E-05 | 0.00013743 | | yes |
| -2.17651 | -6.83418 | 5.00E-05 | 0.00013743 | | yes |
| -2.20594 | -5.58714 | 5.00E-05 | 0.00013743 | | yes |
| -2.2123 | -8.86939 | 5.00E-05 | 0.00013743 | | yes |
| -2.22329 | -1.8125 | 0.0271 | 0.0464571 | | yes |
| -2.23598 | -9.02477 | 5.00E-05 | 0.00013743 | | yes |
| -2.23831 | -8.22517 | 5.00E-05 | 0.00013743 | | yes |
| -2.2466 | -9.14825 | 5.00E-05 | 0.00013743 | | yes |
| -2.2533 | -9.20132 | 5.00E-05 | 0.00013743 | | yes |
| -2.26436 | -8.39801 | 5.00E-05 | 0.00013743 | | yes |
| -2.27472 | -9.2624 | 5.00E-05 | 0.00013743 | | yes |
| -2.28021 | -9.2897 | 5.00E-05 | 0.00013743 | | yes |
| -2.28594 | -8.78394 | 5.00E-05 | 0.00013743 | | yes |
| -2.31719 | -9.55443 | 5.00E-05 | 0.00013743 | | yes |
| -2.321 | -2.25264 | 0.0011 | 0.00246224 | | yes |
| -2.3334 | -8.8591 | 5.00E-05 | 0.00013743 | | yes |
| -2.35015 | -7.99891 | 5.00E-05 | 0.00013743 | | yes |
| -2.35823 | -5.98769 | 5.00E-05 | 0.00013743 | | yes |
| -2.38727 | -5.78595 | 5.00E-05 | 0.00013743 | | yes |
| -2.38783 | -7.78887 | 5.00E-05 | 0.00013743 | | yes |
| -2.38895 | -7.4767 | 5.00E-05 | 0.00013743 | | yes |
| -2.39039 | -9.81991 | 5.00E-05 | 0.00013743 | | yes |
| -2.41527 | -5.64883 | 5.00E-05 | 0.00013743 | | yes |
| -2.43829 | -7.16874 | 5.00E-05 | 0.00013743 | | yes |
| -2.45555 | -10.0601 | 5.00E-05 | 0.00013743 | | yes |
| -2.46537 | -4.09467 | 5.00E-05 | 0.00013743 | | yes |
| -2.48513 | -10.2548 | 5.00E-05 | 0.00013743 | | yes |
| -2.4986 | -4.7162 | 5.00E-05 | 0.00013743 | | yes |
| -2.52554 | -3.87137 | 5.00E-05 | 0.00013743 | | yes |
| -2.54049 | -3.46342 | 5.00E-05 | 0.00013743 | | yes |
| -2.55792 | -6.45476 | 5.00E-05 | 0.00013743 | | yes |
| -2.5749 | -10.4889 | 5.00E-05 | 0.00013743 | | yes |

| | | | | |
|---|---|---|---|---|
| -2.62428 | -10.1917 | 5.00E-05 | 0.00013743 | yes |
| -2.63892 | -9.96797 | 5.00E-05 | 0.00013743 | yes |
| -2.63968 | -10.614 | 5.00E-05 | 0.00013743 | yes |
| -2.64082 | -10.4281 | 5.00E-05 | 0.00013743 | yes |
| -2.65983 | -8.43356 | 5.00E-05 | 0.00013743 | yes |
| -2.69741 | -11.0759 | 5.00E-05 | 0.00013743 | yes |
| -2.70187 | -10.677 | 5.00E-05 | 0.00013743 | yes |
| -2.71447 | -10.7019 | 5.00E-05 | 0.00013743 | yes |
| -2.72795 | -2.93604 | 0.00155 | 0.00336243 | yes |
| -2.76162 | -11.1353 | 5.00E-05 | 0.00013743 | yes |
| -2.76582 | -9.86253 | 5.00E-05 | 0.00013743 | yes |
| -2.83978 | -9.14485 | 5.00E-05 | 0.00013743 | yes |
| -3.02992 | -11.6345 | 5.00E-05 | 0.00013743 | yes |
| -3.11263 | -12.7209 | 5.00E-05 | 0.00013743 | yes |
| -3.15327 | -10.3896 | 5.00E-05 | 0.00013743 | yes |
| -3.16874 | -12.4214 | 5.00E-05 | 0.00013743 | yes |
| -3.21081 | -10.5843 | 5.00E-05 | 0.00013743 | yes |
| -3.29776 | -12.3159 | 5.00E-05 | 0.00013743 | yes |
| -3.55991 | -14.5762 | 5.00E-05 | 0.00013743 | yes |
| -3.57355 | -14.1415 | 5.00E-05 | 0.00013743 | yes |
| -3.7736 | -10.2546 | 5.00E-05 | 0.00013743 | yes |
| -3.83831 | -15.4736 | 5.00E-05 | 0.00013743 | yes |
| -3.90254 | -15.5741 | 5.00E-05 | 0.00013743 | yes |
| -3.95327 | -11.2253 | 5.00E-05 | 0.00013743 | yes |
| -4.19358 | -6.37317 | 5.00E-05 | 0.00013743 | yes |
| -4.7749 | -19.5074 | 5.00E-05 | 0.00013743 | yes |
| -4.83876 | -18.3226 | 5.00E-05 | 0.00013743 | yes |
| -4.8476 | -18.9662 | 5.00E-05 | 0.00013743 | yes |
| -4.85097 | -19.8652 | 5.00E-05 | 0.00013743 | yes |
| -5.6553 | -17.9499 | 5.00E-05 | 0.00013743 | yes |
| -7.76437 | -28.3053 | 5.00E-05 | 0.00013743 | yes |
| -7.81752 | -27.4723 | 5.00E-05 | 0.00013743 | yes |
| -7.86836 | -27.5681 | 5.00E-05 | 0.00013743 | yes |
| -8.39088 | -29.0079 | 5.00E-05 | 0.00013743 | yes |
| -8.45389 | -29.4092 | 5.00E-05 | 0.00013743 | yes |
| -9.26452 | -31.8054 | 5.00E-05 | 0.00013743 | yes |

| gene | | | UNIPROTID |
|---|---|---|---|
| SAOUHSC_02416 | | hypothetical protein | Q2FW87 |
| SAOUHSC_01504 | ferredoxin | fer | Q2FYE8 |
| SAOUHSC_01062 | | | Q2G2C3 |
| SAOUHSC_00580 | | | Q2G0I5 |
| SAOUHSC_02093 | | | Q2FX13 |
| SAOUHSC_00244 | | | Q2G1A1 |
| SAOUHSC_02825 | | | Q2FVA8 |
| SAOUHSC_02695 | | | Q2FVL6 |
| SAOUHSC_00863 | | | Q2FZX2 |
| SAOUHSC_00910 | | | Q2FZT0 |
| SAOUHSC_01756 | | | Q2FXS9 |
| SAOUHSC_01835 | | | Q2FXJ9 |
| SAOUHSC_02694 | | | Q2FVL7 |
| SAOUHSC_02271 | | | Q2FWL8 |
| SAOUHSC_02576 | | | Q2G1W1 |
| SAOUHSC_02372 | | | Q2FWC6 |
| SAOUHSC_02779 | | | Q2FVD4 |
| SAOUHSC_00964 | | | Q2FZN6 |
| SAOUHSC_02896 | | | Q2FV43 |
| SAOUHSC_00753 | | | Q2G068 |
| SAOUHSC_01295 | | | Q2FYX8 |
| SAOUHSC_00551 | | | Q2G0K9 |
| SAOUHSC_01438 | | | Q2FYK2 |
| SAOUHSC_02014 | | | Q2G2H1 |
| SAOUHSC_00330 | | | Q2G126 |
| SAOUHSC_01077 | | | Q2FZF3 |
| SAOUHSC_00267 | | | Q2G180 |
| SAOUHSC_00662 | | | Q2G0E3 |
| SAOUHSC_01439 | | | Q2FYK1 |
| SAOUHSC_00669 | | | Q2G0D6 |
| SAOUHSC_02335 | | | Q2FWF6 |
| SAOUHSC_02004 | | | Q2G2M8 |
| SAOUHSC_01710 | | | Q2FXX0 |
| SAOUHSC_00452 | | | Q2G229 |
| SAOUHSC_00345 | | | Q2G116 |
| SAOUHSC_00702 | | | Q2G0A3 |
| SAOUHSC_01332 | | | Q2FYU2 |
| SAOUHSC_02973 | | | Q2FUX2 |
| SAOUHSC_00274 | | | Q2G2Q9 |
| SAOUHSC_02702 | | | Q2FVK9 |
| SAOUHSC_03021 | | | Q2FUT0 |
| SAOUHSC_02529 | | | Q2FVZ2 |
| SAOUHSC_03041 | | | Q2FUR2 |
| SAOUHSC_01930 | | | Q2FXD3 |
| SAOUHSC_01436 | | | Q2FYK4 |
| SAOUHSC_01479 | | | Q2FYH3 |
| SAOUHSC_01874 | | | Q2FXH3 |
| SAOUHSC_02894 | | | Q2FV45 |
| SAOUHSC_02895 | | | Q2FV44 |
| SAOUHSC_01261 | | | Q2FZ10 |
| SAOUHSC_01060 | | | Q2G1Y9 |
| SAOUHSC_02121 | | | Q2FWY6 |
| SAOUHSC_01412 | | | Q2G2J9 |

| | | | |
|---|---|---|---|
| SAOUHSC_00106 | | | Q2G1L6 |
| SAOUHSC_00275 | | | Q2G2Q8 |
| SAOUHSC_00876 | | | Q2FZV9 |
| SAOUHSC_02912 | | | Q2FV27 |
| SAOUHSC_02013 | | | P0A0K3 |
| SAOUHSC_00301 | | | Q2G154 |
| SAOUHSC_01061 | | | Q2G1Y7 |
| SAOUHSC_01480 | | | Q2FYH2 |
| SAOUHSC_01044 | | | Q2G2A2 |
| SAOUHSC_00755 | | | Q2G066 |
| SAOUHSC_00414 | | | Q2G0W0 |
| SAOUHSC_02737 | | | Q2FVH5 |
| SAOUHSC_01192 | | | Q2FZ59 |
| SAOUHSC_00276 | | | Q2G2Q7 |
| SAOUHSC_01752 | | | Q2FXT2 |
| SAOUHSC_00874 | | | Q2FZW1 |
| SAOUHSC_00022 | | | Q2G2U3 |
| SAOUHSC_01813 | | | Q2FXM2 |
| SAOUHSC_00962 | | | Q2FZN8 |
| SAOUHSC_00264 | | | Q2G183 |
| SAOUHSC_02383 | | | Q2FWB5 |
| SAOUHSC_01896 | | | Q2FXF3 |
| SAOUHSC_01073 | | | Q2FZF7 |
| SAOUHSC_01265 | | | Q2FZ06 |
| SAOUHSC_01877 | hypothetical protein | hypothetical protein | Q2FXH0 |
| SAOUHSC_01990 | | | Q2FX87 |
| SAOUHSC_02697 | | | Q2FVL4 |
| SAOUHSC_01709 | | | Q2FXX1 |
| SAOUHSC_02652 | | | Q2FVQ0 |
| SAOUHSC_02625 | | | Q2FVR7 |
| SAOUHSC_01425 | | | Q2FYL4 |
| SAOUHSC_02686 | | | Q2G1V1 |
| SAOUHSC_00441 | | | Q2G0T6 |
| SAOUHSC_02836 | | | Q2FV98 |
| SAOUHSC_00430 | | | Q2G0U6 |
| SAOUHSC_00811 | | | Q2G016 |
| SAOUHSC_00663 | | | Q2G0E2 |
| SAOUHSC_02370 | | | Q2FWC9 |
| SAOUHSC_01392 | | | Q2FYP2 |
| SAOUHSC_00748 | | | Q2G072 |
| SAOUHSC_02820 | | hypothetical protein | Q2FVB4 |
| SAOUHSC_01413 | | | Q2G2J8 |
| SAOUHSC_00847 | | | Q2FZY7 |
| SAOUHSC_00729 | | | Q2G091 |
| SAOUHSC_02274 | | | Q2FWL5 |
| SAOUHSC_02373 | | | Q2FWC5 |
| SAOUHSC_01046 | | | Q2G2A7 |
| SAOUHSC_02483 | | | Q2FW34 |
| SAOUHSC_01657 | | | Q2FY17 |
| SAOUHSC_00794 | | | Q2G033 |
| SAOUHSC_01825 | | | Q2FXL0 |
| SAOUHSC_01284 | | | Q2FYY8 |
| SAOUHSC_02613 | | | Q2FVS9 |
| SAOUHSC_01267 | | | Q2FZ04 |

| | | | |
|---|---|---|---|
| SAOUHSC_01072 | | | Q2FZF8 |
| SAOUHSC_00346 | | | Q2G115 |
| SAOUHSC_02874 | | | Q2FV63 |
| SAOUHSC_01283 | | | Q2FYY9 |
| SAOUHSC_01660 | | | Q2FY14 |
| SAOUHSC_00027 | | | Q2G252 |
| SAOUHSC_00771 | | | Q2G052 |
| SAOUHSC_02359 | | | Q2FWE0 |
| SAOUHSC_00881 | | | Q2FZV4 |
| SAOUHSC_02460 | | | Q2FW57 |
| SAOUHSC_00462 | | | Q2G1S2 |
| SAOUHSC_00803 | | | Q2G024 |
| SAOUHSC_02831 | | | Q2FVA2 |
| SAOUHSC_00909 | | | Q2FZT1 |
| SAOUHSC_03051 | | | Q2FUQ4 |
| SAOUHSC_02915 | | | Q2FV24 |
| SAOUHSC_02830 | D-lactate dehydrogena | ddh | Q2FVA3 |
| SAOUHSC_02897 | | | Q2FV42 |
| SAOUHSC_02935 | | | Q2FV08 |
| SAOUHSC_02867 | | | Q2FV69 |
| SAOUHSC_02388 | | | Q2FWB0 |
| SAOUHSC_02982 | | | Q2FUW9 |
| SAOUHSC_02580 | | N-acetylmuramoyl-L-alanine a | Q2FVW2 |
| SAOUHSC_01843 | | | Q2FXJ2 |
| SAOUHSC_01895 | | | Q2FXF4 |
| SAOUHSC_01110 | | | Q2FZC2 |
| SAOUHSC_02579 | | | Q2FVW3 |
| SAOUHSC_00039 | | | Q2G1R3 |
| SAOUHSC_00302 | | | Q2G153 |
| SAOUHSC_01700 | | | Q2FXY0 |
| SAOUHSC_02824 | | | Q2FVA9 |
| SAOUHSC_02139 | | | Q2FWY2 |
| SAOUHSC_02605 | | | Q2FVT7 |
| SAOUHSC_00531 | | | Q2G0M9 |
| SAOUHSC_02936 | | | Q2FV07 |
| SAOUHSC_00024 | | | Q2G2U5 |
| SAOUHSC_02901 | | | Q2FV38 |
| SAOUHSC_01696 | | | Q2G297 |
| SAOUHSC_00550 | | | Q2G0L0 |
| SAOUHSC_02924 | | | Q2G1Y4 |
| SAOUHSC_02627 | | | Q2G2W3 |
| SAOUHSC_00764 | | | Q2G058 |
| SAOUHSC_02685 | | | Q2G1V2 |
| SAOUHSC_01718 | | | Q2FXW3 |
| SAOUHSC_02821 | | membrane spanning prote | Q2FVB3 |
| SAOUHSC_00253 | hypothetical protein | hypothetical protein | Q2G193 |
| SAOUHSC_02866 | hypothetical protein | farE | Q2FV70 |
| SAOUHSC_00255 | hypothetical protein | hypothetical protein | Q2G191 |
| SAOUHSC_02823 | | | Q2FVB0 |
| SAOUHSC_02630 | | | Q2G2W5 |
| SAOUHSC_00853 | | | Q2FZY1 |
| SAOUHSC_00701 | | | Q2G0A4 |
| SAOUHSC_02448 | | | Q2G2D6 |
| SAOUHSC_00700 | | | Q2G0A5 |

| | | | |
|---|---|---|---|
| SAOUHSC_00844 | | | Q2FZZ0 |
| SAOUHSC_01984 | | | Q2FX93 |
| SAOUHSC_00660 | | | Q2G0E6 |
| SAOUHSC_02978 | | | Q2G221 |
| SAOUHSC_00428 | | | Q2G0U8 |
| SAOUHSC_01022 | | | Q2FZI2 |
| SAOUHSC_02628 | | | Q2G2W2 |
| SAOUHSC_00012 | | | Q2G2T5 |
| SAOUHSC_01908 | | | Q2G1W5 |
| SAOUHSC_00299 | | | Q2G156 |
| SAOUHSC_00254 | | | Q2G192 |
| SAOUHSC_02338 | | | Q2FWF3 |
| SAOUHSC_00050 | | | Q2G1Q3 |
| SAOUHSC_00014 | | | Q2G2T2 |
| SAOUHSC_02524 | | | Q2FVZ6 |
| SAOUHSC_01899 | | | Q2FXF0 |
| SAOUHSC_01630 | | | Q2FY36 |
| SAOUHSC_01050 | | | Q2G2G6 |
| SAOUHSC_01423 | | | Q2FYL6 |
| SAOUHSC_00740 | | | Q2G080 |
| SAOUHSC_00725 | | | Q2G096 |
| SAOUHSC_02668 | | | Q2FVN4 |
| SAOUHSC_02426 | | | Q2FW78 |
| SAOUHSC_00335 | | | Q2G1V3 |
| SAOUHSC_00334 | | | Q2G1V6 |
| SAOUHSC_00683 | | | Q2G0C2 |
| SAOUHSC_01762 | | | Q2FXS3 |
| SAOUHSC_02591 | | | Q2FVV0 |
| SAOUHSC_00718 | | | Q2G2F9 |
| SAOUHSC_01271 | | | Q2FZ00 |
| SAOUHSC_00991 | | | Q2FZL0 |
| SAOUHSC_01981 | | | Q2FX96 |
| SAOUHSC_00802 | | | Q2G025 |
| SAOUHSC_01316 | | | Q2FYV8 |
| SAOUHSC_00661 | | | Q2G0E5 |
| SAOUHSC_01103 | | | Q2FZC9 |
| SAOUHSC_02660 | | | Q2FVP2 |
| SAOUHSC_00174 | | | Q2G1F1 |
| SAOUHSC_02629 | | | Q2G2W1 |
| SAOUHSC_02700 | | | Q2FVL1 |
| SAOUHSC_02750 | | | Q2FVG4 |
| SAOUHSC_02733 | | | Q2FVH8 |
| SAOUHSC_00569 | | | Q2G0J4 |
| SAOUHSC_02321 | | | Q2FWH0 |
| SAOUHSC_00093 | superoxide dismutase | sodM | Q2G261 |
| SAOUHSC_01653 | | | P0A0J3 |
| SAOUHSC_00457 | | | Q2G2X0 |
| SAOUHSC_02855 | | | Q2FV81 |
| SAOUHSC_02886 | | | Q2FV53 |
| SAOUHSC_00483 | | | Q2G0R3 |
| SAOUHSC_01812 | | | Q2FXM3 |
| SAOUHSC_02554 | | | Q2FVW9 |
| SAOUHSC_02898 | | | Q2FV41 |
| SAOUHSC_01701 | | | Q2FXX9 |

| | | | |
|---|---|---|---|
| SAOUHSC_00880 | hypothetical protein | hypothetical protein | Q2FZV5 |
| SAOUHSC_02272 | | | Q2FWL7 |
| SAOUHSC_01823 | | | Q2FXL2 |
| SAOUHSC_02611 | | | Q2FVT1 |
| SAOUHSC_02789 | | | Q2FVC4 |
| SAOUHSC_01019 | | | Q2FZI4 |
| SAOUHSC_01978 | | | Q2FX99 |
| SAOUHSC_02351 | | | Q2G2F7 |
| SAOUHSC_02615 | | | Q2FVS6 |
| SAOUHSC_02641 | | | Q2FVR0 |
| SAOUHSC_01338 | | | Q2FYT7 |
| SAOUHSC_01871 | | | Q2FXH6 |
| SAOUHSC_00404 | | | Q2G0X0 |
| SAOUHSC_00402 | | | Q2G0X1 |
| SAOUHSC_03020 | | | Q2FUT1 |
| SAOUHSC_01747 | | | Q2FXT7 |
| SAOUHSC_01361 | | | Q7BHL7 |
| SAOUHSC_00854 | | | Q2FZY0 |
| SAOUHSC_02131 | | | Q2G2R9 |
| SAOUHSC_00842 | | | Q2FZZ2 |
| SAOUHSC_00423 | | | Q2G0V2 |
| SAOUHSC_01354 | | | Q2G2N5 |
| SAOUHSC_00949 | | | Q2FZQ1 |
| SAOUHSC_02573 | | | Q2G2J3 |
| SAOUHSC_01385 | | | Q2FYQ0 |
| SAOUHSC_02520 | | | Q2FW00 |
| SAOUHSC_02624 | | | Q2FVR8 |
| SAOUHSC_03033 | | | Q2FUR9 |
| SAOUHSC_01032 | | | Q2FZH2 |
| SAOUHSC_02341 | | | Q2FWF0 |
| SAOUHSC_02347 | | | Q2G2F8 |
| SAOUHSC_02516 | | | Q2FW04 |
| SAOUHSC_02815 | hypothetical protein | hypothetical protein | Q2FVB9 |
| SAOUHSC_00556 | | | Q2G0K4 |
| SAOUHSC_01876 | | | Q2FXH1 |
| SAOUHSC_02531 | | | Q2FVZ0 |
| SAOUHSC_00703 | | | Q2G0A2 |
| SAOUHSC_02525 | | | Q2FVZ5 |
| SAOUHSC_00413 | | | Q2G0W1 |
| SAOUHSC_02382 | | | Q2FWB6 |
| SAOUHSC_01041 | | | Q2G2A5 |
| SAOUHSC_00526 | | | P0A0G6 |
| SAOUHSC_01698 | | | Q2FXY2 |
| SAOUHSC_00049 | | | Q2G1Q4 |
| SAOUHSC_00523 | | | Q2G0N7 |
| SAOUHSC_01661 | | | Q2FY13 |
| SAOUHSC_00129 | | | Q2G1J3 |
| SAOUHSC_01727 | | | Q2FXV4 |
| SAOUHSC_02829 | | | Q2FVA4 |
| SAOUHSC_00549 | | | Q2G0L1 |
| SAOUHSC_02921 | | | Q2FV19 |
| SAOUHSC_02467 | | | Q2FW50 |
| SAOUHSC_01040 | | | Q2FZG4 |
| SAOUHSC_02860 | | | Q2FV76 |

| | | | |
|---|---|---|---|
| SAOUHSC_02091 | | | Q2FX15 |
| SAOUHSC_01743 | | | Q2FXU0 |
| SAOUHSC_01031 | cytochrome d ubiquinol | cydA | Q2FZH3 |
| SAOUHSC_00878 | | | Q2FZV7 |
| SAOUHSC_00999 | | | Q2FZK2 |
| SAOUHSC_00412 | | | Q2G0W2 |
| SAOUHSC_02916 | | | Q2FV23 |
| SAOUHSC_01452 | | | Q2FYJ2 |
| SAOUHSC_01326 | | | Q2FYU8 |
| SAOUHSC_01450 | | | Q2FYJ4 |
| SAOUHSC_02923 | hypothetical protein | hypothetical protein | Q2G1Y2 |
| SAOUHSC_00843 | | | Q2FZZ1 |
| SAOUHSC_00828 | | | Q2G1T0 |
| SAOUHSC_01991 | | | Q2FX86 |
| SAOUHSC_02698 | | | Q2FVL3 |
| SAOUHSC_02590 | | | Q2FVV1 |
| SAOUHSC_00435 | glutamate synthase larg | gltB | Q2G0U2 |
| SAOUHSC_01104 | | | Q2FZC8 |
| SAOUHSC_02680 | | | Q2FVM2 |
| SAOUHSC_00150 | | | Q2G1H3 |
| SAOUHSC_01635 | | | Q2FY32 |
| SAOUHSC_01483 | | | Q2FYG9 |
| SAOUHSC_01699 | | | Q2FXY1 |
| SAOUHSC_01497 | | | Q2FYF4 |
| SAOUHSC_01471 | | | Q2FYH6 |
| SAOUHSC_01737 | | | Q2FXU5 |
| SAOUHSC_00886 | | | Q2G2H7 |
| SAOUHSC_02340 | | | Q2FWF1 |
| SAOUHSC_02349 | | | Q2G2F6 |
| SAOUHSC_02345 | | | Q2FWE8 |
| SAOUHSC_02346 | | | Q2FWE7 |
| SAOUHSC_02343 | | | Q2FWE9 |
| SAOUHSC_01658 | | | Q2FY16 |
| SAOUHSC_01768 | | | Q2FXR7 |
| SAOUHSC_02005 | | | Q2G2W0 |
| SAOUHSC_02122 | | | Q2G1Y0 |
| SAOUHSC_03012 | | | Q2FUT9 |
| SAOUHSC_00989 | | | Q2FZL1 |
| SAOUHSC_02158 | | | Q2FWW3 |
| SAOUHSC_02286 | | | Q2FWK2 |
| SAOUHSC_02285 | | | Q2FWK3 |
| SAOUHSC_02288 | | | Q2FWK0 |
| SAOUHSC_02287 | | | Q2FWK1 |
| SAOUHSC_02468 | | | Q2FW49 |
| SAOUHSC_00965 | | | Q2FZN5 |
| SAOUHSC_01807 | | | Q2FXM8 |
| SAOUHSC_02449 | | | Q2G2D5 |
| SAOUHSC_03016 | | | Q2FUT5 |
| SAOUHSC_01424 | | | Q2FYL5 |
| SAOUHSC_01491 | | | Q2FYG1 |
| SAOUHSC_00545 | | | Q2G0L4 |
| SAOUHSC_02802 | | | Q2G1T5 |
| SAOUHSC_02963 | | | Q2FUY2 |
| SAOUHSC_02690 | | | Q2G1U8 |

| | | | |
|---|---|---|---|
| SAOUHSC_03002 | | icaA | Q9RQP9 |
| SAOUHSC_01158 | | | Q2FZ83 |
| SAOUHSC_00954 | | | Q2FZP6 |
| SAOUHSC_01856 | | | Q2FXJ0 |
| SAOUHSC_02337 | | | Q2FWF4 |
| SAOUHSC_00482 | | | Q2G0R4 |
| SAOUHSC_01150 | | | Q2FZ89 |
| SAOUHSC_01149 | | | O07325 |
| SAOUHSC_01778 | | | Q2FXQ7 |
| SAOUHSC_00471 | | | Q2G0S3 |
| SAOUHSC_00581 | hypothetical protein | hypothetical protein | Q2G0I4 |
| SAOUHSC_01030 | | | Q2FZH4 |
| SAOUHSC_01043 | | | Q2G2A3 |
| SAOUHSC_01652 | | | Q2FY21 |
| SAOUHSC_02816 | | | Q2FVB8 |
| SAOUHSC_00427 | | | Q2G0U9 |
| SAOUHSC_00773 | | | Q2G278 |
| SAOUHSC_00671 | | | Q2G0D4 |
| SAOUHSC_00691 | | | Q2G0B4 |
| SAOUHSC_00248 | | | O33599 |
| SAOUHSC_00646 | | | Q2G2X6 |
| SAOUHSC_02696 | | | Q2FVL5 |
| SAOUHSC_02318 | | | Q2FWH3 |
| SAOUHSC_01106 | | | Q2FZC6 |
| SAOUHSC_02012 | | | Q93Q23 |
| SAOUHSC_01374 | | | Q2FYR1 |
| SAOUHSC_02527 | | | Q2FVZ4 |
| SAOUHSC_01373 | | | Q2FYR2 |
| SAOUHSC_00994 | | | Q2FZK7 |
| SAOUHSC_00499 | pyridoxal biosynthesis | pdxS | Q2G0Q1 |
| SAOUHSC_01384 | | | Q2FYQ1 |
| SAOUHSC_01066 | | | Q2G2B9 |
| SAOUHSC_00912 | | | Q2FZS8 |
| SAOUHSC_00344 | | | Q2G117 |
| SAOUHSC_02255 | | | Q2FWN3 |
| SAOUHSC_01682 | | | Q2FXZ3 |
| SAOUHSC_02254 | | | Q2FWN4 |
| SAOUHSC_01857 | | | Q2FXI9 |
| SAOUHSC_03049 | | | Q2FUQ5 |
| SAOUHSC_01351 | | | Q2FYS5 |
| SAOUHSC_01352 | | | Q2FYS4 |
| SAOUHSC_00342 | | | Q2G118 |
| SAOUHSC_00083 | | | Q2G1M5 |
| SAOUHSC_01075 | | | Q2FZF5 |
| SAOUHSC_01038 | | | Q2FZG6 |
| SAOUHSC_01715 | | | Q2FXW6 |
| SAOUHSC_01735 | | | Q2FXU7 |
| SAOUHSC_00341 | | | Q2G119 |
| SAOUHSC_02850 | hypothetical protein | cidB | P60639 |
| SAOUHSC_02851 | hypothetical protein | cidA | P60647 |
| SAOUHSC_00480 | | | Q2G0R6 |
| SAOUHSC_00019 | | | Q2G1S3 |
| SAOUHSC_01779 | | | Q2FXQ6 |
| SAOUHSC_02368 | | | Q2FWD1 |

| | | | |
|---|---|---|---|
| SAOUHSC_01008 | | | Q2FZJ5 |
| SAOUHSC_01009 | | | Q2FZJ4 |
| SAOUHSC_01010 | | | Q2FZJ3 |
| SAOUHSC_01018 | | | Q2FZI5 |
| SAOUHSC_01395 | | | Q2FYP0 |
| SAOUHSC_01398 | | | Q2FYN7 |
| SAOUHSC_01396 | | | Q9EZ12 |
| SAOUHSC_01777 | | | Q2FXQ8 |
| SAOUHSC_02360 | | | Q2FWD9 |
| SAOUHSC_02861 | | | Q2FV75 |
| SAOUHSC_00730 | | | Q2G090 |
| SAOUHSC_03042 | integrase/recombinasease/recombinase core s | | Q2FUR1 |
| SAOUHSC_02250 | | | Q2G2N7 |
| SAOUHSC_00445 | | | Q2G0T3 |
| SAOUHSC_00349 | bacteriophage L54a sin | ssb | Q2G112 |
| SAOUHSC_01751 | | | Q2FXT3 |
| SAOUHSC_01750 | | | Q2FXT4 |
| SAOUHSC_00533 | | | Q2G0M7 |
| SAOUHSC_01098 | | | Q2FZD4 |
| SAOUHSC_00004 | | | Q2G275 |
| SAOUHSC_02110 | | | Q2FWZ6 |
| SAOUHSC_01454 | | | Q2FYJ1 |
| SAOUHSC_01215 | | | Q2FZ38 |
| SAOUHSC_00018 | | | Q2G1S4 |
| SAOUHSC_01222 | | | Q2FZ32 |
| SAOUHSC_02517 | | | Q2FW03 |
| SAOUHSC_02123 | | | Q53727 |
| SAOUHSC_01241 | | | Q2G1Z8 |
| SAOUHSC_00442 | | | Q2G0T5 |
| SAOUHSC_00005 | | | Q2G274 |
| SAOUHSC_00006 | | | Q2G2Q0 |
| SAOUHSC_01714 | | | Q2FXW7 |
| SAOUHSC_01897 | | | Q2FXF2 |
| SAOUHSC_02362 | | | Q2FWD7 |
| SAOUHSC_00707 | | | Q2G238 |
| SAOUHSC_00451 | | | Q2G230 |
| SAOUHSC_01163 | | | Q2FZ78 |
| SAOUHSC_01363 | | | Q2FYS2 |
| SAOUHSC_01898 | | | Q2FXF1 |
| SAOUHSC_01249 | | | Q2G2Q2 |
| SAOUHSC_00195 | | | Q2G1D0 |
| SAOUHSC_00920 | 3-oxoacyl-ACP syntha | fabH | Q2FZS0 |
| SAOUHSC_00921 | | | Q2FZR9 |
| SAOUHSC_01198 | | | Q2FZ54 |
| SAOUHSC_02336 | | | Q2FWF5 |
| SAOUHSC_01196 | | | Q2FZ56 |
| SAOUHSC_01197 | | | Q2FZ55 |
| SAOUHSC_01199 | | | Q2FZ53 |
| SAOUHSC_00491 | | | Q2G0Q5 |
| SAOUHSC_02374 | | | Q2FWC4 |
| SAOUHSC_01766 | | | Q2FXR9 |
| SAOUHSC_02582 | | | Q2FVV9 |
| SAOUHSC_00281 | | | Q2G172 |
| SAOUHSC_02614 | | | Q2FVS7 |

| | | | |
|---|---|---|---|
| SAOUHSC_01794 | | | Q2FXP2 |
| SAOUHSC_00795 | | | Q2G032 |
| SAOUHSC_00317 | | | Q2G138 |
| SAOUHSC_01287 | | | Q2FYY6 |
| SAOUHSC_00500 | glutamine amidotransf | pdxT | Q2G0Q0 |
| SAOUHSC_00375 | | | Q2G0Y6 |
| SAOUHSC_03010 | | | Q2FUU1 |
| SAOUHSC_01193 | | | Q2FZ58 |
| SAOUHSC_02933 | | | Q2FV10 |
| SAOUHSC_02932 | | | Q2FV11 |
| SAOUHSC_01634 | | | Q2FY33 |
| SAOUHSC_01806 | | | Q2FXM9 |
| SAOUHSC_01666 | | | Q2FY08 |
| SAOUHSC_00374 | | | Q2G0Y7 |
| SAOUHSC_01742 | | | Q2FXU1 |
| SAOUHSC_03015 | | | Q2FUT6 |
| SAOUHSC_03008 | | | Q2FUU3 |
| SAOUHSC_00733 | | | Q2G087 |
| SAOUHSC_03013 | | | Q2FUT8 |
| SAOUHSC_03009 | | | Q2FUU2 |
| SAOUHSC_01394 | | | Q2FYP1 |
| SAOUHSC_00013 | | | Q2G2T4 |
| SAOUHSC_00801 | | | Q2G026 |
| SAOUHSC_00749 | | | Q2G071 |
| SAOUHSC_02417 | | | Q2FW86 |
| SAOUHSC_00873 | | | Q2FZW2 |
| SAOUHSC_00536 | | | Q2G0M4 |
| SAOUHSC_01320 | | | Q2FYV4 |
| SAOUHSC_02284 | | | Q2FWK4 |
| SAOUHSC_02281 | | | Q2FWK7 |
| SAOUHSC_00282 | | | Q2G171 |
| SAOUHSC_02623 | | | Q2FVR9 |
| SAOUHSC_01875 | | | Q2FXH2 |
| SAOUHSC_02622 | | | Q2FVS0 |
| SAOUHSC_01201 | | | Q2FZ51 |
| SAOUHSC_00872 | | | Q2FZW3 |
| SAOUHSC_01702 | | | Q2FXX8 |
| SAOUHSC_01451 | | | Q2FYJ3 |
| SAOUHSC_00010 | | | Q2G2P6 |
| SAOUHSC_00493 | | | Q2G0Q3 |
| SAOUHSC_02486 | 30S ribosomal protein | rpsK | Q2FW31 |
| SAOUHSC_01695 | | | Q2G298 |
| SAOUHSC_02849 | pyruvate oxidase | cidC | Q2FV86 |
| SAOUHSC_00985 | | | Q2FZL5 |
| SAOUHSC_00980 | | | Q2FZM0 |
| SAOUHSC_02887 | immunodominant antig | isaA | Q2FV52 |
| SAOUHSC_00007 | hypothetical protein | nnrD | Q2G2P8 |
| SAOUHSC_00959 | | | Q2FZP1 |
| SAOUHSC_00461 | | | Q2G1R9 |
| SAOUHSC_02682 | | | Q2FVM0 |
| SAOUHSC_01878 | | | Q2FXG9 |
| SAOUHSC_02550 | | | Q2FVX3 |
| SAOUHSC_02536 | | | P69848 |
| SAOUHSC_02538 | | | Q2FVY3 |

| | | | |
|---|---|---|---|
| SAOUHSC_02537 | | | Q2FVY4 |
| SAOUHSC_01263 | | | Q2FZ08 |
| SAOUHSC_01251 | | | Q2FZ20 |
| SAOUHSC_01035 | | | Q2FZG9 |
| SAOUHSC_01248 | | | Q2G2Q3 |
| SAOUHSC_01697 | | | Q2FXY3 |
| SAOUHSC_01685 | | | Q2FXZ0 |
| SAOUHSC_01793 | | | Q2FXP3 |
| SAOUHSC_00934 | | | Q2G1U6 |
| SAOUHSC_02273 | | | Q2FWL6 |
| SAOUHSC_01386 | | | Q2FYP9 |
| SAOUHSC_01387 | | | Q2FYP8 |
| SAOUHSC_01048 | | | Q2G2A9 |
| SAOUHSC_01047 | | | Q2G2B0 |
| SAOUHSC_02684 | | | Q2FVL8 |
| SAOUHSC_02937 | | | Q2FV06 |
| SAOUHSC_01346 | | | Q2FYT0 |
| SAOUHSC_02270 | | | Q2FWL9 |
| SAOUHSC_00324 | | | Q2G132 |
| SAOUHSC_01496 | | | Q2FYF5 |
| SAOUHSC_01496 | | | Q2FYF5 |
| SAOUHSC_00738 | | | Q2G082 |
| SAOUHSC_01909 | | | Q2G1W4 |
| SAOUHSC_01716 | | | Q2FXW5 |
| SAOUHSC_01717 | | | Q2FXW4 |
| SAOUHSC_01585 | | | Q2FY80 |
| SAOUHSC_00021 | | | Q2G2U4 |
| SAOUHSC_02773 | | transporter | Q2FVE1 |
| SAOUHSC_02919 | | | Q2FV21 |
| SAOUHSC_02918 | | | Q2FV22 |
| SAOUHSC_02739 | | | Q2FVH3 |
| SAOUHSC_02920 | | | Q2FV20 |
| SAOUHSC_00069 | protein A | spa | P02976 |
| SAOUHSC_02571 | secretory antigen | ssaA | Q2G2J2 |
| SAOUHSC_00391 | | | Q2G1S5 |
| SAOUHSC_00394 | | | Q2G2X8 |
| SAOUHSC_00265 | | | Q2G182 |
| SAOUHSC_01437 | | | Q2FYK3 |
| SAOUHSC_00392 | | | Q2G2Y0 |
| SAOUHSC_00268 | | | Q2G179 |
| SAOUHSC_00383 | | | Q2G0X9 |
| SAOUHSC_02883 | | | Q2FV55 |
| SAOUHSC_00266 | | | Q2G181 |
| SAOUHSC_00393 | | | Q2G2X9 |
| SAOUHSC_00395 | | | Q2G2X7 |
| SAOUHSC_01964 | | | Q2G2F3 |
| SAOUHSC_00070 | accessory regulator-lik | sarS | Q2G1N7 |
| SAOUHSC_02669 | | | Q2FVN3 |
| SAOUHSC_00620 | | | Q2G2U9 |
| SAOUHSC_00666 | | | Q2G0D9 |
| SAOUHSC_02798 | | | Q2G2B2 |
| SAOUHSC_00928 | | | Q2FZR2 |
| SAOUHSC_00927 | | | Q2FZR3 |
| SAOUHSC_00167 | | | Q2G1F8 |

| | |
|---|---|
| SAOUHSC_00926 | Q2FZR4 |
| SAOUHSC_00925 | Q2FZR5 |
| SAOUHSC_01467 | Q2FYI0 |
| SAOUHSC_02358 | Q2FWE1 |
| SAOUHSC_01093 | Q2FZD8 |
| SAOUHSC_00670 | Q2G0D5 |
| SAOUHSC_01389 | Q2FYP6 |
| SAOUHSC_00504 | Q2G0P6 |
| SAOUHSC_01992 | Q2FX85 |
| SAOUHSC_00290 | Q2G165 |
| SAOUHSC_02848 | Q2FV87 |
| SAOUHSC_00235 | Q2G1B0 |
| SAOUHSC_02597 | Q2FVU4 |
| SAOUHSC_01028 | Q2FZH6 |
| SAOUHSC_01029 | Q2FZH5 |
| SAOUHSC_00708 | Q2G239 |
| SAOUHSC_00155 | Q2G1G8 |
| SAOUHSC_01350 | Q2FYS6 |
| SAOUHSC_02114 | Q2FWZ2 |
| SAOUHSC_01980 | Q2FX97 |
| SAOUHSC_01176 | Q2G1U0 |
| SAOUHSC_02350 | Q2G2F4 |
| SAOUHSC_01049 | Q2G2A8 |
| SAOUHSC_02140 | Q2FWY1 |
| SAOUHSC_02000 | Q2G283 |
| SAOUHSC_02731 | Q2FVI0 |
| SAOUHSC_01034 | Q2FZH1 |
| SAOUHSC_02119 | Q2FWY7 |
| SAOUHSC_00486 | Q2G0R0 |
| SAOUHSC_01684 | Q2FXZ1 |
| SAOUHSC_01683 | Q2FXZ2 |
| SAOUHSC_02983 | Q2FUW8 |
| SAOUHSC_02327 | Q2FWG4 |
| SAOUHSC_00861 | Q2FZX4 |
| SAOUHSC_02984 | Q2FUW7 |
| SAOUHSC_00891 | Q2FZU9 |
| SAOUHSC_00229 | P72360 |
| SAOUHSC_01205 | Q2FZ48 |
| SAOUHSC_03025 | Q2FUS6 |
| SAOUHSC_02092 | Q2FX14 |
| SAOUHSC_00885 | Q2G2H8 |
| SAOUHSC_01330 | Q2FYU4 |
| SAOUHSC_00372 | Q2G0Y9 |
| SAOUHSC_00734 | Q2G086 |
| SAOUHSC_00721 | Q2G1X6 |
| SAOUHSC_00720 | Q2G1X5 |
| SAOUHSC_00719 | Q2G1X7 |
| SAOUHSC_00739 | Q2G081 |
| SAOUHSC_01989 | Q2FX88 |
| SAOUHSC_02911 | Q2FV28 |
| SAOUHSC_01749 | Q2FXT5 |
| SAOUHSC_01748 | Q2FXT6 |
| SAOUHSC_01440 | Q2FYK0 |
| SAOUHSC_00020 | Q2G2U6 |

| | | | |
|---|---|---|---|
| SAOUHSC_00582 | hypothetical protein | hypothetical protein | Q2G0I3 |
| SAOUHSC_01891 | | | Q2FXF7 |
| SAOUHSC_01045 | | | Q2G2A6 |
| SAOUHSC_01285 | | | Q2FYY7 |
| SAOUHSC_02530 | | | Q2FVZ1 |
| SAOUHSC_02322 | | | Q2FWG9 |
| SAOUHSC_02461 | | | Q2FW56 |
| SAOUHSC_01732 | | | Q2FXV0 |
| SAOUHSC_00992 | | | Q2FZK9 |
| SAOUHSC_03001 | | | Q9RQQ0 |
| SAOUHSC_02826 | | | Q2FVA7 |
| SAOUHSC_03001 | | | Q9RQQ0 |
| SAOUHSC_00502 | | | Q2G0P8 |
| SAOUHSC_02369 | | | Q2FWD0 |
| SAOUHSC_01067 | | | Q2FZG3 |
| SAOUHSC_01185 | | | Q2FZ66 |
| SAOUHSC_01448 | | | Q2FYJ5 |
| SAOUHSC_01893 | | | Q2FXF6 |
| SAOUHSC_01894 | | | Q2FXF5 |
| SAOUHSC_01659 | | | Q2FY15 |
| SAOUHSC_01266 | | | Q2FZ05 |
| SAOUHSC_01282 | | | Q2FYZ0 |
| SAOUHSC_01887 | riboflavin biosynthesis | ribA | Q2FXG1 |
| SAOUHSC_01888 | riboflavin synthase sub | ribB | Q2FXG0 |
| SAOUHSC_01889 | riboflavin biosynthesis | ribD | Q2FXF9 |
| SAOUHSC_01505 | | | Q2FYE7 |
| SAOUHSC_01058 | GTP-binding protein Ty | typA | Q2G1Y6 |
| SAOUHSC_01492 | | | Q2FYG0 |
| SAOUHSC_02316 | | | Q2FWH5 |
| SAOUHSC_00528 | | | P48940 |
| SAOUHSC_01188 | | | Q2FZ63 |
| SAOUHSC_01156 | | | Q2FZ84 |
| SAOUHSC_01753 | | | Q2FXT1 |
| SAOUHSC_01091 | | | Q2FZE0 |
| SAOUHSC_01680 | | | Q2FXZ5 |
| SAOUHSC_02113 | | | Q2FWZ3 |
| SAOUHSC_00464 | | | Q2G0T0 |
| SAOUHSC_00463 | | | Q2G0T1 |
| SAOUHSC_00009 | | | P95689 |
| SAOUHSC_00747 | | | Q2G073 |
| SAOUHSC_00746 | | | Q2G074 |
| SAOUHSC_02427 | | | Q2FW77 |
| SAOUHSC_02428 | | | Q2FW76 |
| SAOUHSC_00903 | | | Q2FZT7 |
| SAOUHSC_00060 | | | Q2G1P4 |
| SAOUHSC_01207 | | | Q2FZ46 |
| SAOUHSC_00503 | | | Q2G0P7 |
| SAOUHSC_00572 | | | Q2G0J2 |
| SAOUHSC_00571 | | | Q2G0J3 |
| SAOUHSC_00562 | | | Q2G0J8 |
| SAOUHSC_01824 | | | Q2FXL1 |
| SAOUHSC_01788 | | | Q2FXP7 |
| SAOUHSC_01662 | | | P0A0J0 |
| SAOUHSC_02485 | DNA-directed RNA pol | rpoA | Q2FW32 |

| | | | |
|---|---|---|---|
| SAOUHSC_00525 | DNA-directed RNA poly | rpoC | Q2G0N5 |
| SAOUHSC_00524 | | | P47768 |
| SAOUHSC_02504 | 50S ribosomal protein | rpmC | Q2FW14 |
| SAOUHSC_01651 | | | Q2FY22 |
| SAOUHSC_00474 | | | Q2G0S0 |
| SAOUHSC_01757 | | | Q2FXS8 |
| SAOUHSC_01078 | | | Q2FZF1 |
| SAOUHSC_01755 | | | Q2FXT0 |
| SAOUHSC_01689 | | | Q2FXY6 |
| SAOUHSC_01191 | | | Q2FZ60 |
| SAOUHSC_02499 | | | Q2FW19 |
| SAOUHSC_01214 | | | Q2FZ39 |
| SAOUHSC_01234 | | | Q2FZ23 |
| SAOUHSC_00246 | | drug transporter | Q2G199 |
| SAOUHSC_02740 | | | Q2FVH2 |
| SAOUHSC_02658 | | | Q2FVP4 |
| SAOUHSC_01021 | | | Q2FZI3 |
| SAOUHSC_02003 | | | Q2G2M9 |
| SAOUHSC_02640 | | | Q2FVR1 |
| SAOUHSC_02752 | | | Q2FVG2 |
| SAOUHSC_02618 | | | Q2FVS4 |
| SAOUHSC_00948 | | | Q2FZQ2 |
| SAOUHSC_00804 | | | Q2G023 |
| SAOUHSC_02927 | | | Q2FV16 |
| SAOUHSC_01042 | | | Q2G2A4 |
| SAOUHSC_01839 | | | Q2FXJ5 |
| SAOUHSC_03053 | | | Q2FUQ2 |
| SAOUHSC_03052 | | | Q2FUQ3 |
| SAOUHSC_01679 | | | Q2FXZ6 |
| SAOUHSC_01269 | | | Q2FZ02 |
| SAOUHSC_03024 | | hypothetical protein | Q2FUS7 |
| SAOUHSC_01076 | | | Q2FZF4 |
| SAOUHSC_02480 | | | Q2FW37 |
| SAOUHSC_01368 | | | Q2FYR7 |
| SAOUHSC_00933 | | | Q2FZQ7 |
| SAOUHSC_00995 | hypothetical protein | hypothetical protein | P0A0M9 |
| SAOUHSC_01767 | | | Q2FXR8 |
| SAOUHSC_01988 | | | Q2FX89 |
| SAOUHSC_00373 | | | Q2G0Y8 |
| SAOUHSC_02725 | hypothetical protein | hypothetical protein | Q2FVI5 |
| SAOUHSC_02389 | | | Q2FWA9 |
| SAOUHSC_02512a | 30S ribosomal protein S | rpsJ | #N/A |
| SAOUHSC_01993 | hypothetical protein | tnp3 | #N/A |
| trmD | | | #N/A |
| rimM | | | #N/A |
| SAOUHSC_02410 | | | #N/A |
| rplA | | | #N/A |
| SAOUHSC_A02450 | | | #N/A |
| hslO | | | #N/A |
| rpsL | | | #N/A |
| rpsD | | | #N/A |
| rplK | | | #N/A |
| rplS | | | #N/A |
| rpsO | | | #N/A |

| | |
|---|---|
| rnpA | #N/A |
| gid | #N/A |
| rpmH | #N/A |
| prfC | #N/A |
| hisS | #N/A |
| SAOUHSC_01726 | #N/A |
| SAOUHSC_1342a | #N/A |
| SAOUHSC_A00992 | #N/A |
| SAUSA300_RS08385 | #N/A |
| SAOUHSC_01055a | #N/A |

GOBIO

0
0
0
0
0
0
0
0
0
0
0
0
0
0
0
0
0
0
0
0
0
0
0
0
0
0
0
0
0
0
0
0
0
0
0
0
0
0
0
0
0
0
0
0
0
0
0
0
0
0
0
0
0

0
0
0
0
0
0
0
0
0
0
0
0
0
0
0
0
0
0
0
0
0
0
0
0
0
0
0
0
0
0
0
0
0
0
0
0
0
0
0
0
0
0
0
0
0
0
0
[2Fe-2S] cluster assembly [GO:0044571]
2,4,6-trinitrotoluene catabolic process [GO:0046256]
7,8-dihydroneopterin 3'-triphosphate biosynthetic process [GO:0035998]
acetoin biosynthetic process [GO:0045151]
acetoin biosynthetic process [GO:0045151]
acetyl-CoA biosynthetic process from pyruvate [GO:0006086]; glycolytic process [GO:0006096]
acetyl-CoA metabolic process [GO:0006084]; farnesyl diphosphate biosynthetic process, mevalonate pathw

acyl-CoA metabolic process [GO:0006637]; fatty acid metabolic process [GO:0006631]
adenine salvage [GO:0006168]; AMP salvage [GO:0044209]; purine ribonucleoside salvage [GO:0006166]
aerobic electron transport chain [GO:0019646]
aerobic electron transport chain [GO:0019646]
aerobic electron transport chain [GO:0019646]; ATP synthesis coupled electron transport [GO:0042773]; ele
aerobic electron transport chain [GO:0019646]; electron transport coupled proton transport [GO:0015990]; p
alanine biosynthetic process [GO:0006523]; pantothenate biosynthetic process [GO:0015940]
alanine catabolic process [GO:0006524]; L-alanine catabolic process [GO:0042853]
amino acid transmembrane transport [GO:0003333]
amino acid transmembrane transport [GO:0003333]
amino acid transport [GO:0006865]
amino acid transport [GO:0006865]; D-methionine transport [GO:0048473]; nickel cation transport [GO:001
amino acid transport [GO:0006865]; L-arginine transport [GO:1902023]
amino acid transport [GO:0006865]; nickel cation transport [GO:0015675]
amino acid transport [GO:0006865]; nickel cation transport [GO:0015675]
amino acid transport [GO:0006865]; transmembrane transport [GO:0055085]
ammonia assimilation cycle [GO:0019676]; glutamate biosynthetic process [GO:0006537]
anaerobic respiration [GO:0009061]
anaerobic respiration [GO:0009061]; nitrate metabolic process [GO:0042126]
arginine metabolic process [GO:0006525]; L-proline biosynthetic process [GO:0055129]
aromatic amino acid family biosynthetic process [GO:0009073]; cellular amino acid biosynthetic process [G
aromatic amino acid family biosynthetic process [GO:0009073]; cellular amino acid biosynthetic process [G
aromatic amino acid family biosynthetic process [GO:0009073]; cellular amino acid biosynthetic process [G
asparagine catabolic process [GO:0006530]
asparaginyl-tRNA aminoacylation [GO:0006421]
aspartyl-tRNA aminoacylation [GO:0006422]
ATP synthesis coupled electron transport [GO:0042773]; sodium ion transport [GO:0006814]
ATP synthesis coupled proton transport [GO:0015986]
ATP synthesis coupled proton transport [GO:0015986]
ATP synthesis coupled proton transport [GO:0015986]
ATP synthesis coupled proton transport [GO:0015986]
ATP synthesis coupled proton transport [GO:0015986]; plasma membrane ATP synthesis coupled proton tra
base-excision repair [GO:0006284]
base-excision repair [GO:0006284]; DNA dealkylation involved in DNA repair [GO:0006307]
base-excision repair [GO:0006284]; mismatch repair [GO:0006298]
base-excision repair, DNA ligation [GO:0006288]; DNA replication [GO:0006260]
biosynthetic process [GO:0009058]
biosynthetic process [GO:0009058]
biosynthetic process [GO:0009058]
branched-chain amino acid biosynthetic process [GO:0009082]; leucine biosynthetic process [GO:0009098]
branched-chain amino acid biosynthetic process [GO:0009082]; leucine biosynthetic process [GO:0009098]
branched-chain amino acid biosynthetic process [GO:0009082]; leucine biosynthetic process [GO:0009098]
branched-chain amino acid biosynthetic process [GO:0009082]; leucine biosynthetic process [GO:0009098]
butanediol metabolic process [GO:0034077]; isoleucine biosynthetic process [GO:0009097]; menaquinone b
CAAX-box protein processing [GO:0071586]
canonical glycolysis [GO:0061621]; fructose 1,6-bisphosphate metabolic process [GO:0030388]; fructose 6-
carbohydrate catabolic process [GO:0016052]; lactose catabolic process via tagatose-6-phosphate [GO:001
carbohydrate metabolic process [GO:0005975]
carbohydrate metabolic process [GO:0005975]; cell cycle [GO:0007049]; cell division [GO:0051301]; cell wa
carbohydrate metabolic process [GO:0005975]; glycerol-3-phosphate biosynthetic process [GO:0046167]; g
cell adhesion [GO:0007155]
cell adhesion [GO:0007155]
cell adhesion [GO:0007155]; pathogenesis [GO:0009405]
cell adhesion [GO:0007155]; zinc ion transport [GO:0006829]

cell adhesion involved in biofilm formation [GO:0043708] icaA
cell cycle [GO:0007049]; cell division [GO:0051301]
cell cycle [GO:0007049]; cell division [GO:0051301]; cell wall organization [GO:0071555]; peptidoglycan bic
cell cycle [GO:0007049]; cell division [GO:0051301]; cell wall organization [GO:0071555]; peptidoglycan bic
cell cycle [GO:0007049]; cell division [GO:0051301]; cell wall organization [GO:0071555]; peptidoglycan bic
cell division [GO:0051301]
cell division [GO:0051301]; division septum assembly [GO:0000917]; FtsZ-dependent cytokinesis [GO:004
cell division [GO:0051301]; FtsZ-dependent cytokinesis [GO:0043093]
cell division [GO:0051301]; protein catabolic process [GO:0030163]; protein folding [GO:0006457]; proteoly
cell morphogenesis [GO:0000902]; cell wall organization [GO:0071555]; lipid A biosynthetic process [GO:00
cell redox homeostasis [GO:0045454]
cell redox homeostasis [GO:0045454]
cell redox homeostasis [GO:0045454]
cell wall organization [GO:0071555]
cell wall organization [GO:0071555]
cell wall organization [GO:0071555]; cytolysis [GO:0019835]; defense response to bacterium [GO:0042742]
cell wall organization [GO:0071555]; cytolysis [GO:0019835]; defense response to bacterium [GO:0042742]
cell wall organization [GO:0071555]; cytolysis [GO:0019835]; defense response to bacterium [GO:0042742]
cell wall organization [GO:0071555]; dephosphorylation [GO:0016311]; peptidoglycan biosynthetic process [
cell wall organization [GO:0071555]; pathogenesis [GO:0009405]; peptide catabolic process [GO:0043171]
cell wall organization [GO:0071555]; peptidoglycan biosynthetic process [GO:0009252]; regulation of cell sh
cell wall organization [GO:0071555]; peptidoglycan biosynthetic process [GO:0009252]; regulation of cell sh
cell wall organization [GO:0071555]; peptidoglycan biosynthetic process [GO:0009252]; regulation of cell sh
cell wall organization [GO:0071555]; peptidoglycan biosynthetic process [GO:0009252]; regulation of cell sh
cell wall organization [GO:0071555]; peptidoglycan biosynthetic process [GO:0009252]; regulation of cell sh
cell wall organization [GO:0071555]; peptidoglycan biosynthetic process [GO:0009252]; regulation of cell sh
cell wall organization [GO:0071555]; peptidoglycan biosynthetic process [GO:0009252]; regulation of cell sh
cell wall organization [GO:0071555]; peptidoglycan biosynthetic process [GO:0009252]; regulation of cell sh
cell wall organization [GO:0071555]; peptidoglycan catabolic process [GO:0009253]
cellular amino acid metabolic process [GO:0006520]; pyridoxal phosphate biosynthetic process [GO:004282
cellular phosphate ion homeostasis [GO:0030643]; negative regulation of phosphate metabolic process [GC
cellular respiration [GO:0045333]; heme biosynthetic process [GO:0006783]; heme O biosynthetic process [
cellular response to heat [GO:0034605]; protein refolding [GO:0042026]
cellular response to osmotic stress [GO:0071470]
chaperone cofactor-dependent protein refolding [GO:0051085]
chaperone cofactor-dependent protein refolding [GO:0051085]; DNA replication [GO:0006260]; protein refol
chaperone cofactor-dependent protein refolding [GO:0051085]; protein folding [GO:0006457]; protein refold
chromosome segregation [GO:0007059]
chromosome segregation [GO:0007059]; division septum assembly [GO:0000917]; positive regulation of sp
chromosome segregation [GO:0007059]; DNA topological change [GO:0006265]
chromosome segregation [GO:0007059]; DNA topological change [GO:0006265]; response to antibiotic [GC
chromosome segregation [GO:0007059]; positive regulation of sporulation resulting in formation of a cellula
chromosome segregation [GO:0007059]; positive regulation of sporulation resulting in formation of a cellula
coenzyme A biosynthetic process [GO:0015937]
co-translational protein modification [GO:0043686]; N-terminal protein amino acid modification [GO:003136
CTP salvage [GO:0044211]; UMP salvage [GO:0044206]
cyclic threonylcarbamoyladenosine biosynthetic process [GO:0061504]
cysteine biosynthetic process via cystathionine [GO:0019343]; transsulfuration [GO:0019346]
cytolysis [GO:0019835]
cytolysis [GO:0019835]; programmed cell death [GO:0012501]
dATP catabolic process [GO:0046061]; dGTP catabolic process [GO:0006203]; dTTP catabolic process [GC
'de novo' AMP biosynthetic process [GO:0044208]; IMP metabolic process [GO:0046040]
'de novo' cotranslational protein folding [GO:0051083]; cell cycle [GO:0007049]; cell division [GO:0051301]
'de novo' CTP biosynthetic process [GO:0044210]; CTP biosynthetic process [GO:0006241]; glutamine met

'de novo' IMP biosynthetic process [GO:0006189]
'de novo' IMP biosynthetic process [GO:0006189]
'de novo' IMP biosynthetic process [GO:0006189]; cobalamin biosynthetic process [GO:0009236]
'de novo' IMP biosynthetic process [GO:0006189]; purine nucleobase biosynthetic process [GO:0009113]
'de novo' L-methionine biosynthetic process [GO:0071266]; diaminopimelate biosynthetic process [GO:001
diaminopimelate biosynthetic process [GO:0019877]; lysine biosynthetic process via diaminopimelate [GO:
diaminopimelate biosynthetic process [GO:0019877]; lysine biosynthetic process via diaminopimelate [GO:
division septum assembly [GO:0000917]
DNA biosynthetic process [GO:0071897]; thymidine metabolic process [GO:0046104]
DNA dealkylation involved in DNA repair [GO:0006307]; methylation [GO:0032259]
DNA duplex unwinding [GO:0032508]; DNA recombination [GO:0006310]; DNA repair [GO:0006281]; DNA
DNA integration [GO:0015074]; transposition [GO:0032196]
DNA packaging [GO:0006323]
DNA recombination [GO:0006310]; DNA repair [GO:0006281]
DNA recombination [GO:0006310]; DNA repair [GO:0006281]; DNA replication [GO:0006260]; positive regu
DNA recombination [GO:0006310]; DNA repair [GO:0006281]; SOS response [GO:0009432]
DNA recombination [GO:0006310]; DNA repair [GO:0006281]; SOS response [GO:0009432]
DNA repair [GO:0006281]
DNA repair [GO:0006281]
DNA replication [GO:0006260]; DNA synthesis involved in DNA repair [GO:0000731]; double-strand break r
DNA replication proofreading [GO:0045004]
DNA replication, Okazaki fragment processing [GO:0033567]
DNA replication, removal of RNA primer [GO:0043137]; mismatch repair [GO:0006298]
DNA replication, synthesis of RNA primer [GO:0006269]; DNA unwinding involved in DNA replication [GO:
DNA topological change [GO:0006265]
DNA topological change [GO:0006265]
DNA unwinding involved in DNA replication [GO:0006268]; recombinational repair [GO:0000725]
DNA-dependent DNA replication [GO:0006261]
DNA-dependent DNA replication [GO:0006261]
DNA-dependent DNA replication [GO:0006261]; DNA topological change [GO:0006265]
DNA-dependent DNA replication [GO:0006261]; DNA topological change [GO:0006265]; response to antibic
DNA-templated transcription, elongation [GO:0006354]; regulation of DNA-templated transcription, elongati
DNA-templated transcription, initiation [GO:0006352]
DNA-templated transcription, termination [GO:0006353]
D-tagatose 6-phosphate catabolic process [GO:2001059]; lactose metabolic process [GO:0005988]
dTDP biosynthetic process [GO:0006233]; dTTP biosynthetic process [GO:0006235]; dUDP biosynthetic pro
enzyme-directed rRNA pseudouridine synthesis [GO:0000455]
error-prone translesion synthesis [GO:0042276]; SOS response [GO:0009432]
establishment of competence for transformation [GO:0030420]
FAD biosynthetic process [GO:0006747]; FMN biosynthetic process [GO:0009398]; riboflavin biosynthetic p
fatty acid beta-oxidation [GO:0006635]; phenylacetate catabolic process [GO:0010124]
fatty acid biosynthetic process [GO:0006633]
fatty acid biosynthetic process [GO:0006633]
fatty acid biosynthetic process [GO:0006633]
fatty acid biosynthetic process [GO:0006633]; lipid A biosynthetic process [GO:0009245]
fatty acid biosynthetic process [GO:0006633]; negative regulation of fatty acid biosynthetic process [GO:00
fatty acid biosynthetic process [GO:0006633]; phospholipid biosynthetic process [GO:0008654]
fatty acid elongation [GO:0030497]
folic acid biosynthetic process [GO:0046656]; tetrahydrofolate biosynthetic process [GO:0046654]
folic acid catabolic process [GO:0046657]
folic acid-containing compound biosynthetic process [GO:0009396]
formate metabolic process [GO:0015942]
formate transport [GO:0015724]
galactose catabolic process via UDP-galactose [GO:0033499]; glucose metabolic process [GO:0006006]

glucose metabolic process [GO:0006006]
glucose metabolic process [GO:0006006]
glucose-6-phosphate transport [GO:0015760]; phosphate ion transmembrane transport [GO:0035435]
glutamine biosynthetic process [GO:0006542]
glutamine catabolic process [GO:0006543]; pyridoxal phosphate biosynthetic process [GO:0042823]; pyrido
glutamine metabolic process [GO:0006541]; GMP biosynthetic process [GO:0006177]
glutamine metabolic process [GO:0006541]; histidine biosynthetic process [GO:0000105]
glycerol metabolic process [GO:0006071]
glycine betaine biosynthetic process from choline [GO:0019285]
glycine betaine biosynthetic process from choline [GO:0019285]
glycine decarboxylation via glycine cleavage system [GO:0019464]
glycolytic process [GO:0006096]
glycyl-tRNA aminoacylation [GO:0006426]
GMP biosynthetic process [GO:0006177]; GTP biosynthetic process [GO:0006183]
guanosine tetraphosphate biosynthetic process [GO:0015970]
histidine biosynthetic process [GO:0000105]
histidine biosynthetic process [GO:0000105]
histidine biosynthetic process [GO:0000105]
histidine biosynthetic process [GO:0000105]
histidine biosynthetic process [GO:0000105]; tryptophan biosynthetic process [GO:0000162]
homoserine biosynthetic process [GO:0009090]; lysine biosynthetic process via diaminopimelate [GO:0009
homoserine metabolic process [GO:0009092]; methionine biosynthetic process [GO:0009086]
intracellular protein transmembrane transport [GO:0065002]; protein secretion [GO:0009306]; protein transp
iron ion transport [GO:0006826]
iron-sulfur cluster assembly [GO:0016226]
iron-sulfur cluster assembly [GO:0016226]
isoleucine biosynthetic process [GO:0009097]; leucine biosynthetic process [GO:0009098]; valine biosynthe
isoleucine biosynthetic process [GO:0009097]; methionine biosynthetic process [GO:0009086]; threonine bi
isoleucine biosynthetic process [GO:0009097]; valine biosynthetic process [GO:0009099]
isoleucine biosynthetic process [GO:0009097]; valine biosynthetic process [GO:0009099]
isoleucine transport [GO:0015818]; leucine transport [GO:0015820]; valine transport [GO:0015829]
isoprenoid biosynthetic process [GO:0008299]
leucyl-tRNA aminoacylation [GO:0006429]
L-glutamate transmembrane transport [GO:0015813]
lipid A biosynthetic process [GO:0009245]
lipoteichoic acid biosynthetic process [GO:0070395]
L-methionine salvage from methylthioadenosine [GO:0019509]; L-methionine salvage from S-adenosylmeth
L-serine catabolic process [GO:0006565]; L-threonine catabolic process to propionate [GO:0070689]; threon
L-valine transmembrane transport [GO:1903785]
lysyl-tRNA aminoacylation [GO:0006430]
maturation of SSU-rRNA from tricistronic rRNA transcript (SSU-rRNA, 5.8S rRNA, LSU-rRNA) [GO:000046
mature ribosome assembly [GO:0042256]; negative regulation of ribosome biogenesis [GO:0090071]; nega
menaquinone biosynthetic process [GO:0009234]
menaquinone biosynthetic process [GO:0009234]
menaquinone biosynthetic process [GO:0009234]; ubiquinone biosynthetic process via 3,4-dihydroxy-5-poly
metabolic process [GO:0008152]
metabolite repair [GO:0110051]; nicotinamide nucleotide metabolic process [GO:0046496]
metal ion transport [GO:0030001]
methionyl-tRNA aminoacylation [GO:0006431]
methylation [GO:0032259]; siroheme biosynthetic process [GO:0019354]
methylation [GO:0032259]; tRNA processing [GO:0008033]
Mo-molybdopterin cofactor biosynthetic process [GO:0006777]
Mo-molybdopterin cofactor biosynthetic process [GO:0006777]
Mo-molybdopterin cofactor biosynthetic process [GO:0006777]

Mo-molybdopterin cofactor biosynthetic process [GO:0006777]
mRNA catabolic process [GO:0006402]; pathogenesis [GO:0009405]
mRNA catabolic process [GO:0006402]; RNA catabolic process [GO:0006401]; RNA processing [GO:00063
mRNA processing [GO:0006397]; rRNA processing [GO:0006364]
mRNA pseudouridine synthesis [GO:1990481]; tRNA modification [GO:0006400]; tRNA pseudouridine synth
NAD biosynthetic process [GO:0009435]
negative regulation of transcription, DNA-templated [GO:0045892]
negative regulation of transcription, DNA-templated [GO:0045892]
negative regulation of transcription, DNA-templated [GO:0045892]
negative regulation of transcription, DNA-templated [GO:0045892]; response to redox state [GO:0051775]
nickel cation transport [GO:0015675]; phosphate ion transmembrane transport [GO:0035435]
nickel cation transport [GO:0015675]; phosphate ion transport [GO:0006817]
nickel cation transport [GO:0015675]; transmembrane transport [GO:0055085]
nickel cation transport [GO:0015675]; transmembrane transport [GO:0055085]
nitrate assimilation [GO:0042128]
nitrogen compound transport [GO:0071705]
nitrogen compound transport [GO:0071705]
nitrogen utilization [GO:0019740]
N-terminal peptidyl-serine acetylation [GO:0017198]
nucleobase-containing small molecule interconversion [GO:0015949]
nucleobase-containing small molecule interconversion [GO:0015949]
oligopeptide transport [GO:0006857]
one-carbon metabolic process [GO:0006730]; S-adenosylmethionine biosynthetic process [GO:0006556]
organic substance metabolic process [GO:0071704]
organic substance metabolic process [GO:0071704]
osmosensory signaling via phosphorelay pathway [GO:0007234]
osmosensory signaling via phosphorelay pathway [GO:0007234]
p-aminobenzoyl-glutamate transmembrane transport [GO:1902604]; p-aminobenzoyl-glutamate transport [G
pantothenate biosynthetic process [GO:0015940]
pantothenate biosynthetic process [GO:0015940]
pantothenate biosynthetic process [GO:0015940]
pantothenate biosynthetic process [GO:0015940]
pathogenesis [GO:0009405]
pathogenesis [GO:0009405]
pathogenesis [GO:0009405]
pathogenesis [GO:0009405]
pathogenesis [GO:0009405]
pathogenesis [GO:0009405]
pathogenesis [GO:0009405]
pathogenesis [GO:0009405]
pathogenesis [GO:0009405]
pathogenesis [GO:0009405]
pathogenesis [GO:0009405]
pathogenesis [GO:0009405]
pathogenesis [GO:0009405]
pathogenesis [GO:0009405]
pathogenesis [GO:0009405]; regulation of transcription, DNA-templated [GO:0006355]
pathogenesis [GO:0009405]; regulation of transcription, DNA-templated [GO:0006355]
pathogenesis [GO:0009405]; regulation of transcription, DNA-templated [GO:0006355]
pathogenesis [GO:0009405]; response to antibiotic [GO:0046677]
pathogenesis [GO:0009405]; single-species submerged biofilm formation [GO:0090609]
peptide transport [GO:0015833]
peptide transport [GO:0015833]
peptide transport [GO:0015833]

peptide transport [GO:0015833]
peptide transport [GO:0015833]; transmembrane transport [GO:0055085]
peptidoglycan biosynthetic process [GO:0009252]; response to antibiotic [GO:0046677]
peptidyl-glutamine methylation [GO:0018364]
phenylalanyl-tRNA aminoacylation [GO:0006432]
phosphate ion transmembrane transport [GO:0035435]
phosphate ion transport [GO:0006817]
phosphocreatine biosynthetic process [GO:0046314]; stress response to cadmium ion [GO:1990170]; stress
phosphoenolpyruvate-dependent sugar phosphotransferase system [GO:0009401]
phosphoenolpyruvate-dependent sugar phosphotransferase system [GO:0009401]
phosphoenolpyruvate-dependent sugar phosphotransferase system [GO:0009401]
phosphoenolpyruvate-dependent sugar phosphotransferase system [GO:0009401]
phosphoenolpyruvate-dependent sugar phosphotransferase system [GO:0009401]
phosphoenolpyruvate-dependent sugar phosphotransferase system [GO:0009401]
phosphoenolpyruvate-dependent sugar phosphotransferase system [GO:0009401]
phosphoenolpyruvate-dependent sugar phosphotransferase system [GO:0009401]
phosphoenolpyruvate-dependent sugar phosphotransferase system [GO:0009401]
phospholipid biosynthetic process [GO:0008654]
phospholipid biosynthetic process [GO:0008654]
phosphorelay signal transduction system [GO:0000160]; regulation of transcription, DNA-templated [GO:00(
phosphorylation [GO:0016310]
plasma membrane ATP synthesis coupled proton transport [GO:0042777]
polyamine transport [GO:0015846]
polyphosphate catabolic process [GO:0006798]
porphyrin-containing compound biosynthetic process [GO:0006779]; protoporphyrinogen IX biosynthetic pro
potassium ion transmembrane transport [GO:0071805]; regulation of intracellular pH [GO:0051453]; sodium
potassium ion transport [GO:0006813]
proline transmembrane transport [GO:0035524]; proline transport [GO:0015824]
protein catabolic process [GO:0030163]; proteolysis [GO:0006508]
protein folding [GO:0006457]
protein folding [GO:0006457]
protein glycosylation [GO:0006486]; regulation of protein stability [GO:0031647]
protein insertion into membrane [GO:0051205]; protein transport [GO:0015031]
protein lipoylation [GO:0009249]
protein O-linked glycosylation via serine [GO:0018242]
protein peptidyl-prolyl isomerization [GO:0000413]
protein repair [GO:0030091]; response to nitrosative stress [GO:0051409]; response to oxidative stress [GO
protein targeting [GO:0006605]; SRP-dependent cotranslational protein targeting to membrane [GO:000661
proteolysis [GO:0006508]
proteolysis [GO:0006508]
proton transmembrane transport [GO:1902600]; sodium ion transport [GO:0006814]
purine nucleotide metabolic process [GO:0006163]
purine ribonucleoside salvage [GO:0006166]; xanthine metabolic process [GO:0046110]; XMP salvage [GC
pyrimidine deoxyribonucleotide catabolic process [GO:0009223]
queuosine biosynthetic process [GO:0008616]
queuosine biosynthetic process [GO:0008616]
queuosine biosynthetic process [GO:0008616]
queuosine biosynthetic process [GO:0008616]
queuosine biosynthetic process [GO:0008616]; tRNA processing [GO:0008033]
queuosine biosynthetic process [GO:0008616]; tRNA processing [GO:0008033]
queuosine biosynthetic process [GO:0008616]; tRNA wobble guanine modification [GO:0002099]
queuosine biosynthetic process [GO:0008616]; tRNA-guanine transglycosylation [GO:0101030]; tRNA wobb
queuosine salvage [GO:1990397]
regulation of DNA-templated transcription [GO:0006355]

| | |
|---|---|
| regulation of transcription, DNA-templated [GO:0006355] | |
| regulation of transcription, DNA-templated [GO:0006355] | |
| regulation of transcription, DNA-templated [GO:0006355] | |
| regulation of transcription, DNA-templated [GO:0006355] | |
| regulation of transcription, DNA-templated [GO:0006355] | |
| regulation of transcription, DNA-templated [GO:0006355] | |
| regulation of transcription, DNA-templated [GO:0006355] | |
| regulation of transcription, DNA-templated [GO:0006355] | |
| regulation of transcription, DNA-templated [GO:0006355] | |
| regulation of transcription, DNA-templated [GO:0006355] | icaR |
| regulation of transcription, DNA-templated [GO:0006355] | |
| regulation of transcription, DNA-templated [GO:0006355] | |
| regulation of transcription, DNA-templated [GO:0006355]; stress response to cadmium ion [GO:1990170]; s | |
| regulation of transcription, DNA-templated [GO:0006355]; transcription, DNA-templated [GO:0006351] | |
| respiratory electron transport chain [GO:0022904] | |
| response to antibiotic [GO:0046677]; rRNA base methylation [GO:0070475]; tRNA methylation [GO:003048 | |
| response to antibiotic [GO:0046677]; transmembrane transport [GO:0055085] | |
| response to arsenic-containing substance [GO:0046685] | |
| response to arsenic-containing substance [GO:0046685] | |
| response to cold [GO:0009409]; ribosomal large subunit assembly [GO:0000027]; RNA catabolic process [G | |
| response to oxidative stress [GO:0006979] | |
| response to oxidative stress [GO:0006979] | |
| riboflavin biosynthetic process [GO:0009231] | |
| riboflavin biosynthetic process [GO:0009231] | |
| riboflavin biosynthetic process [GO:0009231] | |
| riboflavin transport [GO:0032218] | |
| ribosomal large subunit assembly [GO:0000027] | |
| ribosomal large subunit assembly [GO:0000027] | |
| ribosomal large subunit assembly [GO:0000027]; RNA catabolic process [GO:0006401]; RNA secondary str | |
| ribosomal small subunit assembly [GO:0000028]; translation [GO:0006412] | |
| ribosomal small subunit biogenesis [GO:0042274] | |
| ribosomal small subunit biogenesis [GO:0042274] | |
| ribosome biogenesis [GO:0042254] | |
| RNA processing [GO:0006396] | |
| rRNA base methylation [GO:0070475] | |
| rRNA base methylation [GO:0070475] | |
| rRNA methylation [GO:0031167] | |
| rRNA processing [GO:0006364] | |
| selenocysteine biosynthetic process [GO:0016260]; selenocysteinyl-tRNA(Sec) biosynthetic process [GO:00 | |
| siderophore-dependent iron import into cell [GO:0033214] | |
| siderophore-dependent iron import into cell [GO:0033214] | |
| siderophore-dependent iron import into cell [GO:0033214] | |
| siderophore-dependent iron import into cell [GO:0033214] | |
| signal peptide processing [GO:0006465] | |
| sodium-dependent phosphate transport [GO:0044341] | |
| SRP-dependent cotranslational protein targeting to membrane [GO:0006614] | |
| stress response to cadmium ion [GO:1990170]; stress response to copper ion [GO:1990169] | |
| succinate transport [GO:0015744] | |
| succinate transport [GO:0015744] | |
| thiamine biosynthetic process [GO:0009228] | |
| thiamine biosynthetic process [GO:0009228]; thiamine diphosphate biosynthetic process [GO:0009229]; thia | |
| threonyl-tRNA aminoacylation [GO:0006435] | |
| transcription initiation from bacterial-type RNA polymerase promoter [GO:0001123] | |
| transcription, DNA-templated [GO:0006351] | |

transcription, DNA-templated [GO:0006351]
transcription, DNA-templated [GO:0006351]
translation [GO:0006412]
translation [GO:0006412]
translation [GO:0006412]
translation [GO:0006412]
translation [GO:0006412]
translation [GO:0006412]
translation [GO:0006412]
translation [GO:0006412]
translation [GO:0006412]
translation [GO:0006412]
translational elongation [GO:0006414]
transmembrane transport [GO:0055085]
transmembrane transport [GO:0055085]
transmembrane transport [GO:0055085]
transmembrane transport [GO:0055085]
transmembrane transport [GO:0055085]
transmembrane transport [GO:0055085]
transmembrane transport [GO:0055085]
transmembrane transport [GO:0055085]
transmembrane transport [GO:0055085]
trans-translation [GO:0070929]; trans-translation-dependent protein tagging [GO:0070930]
tricarboxylic acid cycle [GO:0006099]
tricarboxylic acid cycle [GO:0006099]
tRNA aminoacylation [GO:0043039]; tyrosyl-tRNA aminoacylation [GO:0006437]
tRNA methylation [GO:0030488]; tRNA wobble uridine modification [GO:0002098]
tRNA methylation [GO:0030488]; tRNA wobble uridine modification [GO:0002098]
tRNA methylthiolation [GO:0035600]
tRNA methylthiolation [GO:0035600]
tRNA modification [GO:0006400]
tRNA modification [GO:0006400]
tRNA pseudouridine synthesis [GO:0031119]
tryptophan biosynthetic process [GO:0000162]
tryptophanyl-tRNA aminoacylation [GO:0006436]
UDP-N-acetylglucosamine biosynthetic process [GO:0006048]
valyl-tRNA aminoacylation [GO:0006438]
wobble position cytosine ribose methylation [GO:0002131]; wobble position uridine ribose methylation [GO:(
xanthine transport [GO:0042906]
xenobiotic detoxification by transmembrane export across the plasma membrane [GO:1990961]
zinc ion transmembrane transport [GO:0071577]
#N/A
#N/A
#N/A
#N/A
#N/A
#N/A
#N/A
#N/A
#N/A
#N/A
#N/A
#N/A
#N/A

#N/A
#N/A
#N/A
#N/A
#N/A
#N/A
#N/A
#N/A
#N/A
#N/A

'ay [GO:0010142]

ectron transport coupled proton transport [GO:0015990]
ositive regulation of membrane potential [GO:0045838]

5675]; transmembrane transport [GO:0055085]

O:0008652]; chorismate biosynthetic process [GO:0009423]
O:0008652]; chorismate biosynthetic process [GO:0009423]
O:0008652]; chorismate biosynthetic process [GO:0009423]; shikimate metabolic process [GO:0019632]

ansport [GO:0042777]

iosynthetic process [GO:0009234]; valine biosynthetic process [GO:0009099]

phosphate metabolic process [GO:0006002]
9512]

all organization [GO:0071555]; lipid glycosylation [GO:0030259]; peptidoglycan biosynthetic process [GO:0009252
lycerol-3-phosphate catabolic process [GO:0046168]; glycerol-3-phosphate metabolic process [GO:0006072]; glyc

›synthetic process [GO:0009252]; regulation of cell shape [GO:0008360]
›synthetic process [GO:0009252]; regulation of cell shape [GO:0008360]
›synthetic process [GO:0009252]; regulation of cell shape [GO:0008360]; UDP-N-acetylgalactosamine biosynthetic

3093]; protein polymerization [GO:0051258]

sis [GO:0006508]
009245]; peptidoglycan biosynthetic process [GO:0009252]; regulation of cell shape [GO:0008360]; UDP-N-acetylg

; division septum assembly [GO:0000917]; metabolic process [GO:0008152]; pathogenesis [GO:0009405]
; metabolic process [GO:0008152]
; metabolic process [GO:0008152]
[GO:0009252]; regulation of cell shape [GO:0008360]; response to antibiotic [GO:0046677]

ape [GO:0008360]
ape [GO:0008360]
ape [GO:0008360]
ape [GO:0008360]
ape [GO:0008360]; response to antibiotic [GO:0046677]
ape [GO:0008360]; response to antibiotic [GO:0046677]
ape [GO:0008360]; response to antibiotic [GO:0046677]
ape [GO:0008360]; response to antibiotic [GO:0046677]

23]; pyridoxine biosynthetic process [GO:0008615]
):0045936]; phosphate ion transport [GO:0006817]
[GO:0048034]

lding [GO:0042026]; response to heat [GO:0009408]
ing [GO:0042026]; response to heat [GO:0009408]

orulation resulting in formation of a cellular spore [GO:0045881]

):0046677]
r spore [GO:0045881]
r spore [GO:0045881]; siderophore biosynthetic process [GO:0019290]

5]; peptidyl-methionine modification [GO:0018206]; translation [GO:0006412]

):0046076]; dUTP catabolic process [GO:0046081]; TTP catabolic process [GO:0046047]; UTP catabolic process

; chaperone-mediated protein folding [GO:0061077]; protein transport [GO:0015031]; protein unfolding [GO:00433
abolic process [GO:0006541]; pyrimidine nucleobase biosynthetic process [GO:0019856]

)877]; isoleucine biosynthetic process [GO:0009097]; lysine biosynthetic process via diaminopimelate [GO:000908
)009089]
)009089]

replication [GO:0006260]; SOS response [GO:0009432]

ulation of helicase activity [GO:0051096]

epair [GO:0006302]; SOS response [GO:0009432]

)006268]

otic [GO:0046677]
on [GO:0032784]

ocess [GO:0006227]

rocess [GO:0009231]; riboflavin metabolic process [GO:0006771]

45717]; negative regulation of transcription, DNA-templated [GO:0045892] xine metabolic process [GO:0008614]

)89]; threonine biosynthetic process [GO:0009088]

ort by the Sec complex [GO:0043952]

tic process [GO:0009099]
osynthetic process [GO:0009088]

nionine [GO:0019284]; nucleoside catabolic process [GO:0009164]
nine catabolic process [GO:0006567]

2]; ribosomal small subunit assembly [GO:0000028]; translation [GO:0006412]
tive regulation of translation [GO:0017148]

prenylbenzoate [GO:0032194]; vitamin K biosynthetic process [GO:0042371]

396]

nesis [GO:0031119]

GO:0015814]

; response to copper ion [GO:1990169]

)6355]

cess [GO:0006782]
ı ion import across plasma membrane [GO:0098719]

):0006979]
4]

):0032265]

ɔle guanine modification [GO:0002099]

tress response to copper ion [GO:1990169]

8]

3O:0006401]

ucture unwinding [GO:0010501]

)97056]; seryl-tRNA aminoacylation [GO:0006434]

azole biosynthetic process [GO:0052837]; tRNA 4-thiouridine biosynthesis [GO:0002937]

)002132]

2]; regulation of cell shape [GO:0008360]
cerophospholipid metabolic process [GO:0006650]; phospholipid biosynthetic process [GO:0008654]

c process [GO:0019277]

glucosamine biosynthetic process [GO:0006048]

[GO:0046052]

35]

39]; threonine biosynthetic process [GO:0009088]

| gene | UNIPROTID |
|---|---|
| ureC | #N/A |
| ureB | #N/A |
| murQ | #N/A |
| ureA | #N/A |
| SAOUHSC_02562 | #N/A |
| SAOUHSC_R0005 | #N/A |
| adhA | #N/A |
| SAOUHSC_01761a | #N/A |
| SAOUHSC_A01081 | #N/A |
| SAOUHSC_1307a | #N/A |
| SAOUHSC_R0004 | #N/A |
| SAOUHSC_R0003 | #N/A |
| SAOUHSC_00133 | Q2G1I9 |
| SAOUHSC_02653 | Q2FVP9 |
| SAOUHSC_01280 | Q2FYZ2 |
| SAOUHSC_02647 | Q2FVQ5 |
| SAOUHSC_01611 | Q2FY54 |
| SAOUHSC_00378 | Q2G0Y3 |
| SAOUHSC_02813 | Q2G2L5 |
| SAOUHSC_02151 | Q2FWX0 |
| SAOUHSC_00647 | Q2G2E5 |
| SAOUHSC_01378 | Q2FYQ7 |
| SAOUHSC_02103 | Q2FX03 |
| SAOUHSC_02718 | Q2FVJ2 |
| SAOUHSC_01377 | Q2FYQ8 |
| SAOUHSC_02001 | Q2G2N1 |
| SAOUHSC_02420 | Q2FW83 |
| SAOUHSC_03037 | Q2FUR5 |
| SAOUHSC_02953 | Q2FUZ2 |
| SAOUHSC_00099 | Q2G226 |
| SAOUHSC_00105 | Q2G1L7 |
| SAOUHSC_01493 | Q2FYF9 |
| SAOUHSC_00535 | Q2G0M5 |
| SAOUHSC_00186 | Q2G1D9 |
| SAOUHSC_02331 | Q2FWG0 |
| SAOUHSC_02329 | Q2FWG2 |
| SAOUHSC_01845 | Q2G296 |
| SAOUHSC_00037 | Q2G1R5 |
| SAOUHSC_00758 | Q2G063 |
| SAOUHSC_00759 | Q2G062 |
| SAOUHSC_00309 | Q2G146 |
| SAOUHSC_00294 | Q2G161 |
| SAOUHSC_01764 | Q2FXS1 |
| SAOUHSC_01087 | Q2FZE4 |
| SAOUHSC_01086 | Q2FZE5 |
| SAOUHSC_00072 | Q2G1N5 |
| SAOUHSC_00071 | Q2G1N6 |
| SAOUHSC_02436 | Q2FW70 |
| SAOUHSC_02434 | Q2FW72 |
| SAOUHSC_00636 | Q2G2D9 |
| SAOUHSC_00065 | Q2G1P0 |
| SAOUHSC_00364 | O05204 |
| SAOUHSC_00831 | Q2G1T3 |

| | |
|---|---|
| SAOUHSC_02609 | Q2FVT3 |
| SAOUHSC_00315 | Q2G140 |
| SAOUHSC_02635 | Q2G2I1 |
| SAOUHSC_01602 | Q2FY63 |
| SAOUHSC_02608 | Q2FVT4 |
| SAOUHSC_02810 | Q2G2L8 |
| SAOUHSC_00465 | Q2G0S9 |
| SAOUHSC_02852 | Q2FV83 |
| SAOUHSC_00231 | P60611 |
| SAOUHSC_00434 | Q2G0U3 |
| SAOUHSC_00242 | Q2G1A3 |
| SAOUHSC_00913 | Q2FZS7 |
| SAOUHSC_02269 | Q2FWM0 |
| SAOUHSC_02315 | Q2FWH6 |
| SAOUHSC_01800 | Q2FXN6 |
| SAOUHSC_02011 | Q2G2G9 |
| SAOUHSC_02377 | Q2FWC1 |
| SAOUHSC_00211 | Q2G217 |
| SAOUHSC_01936 | Q2FXC7 |
| SAOUHSC_01938 | Q2FXC5 |
| SAOUHSC_01935 | Q2FXC8 |
| SAOUHSC_01942 | Q2FXC2 |
| SAOUHSC_01941 | Q2FXC3 |
| SAOUHSC_01939 | Q2FXC4 |
| SAOUHSC_01949 | Q2FXB6 |
| SAOUHSC_01763 | P31337 |
| SAOUHSC_00879 | Q2FZV6 |
| SAOUHSC_01838 | Q2FXJ6 |
| SAOUHSC_01912 | Q2G2V6 |
| SAOUHSC_02971 | Q2FUX4 |
| SAOUHSC_01861 | Q2FXI5 |
| SAOUHSC_00958 | Q2FZP2 |
| SAOUHSC_01226 | Q2FZ28 |
| SAOUHSC_02491 | O08387 |
| SAOUHSC_00963 | Q2FZN7 |
| SAOUHSC_02102 | Q2FX05 |
| SAOUHSC_01972 | Q2G2S6 |
| SAOUHSC_02356 | Q2FWE3 |
| SAOUHSC_01473 | Q2G258 |
| SAOUHSC_02314 | Q2FWH7 |
| SAOUHSC_01799 | Q2FXN7 |
| SAOUHSC_01884 | Q2FXG3 |
| SAOUHSC_00633 | Q2G2E0 |
| SAOUHSC_02312 | Q2FWH9 |
| SAOUHSC_02908 | Q2FV31 |
| SAOUHSC_02264 | Q2FWM5 |
| SAOUHSC_00184 | Q2G1E1 |
| SAOUHSC_00736 | Q2G084 |
| SAOUHSC_02974 | Q2FUX1 |
| SAOUHSC_00313 | Q2G142 |
| SAOUHSC_02401 | Q2FW98 |
| SAOUHSC_02841 | Q2FV93 |
| SAOUHSC_02975 | Q2FUX0 |
| SAOUHSC_00310 | Q2G145 |

| | | | |
|---|---|---|---|
| SAOUHSC_00311 | | | Q2G144 |
| SAOUHSC_00209 | | | Q2G219 |
| SAOUHSC_01430 | | | Q2FYL0 |
| SAOUHSC_02754 | | | Q2FVG0 |
| SAOUHSC_01840 | | | Q2FXJ4 |
| SAOUHSC_00170 | | | Q2G1F5 |
| SAOUHSC_01951 | | | Q2FXB4 |
| SAOUHSC_01383 | | | Q2FYQ2 |
| SAOUHSC_00937 | | | Q2G1U3 |
| SAOUHSC_00757 | | | Q2G064 |
| SAOUHSC_02566 | sarR | | Q9F0R1 |
| SAOUHSC_01879 | rot | | Q9RFJ6 |
| SAOUHSC_00674 | sarX | | Q2G0D1 |
| SAOUHSC_02261 | agrB | | Q2FWM7 |
| SAOUHSC_01419 | arlS | | Q9KJN3 |
| SAOUHSC_00987 | cysteine protease | sspB | Q2FZL3 |
| SAOUHSC_00986 | cysteine protease | sspC | Q9EYW6 |
| SAOUHSC_00988 | glutamyl endopeptidase | sspA | Q2FZL2 |
| SAOUHSC_01136 | Phenol-soluble modulins | psmβ2 | Q2FZA3 |
| SAOUHSC_01135 | Phenol-soluble modulins | psmβ1 | Q2FZA4 |
| SAOUHSC_02127 | staphopain thiol proteina | scpA | Q2G2R8 |
| SAOUHSC_01112 | formyl peptide receptor- | flr | Q2FZC0 |
| SAOUHSC_02160 | | | Q2FWW2 |
| SAOUHSC_01125 | | | Q2FZB2 |
| SAOUHSC_01127 | | | Q2FZB1 |
| SAOUHSC_01124 | | | Q2FZB3 |
| SAOUHSC_00399 | | | Q2G0X4 |
| SAOUHSC_02706 | | | Q2FVK5 |
| SAOUHSC_02862 | | | Q2FV74 |
| SAOUHSC_01420 | | | Q9KJN4 |
| SAOUHSC_02532 | | | Q2FVY9 |
| SAOUHSC_00715 | | | Q2G2G2 |
| SAOUHSC_00389 | | | Q2G1S8 |
| SAOUHSC_00354 | | | Q2G107 |
| SAOUHSC_00258 | | | Q2G188 |
| SAOUHSC_00714 | | | Q2G2U1 |
| SAOUHSC_02955 | | | Q2FUZ0 |
| SAOUHSC_00468 | | | Q2G0S6 |
| SAOUHSC_00438 | | | Q2G0T9 |
| SAOUHSC_01601 | | | Q2FY64 |
| SAOUHSC_00780 | | | Q2G046 |
| SAOUHSC_00779 | | | Q2G047 |
| SAOUHSC_00776 | | | Q2G050 |
| SAOUHSC_00860 | | | Q2FZX5 |
| SAOUHSC_00025 | | | Q2G253 |
| SAOUHSC_00501 | | | Q2G0P9 |
| SAOUHSC_02444 | | | Q2FW63 |
| SAOUHSC_02563 | | | Q2G2K7 |
| SAOUHSC_02564 | | | Q2G273 |
| SAOUHSC_02565 | | | Q2G272 |
| SAOUHSC_02134 | | | Q2G234 |
| SAOUHSC_02743 | | | Q2FVG9 |
| SAOUHSC_01380 | | | Q2FYQ5 |
| SAOUHSC_01379 | | | Q2FYQ6 |

| | | | |
|---|---|---|---|
| SAOUHSC_00169 | | | Q2G1F6 |
| SAOUHSC_00931 | | | Q2FZQ9 |
| SAOUHSC_00201 | | | Q2G2P5 |
| SAOUHSC_00138 | | | Q2G1I4 |
| SAOUHSC_02547 | | | Q2FVX5 |
| SAOUHSC_01228 | | | Q2FZ27 |
| SAOUHSC_01850 | | | Q2G295 |
| SAOUHSC_01997 | | | Q2G282 |
| SAOUHSC_00234 | | | Q2G1B1 |
| SAOUHSC_01254 | | | Q2FZ17 |
| SAOUHSC_03045 | | | Q2FUQ9 |
| SAOUHSC_01142 | | | Q2FZ97 |
| SAOUHSC_00439 | | | Q2G0T8 |
| SAOUHSC_00943 | | | Q2G1Z9 |
| SAOUHSC_01203 | | | Q2FZ50 |
| SAOUHSC_02549 | | | Q2FVX4 |
| SAOUHSC_02276 | | | Q2FWL3 |
| SAOUHSC_02245 | | | Q2FWN7 |
| SAOUHSC_00306 | | | Q2G149 |
| SAOUHSC_01916 | | | Q2G2V3 |
| SAOUHSC_01915 | | | Q2G2V2 |
| SAOUHSC_01488 | | | Q2FYG4 |
| SAOUHSC_02403 | | | Q2FW96 |
| SAOUHSC_00177 | | | Q2G1E8 |
| SAOUHSC_00178 | | | Q2G1E7 |
| SAOUHSC_00176 | | | Q2G1E9 |
| SAOUHSC_00145 | | | Q2G1H8 |
| SAOUHSC_02135 | | | Q2G233 |
| SAOUHSC_01416 | | | Q2FYM2 |
| SAOUHSC_00782 | | | Q2G044 |
| SAOUHSC_00114 | | | Q2G1K8 |
| SAOUHSC_00118 | | | Q2G1K4 |
| SAOUHSC_01071 | | | Q2FZF9 |
| SAOUHSC_01830 | | | Q2FXK4 |
| SAOUHSC_00897 | | | Q2FZU3 |
| SAOUHSC_00051 | 1-phosphatidylinositol ph | plc | Q2G1Q2 |
| SAOUHSC_00300 | | | Q2G155 |
| SAOUHSC_03006 | | | Q2FUU5 |
| SAOUHSC_00534 | | | Q2G0M6 |
| SAOUHSC_02585 | | | Q2FVV6 |
| SAOUHSC_00199 | | | Q2G1C6 |
| SAOUHSC_00851 | | | Q2FZY3 |
| SAOUHSC_00327 | | | Q2G129 |
| SAOUHSC_02864 | | | Q2FV72 |
| SAOUHSC_00326 | | | Q2G130 |
| SAOUHSC_00625 | | | Q2G2U8 |
| SAOUHSC_02108 | | | Q2FWZ8 |
| SAOUHSC_00328 | | | Q2G128 |
| SAOUHSC_01055 | | | Q2G2G4 |
| SAOUHSC_01327 | | | Q2FYU7 |
| SAOUHSC_01319 | | | Q2FYV5 |
| SAOUHSC_02606 | | | Q2FVT6 |
| SAOUHSC_02607 | | | Q2FVT5 |
| SAOUHSC_00008 | | | Q2G2P7 |

| | | | |
|---|---|---|---|
| SAOUHSC_02709 | Gamma-hemolysin comp | hlgC | Q2FVK2 |
| SAOUHSC_02710 | Gamma-hemolysin comp | hlgB | Q2FVK1 |
| SAOUHSC_01955 | leukotoxin LukE | lukE | Q2FXB0 |
| SAOUHSC_01954 | leukotoxin LukD | lukD | Q2FXB1 |
| SAOUHSC_01121 | alpha-hemolysin | hla | Q2G1X0 |
| SAOUHSC_02260 | delta-hemolysin | hld | Q2FWM8 |
| SAOUHSC_01085 | | | Q2FZE6 |
| SAOUHSC_01082 | | | Q8KQR1 |
| SAOUHSC_01089 | | | Q2FZE2 |
| SAOUHSC_00130 | | | Q2G1J2 |
| SAOUHSC_01774 | | | O34090 |
| SAOUHSC_02811 | | | Q2G2L7 |
| SAOUHSC_01801 | | | Q2FXN4 |
| SAOUHSC_01418 | | | Q2FYM1 |
| SAOUHSC_00799 | | | Q2G028 |
| SAOUHSC_02922 | | | Q2G1Y5 |
| SAOUHSC_00206 | | | Q2G218 |
| SAOUHSC_00305 | | | Q2G150 |
| SAOUHSC_02741 | | | Q2FVH1 |
| SAOUHSC_02742 | | | Q2FVH0 |
| SAOUHSC_02744 | | | Q2FVG8 |
| SAOUHSC_01276 | | | Q2FYZ5 |
| SAOUHSC_01278 | | | Q2FYZ4 |
| SAOUHSC_01275 | | | Q2FYZ6 |
| SAOUHSC_00658 | | | Q2G0E8 |
| SAOUHSC_00656 | | | Q2G0E9 |
| SAOUHSC_00655 | | | Q2G1Z0 |
| SAOUHSC_02244 | | | Q2FWN8 |
| SAOUHSC_02118 | | | Q2FWY8 |
| SAOUHSC_02869 | | | Q2FV67 |
| SAOUHSC_02760 | | | Q2FVF4 |
| SAOUHSC_00188 | | | Q2G1D7 |
| SAOUHSC_02703 | | | Q2FVK8 |
| SAOUHSC_02840 | | | Q2FV94 |
| SAOUHSC_02839 | | | Q2FV95 |
| SAOUHSC_02822 | | | Q2FVB2 |
| SAOUHSC_01910 | | | Q2G1W2 |
| SAOUHSC_02806 | | | Q2G206 |
| SAOUHSC_01144 | | | Q2FZ95 |
| SAOUHSC_00142 | | | Q2G1I1 |
| SAOUHSC_01650 | | | Q2FY23 |
| SAOUHSC_00489 | | | Q2G0Q7 |
| SAOUHSC_01904 | | | Q2FXE5 |
| SAOUHSC_01903 | | | Q2FXE6 |
| SAOUHSC_00198 | | | Q2G1C7 |
| SAOUHSC_00061 | | | Q2G1P3 |
| SAOUHSC_00557 | | | Q2G0K3 |
| SAOUHSC_01846 | | | Q2G294 |
| SAOUHSC_00139 | | | Q2G1I3 |
| SAOUHSC_00115 | | | Q2G1K7 |
| SAOUHSC_00961 | | | Q2FZN9 |
| SAOUHSC_01691 | | | Q2FXY4 |
| SAOUHSC_02793 | | | Q2FVC1 |
| SAOUHSC_00239 | | | Q2G1A6 |

| | | | |
|---|---|---|---|
| SAOUHSC_00240 | | | Q2G1A5 |
| SAOUHSC_01615 | | | Q2FY50 |
| SAOUHSC_01164 | | | Q2FZ77 |
| SAOUHSC_02298 | | | Q2FWJ4 |
| SAOUHSC_02111 | | | Q2FWZ5 |
| SAOUHSC_01797 | | | Q2FXN9 |
| SAOUHSC_00454 | | | Q2G2X1 |
| SAOUHSC_01095 | | | Q2FZD7 |
| SAOUHSC_01472 | | | Q2FYH5 |
| SAOUHSC_02334 | | | Q2FWF7 |
| SAOUHSC_02942 | | | Q2FV02 |
| SAOUHSC_00765 | | | Q2G057 |
| SAOUHSC_01341 | | | Q2FYT4 |
| SAOUHSC_01723 | | | Q2FXV8 |
| SAOUHSC_01821 | | | Q2FXL4 |
| SAOUHSC_01221 | | | Q2FZ33 |
| SAOUHSC_00624 | | | Q2G2V0 |
| SAOUHSC_01502 | | | Q2FYF0 |
| SAOUHSC_00469 | | | Q2G0S5 |
| SAOUHSC_02808 | | | Q2G205 |
| SAOUHSC_00538 | | | Q2G0M2 |
| SAOUHSC_00743 | | | Q2G077 |
| SAOUHSC_01170 | | | Q2FZ72 |
| SAOUHSC_01168 | | | Q2FZ74 |
| SAOUHSC_02909 | | | Q2FV30 |
| SAOUHSC_01166 | | | Q2FZ75 |
| SAOUHSC_02243 | Leukocidin; Leukocidin/H | lukH | Q2FWN9 |
| SAOUHSC_02241 | hypothetical protein | lukG | Q2FWP0 |
| SAOUHSC_02708 | gamma-hemolysin h-gar | hlgA | Q2FVK3 |
| SAOUHSC_00233 | | | P60643 |
| SAOUHSC_01953 | | | Q2FXB2 |
| SAOUHSC_02422 | | | Q2FW82 |
| SAOUHSC_01775 | | | Q2FXR0 |
| SAOUHSC_01485 | | | Q2FYG7 |
| SAOUHSC_00182 | | | Q2G1E3 |
| SAOUHSC_01950 | | | Q2FXB5 |
| SAOUHSC_02765 | | | Q2FVE9 |
| SAOUHSC_02763 | | | Q2FVF1 |
| SAOUHSC_02766 | | | Q2FVE8 |
| SAOUHSC_02764 | | | Q2FVF0 |
| SAOUHSC_02767 | | | Q2FVE7 |
| SAOUHSC_02996 | | | Q2FUV5 |
| SAOUHSC_01053 | | | Q2G2G3 |
| SAOUHSC_02472 | | | Q2FW45 |
| SAOUHSC_02885 | | | Q2FV54 |
| SAOUHSC_02753 | | | Q2FVG1 |
| SAOUHSC_01307 | | | Q2FYW6 |
| SAOUHSC_00640 | | | Q2G2L3 |
| SAOUHSC_00643 | | | Q2G2X4 |
| SAOUHSC_00222 | | | Q2G1C2 |
| SAOUHSC_00223 | | | Q2G1C1 |
| SAOUHSC_00998 | | | Q2FZK3 |
| SAOUHSC_01220 | | | Q2FZ34 |
| SAOUHSC_01219 | | | Q9ZNI1 |

| | | | |
|---|---|---|---|
| SAOUHSC_00365 | | | P0A0B7 |
| SAOUHSC_00939 | | | Q2G202 |
| SAOUHSC_01614 | | | Q2FY51 |
| SAOUHSC_02654 | | | Q2FVP8 |
| SAOUHSC_02319 | | | Q2FWH2 |
| SAOUHSC_01146 | | | Q2FZ93 |
| SAOUHSC_00634 | | | Q2G2D8 |
| SAOUHSC_00544 | | | Q2G0L5 |
| SAOUHSC_01238 | | | Q2G1Z6 |
| SAOUHSC_02880 | | | Q2FV58 |
| SAOUHSC_02881 | hypothetical protein | crtP | Q2FV57 |
| SAOUHSC_02879 | squalene desaturase | crtM | Q2FV59 |
| SAOUHSC_02877 | | | Q2FV60 |
| SAOUHSC_02142 | | | Q2FWX9 |
| SAOUHSC_02882 | | | Q2FV56 |
| SAOUHSC_01310 | | | Q2FYW4 |
| SAOUHSC_01867 | | | Q2FXI0 |
| SAOUHSC_00175 | | | Q2G1F0 |
| SAOUHSC_00158 | | | Q2G1G5 |
| SAOUHSC_02976 | | | Q2G223 |
| SAOUHSC_00295 | | | Q2G160 |
| SAOUHSC_00710 | | | Q2G2U0 |
| SAOUHSC_02463 | | | Q59801 |
| SAOUHSC_00160 | | | Q2G1G4 |
| SAOUHSC_02599 | | | Q2FVU3 |
| SAOUHSC_02379 | | | Q2FWB9 |
| SAOUHSC_00100 | | | Q2G224 |
| SAOUHSC_02587 | | | Q2FVV4 |
| SAOUHSC_02256 | | | Q2FWN2 |
| SAOUHSC_01900 | | | Q2FXE9 |
| SAOUHSC_01612 | | | Q2FY53 |
| SAOUHSC_01796 | | | Q2FXP0 |
| SAOUHSC_00612 | | | Q2G0F7 |
| SAOUHSC_00628 | | | Q2G212 |
| SAOUHSC_01129 | | | Q2FZA9 |
| SAOUHSC_02965 | | | Q7X2S2 |
| SAOUHSC_02409 | | | Q2FW91 |
| SAOUHSC_02969 | | | Q2FUX7 |
| SAOUHSC_02967 | | | Q2FUX9 |
| SAOUHSC_01128 | | | Q2FZB0 |
| SAOUHSC_02968 | | | Q2FUX8 |
| SAOUHSC_01868 | | | Q2FXH9 |
| SAOUHSC_02970 | | | Q2FUX5 |
| SAOUHSC_01617 | | | Q2FY49 |
| SAOUHSC_02137 | | | Q2FWY4 |
| SAOUHSC_00144 | | | Q2G1H9 |
| SAOUHSC_00064 | | | Q2G1P1 |
| SAOUHSC_00113 | | | Q2G1K9 |
| SAOUHSC_00812 | | | Q2G015 |
| SAOUHSC_02929 | | | Q2FV14 |
| SAOUHSC_01849 | | | Q2G292 |
| SAOUHSC_01847 | | | Q2G293 |
| SAOUHSC_00086 | | | Q2G1M1 |
| SAOUHSC_00101 | | | Q2G227 |

| | |
|---|---|
| SAOUHSC_00622 | Q2G2V1 |
| SAOUHSC_01944 | Q2FXC0 |
| SAOUHSC_00845 | Q2FZY9 |
| SAOUHSC_02442 | Q2FW65 |
| SAOUHSC_02443 | Q2FW64 |
| SAOUHSC_00156 | Q2G1G7 |
| SAOUHSC_01952 | Q2FXB3 |
| SAOUHSC_02441 | Q2FW66 |
| SAOUHSC_01306 | Q2FYW7 |
| SAOUHSC_00560 | Q2G0K0 |
| SAOUHSC_02795 | Q2G2B5 |
| SAOUHSC_01456 | Q2FYI9 |
| SAOUHSC_00143 | Q2G1I0 |
| SAOUHSC_00818 | Q2G010 |
| SAOUHSC_00284 | Q2G170 |
| SAOUHSC_00358 | Q2G103 |
| SAOUHSC_00116 | Q2G1K6 |
| SAOUHSC_02863 | Q2FV73 |
| SAOUHSC_02666 | Q2FVN6 |
| SAOUHSC_01913 | Q2G2V5 |
| SAOUHSC_00356 | Q2G105 |
| SAOUHSC_02519 | Q2FW01 |
| SAOUHSC_02387 | Q2FWB1 |
| SAOUHSC_00827 | Q2G1T1 |
| SAOUHSC_00410 | Q2G0W4 |
| SAOUHSC_00117 | Q2G1K5 |
| SAOUHSC_01081 | Q2FZE9 |
| SAOUHSC_00637 | Q2G2L2 |
| SAOUHSC_02865 | Q2FV71 |
| SAOUHSC_02688 | Q2G1U7 |
| SAOUHSC_00179 | Q2G1E6 |
| SAOUHSC_01297 | Q2FYX6 |
| SAOUHSC_01948 | Q2FXB7 |
| SAOUHSC_02774 | Q2FVE0 |
| SAOUHSC_00585 | Q2G0I0 |
| SAOUHSC_02777 | Q2FVD6 |
| SAOUHSC_02153 | Q2FWW8 |
| SAOUHSC_01643 | Q2FY30 |
| SAOUHSC_03023 | Q2FUS8 |
| SAOUHSC_01729 | Q2FXV2 |
| SAOUHSC_01318 | Q2FYV6 |
| SAOUHSC_00716 | Q2G2G1 |
| SAOUHSC_02812 | Q2G2L6 |
| SAOUHSC_00181 | Q2G1E4 |
| SAOUHSC_00180 | Q2G1E5 |
| SAOUHSC_02129 | Q2G2R7 |
| SAOUHSC_01730 | Q2FXV1 |
| SAOUHSC_01130 | Q2FZA8 |
| SAOUHSC_01947 | Q2FXB8 |
| SAOUHSC_02778 | Q2FVD5 |
| SAOUHSC_02161 | Q2FWW1 |
| SAOUHSC_02751 | Q2FVG3 |
| SAOUHSC_02154 | Q2FWW7 |
| SAOUHSC_01088 | Q2FZE3 |

| | |
|---|---|
| SAOUHSC_01920 | Q2G2Y5 |
| SAOUHSC_00687 | Q2G0B8 |
| SAOUHSC_02719 | Q2FVJ1 |
| SAOUHSC_02152 | Q2FWW9 |
| SAOUHSC_01923 | Q2FXE0 |
| SAOUHSC_00085 | Q2G1M2 |
| SAOUHSC_01054 | Q2G2G7 |
| SAOUHSC_02265 | Q2FWM4 |
| SAOUHSC_01921 | Q2G2Y2 |
| SAOUHSC_00588 | Q2G0H7 |
| SAOUHSC_01165 | Q2FZ76 |
| SAOUHSC_03035 | Q2FUR7 |
| SAOUHSC_02794 | Q2FVC0 |
| SAOUHSC_00775 | Q2G051 |
| SAOUHSC_00586 | Q2G0H9 |
| SAOUHSC_03028 | Q2FUS3 |
| SAOUHSC_02771 | Q2FVE3 |
| SAOUHSC_01945 | Q2FXB9 |
| SAOUHSC_00119 | Q2G1K3 |
| SAOUHSC_00325 | Q2G131 |
| SAOUHSC_00123 | Q2G1J9 |
| SAOUHSC_00371 | Q2G0Z0 |
| SAOUHSC_02645 | Q2FVQ7 |
| SAOUHSC_00241 | Q2G1A4 |
| SAOUHSC_00859 | Q2FZX6 |
| SAOUHSC_01255 | Q2FZ16 |
| SAOUHSC_00074 | Q2G1N4 |
| SAOUHSC_00363 | Q2G0Z8 |
| SAOUHSC_00185 | Q2G1E0 |
| SAOUHSC_01641 | Q2G2I6 |
| SAOUHSC_03046 | Q2FUQ8 |
| SAOUHSC_02155 | Q2FWW6 |
| SAOUHSC_00304 | Q2G151 |
| SAOUHSC_01595 | Q2FY70 |
| SAOUHSC_00559 | Q2G0K1 |
| SAOUHSC_00717 | Q2G2G0 |
| SAOUHSC_00596 | Q2G0H1 |
| SAOUHSC_00043 | Q2G1Q9 |
| SAOUHSC_00592 | Q2G0H4 |
| SAOUHSC_01181 | Q2G267 |
| SAOUHSC_01890 | Q2FXF8 |
| SAOUHSC_00807 | Q2G020 |
| SAOUHSC_00808 | Q2G019 |
| SAOUHSC_00858 | Q2FZX7 |
| SAOUHSC_01303 | Q2FYX0 |
| SAOUHSC_00120 | Q2G1K2 |
| SAOUHSC_00864 | Q2FZX1 |
| SAOUHSC_01443 | Q2FYJ7 |
| SAOUHSC_00237 | Q2G1A8 |
| SAOUHSC_00124 | Q2G1J8 |
| SAOUHSC_02604 | Q2FVT8 |
| SAOUHSC_00377 | Q2G0Y4 |
| SAOUHSC_00593 | Q2G0H3 |
| SAOUHSC_02246 | Q2FWN6 |

| | |
|---|---|
| SAOUHSC_01596 | Q2FY69 |
| SAOUHSC_00806 | Q2G021 |
| SAOUHSC_02557 | Q2FVW6 |
| SAOUHSC_02872 | Q2FV65 |
| SAOUHSC_00122 | Q2G1K0 |
| SAOUHSC_02262 | Q2FWM6 |
| SAOUHSC_01922 | Q2FXE1 |
| SAOUHSC_00121 | Q2G1K1 |
| SAOUHSC_03034 | Q2FUR8 |
| SAOUHSC_02646 | Q2FVQ6 |
| SAOUHSC_00938 | Q2G203 |
| SAOUHSC_00972 | Q2FZM8 |
| SAOUHSC_02759 | Q2FVF5 |
| SAOUHSC_01301 | Q2FYX2 |
| SAOUHSC_02994 | Q2FUV7 |
| SAOUHSC_00712 | Q2G2T8 |
| SAOUHSC_00312 | Q2G143 |
| SAOUHSC_00319 | Q2G136 |
| SAOUHSC_00320 | Q2G135 |
| SAOUHSC_00686 | Q2G0B9 |
| SAOUHSC_00975 | Q2FZM5 |
| SAOUHSC_01924 | Q2FXD9 |
| SAOUHSC_00256 | Q2G190 |
| SAOUHSC_02776 | Q2FVD8 |
| SAOUHSC_01213 | Q2FZ40 |
| SAOUHSC_00826 | Q2G1T2 |
| SAOUHSC_02097 | Q2FX10 |
| SAOUHSC_00617 | Q2G0F2 |
| SAOUHSC_00360 | Q2G100 |
| SAOUHSC_02381 | Q2FWB7 |
| SAOUHSC_00433 | Q2G0U4 |
| SAOUHSC_00591 | Q2G0H5 |
| SAOUHSC_02663 | Q2FVN9 |
| SAOUHSC_00973 | Q2FZM7 |
| SAOUHSC_02832 | Q2FVA1 |
| SAOUHSC_02419 | Q2FW84 |
| SAOUHSC_03022 | Q2FUS9 |
| SAOUHSC_00409 | Q2G0W5 |
| SAOUHSC_00940 | Q2G201 |
| SAOUHSC_00202 | Q2G2P4 |
| SAOUHSC_02844 | Q2FV90 |
| SAOUHSC_02465 | Q2FW52 |
| SAOUHSC_00673 | Q2G0D2 |
| SAOUHSC_01805 | Q2FXN0 |
| SAOUHSC_01137 | Q2FZA2 |
| SAOUHSC_00318 | Q2G137 |
| SAOUHSC_02761 | Q2FVF3 |
| SAOUHSC_02104 | Q2FX02 |
| SAOUHSC_02149 | Q2FWX2 |
| SAOUHSC_02772 | Q2FVE2 |
| SAOUHSC_01598 | Q2FY67 |
| SAOUHSC_01305 | Q2FYW8 |
| SAOUHSC_00917 | Q2FZS3 |
| SAOUHSC_02972 | Q2FUX3 |

| | |
|---|---|
| SAOUHSC_02762 | Q2FVF2 |
| SAOUHSC_02101 | Q2FX06 |
| SAOUHSC_00587 | Q2G0H8 |
| SAOUHSC_02998 | Q2FUV3 |
| SAOUHSC_00823 | Q2G005 |
| SAOUHSC_00415 | Q2G0V9 |
| SAOUHSC_02009 | Q2G2H3 |
| SAOUHSC_01292 | Q2FYY1 |
| SAOUHSC_00271 | Q2G176 |
| SAOUHSC_00146 | Q2G1H7 |
| SAOUHSC_01606 | Q2FY59 |
| SAOUHSC_00307 | Q2G148 |
| SAOUHSC_00825 | Q2G003 |
| SAOUHSC_00408 | Q2G0W6 |
| SAOUHSC_01626 | Q2FY40 |
| SAOUHSC_02596 | Q2FVU5 |
| SAOUHSC_02144 | Q2FWX7 |
| SAOUHSC_01926 | Q2FXD7 |
| SAOUHSC_01084 | Q2FZE7 |
| SAOUHSC_02726 | Q2FVI4 |
| SAOUHSC_00197 | Q2G1C8 |
| SAOUHSC_00272 | Q2G175 |
| SAOUHSC_02376 | Q2FWC2 |
| SAOUHSC_02464 | Q2FW53 |
| SAOUHSC_00431 | Q2G0U5 |
| SAOUHSC_00865 | Q2FZX0 |
| SAOUHSC_03017 | Q2FUT4 |
| SAOUHSC_00713 | Q2G2T7 |
| SAOUHSC_00697 | Q2G0A8 |
| SAOUHSC_01113 | Q2FZB9 |
| SAOUHSC_01376 | Q2FYQ9 |
| SAOUHSC_00810 | Q2G017 |
| SAOUHSC_01966 | Q2G2F2 |
| SAOUHSC_02979 | Q2G222 |
| SAOUHSC_00696 | Q2G0A9 |
| SAOUHSC_01860 | Q2FXI6 |
| SAOUHSC_02008 | Q2G2V7 |
| SAOUHSC_02471 | Q2FW46 |
| SAOUHSC_01511 | Q2FYE3 |
| SAOUHSC_02809 | Q2G208 |
| SAOUHSC_02781 | Q2FVD2 |
| SAOUHSC_00042 | Q2G1R0 |
| SAOUHSC_01798 | Q2FXN8 |
| SAOUHSC_01134 | Q2FZA5 |
| SAOUHSC_02941 | Q2FV03 |
| SAOUHSC_02711 | Q2FVK0 |
| SAOUHSC_02797 | Q2G2B3 |
| SAOUHSC_02768 | Q2FVE6 |
| SAOUHSC_02775 | Q2FVD9 |
| SAOUHSC_03047 | Q2FUQ7 |
| SAOUHSC_00598 | Q2G0H0 |
| SAOUHSC_01296 | Q2FYX7 |
| SAOUHSC_01464 | Q2FYI3 |
| SAOUHSC_00672 | Q2G0D3 |

| | |
|---|---|
| SAOUHSC_01508 | Q2FYE5 |
| SAOUHSC_01649 | Q2FY24 |
| SAOUHSC_02827 | Q2FVA6 |
| SAOUHSC_02433 | Q2FW73 |
| SAOUHSC_00602 | Q2G0G7 |
| SAOUHSC_00621 | Q2G2U7 |
| SAOUHSC_02393 | Q2FWA5 |
| SAOUHSC_01463 | Q2FYI4 |
| SAOUHSC_01971 | Q2G2S9 |
| SAOUHSC_00816 | Q2G012 |
| SAOUHSC_00125 | Q2G1J7 |
| SAOUHSC_00270 | Q2G177 |
| SAOUHSC_00976 | Q2FZM4 |
| SAOUHSC_02845 | Q2FV89 |
| SAOUHSC_02913 | Q2FV26 |
| SAOUHSC_00922 | Q2FZR8 |
| SAOUHSC_01256 | Q2FZ15 |
| SAOUHSC_02650 | Q2FVQ2 |
| SAOUHSC_01859 | Q2FXI7 |
| SAOUHSC_00367 | Q2G0Z4 |
| SAOUHSC_01304 | Q2FYW9 |
| SAOUHSC_02727 | Q2FVI3 |
| SAOUHSC_00084 | Q2G1M3 |
| SAOUHSC_00711 | Q2G2T9 |
| SAOUHSC_00508 | Q2G242 |
| SAOUHSC_02099 | Q2FX08 |
| SAOUHSC_01967 | Q2G2F1 |
| SAOUHSC_02995 | Q2FUV6 |
| SAOUHSC_02892 | Q2FV47 |
| SAOUHSC_00599 | Q2G0G9 |
| SAOUHSC_00280 | Q2G173 |
| SAOUHSC_02583 | Q2FVV8 |
| SAOUHSC_01279 | Q2FYZ3 |
| SAOUHSC_01349 | Q2FYS7 |
| SAOUHSC_02396 | Q2FWA2 |
| SAOUHSC_02143 | Q2FWX8 |
| SAOUHSC_01005 | Q2FZJ7 |
| SAOUHSC_00565 | Q2G0J6 |
| SAOUHSC_01822 | Q2FXL3 |
| SAOUHSC_01324 | Q2FYV0 |
| SAOUHSC_01831 | Q2FXK3 |
| SAOUHSC_01957 | Q2FXA8 |
| SAOUHSC_02900 | Q2FV39 |
| SAOUHSC_00322 | Q2G134 |
| SAOUHSC_00261 | Q2G185 |
| SAOUHSC_00830 | Q2G1T4 |
| SAOUHSC_01761 | Q2FXS4 |
| SAOUHSC_01414 | Q2G2J7 |
| SAOUHSC_00774 | Q2G277 |
| SAOUHSC_02006 | Q2G2V9 |
| SAOUHSC_00141 | Q2G1I2 |
| SAOUHSC_00172 | Q2G1F3 |
| SAOUHSC_02853 | Q2FV82 |
| SAOUHSC_00561 | Q2G0J9 |

| | |
|---|---|
| SAOUHSC_00030 | Q2G2E9 |
| SAOUHSC_00094 | Q2G260 |
| SAOUHSC_02112 | Q2FWZ4 |
| SAOUHSC_01433 | Q2FYK7 |
| SAOUHSC_03031 | Q2FUS1 |
| SAOUHSC_01069 | Q2FZG1 |
| SAOUHSC_01405 | Q2FYN0 |
| SAOUHSC_00890 | Q2FZV0 |
| SAOUHSC_00401 | Q2G0X2 |
| SAOUHSC_02515 | Q2FW05 |
| SAOUHSC_02555 | Q2FVW8 |
| SAOUHSC_02147 | Q2FWX4 |
| SAOUHSC_02842 | Q2FV92 |
| SAOUHSC_00052 | Q2G1Q1 |
| SAOUHSC_02980 | Q2G220 |
| SAOUHSC_01819 | Q2FXL6 |
| SAOUHSC_01133 | Q2FZA6 |
| SAOUHSC_00919 | Q2FZS1 |
| SAOUHSC_02577 | Q2FVW4 |
| SAOUHSC_02736 | Q2FVH6 |
| SAOUHSC_01781 | Q2FXQ4 |
| SAOUHSC_02324 | Q2FWG7 |
| SAOUHSC_02568 | Q2G270 |
| SAOUHSC_00200 | Q2G1C5 |
| SAOUHSC_01724 | Q2FXV7 |
| SAOUHSC_02367 | Q2FWD2 |
| SAOUHSC_01958 | Q2FXA7 |
| SAOUHSC_03030 | Q2FUS2 |
| SAOUHSC_00866 | Q2FZW9 |
| SAOUHSC_01317 | Q2FYV7 |
| SAOUHSC_02570 | Q2G2J4 |
| SAOUHSC_00604 | Q2G0G5 |
| SAOUHSC_00406 | Q2G0W8 |
| SAOUHSC_02138 | Q2FWY3 |
| SAOUHSC_01408 | Q2FYM7 |
| SAOUHSC_00203 | Q2G2P3 |
| SAOUHSC_01512 | Q2FYE2 |
| SAOUHSC_00659 | Q2G0E7 |
| SAOUHSC_02150 | Q2FWX1 |
| SAOUHSC_02875 | Q2FV62 |
| SAOUHSC_00632 | Q2G2E1 |
| SAOUHSC_01998 | Q2G281 |
| SAOUHSC_00996 | P0A0Q4 |
| SAOUHSC_00056 | Q2G1P7 |
| SAOUHSC_02424 | Q2FW80 |
| SAOUHSC_00055 | Q2G1P8 |
| SAOUHSC_00618 | Q2G0F1 |
| SAOUHSC_00163 | Q2G1G1 |

GOBIO

#N/A
#N/A
#N/A
#N/A
#N/A
#N/A
#N/A
#N/A
#N/A
#N/A
#N/A
#N/A
zinc ion transmembrane transport [GO:0071577]
UDP-N-acetylglucosamine biosynthetic process [GO:0006048]
tRNA modification [GO:0006400]
tricarboxylic acid cycle [GO:0006099]
tricarboxylic acid cycle [GO:0006099]
transposition, DNA-mediated [GO:0006313]
transmembrane transport [GO:0055085]
transmembrane transport [GO:0055085]
transmembrane transport [GO:0055085]
transmembrane transport [GO:0055085]
transmembrane transport [GO:0055085]
transmembrane transport [GO:0055085]
transmembrane transport [GO:0055085]
transmembrane transport [GO:0055085]
transmembrane transport [GO:0055085]
transmembrane transport [GO:0055085]
transmembrane transport [GO:0055085]
transmembrane transport [GO:0055085]
transmembrane transport [GO:0055085]
translation [GO:0006412]
threonine catabolic process [GO:0006567]
thiamine transport [GO:0015888]
thiamine biosynthetic process [GO:0009228]; thiamine diphosphate biosynthetic process [GO:00092
thiamine biosynthetic process [GO:0009228]; thiamine diphosphate biosynthetic process [GO:00092
tetrahydrofolate interconversion [GO:0035999]
sulfide oxidation, using sulfide:quinone oxidoreductase [GO:0070221]
succinate transport [GO:0015744]
succinate transport [GO:0015744]
steroid biosynthetic process [GO:0006694]
sodium ion transport [GO:0006814]
signal peptide processing [GO:0006465]
siderophore-dependent iron import into cell [GO:0033214]
siderophore-dependent iron import into cell [GO:0033214]
siderophore-dependent iron import into cell [GO:0033214]
siderophore-dependent iron import into cell [GO:0033214]
siderophore biosynthetic process [GO:0019290]
siderophore biosynthetic process [GO:0019290]
response to zinc ion [GO:0010043]
response to toxic substance [GO:0009636]
response to reactive oxygen species [GO:0000302]
response to oxidative stress [GO:0006979]

response to antibiotic [GO:0046677]
response to antibiotic [GO:0046677]
response to antibiotic [GO:0046677]
regulation of transcription, DNA-templated [GO:0006355]
regulation of transcription, DNA-templated [GO:0006355]
regulation of transcription, DNA-templated [GO:0006355]
regulation of transcription, DNA-templated [GO:0006355]
regulation of transcription, DNA-templated [GO:0006355]
regulation of transcription, DNA-templated [GO:0006355]
regulation of transcription, DNA-templated [GO:0006355]
regulation of transcription, DNA-templated [GO:0006355]
regulation of transcription, DNA-templated [GO:0006355]
regulation of transcription, DNA-templated [GO:0006355]
regulation of DNA-templated transcription [GO:0006355]
regulation of DNA-templated transcription [GO:0006355]
regulation of DNA repair [GO:0006282]
pyrimidine nucleobase metabolic process [GO:0006206]; pyrimidine nucleoside metabolic process [G
purine nucleoside catabolic process [GO:0006152]
proteolysis [GO:0006508]
proteolysis [GO:0006508]
proteolysis [GO:0006508]
proteolysis [GO:0006508]
proteolysis [GO:0006508]
proteolysis [GO:0006508]
proteolysis [GO:0006508]
proteolysis [GO:0006508]
proteolysis [GO:0006508]
proteolysis [GO:0006508]
proteolysis [GO:0006508]
proteolysis [GO:0006508]
proteolysis [GO:0006508]
proteolysis [GO:0006508]
protein unfolding [GO:0043335]
protein transport by the Sec complex [GO:0043952]; SRP-dependent cotranslational protein targeting
protein lipoylation [GO:0009249]
protein initiator methionine removal [GO:0070084]
protein folding [GO:0006457]
protein dephosphorylation [GO:0006470]
protein biotinylation [GO:0009305]; regulation of transcription, DNA-templated [GO:0006355]
protein autophosphorylation [GO:0046777]
protein autophosphorylation [GO:0046777]
proline catabolic process to glutamate [GO:0010133]
potassium ion transmembrane transport [GO:0071805]; regulation of intracellular pH [GO:0051453];
potassium ion transmembrane transport [GO:0071805]
phosphorylation [GO:0016310]
phosphorelay signal transduction system [GO:0000160]
phosphorelay signal transduction system [GO:0000160]
phospholipid biosynthetic process [GO:0008654]
phosphoenolpyruvate-dependent sugar phosphotransferase system [GO:0009401]; regulation of tran
phosphoenolpyruvate-dependent sugar phosphotransferase system [GO:0009401]; regulation of tran
phosphoenolpyruvate-dependent sugar phosphotransferase system [GO:0009401]; regulation of tran
phosphoenolpyruvate-dependent sugar phosphotransferase system [GO:0009401]
phosphoenolpyruvate-dependent sugar phosphotransferase system [GO:0009401]
phosphoenolpyruvate-dependent sugar phosphotransferase system [GO:0009401]

phosphoenolpyruvate-dependent sugar phosphotransferase system [GO:0009401]
phosphoenolpyruvate-dependent sugar phosphotransferase system [GO:0009401]
phosphoenolpyruvate-dependent sugar phosphotransferase system [GO:0009401]
phosphate ion transmembrane transport [GO:0035435]
peptidoglycan biosynthetic process [GO:0009252]; response to antibiotic [GO:0046677]
peptide transport [GO:0015833]
peptide modification [GO:0031179]
peptide metabolic process [GO:0006518]; proteolysis [GO:0006508]
peptide metabolic process [GO:0006518]; proteolysis [GO:0006508]
peptide catabolic process [GO:0043171]
pathogenesis [GO:0009405]; regulation of transcription, DNA-templated [GO:0006355]
pathogenesis [GO:0009405]; regulation of transcription, DNA-templated [GO:0006355]
pathogenesis [GO:0009405]; regulation of transcription, DNA-templated [GO:0006355]
pathogenesis [GO:0009405]; quorum sensing [GO:0009372]
pathogenesis [GO:0009405]; protein autophosphorylation [GO:0046777]
pathogenesis [GO:0009405]
pathogenesis [GO:0009405]
pathogenesis [GO:0009405]
pathogenesis [GO:0009405]
pathogenesis [GO:0009405]
pathogenesis [GO:0009405]
pathogenesis [GO:0009405]
pathogenesis [GO:0009405]
pathogenesis [GO:0009405]
pathogenesis [GO:0009405]
pathogenesis [GO:0009405]
pathogenesis [GO:0009405]
pathogenesis [GO:0009405]
pathogenesis [GO:0009405]
pathogenesis [GO:0009405]
pathogenesis [GO:0009405]
pathogenesis [GO:0009405]
pathogenesis [GO:0009405]
pathogenesis [GO:0009405]
pathogenesis [GO:0009405]
osmosensory signaling via phosphorelay pathway [GO:0007234]; pathogenesis [GO:0009405]
osmosensory signaling via phosphorelay pathway [GO:0007234]
organonitrogen compound catabolic process [GO:1901565]
oligosaccharide catabolic process [GO:0009313]; trehalose catabolic process [GO:0005993]
oligosaccharide catabolic process [GO:0009313]
nucleotide-excision repair [GO:0006289]; SOS response [GO:0009432]
nucleotide-excision repair [GO:0006289]
nucleotide-excision repair [GO:0006289]
nucleotide catabolic process [GO:0009166]
nucleotide catabolic process [GO:0009166]
nucleoside transmembrane transport [GO:1901642]
nitrogen compound transport [GO:0071705]
nitrogen compound metabolic process [GO:0006807]
nitrogen compound metabolic process [GO:0006807]
nitrogen compound metabolic process [GO:0006807]
nitric oxide biosynthetic process [GO:0006809]
nickel cation transport [GO:0015675]; transmembrane transport [GO:0055085]
nickel cation transport [GO:0015675]; transmembrane transport [GO:0055085]
nickel cation transport [GO:0015675]; transmembrane transport [GO:0055085]

nickel cation transport [GO:0015675]; transmembrane transport [GO:0055085]
nickel cation transport [GO:0015675]; transmembrane transport [GO:0055085]
nickel cation transport [GO:0015675]; peptide transport [GO:0015833]
nickel cation transport [GO:0015675]
nickel cation transport [GO:0015675]
negative regulation of transcription, DNA-templated [GO:0045892]; regulation of transcription, DNA-t
negative regulation of transcription, DNA-templated [GO:0045892]; regulation of transcription, DNA-t
negative regulation of transcription, DNA-templated [GO:0045892]; regulation of secondary metaboli
negative regulation of transcription, DNA-templated [GO:0045892]
negative regulation of transcription, DNA-templated [GO:0045892]
negative regulation of DNA-templated transcription, termination [GO:0060567]; regulation of gene ex
negative regulation of DNA-templated transcription, initiation [GO:2000143]
negative regulation of DNA-templated transcription [GO:0045892]
NAD metabolic process [GO:0019674]; NADP biosynthetic process [GO:0006741]
mRNA processing [GO:0006397]; RNA processing [GO:0006396]; rRNA catabolic process [GO:0016
molybdate ion transport [GO:0015689]
mismatch repair [GO:0006298]
methylation [GO:0032259]
metabolic process [GO:0008152]
menaquinone biosynthetic process [GO:0009234]
menaquinone biosynthetic process [GO:0009234]
menaquinone biosynthetic process [GO:0009234]
mannitol catabolic process [GO:0019592]
maltodextrin transport [GO:0042956]; nickel cation transport [GO:0015675]
maltodextrin transport [GO:0042956]; maltose transport [GO:0015768]; nickel cation transport [GO:0
maltodextrin transport [GO:0042956]; maltose transport [GO:0015768]
lysine biosynthetic process via aminoadipic acid [GO:0019878]
L-phenylalanine biosynthetic process [GO:0009094]
L-lysine catabolic process to acetyl-CoA via saccharopine [GO:0033512]; tricarboxylic acid cycle [GC
lipoprotein biosynthetic process [GO:0042158]
lipopolysaccharide biosynthetic process [GO:0009103]; polysaccharide transport [GO:0015774]
lipopolysaccharide biosynthetic process [GO:0009103]
lipid metabolic process [GO:0006629]
lipid metabolic process [GO:0006629]
lipid metabolic process [GO:0006629]
lipid catabolic process [GO:0016042]; pathogenesis [GO:0009405]; signal transduction [GO:0007165
lipid catabolic process [GO:0016042]
lipid catabolic process [GO:0016042]
L-arabinose catabolic process to xylulose 5-phosphate [GO:0019569]
lactose metabolic process [GO:0005988]
ketone body catabolic process [GO:0046952]
iron-sulfur cluster assembly [GO:0016226]
iron ion transmembrane transport [GO:0034755]
iron ion homeostasis [GO:0055072]
iron import into cell [GO:0033212]
ion transport [GO:0006811]
intracellular sequestering of iron ion [GO:0006880]; iron ion transport [GO:0006826]
intracellular protein transmembrane transport [GO:0065002]; protein transport by the Tat complex [G
inositol metabolic process [GO:0006020]; inositol phosphate dephosphorylation [GO:0046855]; signa
hydrogen peroxide catabolic process [GO:0042744]; response to hydrogen peroxide [GO:0042542]
homoserine biosynthetic process [GO:0009090]; lysine biosynthetic process via diaminopimelate [GC
histidine catabolic process [GO:0006548]; histidine catabolic process to glutamate and formamide [G
histidine catabolic process [GO:0006548]; histidine catabolic process to glutamate and formamide [G
histidine catabolic process [GO:0006548]; histidine catabolic process to glutamate and formamide [G

hemolysis in other organism [GO:0044179]; pathogenesis [GO:0009405]
hemolysis in other organism [GO:0044179]; pathogenesis [GO:0009405]
hemolysis in other organism [GO:0044179]; pathogenesis [GO:0009405]
hemolysis in other organism [GO:0044179]; pathogenesis [GO:0009405]
hemolysis in other organism [GO:0044179]; pathogenesis [GO:0009405]
hemolysis by symbiont of host erythrocytes [GO:0019836]; pathogenesis [GO:0009405]
heme transport [GO:0015886]
heme transport [GO:0015886]
heme catabolic process [GO:0042167]; iron import into cell [GO:0033212]
heme catabolic process [GO:0042167]; iron import into cell [GO:0033212]
heme biosynthetic process [GO:0006783]; peptidyl-pyrromethane cofactor linkage [GO:0018160]; pro
guanosine tetraphosphate biosynthetic process [GO:0015970]
glyoxylate cycle [GO:0006097]; tricarboxylic acid cycle [GO:0006099]
glycolytic process [GO:0006096]; tricarboxylic acid cycle [GO:0006099]
glycolytic process [GO:0006096]; pathogenesis [GO:0009405]; regulation of vacuole fusion, non-auto
glycolytic process [GO:0006096]
glycolytic process [GO:0006096]
glycine decarboxylation via glycine cleavage system [GO:0019464]; protein lipoylation [GO:0009249
glycine betaine transport [GO:0031460]; nickel cation transport [GO:0015675]
glycine betaine transport [GO:0031460]
glycine betaine transport [GO:0031460]
glycerol-3-phosphate metabolic process [GO:0006072]; glycerol catabolic process [GO:0019563]; gly
glycerol-3-phosphate catabolic process [GO:0046168]; glycerol catabolic process [GO:0019563]
glycerol transmembrane transport [GO:0015793]
glycerol catabolic process [GO:0019563]; phosphoenolpyruvate-dependent sugar phosphotransferase
glycerol catabolic process [GO:0019563]
glycerol catabolic process [GO:0019563]
glutathione catabolic process [GO:0006751]; lysine biosynthetic process via diaminopimelate [GO:00
glutaminyl-tRNAGln biosynthesis via transamidation [GO:0070681]; regulation of translational fidelity
glutamate biosynthetic process [GO:0006537]; proline catabolic process to glutamate [GO:0010133]
glutamate biosynthetic process [GO:0006537]
glucose metabolic process [GO:0006006]
gluconeogenesis [GO:0006094]; glycolytic process [GO:0006096]
gluconeogenesis [GO:0006094]
gluconeogenesis [GO:0006094]
gluconeogenesis [GO:0006094]
gluconeogenesis [GO:0006094]
gluconate transmembrane transport [GO:0035429]
FtsZ-dependent cytokinesis [GO:0043093]
formate catabolic process [GO:0042183]
folic acid-containing compound biosynthetic process [GO:0009396]; tetrahydrofolate interconversion
folic acid biosynthetic process [GO:0046656]; tetrahydrofolate biosynthetic process [GO:0046654]
fluoride transmembrane transport [GO:1903424]
fluoride transmembrane transport [GO:1903424]
fatty acid metabolic process [GO:0006631]
fatty acid metabolic process [GO:0006631]
fatty acid metabolic process [GO:0006631]
fatty acid metabolic process [GO:0006631]
fatty acid beta-oxidation using acyl-CoA dehydrogenase [GO:0033539]
extracellular polysaccharide biosynthetic process [GO:0045226]
establishment of competence for transformation [GO:0030420]
establishment of competence for transformation [GO:0030420]
enterobacterial common antigen biosynthetic process [GO:0009246]; glucose metabolic process [GO
D-ribose catabolic process [GO:0019303]

D-ribose catabolic process [GO:0019303]
double-strand break repair via homologous recombination [GO:0000724]; response to radiation [GO:
DNA-templated transcription, termination [GO:0006353]; nucleoside metabolic process [GO:000911
DNA-templated transcription, initiation [GO:0006352]; regulation of transcription, DNA-templated [G
DNA-dependent DNA replication [GO:0006261]; error-prone translesion synthesis [GO:0042276]; SO
DNA-dependent DNA replication [GO:0006261]; DNA repair [GO:0006281]
DNA-dependent DNA replication [GO:0006261]
DNA replication, removal of RNA primer [GO:0043137]; mismatch repair [GO:0006298]
DNA replication proofreading [GO:0045004]
DNA replication [GO:0006260]; positive regulation of helicase activity [GO:0051096]
DNA replication [GO:0006260]
DNA recombination [GO:0006310]; DNA replication initiation [GO:0006270]; DNA unwinding involve
DNA recombination [GO:0006310]; DNA repair [GO:0006281]; DNA replication termination [GO:000
DNA recombination [GO:0006310]
DNA methylation [GO:0006306]
DNA mediated transformation [GO:0009294]
DNA integration [GO:0015074]; DNA recombination [GO:0006310]
DNA duplex unwinding [GO:0032508]; DNA recombination [GO:0006310]; DNA repair [GO:0006281]
division septum assembly [GO:0000917]; sporulation resulting in formation of a cellular spore [GO:0
D-gluconate metabolic process [GO:0019521]
dephosphorylation [GO:0016311]; DNA repair [GO:0006281]
deoxyribonucleotide biosynthetic process [GO:0009263]; DNA replication [GO:0006260]
'de novo' UMP biosynthetic process [GO:0044205]; arginine biosynthetic process [GO:0006526]; glut
'de novo' UMP biosynthetic process [GO:0044205]
'de novo' pyrimidine nucleobase biosynthetic process [GO:0006207]; 'de novo' UMP biosynthetic pro
'de novo' pyrimidine nucleobase biosynthetic process [GO:0006207]; 'de novo' UMP biosynthetic pro
cytolysis in other organism [GO:0051715]; pathogenesis [GO:0009405]
cytolysis in other organism [GO:0051715]; pathogenesis [GO:0009405]
cytolysis in other organism [GO:0051715]; pathogenesis [GO:0009405]
cytolysis [GO:0019835]; programmed cell death [GO:0012501]; regulation of single-species biofilm f
cytolysis [GO:0019835]; defense response to bacterium [GO:0042742]
cytolysis [GO:0019835]
cytochrome complex assembly [GO:0017004]; heme transport [GO:0015886]
CTP biosynthetic process [GO:0006241]; GTP biosynthetic process [GO:0006183]; purine nucleotide
C-terminal protein methylation [GO:0006481]
coenzyme A biosynthetic process [GO:0015937]
cobalt ion transport [GO:0006824]; nickel cation transport [GO:0015675]; zinc ion transport [GO:000
cobalt ion transport [GO:0006824]; nickel cation transport [GO:0015675]; zinc ion transport [GO:000
cobalt ion transport [GO:0006824]; nickel cation transport [GO:0015675]; zinc ion transport [GO:000
cobalt ion transport [GO:0006824]; nickel cation transport [GO:0015675]; transmembrane transport [
cobalt ion transport [GO:0006824]; nickel cation transport [GO:0015675]; peptide transport [GO:0015
cellular response to oxidative stress [GO:0034599]
cellular response to iron ion [GO:0071281]
cellular response to heat [GO:0034605]
cellular polysaccharide biosynthetic process [GO:0033692]; lipopolysaccharide biosynthetic process [
cellular iron ion homeostasis [GO:0006879]
cellular amino acid metabolic process [GO:0006520]
cell wall organization [GO:0071555]; teichoic acid biosynthetic process [GO:0019350]
cell wall organization [GO:0071555]; teichoic acid biosynthetic process [GO:0019350]
cell wall organization [GO:0071555]; teichoic acid biosynthetic process [GO:0019350]
cell wall organization [GO:0071555]; teichoic acid biosynthetic process [GO:0019350]
cell wall organization [GO:0071555]; response to antibiotic [GO:0046677]
cell wall organization [GO:0071555]; peptidoglycan biosynthetic process [GO:0009252]; regulation of
cell wall organization [GO:0071555]

cell redox homeostasis [GO:0045454]; response to oxidative stress [GO:0006979]
cell redox homeostasis [GO:0045454]; cellular response to oxidative stress [GO:0034599]; oxygen tr
cell redox homeostasis [GO:0045454]
cell redox homeostasis [GO:0045454]
cell division [GO:0051301]; regulation of cell shape [GO:0008360]
cell cycle [GO:0007049]; cell division [GO:0051301]; cell wall macromolecule biosynthetic process [G
cell adhesion [GO:0007155]; metal ion transport [GO:0030001]
cell adhesion [GO:0007155]
CDP-diacylglycerol biosynthetic process [GO:0016024]
carotenoid biosynthetic process [GO:0016117]; protein N-linked glycosylation [GO:0006487]
carotenoid biosynthetic process [GO:0016117]; pathogenesis [GO:0009405]
carotenoid biosynthetic process [GO:0016117]; pathogenesis [GO:0009405]
carotenoid biosynthetic process [GO:0016117]; pathogenesis [GO:0009405]
carotenoid biosynthetic process [GO:0016117]; cellular aldehyde metabolic process [GO:0006081]; p
carotenoid biosynthetic process [GO:0016117]
cardiolipin biosynthetic process [GO:0032049]
carboxylic acid metabolic process [GO:0019752]; D-amino acid metabolic process [GO:0046416]
carbohydrate transport [GO:0008643]
carbohydrate transmembrane transport [GO:0034219]; phosphoenolpyruvate-dependent sugar phosp
carbohydrate metabolic process [GO:0005975]; negative regulation of transcription, DNA-templated [
carbohydrate metabolic process [GO:0005975]; N-acetylneuraminate catabolic process [GO:001926
carbohydrate metabolic process [GO:0005975]; N-acetylglucosamine catabolic process [GO:000604
carbohydrate metabolic process [GO:0005975]
carbohydrate derivative metabolic process [GO:1901135]; regulation of transcription, DNA-templatec
carbohydrate derivative metabolic process [GO:1901135]; regulation of transcription, DNA-templatec
carbohydrate catabolic process [GO:0016052]; deoxyribonucleotide catabolic process [GO:0009264]
carbohydrate catabolic process [GO:0016052]; deoxyribonucleotide catabolic process [GO:0009264]
CAAX-box protein processing [GO:0071586]
CAAX-box protein processing [GO:0071586]
CAAX-box protein processing [GO:0071586]
branched-chain amino acid catabolic process [GO:0009083]; response to nutrient [GO:0007584]
base-excision repair [GO:0006284]; base-excision repair, AP site formation [GO:0006285]
base-excision repair [GO:0006284]
ATP synthesis coupled electron transport [GO:0042773]; ion transport [GO:0006811]
arginine deiminase pathway [GO:0019546]; carbamoyl phosphate catabolic process [GO:0035975]
arginine deiminase pathway [GO:0019546]; carbamoyl phosphate catabolic process [GO:0035975]
arginine catabolic process to ornithine [GO:0019547]; urea cycle [GO:0000050]
arginine catabolic process to ornithine [GO:0019547]; arginine deiminase pathway [GO:0019546]
arginine catabolic process [GO:0006527]
arginine biosynthetic process via ornithine [GO:0042450]; citrulline biosynthetic process [GO:001924
arginine biosynthetic process via ornithine [GO:0042450]; citrulline biosynthetic process [GO:001924
arginine biosynthetic process [GO:0006526]; proteolysis [GO:0006508]
arginine biosynthetic process [GO:0006526]; protein complex oligomerization [GO:0051259]
arginine biosynthetic process [GO:0006526]; protein complex oligomerization [GO:0051259]
anion transmembrane transport [GO:0098656]; sodium ion transport [GO:0006814]
amino acid activation for nonribosomal peptide biosynthetic process [GO:0043041]; fatty acid metab
alpha-amino acid metabolic process [GO:1901605]; biosynthetic process [GO:0009058]
alcohol metabolic process [GO:0006066]; carbon utilization [GO:0015976]
aggregation of unicellular organisms [GO:0098630]; cell adhesion [GO:0007155]; pathogenesis [GO:
acyl-CoA metabolic process [GO:0006637]; fatty acid biosynthetic process [GO:0006633]
acetoin catabolic process [GO:0045150]
acetoin catabolic process [GO:0045150]
acetoin catabolic process [GO:0045150]
5-phosphoribose 1-diphosphate biosynthetic process [GO:0006015]; cellular metabolic compound sa

;O:0006213]

; to membrane, translocation [GO:0006616]

sodium ion import across plasma membrane [GO:0098719]

scription, DNA-templated [GO:0006355]
scription, DNA-templated [GO:0006355]
scription, DNA-templated [GO:0006355]

:emplated [GO:0006355]
:emplated [GO:0006355]
te biosynthetic process [GO:1900376]

:pression [GO:0010468]

075]; rRNA processing [GO:0006364]; tRNA processing [GO:0008033]

015675]

):0006099]

]

O:0043953]
l transduction [GO:0007165]

):0009089]; threonine biosynthetic process [GO:0009088]
O:0019556]; histidine catabolic process to glutamate and formate [GO:0019557]
O:0019556]; histidine catabolic process to glutamate and formate [GO:0019557]
O:0019556]; histidine catabolic process to glutamate and formate [GO:0019557]

ɔtoporphyrinogen IX biosynthetic process [GO:0006782]

ɔphagic [GO:0032889]

]

ycerol metabolic process [GO:0006071]

e system [GO:0009401]

)09089]; proteolysis [GO:0006508]
/ [GO:0006450]; translation [GO:0006412]

[GO:0035999]

):0006006]

0009314]; SOS response [GO:0009432]
5]
D:0006355]
S response [GO:0009432]

d in DNA replication [GO:0006268]; double-strand break repair [GO:0006302]
5274]; SOS response [GO:0009432]

|
030435]

amine metabolic process [GO:0006541]; nitrogen compound metabolic process [GO:0006807]

cess [GO:0044205]; pyrimidine ribonucleotide biosynthetic process [GO:0009220]
cess [GO:0044205]; cellular amino acid metabolic process [GO:0006520]; nitrogen compound metabolic p

ormation on inanimate substrate [GO:1900231]

metabolic process [GO:0006163]; pyrimidine nucleotide metabolic process [GO:0006220]; UTP biosynthe

5829]
5829]
5829]
GO:0055085]; zinc ion transport [GO:0006829]
5833]; zinc ion transport [GO:0006829]

[GO:0009103]; pathogenesis [GO:0009405]

cell shape [GO:0008360]

ansport [GO:0015671]

GO:0044038]; cell wall organization [GO:0071555]; peptidoglycan biosynthetic process [GO:0009252]; reg

pathogenesis [GO:0009405]

photransferase system [GO:0009401]
[GO:0045892]
2]
6]

d [GO:0006355]
d [GO:0006355]
; deoxyribose phosphate catabolic process [GO:0046386]
; deoxyribose phosphate catabolic process [GO:0046386]

0]
0]

olic process [GO:0006631]; secondary metabolite biosynthetic process [GO:0044550]

0009405]

lvage [GO:0043094]; deoxyribonucleotide catabolic process [GO:0009264] ɔrocess [GO:0006807]

ɛtic process [GO:0006228]

ulation of cell shape [GO:0008360]

| GOBIO UP | #UP | #Down |
|---|---|---|
| pathogenesis [GO:0009405] | | 43 |
| regulation of transcription, DNA-templated [GO:000635 | | 22 |
| transmembrane transport [GO:0055085] | | 19 |
| proteolysis [GO:0006508] | | 18 |
| nickel cation transport [GO:0015675] | | 16 |
| phosphoenolpyruvate-dependent sugar phosphotransfe | | 11 |
| cell wall organization [GO:0071555] | | 8 |
| carotenoid biosynthetic process [GO:0016117] | | 6 |
| negative regulation of transcription, DNA-templated [GC | | 6 |
| cobalt ion transport [GO:0006824] | | 5 |
| DNA recombination [GO:0006310] | | 5 |
| fatty acid metabolic process [GO:0006631] | | 5 |
| gluconeogenesis [GO:0006094] | | 5 |
| glycerol catabolic process [GO:0019563] | | 5 |
| glycolytic process [GO:0006096] | | 5 |
| hemolysis in other organism [GO:0044179] | | 5 |
| nitrogen compound metabolic process [GO:0006807] | | 5 |
| response to antibiotic [GO:0046677] | | 5 |
| tricarboxylic acid cycle [GO:0006099] | | 5 |
| zinc ion transport [GO:0006829] | | 5 |
| arginine biosynthetic process [GO:0006526] | | 4 |
| carbohydrate metabolic process [GO:0005975] | | 4 |
| cell redox homeostasis [GO:0045454] | | 4 |
| 'de novo' UMP biosynthetic process [GO:0044205] | | 4 |
| DNA repair [GO:0006281] | | 4 |
| siderophore-dependent iron import into cell [GO:003321 | | 4 |
| SOS response [GO:0009432] | | 4 |
| teichoic acid biosynthetic process [GO:0019350] | | 4 |
| acetoin catabolic process [GO:0045150] | | 3 |
| arginine deiminase pathway [GO:0019546] | | 3 |
| CAAX-box protein processing [GO:0071586] | | 3 |
| cell adhesion [GO:0007155] | | 3 |
| cytolysis [GO:0019835] | | 3 |
| cytolysis in other organism [GO:0051715] | | 3 |
| deoxyribonucleotide catabolic process [GO:0009264] | | 3 |
| DNA replication [GO:0006260] | | 3 |
| DNA-dependent DNA replication [GO:0006261] | | 3 |
| glycine betaine transport [GO:0031460] | | 3 |
| heme transport [GO:0015886] | | 3 |
| histidine catabolic process [GO:0006548] | | 3 |
| histidine catabolic process to glutamate and formamide | | 3 |
| histidine catabolic process to glutamate and formate [G | | 3 |
| iron import into cell [GO:0033212] | | 3 |
| lipid catabolic process [GO:0016042] | | 3 |
| lipid metabolic process [GO:0006629] | | 3 |
| lipopolysaccharide biosynthetic process [GO:0009103] | | 3 |
| maltodextrin transport [GO:0042956] | | 3 |
| menaquinone biosynthetic process [GO:0009234] | | 3 |
| nucleotide-excision repair [GO:0006289] | | 3 |
| peptide transport [GO:0015833] | | 3 |
| peptidoglycan biosynthetic process [GO:0009252] | | 3 |
| protein autophosphorylation [GO:0046777] | | 3 |
| regulation of cell shape [GO:0008360] | | 3 |
| arginine biosynthetic process via ornithine [GO:004245 | | 2 |

| | |
|---|---|
| arginine catabolic process to ornithine [GO:0019547] | 2 |
| base-excision repair [GO:0006284] | 2 |
| carbamoyl phosphate catabolic process [GO:0035975] | 2 |
| carbohydrate catabolic process [GO:0016052] | 2 |
| carbohydrate derivative metabolic process [GO:190113 | 2 |
| cell division [GO:0051301] | 2 |
| cellular amino acid metabolic process [GO:0006520] | 2 |
| cellular response to oxidative stress [GO:0034599] | 2 |
| citrulline biosynthetic process [GO:0019240] | 2 |
| 'de novo' pyrimidine nucleobase biosynthetic process [G | 2 |
| deoxyribose phosphate catabolic process [GO:0046386 | 2 |
| D-ribose catabolic process [GO:0019303] | 2 |
| establishment of competence for transformation [GO:00 | 2 |
| fluoride transmembrane transport [GO:1903424] | 2 |
| glucose metabolic process [GO:0006006] | 2 |
| glutamate biosynthetic process [GO:0006537] | 2 |
| heme catabolic process [GO:0042167] | 2 |
| ion transport [GO:0006811] | 2 |
| lysine biosynthetic process via diaminopimelate [GO:00 | 2 |
| maltose transport [GO:0015768] | 2 |
| mismatch repair [GO:0006298] | 2 |
| nucleotide catabolic process [GO:0009166] | 2 |
| oligosaccharide catabolic process [GO:0009313] | 2 |
| osmosensory signaling via phosphorelay pathway [GO:( | 2 |
| peptide metabolic process [GO:0006518] | 2 |
| phosphorelay signal transduction system [GO:0000160] | 2 |
| potassium ion transmembrane transport [GO:0071805] | 2 |
| proline catabolic process to glutamate [GO:0010133] | 2 |
| protein complex oligomerization [GO:0051259] | 2 |
| protein lipoylation [GO:0009249] | 2 |
| regulation of DNA-templated transcription [GO:0006355 | 2 |
| response to oxidative stress [GO:0006979] | 2 |
| siderophore biosynthetic process [GO:0019290] | 2 |
| signal transduction [GO:0007165] | 2 |
| sodium ion transport [GO:0006814] | 2 |
| succinate transport [GO:0015744] | 2 |
| tetrahydrofolate interconversion [GO:0035999] | 2 |
| thiamine biosynthetic process [GO:0009228] | 2 |
| thiamine diphosphate biosynthetic process [GO:000922 | 2 |
| translation [GO:0006412] | 2 |
| 5-phosphoribose 1-diphosphate biosynthetic process [G | 1 |
| acyl-CoA metabolic process [GO:0006637] | 1 |
| aggregation of unicellular organisms [GO:0098630] | 1 |
| alcohol metabolic process [GO:0006066] | 1 |
| alpha-amino acid metabolic process [GO:1901605] | 1 |
| amino acid activation for nonribosomal peptide biosynth | 1 |
| anion transmembrane transport [GO:0098656] | 1 |
| arginine catabolic process [GO:0006527] | 1 |
| ATP synthesis coupled electron transport [GO:0042773 | 1 |
| base-excision repair, AP site formation [GO:0006285] | 1 |
| biosynthetic process [GO:0009058] | 1 |
| branched-chain amino acid catabolic process [GO:0009 | 1 |
| carbohydrate transmembrane transport [GO:0034219] | 1 |
| carbohydrate transport [GO:0008643] | 1 |
| carbon utilization [GO:0015976] | 1 |

| | |
|---|---|
| carboxylic acid metabolic process [GO:0019752] | 1 |
| cardiolipin biosynthetic process [GO:0032049] | 1 |
| CDP-diacylglycerol biosynthetic process [GO:0016024] | 1 |
| cell cycle [GO:0007049] | 1 |
| cell wall macromolecule biosynthetic process [GO:0044 | 1 |
| cellular aldehyde metabolic process [GO:0006081] | 1 |
| cellular iron ion homeostasis [GO:0006879] | 1 |
| cellular metabolic compound salvage [GO:0043094] | 1 |
| cellular polysaccharide biosynthetic process [GO:00336 | 1 |
| cellular response to heat [GO:0034605] | 1 |
| cellular response to iron ion [GO:0071281] | 1 |
| coenzyme A biosynthetic process [GO:0015937] | 1 |
| C-terminal protein methylation [GO:0006481] | 1 |
| CTP biosynthetic process [GO:0006241] | 1 |
| cytochrome complex assembly [GO:0017004] | 1 |
| D-amino acid metabolic process [GO:0046416] | 1 |
| defense response to bacterium [GO:0042742] | 1 |
| deoxyribonucleotide biosynthetic process [GO:0009263 | 1 |
| dephosphorylation [GO:0016311] | 1 |
| D-gluconate metabolic process [GO:0019521] | 1 |
| division septum assembly [GO:0000917] | 1 |
| DNA duplex unwinding [GO:0032508] | 1 |
| DNA integration [GO:0015074] | 1 |
| DNA mediated transformation [GO:0009294] | 1 |
| DNA methylation [GO:0006306] | 1 |
| DNA replication initiation [GO:0006270] | 1 |
| DNA replication proofreading [GO:0045004] | 1 |
| DNA replication termination [GO:0006274] | 1 |
| DNA replication, removal of RNA primer [GO:0043137] | 1 |
| DNA unwinding involved in DNA replication [GO:00062 | 1 |
| DNA-templated transcription, initiation [GO:0006352] | 1 |
| DNA-templated transcription, termination [GO:0006353] | 1 |
| double-strand break repair [GO:0006302] | 1 |
| double-strand break repair via homologous recombinati | 1 |
| enterobacterial common antigen biosynthetic process [ | 1 |
| error-prone translesion synthesis [GO:0042276] | 1 |
| extracellular polysaccharide biosynthetic process [GO:0 | 1 |
| fatty acid beta-oxidation using acyl-CoA dehydrogenase | 1 |
| fatty acid biosynthetic process [GO:0006633] | 1 |
| folic acid biosynthetic process [GO:0046656] | 1 |
| folic acid-containing compound biosynthetic process [G | 1 |
| formate catabolic process [GO:0042183] | 1 |
| FtsZ-dependent cytokinesis [GO:0043093] | 1 |
| gluconate transmembrane transport [GO:0035429] | 1 |
| glutamine metabolic process [GO:0006541] | 1 |
| glutaminyl-tRNAGln biosynthesis via transamidation [G | 1 |
| glutathione catabolic process [GO:0006751] | 1 |
| glycerol metabolic process [GO:0006071] | 1 |
| glycerol transmembrane transport [GO:0015793] | 1 |
| glycerol-3-phosphate catabolic process [GO:0046168] | 1 |
| glycerol-3-phosphate metabolic process [GO:0006072] | 1 |
| glycine decarboxylation via glycine cleavage system [G | 1 |
| glyoxylate cycle [GO:0006097] | 1 |
| GTP biosynthetic process [GO:0006183] | 1 |
| guanosine tetraphosphate biosynthetic process [GO:00 | 1 |

| | |
|---|---|
| heme biosynthetic process [GO:0006783] | 1 |
| hemolysis by symbiont of host erythrocytes [GO:00198: | 1 |
| homoserine biosynthetic process [GO:0009090] | 1 |
| hydrogen peroxide catabolic process [GO:0042744] | 1 |
| inositol metabolic process [GO:0006020] | 1 |
| inositol phosphate dephosphorylation [GO:0046855] | 1 |
| intracellular protein transmembrane transport [GO:0065 | 1 |
| intracellular sequestering of iron ion [GO:0006880] | 1 |
| iron ion homeostasis [GO:0055072] | 1 |
| iron ion transmembrane transport [GO:0034755] | 1 |
| iron ion transport [GO:0006826] | 1 |
| iron-sulfur cluster assembly [GO:0016226] | 1 |
| ketone body catabolic process [GO:0046952] | 1 |
| lactose metabolic process [GO:0005988] | 1 |
| L-arabinose catabolic process to xylulose 5-phosphate | | 1 |
| lipoprotein biosynthetic process [GO:0042158] | 1 |
| L-lysine catabolic process to acetyl-CoA via saccharopi | 1 |
| L-phenylalanine biosynthetic process [GO:0009094] | 1 |
| lysine biosynthetic process via aminoadipic acid [GO:00 | 1 |
| mannitol catabolic process [GO:0019592] | 1 |
| metabolic process [GO:0008152] | 1 |
| metal ion transport [GO:0030001] | 1 |
| methylation [GO:0032259] | 1 |
| molybdate ion transport [GO:0015689] | 1 |
| mRNA processing [GO:0006397] | 1 |
| N-acetylglucosamine catabolic process [GO:0006046] | 1 |
| N-acetylneuraminate catabolic process [GO:0019262] | 1 |
| NAD metabolic process [GO:0019674] | 1 |
| NADP biosynthetic process [GO:0006741] | 1 |
| negative regulation of DNA-templated transcription [GO | 1 |
| negative regulation of DNA-templated transcription, initi | 1 |
| negative regulation of DNA-templated transcription, tern | 1 |
| nitric oxide biosynthetic process [GO:0006809] | 1 |
| nitrogen compound transport [GO:0071705] | 1 |
| nucleoside metabolic process [GO:0009116] | 1 |
| nucleoside transmembrane transport [GO:1901642] | 1 |
| organonitrogen compound catabolic process [GO:1901! | 1 |
| oxygen transport [GO:0015671] | 1 |
| peptide catabolic process [GO:0043171] | 1 |
| peptide modification [GO:0031179] | 1 |
| peptidyl-pyrromethane cofactor linkage [GO:0018160] | 1 |
| phosphate ion transmembrane transport [GO:0035435] | 1 |
| phospholipid biosynthetic process [GO:0008654] | 1 |
| phosphorylation [GO:0016310] | 1 |
| polysaccharide transport [GO:0015774] | 1 |
| positive regulation of helicase activity [GO:0051096] | 1 |
| programmed cell death [GO:0012501] | 1 |
| protein biotinylation [GO:0009305] | 1 |
| protein dephosphorylation [GO:0006470] | 1 |
| protein folding [GO:0006457] | 1 |
| protein initiator methionine removal [GO:0070084] | 1 |
| protein N-linked glycosylation [GO:0006487] | 1 |
| protein transport by the Sec complex [GO:0043952] | 1 |
| protein transport by the Tat complex [GO:0043953] | 1 |
| protein unfolding [GO:0043335] | 1 |

| | | |
|---|---|---|
| protoporphyrinogen IX biosynthetic process [GO:00067 | | 1 |
| purine nucleoside catabolic process [GO:0006152] | | 1 |
| purine nucleotide metabolic process [GO:0006163] | | 1 |
| pyrimidine nucleobase metabolic process [GO:0006206 | | 1 |
| pyrimidine nucleoside metabolic process [GO:0006213] | | 1 |
| pyrimidine nucleotide metabolic process [GO:0006220] | | 1 |
| pyrimidine ribonucleotide biosynthetic process [GO:000 | | 1 |
| quorum sensing [GO:0009372] | | 1 |
| regulation of DNA repair [GO:0006282] | | 1 |
| regulation of gene expression [GO:0010468] | | 1 |
| regulation of intracellular pH [GO:0051453] | | 1 |
| regulation of secondary metabolite biosynthetic process | | 1 |
| regulation of single-species biofilm formation on inanim | | 1 |
| regulation of translational fidelity [GO:0006450] | | 1 |
| regulation of vacuole fusion, non-autophagic [GO:0032 | | 1 |
| response to hydrogen peroxide [GO:0042542] | | 1 |
| response to nutrient [GO:0007584] | | 1 |
| response to radiation [GO:0009314] | | 1 |
| response to reactive oxygen species [GO:0000302] | | 1 |
| response to toxic substance [GO:0009636] | | 1 |
| response to zinc ion [GO:0010043] | | 1 |
| RNA processing [GO:0006396] | | 1 |
| rRNA catabolic process [GO:0016075] | | 1 |
| rRNA processing [GO:0006364] | | 1 |
| secondary metabolite biosynthetic process [GO:004455 | | 1 |
| signal peptide processing [GO:0006465] | | 1 |
| sodium ion import across plasma membrane [GO:0098 | | 1 |
| sporulation resulting in formation of a cellular spore [GC | | 1 |
| SRP-dependent cotranslational protein targeting to men | | 1 |
| steroid biosynthetic process [GO:0006694] | | 1 |
| sulfide oxidation, using sulfide:quinone oxidoreductase | | 1 |
| tetrahydrofolate biosynthetic process [GO:0046654] | | 1 |
| thiamine transport [GO:0015888] | | 1 |
| threonine biosynthetic process [GO:0009088] | | 1 |
| threonine catabolic process [GO:0006567] | | 1 |
| transposition, DNA-mediated [GO:0006313] | | 1 |
| trehalose catabolic process [GO:0005993] | | 1 |
| tRNA modification [GO:0006400] | | 1 |
| tRNA processing [GO:0008033] | | 1 |
| UDP-N-acetylglucosamine biosynthetic process [GO:00 | | 1 |
| urea cycle [GO:0000050] | | 1 |
| UTP biosynthetic process [GO:0006228] | | 1 |
| zinc ion transmembrane transport [GO:0071577] | | 1 |
| pathogenesis [GO:0009405] | 23 | |
| cell wall organization [GO:0071555] | 21 | |
| regulation of transcription, DNA-template | 18 | |
| peptidoglycan biosynthetic process [GO:0 | 15 | |
| transmembrane transport [GO:0055085] | 15 | |
| regulation of cell shape [GO:0008360] | 14 | |
| translation [GO:0006412] | 13 | |
| response to antibiotic [GO:0046677] | 11 | |
| cell division [GO:0051301] | 10 | |
| phosphoenolpyruvate-dependent sugar p | 9 | |
| queuosine biosynthetic process [GO:000 | 8 | |
| nickel cation transport [GO:0015675] | 7 | |

| | |
|---|---|
| DNA repair [GO:0006281] | 7 |
| cell cycle [GO:0007049] | 6 |
| amino acid transport [GO:0006865] | 6 |
| fatty acid biosynthetic process [GO:0006[illegible] | 6 |
| chromosome segregation [GO:0007059] | 6 |
| isoleucine biosynthetic process [GO:000[illegible] | 6 |
| histidine biosynthetic process [GO:00001[illegible] | 6 |
| DNA topological change [GO:0006265] | 6 |
| DNA replication [GO:0006260] | 5 |
| DNA recombination [GO:0006310] | 5 |
| ATP synthesis coupled proton transport [[illegible] | 5 |
| cytolysis [GO:0019835] | 5 |
| negative regulation of transcription, DNA-[illegible] | 5 |
| peptide transport [GO:0015833] | 5 |
| pantothenate biosynthetic process [GO:0[illegible] | 5 |
| leucine biosynthetic process [GO:000909[illegible] | 5 |
| SOS response [GO:0009432] | 5 |
| aerobic electron transport chain [GO:001[illegible] | 4 |
| DNA-dependent DNA replication [GO:000[illegible] | 4 |
| ribosomal large subunit assembly [GO:00[illegible] | 4 |
| menaquinone biosynthetic process [GO:0[illegible] | 4 |
| 'de novo' IMP biosynthetic process [GO:0[illegible] | 4 |
| branched-chain amino acid biosynthetic [illegible] | 4 |
| cell adhesion [GO:0007155] | 4 |
| division septum assembly [GO:0000917] | 4 |
| metabolic process [GO:0008152] | 4 |
| Mo-molybdopterin cofactor biosynthetic p[illegible] | 4 |
| phospholipid biosynthetic process [GO:00[illegible] | 4 |
| protein folding [GO:0006457] | 4 |
| proteolysis [GO:0006508] | 4 |
| riboflavin biosynthetic process [GO:0009[illegible] | 4 |
| siderophore-dependent iron import into c[illegible] | 4 |
| transcription, DNA-templated [GO:00063[illegible] | 4 |
| lysine biosynthetic process via diaminopi[illegible] | 4 |
| valine biosynthetic process [GO:0009099[illegible] | 4 |
| carbohydrate metabolic process [GO:000[illegible] | 3 |
| aromatic amino acid family biosynthetic p[illegible] | 3 |
| chaperone cofactor-dependent protein re[illegible] | 3 |
| base-excision repair [GO:0006284] | 3 |
| glutamine metabolic process [GO:00065[illegible] | 3 |
| diaminopimelate biosynthetic process [G[illegible] | 3 |
| methylation [GO:0032259] | 3 |
| stress response to cadmium ion [GO:199[illegible] | 3 |
| tRNA methylation [GO:0030488] | 3 |
| biosynthetic process [GO:0009058] | 3 |
| cell redox homeostasis [GO:0045454] | 3 |
| glucose metabolic process [GO:0006006[illegible] | 3 |
| lipid A biosynthetic process [GO:0009245[illegible] | 3 |
| phosphate ion transmembrane transport [illegible] | 3 |
| phosphate ion transport [GO:0006817] | 3 |
| response to oxidative stress [GO:000697[illegible] | 3 |
| rRNA base methylation [GO:0070475] | 3 |
| tRNA modification [GO:0006400] | 3 |
| cellular amino acid biosynthetic process [[illegible] | 3 |
| positive regulation of sporulation resulting [illegible] | 3 |

| | |
|---|---|
| protein refolding [GO:0042026] | 3 |
| RNA catabolic process [GO:0006401] | 3 |
| stress response to copper ion [GO:19901 | 3 |
| tRNA processing [GO:0008033] | 3 |
| chorismate biosynthetic process [GO:000 | 3 |
| defense response to bacterium [GO:0042 | 3 |
| threonine biosynthetic process [GO:0009 | 3 |
| thiamine biosynthetic process [GO:00092 | 2 |
| mRNA catabolic process [GO:0006402] | 2 |
| purine ribonucleoside salvage [GO:00061 | 2 |
| GMP biosynthetic process [GO:0006177] | 2 |
| DNA dealkylation involved in DNA repair | 2 |
| anaerobic respiration [GO:0009061] | 2 |
| protein catabolic process [GO:0030163] | 2 |
| DNA unwinding involved in DNA replicatio | 2 |
| ATP synthesis coupled electron transport | 2 |
| ribosomal small subunit assembly [GO:00 | 2 |
| acetoin biosynthetic process [GO:004515 | 2 |
| amino acid transmembrane transport [GO | 2 |
| glycine betaine biosynthetic process from | 2 |
| glycolytic process [GO:0006096] | 2 |
| iron-sulfur cluster assembly [GO:0016226 | 2 |
| nitrogen compound transport [GO:00717( | 2 |
| nucleobase-containing small molecule int | 2 |
| organic substance metabolic process [GO | 2 |
| osmosensory signaling via phosphorelay | 2 |
| plasma membrane ATP synthesis couple | 2 |
| response to arsenic-containing substance | 2 |
| ribosomal small subunit biogenesis [GO:( | 2 |
| RNA processing [GO:0006396] | 2 |
| rRNA processing [GO:0006364] | 2 |
| SRP-dependent cotranslational protein ta | 2 |
| succinate transport [GO:0015744] | 2 |
| tricarboxylic acid cycle [GO:0006099] | 2 |
| tRNA methylthiolation [GO:0035600] | 2 |
| tRNA pseudouridine synthesis [GO:0031 | 2 |
| tryptophan biosynthetic process [GO:000 | 2 |
| UDP-N-acetylglucosamine biosynthetic pr | 2 |
| electron transport coupled proton transpo | 2 |
| FtsZ-dependent cytokinesis [GO:004309 | 2 |
| methionine biosynthetic process [GO:000 | 2 |
| mismatch repair [GO:0006298] | 2 |
| protein transport [GO:0015031] | 2 |
| pyridoxal phosphate biosynthetic process | 2 |
| sodium ion transport [GO:0006814] | 2 |
| tRNA wobble guanine modification [GO:0 | 2 |
| tRNA wobble uridine modification [GO:00 | 2 |
| response to heat [GO:0009408] | 2 |
| 'de novo' cotranslational protein folding [G | 1 |
| cell morphogenesis [GO:0000902] | 1 |
| dATP catabolic process [GO:0046061] | 1 |
| DNA duplex unwinding [GO:0032508] | 1 |
| 'de novo' L-methionine biosynthetic proce | 1 |
| butanediol metabolic process [GO:00340 | 1 |
| co-translational protein modification [GO: | 1 |

| | |
|---|---|
| FAD biosynthetic process [GO:0006747] | 1 |
| 'de novo' CTP biosynthetic process [GO:( | 1 |
| dTDP biosynthetic process [GO:0006233 | 1 |
| canonical glycolysis [GO:0061621] | 1 |
| cellular respiration [GO:0045333] | 1 |
| mature ribosome assembly [GO:0042256 | 1 |
| L-methionine salvage from methylthioade | 1 |
| cellular phosphate ion homeostasis [GO:( | 1 |
| intracellular protein transmembrane trans | 1 |
| adenine salvage [GO:0006168] | 1 |
| cellular amino acid metabolic process [G( | 1 |
| glutamine catabolic process [GO:000654 | 1 |
| protein repair [GO:0030091] | 1 |
| response to cold [GO:0009409] | 1 |
| selenocysteine biosynthetic process [GO: | 1 |
| potassium ion transmembrane transport [ | 1 |
| phosphocreatine biosynthetic process [G( | 1 |
| homoserine biosynthetic process [GO:00( | 1 |
| L-serine catabolic process [GO:0006565] | 1 |
| maturation of SSU-rRNA from tricistronic | 1 |
| mRNA pseudouridine synthesis [GO:199( | 1 |
| isoleucine transport [GO:0015818] | 1 |
| base-excision repair, DNA ligation [GO:0( | 1 |
| DNA replication, synthesis of RNA primer | 1 |
| acetyl-CoA metabolic process [GO:0006( | 1 |
| acyl-CoA metabolic process [GO:000663 | 1 |
| galactose catabolic process via UDP-gala | 1 |
| ammonia assimilation cycle [GO:0019676 | 1 |
| acetyl-CoA biosynthetic process from pyr | 1 |
| 'de novo' AMP biosynthetic process [GO:( | 1 |
| carbohydrate catabolic process [GO:0016 | 1 |
| D-tagatose 6-phosphate catabolic proces | 1 |
| alanine catabolic process [GO:0006524] | 1 |
| arginine metabolic process [GO:0006525 | 1 |
| homoserine metabolic process [GO:0009 | 1 |
| DNA replication, removal of RNA primer [ | 1 |
| metabolite repair [GO:0110051] | 1 |
| p-aminobenzoyl-glutamate transmembrar | 1 |
| alanine biosynthetic process [GO:000652 | 1 |
| fatty acid beta-oxidation [GO:0006635] | 1 |
| glucose-6-phosphate transport [GO:0015 | 1 |
| proline transmembrane transport [GO:00: | 1 |
| cellular response to heat [GO:0034605] | 1 |
| protein insertion into membrane [GO:005 | 1 |
| porphyrin-containing compound biosynthe | 1 |
| DNA-templated transcription, elongation [ | 1 |
| protein glycosylation [GO:0006486] | 1 |
| phosphorelay signal transduction system | 1 |
| mRNA processing [GO:0006397] | 1 |
| one-carbon metabolic process [GO:0006 | 1 |
| proton transmembrane transport [GO:19( | 1 |
| error-prone translesion synthesis [GO:00 | 1 |
| protein targeting [GO:0006605] | 1 |
| folic acid biosynthetic process [GO:0046( | 1 |
| DNA biosynthetic process [GO:0071897] | 1 |

| | |
|---|---|
| DNA integration [GO:0015074] | 1 |
| cysteine biosynthetic process via cystathi | 1 |
| trans-translation [GO:0070929] | 1 |
| tRNA aminoacylation [GO:0043039] | 1 |
| CTP salvage [GO:0044211] | 1 |
| wobble position cytosine ribose methylati | 1 |
| [2Fe-2S] cluster assembly [GO:0044571] | 1 |
| 2,4,6-trinitrotoluene catabolic process [G | 1 |
| 7,8-dihydroneopterin 3'-triphosphate bios | 1 |
| asparagine catabolic process [GO:00065 | 1 |
| asparaginyl-tRNA aminoacylation [GO:00 | 1 |
| aspartyl-tRNA aminoacylation [GO:00064 | 1 |
| CAAX-box protein processing [GO:00715 | 1 |
| cell adhesion involved in biofilm formatior | 1 |
| cellular response to osmotic stress [GO:0 | 1 |
| coenzyme A biosynthetic process [GO:00 | 1 |
| cyclic threonylcarbamoyladenosine biosy | 1 |
| DNA packaging [GO:0006323] | 1 |
| DNA replication proofreading [GO:00450 | 1 |
| DNA replication, Okazaki fragment proce | 1 |
| DNA-templated transcription, initiation [G | 1 |
| DNA-templated transcription, termination | 1 |
| enzyme-directed rRNA pseudouridine syr | 1 |
| establishment of competence for transfor | 1 |
| fatty acid elongation [GO:0030497] | 1 |
| folic acid catabolic process [GO:0046657 | 1 |
| folic acid-containing compound biosynthe | 1 |
| formate metabolic process [GO:0015942 | 1 |
| formate transport [GO:0015724] | 1 |
| glutamine biosynthetic process [GO:0006 | 1 |
| glycerol metabolic process [GO:0006071 | 1 |
| glycine decarboxylation via glycine cleava | 1 |
| glycyl-tRNA aminoacylation [GO:0006426 | 1 |
| guanosine tetraphosphate biosynthetic pr | 1 |
| iron ion transport [GO:0006826] | 1 |
| isoprenoid biosynthetic process [GO:000 | 1 |
| leucyl-tRNA aminoacylation [GO:0006429 | 1 |
| L-glutamate transmembrane transport [G | 1 |
| lipoteichoic acid biosynthetic process [GC | 1 |
| L-valine transmembrane transport [GO:1 | 1 |
| lysyl-tRNA aminoacylation [GO:0006430] | 1 |
| metal ion transport [GO:0030001] | 1 |
| methionyl-tRNA aminoacylation [GO:000 | 1 |
| NAD biosynthetic process [GO:0009435] | 1 |
| nitrate assimilation [GO:0042128] | 1 |
| nitrogen utilization [GO:0019740] | 1 |
| N-terminal peptidyl-serine acetylation [GC | 1 |
| oligopeptide transport [GO:0006857] | 1 |
| peptidyl-glutamine methylation [GO:0018 | 1 |
| phenylalanyl-tRNA aminoacylation [GO:0 | 1 |
| phosphorylation [GO:0016310] | 1 |
| polyamine transport [GO:0015846] | 1 |
| polyphosphate catabolic process [GO:00 | 1 |
| potassium ion transport [GO:0006813] | 1 |
| protein lipoylation [GO:0009249] | 1 |

| | |
|---|---|
| protein O-linked glycosylation via serine [ | 1 |
| protein peptidyl-prolyl isomerization [GO:( | 1 |
| purine nucleotide metabolic process [GO | 1 |
| pyrimidine deoxyribonucleotide catabolic | 1 |
| queuosine salvage [GO:1990397] | 1 |
| regulation of DNA-templated transcriptior | 1 |
| respiratory electron transport chain [GO:( | 1 |
| riboflavin transport [GO:0032218] | 1 |
| ribosome biogenesis [GO:0042254] | 1 |
| rRNA methylation [GO:0031167] | 1 |
| signal peptide processing [GO:0006465] | 1 |
| sodium-dependent phosphate transport [( | 1 |
| threonyl-tRNA aminoacylation [GO:00064 | 1 |
| transcription initiation from bacterial-type | 1 |
| translational elongation [GO:0006414] | 1 |
| tryptophanyl-tRNA aminoacylation [GO:0( | 1 |
| valyl-tRNA aminoacylation [GO:0006438] | 1 |
| xanthine transport [GO:0042906] | 1 |
| xenobiotic detoxification by transmembra | 1 |
| zinc ion transmembrane transport [GO:0( | 1 |
| AMP salvage [GO:0044209] | 1 |
| cobalamin biosynthetic process [GO:000( | 1 |
| CTP biosynthetic process [GO:0006241] | 1 |
| dephosphorylation [GO:0016311] | 1 |
| dGTP catabolic process [GO:0006203] | 1 |
| D-methionine transport [GO:0048473] | 1 |
| DNA synthesis involved in DNA repair [G( | 1 |
| dTTP biosynthetic process [GO:000623( | 1 |
| farnesyl diphosphate biosynthetic proces: | 1 |
| fatty acid metabolic process [GO:000663 | 1 |
| FMN biosynthetic process [GO:0009398] | 1 |
| fructose 1,6-bisphosphate metabolic proc | 1 |
| glutamate biosynthetic process [GO:0006 | 1 |
| glycerol-3-phosphate biosynthetic proces | 1 |
| GTP biosynthetic process [GO:0006183] | 1 |
| heme biosynthetic process [GO:000678( | 1 |
| IMP metabolic process [GO:0046040] | 1 |
| lactose catabolic process via tagatose-6-( | 1 |
| lactose metabolic process [GO:0005988] | 1 |
| L-alanine catabolic process [GO:004285( | 1 |
| L-arginine transport [GO:1902023] | 1 |
| leucine transport [GO:0015820] | 1 |
| L-methionine salvage from S-adenosylme | 1 |
| L-proline biosynthetic process [GO:00551 | 1 |
| L-threonine catabolic process to propiona | 1 |
| negative regulation of fatty acid biosynthe | 1 |
| negative regulation of phosphate metabo | 1 |
| negative regulation of ribosome biogenes | 1 |
| nicotinamide nucleotide metabolic proces | 1 |
| nitrate metabolic process [GO:0042126] | 1 |
| N-terminal protein amino acid modificatio | 1 |
| p-aminobenzoyl-glutamate transport [GO | 1 |
| peptidoglycan catabolic process [GO:000 | 1 |
| phenylacetate catabolic process [GO:001 | 1 |
| programmed cell death [GO:0012501] | 1 |

| | |
|---|---|
| proline transport [GO:0015824] | 1 |
| protein secretion [GO:0009306] | 1 |
| protoporphyrinogen IX biosynthetic proce | 1 |
| purine nucleobase biosynthetic process [ | 1 |
| recombinational repair [GO:0000725] | 1 |
| regulation of DNA-templated transcription | 1 |
| regulation of intracellular pH [GO:005145 | 1 |
| regulation of protein stability [GO:003164 | 1 |
| response to nitrosative stress [GO:00514 | 1 |
| response to redox state [GO:0051775] | 1 |
| S-adenosylmethionine biosynthetic proce | 1 |
| selenocysteinyl-tRNA(Sec) biosynthetic p | 1 |
| single-species submerged biofilm formati | 1 |
| siroheme biosynthetic process [GO:0019 | 1 |
| tetrahydrofolate biosynthetic process [GC | 1 |
| thiamine diphosphate biosynthetic proces | 1 |
| thymidine metabolic process [GO:004610 | 1 |
| transposition [GO:0032196] | 1 |
| transsulfuration [GO:0019346] | 1 |
| trans-translation-dependent protein taggi | 1 |
| tRNA-guanine transglycosylation [GO:010 | 1 |
| tyrosyl-tRNA aminoacylation [GO:000643 | 1 |
| ubiquinone biosynthetic process via 3,4-d | 1 |
| UMP salvage [GO:0044206] | 1 |
| wobble position uridine ribose methylation | 1 |
| xanthine metabolic process [GO:0046110 | 1 |
| zinc ion transport [GO:0006829] | 1 |
| double-strand break repair [GO:0006302] | 1 |
| dTTP catabolic process [GO:0046076] | 1 |
| dUDP biosynthetic process [GO:0006227 | 1 |
| fructose 6-phosphate metabolic process [ | 1 |
| glycerol-3-phosphate catabolic process [( | 1 |
| heme O biosynthetic process [GO:00480 | 1 |
| negative regulation of translation [GO:001 | 1 |
| nucleoside catabolic process [GO:000916 | 1 |
| peptide catabolic process [GO:0043171] | 1 |
| peptidyl-methionine modification [GO:001 | 1 |
| positive regulation of membrane potential | 1 |
| protein transport by the Sec complex [GC | 1 |
| pyridoxine biosynthetic process [GO:0008 | 1 |
| pyridoxine metabolic process [GO:00086 | 1 |
| RNA secondary structure unwinding [GO: | 1 |
| seryl-tRNA aminoacylation [GO:0006434] | 1 |
| siderophore biosynthetic process [GO:00 | 1 |
| sodium ion import across plasma membr | 1 |
| thiazole biosynthetic process [GO:005283 | 1 |
| threonine catabolic process [GO:0006567 | 1 |
| valine transport [GO:0015829] | 1 |
| vitamin K biosynthetic process [GO:0042 | 1 |
| XMP salvage [GO:0032265] | 1 |
| tRNA 4-thiouridine biosynthesis [GO:0002 | 1 |
| shikimate metabolic process [GO:001963 | 1 |
| riboflavin metabolic process [GO:000677 | 1 |
| pyrimidine nucleobase biosynthetic proce | 1 |
| protein polymerization [GO:0051258] | 1 |

| | | |
|---|---|---|
| positive regulation of helicase activity [GC | 1 | |
| glycerol-3-phosphate metabolic process [ | 1 | |
| dUTP catabolic process [GO:0046081] | 1 | |
| chaperone-mediated protein folding [GO: | 1 | |
| glycerophospholipid metabolic process [G | 1 | |
| lipid glycosylation [GO:0030259] | 1 | |
| TTP catabolic process [GO:0046047] | 1 | |
| protein unfolding [GO:0043335] | 1 | |
| UDP-N-acetylgalactosamine biosynthetic | 1 | |
| UTP catabolic process [GO:0046052] | 1 | |
| Suma | 730 | 561 |

| GOBIO UP | #UP | %up | #Down |
|---|---|---|---|
| pathogenesis [GO:0009405] | 23 | 3.15 | 43 |
| regulation of transcription, DNA-templated [GO:0006355] | 18 | 2.47 | 22 |
| transmembrane transport [GO:0055085] | 15 | 2.05 | 19 |
| proteolysis [GO:0006508] | 4 | 0.55 | 18 |
| nickel cation transport [GO:0015675] | 7 | 0.96 | 16 |
| phosphoenolpyruvate-dependent sugar phosphotransferase system | 9 | 1.23 | 11 |
| cell wall organization [GO:0071555] | 21 | 2.88 | 8 |
| negative regulation of transcription, DNA-templated [GO:0045892] | 5 | 0.68 | 6 |
| carotenoid biosynthetic process [GO:0016117] | 0 | 0.00 | 6 |
| response to antibiotic [GO:0046677] | 11 | 1.51 | 5 |
| DNA recombination [GO:0006310] | 5 | 0.68 | 5 |
| cobalt ion transport [GO:0006824] | 0 | 0.00 | 5 |
| fatty acid metabolic process [GO:0006631] | 0 | 0.00 | 5 |
| gluconeogenesis [GO:0006094] | 0 | 0.00 | 5 |
| glycerol catabolic process [GO:0019563] | 0 | 0.00 | 5 |
| glycolytic process [GO:0006096] | 0 | 0.00 | 5 |
| hemolysis in other organism [GO:0044179] | 0 | 0.00 | 5 |
| nitrogen compound metabolic process [GO:0006807] | 0 | 0.00 | 5 |
| tricarboxylic acid cycle [GO:0006099] | 0 | 0.00 | 5 |
| zinc ion transport [GO:0006829] | 0 | 0.00 | 5 |
| DNA repair [GO:0006281] | 7 | 0.96 | 4 |
| SOS response [GO:0009432] | 5 | 0.68 | 4 |
| 'de novo' IMP biosynthetic process [GO:0006189] | 4 | 0.55 | 4 |
| siderophore-dependent iron import into cell [GO:0033214] | 4 | 0.55 | 4 |
| arginine biosynthetic process [GO:0006526] | 0 | 0.00 | 4 |
| carbohydrate metabolic process [GO:0005975] | 0 | 0.00 | 4 |
| cell redox homeostasis [GO:0045454] | 0 | 0.00 | 4 |
| teichoic acid biosynthetic process [GO:0019350] | 0 | 0.00 | 4 |
| peptidoglycan biosynthetic process [GO:0009252] | 15 | 2.05 | 0 |
| regulation of cell shape [GO:0008360] | 14 | 1.92 | 0 |
| translation [GO:0006412] | 13 | 1.78 | 0 |
| cell division [GO:0051301] | 10 | 1.37 | 0 |
| queuosine biosynthetic process [GO:0008616] | 8 | 1.10 | 0 |
| amino acid transport [GO:0006865] | 6 | 0.82 | 0 |
| cell cycle [GO:0007049] | 6 | 0.82 | 0 |
| chromosome segregation [GO:0007059] | 6 | 0.82 | 0 |
| DNA topological change [GO:0006265] | 6 | 0.82 | 0 |
| fatty acid biosynthetic process [GO:0006633] | 6 | 0.82 | 0 |
| histidine biosynthetic process [GO:0000105] | 6 | 0.82 | 0 |
| isoleucine biosynthetic process [GO:0009097] | 6 | 0.82 | 0 |
| ATP synthesis coupled proton transport [GO:0015986] | 5 | 0.68 | 0 |
| cytolysis [GO:0019835] | 5 | 0.68 | 0 |
| DNA replication [GO:0006260] | 5 | 0.68 | 0 |
| leucine biosynthetic process [GO:0009098] | 5 | 0.68 | 0 |
| pantothenate biosynthetic process [GO:0015940] | 5 | 0.68 | 0 |
| peptide transport [GO:0015833] | 5 | 0.68 | 0 |
| aerobic electron transport chain [GO:0019646] | 4 | 0.55 | 0 |
| branched-chain amino acid biosynthetic process [GO:0009082] | 4 | 0.55 | 0 |
| cell adhesion [GO:0007155] | 5 | 0.68 | 0 |
| division septum assembly [GO:0000917] | 4 | 0.55 | 0 |
| DNA-dependent DNA replication [GO:0006261] | 4 | 0.55 | 0 |
| lysine biosynthetic process via diaminopimelate [GO:0009089] | 4 | 0.55 | 0 |
| menaquinone biosynthetic process [GO:0009234] | 4 | 0.55 | 0 |
| metabolic process [GO:0008152] | 4 | 0.55 | 0 |

| | | | |
|---|---|---|---|
| Mo-molybdopterin cofactor biosynthetic process [GO:0006777] | 4 | 0.55 | 0 |
| phospholipid biosynthetic process [GO:0008654] | 4 | 0.55 | 0 |
| protein folding [GO:0006457] | 4 | 0.55 | 0 |
| riboflavin biosynthetic process [GO:0009231] | 4 | 0.55 | 0 |
| ribosomal large subunit assembly [GO:0000027] | 4 | 0.55 | 0 |
| transcription, DNA-templated [GO:0006351] | 4 | 0.55 | 0 |
| | 327 | | |

| %Down |
|---|
| 7.66 |
| 3.92 |
| 3.39 |
| 3.21 |
| 2.85 |
| 1.96 |
| 1.43 |
| 1.07 |
| 1.07 |
| 0.89 |
| 0.89 |
| 0.89 |
| 0.89 |
| 0.89 |
| 0.89 |
| 0.89 |
| 0.89 |
| 0.89 |
| 0.89 |
| 0.89 |
| 0.71 |
| 0.71 |
| 0.71 |
| 0.71 |
| 0.71 |
| 0.71 |
| 0.71 |
| 0.71 |
| 0.00 |
| 0.00 |
| 0.00 |
| 0.00 |
| 0.00 |
| 0.00 |
| 0.00 |
| 0.00 |
| 0.00 |
| 0.00 |
| 0.00 |
| 0.00 |
| 0.00 |
| 0.00 |
| 0.00 |
| 0.00 |
| 0.00 |
| 0.00 |
| 0.00 |
| 0.00 |
| 0.00 |
| 0.00 |
| 0.00 |
| 0.00 |
| 0.00 |
| 0.00 |

| GOBIO UP | #UP |
|---|---|
| Virulence | 23 |
| Regulation of transcription, DNA-dependent | 18 |
| Membrane transport | 15 |
| Peptidolysis | 4 |
| Nickel cation transmembrane transport | 7 |
| Phosphotransferase system | 9 |
| Cell wall organization in other organism | 21 |
| Carotenoid biosynthetic process | 0 |
| Response to antibiotic | 11 |
| DNA recombination | 5 |
| hemolysis in another organism | 0 |
| Peptidoglycan biosynthetic process | 15 |
| Regulation of cell shape | 14 |
| Translation | 13 |
| Cell division | 10 |
| queuosine biosynthetic process | 8 |
| Cell adhesion | 5 |

0.00
0.00
0.00
0.00
0.00
0.00

| %up | #Down | %Down |
|---|---|---|
| 3.15 | 43 | 7.66 |
| 2.47 | 22 | 3.92 |
| 2.05 | 19 | 3.39 |
| 0.55 | 18 | 3.21 |
| 0.96 | 16 | 2.85 |
| 1.23 | 11 | 1.96 |
| 2.88 | 8 | 1.43 |
| 0.00 | 6 | 1.07 |
| 1.51 | 5 | 0.89 |
| 0.68 | 5 | 0.89 |
| 0.00 | 5 | 0.89 |
| 2.05 | 0 | 0.00 |
| 1.92 | 0 | 0.00 |
| 1.78 | 0 | 0.00 |
| 1.37 | 0 | 0.00 |
| 1.10 | 0 | 0.00 |
| 0.68 | 0 | 0.00 |
